# Adaptive-Resolution Multi-Orientation Analysis of Complex Filamentous Network Images

**DOI:** 10.1101/757005

**Authors:** Mark Kittisopikul, Amir Vahabikashi, Takeshi Shimi, Robert D. Goldman, Khuloud Jaqaman

**Affiliations:** Department of Biophysics, UT Southwestern Medical Center, Dallas, TX 75390; Department of Cell and Developmental Biology, Feinberg School of Medicine, Northwestern University, Chicago, IL 60611; Institute of Innovative Research, Tokyo Institute of Technology, Yokohama, Japan; Department of Bioinformatics, UT Southwestern Medical Center, Dallas, TX 75390

## Abstract

Microscopy images of cytoskeletal, nucleoskeletal, and other filamentous structures contain complex junctions where multiple filaments with distinct orientations overlap, yet state-of-the-art software generally uses single orientation analysis to segment these structures. We describe an image analysis approach to simultaneously segment both filamentous structures and their intersections in microscopy images, based on analytically resolving coincident multiple orientations upon image filtering in a manner that balances orientation resolution and spatial localization.

## Introduction

Our approach uses steerable ridge filters to enhance curvilinear structures in images^1,2^. While most techniques using ridge filters identify the single best orientation from the filter response per point in space (e.g. pixel)^2-6^, our approach is based on the extraction of multiple orientations per point in space. This allows the full segmentation of junctions without producing gaps, an inherent limitation of the commonly used single-orientation identification followed by non-maximum suppression (NMS)^2,7^ (Fig. 1A-F). A common approach to complete junctions in this case has been to close the gaps using inference methods^3,8^, but these tend to involve heuristics. Additionally, unlike prior attempts to detect junctions explicitly, our approach does not need predefined angles^2^ or symmetries^9^. Our approach applies generally to steerable ridge filters, including to a unifying parametric framework for steerable wavelets^10^.

**Figure 1:**
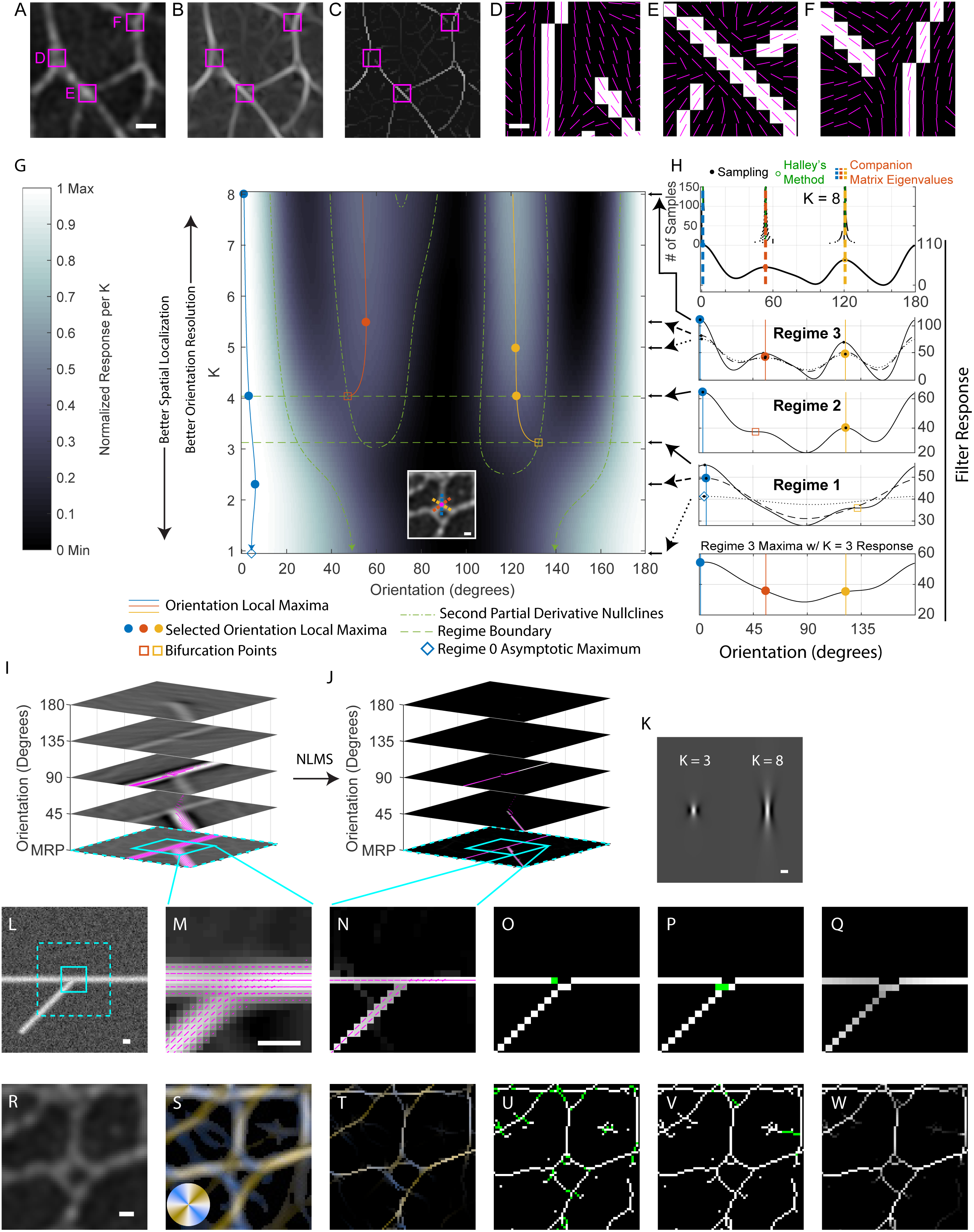
Multi-orientation detection and non-local maxima suppression (NLMS) segments simultaneously lines and junctions of arbitrary geometry. (**A**) Raw fluorescence intensity SIM image of Lamin A in a Lmnb1-/-Mouse Embryonic Fibroblast. Magenta boxes, also shown in B-C, are magnified in D, E, and F. Scale bar is 380 nm, applicable to B and C as well. (**B**) Response to Canny-optimized steerable filter. (**C**) Non-maximum suppression of steerable filter response. (**D-F**) Zoom-in of binarized non-maximum suppression (using threshold = 0) for boxes in A. Magenta lines indicate the best orientation per pixel detected by the Canny-optimized steerable filter (response shown in B). Scale bar is 60 nm. (**G**) Kymograph of filter response curves across the parameter K, with each row false-colored between its minimum and maximum values (as shown in color legend to the left of the kymograph). Filter response local maxima per K are depicted as blue, red, or yellow solid lines. Regimes of K containing distinct numbers of local maxima are demarked by horizontal dashed green lines. Filled-in circles indicate the selected filter local maxima according to the slope of the solid line. Empty squares indicate bifurcation points that occur at the intersection of the solid lines and the second partial derivative nullclines (green dot-dashed lines). Blue open diamond indicates location of asymptotic orientation maximum as K approaches −0.5. **Inset**: Raw image (same as in R), the center pixel of which is being analyzed in the kymograph. The center pixel is at the intersection of the three colored dashed lines, indicating the detected orientations at K=8, using the same color-coding as in the kymograph. Inset scale bar is 200 nm. (**H**) Top: Orientations detected using a companion matrix root-solver approach, indicated by the blue, red, and yellow lines, compared to those detected by sampling with a fixed number of samples (indicated on the left axis) or with Halley’s refinement method. Middle: Filter response curves for Regimes 3, 2, and 1, numbered according to the number of detected orientations (i.e. filter response local maxima), which are shown with the same symbols used in G. Bottom: Orientations detected in Regime 3 superimposed on the K = 3 filter response curve. (**I**) Orientation space construction where the x and y dimensions of a 2D image of a synthetic y-junction are augmented with an orientation dimension. Magenta lines indicate detected filter response local maxima at the indicated orientations. (**J**) Non-local maxima suppression (NLMS) applied to the orientation space in I. Magenta lines indicate non-suppressed pixels and their detected orientations. (**K**) Filter at K = 3 and K = 8. Scale bar is 200 nm. (**L**) Synthetic junction analyzed in I and J. Dashed cyan box indicates full extent of the representation in I and J. Solid cyan box indicates area depicted in M-Q. Scale bar is 200 nm. (**M**) Maximum response projection of orientation space in I with detected multiple orientations (magenta). Scale bar is 200 nm, applicable also to N-Q. (**N**) Maximum response projection of orientation space in J, in which the NLMS procedure has been applied. (**O, P**) Output of Step 1 (**O**) and Step 2 (**P**) of the minimal bridging segmentation algorithm, for junction in L. Bridges are shown as green pixels. (**Q**) Response-weighted segmentation of the y-junction (final output of the minimal bridging algorithm). (**R**) Raw fluorescence intensity SIM image of Lamin A immunofluorescence in a Lmnb1-/-Mouse Embryonic Fibroblast (same as inset in G). Scale bar is 200 nm, applicable also to S-W. (**S**) Maximum response projection color-coded by orientation according to the color wheel in the lower left. Colors are blended proportional to the orientation response if multiple orientations are detected. (**T-W**) Same as N-Q but for the image in R.

## Results and Discussion

The specific filter used in our work is based on that proposed by van Ginkel and van Vliet^11,12^. This filter is well-suited for multiple orientation detection because its aspect ratio, and thus orientation resolution, can be controlled via an explicit parameter, *K* (Supplemental Notes 1-3; specifically Eqs 25-29). However, the original filter suffered from discontinuities for values of *K* < 3 (Supplemental Fig. S1). By redefining the angular part of the filter in terms of angular frequency space instead of direct angular space (Supplemental Note 2.3; specifically Eqs 34-36), we were able to extend its applicability to the full range of *K*, namely *K* > −0.5 (Supplemental Fig. S1). This redefinition allowed us to develop a line and junction segmentation approach that integrates orientation information from multiple *K* values in order to balance orientation resolution (Supplemental Note 3) and spatial localization (discussed shortly). The final form of the filter employed in our study is given in Supplemental Note 2.4, Eqs 43 and 44.

To identify multiple coincident orientations from the filter response, we treated the steerable filter response, which is a finite Fourier series, as a trigonometric polynomial^1^. Therefore, the orientation(s) maximizing the filter response (i.e. filter response local maxima), along with all other extrema, were determined by solving a general eigenvalue problem using the polynomial’s companion matrix^13,14^ (Supplemental Notes 4 and 5). Unlike other eigensystem approaches, such as using a Hessian matrix^15^, our approach applies to any steerable filter response and is not constrained by the degree of the polynomial. The analytical and global nature of our approach allowed us to identify orientation(s) much more precisely than would be achievable by sampling approaches, at a fraction of the computational cost (Supplemental Fig. S2).

As mentioned above, the orientation resolution of our filter is controlled via the parameter *K*. Specifically, higher *K* produces higher aspect ratio filters that provide higher orientation resolution by integrating more signal along their length^1^. However, by construction, this increased orientation resolution comes at the expense of spatial localization. To balance the conflicting requirements of orientation resolution and spatial localization, we did not employ a single *K* for ridge filtering and subsequent analysis. Rather, we calculated the filter response for a range of *K* values, namely −0.5 < *K* ≤ *K*_*h*_ (*K*_*h*_ typically taken as 8, resulting in an orientation resolution = π/8; Supplemental Note 3). Noting that steerable filters at different *K* values are related by the heat-diffusion partial differentiation equation with periodic boundary conditions, here again we used analytical methods to calculate the ridge filter response at any *K* from the response at the maximum value *K*_*h*_ (Supplemental Note 4). In essence, the filter responses at distinct *K* levels are related by convolution/deconvolution with a known Gaussian kernel.

Having the filter response as a function of *K*, we devised a procedure of adaptive-orientation resolution ridge filtering (Fig. 1G, H; Supplemental Fig. S3; Supplemental Note 4). First, we determined the regimes of *K* in which different numbers of orientations are identified, starting with the maximum number in the highest regime (Regime H, which includes *K*_*h*_), and ending with asymptotically diminishing orientation information in the lowest regime (Regime 0, *K* in the open interval −0.5 and 1). Second, within each regime, we selected for each surviving orientation its optimal *K*, defined as the *K* at which the identified orientation changes the least with respect to *K*. The orientation values at their optimal *K*’s were then taken as the identified orientations in each regime. Altogether, adaptive resolution ridge filtering provided us with multi-orientation information at every pixel in the image in a manner that balances orientation resolution and spatial localization.

To then employ this information for image segmentation, we extended the traditional NMS concept in multiple ways, collectively resulting in a novel algorithm that we term Adaptive-Resolution Non-Local-Maxima Suppression (AR-NLMS; Fig. 1I-N; Movies S1, S2). NLMS is the core of this algorithm: Building on the concept of orientation space^11,16,17^, the algorithm performs NMS separately in each plane where signal orientation (i.e. filter response local maximum) has been identified (thus the term NLMS). The adaptive-resolution feature of the algorithm stems from (i) employing the adaptive-resolution ridge filtering procedure described above to identify signal orientations at their optimal *K* values within a selected resolution regime, and (ii) uncoupling the *K* used for identifying signal orientations from the *K* used to calculate the response to which NLMS is applied. For example, Fig. 1I-K shows AR-NLMS obtained by combining orientations identified from high *K* (*K*_*h*_ = 8) with a medium *K* response (*K*_*m*_ = 3), in order to balance the conflicting needs of orientation resolution and spatial localization. Performing a maximum response projection (MRP) along the orientation axis of all the AR-NLMS outputs then produced a 2D image resembling NMS. However, unlike NMS, the projected AR-NLMS retained multiple orientation information per pixel, and thus gaps did not appear at junctions (Fig. 1L-N).

While the projected AR-NLMS already yielded complete junctions without gaps, its output was not guaranteed to be a one-pixel-wide topological skeleton^18^ (Fig. 1N), as needed to identify the centers of lines and their intersections. Additionally, the AR-NLMS from different orientation resolutions (and, conversely, spatial localizations) may reveal different information about the signal in the image. Therefore, we devised a minimal bridging algorithm to produce a parsimonious segmentation of the centers of lines and junctions by integrating the multi-scale information available through AR-NLMS from different orientation resolution combinations. In a nutshell, starting with “simple” lines (i.e. no junctions) as identified from lower orientation resolution (but higher spatial localization) filters, this algorithm determines the shortest paths to bridge between these lines by using multi-orientation information revealed by higher orientation resolution filters (Fig. 1O-W; Supplemental Note 6; Supplemental Figs S4-S9). The output of the minimal bridging algorithm is a 1-pixel wide response-weighted segmentation (Fig. 1Q, W) incorporating orientation information across both low and high orientation resolution filters, allowing the algorithm to balance the trade-off between orientation resolution and spatial localization. This output resembles the NMS output of traditional single-orientation analysis^2,7^, and can be subsequently thresholded to segment the lines and junctions of interest (Supplemental Fig. S9).

To evaluate the performance of our AR-NLMS analysis and minimal bridging algorithm, we applied it to synthetic images of symmetric, asymmetric, and curved junctions over the full range of angles and at two distinct signal-to-noise ratios (5 and 10) (Fig. 2A-I; Supplemental Note 7; Supplemental Fig. S10; Movies S3-S8). We compared the number of identified orientations, their values, and the junction location (if detected) to the ground truth. We found that the algorithm performed as expected from theoretical predictions based on orientation and spatial resolution (Supplemental Note 3). The detected junction location deviated from the ground truth location for small angles, but then was very close to it when the intersecting orientations were separated by about π/8 (22.5 degrees). Performance for SNR 5 was slightly less robust than for SNR 10, as reflected by slightly larger performance metric standard deviations (Fig. 2G, I; Supplemental Fig. S10).

**Figure 2:**
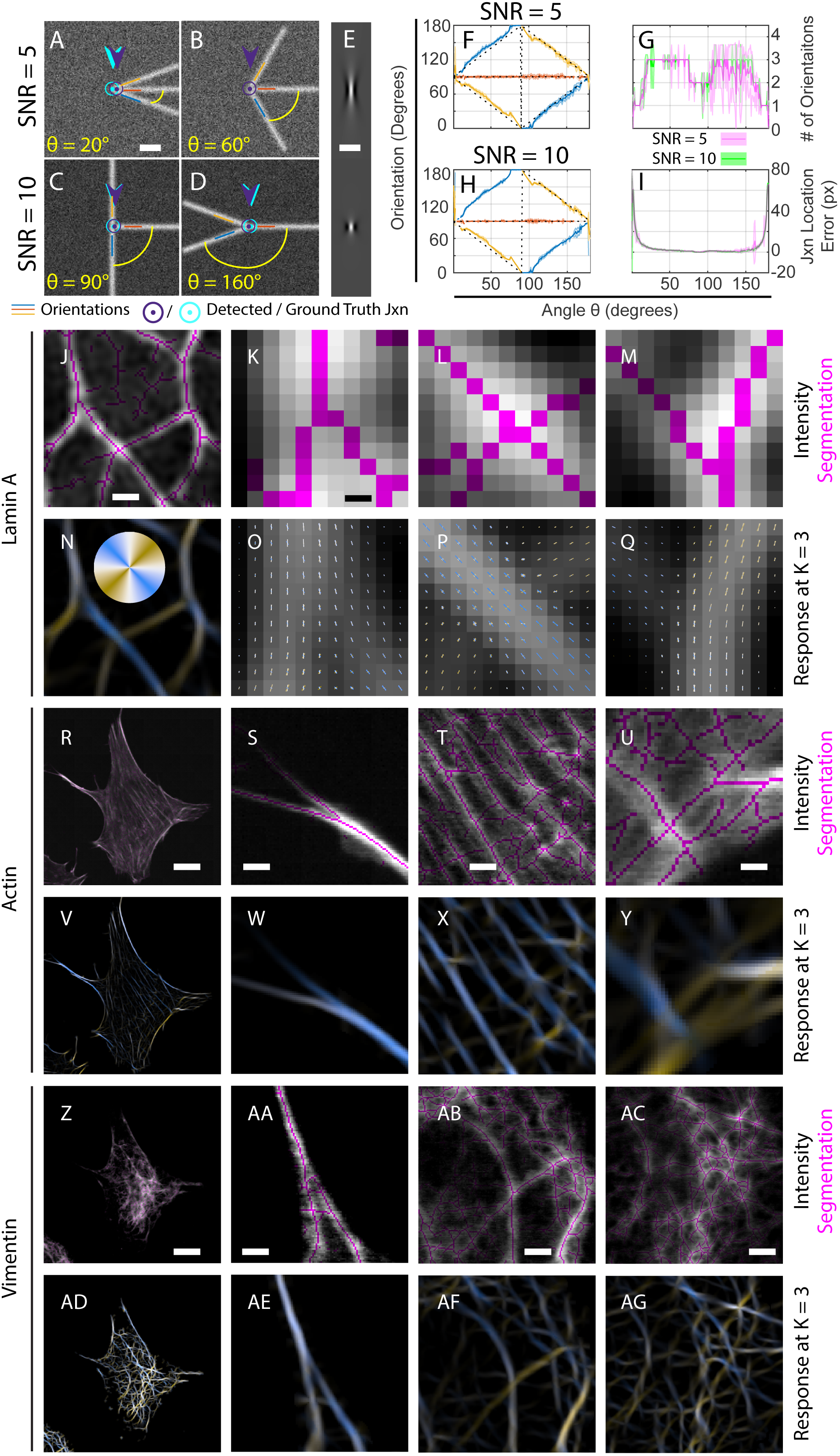
AR-NLMS and minimal bridging based segmentation of synthetic images and light microscopy images of biological filaments. (**A-D**) Examples of asymmetric, three radial line synthetic junctions with the indicated intervening angles. Scale bar in A is 600 nm, applicable to Panels B-D as well. (**E**) Ridge filters at K = 8 (top) and K = 3 (bottom). Scale bar is 600 nm. (**F-I**) Algorithm performance analysis for the three radial line junction case, as captured by comparing the detected orientations with the ground truth orientations (**F** for SNR 5 and **H** for SNR 10), the number of orientations at the detected junction in (**G**; SNR 5 in pink, SNR 10 in green), and the distance between the detected junction and the ground truth junction in (**I**; same color coding as in G). The results are the mean (dark lines) ± standard deviation (shaded area around dark lines) over 10 tests per angle and SNR. See Fig. S10 for further discussion of performance analysis results. (**J-Q**) Lamin A analyzed in a Lmnb1-/-MEF, corresponding to images in Fig. 1A-F. In **J-M**, the segmentation (magenta) is overlaid on the raw images. In **N-Q**, the filter response (brightness) and detected orientations (colored vectors) are shown, with colors corresponding to color wheel in N. Scale bars: 380 nm in J and N; 50 nm is K-M and O-Q. (**R-Y**) Orientation analysis and segmentation of phalloidin-stained actin in a MEF imaged via confocal microscopy. Panel descriptions as in J-Q. Scale bars: 11.4 μm in R and V; 1.4 μm in S, T, W and X; 0.72 μm in U and Y. (**Z-AG**) Orientation analysis and segmentation of vimentin immunofluorescence imaging via confocal microscopy. Cell same as in R. Panel descriptions as in J-Q. Scale bars: 11.4 μm in Z and AD; 1.5 μm in AA and AE; 2.2 μm in AB and AF; 2.7 μm in AC and AG.

To demonstrate the broad applicability of our approach, we applied it to complex networks formed by nuclear lamins (Fig. 2J-Q), the actin cytoskeleton (Fig. 2R-Y), and vimentin filaments (Fig. 2Z-AG) in samples of fixed mouse embryonic fibroblasts acquired through structured illumination microscopy (Fig. 2J-Q) or confocal microscopy (Fig. 2R-AG). We found that our approach was able to segment successfully both the lines and junctions in these complex networks, without producing gaps at junctions (e.g., compare Fig. 2K-M vs. Fig. 1D-F).

Overall, our work demonstrates that orientation response curves produced by ridge-like steerable filters contain sufficient information to completely segment complex curvilinear structures including junctions of arbitrary geometry without the need for inference and heuristics. Our method for extracting multi-orientation information with adaptive orientation resolution could thus be useful for deep learning approaches for image segmentation^19^. Of note, because the orientation resolution of our filter is determined by its aspect ratio, improved spatial resolution (e.g. in SIM images vs. conventional light microscopy images) allows the use of narrower filters and thus improved orientation resolution. Furthermore, this multi-orientation information could be used to localize lines and junctions with sub-pixel precision^4^. Our approach is generic, and can be applied to segment network-like biological structures other than cytoskeletal and nucleoskeletal filaments, such as stained membranes in cell sheets or network-like organelles such as the endoplasmic reticulum.

## Supporting information

MATLAB code from Github

URL for Uncompressed Movies

Movie S1: A Scan Through the Orientation Space Response

Movie S2: A Scan Through Non-Local Maxima Suppressed Orientation Space

Movie S3: Analysis of Two Symmetric Lines, SNR 5

Movie S4: Analysis of Two Symmetric Lines, SNR 10

Movie S5: Analysis of Three Radial Lines, SNR 5

Movie S6: Analysis of Three Radial Lines, SNR 10

Movie S7: Analysis of Two Arcs, SNR 5

Movie S8: Analysis of Two Arcs, SNR 10

## Online methods

Please see Section entitled “Online methods” at the end of the main text file.

## Acknowledgements

This work was supported by NIH/NCI (T32CA080621 for M.K. and A.V.), NIH/NICHD (U54HD087351, via UTSW Wellstone MDCRC Training Pilot Grant to M.K.), NIH/NIGMS (R35 GM119619 to K.J. and R01GM106023 to R.D.G.), CPRIT (R1216 to K.J.), and the UTSW Endowed Scholars program (to K.J.).

## Author contributions

M.K. and K.J. conceived of the project and wrote the manuscript. M.K. developed the software, tested it and applied it to experimental data, and prepared the figures. A.V., T.S. and R.D.G. provided the microscopy images.

## Additional information

This manuscript is accompanied by a Supplemental Information PDF file consisting of 8 notes and 10 figures, and by 8 movies (movie legends are Note 8 in Supplemental Information PDF).

Custom software written in MATLAB to implement the algorithms described in this work is available online on Github at the following URL: https://github.com/mkitti/AdaptiveResolutionOrientationSpace.

## Online methods

### Cell culture

The cell samples for actin and vimentin staining were Mouse embryonic fibroblasts (MEFs) provided by John Eriksson (Abo Akademi University, Turku, Finland). For lamin staining studies, the MEFs were provided by Yixian Zheng (Carnegie Institution for Science, Baltimore, MD, USA). Cells were maintained in DMEM with 25 mM HEPES and sodium pyruvate (Life Technologies; Grand Island, NY) supplemented by 10% fetal bovine serum, 1% penicillin streptomycin, and nonessential amino acids. All cell cultures were maintained at 37°C and 5% CO_2_.

### Immunofluorescence and Antibodies

Cold methanol fixation was used for 10 min at -20°C to fix the MEFs. Following fixation, initial blocking was done with 1% bovine serum albumin (BSA) in phosphate buffered saline (PBS; ThermoFisher 70011044) for 30 min at RT. Primary antibody incubation was done either at room temperature for 1 hr or overnight at 4°C for single or double label immunofluorescence. Antibody dilutions: chicken anti-vimentin (Biolegend 919101) at 1:200; rabbit anti-Lamin A (#323^1^) at 1:500. Cells were washed in PBS at RT and then incubated with secondary antibodies: goat anti-chicken Alexa Fluor 488 (ThermoFisher A-11039) at 1:1000 dilution; goat anti-rabbit Alexa Fluor 488 (ThermoFisher A-11008) at 1:1000 dilution. Cells were washed and stained with Hoechst 33342 (ThermoFisher H-1399) to help locate cells, and in some cases Alexa Fluor 647 phalloidin (ThermoFisher 8940S) for 1 hr according to manufacturer’s instructions.

### Confocal Microscopy

Images of MEFs stained for vimentin and actin were imaged using a scanning confocal Zeiss LSM 510 META microscope (Carl Zeiss, Jena, Germany) using an oil immersion objective lens (PlanApochromat, 63x, 1.40 numerical aperture).

### Structured Illumination Microscopy

Images of nuclear lamin A were acquired using a Nikon Structured Illumination Microscope (N-SIM; Nikon, Tokyo, Japan) using an oil immersion objective lens CFI SR (Apochromat TIRF 100x, 1.49 NA, Nikon). N-SIM images were reconstructed using Nikon Elements with the following reconstruction parameters: Illumination Contrast Modulation 1.00, High Frequency Noise Suppression 0.75, and Out-of-Focus Blur Suppression 0.15.

### Image Analysis and Computing Environment

Images acquired by microscopy were loaded into MATLAB 2017a (Natick, MA) using Bioformats^2^. All acquired images are shown with linear contrast between the maximum and minimum value unless otherwise noted. The steerable filter code based on Jacobs and Unser was implemented by Francois Aguet, and was previously distributed^3^. Image analysis code was executed on the UT Southwestern Biological High Performance Computing (BioHPC) cluster. Custom software written in MATLAB to implement the algorithms described in this work is available online on Github at the following URL: https://github.com/mkitti/AdaptiveResolutionOrientationSpace.

## References

1 Freeman, W. T. & Adelson, E. H. The Design and Use of Steerable Filters. Ieee T Pattern Anal 13, 891–906, doi:Doi 10.1109/34.93808 (1991).

2 Jacob, M. & Unser, M. Design of steerable filters for feature detection using Canny- like criteria. Ieee T Pattern Anal 26, 1007–1019, doi:Doi 10.1109/Tpami.2004.44 (2004).

3 Gan, Z. et al. Vimentin Intermediate Filaments Template Microtubule Networks to Enhance Persistence in Cell Polarity and Directed Migration. Cell Syst 3, 252-+, doi:10.1016/j.cels.2016.08.007 (2016).

4 Steger, C. Subpixel-precise extraction of lines and edges. International Archives of Photogrammetry and Remote Sensing 33, 141–156 (2000).

5 Peters, R., Griffie, J., Burn, G. L., Williamson, D. J. & Owen, D. M. Quantitative fibre analysis of single-molecule localization microscopy data. Sci Rep 8, 10418, doi:10.1038/s41598-018-28691-5 (2018).

6 Arganda-Carreras, I. et al. Trainable Weka Segmentation: a machine learning tool for microscopy pixel classification. Bioinformatics 33, 2424–2426, doi:10.1093/bioinformatics/btx180 (2017).

7 Canny, J. A Computational Approach to Edge-Detection. Ieee T Pattern Anal 8, 679–698, doi:Doi 10.1109/Tpami.1986.4767851 (1986).

8 Shimi, T. et al. Structural organization of nuclear lamins A, C, B1, and B2 revealed by superresolution microscopy. Mol Biol Cell 26, 4075–4086, doi:10.1091/mbc.E15-07-0461 (2015).

9 Puspoki, Z., Uhlmann, V., Vonesch, C. & Unser, M. Design of Steerable Wavelets to Detect Multifold Junctions. IEEE Trans Image Process 25, 643–657, doi:10.1109/TIP.2015.2507981 (2016).

10 Unser, M. & Chenouard, N. A Unifying Parametric Framework for 2D Steerable Wavelet Transforms. Siam Journal on Imaging Sciences 6, 102–135, doi:10.1137/120866014 (2013).

11 Ginkel, M. v. Image analysis using orientation space based on steerable filters. PhD thesis, Technische Universiteit Delft, (2002).

12 Nieuwenhuizen, R. P. et al. Co-Orientation: Quantifying Simultaneous Co-Localization and Orientational Alignment of Filaments in Light Microscopy. PLoS One 10, e0131756, doi:10.1371/journal.pone.0131756 (2015).

13 Boyd, J. P. Computing the zeros, maxima and inflection points of Chebyshev, Legendre and Fourier series: solving transcendental equations by spectral interpolation and polynomial rootfinding. J Eng Math 56, 203–219, doi:10.1007/s10665-006-9087-5 (2006).

14 Marchant, R. & Jackway, P. Local Feature Analysis Using a Sinusoidal Signal Model Derived from Higher-Order Riesz Transforms. Ieee Image Proc, 3489–3493 (2013).

15 Frangi, A. F., Niessen, W. J., Vincken, K. L. & Viergever, M. A. Multiscale vessel enhancement filtering. Medical Image Computing and Computer-Assisted Intervention-Miccai’98 1496, 130–137 (1998).

16 Chen, J., Sato, Y. & Tamura, S. Orientation space filtering for multiple orientation line segmentation. Ieee T Pattern Anal 22, 417–429 (2000).

17 vanGinkel, M., Verbeek, P. W. & vanVliet, L. J. Improved orientation selectivity for orientation estimation. Scia’97 -Proceedings of the 10th Scandinavian Conference on Image Analysis, Vols 1 and 2, 533–537 (1997).

18 Lam, L., Lee, S. & Suen, C. Y. Thinning methodologies-a comprehensive survey. Ieee T Pattern Anal 14, 869–885, doi:10.1109/34.161346 (1992).

19 Depeursinge, A., Puspoki, Z., Ward, J. P. & Unser, M. Steerable Wavelet Machines (SWM): Learning Moving Frames for Texture Classification. Ieee T Image Process 26, 1626–1636, doi:10.1109/Tip.2017.2655438 (2017).

## References

1 Dechat, T. et al. Alterations in mitosis and cell cycle progression caused by a mutant lamin A known to accelerate human aging. Proc Natl Acad Sci U S A 104, 4955–4960, doi:10.1073/pnas.0700854104 (2007).

2 Linkert, M. et al. Metadata matters: access to image data in the real world. J Cell Biol 189, 777–782, doi:10.1083/jcb.201004104 (2010).

