## Supplemental Materials for "Adaptive-Resolution Multi-Orientation Analysis of Complex Filamentous Network Images"

**8 Movie Legends** **43**

### Introduction

In this work, we make use of Fourier analysis to detect orientation information embedded in an image. This supplement contains a description of Fourier Transforms, the Fourier domain descriptions of the steerable filters used, and the analytical routines used to extract orientation information. Additionally, a detailed explanation of the segmentation algorithm based upon this information is provided as well as the evaluation procedure used.

### 1 Mathematical preliminaries

The below definitions are stated as applied to a discretely-sampled image, $I(x, y)$, with a domain $(x, y) \in [1, N] \times [1, M]$ with $I(x, y) \in \mathbf{R}$.

#### 1.1 2-D Fourier transforms

Fourier transforms are used extensively in this work for image processing. The forward two dimensional continuous Fourier transform is defined as:

$$\hat{F}(f_x, f_y) = \mathcal{F}_{xy}\{F(x, y)\}(f_x, f_y) \tag{1}$$

$$= \iint_{-\infty}^{+\infty} F(x, y) \exp\{(-2\pi i(xf_x + yf_y))\} dx\, dy \tag{2}$$

The corresponding reverse two dimensional continuous Fourier transform is then defined as:

$$F(x, y) = \mathcal{F}_{xy}^{-1}\{\hat{F}(f_x, f_y)\}(x, y) \tag{3}$$

$$= \iint_{-\infty}^{+\infty} \hat{F}(f_x, f_y) \exp\{(2\pi i(xf_x + yf_y))\} df_x\, df_y \tag{4}$$

They are discretized according to the normalization conventions used in MATLAB. The discrete forward 2D Fourier Transform is defined as:

$$\hat{F}(f_x, f_y) = \mathcal{F}_{xy}\{F(x, y)\}(f_x, f_y) \tag{5}$$

$$= \sum_{x=0}^{N-1} \sum_{y=0}^{M-1} F(x, y) \exp\{(-2\pi i(x f_x/N + y f_y/M))\} \tag{6}$$

The corresponding discrete reverse 2D Fourier Transform is then defined as:

$$F(x, y) = \mathcal{F}_{xy}^{-1}\{\hat{F}(f_x, f_y)\}(x, y) \tag{7}$$

$$= \frac{1}{NM} \sum_{f_x=0}^{N-1} \sum_{f_y=0}^{M-1} \hat{F}(f_x, f_y) \exp\{(2\pi i(x f_x/N + y f_y/M))\} \tag{8}$$

#### 1.2 Structure of the Fourier domain of a real-valued image

The two dimensional transform of an image is as follows, in terms of the spatial frequency components $(f_x, f_y)$:

$$\hat{I}(f_x, f_y) = \mathcal{F}_{xy}\{I(x, y)\}(f_x, f_y) \tag{9}$$

Because $I(x, y) \in \mathbf{R}$, the Fourier transform is conjugate symmetric around the origin:

$$\hat{I}(f_x, f_y) = \hat{I}^*(-f_x, -f_y) \tag{10}$$

To analyze orientation, it is convenient to address the Fourier domain using polar coordinates $(f, \theta)$ with $f \in [0, \pi/Q]$ and $\theta \in [0, 2\pi)$ where $Q = \min(N, M)$.

$$f^2 = f_x^2 + f_y^2 \tag{11}$$

$$\theta = \arctan \frac{f_y}{f_x} \tag{12}$$

The polar coordinate representation is easily translated to Cartesian coordinates.

$$f_x = f \cos\theta \tag{13}$$

$$f_y = f \sin\theta \tag{14}$$

Thus for the purposes of notation $\hat{I}(f, \theta) \equiv \hat{I}(f \cos\theta, f \sin\theta) = \hat{I}(f_x, f_y)$.

#### 1.3 1-D Fourier transform for orientation

To discuss ridge filter responses, we will also need a one dimensional Fourier transform for a domain with a period of $\pi$, where the Fourier coefficients $c_k$ are given by:

$$c_k = \hat{\rho}(k) = \sum_{n=-K}^{K} \rho(\phi_n) \exp\{(-2ik\phi_n)\} \qquad \phi_n = \frac{n\pi}{2K+1} \tag{15}$$

$[-K, K]$ are the limits for angular samples. The reverse Fourier transform is then given by:

$$\rho(\phi) = \frac{1}{2K+1} \sum_{k=-K}^{K} c_k \exp(2ik\phi) \tag{16}$$

#### 1.4 Complexified Riesz transform

The Riesz transform stated in complex by Unser and Chenuoard [1] allows the 2-D spatial Fourier transform and 1-D angular Fourier transform to be combined into a single mathematical formalism.

$$\mathcal{F}_{xy}\{\mathcal{R}I(x,y)\}(f_x, f_y) \equiv \frac{(f_x + if_y)}{\|f\|} \hat{I}(f_x, f_y) \tag{17}$$

Addressing the Fourier domain in polar coordinates, the nature of the multiplier in the Fourier domain becomes apparent.

$$\mathcal{F}_{xy}\{\mathcal{R}I(x,y)\}(f,\theta) = \exp(i\theta)\hat{I}(f,\theta) \tag{18}$$

The conjugate Riesz transform then just has the conjugate Fourier multiplier.

$$\mathcal{F}_{xy}\{\mathcal{R}^*I(x,y)\}(f,\theta) \equiv \exp(-i\theta)\hat{I}(f,\theta) \tag{19}$$

By extension of the definition, repeated applications of the Riesz transform and its conjugate can be expressed through a multiplier in the Fourier domain.

$$\mathcal{F}_{xy}\{\mathcal{R}^n I(x,y)\}(f,\theta) = \exp(in\theta)\hat{I}(f,\theta) \tag{20}$$

$$\mathcal{F}_{xy}\{\mathcal{R}^{2k} I(x,y)\}(f,\theta) = \exp(2ik\theta)\hat{I}(f,\theta) \tag{21}$$

$$\mathcal{F}_{xy}\{\mathcal{R}^{2k*} I(x,y)\}(f,\theta) = \exp(-2ik\theta)\hat{I}(f,\theta) \tag{22}$$

Note that inverting the 2-D Fourier transform sums the Riesz multiplied Fourier coefficients for each pixel. This in turn yields the angular Fourier coefficients for each pixel:

$$c_k(x, y) = \mathcal{R}^{2k*} I(x, y) \tag{23}$$

$$= \mathcal{F}_{xy}^{-1}\{\exp(-2ik\theta)\hat{I}(f, \theta)\}(x, y) \tag{24}$$

##### 1.5 Summary of symbols

| Symbol | Description |
|---|---|
| $x, y$ | Spatial coordinates of 2-D image |
| $f_x, f_y$ | Spatial frequency coordinates of Fourier Transform of 2D image |
| $f, \theta$ | Spatial frequency in polar coordinates |
| $\mathcal{F}$ | Fourier transform |
| $\mathcal{F}^{-1}$ | Inverse Fourier transform |
| $\mathcal{F}_{xy}$ | 2-D Fourier Transform |
| $I, \hat{I}$ | Image and its 2-D Fourier Transform |
| $N$ | Number of pixels in the x-dimension |
| $M$ | Number of pixels in the y-dimension |
| $Q$ | Least number of pixels in x- or y-dimensions |
| $K$ | $[-K, K]$ are the limits for angular samples |

#### 2 Filter design

In this work, we employ an orientation selective steerable filter that is polar-separable in the Fourier domain, and focus on the orientation component of the filter. The specific filter we have chosen is adapted from that developed by van Ginkel and van Vliet [2] for the purpose of designing an orientation space, and parameterized according to its orientation resolution.

The orientation component of the filter is also compatible with a universal parameterized framework for steerable wavelets when the radial component of the filter is a wavelet (with value zero outside of an interval).

Here we first review the original filter design [2], and then describe our modifications to the radial and orientation components of the filter. We conclude this section by summarizing the filter as used in this study.

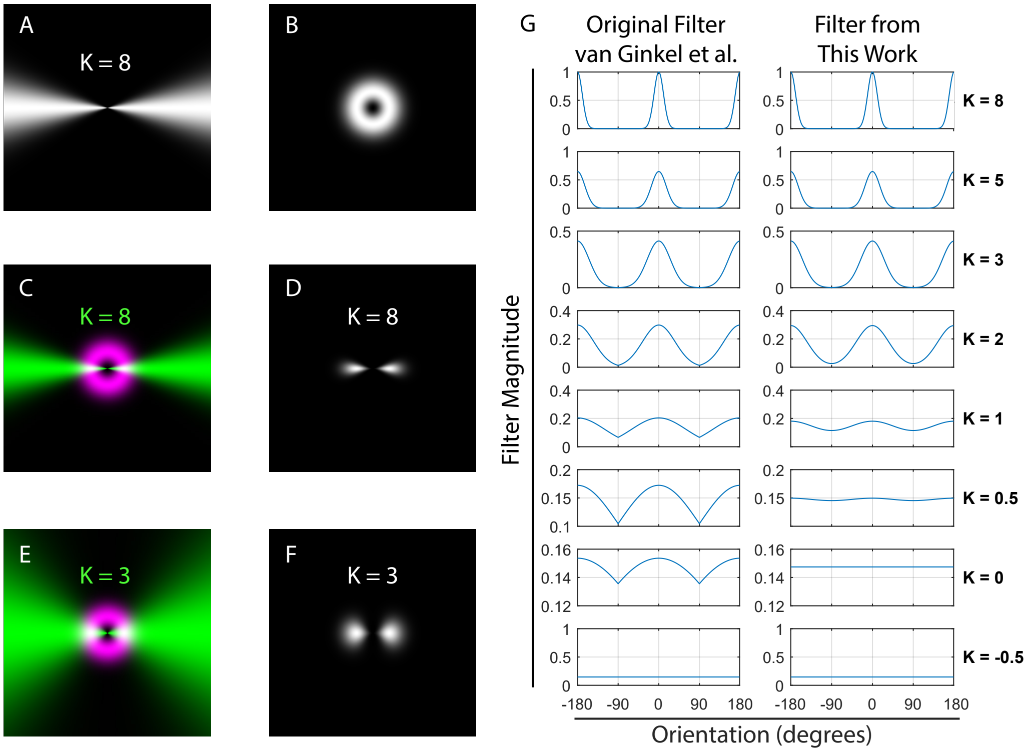

**Figure S1: Filter Design.**
The filter is derived from an original design described by van Ginkel et al. [2], which is polar separable into angular and radial components in the Fourier domain. (A) Angular component of the filter at $K = 8$. (B) Radial component of the filter with $f_c = b_f = \frac{1}{4\pi}$. (C) Color merge of angular (green) and radial (magenta) components of the filter as in A and B. (D) Pointwise product of angular and radial components of the filter. (E) Color merge of angular (green) and radial (magenta) components of the filter with $K = 3$ and $f_c = b_f = \frac{1}{4\pi}$. (F) Pointwise product of angular and radial filter components shown in E. (G) The angular component in this work is defined in terms of angular frequency rather than angle, resulting in the original filter and the filter used in this work becoming distinct for $K < 3$.

#### 2.1 van Ginkel et al. orientation space filter

The orientation space filter described in [2] is stated as follows. The filter is divided into a radial component, $\hat{\Phi}_{[f]}$, and an angular component, $\hat{\Phi}_{[G:\theta]}$ (based on a Gaussian). The filter design is illustrated in Fig. S1A-F.

$$\hat{\Phi}(f, \theta; \phi, f_c, b_f, K) = \hat{\Phi}_{[f]}(f; f_c, b_f)\hat{\Phi}_{[G:\theta]}(\theta - \phi; K) \tag{25}$$

The radial part is constructed with two parameters $f_c$ and $b_f$, which represent the central spatial frequency and spatial frequency bandwidth.

$$\hat{\Phi}_{[f]}(f; f_c, b_f) = \left(\frac{f}{f_c}\right)^{\frac{f_c^2}{b_f^2}} \exp\left(-\frac{f^2 - f_c^2}{2b_f^2}\right) \tag{26}$$

This creates a bandpass filter resembling an annulus with a peak value of 1 at $f = f_c$ and a width controlled by $b_f$ (Fig. S1B). For ridge detection, it suffices that $b_f \approx f_c$ (smaller values of $b_f$ permit texture analysis that is not used in this work). This leads to the simplification of the radial component described in the next section.

Another feature of this filter is that its value is zero at $f = 0$, meaning that the zeroth frequency component is not passed by the filter. In other words, the mean value of the intensities of the image does not affect the filter response.

The exact form of the radial component of the filter does not affect the main results of our work and this could be easily replaced by another radial filter. For example, the radial compoennt could be replaced by radial wavelets that can permit spatial multi-resolution frequency analysis [1].

The angular component of the filter has the form of a Gaussian function, with standard deviation $s_a$ defined by the orientation resolution parameter $K$.

$$\hat{\Phi}_{[G:\theta]}(\theta; K) = 2\exp\left(-\frac{\theta^2}{2s_a^2}\right) \qquad \theta \in [-\frac{\pi}{2}, \frac{\pi}{2}) \tag{27}$$

$$s_a = \frac{\pi}{2K+1} \tag{28}$$

$K$ is a more convenient parameter than $s_a$ in the filter definition because it is more directly used to define the sampling requirements. The subscript $G$ indicates this is one of many of angular functions that van Ginkel considered, with this Gaussian based version being a practical compromise [2].

$\hat{\Phi}_{[G:\theta]}(\theta; K)$ can be extended to the domain $\theta \in (-\pi, \pi)$ by the relation

$$\hat{\Phi}_{[G:\theta]}(\theta + \pi; K) = \hat{\Phi}_{[G:\theta]}(\theta; K) \tag{29}$$

To summarize, the symbols in the filter definition are described in the table below.

| Symbol | Description |
|---|---|
| $f$ | Radial spatial frequency, $f \in [0, +\infty)$ |
| $\theta$ | Polar angle in the Fourier domain, $\theta \in [-\pi, \pi)$ |
| $f_c$ | Central frequency of the spatial bandpass filter |
| $b_f$ | Spatial frequency bandwidth |
| $\phi$ | Orientation angle selected by the filter |
| $K$ | Orientation resolution parameter |
| $s_a$ | Orientation resolution bandwidth |

#### 2.2 Simplified radial spatial filter component

The spatial filter component forms an annulus in the Fourier domain with central radius of $f_c$ and a width of $b_f$. As spatial scale space is not a focus of the current work, we simplified the radial spatial filter component by setting $b_f = f_c$:

$$\hat{\Phi}_{[f]}(f; f_c) = \left(\frac{f}{f_c}\right) \exp\left(-\frac{f^2}{2f_c^2}\right) \exp\left(\frac{1}{2}\right) \tag{30}$$

An important benefit of this simplification is that the integrals of the radial function in this form can be solved analytically (Box S1). Another application is that it allows us to determine the optimal central frequency $f_c$ to respond to a line with a Gaussian cross-section, given its standard deviation $\hat{\sigma}$ in frequency space (Box S2).

**Box S1 : Simplified radial filter has a closed form integral**

One justification for the radial component simplification is that integrals of the radial function can be solved analytically. For example, to solve for the peak central value of the reverse Fourier transform of the radial function, we would need to compute the following polar integral.

$$\text{We want to solve } \int_{-\pi}^{\pi}\int_{0}^{\infty} \hat{\Phi}_{[f]}(f; f_c) f df d\theta$$

First we solve the inner integral.

$$\text{Let } g = \frac{f}{f_c}, \quad dg = \frac{df}{f_c}$$

$$\begin{aligned}
\int_0^\infty \hat{\Phi}_{[f]}(f; f_c) f df &= \int_0^\infty \left(\frac{f^2}{f_c}\right) \exp\left(-\frac{f^2}{2f_c^2}\right) \exp\left(\frac{1}{2}\right) df \\
&= \sqrt{e}\int_0^\infty g^2 f_c \exp\left(-\frac{g^2}{2}\right) f_c dg \\
&= f_c^2\sqrt{e}\int_0^\infty g^2 \exp\left(-\frac{g^2}{2}\right) dg \\
&= f_c^2\sqrt{\frac{e\pi}{2}}
\end{aligned}$$

$$\int_{-\pi}^{\pi}\int_{0}^{\infty} \hat{\Phi}_{[f]}(f; f_c) f df d\theta = 2\pi f_c^2\sqrt{\frac{e\pi}{2}} = f_c^2\sqrt{2e\pi^3}$$

Non-integer ratios of $f_c$ and $b_f$ would involve numerical solutions to the Gamma function [3].

**Box S2 : Optimal radial filter for a Line with a Gaussian cross-section**

If we consider an infinitely long line convolved with a Gaussian, the Fourier domain representation is $\delta(f_x)$ times the reciprocal Gaussian. The delta function reduces the filter response calculation to a 1D integral with respect to $f$.

$$\text{Find } f_c > 0 \text{ s.t. } \frac{\mathrm{d}}{\mathrm{d}f_c}\int_0^\infty \hat{\Phi}_{[f]}(f;f_c)\exp\left(-\frac{f^2}{2\hat{\sigma}^2}\right)df = 0$$

$$\text{Let } s = \frac{\hat{\sigma}}{\sqrt{f_c^2+\hat{\sigma}^2}}, \quad \frac{\mathrm{d}s}{\mathrm{d}f_c} = \frac{-f_c\hat{\sigma}}{(f_c^2+\hat{\sigma}^2)^{3/2}} = \frac{-f_c s^3}{\hat{\sigma}^2}$$

$$\frac{\mathrm{d}}{\mathrm{d}f_c}\int_0^\infty \left(\frac{f}{f_c}\right)\exp\left(-\frac{f^2}{2s^2f_c^2}\right)\exp\left(\frac{1}{2}\right)df = 0$$

$$\frac{\mathrm{d}}{\mathrm{d}f_c}s^2 f_c = s^2 - 2f_c^2s^4/\hat{\sigma}^2 = s^2(1-2f_c^2s^2/\hat{\sigma}^2) = 0$$

$$f_c^2+\hat{\sigma}^2-2f_c^2 = \hat{\sigma}^2 - f_c^2 = 0 \therefore f_c = \hat{\sigma}$$

Therefore, with the simplification $b_f = f_c$, the radial filter reaches an extremum when $f_c = \hat{\sigma}$. The second derivative test classifies this extremum as a maximum:

$$\frac{\mathrm{d}^2}{\mathrm{d}f_c^2}s^2 f_c = -2f_cs^4/\hat{\sigma}^2 - 4f_cs^4/\hat{\sigma}^2 + 8f_c^3s^6/\hat{\sigma}^4$$

$$= -1/(2\hat{\sigma}) < 0 \text{ when } f_c = \hat{\sigma}$$

Thus, the filter is maximally responsive to the line when $f_c = \hat{\sigma}$ .

The following useful properties of the original filter are retained:
1. The zeroth component does not affect the filter response.

$$\hat{\Phi}_{[f]}(0;f_c) = 0 \tag{31}$$

2. The maximum value of the filter is 1 and is obtained at $f = f_c$

$$\hat{\Phi}_{[f]}(f_c;f_c) = 1 \geq \hat{\Phi}_{[f]}(f \neq f_c;f_c) \tag{32}$$

3. The filter approaches zero for large values of $f$.

$$\lim_{f \to +\infty} \hat{\Phi}_{[f]}(f; f_c) = 0 \tag{33}$$

#### 2.3 Redefined angular filter component

The angular filter component as designed by van Ginkel is not well defined for values of $K < 3$, resulting in $s_a > \frac{\pi}{7}$, because $\hat{\Phi}_{[G:\theta]}(\theta; K < 3)$ is noticeably distinct from zero at the periodic boundary, and thus the transfer function of the filter is not well defined in Fourier space. To address this, we redefined the filter explicitly in Fourier space in terms of angular frequency.

$$\hat{\Phi}_{[\hat{G}:\theta]}(\theta; K, K_h) = s_a\sqrt{2\pi} \sum_{k=-K_h}^{K_h} \exp\left(-2\pi^2 k^2 s_a^2\right) \exp(2ik\theta) \tag{34}$$

$$= s_a\sqrt{2\pi} \sum_{j=-2K_h}^{2K_h} c_j \exp(i\omega_j\theta/(2\pi)) \tag{35}$$

$$c_j = \begin{cases} \exp\left(-w_j^2 s_a^2/2\right), & \text{if } j \text{ is even} \\ 0, & \text{if } j \text{ is odd} \end{cases} \tag{36}$$

where $K \leq K_h$, $\theta \in [-\pi, \pi)$, $s_a = \frac{\pi}{2K+1}$ (as defined in Section 2.1 above), and $\omega_j = 2\pi j$. Under this definition, $\hat{\Phi}_{[\hat{G}:\theta]}(\theta; K, K_h)$ is explicitly defined on a domain with a period of $2\pi$, although it has a period of $\pi$. This is because only the even coefficients are non-zero, and thus $\hat{\Phi}_{[\hat{G}:\theta]}$ is an even function, such that $\hat{\Phi}_{[\hat{G}:\theta]}(\theta; K, K_h) = \hat{\Phi}_{[\hat{G}:\theta]}(-\theta; K, K_h)$.

For comparison, the two Gaussian waveforms either in terms of angle, $\theta$, or angular-frequency, $k$, are as follows:

$$G(\theta; s_a(K)) = \exp\left(-\frac{\theta^2}{2s_a^2}\right) \tag{37}$$

$$\hat{G}(k; s_a(K)) = s_a\sqrt{2\pi} \exp\left(-2\pi^2 k^2 s_a^2\right) \tag{38}$$

$\hat{\Phi}_{[\hat{G}:\theta]}$ is closely related to $\hat{\Phi}_{[G:\theta]}$ in the following way if $\hat{\Phi}_{[G:\theta]}(\theta; K)$ were extended directly on to the domain with a period of $2\pi$ rather than defining it on a domain of period $\pi$ and replicating it over another period of $\pi$:

$$\hat{\Phi}_{[G':\theta]}(\theta; K) = 2\exp\left(-\frac{\theta^2}{2s_a^2}\right), \quad \theta \in [-\pi, \pi) \tag{39}$$

$$= \sum_{j=-2K_h}^{2K_h} 2\exp\bigl(-w_j^2 s_a^2/2\bigr)\exp(i\omega_j\theta/(2\pi)) \tag{40}$$

$$\hat{\Phi}_{[G':\theta]}(\theta+\pi; K) = \sum_{j=-2K_h}^{2K_h} 2\exp\bigl(-w_j^2 s_a^2/2\bigr)\exp(i\omega_j\theta/(2\pi))(-1)^j \tag{41}$$

$$\hat{\Phi}_{[\hat{G}:\theta]}(\theta; K, K_h) = \left[\hat{\Phi}_{[G':\theta]}(\theta; K) + \hat{\Phi}_{[G':\theta]}(\theta+\pi; K)\right]/2 \tag{42}$$

For larger values of $s_a$ the periodic domain may have to be extended further in increments of $\pi$ such that $\hat{\Phi}_{[G':\theta]} \approx 0$ at the periodic boundary. Additional terms offset by $\pi$ would then be required in the summation in order to relate the redefined filter to the original one.

Defining the angular part of the filter in terms of angular frequency has several advantages over the original definition in terms of angle [2]:

1. For $K \geq 3$, $\hat{\Phi}_{[\hat{G}:\theta]}(\theta; K, K_h) \approx \hat{\Phi}_{[G:\theta]}(\theta; K)$, but the definition begins to appreciably differ for $K < 3$, where the Gaussian becomes distinct from zero for $\|\theta\| > \pi/2$.

2. The coefficients in this finite Fourier series are the complexified Riesz coefficients in the context of the parametric framework for steerable wavelets [1]. In other words, the form above shows how the filter could be applied by first evaluating the circular harmonic coefficients and then multiplying by the Gaussian term.

3. The definition is compatible with the Heat-diffusion equation relationship exploited in Note 3 below.

The missing odd coefficients comprise the corresponding edge filter which produces an odd orientation filter response function. All of the coefficients would provide for a single Gaussian on the domain spanning $2\pi$ resulting in a quadrature filter generating a complex-valued response.

In this work, our focus is on ridge filters alone, and thus on only the even coefficients of the finite Fourier series.

#### 2.4 Form of filter used in this study

To conclude, we have modified the orientation space steerable filter [2] by: 1) simplifying the parameters for the radial spatial frequency component, and 2) redefining the angular component in terms of angular frequency. For clarity, we state the modified filter design here explicitly:

$$
\begin{aligned}
\hat{\Phi}(f,\theta;\phi,f_c,K) = &\hat{\Phi}_{[f]}(f;f_c)\hat{\Phi}_{\hat{G}:\theta}(\theta-\phi;K) \quad (43)\\
=&\sqrt{e}\left(\frac{f}{f_c}\right)\exp\left(-\frac{f^2}{2f_c^2}\right)\\
&\cdot\left[\sum_{k=-K_h}^{K_h} 2\exp\left(-2\pi^2k^2s_a^2\right)\exp(2ik(\theta-\phi))\right] \quad (44)
\end{aligned}
$$

For the numerical software implementation, the default normalization is set such that the sum across the angular samples remains constant regardless of $K$ or $K_h$.

Below are the parameters as used in our filter.

| Symbol | Description |
|---|---|
| $f$ | Radial spatial frequency, $f \in (0, +\infty)$ |
| $\theta$ | Polar angle in the Fourier domain, $\theta \in [-\pi, \pi)$ |
| $f_c$ | Central frequency and bandwidth of the spatial bandpass filter |
| $\phi$ | Orientation angle selected by the filter |
| $K$ | Orientation resolution parameter |
| $K_h$ | Highest orientation resolution parameter |
| $s_a$ | Orientation resolution bandwidth |

### 3 Orientation resolution

#### 3.1 Definition

Orientation resolution is the minimum separation angle required to distinguish two orientations. In the case of finite discrete Fourier series, the practical resolution is related to the shortest wavelength with a coefficient distinguishable from zero. Therefore, the orientation resolution is the wavelength of the highest frequency component of the Fourier series:

$$\text{Orientation Resolution} = \frac{\pi}{K} \quad (45)$$

#### 3.2 Limits of orientation resolution

The upper limit of orientation resolution is determined by the spatial resolution of the filter. Given the scale of the angular Gaussian $s_a$, we can calculate the arc length swept by the Gaussian:

$$\text{For } s_a = \frac{\pi}{2K+1} \tag{46}$$

$$\text{arc length } = \frac{f_c \pi}{2K+1} \tag{47}$$

The practical limit is that this arc length should be larger than the discrete frequency interval, $\frac{2\pi}{Q}$, at which point the angular Gaussian becomes effectively a delta function. This imposes an upper limit on the orientation resolution parameter K:

$$\frac{f_c \pi}{2K+1} \geq \frac{2\pi}{Q} \tag{48}$$

$$4K + 2 \leq Q f_c \tag{49}$$

$$K \leq \frac{Q f_c - 2}{4} \tag{50}$$

The lower limit for $K$ is $-\frac{1}{2}$. As $K$ approaches $-\frac{1}{2}$, $s_a$ approaches $+\infty$, meaning that no orientation information is obtained. The response to the filter at the lower bound is effectively the mean orientation filter response value which is the same as if the radial component of the filter were applied directly.

#### 3.3 Interplay between orientation resolution and spatial resolution

Besides setting an upper bound to orientation resolution as described in the previous section, spatial resolution limits orientation resolution also by the basic requirement that for two structures with distinct orientations to be detected as such, they must be resolvable in space. Otherwise, the two structures may appear as a single thicker structure, with an orientation equal to the mean of the contributing structures.

In concrete terms, if the spatial resolution is a distance, $d$, then two lines with an intervening angle $\theta$ can only be distinguished at a distance $L$ away from the intersection point, given by $L = \cot(\theta/2)\frac{d}{2}$.

### 4 Ridge response

The ridge response can be obtained by convolving the image and the filter using the Convolution Theorem:

$$\begin{aligned}
R(x, y, \phi_n; f_c, K, K_h) &= \mathcal{F}_{xy}^{-1}\left\{\hat{I}(f_x, f_y)\hat{\Phi}(f, \theta; \phi_n, f_c, K, K_h)\right\}(x, y) \\
&= I(x, y) * \Phi(x, y; \phi_n, f_c, K, K_h) \qquad (51) \\
&\text{for } \phi_n = \frac{n\pi}{2K_h + 1}, n = 0, \dots, 2K_h \\
c(k; x, y, f_c, K, K_h) &= \mathcal{F}_{\phi_n} R(x, y, \phi_n; f_c, K, K_h)(k) \qquad (52) \\
&= \sum_{n=-K_h}^{n=K_h} R(x, y, \phi_n; f_c, K, K_h) \exp(2i\phi_n k) \qquad (53) \\
&\text{for } k = -K_h, \dots, K_h
\end{aligned}$$

#### 4.1 Interpolating the response through steerability

We can then obtain the ridge filter response for any orientation $\phi$ per the steerable property of the filter by the inverse Fourier Transform:

$$R(x, y, \phi; f_c, K, K_h) = \sum_{k=-K_h}^{K_h} c(k; x, y, f_c, K) \exp(2i\phi k) \qquad (54)$$

#### 4.2 Relating the filter response at different K values via the heat-diffusion equation

As the filter response is scaled using a Gaussian, it is possible to relate the orientation filter responses for distinct values of the orientation resolution parameter K through the heat-diffusion partial differential equation. In its traditional physical formulation:

$$\frac{\partial \rho}{\partial t} = D \frac{\partial^2 \rho}{\partial \phi^2} \qquad (55)$$

In the original definition, $\rho, t, \phi,$ and $D$ represent a heat distribution function, time, space, and the diffusion coefficient, respectively. Here we take these parameters as the orientation filter response ($\rho$), a parameter related to resolution ($t$), angle ($\phi$), and a constant ($D$).

To develop the relationship, let $\rho$ be an orientation response at some position $(x, y)$ and at some orientation resolution determined by $K$, which is a function of $t$, and expand it into a finite Fourier series:

$$\rho(\phi, t) = R(x, y, \phi; f_c, K(t), K_h) \tag{56}$$

$$= \sum_{k=-K_h}^{k=K_h} c(k; x, y, f_c, K(t)) \exp(-2i\phi k) \tag{57}$$

$$= \sum_{k=-K_h}^{k=K_h} c(k; x, y, f_c, K_h) \frac{\hat{G}(k; K(t))}{\hat{G}(k; K_h)} \exp(-2i\phi k) \tag{58}$$

The last expression arises from the definition of the angular portion of the filter (Section 2.3) and from the Convolution Theorem, which states that

$$c(k; x, y, f_c, K) = c(k; x, y, f_c, K_h) \frac{\hat{G}(k; K(t))}{\hat{G}(k; K_h)} \tag{59}$$

To derive an expression for $K(t)$ (written as K for brevity), we compute the partial derivatives of $\rho$:

$$\frac{\partial \rho}{\partial \phi} = \sum_{k=-K_h}^{k=K_h} -2ikc(k; x, y, f_c, K_h) \frac{\hat{G}(k; K)}{\hat{G}(k; K_h)} \exp(-2i\phi k) \tag{60}$$

$$\frac{\partial^2 \rho}{\partial \phi^2} = \sum_{k=-K_h}^{k=K_h} -4k^2 c(k; x, y, f_c, K_h) \frac{\hat{G}(k; K)}{\hat{G}(k; K_h)} \exp(-2i\phi k) \tag{61}$$

$$\frac{\partial \rho}{\partial K} = \sum_{k=-K_h}^{k=K_h} \frac{\partial \hat{G}(k; K)}{\partial K} \frac{c(k; x, y, f_c, K_h)}{\hat{G}(k; K_h)} \exp(-2i\phi k) \tag{62}$$

$$= \sum_{k=-K_h}^{k=K_h} \frac{8k^2\pi^2}{(2K+1)^3} \hat{G}(k; K) \frac{c(k; x, y, f_c, K_h)}{\hat{G}(k; K_h)} \exp(-2i\phi k) \tag{63}$$

With the partial derivatives of $\rho$ in hand, we can substitute these into the heat-diffusion equation (55) in order to relate $K$, $t$, and $D$.

$$\frac{\partial \rho}{\partial t} = \frac{\partial \rho}{\partial K}\frac{dK}{dt} = D\frac{\partial^2 \rho}{\partial \phi^2} \tag{64}$$

$$\frac{1}{D}\frac{dK}{dt} = -\frac{(2K+1)^3}{2\pi^2} \tag{65}$$

$$\int \frac{dK}{(2K+1)^3} = -D\int \frac{dt}{2\pi^2} \tag{66}$$

$$\frac{1}{4}\frac{1}{(2K+1)^2} = \frac{D}{2\pi^2}t \tag{67}$$

$$\frac{1}{2}\frac{\pi^2}{(2K+1)^2} = Dt \tag{68}$$

With the relationship between $K$, $t$, and $D$ established, we can now infer an expression for $K(t)$ and the derivative of $K$ with respect to $t$, $\mathrm{d}K/\mathrm{d}t$, and vice-versa, $\mathrm{d}t/\mathrm{d}K$. Taking $D = \frac{1}{2}$ as a convenient choice (there is some ambiguity in the allocation of constants between $K(t)$ and $D$), it follows that:

$$t = \frac{\pi^2}{(2K+1)^2} = s_a^2, \quad K(t) = \frac{\pi}{2\sqrt{t}} - \frac{1}{2} \tag{69}$$

$$\frac{dK}{dt} = -\frac{\pi}{4t^{3/2}}, \quad \frac{dt}{dK} = \frac{-4\pi^2}{(2K+1)^3} \tag{70}$$

With $K(t)$ defined and an expression for $\mathrm{d}t/\mathrm{d}K$, we can now express the heat-diffusion equation (55) in terms of $K$:

$$\frac{\partial \rho}{\partial K} = \frac{\partial \rho}{\partial t}\frac{dt}{dK} = D\frac{dt}{dK}\frac{\partial^2 \rho}{\partial \phi^2} \tag{71}$$

$$= \frac{-2\pi^2}{(2K+1)^3}\frac{\partial^2 \rho}{\partial \phi^2} \tag{72}$$

This equation relates the curvature of the response curve with respect to $\phi$, $\partial^2\rho/\partial\phi^2$, to how the response curve will change with respect to orientation resolution as parameterized by $K$, $\partial\rho/\partial K$.

#### 4.3 Tracing and evaluating orientation local maxima using the heat-diffusion equation

Let $\phi_m(K)$ be an orientation local maximum, i.e.

$$\frac{\partial \rho}{\partial \phi}(\phi_m(K), K) = 0 \text{ and } \frac{\partial^2 \rho}{\partial \phi^2}(\phi_m(K), K) \leq 0 \tag{73}$$

Then

$$\frac{\mathrm{d}}{\mathrm{d}K}\left(\frac{\partial \rho}{\partial \phi}(\phi_m(K), K)\right) = \frac{\partial^2 \rho}{\partial \phi^2}\frac{d\phi_m}{dK} + \frac{\partial^2 \rho}{\partial \phi \partial K} \tag{74}$$

$$= \frac{\partial^2 \rho}{\partial \phi^2}\frac{d\phi_m}{dK} + D\frac{dt}{dK}\frac{\partial^3 \rho}{\partial \phi^3} = 0 \tag{75}$$

$$\frac{d\phi_m}{dK} = -D\frac{dt}{dK}\frac{\dfrac{\partial^3 \rho}{\partial \phi^3}}{\dfrac{\partial^2 \rho}{\partial \phi^2}} \tag{76}$$

The last equation is called Davidenko's equation and indicates how sensitive $\phi_m$ is to changes in $K$. Note that the partial derivatives of $\rho$ can be obtained from the finite Fourier series:

$$\frac{\partial^n \rho}{\partial \phi^n} = \sum_{k=-K_h}^{k=K_h} (-2ik)^n c(k; x, y, f_c, K_h)\frac{\hat{G}(k; K)}{\hat{G}(k; K_h)}\exp(-2i\phi k) \tag{77}$$

The $n$th derivative of $\phi_m$ with respect to $K$, $\mathrm{d}^n\phi_m/\mathrm{d}K^n$, can be found by expanding the $n$th derivative of $\partial\rho(\phi_m(K), K)/\partial\phi$ with respect to $K$ according to Faa di Bruno's formula. As an example, the second derivative with respect to $K$ of the first partial derivative with respect to $\phi$ yields the following expansion. We start by applying $\mathrm{d}/\mathrm{d}K$ across Equation 74.

$$\frac{\mathrm{d}^2}{\mathrm{d}K^2}\left(\frac{\partial \rho(\phi_m(K), K)}{\partial \phi}\right) = 0 \tag{78}$$

$$= \frac{\mathrm{d}}{\mathrm{d}K}\left(\frac{\partial^2 \rho}{\partial \phi^2}\right)\frac{\mathrm{d}\phi_m}{\mathrm{d}K} + \frac{\partial^2 \rho}{\partial \phi^2}\frac{\mathrm{d}^2\phi_m}{\mathrm{d}K^2} + D\frac{\mathrm{d}}{\mathrm{d}K}\left(\frac{\mathrm{d}t}{\mathrm{d}K}\frac{\partial^3 \rho}{\partial \phi^3}\right) \tag{79}$$

Then we isolate the term containing $\frac{\mathrm{d}^2\phi_m}{\mathrm{d}K^2}$ on the left side:

$$-\frac{\partial^2\rho}{\partial\phi^2}\frac{\mathrm{d}^2\phi_m}{\mathrm{d}K^2} = \frac{\mathrm{d}}{\mathrm{d}K}\left(\frac{\partial^2\rho}{\partial\phi^2}\right)\frac{\mathrm{d}\phi_m}{\mathrm{d}K} + D\frac{\mathrm{d}}{\mathrm{d}K}\left(\frac{\mathrm{d}t}{\mathrm{d}K}\frac{\partial^3\rho}{\partial\phi^3}\right) \tag{80}$$

$$= \left(\frac{\partial^3\rho}{\partial\phi^3}\frac{\mathrm{d}\phi_m}{\mathrm{d}K} + D\frac{\mathrm{d}t}{\mathrm{d}K}\frac{\partial^4\rho}{\partial\phi^4}\right)\frac{\mathrm{d}\phi_m}{\mathrm{d}K} + D\frac{\mathrm{d}^2t}{\mathrm{d}K^2}\frac{\partial^3\rho}{\partial\phi^3} + D\frac{\mathrm{d}t}{\mathrm{d}K}\frac{\partial^4\rho}{\partial\phi^4}\frac{\mathrm{d}\phi_m}{\mathrm{d}K} + D^2\left(\frac{\mathrm{d}t}{\mathrm{d}K}\right)^2\frac{\partial^5\rho}{\partial\phi^5} \tag{81}$$

$$\frac{\mathrm{d}^2\phi_m}{\mathrm{d}K^2} = -\left[2D\frac{\mathrm{d}t}{\mathrm{d}K}\frac{\partial^4\rho}{\partial\phi^4}\frac{\mathrm{d}\phi_m}{\mathrm{d}K} + \left(D\frac{\mathrm{d}^2t}{\mathrm{d}K^2} + \left(\frac{\mathrm{d}\phi_m}{\mathrm{d}K}\right)^2\right)\frac{\partial^3\rho}{\partial\phi^3} + D^2\left(\frac{\mathrm{d}t}{\mathrm{d}K}\right)^2\frac{\partial^5\rho}{\partial\phi^5}\right]\Big/\frac{\partial^2\rho}{\partial\phi^2} \tag{82}$$

In practice, we usually compute the derivatives with respect to $t$, and then convert them to derivatives with respect to $K$. With this, we determine for each identified orientation its value at its optimal $K$, defined as the $K$ at which the identified orientation changes the least with respect to $K$ (Fig. 1G, H; Fig. S3).

#### 4.4 Scope of application and generalization

The angular component of the filter design was chosen in part to facilitate the application of the heat-diffusion equation to relate orientation filter responses across orientation resolutions. Diffusion, however, only requires Gaussian scaling but does not require the initial filter to be based on a Gaussian (e.g. $\hat{G}(k; K_h)$). The initial filter as applied here is actually a truncated Gaussian as $\hat{G}(k; K_h) \neq 0$ for $|k| > K_h$. It is only approximately zero beyond the truncation point.

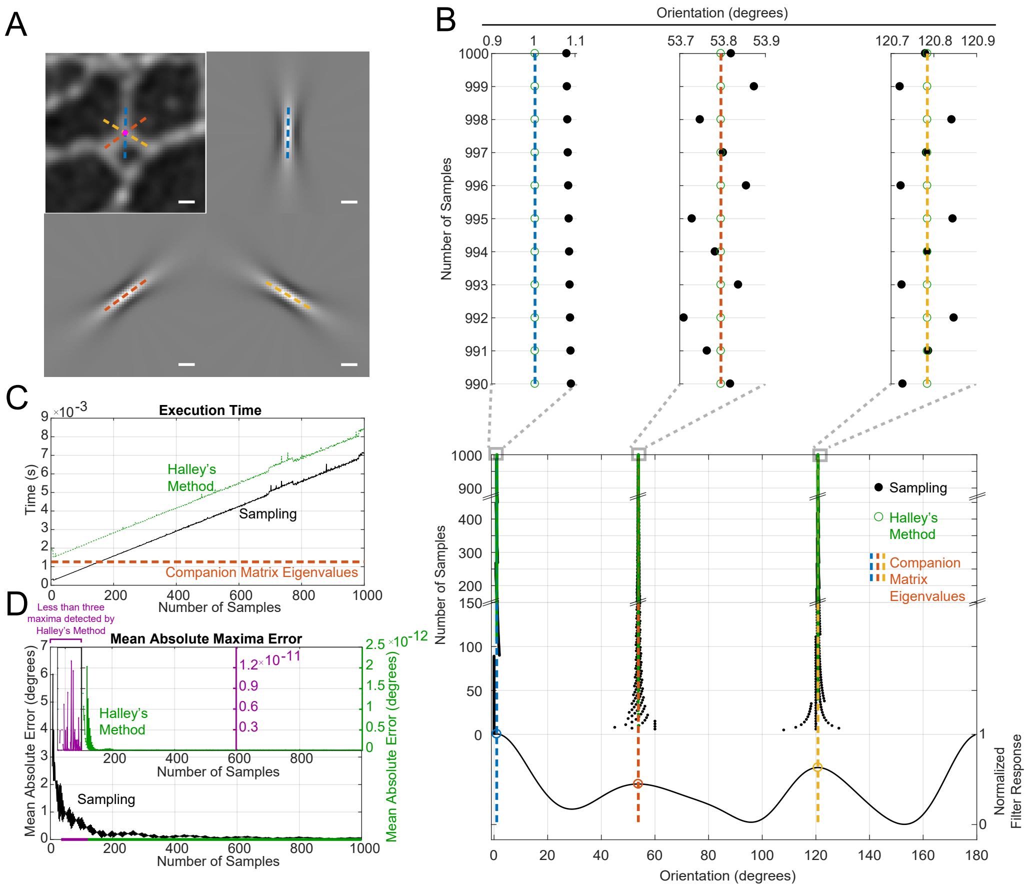

**Figure S2: Companion Matrix Eigenvalue Computation of Filter Local Maxima Versus Sampling.**

(A) Detected orientations overlaid on the original fluorescence image and the oriented $K = 8$ filters. Scale Bar is 1 $\mu$m. (B) Comparison of local maxima detection by sampling (black dots), sampling followed by Halley's method (green open circles), and the companion matrix eigenvalues employed in our work (dashed lines with colors corresponding to Panel A). Top zoom shows a comparison with 990 to 1000 samples. Curve at Bottom is the response curve using filter at $K = 8$. (C) Comparison of execution time between sampling (black), sampling followed by Halley's method (green), and calculating the eigenvalues of the companion matrix (red). (D) Mean absolute error in detecting maxima using sampling, sampling followed by Halley's method, and calculating the eigenvalues of the companion matrix. Purple lines indicate area where less than three orientations are detected by sampling followed by Halley's method.

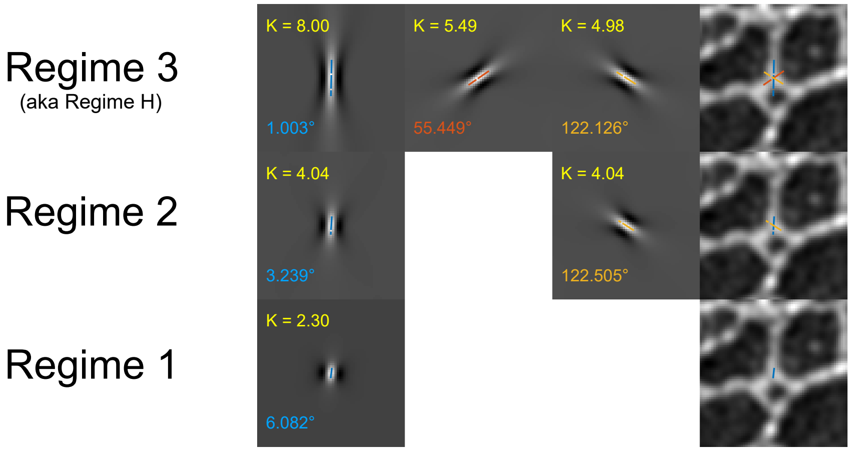

**Figure S3: Expanded Orientation Selection in Regimes.**
Regimes are numbered according to the number of maxima. The regime with the highest number of maxima is also designated Regime H. Left 3 columns: Orientations detected in each regime along with the oriented filters at the appropriate $K$. The values of the parameter $K$ are written in yellow in the top-left of each filter. The orientation (in degrees) of the maximum is written in the lower right in blue, red, or gold. Lines indicating the orientations are superimposed on the filter, with line length corresponding to the distance between half maximum values along the filter axis (equivalent to full width at half maximum, but in this case it is full length at half maximum). Right column: Lines indicating the detected orientations overlaid on the original image of Lamin A in a $Lmnb1^{-/-}$ mouse embyronic fibroblast, centered at the pixel being analyzed.

### 5 Finding the extrema and roots of finite Fourier series

#### 5.1 Companion matrix eigenvalue calculations

In this work, we detect orientations by finding the local maxima of the orientation filter response curve, $R$. To do so, we must find the roots of the first derivative with respect to orientation, $\frac{\partial R}{\partial \phi}$, and evaluate the higher derivatives.

Since a finite Fourier series is a trigonometric polynomial, the roots of the polynomial can be found by forming the companion matrix from the polynomial coefficients and solving for the eigenvalues [4]. We first restate the form of the filter response in terms of a finite Fourier series.

$$\text{Let z } = \exp\left(2i\phi\right), \text{ then } z^k = \exp(2i\phi k) \tag{83}$$

$$R(x, y, \phi; f_c, K, K_h) = \sum_{k=-K_h}^{K_h} c(k; x, y, f_c, K) \exp(2i\phi k) \tag{84}$$

$$= \sum_{k=-K_h}^{K_h} c(k; x, y, f_c, K) z^k \tag{85}$$

We then calculate the first derivative with respect to $\phi$ .

$$\frac{\partial R}{\partial \phi} = \sum_{k=-K_h}^{K_h} c(k; x, y, f_c, K) k z^{k-1} \frac{\partial z}{\partial \phi} \tag{86}$$

$$= \sum_{k=-K_h}^{K_h} c(k; x, y, f_c, K) 2ikz^k \tag{87}$$

$$= z^{-K_h} \sum_{k=0}^{2K_h} c(k - K_h; x, y, f_c, K) 2ikz^k \tag{88}$$

We then transform this into a more traditional polynomial form.

$$\text{Let } p(z(\phi)) = \frac{\partial R}{\partial \phi} z^{K_h} = \sum_{k=0}^{2K_h} c(k - K_h; x, y, f_c, K) 2ikz^k \tag{89}$$

$$\text{We want } \frac{\partial R}{\partial \phi} = 0 \tag{90}$$

$$\therefore \text{ we want } p(z(\phi)) = 0 \text{ such that } \|z\| = \|\exp(2i\phi)\| = 1 \tag{91}$$

We can then construct the companion matrix, $C$, with dimensions $(2K_h + 1) \times (2K_h + 1)$ as follows. Let

$$d(k) = 2ikc(k - K_h; x, y, f_c, K) \tag{92}$$

Then

$$C = \begin{bmatrix} 0 & \frac{d(1-K_h)}{d(-K_h)} & \frac{d(2-K_h)}{d(-K_h)} & \cdots & \frac{d(K_h-1)}{d(-K_h)} & \frac{d(K_h)}{d(-K_h)} \\ 1 & 0 & 0 & \cdots & 0 & 0 \\ 0 & 1 & 0 & & 0 & 0 \\ \vdots & 0 & \ddots & & 0 & 0 \\ 0 & 0 & 0 & & 1 & 0 \end{bmatrix} \tag{93}$$

Note that

$$p(z(\phi)) = \det(z\mathbf{I} - C) \tag{94}$$

Thus if we solve for the eigenvalues $\lambda_n$ of $C$ and select the eigenvalues such that $\|\lambda_n\| = 1$, then $\phi_m = -\frac{i}{2} \ln \lambda_n$ will be a root such that $p(z(\phi_m)) = 0$:

$$Cv = \lambda_n v \tag{95}$$

$$0 = \lambda_n v - Cv \tag{96}$$

$$0 = (\lambda_n \mathbf{I} - C)v \tag{97}$$

for $v \neq \mathbf{0}$

As a practical numerical matter we accept eigenvalues as roots such that $|\ln \|\lambda_n\|| < \epsilon$. $\epsilon$ depends on the precision of the numerical system [4].

In order to classify the type of extrema to which the roots of the derivative correspond, the second derivative and potentially higher roots may need to be computed:

$$\frac{\partial^n R}{\partial \phi^n}(\phi_m) = \sum_{k=-K_h}^{K_h} c(k; x, y, f_c, K)(2ik)^n z(\phi_m)^k \tag{98}$$

$$\text{If } \frac{\partial^2 R}{\partial \phi^2}(\phi_m) < 0, \text{ then } R(\phi_m) \text{ is a local maximum of } R. \tag{99}$$

$$\begin{aligned} \text{If } \frac{\partial^2 R}{\partial \phi^2}(\phi_m) = 0, \text{ then } \phi_m \text{ is a local maximum if and only if} \\ \frac{\partial^n R}{\partial \phi^n}(\phi_m) < 0, \text{ n is even,} \\ \text{and } \frac{\partial^q R}{\partial \phi^q}(\phi_m) = 0 \, \forall \, q \in [2, n) \end{aligned} \tag{100}$$

#### 5.2 Sampling

For performance evaluation, we compare the companion matrix approach described above to a simple sampling approach, in which case extrema or roots are found by querying values of a function at regular intervals and then identifying which one of those queries was greater than the query to the left or right of it. While simple in approach, the method is inaccurate for small numbers of samples and may miss local maxima that occur in the interval between samples. We use Horner's method to evaluate the Fourier series as a trigonometric polynomial at regular intervals to evaluate the sampling approach [5].

#### 5.3 Householder's methods for refinement of maxima

To improve upon the sampling method, we also refine the local maxima found by sampling using an iterative root finding method, initializing the routine with the solution from the sampling approach. Householder's methods are a class of iterative root finding methods that are commonly employed. The first order method is commonly called Newton's method. Here we apply the second order method, Halley's method:

$$\phi_{n+1} = \phi_n - \frac{\partial\rho(\phi_n)/\partial\phi \; \partial^2\rho(\phi_n)/\partial\phi^2)}{(\partial^2\rho(\phi_n)/\partial\phi^2)^2 - \partial\rho(\phi_n)/\partial\phi \; \partial^3\rho(\phi_n)/\partial\phi^3 \; /2} \tag{101}$$

Note that the third partial derivative with respect to $\theta$ is readily accessible via Equation 77. To evaluate the performance of sampling combined with Halley's method, a single round of iteration was performed.

The comparison between the three approaches in terms of ability to find local maxima and computational time is shown in Fig. S2.

### 6 Minimal Bridging Algorithm

Our minimal bridging algorithm combines AR-NLMS outputs from three combinations of orientations and responses at different orientation resolutions (Section 6.1), in order to balance orientation resolution and spatial localization. To achieve this, it employs the mean orientation response (MOR) (Section 6.2) and a complexity-based attenuated MOR (AMOR) (Section 6.3) to threshold the different AR-NLMS outputs and mask the image region of interest, and finally two rounds of minimal bridging to achieve a parsimonious segmentation of both lines and junctions (Section 6.4). The final output is intended to resemble non-maximum suppression outputs produced by prior algorithms [6]. It is in essence a response-weighted segmentation that can be subsequently thresholded as needed to yield a binary segmentation of the lines and junctions of interest.

#### 6.1 The three AR-NLMS branches

The three AR-NLMS branches employed in the minimal bridging algorithm arise from the following orientation and response pairings:

1. Regime 0 orientations combined with response at medium orientation resolution ($K_m$, usually taken as 3; red branch in Fig. S4 and in Fig. S5).

   Regime 0 orientations come from the asymptotic regime, the limit as $K$ approaches -0.5. They are the most spatially localized, and there is typically only a single orientation (i.e. response maximum) at each point in space. As $K \to -0.5$, the location of the maximum asymptotically approaches the phase angle of the 1st Fourier coefficient. Therefore, Regime 0 orientations are determined by evaluating the phase of the 1st Fourier coefficient, which is readily retrieved after evaluation of the discrete Fourier transform. Thus the Regime 0 orientations are calculated without significant processing.

Combining Regime 0 orientations with the medium orientation resolution response allows for determination of a single local orientation for simple neighborhoods within the image, where a single line or curve might exist. In a sense, this orientation is a weighted average of all the orientations that may exist around a pixel. This baseline orientation information is used as the starting point for the determination of more complex structures using the minimal bridging algorithm (Section 6.4) and information from AR-NLMS Branches 2 and 3 described next.

2. Regime H orientations combined with response at medium orientation resolution (green branch in Fig. S4 and in Fig. S6).

   Regime H orientations are from the highest regime, where the number of maxima matches the number of maxima initially detected at the highest orientation resolution used ($K_h$, typically taken as 8), but with potentially increased spatial localization because of $K$ adaptation within the regime (Section 4.2). Combining Regime H orientations with the medium orientation resolution response balances the conflicting needs of both high resolution orientation information and spatial localization.

3. Regime 0 and Regime H orientations combined with response at the highest used orientation resolution ($K_h$, usually taken as 8; blue branch in Fig. S4 and in Fig. S6).

   The last branch uses the previously identified orientations in Regime H and Regime 0, but now evaluates these orientations with the highest resolution filter response values at $K_h$. As $K_h$ is equal to or greater than the $K$ values associated with each identified orientation, this combination offers another opportunity to integrate information across orientation resolutions, thus enabling the complete segmentation of junctions where the other two branches might fail (Fig. S9). The benefits of this branch are threefold.

   First, the longer aspect ratio of the filter at $K_h$ makes the response at $K_h$ more robust to noise in the image since more pixels are considered. For example, while the Regime 0 orientations are the most local, whether these are spatial local maxima that survive NLMS is susceptible to the noise of the surrounding pixels. Thus examining these local orientations again using a long filter allows us to evaluate if the local orientation is supported by a persistent pattern across space along the orientation of interest.

Second, using a long filter allows for junctions to be resolved in space by evaluating areas along each orientation where the underlying intensities of each structure radiating from the junction are more separated. For acute angles, individual curvilinear segments may be blurred together near the junction due to the lack of spatial resolution. Because the filter at $K_h$ is long, this helps to locate the junction by using information further away in space where the component curvilinear segments may be resolvable in space.

Third, the use of a long filter mitigates the issue of the radial component of the filter not being well matched to the width of structures near junctions, as they might get wider because of spatial overal (See Box S2). In this work, we focus on orientation resolution alone with fixed selectivity for width as determined by the radial component of the filter. Interpreted locally by a short filter at lower $K$, the filter response may be attenuated due to the width mismatch. With a long filter the matching width away from the junction can be incorporated into the segmentation process.

Overall, combining maxima detected using the adaptive resolution process with the highest resolution response level assists in the segmentation of the area near junctions where the issue of spatial resolution may otherwise interfere with the segmentation process. Away from junctions the above attributes become a liability because the response values have poor spatial localization and may suggest the inclusion of pixels in the segmentation that are not supported by the local image intensity. However, because the long filter response is used solely for bridging between lines revealed by Branches 1 and 2 above, this branch is applied mainly near junctions where it is most informative, and the effect of lack of spatial localization is kept to a minimum.

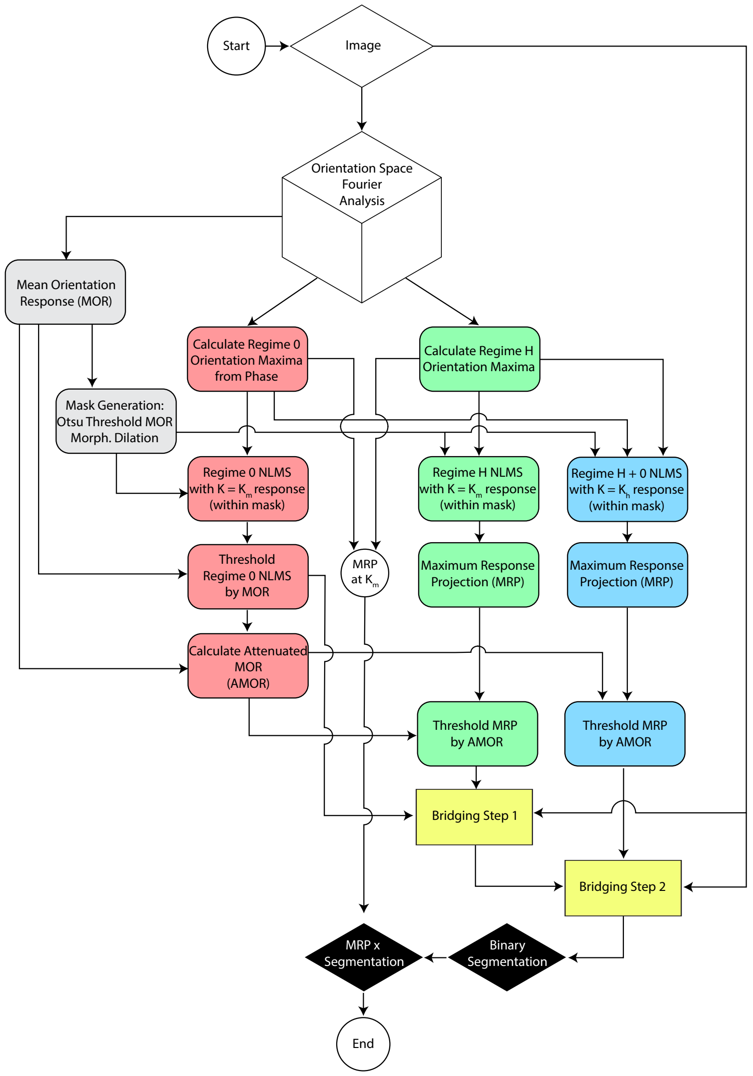

**Figure S4: Segmentation Flow Chart.**
The flow chart starts with the construction of orientation space by convolving the input image with the orientation space filters. Mean, local, and multiple orientations are calculated and combined with response curves at different $K$ values. $K_h$ is the highest $K$ used for initial detection of orientations. $K_m$ is the middle $K$ value used to obtain a local orientation filter response. The final output is a response-weighted segmentation of lines and junctions, which can be subsequently thresholded for a final segmentation of the lines and junctions of interest. See Section 6 for details.

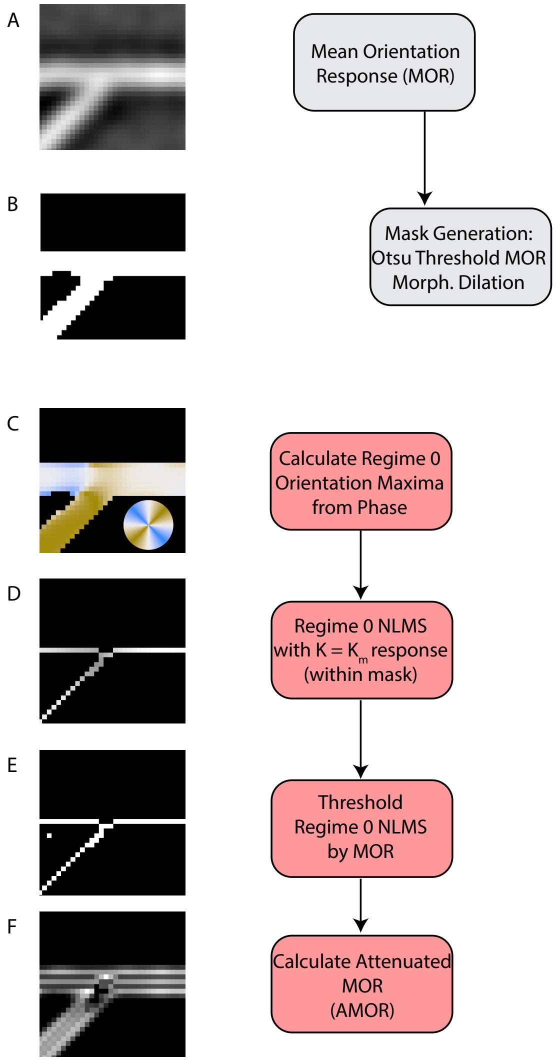

**Figure S5: Example of Low Resolution Orientation Analysis.**
(A) Mean orientation filter response (MOR) reflecting the response to the radial component of the filter in Fourier space. (B) Mask generated by thresholding the MOR using Otsu's algorithm. (C) Regime 0 orientations calculated from phase, drawn as lines color coded by direction as shown in color wheel. (D) NLMS using Regime 0 orientations and the filter response at $K_m = 3$, restricted to the mask in B. (E) Binarization of D by thresholding with MOR. (F) Attenuated MOR calculated from the MOR and 8-connected neighborhood occupancy from (E). See Section 6.3 for details.

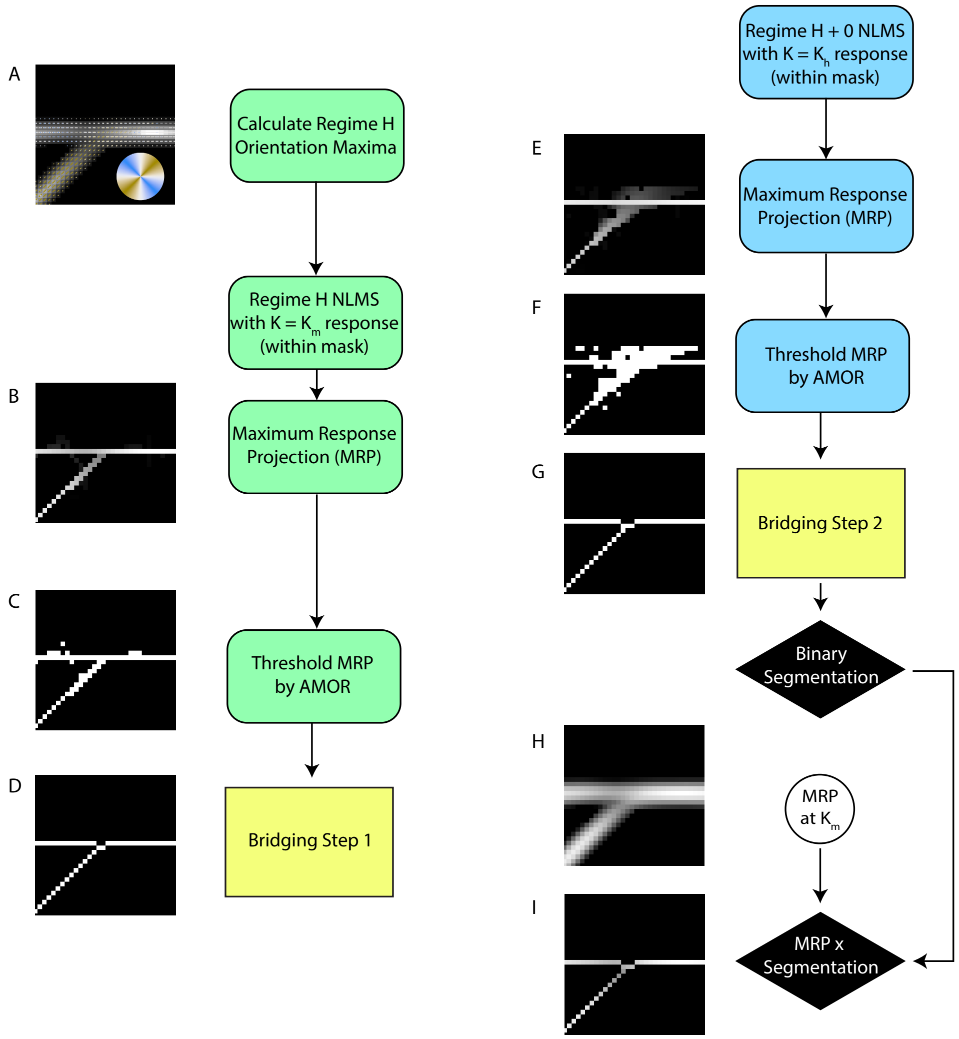

**Figure S6: Examples of High Resolution Orientation Analysis.**
(A) Regime H orientations, drawn as lines color coded by direction as shown in color wheel. (B) Maximum response projection (MRP) of NLMS using Regime H orientations and filter response at $K_m = 3$. (C) Binarization of the MRP in B using the AMOR in Fig. S5F. (D) The result of bridging Step 1 (using the Regime 0 NLMS (Fig. S5E) as left input and binarized MRP in C as top input. (E) MRP of the NLMS using Regime H+0 orientations and filter response at $K_h = 8$. (F) Binarization of the MRP in E using the AMOR in Fig. S5F. (G) The result of bridging Step 2 (using the output of bridging Step 1 as left input and binarized MRP in F as top input). (H) MRP of the filter response at $K = K_m$ over the orientations identified in Regime H and Regime 0. (I) Pixel-wise product of the response MRP in H and the binary from G to produce the response-weighted segmentation (final output).

#### 6.2 Mean orientation response

The mean orientation response (MOR) is evaluated from the 0th Fourier coefficient of the filter response and is independent of the choice of $K$. The independence from $K$ reflects that the MOR corresponds to the response of the image to the radial component of the filter alone, and contains no orientation information. It is calculated immediately upon evaluation of the discrete Fourier transform of the filter response. Based upon the MOR, Otsu's algorithm is employed to automatically determine a threshold used to generate a mask defining the region of interest for further orientation analysis (Fig. S5A,B). The MOR is also used to segment the AR-NLMS of Branch 1 (Regime 0 orientations with $K_m$ response), yielding the initial segmentation of "simple" lines, which serves as the starting point for minimal bridging and complete segmentation (Fig. S5C-E).

#### 6.3 Attenuated mean orientation response

The initial segmentation using Branch 1 AR-NLMS and MOR allows us to estimate the potential complexity of the neighborhood of each pixel. Specifically, for regions with multiple orientations, it often yields many not-suppressed pixels next to each other, while for regions with truly a single orientation it yields a 1-pixel-thin segmentation. Therefore, we use the thresholded Branch 1 AR-NLMS to lower the thresholds for the Branch 2 and Branch 3 AR-NLMS outputs, by calculating an attenuated MOR (AMOR) (Fig. S5F). This is necessary because intersecting lines interfere with each other's filter responses, and because complex areas may have increased thickness due the convergence of many lines, which may reduce the filter response due to mismatch with the scale (i.e. width) of the filter.

To calculate the AMOR, the level of attenuation for each pixel is proportional to the number of pixels in its eight connected neighborhood that are contained in the Branch 1 segmentation, divided by four. If more than four neighboring pixels are within the segmentation, then the value of the AMOR at the center pixel is set to zero (effectively no threshold at that pixel).

#### 6.4 Minimal bridging

The goal of the minimal bridging algorithm is to incorporate the multi-resolution information of the three AR-NLMS branches described above into a complete segmentation. Along the way, it also produces several intermediate segmented structures that could be useful in different applications.

Our overall workflow (Fig. S4) employs two steps of bridging (Fig. S7, Fig. S8). In Step 1, the low orientation resolution segmentation of Branch 1 (Regime 0 orientations, response at $K_m$) is bridged with higher orientation resolution information from Branch 2 (Regime H orientations, response at $K_m$). In Step 2, the output of Bridging Step 1 is bridged further with even higher orientation resolution information from Branch 3 (Regime H+0 orientations, response at $K_h$). Below we describe the input and workflow of the bridging algorithm.

##### 6.4.1 Input

The minimal bridging algorithm requires three inputs (Fig. S7).

1. The parsimonious *low-res binary*, which is the initial lower orientation resolution segmentation serving as the starting point for bridging.

2. The full *high-res binary*, which provides the candidate pixels for bridging between the segments of the parsimonious *low-res binary*. These candidate pixels are generated by orientation analysis at a higher resolution than that used for initial segmentation.

3. The *original image*, which is used for calculating the cost of adding bridges, in order to decide on which bridges to add. The costs are taken as the complement of the grey scale values of the original image.

The lower part of Fig. S4 shows the specific inputs to the bridging algorithm for the two steps of bridging.

##### 6.4.2 Workflow

1. Segments are derived from the *low-res binary* by removing morphological branch points and sharp arrows from it. Sharp arrows are removed by changing the center pixel from 1 to 0 when the 3x3 neighborhood around it appears as the following pattern or its 90 degree rotations.

   | | | |
   |---|---|---|
   | 0 | 0 | 1 |
   | 0 | 1 | 0 |
   | 0 | 0 | 1 |

   The segments are then labeled as individual 8-connected components.

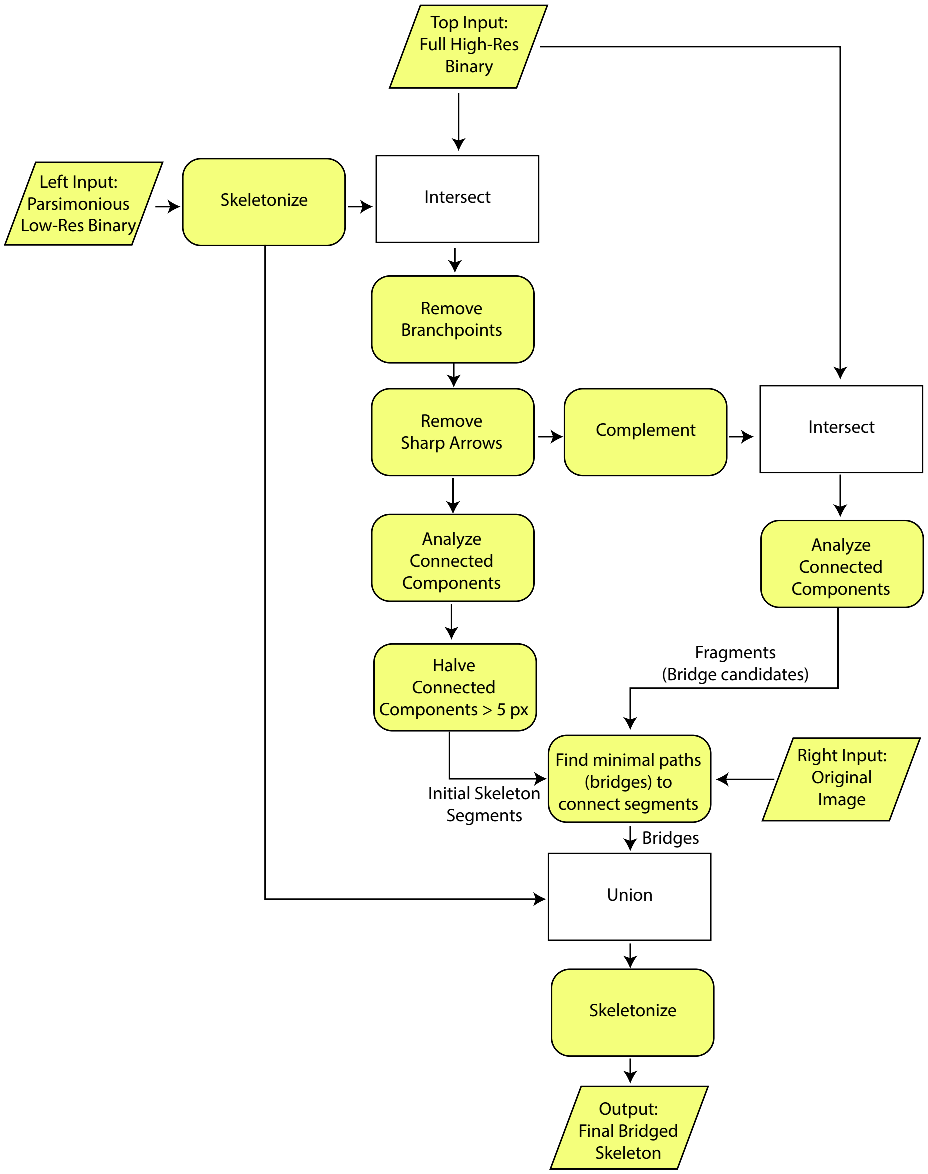

**Figure S7: Minimal Bridging Algorithm Flow Chart.**
Rhomboids indicate input or output. Yellow rectangles with rounded corners represent processes. White rectangles represent set operations. See Section 6.4 for details.

2. Long segments exceeding five pixels in length are divided in half into two separate segments by ordering the pixels by connectivity from endpoint to endpoint and assigning the last half of the pixels to a separate connected component index. This produces the initial skeleton, which does not contain any junctions (Fig. S8C,I).

3. The pixels in the *high-res binary* that are not part of the segments obtained from the *low-res binary* (as described above) are considered fragments and are used as candidate pixels for bridges to connect the *low-res binary* segments. Each fragment is also labeled as an individual 8-connected component (Fig. S8D,J).

4. The minimal bridging algorithm then proceeds by iterating over each fragment, finding the neighboring segments to each fragment, and then finding the path with minimal cost through the fragment for each pair of neighboring segments. The path cost is calculated by using the grey distance along an inverted map of the original image according to a chessboard distance metric [7] relative to each segment. The candidate bridge, a path with the minimal cost between two segments, is identified by adding together the distance transforms relative to each segment in a pair, finding the smallest value in the sum of distance transforms, and taking the path as all pixels where the sum of the distance transforms is within 0.001 of the smallest value [8]. To resolve cases where there are more than two segments bordering a fragment, a graph is constructed with a node for each segment and edges between the nodes corresponding to the candidate bridges. A minimum spanning tree is then calculated to select a set of bridges such that all the neighboring segments are connected with bridges at the least cost.

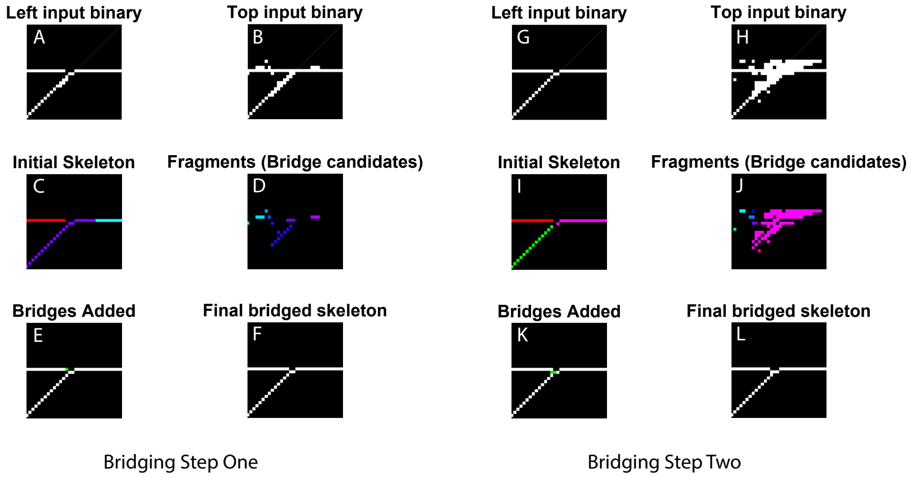

**Figure S8: Minimal Bridging Examples.**
(A-F) Example of Bridging Step 1. (G-L) Example of Bridging Step 2. (A,G) Left-input binary, a parsimonious segmentation from lower resolution orientation analysis. (B,H) Top-input binary, a superset of the final segmentation, from a higher resolution orientation analysis. (C,I) Initial skeleton, derived from the left-input binary. Colors represent distinct connected components (”segments”). (D,J) Fragments for bridging, from top-input binary. Colors indicate distinct connected components. (E,K) Initial skeleton in white with added bridge pixels in green. (F,L) Final bridged skeleton, which is the skeletonization of the images in (E,K).

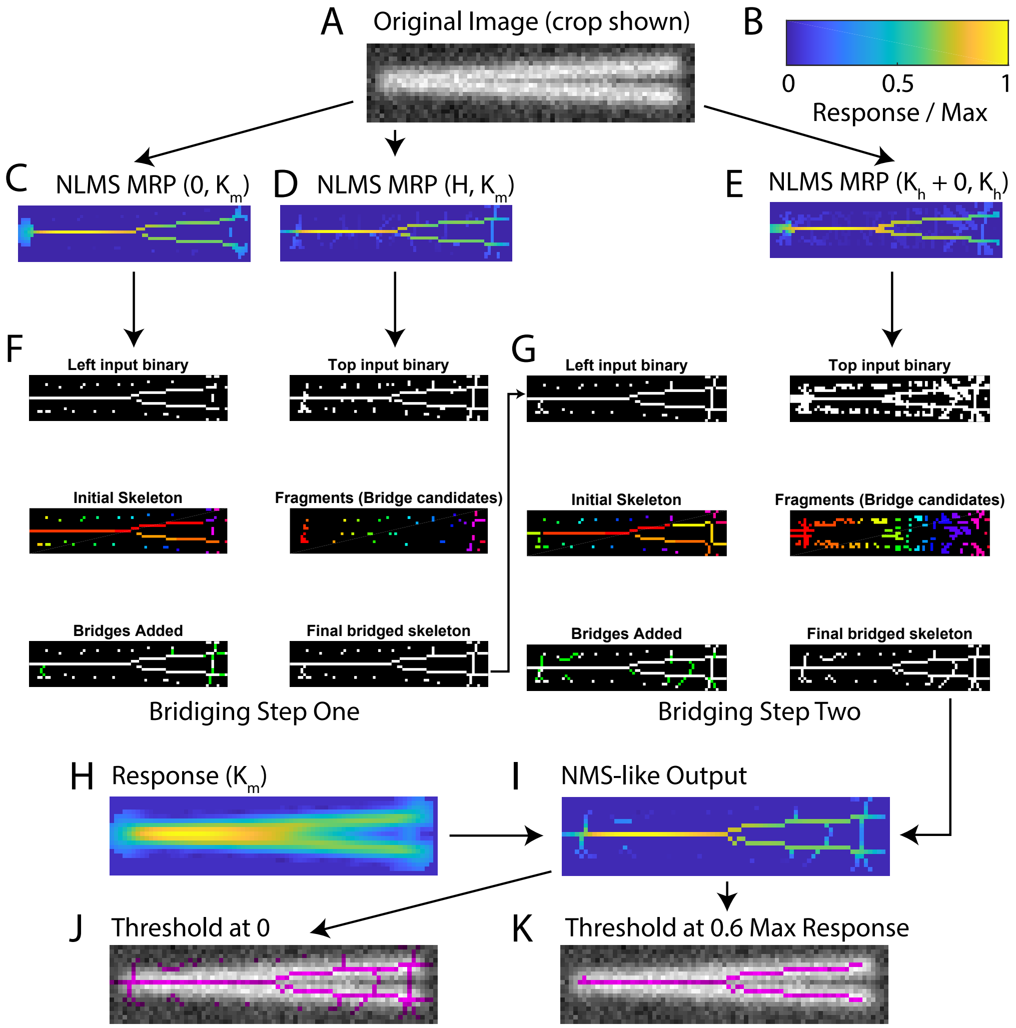

**Figure S9: Application of AR-NLMS and Minimal Bridging: Acute Angle Intersection Example.**
(A) Original synthetic junction with an acute angle of six degrees. (B) Normalized response color bar used for C,D,E, H, and I. (C) MRP of NLMS for Regime 0 orientations combined with $K_m$ response (red branch in Fig. S4). (D) MRP of NLMS for Regime H orientations combined with $K_m$ response (green branch in Fig. S4). (E) MRP of NLMS for Regimes H + 0 orientations combined with $K_h$ response (blue branch in Fig. S4). (F-G) Bridging steps one and two, respectively, as in Fig. S8). (H) MRP of the filter response at $K = K_m$ over the orientations identified in Regime H and Regime 0 (similar to Fig. S6H). *Continued on the next page ...*

**Figure S9: Application of AR-NLMS and Minimal Bridging: Acute Angle Intersection Example (cont.)** (I) Pixel-wise product of the response MRP in H and the output of Bridging Step 2 in G to produce the response-weighted segmentation (final output). (J) Binarization of I with a threshold of 0 (magenta), overlaid on the original image in greyscale. (K) Same as J but with a higher segmentation threshold to eliminate weak, spurious lines.

### 7 Performance evaluation

To evaluate the performance of our line and junction segmentation approach, we tested the algorithm on synthetic junctions by evaluating (1) whether the junction was detected, (2) the location of the detected junction relative to the ground truth junction, and (3) the orientations of the lines at the detected junction relative to their ground truth orientations (colored lines in example images in Fig. S10).

#### 7.1 Generation of synthetic junction images

Three sets of synthetic junctions were generated:

- Symmetric junctions involving two orientations (Fig. S10A-I; Movies S3, S4)
- Asymmetric junctions involving three orientations (Fig. S10J-Q; Movies S5, S6)
- Junctions of two arcs involving two orientations local to the junction (Fig. S10R-Y; Movies S7, S8)

The synthetic junctions were drawn by calculating the smallest distance of each pixel from one of the abstract lines or arcs. Intensities were assigned to each pixel according to a normalized Gaussian function with a standard deviation of two pixels. This procedure of subpixel accuracy was designed to avoid rastering an intermediate binary image to the pixel followed by convolution with a Gaussian kernel, which could skew the results. Gaussian noise was then added to the image using the built-in MATLAB function imnoise. The signal to noise ratio was calculated as the ratio between the maximum intensity above background and the standard deviation of the added Gaussian noise.

#### 7.2 Evaluation procedure

For each junction type, we varied the intervening angles between the lines drawn, repeating each set of intervening angles ten times. This was done for two signal-to-noise ratios: 5 and 10. From the NMS-like output, we obtained a binary, pixel-based segmentation by thresholding the output with zero. We then evaluated the segmentation based on the location of the morphological branch point (”detected junction”) relative to the ground junction location (Fig. S10I,Q,Y), and the orientations identified at the detected junction in comparison to the ground truth orientations (Fig. S10F-H,N-P,V-X). In addition, to separate orientation detection from branch point detection, we also measured the orientation at the location of the ground truth junction (Movies S3-S8).

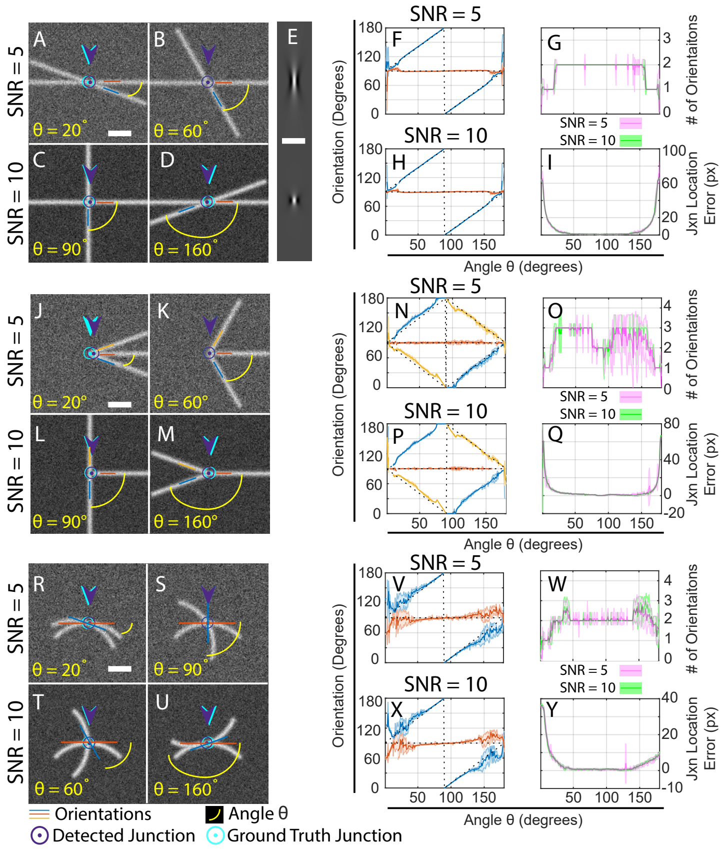

**Figure S10: Orientation and Junction Detection Performance.** (A-D) Examples of symmetric two-line junctions with indicated angles and SNR (5 in top row, 10 in bottom row). (E) Ridge filters with $K = 8$ (top) and $K = 3$ (bottom). (F-I) Performance analysis, as captured by comparing the detected orientations with the ground truth orientations in F (SNR 5) and H (SNR 10), the number of orientations at the detected junction in G (SNR 5 in pink, SNR 10 in green), and the distance between the detected junction and the ground truth junction in I (same color coding as in G). The results are the mean (dark lines) $\pm$ standard deviation (shaded area around dark lines) over 10 tests per angle and SNR. Because junction detection is based on morphological criteria (branchpoints), it is possible to have a detected junction with only 1 orientation in G. (J-Q) Examples of asymmetric three-line junctions and associated performance; image description as in A-D and performance description as in F-I. These are the same results shown in Fig. 2A-I, repeated here to place them in the context of the other performance tests. *Continued on the next page ...*

**Figure S10: Orientation and Junction Detection Performance (cont.)** (R-Y) Examples of intersecting arcs and associated performance; image description as in A-D and performance description as in F-I. Scale bars in A, E, J and R are applicable to all images and indicate that the synthetic images and the filter in E are shown at the same scale. Further performance analysis is shown in Movies S3-S8.

### 8 Movie Legends

Movie S1: A scan through orientation space showing detected maxima in individual orientation planes. Each frame of the movie is a plane in orientation space showing the filter response at the indicated orientation, with brighter pixels reflecting a higher filter response. Magenta lines are drawn at the the filter response maxima with the lines oriented accordingly, and the line length corresponding to the filter response magnitude. Left panel shows a top-down view of each 2D plane, and magenta lines are shown if they are within 15 degrees of the plane orientation. Right panel shows the 2D plane in the context of 3D orientation space.

Movie S2: Non-local maxima suppression (NLMS), as applied to the response and maxima detected in Movie S1. In NLMS, non-maximum suppression (NMS) is applied per orientation plane. Only magenta lines that survive NMS in each orientation plane are shown.

Movies S3-S8: Various validation examples, all displayed in the same manner. Top-left: Synthetic test image with blue text and arcs illustrating the inscribed angle. Top-middle: MRP of AR-NLMS using Regime H maxima and response at $K = 3$. Top-right: Final output of minimal bridging algorithm. Middle row: Orientations detected at the detected junction. Green open circles and black dots indicate the orientations detected at the current frame and at all previous frames, respectively. Blue dashed lines show the ground truth orientations. Bottom row: Same as the middle row except the orientations are detected at the ground truth junction and the orientations detected at the current frame are shown as yellow open circles. The specific validation examples are described next.

Movie S3: A sequence of symmetric junctions at inscribed angles from 0 to 180 degrees with two lines passing through the center at a SNR of 5.

Movie S4: A sequence of symmetric junctions at inscribed angles from 0 to 180 degrees with two lines passing through the center at a SNR of 10.

Movie S5: A sequence of asymmetric junctions at inscribed angles from 0 to 180 degrees with three radial lines projecting from the center at a SNR of 5.

Movie S6: A sequence of asymmetric junctions at inscribed angles from 0 to 180 degrees with three radial lines projecting from the center at a SNR of

10.

Movie S7: A sequence of junctions of two arcs at inscribed angles from 0 to 180 degrees with two lines passing through the center at a SNR of 5.

Movie S8: A sequence of junctions of two arcs at inscribed angles from 0 to 180 degrees with two lines passing through the center at a SNR of 10.

### References

[1] Unser, M. & Chenouard, N. A unifying parametric framework for 2d steerable wavelet transforms. *Siam Journal on Imaging Sciences* **6**, 102–135 (2013).

[2] Ginkel, M. v. *Image analysis using orientation space based on steerable filters*. Diss., technische universiteit delft, 2002., Technische Universiteit Delft (2002).

[3] Olkin, I., Gleser, L. J. & Derman, C. Probability models and applications. Tech. Rep., World Scientific (1980).

[4] Boyd, J. P. Computing the zeros, maxima and inflection points of chebyshev, legendre and fourier series: solving transcendental equations by spectral interpolation and polynomial rootfinding. *Journal of Engineering Mathematics* **56**, 203–219 (2006).

[5] Hildebrand, F. B. *Introduction to numerical analysis* (Dover Publications, New York, 1987), 2nd ed.. edn.

[6] Jacob, M. & Unser, M. Design of steerable filters for feature detection using canny-like criteria. *Ieee Transactions on Pattern Analysis and Machine Intelligence* **26**, 1007–1019 (2004).

[7] Soille, P. Generalized geodesy via geodesic time. *Pattern Recognition Letters* **15**, 1235–1240 (1994).

[8] Soille, P. *Morphological Image Analysis: Principles and Applications* (Springer, 2010).
