## Supplementary material for "Adaptive-Resolution Multi-Orientation Analysis of Complex Filamentous Network Images": MATLAB code from Github: bwRemoveBranchPoints.m

```
function [ bw, count, cumBranchPoints ] = bwRemoveBranchPoints( bw, n )
%BWREMOVEBRANCHPOINTS Remove all branch points
%
% INPUT
% bw - logical image, any non-logical image will be converted using logical
% n  - (optional) Number of times to remove branchpoints using bwmorph
%      default: Inf (until no more branchpoints are found)
%
% OUTPUT
% bw
% count
% branchPoints
%
% Copyright (C) 2019, Jaqaman Lab - UT Southwestern, Goldman Lab - Northwestern 
%
% This file is part of AdaptiveResolutionOrientationSpace.
% 
% AdaptiveResolutionOrientationSpace is free software: you can redistribute it and/or modify
% it under the terms of the GNU General Public License as published by
% the Free Software Foundation, either version 3 of the License, or
% (at your option) any later version.
% 
% AdaptiveResolutionOrientationSpace is distributed in the hope that it will be useful,
% but WITHOUT ANY WARRANTY; without even the implied warranty of
% MERCHANTABILITY or FITNESS FOR A PARTICULAR PURPOSE.  See the
% GNU General Public License for more details.
% 
% You should have received a copy of the GNU General Public License
% along with AdaptiveResolutionOrientationSpace.  If not, see <http://www.gnu.org/licenses/>.
% 
% 

if(nargin < 2)
    n = Inf;
end

done = false;
count = 0;
if(nargout > 2)
    cumBranchPoints = false(size(bw));
end

while(~done)
    branchPoints = bwmorph(bw,'branchpoints');
    bw = bw & ~branchPoints;
    done = ~any(branchPoints(:));
    count = count + 1;
    if(nargout > 2)
        cumBranchPoints = cumBranchPoints | branchPoints;
    end
    if(count > n)
        break;
    end
end


end
```
