## Supplementary material for "Adaptive-Resolution Multi-Orientation Analysis of Complex Filamentous Network Images": MATLAB code from Github: bwRemoveSharpArrows.m

```
function [ bw, sharpArrows ] = bwRemoveSharpArrows( bw )
%bwRemoveSharpArrows Remove 8-connected sharp arrows
%
% INPUT
% bw - binary image
%
% OUTPUT
% bw - binary image without sharp arrows
% sharpArrows - location of sharp arrows
%
% Sharp arrows are
% [ 1 0 0
%   0 1 0
%   1 0 0]
% [ 1 0 1
%   0 1 0
%   0 0 0]
% [ 0 0 0
%   0 1 0
%   1 0 1]
% [ 0 0 1
%   0 1 0
%   0 0 1]
%
% Copyright (C) 2019, Jaqaman Lab - UT Southwestern, Goldman Lab - Northwestern 
%
% This file is part of AdaptiveResolutionOrientationSpace.
% 
% AdaptiveResolutionOrientationSpace is free software: you can redistribute it and/or modify
% it under the terms of the GNU General Public License as published by
% the Free Software Foundation, either version 3 of the License, or
% (at your option) any later version.
% 
% AdaptiveResolutionOrientationSpace is distributed in the hope that it will be useful,
% but WITHOUT ANY WARRANTY; without even the implied warranty of
% MERCHANTABILITY or FITNESS FOR A PARTICULAR PURPOSE.  See the
% GNU General Public License for more details.
% 
% You should have received a copy of the GNU General Public License
% along with AdaptiveResolutionOrientationSpace.  If not, see <http://www.gnu.org/licenses/>.
% 
% 

dbw = double(logical(bw));
neighborCount = imfilter(dbw,[1 1 1; 1 0 1; 1 1 1]);
forwardSlashCount = imfilter(dbw,eye(3));
backwardSlashCount = imfilter(dbw,rot90(eye(3)));
xCount = imfilter(dbw,[1 0 1; 0 0 0; 1 0 1]);

sharpArrows = neighborCount == 2 & forwardSlashCount ~= 3 & backwardSlashCount ~= 3 & xCount == 2;
bw(sharpArrows) = 0;

end
```
