## Supplementary material for "Adaptive-Resolution Multi-Orientation Analysis of Complex Filamentous Network Images": MATLAB code from Github: minimalBridge.m

```
function [ bridges, allway] = minimalBridge( fragment_cc, segment_cc, I )
%minimalBridge Find parsimonious connections over fragments that connect
%segments as guided by image intensity
%
% INPUT
% fragment_cc - connected components structure representing fragments
% segment_cc  - connected components structure representing segments
% I - image or response map
%
% OUTPUT
% bridges - a minimum subset of fragment_cc pixels that connect segment_cc such
% that segment_cc is well connected
%
% Copyright (C) 2019, Jaqaman Lab - UT Southwestern, Goldman Lab - Northwestern 
%
% This file is part of AdaptiveResolutionOrientationSpace.
% 
% AdaptiveResolutionOrientationSpace is free software: you can redistribute it and/or modify
% it under the terms of the GNU General Public License as published by
% the Free Software Foundation, either version 3 of the License, or
% (at your option) any later version.
% 
% AdaptiveResolutionOrientationSpace is distributed in the hope that it will be useful,
% but WITHOUT ANY WARRANTY; without even the implied warranty of
% MERCHANTABILITY or FITNESS FOR A PARTICULAR PURPOSE.  See the
% GNU General Public License for more details.
% 
% You should have received a copy of the GNU General Public License
% along with AdaptiveResolutionOrientationSpace.  If not, see <http://www.gnu.org/licenses/>.
% 
% 

%% Image map processing
% Suppress negative pixels
I(I < 0) = 0;
% Take the complement such that high intensity pixels have low cost
I = imcomplement(mat2gray(double(I)));
I = max(I,1e-3);


import bridging.*;

% Dilate fragments individually to detect which segments are adjacent to
% each fragment
fragment_dilated_cc = connectedComponents.ccDilate(fragment_cc,ones(3));
segment_label = labelmatrix(segment_cc);
fragment_label = labelmatrix(fragment_cc);
% Bridges will be a binary map consisting of a subset of fragment_cc pixels
bridges = false(fragment_cc.ImageSize);

% Do a run length encoding decompression on fragment_dilated_cc
% tic;
% lengths = cellfun('length',fragment_dilated_cc.PixelIdxList);
% cumLengths = cumsum(lengths);
% idx = [1 cumLengths(1:end-1)+1];
% ccIdx = zeros(cumLengths(end),1);
% ccIdx(idx) = 1;
% ccIdx = cumsum(ccIdx);
% 
% % Find which segments are associated with each fragment
% segmentIdx = segment_label(vertcat(fragment_dilated_cc.PixelIdxList{:}));
% s = segmentIdx ~= 0;
% ccIdx = ccIdx(s);
% segmentIdx = segmentIdx(s);
% segmentsPerFragment = accumarray(ccIdx,segmentIdx,[],@(x) {unique(x)});
% toc;

rp = regionprops(fragment_dilated_cc,segment_label,'PixelValues');
[rp.unzPixelValues] = dealcell( arrayfun(@(x) unique(x.PixelValues(x.PixelValues ~= 0)),rp,'UniformOutput',false) );

s = cellfun('length',{rp.unzPixelValues});
s = s > 1 & s < 32;
fragment_cc = connectedComponents.ccFilter(fragment_cc,s);
fragment_dilated_cc = connectedComponents.ccFilter(fragment_dilated_cc,s);
rp = rp(s);

rp2 = regionprops(fragment_dilated_cc,'PixelList','BoundingBox');
% [rp.PixelList] = deal(rp2.PixelList);
% [rp.BoundingBox] = deal(rp2.BoundingBox);
% clear rp2;

[rp.RelativePixelList] = dealcell( arrayfun(@(x) x.PixelList-x.BoundingBox(1:2)+0.5,rp2,'UniformOutput',false) );


bridges = calculateBridges(fragment_cc,fragment_dilated_cc,rp,segment_label,I);

% % Loop over each fragment
% tic
% parfor f = 1:fragment_cc.NumObjects
%     % L is true when the pixels are in fragment f
%     L = fragment_label == f;
%     % Obtain the list of segments adjacent to fragment f
%     segments = unique(segment_label(fragment_dilated_cc.PixelIdxList{f}));
%     % Don't include the background
%     segments = segments(segments ~= 0);
%     % Determine the geodesic distance along the fragment from each segment
%     segment_dist = cell(1,length(segments));
%     for s = 1:length(segments)
%         % Lseg is true when the segment 
%         Lseg = segment_label == segments(s);
% %         segment_dist{s} = bwdistgeodesic(L | Lseg, Lseg,'quasi-euclidean');
%         A = I;
%         % The cost of traveling outside the fragment or segment is infinite
%         % See bwdistgeodesic source code
%         A(~L & ~Lseg) = Inf;
%         % chessboard: The distance for diagonals is the same as up or down
%         segment_dist{s} = graydist(A, Lseg,'chessboard');
%     end
%     % If fragment is adjacent to more than one segment, then ...
%     if(length(segments) > 1)
%         % Measure the distance along the fragment between each pair of
%         % segments
%         % Determine number of segment pairs
%         np = nchoosek(length(segments),2);
%         % Store the connection between each segment
%         pair_bridge = cell(1,np);
%         % For each pair, determine the distance of the smallest path
%         mindist = zeros(1,np);
%         % Counter to track pair number
%         p = 0;
%         % Loop over each pair of segments
%         for sx = 1:length(segments)-1
%             for sy = sx+1:length(segments)
%                 p = p + 1;
%                 % Minimum path distance will be revealed by the sum of the
%                 % distances from each segment in the pair
%                 D = segment_dist{sx}+segment_dist{sy};
%                 % Round to the nearest 1000th
% %                 D = ceil(D*1000)/1000;
%                 % Distance between each segment is the minimum of the
%                 % summed distance
%                 mindist(p) = nanmin(D(:));
%                 % The bridge is the path through the fragment where the
%                 % summed distance is equal the minimum
%                 pair_bridge{p} = abs(D - mindist(p)) < 1e-3;
%             end
%         end
%         % Find the minimum spanning tree
%         % That is the tree with the minimum distance weight that connects
%         % all the segmments
%         min_span_tree = graphminspantree(sparse(squareform(mindist)));
%         % Map square indices to pair number
%         ind2p = squareform(1:np);
%         % Select the bridges that provide the minimal connectivity
%         pair_bridge = pair_bridge(ind2p(min_span_tree > 0));
% %         pair_bridge = pair_bridge(ind2p(find(min_span_tree)));
%         
%         bridges = bridges | any(cat(3,pair_bridge{:},zeros(size(I))),3);
%     end
% end
% toc


end

function bridges = calculateBridges(fragment_cc,fragment_dilated_cc,rp,segment_label,I)
    bridges = false(fragment_cc.ImageSize);

%     rp2 = regionprops(fragment_dilated_cc,'PixelList','BoundingBox');
    rp2 = regionprops(fragment_dilated_cc,'BoundingBox');
    % [rp.PixelList] = deal(rp2.PixelList);
    % [rp.BoundingBox] = deal(rp2.BoundingBox);
    % clear rp2;

%     [rp2.RelativePixelList] = dealcell( arrayfun(@(x) x.PixelList-x.BoundingBox(1:2)+0.5,rp2,'UniformOutput',false) );
    
    rp_undilated = regionprops(fragment_cc,'Image','BoundingBox');
    for i=1:length(rp2)
%         rp_undilated(i).PaddedImage(~rp_undilated(i).Image) = Inf;
        L = padarray(rp_undilated(i).Image,[1 1],false);
        
        % Padding may over extend the image boundary
        % Remove pixels that are outside of the image
        Lrs = (1:(rp_undilated(i).BoundingBox(4)+2))+rp_undilated(i).BoundingBox(2)-0.5-1; 
        Lcs = (1:(rp_undilated(i).BoundingBox(3)+2))+rp_undilated(i).BoundingBox(1)-0.5-1;
        Lrs = Lrs > 0 & Lrs <= fragment_cc.ImageSize(1);
        Lcs = Lcs > 0 & Lcs <= fragment_cc.ImageSize(2);
        L = L(Lrs,Lcs);
        
        segments = rp(i).unzPixelValues;
        rs = (1:rp2(i).BoundingBox(4))+rp2(i).BoundingBox(2)-0.5;
        cs = (1:rp2(i).BoundingBox(3))+rp2(i).BoundingBox(1)-0.5;
        A = I(rs,cs);
        % Determine the geodesic distance along the fragment from each segment
        segment_dist = cell(1,length(segments));
        all_segments = ismember(segment_label(rs,cs),segments);
        A(all_segments) = 0;
        for s = 1:length(segments)
            % Lseg is true when the segment 
            Lseg = segment_label(rs,cs) == segments(s);
            if(any(size(Lseg) ~= size(L)))
                Lseg = L;
            end
    %         segment_dist{s} = bwdistgeodesic(L | Lseg, Lseg,'quasi-euclidean');
            D = A;
            % The cost of traveling outside the fragment or segment is infinite
            % See bwdistgeodesic source code
            % Allow paths through all segments in case segments touch
            D(~L & ~all_segments) = Inf;
            %D(~L & ~Lseg) = Inf
            % chessboard: The distance for diagonals is the same as up or down
            segment_dist{s} = graydist(D, Lseg,'chessboard');
        end
    
        % Measure the distance along the fragment between each pair of
        % segments
        % Determine number of segment pairs
        np = nchoosek(length(segments),2);
        % Store the connection between each segment
        pair_bridge = cell(1,np);
        % For each pair, determine the distance of the smallest path
        mindist = zeros(1,np);
        % Counter to track pair number
        p = 0;
        % Loop over each pair of segments
        for sx = 1:length(segments)-1
            for sy = sx+1:length(segments)
                p = p + 1;
                % Minimum path distance will be revealed by the sum of the
                % distances from each segment in the pair
                D = segment_dist{sx}+segment_dist{sy};
                % Round to the nearest 1000th
    %                 D = ceil(D*1000)/1000;
                % Distance between each segment is the minimum of the
                % summed distance
                mindist(p) = nanmin(D(:));
                % The bridge is the path through the fragment where the
                % summed distance is equal the minimum
                pair_bridge{p} = abs(D - mindist(p)) < 1e-3;
            end
        end
        % Find the minimum spanning tree
        % That is the tree with the minimum distance weight that connects
        % all the segmments
        min_span_tree = graphminspantree(sparse(squareform(mindist)));
        % Map square indices to pair number
        ind2p = squareform(1:np);
        % Select the bridges that provide the minimal connectivity
        pair_bridge = pair_bridge(ind2p(min_span_tree > 0));
    %         pair_bridge = pair_bridge(ind2p(find(min_span_tree)));

        bridges(rs,cs) = bridges(rs,cs) | any(cat(3,pair_bridge{:},false(size(L))),3);
    end
end
```
