## Supplementary material for "Adaptive-Resolution Multi-Orientation Analysis of Complex Filamentous Network Images": MATLAB code from Github: minimalBridgingFromBinary.m

```
function [bridgedSkeleton, bridgingStruct, diagFig] = minimalBridgingFromBinary(I,full_binary,consensus_binary,minLength,diagnosticMode,boundingBox)
% minimalBridgingFromBinary
%
% Copyright (C) 2019, Jaqaman Lab - UT Southwestern, Goldman Lab - Northwestern 
%
% This file is part of AdaptiveResolutionOrientationSpace.
% 
% AdaptiveResolutionOrientationSpace is free software: you can redistribute it and/or modify
% it under the terms of the GNU General Public License as published by
% the Free Software Foundation, either version 3 of the License, or
% (at your option) any later version.
% 
% AdaptiveResolutionOrientationSpace is distributed in the hope that it will be useful,
% but WITHOUT ANY WARRANTY; without even the implied warranty of
% MERCHANTABILITY or FITNESS FOR A PARTICULAR PURPOSE.  See the
% GNU General Public License for more details.
% 
% You should have received a copy of the GNU General Public License
% along with AdaptiveResolutionOrientationSpace.  If not, see <http://www.gnu.org/licenses/>.
% 
% 
   
    if(nargin < 4)
        minLength = 5;
    end
    if(nargin < 5)
        diagnosticMode = false;
    end

    %% Do minimal bridging
    % nlms_binary = nlms_highest_mip_binary;
    % nlms_binary = nlms_highest_mip > 0;

    %% Morphologically skeletonize consensus binary
    consensus_binary_skel = bwmorph(consensus_binary,'skel',Inf);
    
    %% Binary processing of segments
    segments = consensus_binary_skel.* full_binary;
    segments = bridging.bwRemoveBranchPoints(segments);
    segments = bridging.bwRemoveSharpArrows(segments);

    %% Processing of fragments
    fragments = full_binary & ~segments;
    fragments = bwconncomp(fragments);

    %% Connected component processing of segments
    segments = bwconncomp(segments);
    segments = connectedComponents.orderPixelIdxList(segments);
    segments = connectedComponents.halveEachComponent(segments,minLength);
    
    if(segments.NumObjects > 1)

        %% Perform minimal bridging algorithm
        bridges = bridging.minimalBridge(fragments,segments,I);
        bridges = bridges & labelmatrix(segments) == 0;

        %% Combine bridges and consensus skeleton
        bridgedSkeleton = consensus_binary_skel | bridges;
        % Ensure final product is morphologically skeletonized
        bridgedSkeleton = bwmorph(bridgedSkeleton,'skel',Inf);

    else
        bridgedSkeleton = consensus_binary_skel;
        bridges = false(size(consensus_binary_skel));
        
    end
    
    if(diagnosticMode)
        diagFig = figure('Name','bridgedSkeleton ');
        imshow(bridgedSkeleton,[]);
        title(diagFig.Children,diagFig.Name, ...
            'Interpreter','none');
        if(nargin > 5)
            xlim([0 boundingBox(3)]+boundingBox(1));
            ylim([0 boundingBox(4)]+boundingBox(2));
        end
        
        diagFig(2) = figure('Name','bridgedSkeleton Detail ');
        subplot(3,2,1);
        imshow(consensus_binary,[]);
        title('Left input binary');
        if(nargin > 5)
            xlim([0 boundingBox(3)]+boundingBox(1));
            ylim([0 boundingBox(4)]+boundingBox(2));
        end
        
        subplot(3,2,2);

        imshow(full_binary,[]);
        title('Top input binary');
        if(nargin > 5)
            xlim([0 boundingBox(3)]+boundingBox(1));
            ylim([0 boundingBox(4)]+boundingBox(2));
        end
        
        subplot(3,2,3);
        imshow(label2rgb(labelmatrix(segments),hsv(segments.NumObjects+1),'k'),[]);
%         title('Segments');
        title('Initial Skeleton');
        if(nargin > 5)
            xlim([0 boundingBox(3)]+boundingBox(1));
            ylim([0 boundingBox(4)]+boundingBox(2));
        end
        
        subplot(3,2,4);
        imshow(label2rgb(labelmatrix(fragments),hsv(fragments.NumObjects+1),'k'),[]);
        title('Fragments (Bridge candidates)');
        if(nargin > 5)
            xlim([0 boundingBox(3)]+boundingBox(1));
            ylim([0 boundingBox(4)]+boundingBox(2));
        end
        
        subplot(3,2,5);
        imshowpair(consensus_binary_skel | bridges, consensus_binary_skel & ~bridges);
%         imshow(bridges,[]);
        title('Bridges Added');
%         title('Bridges');
        if(nargin > 5)
            xlim([0 boundingBox(3)]+boundingBox(1));
            ylim([0 boundingBox(4)]+boundingBox(2));
        end
        
        subplot(3,2,6);
        imshow(bridgedSkeleton,[]);
%         title('Bridged skeleton');
        title('Final bridged skeleton');
        if(nargin > 5)
            xlim([0 boundingBox(3)]+boundingBox(1));
            ylim([0 boundingBox(4)]+boundingBox(2));
        end
        
    end
    
    if(nargout > 1)
        bridgingStruct.full_binary = full_binary;
        bridgingStruct.consensus_binary = consensus_binary;
        bridgingStruct.segments = segments;
        bridgingStruct.fragments = fragments;
        bridgingStruct.bridges = bridges;
        bridgingStruct.bridgedSkeleton = bridgedSkeleton;
    end
    
    
end
```
