## Supplementary material for "Adaptive-Resolution Multi-Orientation Analysis of Complex Filamentous Network Images": MATLAB code from Github: ccDilate.m

```
function dilatedCC = ccDilate(cc,se)
    import connectedComponents.*;
    if(isa(se,'strel'))
        se = se.getnhood;
    end
    
%     cc.PixelIdxList = distributed(cc.PixelIdxList);
%
% Copyright (C) 2019, Jaqaman Lab - UT Southwestern, Goldman Lab - Northwestern 
%
% This file is part of AdaptiveResolutionOrientationSpace.
% 
% AdaptiveResolutionOrientationSpace is free software: you can redistribute it and/or modify
% it under the terms of the GNU General Public License as published by
% the Free Software Foundation, either version 3 of the License, or
% (at your option) any later version.
% 
% AdaptiveResolutionOrientationSpace is distributed in the hope that it will be useful,
% but WITHOUT ANY WARRANTY; without even the implied warranty of
% MERCHANTABILITY or FITNESS FOR A PARTICULAR PURPOSE.  See the
% GNU General Public License for more details.
% 
% You should have received a copy of the GNU General Public License
% along with AdaptiveResolutionOrientationSpace.  If not, see <http://www.gnu.org/licenses/>.
% 
% 
    
    dilatedCC = ccPad(cc,size(se));
    offsets = mat2offset(dilatedCC.ImageSize,se);
%     test = dilatedCC.PixelIdxList;
%     test = distributed(test);

    dilatedCC.PixelIdxList = cellfun(@applyOffsets,dilatedCC.PixelIdxList,'UniformOutput',false);
%     for i=1:cc.NumObjects
%         dilatedCC.PixelIdxList{i} = bsxfun(@plus,dilatedCC.PixelIdxList{i},offsets);
%         dilatedCC.PixelIdxList{i} = unique(dilatedCC.PixelIdxList{i});
%         dilatedCC.PixelIdxList{i} = dilatedCC.PixelIdxList{i}(:);
%     end
    dilatedCC = ccUnpad(dilatedCC);
%     dilatedCC.PixelIdxList = gather(dilatedCC.PixelIdxList);
    function offetPixelIdx = applyOffsets(pixelIdx)
        offetPixelIdx = bsxfun(@plus,pixelIdx,offsets);
        offetPixelIdx = unique(offetPixelIdx);
        offetPixelIdx = offetPixelIdx(:);
    end
end
```
