## Supplementary material for "Adaptive-Resolution Multi-Orientation Analysis of Complex Filamentous Network Images": MATLAB code from Github: ccFilter.m

```
function [ cc, oldccidx ] = ccFilter( cc, filter )
%ccFilter Filter a connected components structure
%
% cc is a connected components structure from bwconncomp
% filter is a vector representing a subindex into PixelIdxList
%        used to select the new connected components. 
%        This may be a logical index or one selecting specific
%        components of the connected components structure
%
% output:
%   newcc is a new conneccted component struccture with filtere applied
%   oldccidx is a vector containing numeric indices of the original
%     connected component struccture. Useful if logical indexing was used.
%
% Copyright (C) 2019, Jaqaman Lab - UT Southwestern, Goldman Lab - Northwestern 
%
% This file is part of AdaptiveResolutionOrientationSpace.
% 
% AdaptiveResolutionOrientationSpace is free software: you can redistribute it and/or modify
% it under the terms of the GNU General Public License as published by
% the Free Software Foundation, either version 3 of the License, or
% (at your option) any later version.
% 
% AdaptiveResolutionOrientationSpace is distributed in the hope that it will be useful,
% but WITHOUT ANY WARRANTY; without even the implied warranty of
% MERCHANTABILITY or FITNESS FOR A PARTICULAR PURPOSE.  See the
% GNU General Public License for more details.
% 
% You should have received a copy of the GNU General Public License
% along with AdaptiveResolutionOrientationSpace.  If not, see <http://www.gnu.org/licenses/>.
% 
% 

% newcc = cc;
cc.PixelIdxList = cc.PixelIdxList(filter);
cc.NumObjects = length(cc.PixelIdxList);
if(nargout > 1)
    idx = 1:cc.NumObjects;
    oldccidx = idx(filter);
end

end
```
