## Supplementary material for "Adaptive-Resolution Multi-Orientation Analysis of Complex Filamentous Network Images": MATLAB code from Github: ccPad.m

```
function padded = ccPad(cc,padSize)
    padded = cc;
    padded.ImageSize = cc.ImageSize + padSize*2;
    ccImageSize = cc.ImageSize;
    paddedImageSize = padded.ImageSize;
    padded.PixelIdxList = cellfun(@padPixels,cc.PixelIdxList,'UniformOutput',false);
%     for i=1:cc.NumObjects
%         [r,c] = ind2sub(cc.ImageSize , cc.PixelIdxList{i});
%         r = r + padSize(1);
%         c = c + padSize(2);
%         padded.PixelIdxList{i} = sub2ind(padded.ImageSize , r, c);
%     end
%
% Copyright (C) 2019, Jaqaman Lab - UT Southwestern, Goldman Lab - Northwestern 
%
% This file is part of AdaptiveResolutionOrientationSpace.
% 
% AdaptiveResolutionOrientationSpace is free software: you can redistribute it and/or modify
% it under the terms of the GNU General Public License as published by
% the Free Software Foundation, either version 3 of the License, or
% (at your option) any later version.
% 
% AdaptiveResolutionOrientationSpace is distributed in the hope that it will be useful,
% but WITHOUT ANY WARRANTY; without even the implied warranty of
% MERCHANTABILITY or FITNESS FOR A PARTICULAR PURPOSE.  See the
% GNU General Public License for more details.
% 
% You should have received a copy of the GNU General Public License
% along with AdaptiveResolutionOrientationSpace.  If not, see <http://www.gnu.org/licenses/>.
% 
% 
    padded.Pad = padSize;
    function paddedPixels = padPixels(pixelIdx)
        [r,c] = ind2sub(ccImageSize, pixelIdx);
        r = r + padSize(1);
        c = c + padSize(2);
        paddedPixels = sub2ind(paddedImageSize , r, c);
    end
end
```
