## Supplementary material for "Adaptive-Resolution Multi-Orientation Analysis of Complex Filamentous Network Images": MATLAB code from Github: ccUnpad.m

```
function padded = ccUnpad(cc,padSize)
    if(nargin < 2)
        padSize = cc.Pad;
    end
    padded = cc;
    padded.ImageSize = cc.ImageSize - padSize*2;
    ccImageSize = cc.ImageSize;
    paddedImageSize = padded.ImageSize;
    padded.PixelIdxList= cellfun(@unpadPixels,cc.PixelIdxList,'UniformOutput',false);
%     for i=1:cc.NumObjects
%         [r,c] = ind2sub(cc.ImageSize , cc.PixelIdxList{i});
%         r = r - padSize(1);
%         c = c - padSize(2);
%         f = r > 0 & ...
%             c > 0 & ...
%             r <= padded.ImageSize(1) & ...
%             c <= padded.ImageSize(2);
%         padded.PixelIdxList{i} = sub2ind(padded.ImageSize , r(f), c(f));
%     end
%
% Copyright (C) 2019, Jaqaman Lab - UT Southwestern, Goldman Lab - Northwestern 
%
% This file is part of AdaptiveResolutionOrientationSpace.
% 
% AdaptiveResolutionOrientationSpace is free software: you can redistribute it and/or modify
% it under the terms of the GNU General Public License as published by
% the Free Software Foundation, either version 3 of the License, or
% (at your option) any later version.
% 
% AdaptiveResolutionOrientationSpace is distributed in the hope that it will be useful,
% but WITHOUT ANY WARRANTY; without even the implied warranty of
% MERCHANTABILITY or FITNESS FOR A PARTICULAR PURPOSE.  See the
% GNU General Public License for more details.
% 
% You should have received a copy of the GNU General Public License
% along with AdaptiveResolutionOrientationSpace.  If not, see <http://www.gnu.org/licenses/>.
% 
% 
    padded = rmfield(padded,'Pad');
    function unPaddedPixels = unpadPixels(pixelIdx)
        [r,c] = ind2sub(ccImageSize , pixelIdx);
        r = r - padSize(1);
        c = c - padSize(2);
        f = r > 0 & ...
            c > 0 & ...
            r <= paddedImageSize(1) & ...
            c <= paddedImageSize(2);
        unPaddedPixels = sub2ind(paddedImageSize , r(f), c(f));
    end
end
```
