## Supplementary material for "Adaptive-Resolution Multi-Orientation Analysis of Complex Filamentous Network Images": MATLAB code from Github: halveEachComponent.m

```
function [ cc ] = halveEachComponent( cc, minLength )
%halveEachComponent Split each component into two components at the
%midpoint
%
% Use orderPixelIdxList first
%
% Copyright (C) 2019, Jaqaman Lab - UT Southwestern, Goldman Lab - Northwestern 
%
% This file is part of AdaptiveResolutionOrientationSpace.
% 
% AdaptiveResolutionOrientationSpace is free software: you can redistribute it and/or modify
% it under the terms of the GNU General Public License as published by
% the Free Software Foundation, either version 3 of the License, or
% (at your option) any later version.
% 
% AdaptiveResolutionOrientationSpace is distributed in the hope that it will be useful,
% but WITHOUT ANY WARRANTY; without even the implied warranty of
% MERCHANTABILITY or FITNESS FOR A PARTICULAR PURPOSE.  See the
% GNU General Public License for more details.
% 
% You should have received a copy of the GNU General Public License
% along with AdaptiveResolutionOrientationSpace.  If not, see <http://www.gnu.org/licenses/>.
% 
% 

if(nargin < 2)
    minLength = 5;
end

nElements = cellfun('prodofsize',cc.PixelIdxList);
splitMe = nElements >= minLength;
halved = arrayfun(@(x,n) {x{1}(1:n) x{1}(n+1:end)},cc.PixelIdxList(splitMe),ceil(nElements(splitMe)/2),'UniformOutput',false);
if(isempty(halved))
    return;
else
    halved = [halved{:} cc.PixelIdxList{~splitMe}];
end

cc.PixelIdxList = halved;
cc.NumObjects = length(halved);


end
```
