## Supplementary material for "Adaptive-Resolution Multi-Orientation Analysis of Complex Filamentous Network Images": MATLAB code from Github: orderPixelIdxList.m

```
function [ cc ] = orderPixelIdxList( cc )
%orderPixelIdxList orders the pixel indices so that they are sorted by the
%distance from a particular endpoint of each connected component
%
% This is most effective if each connected component only has two endpoints
% in which the case the distance will be monotonic. If branches exist then
% the result is less interpretable.
%
% Copyright (C) 2019, Jaqaman Lab - UT Southwestern, Goldman Lab - Northwestern 
%
% This file is part of AdaptiveResolutionOrientationSpace.
% 
% AdaptiveResolutionOrientationSpace is free software: you can redistribute it and/or modify
% it under the terms of the GNU General Public License as published by
% the Free Software Foundation, either version 3 of the License, or
% (at your option) any later version.
% 
% AdaptiveResolutionOrientationSpace is distributed in the hope that it will be useful,
% but WITHOUT ANY WARRANTY; without even the implied warranty of
% MERCHANTABILITY or FITNESS FOR A PARTICULAR PURPOSE.  See the
% GNU General Public License for more details.
% 
% You should have received a copy of the GNU General Public License
% along with AdaptiveResolutionOrientationSpace.  If not, see <http://www.gnu.org/licenses/>.
% 
% 

    lm = labelmatrix(cc);
    bw = lm > 0;
    % sort PixelIdxList along the edges
    endpts = find(bwmorph(bw,'endpoints'));
    % the startpt will be indixed by the connected component index and will
    % be referenced by it's linear index
    edges_startpt = zeros(1,cc.NumObjects);
    % identify one end point per connected component as the start point
    edges_startpt(lm(endpts)) = endpts;
    % use the geodesic distance to measure the distance along the connected
    % components
    geo = bwdistgeodesic(bw,edges_startpt(edges_startpt ~= 0));
    I = cc.PixelIdxList;
    % sort according to the geodesic distance along the components
    I = cellfun(@(x) sortrows([geo(x) x]),I,'UniformOutput',false);
    I = cellfun(@(x) x(:,2),I,'UniformOutput',false);
    cc.PixelIdxList = I;
end
```
