## Supplementary material for "Adaptive-Resolution Multi-Orientation Analysis of Complex Filamentous Network Images": MATLAB code from Github: alignExtrema.m

```
function [ aligned, events ] = alignExtrema( extrema , period, unwrap, truncate)
%alignExtrema Align extrema tracks as K decreases (time increases)
%
% Copyright (C) 2019, Jaqaman Lab - UT Southwestern, Goldman Lab - Northwestern 
%
% This file is part of AdaptiveResolutionOrientationSpace.
% 
% AdaptiveResolutionOrientationSpace is free software: you can redistribute it and/or modify
% it under the terms of the GNU General Public License as published by
% the Free Software Foundation, either version 3 of the License, or
% (at your option) any later version.
% 
% AdaptiveResolutionOrientationSpace is distributed in the hope that it will be useful,
% but WITHOUT ANY WARRANTY; without even the implied warranty of
% MERCHANTABILITY or FITNESS FOR A PARTICULAR PURPOSE.  See the
% GNU General Public License for more details.
% 
% You should have received a copy of the GNU General Public License
% along with AdaptiveResolutionOrientationSpace.  If not, see <http://www.gnu.org/licenses/>.
% 
% 

if(nargin < 2 || isempty(period))
    period = 2*pi;
end
if(nargin < 3)
    unwrap = false;
end
if(nargin < 4)
    truncate = true;
end

useK = false;
if(ndims(extrema) == 3)
    K = extrema(:,:,2);
    extrema = extrema(:,:,1);
    [aligned,K] = sortMatrices(extrema,K,1);
    useK = true;
else
    % aligned = sort(extrema,'ComparisonMethod','real');
    aligned = sort(extrema);  
end


nExtrema = sum(~isnan(aligned),1);
totalExtrema = max(nExtrema);
aligned = aligned(1:totalExtrema,:);

currentCost = nansum(abs(diff(aligned,1,2)));

% Detect change in number of extrema and crossings of the periodic boundary
events = diff(nExtrema) ~= 0;
events = events | currentCost > period - currentCost;
if(useK)
    events = events | any(abs(diff(K,1,2)) > 1);
end
events = find(events);
% TODO: trigger alignment event if total difference is large due to extrema
% heading over periodic boundary
% events = 1:length(nExtrema)-1;

for e = events
    cost = abs(bsxfun(@minus,real(aligned(:,e).'),real(aligned(:,e+1))));

%     wrap = cost > period;
%     cost(wrap) = mod(cost(wrap),period);
    cost = min(abs(period-cost),cost);
    if(useK)
        cost = cost + abs(bsxfun(@minus,real(K(:,e).'),real(K(:,e+1))));
    end
    max_cost = max(cost(:));
    if(isnan(max_cost))
        % Cost matrix is all NaN
    else
        cost(isnan(cost)) = max(cost(:))+1;
        [link12,link21] = lap(cost);
        aligned(:,e+1:end) = aligned(link21,e+1:end);
    end
end

if(unwrap)
    aligned = orientationSpace.diffusion.unwrapExtrema(aligned, events, period);
end
if(~truncate)
    extrema(1:totalExtrema,:) = aligned;
    aligned = extrema;
end

end
```
