## Supplementary material for "Adaptive-Resolution Multi-Orientation Analysis of Complex Filamentous Network Images": MATLAB code from Github: findRegimeBifurcation.m

```
function [K_high,K_low,extrema_high,extrema_low] = findRegimeBifurcation(response,K_response,K_high,K_low,maxima,minima,maxIterations,tolerance,freq,debugIdx)
%findRegimeBifurcation Find a K interval where the bifurcation of the
%extrema regime at K_response exists
%
% INPUT
% response - response values at equidistant angles, NxM
% K_response - K of the response values, 1xM
% K_high - Initial upper limit of K for bifurcation, 1xM
% K_low - Initial lower limit of K for bifurcation, 1xM
% maxima - (optional) maxima of response at K_response
% minima - (optional) mimia of response at K_response
% maxIterations - Maximum number of times to try to refine interval
% tolerance - K interval around to bound the bifurcation
%
% OUTPUT
% K_high - Upper limit of K bounding bifurcation
% K_low - Lower limit of K bounding bifurcation
% maxima_high - Maxima at K_high
% maxima_low - Mimia at K_low
%
% REMARKS
% This function uses halleyft to quickly find extrema at the midpoint
% between K_high and K_low using the extrema at K_midpoint
%
% Copyright (C) 2019, Jaqaman Lab - UT Southwestern, Goldman Lab - Northwestern 
%
% This file is part of AdaptiveResolutionOrientationSpace.
% 
% AdaptiveResolutionOrientationSpace is free software: you can redistribute it and/or modify
% it under the terms of the GNU General Public License as published by
% the Free Software Foundation, either version 3 of the License, or
% (at your option) any later version.
% 
% AdaptiveResolutionOrientationSpace is distributed in the hope that it will be useful,
% but WITHOUT ANY WARRANTY; without even the implied warranty of
% MERCHANTABILITY or FITNESS FOR A PARTICULAR PURPOSE.  See the
% GNU General Public License for more details.
% 
% You should have received a copy of the GNU General Public License
% along with AdaptiveResolutionOrientationSpace.  If not, see <http://www.gnu.org/licenses/>.
% 
% 

% Mark Kittisopikul
% Northwestern University
% November 30th, 2018
    
    if(isscalar(K_response))
        K_response = repmat(K_response,1,size(response,2));
    end
    
    if(nargin < 4 || isempty(K_low))
        K_low = ones(1,size(response,2));
    end

    if(nargin < 5)
        maxima = [];
    end
    if(nargin < 6 )
        minima = [];
    end
    if(nargin < 7 || isempty(maxIterations))
        maxIterations = 10;
    end
    if(nargin < 8 || isempty(tolerance))
        tolerance = max(eps(K_high),(K_high-K_low)/2^maxIterations);
    end
    if(nargin < 9 || isempty(freq))
        freq = false;
    end
    if(nargin < 10 || isempty(debugIdx))
        debugIdx = [];
        debug = false;
    else
        debug = true;
    end
    
    if(freq)
        response_hat = response;
    else
        response_hat = fft(response);
    end
    
    if(isempty(maxima))
        % Calculate maxima and minima from response_hat if not given
        [maxima,minima] = interpft_extrema(response_hat,1,false,[],false);
    end
    
    % Use minima allows for some error checking since maxima and minima
    % should occur in pairs
    if(isempty(minima))
        useMinima = false;
    else
        useMinima = true;
    end

    % Make K_high match 2nd dimension of response
    if(isscalar(K_high))
        K_high = repmat(K_high,1,size(response,2));
    end
    
    % Make K_low match 2nd dimension of response
    if(size(K_low,2) == 1)
        K_low = repmat(K_low,1,size(response,2));
    end
    

    if(useMinima)
        % Combine maxima and minima into single array
        extrema_working = sort([maxima; minima]);
        % Pad extrema so that it matches first dimension of response
        extrema_working(end+1:size(response,1),:) = NaN;
%         isMaxima = [true(size(maxima)); false(size(minima))];
    else
        extrema_working = maxima;     
%         isMaxima = true(size(maxima));
    end
    % Initialize output to input, should be the same size
    extrema_high = extrema_working;
    
    nExtrema_working = size(extrema_working,1) - sum(isnan(extrema_working));
    % If there are one or less extrema we are done
    not_done = nExtrema_working > 1;
    
    % The working variables represent the data we are not done with
    K_high_working = K_high(:,not_done);
    K_low_working = K_low(:,not_done);
    K_response_working = K_response(:,not_done);
    extrema_working = extrema_working(:,not_done);  
    response_working_hat = response_hat(:,not_done);
    
    for i=1:maxIterations
        % Delegate the real work to a stripped down function
        [K_high_working,K_low_working,extrema_working] = ...
            findRegimeBifurcationHat( ...
                response_working_hat,K_response_working, ...
                K_high_working,K_low_working, ...
                extrema_working,useMinima);
        
        % Update the output variables with the data we are not done with
        K_high(not_done) = K_high_working;
        K_low(not_done) = K_low_working;
        extrema_high(:,not_done) = extrema_working;
        
        % We are not done if the difference in K exceeds the tolerance
        not_done_working = K_high_working - K_low_working > tolerance;
        not_done(not_done) = not_done_working;
        
        % Update the working variables
        K_response_working = K_response_working(not_done_working);
        K_high_working = K_high_working(not_done_working);
        K_low_working = K_low_working(not_done_working);
        response_working_hat = response_working_hat(:,not_done_working);
        extrema_working = extrema_working(:,not_done_working);
        
        if(debug)
            K_high(debugIdx)
            K_low(debugIdx)
            extrema_high(:,debugIdx)
        end
        
        if(isempty(K_high_working))
            if(debug)
                i
            end
            break;
        elseif(debug)
            length(K_high_working)
        end
        

    end
    
%         K_high(not_done) = K_high_working;
%         K_low(not_done) = K_low_working;
%         extrema_high(:,not_done) = extrema_working;
    
    if(nargout > 3)
        response_low_hat = orientationSpace.getResponseAtOrderVecHat(response_hat,K_response,K_low);
        [maxima_low,minima_low] =  interpft_extrema(response_low_hat,1,false,[],false);
        extrema_low = sort([maxima_low,minima_low]);
    end
end

function [K_high,K_low,extrema_high] = findRegimeBifurcationHat(response_hat,K_response,K_high,K_low,extrema,useMinima)
    nExtrema = size(extrema,1) - sum(isnan(extrema));
    K_midpoint = (K_high+K_low)/2;
    response_midpoint_hat = orientationSpace.getResponseAtOrderVecHat(response_hat,K_response,K_midpoint);
    
    [extrema_midpoint,xdg] = halleyft(response_midpoint_hat,extrema,true,1,1e-12,10,true,1e-4);
%     [maxima_midpoint,xdg] = halleyft_parallel_by_guess(response_midpoint_hat,maxima,true,1,1e-6,10,true,true);

    % Only keep maxima
    if(~useMinima)
        extrema_midpoint(xdg(:,:,2) > 0) = NaN;
    end

    % Should be done by halleyft with uniq = true
    % Eliminate duplicates
    extrema_midpoint = sort(extrema_midpoint);
    max_extrema_midpoint = max(extrema_midpoint)-2*pi;
    extrema_midpoint(diff([max_extrema_midpoint; extrema_midpoint]) < 1e-4) = NaN;
    extrema_midpoint = sort(extrema_midpoint);

    
%     maxima_midpoint = interpft_extrema(response_midpoint_hat,1,false,[],false);
    
    nExtremaMidpoint = size(extrema_midpoint,1) - sum(isnan(extrema_midpoint));
    
    % Do error correction
    if(useMinima)
        % Maxima and minima should occur in pairs.
        % An odd number of extrema would indicate an error
        oddIdx = mod(nExtremaMidpoint,2) == 1;
        % Find extrema that are close together, which may indicate an error
        closeExtremaIdx = any(diff([max(extrema_midpoint)-2*pi; extrema_midpoint]) < 0.02);
        
        oddIdx(closeExtremaIdx) = true;
        if(any(oddIdx))
            [odd_maxima,odd_minima] = interpft_extrema(response_midpoint_hat(:,oddIdx),1,true,[],false);
            oddExtrema = [odd_maxima; odd_minima];
            oddExtrema = sort(oddExtrema);
            oddExtrema = oddExtrema(1:min(size(extrema_midpoint,1),end),:);
            sExtrema = size(oddExtrema,1);
            extrema_midpoint(1:sExtrema,oddIdx) = oddExtrema;
            extrema_midpoint(sExtrema+1:end,oddIdx) = NaN;
            nExtremaMidpoint(oddIdx) = size(extrema_midpoint,1) - sum(isnan(extrema_midpoint(:,oddIdx)));
        end
    end
    
%     response_low_hat = orientationSpace.getResponseAtOrderVecHat(response_hat,K_response,K_low);
%     maxima_low = interpft_extrema(response_low_hat,1,false,[],false);
%     nMaximaLow = size(maxima,1) - sum(isnan(maxima_low));
%     assert(all(nMaximaLow ~= nMaxima))
    
    if(useMinima)
        bifurcationInHigh = nExtrema - nExtremaMidpoint >= 2;
    else
        bifurcationInHigh = nExtremaMidpoint ~= nExtrema;
    end
    bifurcationInLow = ~bifurcationInHigh;
    K_low(bifurcationInHigh) = K_midpoint(bifurcationInHigh);
    K_high(bifurcationInLow) = K_midpoint(bifurcationInLow);
    extrema_high = extrema;
    extrema_high(:,bifurcationInLow) = sort(extrema_midpoint(:,bifurcationInLow));
end
```
