## Supplementary material for "Adaptive-Resolution Multi-Orientation Analysis of Complex Filamentous Network Images": MATLAB code from Github: orientationMaximaDerivatives.m

```
function [ dnm_dKn, dnm_dtn ] = orientationMaximaDerivatives( rho, K, derivOrder, lm , period)
%ORIENTATIONMAXIMADERIVATIVES Find the K derivatives of local maxima
%
% INPUT
% rho - regularly spaced samples of orientation response at K,
%       nSamples x numel(K)
% K - angular order, vector of K values correpsond to rho and lm
%     If scalar, then K is expanded to match size(rho,2)
% derivOrder - highest derivative desired
% lm  - orientation local maxima of rho, via interpft_extrema
%       maxNLocalMaxima x numel(K)
%
% OUTPUT
% full derivatives of the orientation local maxima with respect to K
%
% Copyright (C) 2019, Jaqaman Lab - UT Southwestern, Goldman Lab - Northwestern 
%
% This file is part of AdaptiveResolutionOrientationSpace.
% 
% AdaptiveResolutionOrientationSpace is free software: you can redistribute it and/or modify
% it under the terms of the GNU General Public License as published by
% the Free Software Foundation, either version 3 of the License, or
% (at your option) any later version.
% 
% AdaptiveResolutionOrientationSpace is distributed in the hope that it will be useful,
% but WITHOUT ANY WARRANTY; without even the implied warranty of
% MERCHANTABILITY or FITNESS FOR A PARTICULAR PURPOSE.  See the
% GNU General Public License for more details.
% 
% You should have received a copy of the GNU General Public License
% along with AdaptiveResolutionOrientationSpace.  If not, see <http://www.gnu.org/licenses/>.
% 
% 

% Mark Kittisopikul, June 2017

    if(nargin < 5)
        period = 2*pi;
    end
    
    D = period.^2/2;
    
    % For period of 2*pi
%     D = 2*pi^2;
    % For period of pi
    % D = pi^2/2;

%% Calculate derivative in terms of t
%t = 1./(2*K+1).^2;

% n = derivOrder
% m = theta_{m} (local maxima location)

if(nargin < 4 || isempty(lm))
    lm = interpft_extrema(rho);
    lm = orientationSpace.diffusion.alignExtrema(lm,period);
end

if(~ismatrix(rho))
%     rho_sz = size(rho);
    rho = rho(:,:);
end
if(~ismatrix(lm))
    lm_sz = size(lm);
    lm = lm(:,:);
end

rho_derivs = interpft1_derivatives(rho,lm,2:derivOrder*2+1,period);

[dnm_dtn,maximaDerivatives] = dqm_dtq(derivOrder,D,rho_derivs);
if(nargout > 1)
    dnm_dtn = cat(3,maximaDerivatives{:});
end
    
% if(derivOrder == 1)
% %     rho_derivs = interpft1_derivatives(rho,lm,[2 3]);
%     dnm_dtn = -D.*rho_derivs(:,:,2)./rho_derivs(:,:,1);
% else
% end

%% Calculate derivatives of t by K

% n = shiftdim(1:derivOrder,-1);
% dnt_dKn = factorial(n+1) .* (-2).^(n );
% dnt_dKn = bsxfun(@times,dnt_dKn, ...
%     bsxfun(@power,1./(2*K+1),n+2));
% keyboard

%% Translate derivative with respect to t to with respect to K

% if(derivOrder == 1)
% %     keyboard;
%     dnm_dKn = bsxfun(@times,dnm_dtn,dnt_dKn(:,:,1));
% else
%     dnm_dKn = dnm_dtn;
% end

dnt_dKn = get_t_derivatives_with_respect_to_K(derivOrder,K);
K_expansion_factor = 1;
if(isscalar(K))
    K_expansion_factor = size(rho,2);
end
dnt_dKn = repmat(dnt_dKn,[size(lm,1) K_expansion_factor 1]);

dnm_dKn = zeros([size(lm),derivOrder]);
for d = 1:derivOrder
    dnm_dKn(:,:,d) = translate_from_t_to_K_hard(d,cat(3,maximaDerivatives{:}),K,dnt_dKn);
%     dnm_dKn(:,:,d) = translate_from_t_to_K(d,cat(3,maximaDerivatives{:}),K);
end

if(exist('lm_sz','var'))
    dnm_dKn = reshape(dnm_dKn,[lm_sz derivOrder]);
end

end

function deriv = total_dq_dtq_partial_dnrho_dmn(q,n,D,rho_derivs,maximaDerivatives)
%     assert(q >= 0);
%     assert(n > 0);
    if(n == 1)
        % Total derivative of 1st partial derivative with respect to
        % orientation. Always zero by definition of orientation local
        % maxima.
        deriv = 0;
%         fprintf('GET   total q=%d, n=%d\n',q,n);
%         fprintf('END total q=%d, n=%d\n',q,n);
        return;
    end
    if(q == 0)
        % No total derivative, answer is just the nth partial derivative with
        % respect to orientation
        deriv = rho_derivs(:,:,n-1);
%         fprintf('END total q=%d, n=%d\n',q,n);
%         fprintf('GET   total q=%d, n=%d\n',q,n);
        return;
    end
%     fprintf('START total q=%d, n=%d\n',q,n);

    % Then q > 0

    deriv = D * total_dq_dtq_partial_dnrho_dmn(q-1,n+2,D,rho_derivs,maximaDerivatives);
    deriv = deriv + total_dq_dtq_partial_dnrho_dmn(q-1,n+1,D,rho_derivs,maximaDerivatives).*maximaDerivatives{1};
    if(q == 1)
        return;
    end
    binom = nchoose_allk(q-1);
    
    for l = 2:q
%         binom = nchoosek(q-1,l-1);
%         deriv = deriv + binom.*total_dq_dtq_partial_dnrho_dmn(q-l,n+1,D,rho_derivs).*dqm_dtq(l,D,rho_derivs);
        deriv = deriv + binom(l).*total_dq_dtq_partial_dnrho_dmn(q-l,n+1,D,rho_derivs,maximaDerivatives).*maximaDerivatives{l};
    end
%     deriv = deriv +  D * total_dq_dtq_partial_dnrho_dmn(q-1,n+2,D,rho_derivs,maximaDerivatives);
%     fprintf('END   total q=%d, n=%d\n',q,n);
end

function [deriv,maximaDerivatives] = dqm_dtq(q,D,rho_derivs,maximaDerivatives)
    % order of the partial derivative is 1
    % need rho_derivs up to 1+q*2;
%     fprintf('START dqm_dtq q=%d\n',q);
    assert(q > 0);
    n = 1;
%     deriv = 0;
    if(nargin < 4)
        maximaDerivatives = cell(1,q);
        for l=1:q-1
            maximaDerivatives{l} = dqm_dtq(l,D,rho_derivs,maximaDerivatives);
        end
    end
    deriv = D * total_dq_dtq_partial_dnrho_dmn(q-1,n+2,D,rho_derivs,maximaDerivatives);
    if(q > 1)
        % binom(1) = 1
        deriv = deriv + total_dq_dtq_partial_dnrho_dmn(q-1,n+1,D,rho_derivs,maximaDerivatives).*maximaDerivatives{1};
        if(q > 2)
            binom = nchoose_allk(q-1);
            for l=2:q-1
        %         binom = nchoosek(q-1,l-1);
                deriv = deriv + binom(l).*total_dq_dtq_partial_dnrho_dmn(q-l,n+1,D,rho_derivs,maximaDerivatives).*maximaDerivatives{l};
            end
        end
    end
%     deriv = deriv + D * total_dq_dtq_partial_dnrho_dmn(q-1,n+2,D,rho_derivs,maximaDerivatives);
    % Divide by second partial derivative with respect to orientation
    deriv = -deriv./rho_derivs(:,:,1);
    maximaDerivatives{q} = deriv;
%     fprintf('END dqm_dtq q=%d\n',q);
end

function tpm = total_partial_matrix()
end

function dqm_dKq = translate_from_t_to_K(q,dqm_dtq_v,K)
    %% Calculate Faa di Bruno coefficients
    part = partitions(q);
    faa_di_bruno = factorial(q);
    faa_di_bruno = faa_di_bruno./prod(bsxfun(@power,factorial(1:q),part).');
    faa_di_bruno = faa_di_bruno./prod(factorial(part).');
    %% Calculate order of the local maxima derivative
    derivOrder = sum(part.');
    %% Calculate derivatives of t with respect to K
    n = shiftdim(1:q,-1);
    dnt_dKn = factorial(n+1) .* (-2).^(n );
    dnt_dKn = bsxfun(@times,dnt_dKn, ...
        bsxfun(@power,1./(2*K+1),n+2));
    dnt_dKn_pow = bsxfun(@power,dnt_dKn,shiftdim(part.',-2));
    dnt_dKn_pow = prod(dnt_dKn_pow,3);
    dqm_dKq = bsxfun(@times,dnt_dKn_pow,shiftdim(faa_di_bruno,-2));
    dqm_dKq = reshape(dqm_dKq,[1 size(dqm_dKq,2) size(dqm_dKq,4)]);
    dqm_dKq = bsxfun(@times,dqm_dKq,dqm_dtq_v(:,:,derivOrder));
    dqm_dKq = sum(dqm_dKq,3);
end

function dnt_dKn = get_t_derivatives_with_respect_to_K(q,K)
        dnt_dKn = zeros(1,length(K),q);
        dnt_dKn(:,:,1) = -4./(2*K+1).^3;
        for i=2:q
            dnt_dKn(:,:,i) = dnt_dKn(:,:,i-1)./(2*K+1)*-2*(i+1);
        end
end

function dqm_dKq = translate_from_t_to_K_hard(q,dqm_dtq_v,K,dnt_dKn)
% Translate derivatives with respect to t to derivatives with respect to K
% Hardcoded version for efficiency
% q - scalar, order of the derivative with respect to K
% dqm_dtq - derivatives with respect to t
% K - 
    if(q > 6)
        dqm_dKq = translate_from_t_to_K(q,dqm_dtq_v,K);
        return;
    end
    
    dnt_dKn = repmat(dnt_dKn,[1 size(dqm_dtq_v,2)./size(dnt_dKn,2) 1]);
    
    switch(q)
        case 1
            % partitions(1)
            % 
            % ans =
            % 
            %      1
            dqm_dKq = bsxfun(@times,dqm_dtq_v(:,:,1),dnt_dKn(:,:,1));
        case 2
            % partitions(2)
            % 
            % ans =
            % 
            %      2     0
            %      0     1
            dqm_dKq = dqm_dtq_v(:,:,2).* dnt_dKn(:,:,1).^2 ... 
                    + dqm_dtq_v(:,:,1).* dnt_dKn(:,:,2);
        case 3
            % partitions(3)
            % 
            % ans =
            % 
            %      3     0     0
            %      1     1     0
            %      0     0     1
            dqm_dKq =   dqm_dtq_v(:,:,3) .* dnt_dKn(:,:,1).^3 ...
                    + 3*dqm_dtq_v(:,:,2) .* dnt_dKn(:,:,1)     .* dnt_dKn(:,:,2) ...
                    +   dqm_dtq_v(:,:,1) .* dnt_dKn(:,:,3);
        case 4
            % partitions(4)
            % 
            % ans =
            % 
            %      4     0     0     0
            %      2     1     0     0
            %      0     2     0     0
            %      1     0     1     0
            %      0     0     0     1
            dqm_dKq =   dqm_dtq_v(:,:,4) .* dnt_dKn(:,:,1).^4 ...
                    + 6*dqm_dtq_v(:,:,3) .* dnt_dKn(:,:,1).^2 .* dnt_dKn(:,:,2) ...
                    + 3*dqm_dtq_v(:,:,2) .* dnt_dKn(:,:,2).^2 ...
                    + 4*dqm_dtq_v(:,:,2) .* dnt_dKn(:,:,1)    .* dnt_dKn(:,:,3) ...
                    +   dqm_dtq_v(:,:,1) .* dnt_dKn(:,:,4);
        case 5
            % partitions(5)
            % 
            % ans =
            % 
            %      5     0     0     0     0
            %      3     1     0     0     0
            %      1     2     0     0     0
            %      2     0     1     0     0
            %      0     1     1     0     0
            %      1     0     0     1     0
            %      0     0     0     0     1
            dqm_dKq =   dqm_dtq_v(:,:,5) .* dnt_dKn(:,:,1).^5 ...
                    +10*dqm_dtq_v(:,:,4) .* dnt_dKn(:,:,1).^3 .* dnt_dKn(:,:,2) ...
                    +15*dqm_dtq_v(:,:,3) .* dnt_dKn(:,:,1)    .* dnt_dKn(:,:,2).^2 ...
                    +10*dqm_dtq_v(:,:,3) .* dnt_dKn(:,:,1).^2 .* dnt_dKn(:,:,3) ...
                    +10*dqm_dtq_v(:,:,2) .* dnt_dKn(:,:,2)    .* dnt_dKn(:,:,3) ...
                    + 5*dqm_dtq_v(:,:,2) .* dnt_dKn(:,:,1)    .* dnt_dKn(:,:,4) ...
                    +   dqm_dtq_v(:,:,1) .* dnt_dKn(:,:,5);
        case 6
            % partitions(6)
            % 
            % ans =
            % 
            %      6     0     0     0     0     0
            %      4     1     0     0     0     0
            %      2     2     0     0     0     0
            %      0     3     0     0     0     0
            %      3     0     1     0     0     0
            %      1     1     1     0     0     0
            %      0     0     2     0     0     0
            %      2     0     0     1     0     0
            %      0     1     0     1     0     0
            %      1     0     0     0     1     0
            %      0     0     0     0     0     1
            dqm_dKq =   dqm_dtq_v(:,:,6) .* dnt_dKn(:,:,1).^6 ...
                    +15*dqm_dtq_v(:,:,5) .* dnt_dKn(:,:,1).^4 .* dnt_dKn(:,:,2) ...
                    +45*dqm_dtq_v(:,:,4) .* dnt_dKn(:,:,1).^2 .* dnt_dKn(:,:,2).^2 ...
                    +15*dqm_dtq_v(:,:,3) .* dnt_dKn(:,:,2).^3  ...
                    +20*dqm_dtq_v(:,:,4) .* dnt_dKn(:,:,1).^3 .* dnt_dKn(:,:,3) ...
                    +60*dqm_dtq_v(:,:,3) .* prod(dnt_dKn(:,:,1:3),3) ...
                    +10*dqm_dtq_v(:,:,2) .* dnt_dKn(:,:,3).^2 ...
                    +15*dqm_dtq_v(:,:,3) .* dnt_dKn(:,:,1).^2 .* dnt_dKn(:,:,4) ...
                    +15*dqm_dtq_v(:,:,2) .* dnt_dKn(:,:,2) .* dnt_dKn(:,:,4) ...
                    + 6*dqm_dtq_v(:,:,2) .* dnt_dKn(:,:,1) .* dnt_dKn(:,:,5) ...
                    +   dqm_dtq_v(:,:,1) .* dnt_dKn(:,:,6);
        otherwise
            error('Should not have gotten here');
    end
end

function out = nchoose_allk(n)
    switch(n)
        case 1
            out = [1 1];
            return;
        case 2
            out = [1 2 1];
            return;
        case 3
            out = [1 3 3 1];
            return;
        case 4
            out = [1 4 6 4 1];
            return;
        case 5
            out = [1 5 10 10 5 1];
            return;
        case 6
            out = [1 6 15 20 15 6 1];
            return;
        case 0
            out = 1;
            return;
        otherwise
    end
    out = zeros(1,n+1);
    for i=0:n
        out(i+1) = nchoosek(n,i);
    end
%     if(mod(n,2))
%         % odd
%         h = ceil(n/2)+1;
%     else
%         % even
%     end
end
```
