## Supplementary material for "Adaptive-Resolution Multi-Orientation Analysis of Complex Filamentous Network Images": MATLAB code from Github: orientationMaximaFirstDerivative.m

```
function [dm_dK,dm_dt,maxima] = orientationMaximaFirstDerivative(response_hat,response_K,maxima,period,refine)
%orientationMaximaFirstDerivative Get first derivative of the maxima with
%respect to K
%
% INPUT
% response_hat - Fourier transform of response
% response_K   - K of response
% maxima       - maxima
% period       - period of the doain
%
% OUTPUT
% dm_dK - first derivative of maxima with respect to K
% dm_dt - first derivative to maxima with respect to t
%
% This is a simplified version of orientationMaximaDerivatives
%
% See also orientationSpace.diffusion.orientationMaximaDerivatives
%
% Copyright (C) 2019, Jaqaman Lab - UT Southwestern, Goldman Lab - Northwestern 
%
% This file is part of AdaptiveResolutionOrientationSpace.
% 
% AdaptiveResolutionOrientationSpace is free software: you can redistribute it and/or modify
% it under the terms of the GNU General Public License as published by
% the Free Software Foundation, either version 3 of the License, or
% (at your option) any later version.
% 
% AdaptiveResolutionOrientationSpace is distributed in the hope that it will be useful,
% but WITHOUT ANY WARRANTY; without even the implied warranty of
% MERCHANTABILITY or FITNESS FOR A PARTICULAR PURPOSE.  See the
% GNU General Public License for more details.
% 
% You should have received a copy of the GNU General Public License
% along with AdaptiveResolutionOrientationSpace.  If not, see <http://www.gnu.org/licenses/>.
% 
% 

% Mark Kittisopikul, Ph.D.
% Goldman Lab
% Northwestern University
% December 2nd, 2018

    if(nargin < 4 || isempty(period))
        period = 2*pi;
    end
    if(nargin < 5)
        refine = false;
    end
    
    D = period.^2/2;
    
    if(refine)
        [new_maxima,partial_derivs] = halleyft(response_hat,maxima,'freq',true,'deriv',1,'uniq',true);
        new_maxima(partial_derivs(:,:,2) > 0) = NaN;
        
        % Do error correction
        nMaxima = size(maxima,1) - sum(isnan(maxima));
        nNewMaxima = size(new_maxima,1) - sum(isnan(new_maxima));
        error = nMaxima ~= nNewMaxima;
        if(any(error))
            temp_maxima = interpft_extrema(response_hat(:,error),1,true,[],false);
            new_maxima(1:size(temp_maxima,1),error) = temp_maxima;
            new_maxima(size(temp_maxima,1)+1:end,error) = NaN;
        end
        maxima = new_maxima;
        partial_derivs = partial_derivs(:,:,2:3);
    else
        partial_derivs = interpft1_derivatives(response_hat,maxima,2:3,period,true);
    end
    dm_dt = -partial_derivs(:,:,2)./partial_derivs(:,:,1)*D;
    dm_dK = dm_dt.*-4./(2*response_K+1).^3;
end
```
