## Supplementary material for "Adaptive-Resolution Multi-Orientation Analysis of Complex Filamentous Network Images": MATLAB code from Github: unwrapExtrema.m

```
function [ extrema ] = unwrapExtrema( extrema,events, period)
%unwrapExtrema Unwrap aligned extrema around the periodic boundary to avoid
%discontinuities
%
% Copyright (C) 2019, Jaqaman Lab - UT Southwestern, Goldman Lab - Northwestern 
%
% This file is part of AdaptiveResolutionOrientationSpace.
% 
% AdaptiveResolutionOrientationSpace is free software: you can redistribute it and/or modify
% it under the terms of the GNU General Public License as published by
% the Free Software Foundation, either version 3 of the License, or
% (at your option) any later version.
% 
% AdaptiveResolutionOrientationSpace is distributed in the hope that it will be useful,
% but WITHOUT ANY WARRANTY; without even the implied warranty of
% MERCHANTABILITY or FITNESS FOR A PARTICULAR PURPOSE.  See the
% GNU General Public License for more details.
% 
% You should have received a copy of the GNU General Public License
% along with AdaptiveResolutionOrientationSpace.  If not, see <http://www.gnu.org/licenses/>.
% 
% 

if(nargin < 3)
    period = 2*pi;
end

change = extrema(:,events+1) - extrema(:,events);
[r,c] = find(abs(change) > period/2);

for e = 1:length(r)
    extrema(r(e),events(c(e))+1:end) = extrema(r(e),events(c(e))+1:end) -sign(change(r(e),c(e)))*period;
end


end
```
