## Supplementary material for "Adaptive-Resolution Multi-Orientation Analysis of Complex Filamentous Network Images": MATLAB code from Github: angularKernel.m

```
function [ filterKernel ] = angularKernel( K, angle, N)
%orientationSpace.kernel
%
% Based on the thesis by Michael van Ginkel. Chapter 3
% "Image Analysis using Orientation Sapce based on Steerable Filters".
% Delft University of Technology. October 7th, 2002.
%
% K: number of rotation angles through 360 degrees
%
% Copyright (C) 2019, Jaqaman Lab - UT Southwestern, Goldman Lab - Northwestern 
%
% This file is part of AdaptiveResolutionOrientationSpace.
% 
% AdaptiveResolutionOrientationSpace is free software: you can redistribute it and/or modify
% it under the terms of the GNU General Public License as published by
% the Free Software Foundation, either version 3 of the License, or
% (at your option) any later version.
% 
% AdaptiveResolutionOrientationSpace is distributed in the hope that it will be useful,
% but WITHOUT ANY WARRANTY; without even the implied warranty of
% MERCHANTABILITY or FITNESS FOR A PARTICULAR PURPOSE.  See the
% GNU General Public License for more details.
% 
% You should have received a copy of the GNU General Public License
% along with AdaptiveResolutionOrientationSpace.  If not, see <http://www.gnu.org/licenses/>.
% 
% 

% Mark Kittisopikul, August 22nd, 2015
% Jaqaman Lab
% UT Southwestern

    import orientationSpace.*;

if(nargin < 1)
    K = 5;
end

n = 2*K+1;

if(nargin < 2 || isempty(angle))
    angle = (0:n-1)*(pi/n);
end

if(nargin < 3)
    N = 1024;
end

%% Moved to getFrequencySpaceCoordinates
% if(nargin < 5)
%     N = 1024;
% end
% 
% if(isscalar(N))
%     N = [N N];
% end
% 
% % [X, Y] = meshgrid(-N:(N-1));
% [X, Y] = meshgrid( (0:N(2)-1) - floor(N(2)/2), (0:N(1)-1) - floor(N(1)/2) );
% [theta, f] = cart2pol(X / floor(N(2)/2) / 2,Y / floor(N(1)/2) / 2);
% 
% theta = ifftshift(theta);
% f = ifftshift(f);
% % rho = ifftshift(rho);

if(isstruct(N))
    coords = N;
else
    coords = orientationSpace.getFrequencySpaceCoordinates(N);
end


% theta_shifted = acos(cos(theta - angle + pi));
% theta = acos(cos(theta - angle));
% 
% theta = theta - angle;
coords.theta = bsxfun(@minus,coords.theta,shiftdim(angle(:),-2));
% theta_shifted = theta + pi;

coords.theta = mod(coords.theta+pi,2*pi)-pi;
% theta = atan2(sin(theta),cos(theta));
% theta_shifted = atan2(sin(theta_shifted),cos(theta_shifted));


% f = rho / N / 2;

%% Radial part
% compute radial order, f_c = sqrt(K_f * b_f^2)
% K_f = (f_c / b_f)^2;

% scale frequency
% f_s = coords.f / f_c;

% Equation 3.11
% Note -(f^2 - f_c^2)/(2*b_f^2) = (1 - (f/f_c)^2)/(2* b_f^2/f_c^2)
%                               = (1 - (f/f_c)^2)/(2 / K_f)
% radialFilter = f_s.^K_f .* exp((1 - f_s.^2)*K_f/2);
% radialFilter2 = f_s^K_f .* exp(-(f.^2-f_c.^2)/2/b_f.^2);
% assertEqual(radialFilter,radialFilter2);

%% Angular part
% scale the angle

s_a = pi/(2*K+1);

theta_s = coords.theta / s_a;
% theta_shifed_s = theta_shifted / s_a;

angularFilter = 2*exp(-theta_s.^2/2);
% angularFilter_shifted = 2*exp(-theta_shifed_s.^2/2);
angularFilter_shifted = angularFilter([1 end:-1:2],[1 end:-1:2],:);

% filterKernel = bsxfun(@times,0.5,angularFilter + angularFilter_shifted);
filterKernel = 0.5*(angularFilter + angularFilter_shifted);

% could we simplify this? neg/pos dividing line does not have to rotate
posMask = abs(coords.theta) < pi/2;
filterKernel = filterKernel.*(1 + 1j.*(posMask.*2-1));

% shift = exp(1i*2*pi*(X/2+Y/2));
% filterKernel = radialFilter .* angularFilter;
% filterKernel = radialFilter .* angularFilter .* 2 .* ( abs(theta) < pi/2 );
% filterKernel = radialFilter .* angularFilter .*exp(1i*2*pi*(X/2+Y/2));
% filterKernel = radialFilter .* angularFilter + 1i * radialFilter .* angularFilter .* (-1).^(abs(theta) >= pi/2);
% filterKernel = filterKernel .* (f > 0);


end
```
