## Supplementary material for "Adaptive-Resolution Multi-Orientation Analysis of Complex Filamentous Network Images": MATLAB code from Github: blendOrientationMap.m

```
function [ blended_map ] = blendOrientationMap( theta, res, cm )
%UNTITLED Summary of this function goes here
%   Detailed explanation goes here
%
% Copyright (C) 2019, Jaqaman Lab - UT Southwestern, Goldman Lab - Northwestern 
%
% This file is part of AdaptiveResolutionOrientationSpace.
% 
% AdaptiveResolutionOrientationSpace is free software: you can redistribute it and/or modify
% it under the terms of the GNU General Public License as published by
% the Free Software Foundation, either version 3 of the License, or
% (at your option) any later version.
% 
% AdaptiveResolutionOrientationSpace is distributed in the hope that it will be useful,
% but WITHOUT ANY WARRANTY; without even the implied warranty of
% MERCHANTABILITY or FITNESS FOR A PARTICULAR PURPOSE.  See the
% GNU General Public License for more details.
% 
% You should have received a copy of the GNU General Public License
% along with AdaptiveResolutionOrientationSpace.  If not, see <http://www.gnu.org/licenses/>.
% 
% 

if(nargin < 2 || isempty(res))
    res = ones(size(theta));
end
if(nargin < 3)
    cm = hsv(256);
end

theta_not_nan = ~isnan(theta);
theta_mapped = NaN([numel(theta) 3]);
theta_mapped(theta_not_nan,:) = cm(floor(theta(theta_not_nan)/pi*(size(cm,1)-1))+1,:);
sz_theta = size(theta);
sz_theta(ndims(theta)+1:3) = 1;
theta_mapped = reshape(theta_mapped,[sz_theta 3]);
res(res < 0) = NaN;
blended_map = squeeze((nansum(theta_mapped.*res,3)./nansum(res,3)));

if(nargout == 0)
    figure;
    imshow(blended_map);
end


% figure;
% imshow(squeeze((nansum(theta_mapped.*res,3)./nansum(res,3))),[]);

end
```
