## Supplementary material for "Adaptive-Resolution Multi-Orientation Analysis of Complex Filamentous Network Images": MATLAB code from Github: getFrequencySpaceCoordinates.m

```
function [ polar, cartesian ] = getFrequencySpaceCoordinates( N )
%GETFREQUENCYSPACECOORDINATES orientationSpace.getFrequencySpaceCoordinates
% obtain polar and cartesian coordinates
%
% Caches the last coordinates for the last N
%
% INPUT
% N - size of space, if scalar a size of [N N] is extrapolated
%
% OUTPUT
% polar - polar coordinates, shifted
%    .f - frequency magnitude, from 0 to sqrt(2)/2
%    .theta - frequency angle (-pi,pi]
% cartesian - cartesian coordinates, shifted
%    .X - X coordinate from -floor(N/2) to N-floor(N/2)
%    .Y - Y coordinate from -floor(N/2) to N-floor(N/2)
% 
% See also ifftshift, fftshift
%
% Copyright (C) 2019, Jaqaman Lab - UT Southwestern, Goldman Lab - Northwestern 
%
% This file is part of AdaptiveResolutionOrientationSpace.
% 
% AdaptiveResolutionOrientationSpace is free software: you can redistribute it and/or modify
% it under the terms of the GNU General Public License as published by
% the Free Software Foundation, either version 3 of the License, or
% (at your option) any later version.
% 
% AdaptiveResolutionOrientationSpace is distributed in the hope that it will be useful,
% but WITHOUT ANY WARRANTY; without even the implied warranty of
% MERCHANTABILITY or FITNESS FOR A PARTICULAR PURPOSE.  See the
% GNU General Public License for more details.
% 
% You should have received a copy of the GNU General Public License
% along with AdaptiveResolutionOrientationSpace.  If not, see <http://www.gnu.org/licenses/>.
% 
% 

persistent lastN lastCartesian lastPolar


if(nargin < 1)
    N = 1024;
end

if(isscalar(N))
    N = [N N];
end

% if(~isempty(lastN) && all(N == lastN))
if(isequal(lastN,N))
    cartesian = lastCartesian;
    polar = lastPolar;
else   
    lastN = N;

    % [X, Y] = meshgrid(-N:(N-1));
    [cartesian.X, cartesian.Y] = meshgrid( (0:N(2)-1) - floor(N(2)/2), (0:N(1)-1) - floor(N(1)/2) );

    cartesian.X = ifftshift(cartesian.X);
    cartesian.Y = ifftshift(cartesian.Y);

    [polar.theta, polar.f] = cart2pol(cartesian.X / floor(N(2)/2) / 2,cartesian.Y / floor(N(1)/2) / 2);
    
    lastCartesian = cartesian;
    lastPolar = polar;

    % polar.theta = ifftshift(polar.theta);
    % polar.f = ifftshift(polar.f);
end


end
```
