## Supplementary material for "Adaptive-Resolution Multi-Orientation Analysis of Complex Filamentous Network Images": MATLAB code from Github: getResponseAtOrderVecHat.m

```
function responseAtOrder = getResponseAtOrderVecHat(response_hat,Korg,Kg)
%getResponseAtOrderVec Get response at order, vectorized so that each pixel
% has an independent order. Response already has Fourier transform applied
%
% Extracted from orientationSpace.diffusion.refineBifurcation
%
% Copyright (C) 2019, Jaqaman Lab - UT Southwestern, Goldman Lab - Northwestern 
%
% This file is part of AdaptiveResolutionOrientationSpace.
% 
% AdaptiveResolutionOrientationSpace is free software: you can redistribute it and/or modify
% it under the terms of the GNU General Public License as published by
% the Free Software Foundation, either version 3 of the License, or
% (at your option) any later version.
% 
% AdaptiveResolutionOrientationSpace is distributed in the hope that it will be useful,
% but WITHOUT ANY WARRANTY; without even the implied warranty of
% MERCHANTABILITY or FITNESS FOR A PARTICULAR PURPOSE.  See the
% GNU General Public License for more details.
% 
% You should have received a copy of the GNU General Public License
% along with AdaptiveResolutionOrientationSpace.  If not, see <http://www.gnu.org/licenses/>.
% 
% 

% Mark Kittisopikul, November 2017
% Northwestern University

    if(isempty(Kg))
%         responseAtOrder = NaN(size(response_hat));
        responseAtOrder = NaN([size(response_hat,1) numel(Kg)]);
        return;
    end
%     x = [0:ceil(R.filter.K)*R.filter.sampleFactor -ceil(R.filter.K)*R.filter.sampleFactor:-1];
%     x = [0:8 -8:-1];
    s = floor(size(response_hat,1)/2);
    x = [0:s -s:-1];
    n_org = 2*Korg+1;

    n_new = 2*Kg+1;
    s_inv = sqrt(n_org.^2.*n_new.^2./(n_org.^2-n_new.^2));
    s_hat = s_inv/(2*pi);
    f_hat = exp(-0.5 * bsxfun(@rdivide,x(:),s_hat).^2);
    responseAtOrder = bsxfun(@times,response_hat,f_hat);
end
```
