## Supplementary material for "Adaptive-Resolution Multi-Orientation Analysis of Complex Filamentous Network Images": MATLAB code from Github: maxima.m

```
function [ theta, response, nms ] = maxima( orientationMatrix, upsample_factor )
%orientationSpace.maxima finds the absolute maxima
%
% Copyright (C) 2019, Jaqaman Lab - UT Southwestern, Goldman Lab - Northwestern 
%
% This file is part of AdaptiveResolutionOrientationSpace.
% 
% AdaptiveResolutionOrientationSpace is free software: you can redistribute it and/or modify
% it under the terms of the GNU General Public License as published by
% the Free Software Foundation, either version 3 of the License, or
% (at your option) any later version.
% 
% AdaptiveResolutionOrientationSpace is distributed in the hope that it will be useful,
% but WITHOUT ANY WARRANTY; without even the implied warranty of
% MERCHANTABILITY or FITNESS FOR A PARTICULAR PURPOSE.  See the
% GNU General Public License for more details.
% 
% You should have received a copy of the GNU General Public License
% along with AdaptiveResolutionOrientationSpace.  If not, see <http://www.gnu.org/licenses/>.
% 
% 
    import orientationSpace.*;
    
    if(nargin < 2)
        % factor to upsample by to increase efficiency of algorithm
        upsample_factor = 8;
    end

    s = size(orientationMatrix);
    
    if(upsample_factor > 1)
        M_upsample = orientationSpace.upsample(orientationMatrix,pi/s(3)/upsample_factor);
        M_upsample = real(reshape(M_upsample,s(1)*s(2),s(3)*upsample_factor));
    end
    
    M = real(reshape(orientationMatrix,s(1)*s(2),s(3)));

    n = (s(3))/2;

    
%     x = [0:n -n:-1];
    x = wraparoundN(0:s(3)-1,-n,n);
    % toeplitz?
    xx = wraparoundN(bsxfun(@minus,x,(0:(2*n-1))'),[-n n]);
    % xx = arrayfun(@(k) circshift(x,k,2),0:s(3),'UniformOutput',false);
    % xx = vertcat(xx{:});

    
    if(upsample_factor > 1)
        % upsample
        x_upsample = wraparoundN(0:1/upsample_factor:s(3)-1/upsample_factor,-n,n);
        [~,ind] = max(M_upsample,[],2);
        xind = x_upsample(ind)';
        clear M_upsample x_upsample
    else
        [~,ind] = max(M,[],2);
        xind = x(ind)';
    end
    
    clear ind

    A = 2*exp(-xx.^2/2);
    M = M/A;
    nM = size(M,1);
    
    % Initialize Batch Structures for Optimization
    batchSize = s(1);
    nBatches  = nM/batchSize;
    Mcell     = cell(1,nBatches);
    xindc     = cell(1,nBatches);
    theta     = cell(1,nBatches);
    
    % group indices into batches
    for i=1:nBatches
        idx = (1:batchSize)+(i-1)*batchSize;
        Mcell{i} = M(idx,:);
        xindc{i} = xind(idx);
    end
    
    clear xind

    options = optimoptions('lsqnonlin','Jacobian','on','Display','off','TolFun',1e-9);
%     progressText(0,'Solving');
    
    parfor i=1:nBatches
        offset = xindc{i};
        f = @(k) orientationDerivWrap(k,x,Mcell{i},n);
        theta{i} = lsqnonlin(f,offset,offset-0.5,offset+0.5,options);
        %         progressText(i/nM*1024);
    end

    % assemble image
    theta = horzcat(theta{:});
    % wrap around one more time, needed in the even case
    theta = wraparoundN(theta,-n,n);
    
    if(nargout > 1)
        T = bsxfun(@minus,x,theta(:));
        T = wraparoundN(T,-n,n);
        T = 2*exp(-T.^2/2);
        response = sum(M.*T,2);
        response = reshape(response,s(1),s(2));
        if(~isreal(orientationMatrix))
            [theta_i,response_i] = orientationSpace.maxima(cat(3,imag(orientationMatrix),-imag(orientationMatrix)),upsample_factor);
            % multiply theta by 2 to expand range to [-pi,pi)
            theta = theta + 2j*theta_i;
            response = response + 1j*response_i;
        end
    elseif(~isreal(orientationMatrix))
        theta_i = orientationSpace.maxima(cat(3,imag(orientationMatrix),-imag(orientationMatrix)),upsample_factor);
        theta = theta + 2j*theta_i;
    end
    
    theta = real(theta)*pi/s(3) + 1j*imag(theta);
   
  
    if(nargout > 2)
        nms = nonMaximumSuppression(real(response),real(theta));
        if(~isreal(theta))
            nms = nms + 1j*nonMaximumSuppression(imag(response),imag(theta));
        end
    end

end
function [F,J] = orientationDerivWrap(k,x,M,n)
    % k is the angle of the candidate maximum
    % x is the angle space
    % M represents the coefficients, the deconvolved response
    % n represents the wraparound limits
    t = wraparoundN(bsxfun(@minus,k,x),-n,n);
    et = exp(-t.^2/2);
    % derivative with respect to k
    F = -sum(M.*(t.*et),2);
    if(nargout > 1)
        % second derivative with respect to k
        J = sum(M.*(t.^2.*et - et),2);
        % along the main diagonal is of interest
        J = spdiags(J(:),0,length(F),length(k));
    end
end
```
