## Supplementary material for "Adaptive-Resolution Multi-Orientation Analysis of Complex Filamentous Network Images": MATLAB code from Github: OrientationSpaceResponseIndex.m

```
classdef OrientationSpaceResponseIndex
    %orietationSpace.OrientationSpaceResponseIndex allow for indexing into a
    %OrientationResponse object
%
% Copyright (C) 2019, Jaqaman Lab - UT Southwestern, Goldman Lab - Northwestern 
%
% This file is part of AdaptiveResolutionOrientationSpace.
% 
% AdaptiveResolutionOrientationSpace is free software: you can redistribute it and/or modify
% it under the terms of the GNU General Public License as published by
% the Free Software Foundation, either version 3 of the License, or
% (at your option) any later version.
% 
% AdaptiveResolutionOrientationSpace is distributed in the hope that it will be useful,
% but WITHOUT ANY WARRANTY; without even the implied warranty of
% MERCHANTABILITY or FITNESS FOR A PARTICULAR PURPOSE.  See the
% GNU General Public License for more details.
% 
% You should have received a copy of the GNU General Public License
% along with AdaptiveResolutionOrientationSpace.  If not, see <http://www.gnu.org/licenses/>.
% 
% 
    
    properties
        response
    end
    
    methods
        function obj =  OrientationSpaceResponseIndex(orientationSpaceResponse)
            obj.response = orientationSpaceResponse;
        end
        function varargout = subsref(obj,S)
            switch(S(1).type)
%                 case '.'
                case '()'
                    varargout{1} = obj.response.getResponseAtPoint(S(1).subs{:});
                case '{}'
                    varargout{1} =  obj.response.getResponseAtOrientation(S(1).subs{:});
                otherwise
                    [varargout{1:nargout}] = builtin('subsref',obj,S);
            end
        end
    end
    
end
```
