## Supplementary material for "Adaptive-Resolution Multi-Orientation Analysis of Complex Filamentous Network Images": MATLAB code from Github: radialKernel.m

```
function [ radialFilter ] = radialKernel( f_c, b_f, N )
%radialKernel Calculate the orientationSpace radial kernel
%
% See also orientationSpace.kernel, orientationSpace.angularKernel,
% orientationSpace.getFrequencySpaceCoordinates
%
% Copyright (C) 2019, Jaqaman Lab - UT Southwestern, Goldman Lab - Northwestern 
%
% This file is part of AdaptiveResolutionOrientationSpace.
% 
% AdaptiveResolutionOrientationSpace is free software: you can redistribute it and/or modify
% it under the terms of the GNU General Public License as published by
% the Free Software Foundation, either version 3 of the License, or
% (at your option) any later version.
% 
% AdaptiveResolutionOrientationSpace is distributed in the hope that it will be useful,
% but WITHOUT ANY WARRANTY; without even the implied warranty of
% MERCHANTABILITY or FITNESS FOR A PARTICULAR PURPOSE.  See the
% GNU General Public License for more details.
% 
% You should have received a copy of the GNU General Public License
% along with AdaptiveResolutionOrientationSpace.  If not, see <http://www.gnu.org/licenses/>.
% 
% 

% Mark Kittisopikul
% Jaqaman Lab
% UT Southwestern
% June 24th, 2016

if(nargin < 2 || isempty(b_f))
    b_f = f_c/sqrt(2);
end
if(nargin < 3)
    N = 1024;
end

assert(length(f_c) == length(b_f),'orientationSpace.radialKernel:f_c and b_f must be of equal size');

f_c = shiftdim(f_c(:),-3);
b_f = shiftdim(b_f(:),-3);

if(isstruct(N))
    coords = N;
else
    coords = orientationSpace.getFrequencySpaceCoordinates(N);
end

%% Radial part
% compute radial order, f_c = sqrt(K_f * b_f^2)
if(f_c)
    K_f = (f_c ./ b_f).^2;

    % scale frequency
    % f_s = coords.f / f_c;
    f_s = bsxfun(@rdivide,coords.f,f_c);

    % Equation 3.11
    % Note -(f^2 - f_c^2)/(2*b_f^2) = (1 - (f/f_c)^2)/(2* b_f^2/f_c^2)
    %                               = (1 - (f/f_c)^2)/(2 / K_f)
    % radialFilter = f_s.^K_f .* exp((1 - f_s.^2)*K_f/2);
    radialFilter = bsxfun(@power,f_s,K_f);
    radialFilter = radialFilter .* exp(bsxfun(@times,(1 - f_s.^2),K_f/2));
    % radialFilter2 = f_s^K_f .* exp(-(f.^2-f_c.^2)/2/b_f.^2);
    % assertEqual(radialFilter,radialFilter2);
else
    radialFilter = exp(-bsxfun(@rdivide,coords.f.^2,2*b_f.^2));
    radialFilter(1) = 0;
end

end
```
