## Supplementary material for "Adaptive-Resolution Multi-Orientation Analysis of Complex Filamentous Network Images": MATLAB code from Github: rainbowOrientationQuivers.m

```
function [h] = rainbowOrientationQuivers(maxima,magnitude,cm,N,threeD,magnitudeThreshold,varargin)
%rainbowQuivers Plot a rainbow plot of angles
%
% Copyright (C) 2019, Jaqaman Lab - UT Southwestern, Goldman Lab - Northwestern 
%
% This file is part of AdaptiveResolutionOrientationSpace.
% 
% AdaptiveResolutionOrientationSpace is free software: you can redistribute it and/or modify
% it under the terms of the GNU General Public License as published by
% the Free Software Foundation, either version 3 of the License, or
% (at your option) any later version.
% 
% AdaptiveResolutionOrientationSpace is distributed in the hope that it will be useful,
% but WITHOUT ANY WARRANTY; without even the implied warranty of
% MERCHANTABILITY or FITNESS FOR A PARTICULAR PURPOSE.  See the
% GNU General Public License for more details.
% 
% You should have received a copy of the GNU General Public License
% along with AdaptiveResolutionOrientationSpace.  If not, see <http://www.gnu.org/licenses/>.
% 
% 

if(nargin < 4  || isempty(N))
    N = 36;
end
if(nargin < 2)
    magnitude = ones(size(maxima));
end
if(nargin < 3 || isempty(cm))
    cm = hsv(N);
else
    N = size(cm,1);
end
if(nargin < 5)
    threeD = false;
end
if(nargin < 6)
    magnitudeThreshold = 0;
end


maxima = wraparoundN(maxima,0,pi);
maximaBinned = floor(maxima/pi*N);

[X,Y] = meshgrid(1:size(maxima,2),1:size(maxima,1));

X = repmat(X,1,1,size(maxima,3));
Y = repmat(Y,1,1,size(maxima,3));

if(threeD)
    W = zeros(size(maxima));
end


s = magnitude >= magnitudeThreshold;
magnitude(s) = mat2gray(magnitude(s))*0.99+0.01;
magnitude(magnitude < magnitudeThreshold) = 0;
magnitude(isnan(magnitude)) = 0;

h(N) = matlab.graphics.GraphicsPlaceholder;

hold on;

for n=1:N
    s = maximaBinned == (n-1) & magnitude ~= 0;
    Xs = X(s);
    Ys = Y(s);
    maxima_s = maxima(s);
    magnitude_s = magnitude(s);
    Us = magnitude_s.*-sin(maxima(s));
    Vs = magnitude_s.*cos(maxima(s));
    Xs = Xs - Us/2;
    Ys = Ys - Vs/2;
    if(threeD)
        Zs = maxima(s);
        Ws = W(s);
        h(n) = quiver3(Xs,Ys,Zs,Us,Vs,Ws,0,'Color',cm(n,:),'ShowArrowHead','off',varargin{:});
    else
        h(n) = quiver(Xs,Ys,Us,Vs,0,'Color',cm(n,:),'ShowArrowHead','off',varargin{:});
    end
end

end
```
