## Supplementary material for "Adaptive-Resolution Multi-Orientation Analysis of Complex Filamentous Network Images": MATLAB code from Github: upsample.m

```
function [ y ] = upsample( orientationMatrix, target )
%orientationSpace.upsample upsample orientation information
%
% INPUT
% orientationMatrix : YxXxOrientation
% target            : Orientation angles to estimate
%
% OUTPUT
% y                 : YxXxTarget
%
% Copyright (C) 2019, Jaqaman Lab - UT Southwestern, Goldman Lab - Northwestern 
%
% This file is part of AdaptiveResolutionOrientationSpace.
% 
% AdaptiveResolutionOrientationSpace is free software: you can redistribute it and/or modify
% it under the terms of the GNU General Public License as published by
% the Free Software Foundation, either version 3 of the License, or
% (at your option) any later version.
% 
% AdaptiveResolutionOrientationSpace is distributed in the hope that it will be useful,
% but WITHOUT ANY WARRANTY; without even the implied warranty of
% MERCHANTABILITY or FITNESS FOR A PARTICULAR PURPOSE.  See the
% GNU General Public License for more details.
% 
% You should have received a copy of the GNU General Public License
% along with AdaptiveResolutionOrientationSpace.  If not, see <http://www.gnu.org/licenses/>.
% 
% 
    import orientationSpace.*;
    % 
    s = size(orientationMatrix);
    M = reshape(orientationMatrix,s(1)*s(2),s(3));
    % todo, deal with even case
    n = (s(3))/2;
%     n = n+0.5;

    
%     x = [0:n -n:-1];
    x = wraparoundN(0:s(3)-1,-n,n);
    % toeplitz?
    xx = wraparoundN(bsxfun(@minus,x,(0:(2*n-1))'),[-n n]);
    % xx = arrayfun(@(k) circshift(x,k,2),0:s(3),'UniformOutput',false);
    % xx = vertcat(xx{:});

    A = exp(-xx.^2/2);

    if(isscalar(target))
        target = 0:target:pi-target;
    end
    target = target./(pi/s(3));
    
    % note we factor out the exp(1/2) constant factor

    T = bsxfun(@minus,x,target')';
    T = wraparoundN(T,[-n n]);
    T = exp(-T.^2/2);
    w = A\T;
    % normalize the weights such that the columns sum to 1
%     w = w./repmat(sum(w),s(3),1);

    y = M*w;
    y = reshape(y,s(1),s(2),[]);

end
```
