## Supplementary material for "Adaptive-Resolution Multi-Orientation Analysis of Complex Filamentous Network Images": MATLAB code from Github: animateAngularOrder.m

```
function [  ] = animateAngularOrder( R, r, c, num_derivatives )
%animateAngularOrder Summary of this function goes here
%   Detailed explanation goes here
%
% Copyright (C) 2019, Jaqaman Lab - UT Southwestern, Goldman Lab - Northwestern 
%
% This file is part of AdaptiveResolutionOrientationSpace.
% 
% AdaptiveResolutionOrientationSpace is free software: you can redistribute it and/or modify
% it under the terms of the GNU General Public License as published by
% the Free Software Foundation, either version 3 of the License, or
% (at your option) any later version.
% 
% AdaptiveResolutionOrientationSpace is distributed in the hope that it will be useful,
% but WITHOUT ANY WARRANTY; without even the implied warranty of
% MERCHANTABILITY or FITNESS FOR A PARTICULAR PURPOSE.  See the
% GNU General Public License for more details.
% 
% You should have received a copy of the GNU General Public License
% along with AdaptiveResolutionOrientationSpace.  If not, see <http://www.gnu.org/licenses/>.
% 
% 

if(nargin < 2)
    r = 127;
end
if(nargin < 3)
    c = 126;
end
if(nargin < 4)
    num_derivatives = 3;
end
% if(nargin < 4)
    da = ifft(bsxfun(@times,fft(R.a,[],3),shiftdim([0:8 -8:-1],-1))*1i,[],3);
    Rd = OrientationSpaceResponse(R.filter,da);
% end

pauseTime = 0.1;
deltaK = 0.1;

h(1) = plot(0:0.5:179.5,interpft(R.getResponseAtOrderFTatPoint(r,c,8),360));
hold on;
h(2) = plot(0:0.5:179.5,interpft(Rd.getResponseAtOrderFTatPoint(r,c,8),360));

h(3) = line;
h(3).LineStyle = 'none';
h(3).Marker = '.';
h(3).MarkerSize = 1;

h(4) = line;
h(4).Marker = 'o';
h(4).LineStyle = 'none';

h(5) = line;
h(5).LineStyle = 'none';
h(5).Marker = '.';
h(5).MarkerSize = 1;
h(5).Color = 'r';

h(6) = line;
h(6).Marker = 's';
h(6).LineStyle = 'none';
h(6).Color = 'r';

grid on;
drawnow;
yl = ylim;
xl = xlim;
ht = text(xl(2)-30,yl(2)-10,['K = ' num2str(R.filter.K)]);
ylim('manual');

for nd = 2:num_derivatives
    hd(nd-1) = plot(0:0.5:179.5,zeros(1,360),'--');
%     hd(nd-1).XData = 0:0.5:179.5;
%     hd(nd-1).YData = zeros(1,360);
end


while all(isvalid(h))
    h(3).XData = [];
    h(3).YData = [];
%     frame(length(R.filter.K:-deltaK:-0.5+deltaK)) = struct();
%     counter = 1;
    for K=R.filter.K:-deltaK:-0.5+deltaK
        temp = R.getResponseAtOrderFTatPoint(r,c,K);
        h(1).YData = interpft(temp,360);
        h(2).YData = interpft(Rd.getResponseAtOrderFTatPoint(r,c,K),360);
        ht.String = ['K = ' num2str(K)];
        [maxima,minima,maxima_value,minima_value] = interpft_extrema(R.getResponseAtOrderFTatPoint(r,c,K));
        h(3).XData = [h(3).XData maxima.'/2/pi*180];
        h(3).YData = [h(3).YData maxima_value.'];
        h(4).XData = maxima.'/2/pi*180;
        h(4).YData = maxima_value.';
        
        h(5).XData = [h(5).XData minima.'/2/pi*180];
        h(5).YData = [h(5).YData minima_value.'];
        h(6).XData = minima.'/2/pi*180;
        h(6).YData = minima_value.';
        for nd = 2:num_derivatives
            da = ifft(bsxfun(@times,fft(temp),(shiftdim([0:R.filter.order -R.filter.order:-1]*1i,1)).^nd));
            hd(nd-1).YData = interpft(da,360);
        end
%         frame(counter) = getframe(gcf);
%         counter = counter+1;
        pause(pauseTime);
    end
end

end
```
