## Supplementary material for "Adaptive-Resolution Multi-Orientation Analysis of Complex Filamentous Network Images": MATLAB code from Github: findRegimesAtPoint.m

```
function [ allMaxima, maximaTraceRegimesInv, maximaTraceRegimesInvK, maximaValue] = findRegimesAtPoint( R, r, c, K )
%findRegimesAtPoint Summary of this function goes here
%   Detailed explanation goes here
%
% Copyright (C) 2019, Jaqaman Lab - UT Southwestern, Goldman Lab - Northwestern 
%
% This file is part of AdaptiveResolutionOrientationSpace.
% 
% AdaptiveResolutionOrientationSpace is free software: you can redistribute it and/or modify
% it under the terms of the GNU General Public License as published by
% the Free Software Foundation, either version 3 of the License, or
% (at your option) any later version.
% 
% AdaptiveResolutionOrientationSpace is distributed in the hope that it will be useful,
% but WITHOUT ANY WARRANTY; without even the implied warranty of
% MERCHANTABILITY or FITNESS FOR A PARTICULAR PURPOSE.  See the
% GNU General Public License for more details.
% 
% You should have received a copy of the GNU General Public License
% along with AdaptiveResolutionOrientationSpace.  If not, see <http://www.gnu.org/licenses/>.
% 
% 

if(nargin < 2)
    r = round(size(R.a,1)/2);
end
if(nargin < 3)
    c = round(size(R.a,2)/2);
end
if(nargin < 4)
    K = 8:-0.1:1;
end

rho = R.getResponseAtOrderFTatPoint(r,c,K);
out = interpft_extrema(rho,1,true)/2;
out = orientationSpace.diffusion.alignExtrema(out,pi);
out_values = interpft1([0 pi],rho,out);
% out = orientationSpace.diffusion.unwrapExtrema(out);
nExtrema = sum(~isnan(out));
maximaTraceDeriv = orientationSpace.diffusion.orientationMaximaDerivatives(rho,K,1,out,pi);

maximaTraceRegimesInv = cell(1,size(out,1));
maximaTraceRegimesInvK = cell(1,size(out,1));

for n=1:size(out,1)
    s = nExtrema == n;
    maximaTraceDerivAbs = abs(maximaTraceDeriv);
    maximaTraceDerivAbs(:,~s) = NaN;
    [minDeriv,minidx] = min(maximaTraceDerivAbs,[],2);
    maximaTraceRegimesInvK{n} = K(minidx);
    maximaTraceRegimesInvK{n}(isnan(minDeriv)) = NaN;
    o = ones(size(minidx));
    minidx = sub2ind(size(out),(1:size(minidx,1)).'.*o,minidx);
    maximaTraceRegimesInv{n} = out(minidx);
    maximaTraceRegimesInvValues{n} = out_values(minidx);
    maximaTraceRegimesInv{n}(isnan(minDeriv)) = NaN;
    maximaTraceRegimesInvValues{n}(isnan(minDeriv)) = NaN;
end
allMaxima = [maximaTraceRegimesInv{:}];
allMaxima = allMaxima(:);
% maximaValue = interpft1([0 pi],rho(:,1),allMaxima);
maximaValue = [maximaTraceRegimesInvValues{:}];
maximaValue = maximaValue(:);
avg = mean(rho(:,1));
allMaxima(maximaValue < avg) = NaN;
if(nargout > 1)
    numPerRegime = cellfun('prodofsize',maximaTraceRegimesInv);
    maximaTraceRegimesInv = mat2cell(allMaxima,numPerRegime);
    if(nargout > 2)
        maximaValue = mat2cell(maximaValue,numPerRegime);
    end
end

end
```
