## Supplementary material for "Adaptive-Resolution Multi-Orientation Analysis of Complex Filamentous Network Images": MATLAB code from Github: OrientationSpaceResponse.m

```
classdef OrientationSpaceResponse < handle
    %OrientationSpaceResponse Response object for OrientationSpaceFilter
    %
    % Response object used by
    % steerableAdaptiveResolutionOrientationSpaceDetector.m
    %
%
% Copyright (C) 2019, Jaqaman Lab - UT Southwestern, Goldman Lab - Northwestern 
%
% This file is part of AdaptiveResolutionOrientationSpace.
% 
% AdaptiveResolutionOrientationSpace is free software: you can redistribute it and/or modify
% it under the terms of the GNU General Public License as published by
% the Free Software Foundation, either version 3 of the License, or
% (at your option) any later version.
% 
% AdaptiveResolutionOrientationSpace is distributed in the hope that it will be useful,
% but WITHOUT ANY WARRANTY; without even the implied warranty of
% MERCHANTABILITY or FITNESS FOR A PARTICULAR PURPOSE.  See the
% GNU General Public License for more details.
% 
% You should have received a copy of the GNU General Public License
% along with AdaptiveResolutionOrientationSpace.  If not, see <http://www.gnu.org/licenses/>.
% 
% 
    
    properties
        filter
        angularResponse
        n
    end
    
    properties (Transient)
        matrix
        idx
        angularGaussians
    end
    
    properties (Transient, Access = protected)
        cache
    end
    
    properties (Dependent, Hidden)
        res
        nms
        nlms
        theta
        a
        NMS
    end
    
    properties (Dependent, Hidden)
        nlms_mip
        nlms_count
        nlms_sum
        nlms_mean
        nlms_std
    end
    
    
    
    methods
        function obj = OrientationSpaceResponse(filter,angularResponse)
            if(~nargin)
                return;
            end
            obj.filter = filter;
            obj.angularResponse = angularResponse;
            obj.n = size(angularResponse,3);
        end
        
        % Response, defaults to maximum response with initial basis
        function set.res(obj,res)
            obj.cache.res = res;
        end
        function res = get.res(obj)
            if(~isfield(obj.cache,'res') || isempty(obj.cache.res))
                obj.cache.res = getMaxResponse(obj);
            end
            res = obj.cache.res;
        end
        
        % Non-Maximal Suppression, defaults to nms based on initial basis
        function set.nms(obj,nms)
            obj.cache.nms = nms;
        end
        function nms = get.nms(obj)
            if(~isfield(obj.cache,'nms') || isempty(obj.cache.nms))
                [obj.cache.nms] = nonMaximumSuppression(real(obj.res),real(obj.theta)) + ...
                                   + 1j .* nonMaximumSuppression(imag(obj.res),imag(obj.theta));
            end
            nms = obj.cache.nms;
        end
        function NMS = get.NMS(obj)
            NMS = OrientationSpaceNMS(obj);
        end
        
        function nlms = get.nlms(obj)
            if(~isfield(obj.cache,'nlms') || isempty(obj.cache.nlms))
                [obj.cache.nlms] = nonLocalMaximaSuppression(obj);
            end
            nlms = obj.cache.nlms;
        end
        
        function nlms_mip = get.nlms_mip(obj)
            nlms_mip = max(real(obj.nlms),[],3) + 1j*max(abs(imag(obj.nlms)),[],3);
        end
        
        function nlms_sum = get.nlms_sum(obj)
            nlms_sum = sum(obj.nlms,3);
        end
               
        function nlms_count = get.nlms_count(obj)
            nlms_count = sum(real(obj.nlms) > 0,3) + 1j*sum(abs(imag(obj.nlms)) > 0,3);
        end
               
        function nlms_mean = get.nlms_mean(obj)
            sum = obj.nlms_sum;
            count = obj.nlms_count;
%             nlms_mean = real(sum)./real(count) + 1j*imag(sum)./imag(count);
            nlms_mean = real(sum)./real(count);
        end
        
        function nlms_std = get.nlms_std(obj)
            m = obj.nlms_mean;
            nlms = obj.nlms;
            count = obj.nlms_count;
            d = bsxfun(@minus,nlms,m);
%             d = real(d).^2.*(real(nlms) > 0) + 1j*imag(d).^2.*(imag(nlms) > 0);
            d = real(d).^2.*(real(nlms) > 0); % + 1j*imag(d).^2.*(imag(nlms) > 0);
            nlms_std = sqrt(sum(d,3)./(count-1));
        end
        
        % Orientation, defaults to best orientation from basis
        function set.theta(obj,theta)
            obj.cache.theta = theta;
        end
        function theta = get.theta(obj)
            if(~isfield(obj.cache,'theta') || isempty(obj.cache.theta))
                [~,obj.cache.theta] = getMaxResponse(obj);
            end
            theta = obj.cache.theta;
        end
        
        % Shortcut for angularResponse
        function set.a(obj,a)
            obj.angularResponse = a;
        end
        function a = get.a(obj)
            a = obj.angularResponse;
        end
        
        function idx = get.idx(obj)
            if(isempty(obj.idx))
                obj.idx = orientationSpace.OrientationSpaceResponseIndex(obj);
            end
            idx = obj.idx;
        end
        
        function a_hat = a_hat(obj, varargin)
            if(~isfield(obj.cache,'a_hat') || isempty(obj.cache.a_hat))
                obj.cache.a_hat = fft(obj.a,[],3);
            end
            a_hat = obj.cache.a_hat(varargin{:});
        end
        
        function nlms = nonLocalMaximaSuppression(obj, theta, suppressionValue)
            A = real(obj.angularResponse);
            if(nargin < 2)
                theta = 0:obj.n-1;
                theta = theta*pi/obj.n;
            elseif(isscalar(theta))
                if(~mod(theta,1))
                    % integer value
                    A = orientationSpace.upsample(A,pi/theta);
                else
                    A = orientationSpace.upsample(A,theta);
                end
            end
            if(nargin < 3)
                suppressionValue = 0;
            end
            nlms = nonLocalMaximaSuppression(A,theta, suppressionValue);
        end
        
        function [nlms_precise,varargout] = nonLocalMaximaSuppressionPrecise(obj, theta, suppressionValue,interpMethod,mask, offsetAngle, angleMultiplier)
            if(nargin < 2 || isempty(theta))
                theta = obj.getRidgeOrientationLocalMaxima;
            end
            if(nargin < 3 || isempty(suppressionValue))
                suppressionValue = 0;
            end
            if(nargin < 4 || isempty(interpMethod))
                interpMethod = 'cubic';
            end
            if(nargin < 5)
                mask = [];
            end
            if(nargin < 6 || isempty(offsetAngle))
                offsetAngle = theta;
            end
	    if(nargin < 7 || isempty(angleMultiplier))
		angleMultiplier = 3;
	    end
            [nlms_precise,varargout{1:nargout-1}] = nonLocalMaximaSuppressionPrecise(real(obj.a),theta,suppressionValue,interpMethod,mask,offsetAngle, angleMultiplier);
        end
               
        function A = getAngularGaussians(obj)
            if(isempty(obj.angularGaussians))
                N = obj.n;
                x = 0:N-1;
                xx = bsxfun(@minus,x,x');
                xx = wraparoundN(xx,-N/2,N/2);
                obj.angularGaussians = exp(-xx.^2/2);
            end
            A = obj.angularGaussians;
        end
        function M = getMatrix(obj)
            if(isempty(obj.matrix))
                A = obj.getAngularGaussians;
                aR = obj.angularResponse;
                obj.matrix = reshape(aR,size(aR,1)*size(aR,2),size(aR,3))/A;
            end
            M = obj.matrix;
        end
        function [response,samples] = getResponseAtPoint(obj,r,c,samples)
            if(nargin < 4)
                samples = obj.n;
            end
            if(isscalar(samples))
                response = interpft(squeeze(real(obj.angularResponse(r,c,:))),samples).';
                if(nargout > 1)
                    samples = (0:samples-1).'.*obj.n/samples;
                end
            else
                response = interpft1(squeeze(real(obj.angularResponse(r,c,:))),samples);
            end
%             siz = size(obj.angularResponse);
%             if(nargin > 2)
%                 linIdx = sub2ind(siz([1 2]),r,c);
%             else
%                 linIdx = r;
%             end
%             if(isscalar(samples) && samples == obj.n)
%                 a = reshape(obj.angularResponse,[],obj.n);
%                 response = squeeze(a(linIdx,:));
%                 samples = (0:samples-1)'*obj.n/samples;
%             else
%                 if(isscalar(samples))
%                     if(mod(samples,1) == 0)
%                         samples = (0:samples-1)'*obj.n/samples;
%                     else
%                         samples = pi/samples;
%                         samples = (0:samples-1)'*obj.n/samples;
%                     end
%                 end
%                 M = obj.getMatrix;
%                 sampling = bsxfun(@minus,0:obj.n-1,samples);
%                 sampling = wraparoundN(sampling,-obj.n/2,obj.n/2)';
%                 sampling = exp(-sampling.^2/2);
%                 response = M(linIdx,:)*sampling;
%             end
        end
        function response = getResponseAtOrientation(obj,angle)
        %getResponseAtOrientation(orientationAngle)
            % Returns the response image plane at the orientation specified
            % in radians
            M = obj.getMatrix;
            if(isinteger(angle) && angle ~= 0)
                %do nothing
                angle = double(angle);
            else
                angle = double(angle)/pi*obj.n;
            end
            xt = bsxfun(@minus,0:obj.n-1,angle)';
            xt = wraparoundN(xt,-obj.n/2,obj.n/2);
            response = M*exp(-xt.^2/2);
            response = reshape(response,size(obj.angularResponse,1),size(obj.angularResponse,2));
            if(abs(wraparoundN(angle,-pi,pi))/pi > 0.5)
                response = real(response) - 1j*imag(response);
            end
        end
        function response = rotateResponse(obj,angle)
            if(isscalar(obj))
                % TODO: Edge response
                a_hat = fft(real(obj.angularResponse),[],3);
                nn = [0:obj.n/2 -fliplr(1:obj.n/2)];
                nn = shiftdim(nn(:),-2);
                a_hat = bsxfun(@times,a_hat,exp(-1i.*nn*angle*2));
                a = ifft(a_hat,[],3);
                response = OrientationSpaceResponse(obj.filter,a);
            else
                response(numel(obj)) = OrientationSpaceResponse;
                for ii = 1:numel(obj)
                    response(ii) = obj(ii).rotateResponse(angle);
                end
                response = reshape(response,size(obj));
            end
        end
        function [response,theta] = getMaxResponse(obj,nn)
            if(nargin < 2)
                nn = obj.n;
            end
            if(isfinite(nn))
                [response,theta] = obj.getMaxFiniteResponse(nn);
            else
                [theta,response] = orientationSpace.maxima(obj.angularResponse);
            end
        end
        function [response,maxima] = getMaxResponseFT(obj,maxima)
            if(nargin < 2)
                maxima = obj.getRidgeOrientationLocalMaxima;
            end
            response = interpft1(obj,maxima(:,:,1));
        end
        function [response,theta] = getMaxFiniteResponse(obj,nn)
            a = obj.angularResponse;
            if(obj.n ~= nn)
                orientationSpace.upsample(a,pi/nn);
            end
            [response,theta] = max(real(a),[],3);
            [response_i,theta_i] = max(cat(3,imag(a),-imag(a)),[],3);
            response = response +1j*response_i;
            theta = theta + 1j*theta_i;
            
            outAnglesRidge = wraparoundN(0:  nn-1,-nn/2,nn/2) * pi/nn;
            outAnglesEdge  = wraparoundN(0:2*nn-1,-nn,  nn)   * pi/nn;
            
            if(nargout > 1)
                theta = outAnglesRidge(real(theta)) + 1j*outAnglesEdge(imag(theta));
            end
        end
        function Response = getResponseAtOrder(obj,K_new)
            assert(~mod(K_new*2,1), ...
                'OrientationSpaceResponse:getResponseAtOrder', ...
                'Kf_new*2 must be an integer value');
            if(~isscalar(obj))
                Response(numel(obj)) = OrientationSpaceResponse;
                for o=1:numel(obj)
                    Response(o) = obj(o).getResponseAtOrder(K_new);
                end
                Response = reshape(Response,size(obj));
                return;
            end
%             A = obj.getAngularGaussians;
            % Calculate new number of angles at new order
            n_new = 2*K_new+1;
            scaleFactor = obj.n / n_new;
            % Do we need to wraparound twice?
            queryPts = wraparoundN((0:n_new-1)*scaleFactor,[-obj.n obj.n]/2);
            tt = wraparoundN(bsxfun(@minus,queryPts,(0:obj.n-1)'),[-obj.n obj.n]/2);
            T = exp(-tt.^2/2/(scaleFactor*scaleFactor));
            % Calculate new angular response at Kf_new order
            M = obj.getMatrix();
            angularResponseSize = size(obj.angularResponse);
            angularResponse_new = real(M) * T;
            angularResponse_new = reshape(angularResponse_new,[angularResponseSize([1 2]) n_new]);
            % Deal with edge response if it exists
            if(~isreal(M))
                imagM = imag(cat(3,obj.angularResponse,-obj.angularResponse));
                imagResponse = OrientationSpaceResponse(obj.filter,imagM);
                imagResponse = imagResponse.getResponseAtOrder(K_new*2+0.5);
                angularResponse_new = angularResponse_new + 1j * imagResponse.angularResponse(:,:,1:end/2);
            end
            % Create objects and return
            filter_new = OrientationSpaceFilter(obj.filter.f_c,obj.filter.b_f,K_new);
            Response = OrientationSpaceResponse(filter_new,angularResponse_new);
        end
        function Response = getResponseAtOrderFT(obj,K_new,normalize)
            if(nargin < 3)
                normalize = 0;
            end
            if(~isscalar(obj))
                Response(numel(obj)) = OrientationSpaceResponse;
                for o=1:numel(obj)
                    Response(o) = obj(o).getResponseAtOrderFT(K_new);
                end
                Response = reshape(Response,size(obj));
                return;
            end
            if(K_new == obj.filter.K)
                % copy the response object
%                 Response = OrientationSpaceResponse(obj.filter,obj.angularResponse);
                % For consistency at the moment, return only the real
                % (ridge) component 2017/11/28
                Response = OrientationSpaceResponse(obj.filter,real(obj.angularResponse));
                return;
            end
%             n_new = 2*obj.filter.sampleFactor*K_new+1;

            % Just for Gaussian calculation;
            n_new = 2*K_new+1;
            n_old = 2*obj.filter.K+1;

            % Shouldn't be based off K and not n
            s_inv = sqrt(n_old^2*n_new.^2/(n_old.^2-n_new.^2));
            s_hat = s_inv/(2*pi);
%             
            if(normalize == 2)
%                 x = -obj.filter.sampleFactor*ceil(obj.filter.K):ceil(obj.filter.K)*obj.filter.sampleFactor;
                x = (1:obj.n)-floor(obj.n/2+1);
            else
                x = -obj.filter.sampleFactor*ceil(K_new):ceil(K_new)*obj.filter.sampleFactor;
            end
            if(s_hat ~= 0)
                f_hat = exp(-0.5 * (x./s_hat).^2); % * obj.n/n_new;
                f_hat = ifftshift(f_hat);
            else
                f_hat = 1;
            end
            f_hat = shiftdim(f_hat,-1);
            a_hat = fft(real(obj.a),[],3);
            % This reduces the number of coefficients in the Fourier domain
            % Beware of the normalization factor, 1/n, done by ifft
            if(normalize < 2)
                a_hat = a_hat(:,:,[1:ceil(K_new)*obj.filter.sampleFactor+1 end-ceil(K_new)*obj.filter.sampleFactor+1:end]);
            end
            a_hat = bsxfun(@times,a_hat,f_hat);
            filter_new = OrientationSpaceFilter(obj.filter.f_c,obj.filter.b_f,K_new);
            % Consider using fft rather than ifft so that the mean is
            % consistent
            if(normalize == 1)
                Response = OrientationSpaceResponse(filter_new,ifft(a_hat*size(a_hat,3)/size(obj.a,3),[],3));
            else
                Response = OrientationSpaceResponse(filter_new,ifft(a_hat,[],3));
            end
        end
        function Response = getDerivativeResponse(obj,deriv,order,normalize)
            if(deriv)
                nFreq = (obj.n-1)/2;
                freq = [0:nFreq -nFreq:-1];
                dda = ifft(bsxfun(@times,fft(real(obj.a),[],3),(shiftdim(freq,-1)*1i).^deriv),[],3);
                Response = OrientationSpaceResponse(obj.filter,dda);
            else
                Response = OrientationSpaceResponse(obj.filter,obj.a);
            end
            if(nargin > 2)
                Response = Response.getResponseAtOrderFT(order,normalize);
            end
        end
        function response = getDerivativeResponseAtPoint(obj,r,c,deriv,order)
            response = obj.getResponseAtOrderFTatPoint(r,c,order);
            nFreq = (obj.n-1)/2;
            freq = [0:nFreq -nFreq:-1];
            deriv = shiftdim(deriv,-1);
            response = ifft(bsxfun(@times,fft(real(response)),(shiftdim(freq,1)*1i).^deriv));
        end
        function response = getResponseAtOrderFTatPoint(obj,r,c,K_new)
            % Get response at a lower order using Fourier Transform at a particular point
            % INPUT
            % r - row, scalar integer
            % c - column, scalar integer
            % K_new - new angular order
            % OUTPUT
            % response at pixel (r,c) at angular order K_new

            % New number of coefficients
%             n_new = 2*ceil(K_new)+1;
            n_new = 2*K_new+1;
            n_old = 2*obj.filter.K+1;

            % The convolution of two Gaussians results in a Gaussian
            % The multiplication of two Gaussians results in a Gaussian
            % The signal has been convolved with a Gaussian with sigma = pi/obj.n
            % Compute the signal convoled with a Gaussian with sigma = pi/n_new
            % Note if n == n_new, we divide by 0. Then s_inv = Inf
            
            s_inv = sqrt(n_old^2.*n_new.^2./(n_old.^2-n_new.^2));
            s_hat = s_inv/(2*pi);
%             x = -ceil(obj.filter.K)*obj.filter.sampleFactor:ceil(obj.filter.K)*obj.filter.sampleFactor;
            x = (1:obj.n)-floor(obj.n/2+1);
            
            % Each column represents a Gaussian with sigma set to s_hat(column)
            f_hat = exp(-0.5 * bsxfun(@rdivide,x(:),s_hat).^2); % * obj.n/n_new;
            f_hat = ifftshift(f_hat,1);
            
            % Angular response will be in a column
            a_hat = fft(squeeze(real(obj.a(r,c,:))));
            % Each column represents an angular response with order K_new(column)
            a_hat = bsxfun(@times,a_hat,f_hat);
            % Condition step added 2017/11/03
            a_hat(abs(a_hat) < eps*1e3) = 0;
            response = ifft(a_hat);
        end
        function varargout = getRidgeOrientationLocalMaxima(obj,sorted)
            if(nargin < 2)
                sorted = true;
            end
            % TODO: edge local maxima
            [varargout{1:nargout}] = interpft_extrema(real(obj.a),3,sorted);
            varargout{1} = varargout{1}/2;
            if(nargout > 1)
                varargout{2} = varargout{2}/2;
            end
            if(nargout > 4)
                varargout{5} = varargout{5}/2;
            end
        end
        function singleOrientation = getSingleRidgeOrientation(obj)
            a_hat = fft(real(obj.a),[],3);
            singleOrientation = -angle(a_hat(:,:,2))/2;
            singleOrientation = wraparoundN(singleOrientation,0,pi);
        end
        function vq  = interpft1(obj,xq)
            % interpft1 - Interpolate
            if(ndims(xq) == 3)
                xq = shiftdim(xq,2);
            else
                xq = shiftdim(xq,-1);
            end
            if(isreal(xq))
                % Only ridge interpolation is requested
                vq = interpft1([0 pi],shiftdim(obj.a,2),xq,'horner');
            else
                % Ridge
                vq = interpft1([0 pi],shiftdim(real(obj.a),2),xq,'horner');
                % and Edge interpolation is requested
                vq = interpft1([0 2*pi],shiftdim(imag(obj.a),2),imag(xq),'horner')*1i + vq;
            end
            vq = shiftdim(vq,1);
        end
        function fineResponseGrid = getResponseForInterpolation(obj,scaleFactor)
            %getResponseForInterpolation
            % INPUT
            % scaleFactor - multiplier for how grid to create
            % OUTPUT
            % fineResponseGrid - response interpolated using interpft at
            % scaleFactor
            if(nargin < 2)
                scaleFactor = 3;
            end
            % Create fine grid at scaleFactor times canonical grid
            % See Boyd, Chebyshev and Fourier Spectral Methods, Second Edition
            % (Revised), Dover, 2001. Toronto. ISBN 0-486-41183-4, page 198
            fineResponseGrid = interpft(real(obj.a),scaleFactor*obj.n,3);
            % Append last and first since orientation is periodic
            fineResponseGrid = fineResponseGrid(:,:,[end 1:end 1]);
            % TODO: pad spatial dimensions like in N(L)MS
        end
        function asymSignal = evaluateAssymetricSignalAtPoint(obj,r,c,distance,nSamples)
            if(nargin < 4)
                distance = 2/obj.filter.f_c/2/pi;
            end
            if(nargin < 5)
                nSamples = obj.n*6;
                scaleFactor = 3;
            else
                scaleFactor = max(nSamples/obj.n/2,3);
            end
            % Angle in space from point
            theta = (0:nSamples-1)*(2*pi/nSamples);
            % Orientation angle index
            orientation = mod(0:nSamples-1,nSamples/2) .* scaleFactor*obj.n/nSamples*2;
            % Add 2 for wrapping elements
            orientation = orientation+2;
%             orientation = [orientation orientation]+2;
            coords = [r-cos(theta')*distance c+sin(theta')*distance orientation'];
            % TODO: pad spatial dimensions like in N(L)MS
            A = obj.getResponseForInterpolation(scaleFactor);
            asymSignal = interp3(A,coords(:,2),coords(:,1),coords(:,3),'cubic');
            if(nargout == 0)
                plot((0:nSamples-1)/nSamples*2,asymSignal);
                xlabel('Orientation (\pi radians)');
            end
        end
        function R = real(obj)
            R = OrientationSpaceResponse(real(obj.filter),real(obj.angularResponse));
        end
        function R = imag(obj)
            R = OrientationSpaceResponse(imag(obj.filter),imag(obj.angularResponse));
        end
        function h = imshow(obj,varargin)
            normalize = false;
            if(~isempty(varargin) && isempty(varargin{1}) && numel(obj) > 1)
                    normalize = true;
            end
            outI = cell(size(obj));
            for o=1:numel(obj)
                outI{o} = obj(o).getMaxResponse;
                if(normalize)
                    outI{o} = mat2gray(real(outI{o}));
                end
            end
            h = imshow(cell2mat(outI),varargin{:});
        end
        function h = imshowpair(A,B)
            if(nargin > 1)
                if(isa(B,'OrientationSpaceResponse'))
                    B = real(B.res);
                end
            else
                B = imag(A.res);
            end
            if(isa(A,'OrientationSpaceResponse'))
                A = real(A.res);
            end
            h = imshowpair(A,B);
        end
        function h = plot(obj,angles,r,c,varargin)
            holdState = ishold;
            hold on;
            for o=1:numel(obj)
                [Y,samples] = obj(o).getResponseAtPoint(r,c,angles);
                h = plot(samples/obj(o).n,Y,varargin{:});
            end
            if(~holdState)
                hold off;
            end
        end
        function h = polar(obj,angles,r,c,varargin)           
            holdState = ishold;
            for o = 1:numel(obj)
                [Y,samples] = obj(o).getResponseAtPoint(r,c,angles);
                samples = [samples ; samples+obj(o).n];
                Y = [ Y Y ];
                h = polarplot(samples'/obj(o).n*pi,Y,varargin{:});
                ax = gca;
                set(ax,'ThetaZeroLocation','top')
                set(ax,'ThetaDir','clockwise')
                hold on;
                select = Y < 0;
                if(any(select))
                    addBreaks = diff(select) == 1;
                    select(addBreaks) = true;
                    Y(addBreaks) = NaN;
                    h(2) = polarplot(samples(select)'/obj(o).n*pi,Y(select),varargin{:});
                    set(h(2),'LineStyle','--','Color','w');
                end
            end
            if(~holdState)
                hold off;
            end
        end
        function A = getArraySpace(obj,varargin)
            % Useful if using a multiscale filter array
            % Concatenate along the next available dimension
            d = ndims(obj(1).a)+1;
            d = max(d,4);
            if(nargin < 2)
                A = cat(d,obj.a);
                sA = size(A);
                A = reshape(A,[sA(1:d-1) size(obj)]);
                A = A(varargin{:});
            else
                varargin(nargin:4) = {':'};
                A = arrayfun(@(R) R.a(varargin{1:3}),obj,'UniformOutput',false);
                A = cat(d,A{varargin{4}});
                sA = size(A);
                if(varargin{4}(1) == ':')
                    A = reshape(A,[sA(1:d-1) size(obj)]);
                end
            end
        end
        function E = getRidgeAngularEnergy(obj)
            a_hat = fft(real(obj.a),[],3);
            E = sum(abs(a_hat(:,:,2:end)).^2,3)./(obj.n.^2);
        end
        [] = animateAngularOrder(R, r, c, Rd);
        [ localMaxima, localMaximaValue, K ] = traceLocalMaxima( obj, r, c, K, polish );
        [ localMaxima, localMaximaValue, K ] = traceLocalMaximaHouseholder( obj, r, c, K );
        [ maxima, iter ] = traceAllLocalMaxima( obj, K, maxima_org);
        [ maxima, iter ] = traceAllLocalMaximaAtPoint( obj, K , r, c);
        [ allMaxima, maximaTraceRegimesInv, maximaTraceRegimesInvK ] = findRegimesAtPoint( R, r, c, K );
    end
    
end
```
