## Supplementary material for "Adaptive-Resolution Multi-Orientation Analysis of Complex Filamentous Network Images": MATLAB code from Github: traceAllLocalMaxima.m

```
function [ maxima, iter ] = traceAllLocalMaxima( obj, K , maxima_org)
%traceAllLocalMaxima Summary of this function goes here
%   Detailed explanation goes here
%
% Copyright (C) 2019, Jaqaman Lab - UT Southwestern, Goldman Lab - Northwestern 
%
% This file is part of AdaptiveResolutionOrientationSpace.
% 
% AdaptiveResolutionOrientationSpace is free software: you can redistribute it and/or modify
% it under the terms of the GNU General Public License as published by
% the Free Software Foundation, either version 3 of the License, or
% (at your option) any later version.
% 
% AdaptiveResolutionOrientationSpace is distributed in the hope that it will be useful,
% but WITHOUT ANY WARRANTY; without even the implied warranty of
% MERCHANTABILITY or FITNESS FOR A PARTICULAR PURPOSE.  See the
% GNU General Public License for more details.
% 
% You should have received a copy of the GNU General Public License
% along with AdaptiveResolutionOrientationSpace.  If not, see <http://www.gnu.org/licenses/>.
% 
% 

if(nargin < 3)
    maxima_org = obj.getRidgeOrientationLocalMaxima;
end

    maxima_org = shiftdim(maxima_org,2)*2;
    
    maxima = cell(1,length(K));
    iter = cell(1,length(K));
    
    [K,sort_idx] = sort(K,'descend');
    
    Kidx = 1:length(K);
    Kidx(obj.filter.K ==K) = [];

    pool = gcp('nocreate');
    if(isempty(pool))
        prev_maxima = maxima_org;
        % Consider sorting to avoidNans per halleyft
        for ki=Kidx
            [maxima{ki},xgd,iter{ki}] = halleyft(shiftdim(obj.getResponseAtOrderFT(K(ki),2).a,2),prev_maxima,false,1,1e-12,10,false);
            maxima{ki}(xgd(:,:,2) > 0) = NaN;
            prev_maxima = maxima{ki};
        end
%         maxima{obj.filter.K ==K} = maxima_org;
    else
        maxima_org_sz = size(maxima_org);
        prev_maxima = maxima_org(:,:);

        pool_size = pool.NumWorkers;
        block_size = ceil(size(prev_maxima,2)/pool_size);
        partition = [repmat(block_size,1,floor(size(prev_maxima,2)/block_size)) mod(size(prev_maxima,2),block_size)];
        partition = partition(partition ~= 0);
        
        prev_maxima = mat2cell(prev_maxima,maxima_org_sz(1),partition);
        
        response = cell(length(K),length(partition));
        for ki=Kidx
            response{ki,1} = shiftdim(obj.getResponseAtOrderFT(K(ki),2).a,2);
            response{ki,1} = response{ki,1}(:,:);
            response(ki,:) = mat2cell(response{ki,1},size(response{ki},1),partition);
        end
        response = num2cell(response,1);
        
        maxima = parcellfun_progress(@(r,m) halleyftDescent(r,m,Kidx),response,prev_maxima,'UniformOutput',false);
        maxima = vertcat(maxima{:});
        maxima = num2cell(maxima,1);
        
%         for ki=Kidx
%             response = shiftdim(obj.getResponseAtOrderFT(K(ki),2).a,2);
%             response = response(:,:);
%             response = mat2cell(response,size(response,1),partition);
%             maxima{ki} = parcellfun_progress(@(r,m) halleyft(r,m,false,1),response,prev_maxima,'UniformOutput',false);
%             prev_maxima = maxima{ki};
%         end
        for ki=Kidx
            maxima{ki} = [maxima{ki}{:}];
            maxima{ki} = reshape(maxima{ki},maxima_org_sz);
        end
%         maxima{obj.filter.K ==K} = maxima_org;
    end
    isOrgK = obj.filter.K == K;
    if(any(isOrgK))
        maxima{obj.filter.K ==K} = maxima_org;
    end
    
    maxima(sort_idx) = maxima;
    maxima = cat(4,maxima{:});
    maxima = maxima/2;
    maxima = permute(maxima,[2 3 1 4]);
    
    %% Eliminate duplicates
    duplicateThreshold = 1e-6;
    [maxima,sort_idx] = sort(maxima,3);
    maximaDiff = diff(maxima,1,3);
    maximaDiff = cat(3,maximaDiff,min(maxima,[],3)+pi-max(maxima,[],3));
    maximaDuplicates = abs(maximaDiff) < duplicateThreshold;
%     maximaDuplicates(:,:,end+1,:) = false;
    maxima(maximaDuplicates) = NaN;
    
    %% Re-sort according to orignal alignment
    o = ones(size(sort_idx));
    sort_idx = sub2ind(size(sort_idx),(1:size(sort_idx,1)).'.*o,(1:size(sort_idx,2)).*o,sort_idx,shiftdim(1:size(sort_idx,4),-2).*o);
    maxima(sort_idx) = maxima;
    
    %% Remove interstitial NaN
    [~,sort_idx] = sort(isnan(maxima),3);
    sort_idx = sub2ind(size(sort_idx),(1:size(sort_idx,1)).'.*o,(1:size(sort_idx,2)).*o,sort_idx,shiftdim(1:size(sort_idx,4),-2).*o);
    maxima = maxima(sort_idx);
%     maxima = sort(maxima,3);
    
    if(nargout > 1)
        iter{obj.filter.K == K} = NaN(size(maxima_org));
        iter(sort_idx) = iter;
        iter = cat(4,iter{:});
        iter = permute(iter,[2 3 1 4]);
    end

end
function maxima = halleyftDescent(response,prev_maxima,Kidx)
    maxima = cell(1,length(response));
    for ki=Kidx
        [maxima{ki},xgd] = halleyft(response{ki},prev_maxima,false,1,1e-12,10,false);
        maxima{ki}(xgd(:,:,2) > 0) = NaN;
        prev_maxima = maxima{ki};
    end
end
```
