## Supplementary material for "Adaptive-Resolution Multi-Orientation Analysis of Complex Filamentous Network Images": MATLAB code from Github: traceLocalMaxima.m

```
function [ localMaxima, localMaximaValue, K ] = traceLocalMaxima( obj, r, c, K, polish )
%traceLocalMaxima Traces local maxima through lower angular orders
%
% Copyright (C) 2019, Jaqaman Lab - UT Southwestern, Goldman Lab - Northwestern 
%
% This file is part of AdaptiveResolutionOrientationSpace.
% 
% AdaptiveResolutionOrientationSpace is free software: you can redistribute it and/or modify
% it under the terms of the GNU General Public License as published by
% the Free Software Foundation, either version 3 of the License, or
% (at your option) any later version.
% 
% AdaptiveResolutionOrientationSpace is distributed in the hope that it will be useful,
% but WITHOUT ANY WARRANTY; without even the implied warranty of
% MERCHANTABILITY or FITNESS FOR A PARTICULAR PURPOSE.  See the
% GNU General Public License for more details.
% 
% You should have received a copy of the GNU General Public License
% along with AdaptiveResolutionOrientationSpace.  If not, see <http://www.gnu.org/licenses/>.
% 
% 

if(nargin < 4 || isempty(K))
    K = obj.filter.K:-0.1:-0.1;
end
if(nargin < 5)
    polish = false;
end

% Orders of each polynomial term
orders = [0:ceil(obj.filter.K) -ceil(obj.filter.K):-1].';
orders2 = orders.^2;
orders_1i = orders.*1i;

p = squeeze(real(obj.a(r,c,:)));
p_hat = fft(p);

% % First derivative
p_hat(:,2) = p_hat.*orders_1i;
% % Second derivative
% p_hat(:,3) = p_hat(:,2).*orders_1i;
% % Third Derivative
% p_hat(:,4) = p_hat(:,3).*orders_1i;
% 
% maxima = interpft_extrema(p);
% maxima = maxima(~isnan(maxima));

% Polish zeros of first derivative
% for ii=1:length(maxima)
%     maxima(ii) = fzero(@(m) interpft1([0 2*pi],p_hat(:,2),m,'horner_freq'),maxima(ii));
% end

buffer = zeros(length(p_hat),length(K));
bufferd = buffer;


for k =1:length(K)
    n_new = 2*K(k)+1;
    s_hat_2 = obj.n^2.*n_new.^2./(obj.n.^2-n_new.^2);
    s_hat_2 = s_hat_2/(4*pi*pi);
    s_hat_2 = s_hat_2.';
    f_hat = exp(-0.5 * bsxfun(@rdivide,orders2,s_hat_2));
    p_hat_new = bsxfun(@times,p_hat,f_hat);
    buffer(:,k) = p_hat_new(:,1);
    bufferd(:,k) = p_hat_new(:,2);
end

% maxima_all = interpft_extrema(ifft(buffer));
maxima_all = interpft_extrema(buffer,[],[],[],false);

maxima = maxima_all(~isnan(maxima_all(:,1)),1);

localMaxima = NaN(length(K),length(maxima));
localMaximaValue = localMaxima;

for k =1:length(K)    
%     maxima = interpft_extrema(ifft(p_hat_new(:,1)));
    maxima = maxima_all(:,k); 
    maxima = maxima(~isnan(maxima));
    
    
%     tic;
    if(polish)
        for ii=1:length(maxima)
            localMaxima(k,ii) = fzero(@(m) interpft1([0 2*pi],bufferd(:,k),m,'horner_freq'),maxima(ii));
        end
    else
        localMaxima(k,1:length(maxima)) = maxima;
    end
    
%     if(nargout > 1)
%         localMaximaValue(k,1:length(maxima)) = interpft1([0 2*pi],buffer(:,k),localMaxima(k,1:length(maxima)),'horner_freq');
%     end

%     toc
%     tic
%     actual = interpft_extrema(ifft(p_hat_new(:,1)));
%     toc
%     maxima = unique(localMaxima(k,:));
%     hold on;
%     plot(interpft(ifft(p_hat_new(:,1)),360));
%     pause(0.1);
end

if(nargout > 1)
    localMaximaValue = interpft1([0 2*pi],buffer,localMaxima.','horner_freq').';
end

end
```
