## Supplementary material for "Adaptive-Resolution Multi-Orientation Analysis of Complex Filamentous Network Images": MATLAB code from Github: traceLocalMaximaHouseholder.m

```
function [ localMaxima, localMaximaValue, K ] = traceLocalMaximaHouseholder( obj, r, c, K )
%traceLocalMaxima Traces local maxima through lower angular orders
%
% Copyright (C) 2019, Jaqaman Lab - UT Southwestern, Goldman Lab - Northwestern 
%
% This file is part of AdaptiveResolutionOrientationSpace.
% 
% AdaptiveResolutionOrientationSpace is free software: you can redistribute it and/or modify
% it under the terms of the GNU General Public License as published by
% the Free Software Foundation, either version 3 of the License, or
% (at your option) any later version.
% 
% AdaptiveResolutionOrientationSpace is distributed in the hope that it will be useful,
% but WITHOUT ANY WARRANTY; without even the implied warranty of
% MERCHANTABILITY or FITNESS FOR A PARTICULAR PURPOSE.  See the
% GNU General Public License for more details.
% 
% You should have received a copy of the GNU General Public License
% along with AdaptiveResolutionOrientationSpace.  If not, see <http://www.gnu.org/licenses/>.
% 
% 

if(nargin < 4 || isempty(K))
    K = obj.filter.K:-0.1:-0.1;
end

% Orders of each polynomial term
orders = [0:ceil(obj.filter.K) -ceil(obj.filter.K):-1].';
orders2 = orders.^2;
orders_1i = orders.*1i;

p = squeeze(real(obj.a(r,c,:)));
p_hat = fft(p);

% % First derivative
p_hat(:,2) = p_hat.*orders_1i;
% % Second derivative
p_hat(:,3) = p_hat(:,2).*orders_1i;
% % Third Derivative
p_hat(:,4) = p_hat(:,3).*orders_1i;
% 
maxima = interpft_extrema(p);
maxima = maxima(~isnan(maxima));

% Polish zeros of first derivative
maxima = halleyRootFinder(maxima(:),p_hat(:,2),p_hat(:,3),p_hat(:,4),eps);
% for ii=1:length(maxima)
%     maxima(ii) = halleyRootFinder(maxima(ii),p_hat(:,2),p_hat(:,3),p_hat(:,4),1e-14);
% end

buffer = zeros(length(p_hat),length(K));
bufferd = buffer;

lastK = obj.filter.K;

p_hat_old = p_hat;

localMaxima = NaN(length(K),length(maxima));

for k =1:length(K)
    n_new = 2*K(k)+1;
    s_hat_2 = obj.n^2.*n_new.^2./(obj.n.^2-n_new.^2);
    s_hat_2 = s_hat_2/(4*pi*pi);
    s_hat_2 = s_hat_2.';
    f_hat = exp(-0.5 * bsxfun(@rdivide,orders2,s_hat_2));
    p_hat_new = bsxfun(@times,p_hat,f_hat);
    buffer(:,k) = p_hat_new(:,1);
%     bufferd(:,k) = p_hat_new(:,2);
    deriv_values = interpft1([0 2*pi],p_hat_old(:,3:4),maxima,'horner_freq');
    guess = maxima(:)-deriv_values(:,2)./deriv_values(:,1)./(2*k+1).^3*8*pi.^2*(K(k)-lastK);
%     guess = maxima(:)-interpft1([0 2*pi],p_hat_old(:,4),maxima,'horner_freq')./interpft1([0 2*pi],p_hat_old(:,3),maxima,'horner_freq')./(2*k+1).^3*8*pi.^2*(K(k)-lastK);
    maxima = halleyRootFinder(guess(:),p_hat_new(:,2),p_hat_new(:,3),p_hat_new(:,4),eps);
    localMaxima(k,:) = maxima;
end

if(nargout > 1)
    localMaximaValue = interpft1([0 2*pi],buffer,localMaxima.','horner_freq').';
end

% maxima_all = interpft_extrema(ifft(buffer));
% % buffer = interpft_extrema(buffer,[],[],[],false);
% 
% maxima = maxima_all(~isnan(maxima_all(:,1)),1);
% 
% localMaxima = NaN(length(K),length(maxima));
% localMaximaValue = localMaxima;
% 
% for k =1:length(K)    
% %     maxima = interpft_extrema(ifft(p_hat_new(:,1)));
%     maxima = maxima_all(:,k); 
%     maxima = maxima(~isnan(maxima));
%     
%     
% %     tic;
%     if(polish)
%         for ii=1:length(maxima)
%             localMaxima(k,ii) = fzero(@(m) interpft1([0 2*pi],bufferd(:,k),m,'horner_freq'),maxima(ii));
%         end
%     else
%         localMaxima(k,1:length(maxima)) = maxima;
%     end
%     
% %     if(nargout > 1)
% %         localMaximaValue(k,1:length(maxima)) = interpft1([0 2*pi],buffer(:,k),localMaxima(k,1:length(maxima)),'horner_freq');
% %     end
% 
% %     toc
% %     tic
% %     actual = interpft_extrema(ifft(p_hat_new(:,1)));
% %     toc
% %     maxima = unique(localMaxima(k,:));
% %     hold on;
% %     plot(interpft(ifft(p_hat_new(:,1)),360));
% %     pause(0.1);
% end
% 
% if(nargout > 1)
%     localMaximaValue = interpft1([0 2*pi],buffer,localMaxima.','horner_freq').';
% end

end

function [root, root_v] = halleyRootFinder(guess,p_hat,pd_hat,pdd_hat,TOL)
% Use Halley's method to polish root guess
% Second order Householder Method
    if(nargin < 5)
        TOL = 1e-12;
    end
    MAXITER = 10;
%     p_v = interpft1([0 2*pi],p_hat,guess,'horner_freq');
    done = false(size(guess));
    
    root = guess;
    % place holder
%     root_v = guess;

    v = zeros(length(guess),3);
    
    p_hat = [p_hat pd_hat pdd_hat];
    for i=1:MAXITER
        % only size of roots being refined
        v(~done,:) = interpft1([0 2*pi],p_hat,guess(~done),'horner_freq');
        
        if(i == 1)
            root_v = v(:,1);
        else
            
            % Find values that moved away from zero
            regressed = abs(root_v) <= abs(v(:,1)) | v(:,2) > 0;
            done(regressed) = true;
            
            % Only update values that did regress
            root(~regressed) = guess(~regressed);
            root_v(~regressed) = v(~regressed,1);
            
            % Check if update is below tolerance
            done(~done) = abs(root_v(~done)) < TOL;
            
%             v = v(~done,:);
            
            if(all(done))
                break;
            end
        end
        
        p_v = v(~done,1);
        pd_v = v(~done,2);
        pdd_v = v(~done,3);
        
        % only size of roots being refined
%         pdd_v = interpft1([0 2*pi],pdd_hat,guess(~done),'horner_freq');
        % root is same size as guess
        guess(~done) = root(~done) - 2*p_v.*pd_v./( 2*pd_v.*pd_v - p_v.*pdd_v);
        % root_v is same size as guess
%         root_v(~done) = interpft1([0 2*pi],p_hat,root(~done),'horner_freq');

%         regressed = abs(root_v) >= abs(p_v);
%         root(regressed) = guess(regressed);
%         root_v(regressed) = p_v(regressed);
%         done(regressed) = true;
%         
%         done(~done) = abs(root_v(~done)) >= abs(p_v(~done));
%         done(~done) = abs(root_v(~done)) < TOL;
%         
%         guess = root;
%         p_v = root_v;
%         if(all(done))
%             break;
%         end
%         if(abs(root_v) >= abs(p_v))
%             warning('OrientationSpaceResponse.traceLocalMaximaHouseholder/halleyRootFinder Not monotically decreasing');
%             root = guess;
%             root_v = p_v;
%             break;
%         end
%         guess = root;
%         p_v = root_v;
%         if(abs(root_v) < TOL)
%             break;
%         end
    end
%     disp(i);
    if(i == MAXITER)
        warning('OrientationSpaceResponse.traceLocalMaximaHouseholder/halleyRootFinder MAXITER reached');
    end
end
```
