## Supplementary material for "Adaptive-Resolution Multi-Orientation Analysis of Complex Filamentous Network Images": MATLAB code from Github: optimalHistogram.m

```
function [N,X,sp] = optimalHistogram(varargin)
% OPTIMALHISTOGRAM generates a histogram using the "optimal" number of bins
%
% This function was previously called histogram prior to R2014B
%
% If called with no output argument, optimalHistogram plots into the current axes
%
% SYNOPSIS [N,X,sp] = optimalHistogram(data,factor,normalize)
%          [...] = optimalHistogram(data,'smooth')
%          [...] = optimalHistogram(axesHandle,...)
%
% INPUT    data: vector of input data
%          factor: (opt) factor by which the bin-widths are multiplied
%                   if 'smooth' (or 's'), a smooth histogram will be formed.
%                   (requires the spline toolbox). For an alternative
%                   approach to a smooth histogram, see ksdensity.m
%                   if 'discrete' (or 'd'), the data is assumed to be a discrete
%                   collection of values. Note that if every data point is,
%                   on average, repeated at least 3 times, histogram will
%                   consider it a discrete distribution automatically.
%                   if 'continuous' (or 'c'), histogram is not automatically
%                   checking for discreteness.
%          normalize : if 1 (default), integral of histogram equals number
%                       data points. If 0, height of bins equals counts.
%                       This option is exclusive to non-"smooth" histograms
%          axesHandle: (opt) if given, histogram will be plotted into these
%                       axes, even if output arguments are requested
%
% OUTPUT   N   : number of points per bin (value of spline)
%          X   : center position of bins (sorted input data)
%          sp  : definition of the smooth spline
%
% REMARKS: The smooth histogram is formed by calculating the cumulative
%           histogram, fitting it with a smoothening spline and then taking
%           the analytical derivative. If the number of data points is
%           markedly above 1000, the spline is fitting the curve too
%           locally, so that the derivative can have huge peaks. Therefore,
%           only 1000-1999 points are used for estimation.
%           Note that the integral of the spline is almost exactly the
%           total number of data points. For a standard histogram, the sum
%           of the hights of the bins (but not their integral) equals the
%           total number of data points. Therefore, the counts might seem
%           off.
%
%           WARNING: If there are multiples of the minimum value, the
%           smooth histogram might get very steep at the beginning and
%           produce an unwanted peak. In such a case, remove the
%           multiple small values first (for example, using isApproxEqual)
%
%
% c: 2/05 jonas
%%%%%%%%%%%%%%%%%%%%%%%%%%%%%%%%%%%%%%%%%%%%%%%%%%%%%%%%%%%%%%%%%%%%%%%%%%%
%
% Copyright (C) 2019, Jaqaman Lab - UT Southwestern, Goldman Lab - Northwestern 
%
% This file is part of AdaptiveResolutionOrientationSpace.
% 
% AdaptiveResolutionOrientationSpace is free software: you can redistribute it and/or modify
% it under the terms of the GNU General Public License as published by
% the Free Software Foundation, either version 3 of the License, or
% (at your option) any later version.
% 
% AdaptiveResolutionOrientationSpace is distributed in the hope that it will be useful,
% but WITHOUT ANY WARRANTY; without even the implied warranty of
% MERCHANTABILITY or FITNESS FOR A PARTICULAR PURPOSE.  See the
% GNU General Public License for more details.
% 
% You should have received a copy of the GNU General Public License
% along with AdaptiveResolutionOrientationSpace.  If not, see <http://www.gnu.org/licenses/>.
% 
% 

% test input
if nargin < 1
    error('not enough input arguments for optimalHistogram')
end

% check for axes handle
if length(varargin{1}) == 1 && ishandle(varargin{1});
    axesHandle = varargin{1};
    varargin(1) = [];
else
    % ensure compatibility to when axesHandle was given as last input
    if nargin == 3 && ishandle(varargin{end}) && varargin{end} ~= 0
        axesHandle = varargin{end};
        varargin(end) = [];
    else
        axesHandle = 0;
    end
end

% assign data
numArgIn = length(varargin);
data = varargin{1};
data = data(:);

% check for non-finite data points
data(~isfinite(data)) = [];

% check for "factor"
if numArgIn < 2 || isempty(varargin{2})
    factor = 1;
else
    factor = varargin{2};
end
if ischar(factor)
    switch factor
        case {'smooth','s'}
        factor = -1;
        case {'discrete','d'}
            factor = -2;
        case {'continuous','c'}
            factor = -3;
    otherwise
        error('The only string inputs permitted for optimalHistogram.m are ''smooth'',''discrete'', or ''continuous''')
    end
else
    % check for normalize, but do so only if there is no "smooth". Note
    % that numArgIn is not necessarily equal to nargin
    if numArgIn < 3 || isempty(varargin{3})
        normalize = true;
    else
        normalize = varargin{3};
    end
end

% doPlot is set to 1 for now. We change it to 0 below if necessary.
doPlot = 1;

nData = length(data);
% check whether we do a standard or a smooth histogram
if factor ~= -1
    % check for discrete distribution
    [nn,xx] = getMultiplicity(data);
    % consider the distribution discrete if there are, on average, 3
    % entries per bin
    nBins = length(xx);
    if factor == -2 || (factor ~= -3 && nBins*3 < nData) 
        % discrete distribution. 
        nn = nn';
        xx = xx';
    else
        % not a discrete distribution
        if nData < 20
            warning('HISTOGRAM:notEnoughDataPoints','Less than 20 data points!')
            nBins = ceil(nData/4);
        else
            
            % create bins with the optimal bin width
            % W = 2*(IQD)*N^(-1/3)
            interQuartileDist = diff(prctile(data,[25,75]));
            binLength = 2*interQuartileDist*length(data)^(-1/3)*factor;
            
            % number of bins: divide data range by binLength
            nBins = round((max(data)-min(data))/binLength);
            
            if ~isfinite(nBins)
                nBins = length(unique(data));
            end
            
        end
        
        
        
        % histogram
        [nn,xx] = hist(data,nBins);
        % adjust the height of the histogram
        if normalize
            Z = trapz(xx,nn);
            nn = nn * nData/Z;
        end
        
    end
    if nargout > 0
        N = nn;
        X = xx;
        doPlot = axesHandle;
    end
    if doPlot
        if axesHandle
            bar(axesHandle,xx,nn,1);
        else
            bar(xx,nn,1);
        end
    end
    
else
    % make cdf, smooth with spline, then take the derivative of the spline
    
    % cdf
    xData = sort(data);
    yData = 1:nData;
    
    % when using too many data points, the spline fits very locally, and
    % the derivatives can still be huge. Good results can be obtained with
    % 500-1000 points. Use 1000 for now
    myFactor=1000;
    step = max(floor(nData/myFactor),1);
    xData2 = xData(1:step:end);
    yData2 = yData(1:step:end);
    
    % spline. Use strong smoothing
    cdfSpline = csaps(xData2,yData2,1./(1+mean(diff(xData2))^3/0.0006));
    
    % pdf is the derivative of the cdf
    pdfSpline = fnder(cdfSpline);
    
    % histogram
    if nargout > 0
        xDataU = unique(xData);
        N = fnval(pdfSpline,xDataU);
        X = xDataU;
        % adjust the height of the histogram
        Z = trapz(X,N);
        N = N * nData/Z;
        sp = pdfSpline;
        % set doPlot. If there is an axesHandle, we will plot
        doPlot = axesHandle;
    end
    % check if we have to plot. If we assigned an output, there will only
    % be plotting if there is an axesHandle.
    if doPlot
        if axesHandle
            plot(axesHandle,xData,fnval(pdfSpline,xData));
        else
            plot(xData,fnval(pdfSpline,xData));
        end
    end
end
```
