## Supplementary material for "Adaptive-Resolution Multi-Orientation Analysis of Complex Filamentous Network Images": MATLAB code from Github: nonMaximumSuppression.m

```
%nms = nonMaximumSuppression(res, th, sup)
%
% Inputs:   res : response
%            th : orientation
%           sup : value to set suppressed pixels to
%
% Uses the grid conventions of steerableDetector()
%
% Copyright (C) 2019, Jaqaman Lab - UT Southwestern, Goldman Lab - Northwestern 
%
% This file is part of AdaptiveResolutionOrientationSpace.
% 
% AdaptiveResolutionOrientationSpace is free software: you can redistribute it and/or modify
% it under the terms of the GNU General Public License as published by
% the Free Software Foundation, either version 3 of the License, or
% (at your option) any later version.
% 
% AdaptiveResolutionOrientationSpace is distributed in the hope that it will be useful,
% but WITHOUT ANY WARRANTY; without even the implied warranty of
% MERCHANTABILITY or FITNESS FOR A PARTICULAR PURPOSE.  See the
% GNU General Public License for more details.
% 
% You should have received a copy of the GNU General Public License
% along with AdaptiveResolutionOrientationSpace.  If not, see <http://www.gnu.org/licenses/>.
% 
% 

% Francois Aguet

function res = nonMaximumSuppression(res, th, sup)

if(nargin < 3)
    sup = 0;
end

[ny,nx] = size(res);

res = padarrayXT(res, [1 1], 'symmetric');

[x,y] = meshgrid(1:nx,1:ny);

% +1 interp
A1 = interp2(res, x+1+cos(th), y+1+sin(th),'linear',0);

% -1 interp
A2 = interp2(res, x+1-cos(th), y+1-sin(th),'linear',0);

res = res(2:end-1,2:end-1);

res(res<A1 | res<A2) = sup;
```
