## Supplementary material for "Adaptive-Resolution Multi-Orientation Analysis of Complex Filamentous Network Images": MATLAB code from Github: thresholdOtsu.m

```
function [level, varargout] = thresholdOtsu(imageIn,varargin)
% Select a thresholding level using Otsu's method
%
% level = thresholdOtsu(imageIn)
% level = thresholdOtsu(imageIn,showPlots)
% 
% This function selects a threshold for the input fluorescence image by
% analyzing the image's intensity distribution. This requires good signal-
% to-noise, and a significant amount of background in the image. The
% threshold is selected using Otsu's method
% 
% Input:
% 
%   imageIn - The N-Dimensional image to be thresholded.
% 
% 
%   showPlots - If true, a plot of the histogram and an overlay of the mask
%   on the image will be shown. The overlay plot only works if the image is
%   2D.
% 
% 
% Output:
% 
% 
%   level - The intensity value selected for thresholding.
%   mask (optional) -- the thresholded foreground mask is returned as 
%                      an optional output argument
%                      (added by Deepak on Mar, 2013) 
%
% Revamped from OtsuSeg
%
% Sebastien Besson, 5/2011
%
% Copyright (C) 2019, Jaqaman Lab - UT Southwestern, Goldman Lab - Northwestern 
%
% This file is part of AdaptiveResolutionOrientationSpace.
% 
% AdaptiveResolutionOrientationSpace is free software: you can redistribute it and/or modify
% it under the terms of the GNU General Public License as published by
% the Free Software Foundation, either version 3 of the License, or
% (at your option) any later version.
% 
% AdaptiveResolutionOrientationSpace is distributed in the hope that it will be useful,
% but WITHOUT ANY WARRANTY; without even the implied warranty of
% MERCHANTABILITY or FITNESS FOR A PARTICULAR PURPOSE.  See the
% GNU General Public License for more details.
% 
% You should have received a copy of the GNU General Public License
% along with AdaptiveResolutionOrientationSpace.  If not, see <http://www.gnu.org/licenses/>.
% 
% 

ip=inputParser;
ip.addRequired('imageIn',@isnumeric);
ip.addOptional('showPlots',0,@isnumeric)
ip.parse(imageIn,varargin{:});
showPlots=ip.Results.showPlots;

%Convert to double if necessary
imageIn = double(imageIn);

%find nonzero values (due to masking)
nzInd = find(imageIn);

%get minumum and maximum pixel values in image
minSignal = min(imageIn(nzInd));
maxSignal = max(imageIn(nzInd));

%normalize nonzero value between 0 and 1
imageInNorm = zeros(size(imageIn));
imageInNorm(nzInd) = (imageIn(nzInd)- minSignal) / (maxSignal - minSignal);

level = graythresh(imageInNorm);

level = level*(maxSignal - minSignal)+minSignal;

if showPlots 
    imageMask = imageIn >= level;
    figure;
    imagesc(imageIn);
    hold on
    contour(imageMask,'w')
    colormap hot
end

if nargout > 1
    varargout{1} = double(imageIn >= level);
end
```
