## Supplementary material for "Adaptive-Resolution Multi-Orientation Analysis of Complex Filamentous Network Images": MATLAB code from Github: cutFirstHistMode.m

```
function [cutoffIndex, cutoffValue, sp, axesH,maxBinValue] = cutFirstHistMode(varargin)
%CUTFIRSTHISTMODE finds the end of the first mode of a histogram using unimodal thresholding (Rosin).
%
% cutFirstHistMode is an implementation of the algorithm presented in "uni-
% modal thresholding" by P.L. Rosin, Pattern Recognition (2001); 34:2083.
% It assumes  that the first mode in the histogram (noise/background values
% for most applications) is strongest, and places the cutoff where distance
% between the line from the largest bin in the first mode to the first
% empty bin after the last nonempty bin and the histogram is largest.
%
% SYNOPSIS [cutoffIndex, cutoffValue,sp] = cutFirstHistMode(counts, bins);
%          [cutoffIndex, cutoffValue,sp] = cutFirstHistMode(data);
%          [...] = cutFirstHistMode(...,verbose)
%          [...] = cutFirstHistMode(axesH,...)
%
% INPUT    counts, bins : counts in and center of histogram bins (output
%               of functions such as "hist" or "histogram". There has to be
%               more than one bin.
%          alternatively, you can pass the data directly, and the program
%               set up the histogram
%
%          verbose (optional) decides whether the function will open a
%               figure or not (default = 1).
%          axesH (optional) axes handle into which the function will plot.
%               Supplying ax automatically sets verbose to 1. (Of course,
%               setting verbose = 0 will overrule that, but this would be a
%               bit nonsensical)
%
%
% OUTPUT   cutoffIndex : Index into list of bins/data of the placement of
%               cutoff
%          cutoffValue : position of bin/data point where the histogram is
%               cut off
%          sp : definition of the histogram spline (only possible of
%               cutFirstHistMode had to calculate a histogram)
%          axesH : plot axes. Empty if verbose is 0
%
% REMARKS  If data is supplied, cutoffIndex/cutoffData will point to the
%               data point just at or above the center of the bin
%          The function works only on 1D data so far (multidimensional data
%          is handled as a vector)
%
%          Warning: If there are multiples of the minimum value, the
%          smooth histogram might get very steep at the beginning and
%          produce an unwanted highest peak. In such a case, remove the
%          multiple small values first (for example, using isApproxEqual)
%
%
% c: jonas, 8/06   
%%%%%%%%%%%%%%%%%%%%%%%%%%%%%%%%%%%%%%%%%%%%%%%%%%%%%%%%%%%%%%%%%%%%%%%%%%%
%
% Copyright (C) 2019, Jaqaman Lab - UT Southwestern, Goldman Lab - Northwestern 
%
% This file is part of AdaptiveResolutionOrientationSpace.
% 
% AdaptiveResolutionOrientationSpace is free software: you can redistribute it and/or modify
% it under the terms of the GNU General Public License as published by
% the Free Software Foundation, either version 3 of the License, or
% (at your option) any later version.
% 
% AdaptiveResolutionOrientationSpace is distributed in the hope that it will be useful,
% but WITHOUT ANY WARRANTY; without even the implied warranty of
% MERCHANTABILITY or FITNESS FOR A PARTICULAR PURPOSE.  See the
% GNU General Public License for more details.
% 
% You should have received a copy of the GNU General Public License
% along with AdaptiveResolutionOrientationSpace.  If not, see <http://www.gnu.org/licenses/>.
% 
% 


%======================
% TEST INPUT
%======================

% defaults
verbose = 1;

% goodDataIdx is used in case data contains nans
goodDataIdx = [];

% check for axesHandle
if ishandle(varargin{1})
    axesH = varargin{1};
    varargin(1) = [];
    verbose = 1;
else
    axesH = [];
end

switch length(varargin) - isscalar(varargin{end})
    case 1 % data
        doHistogram = 1;
        data = varargin{1};
        data = data(:);

        if length(varargin) == 2
            verbose = varargin{2};
        end

        % make sure that there are no nans or infs
        goodDataIdx = find(isfinite(data));
        if ~isempty(goodDataIdx)
            data = data(goodDataIdx);
        else
            error('all data is NaN or Inf or empty!')
        end

    case 2 % conts,bins
        doHistogram = 0;
        counts = varargin{1};
        bins = varargin{2};
        counts = counts(:);
        bins = bins(:);
        nBins = length(bins);
        sp = [];

        if nargin == 3
            verbose = varargin{3};
        end

    otherwise
        error('wrong number of input arguments or non-scalar ''verbose''')
end

%===========================


%==========================
% BUILD HISTOGRAM
%==========================
if doHistogram
    %     [counts,bins] = optimalHistogram(data);
    %     counts = counts(:);
    %     bins = bins(:);
    % instead of a histogram, make a cumulative histogram and flip it along the
    % horizontal. The number-count becomes counts, the actual value becomes the
    % bin position
    % we still need a histogram to find the maximum!

    %     [bins,sortIdx] = sort(data(:));
    %     [counts] = optimalHistogram(bins);
    %     [maxVal, maxIdx] = max(counts);
    %     bins = bins(maxIdx:end);
    %     nBins = length(bins);
    %     counts = [nBins:-1:1]';

    % the continuous histogram can potentially place the maximum elsewhere
    % (where it might belong, but that's not the point)
    % Do discrete histogram first, find max-bin, and then look for the
    % maximum in the continuous histogram only up to the end of the
    % maxBin+0.5
    [counts, bins] = optimalHistogram(data);
    % if there is only one bin, we have a problem. Therefore, force at
    % least 3 bins.
    if length(bins)<3
        [counts,bins] = hist(data,3);
    end
    % the last bin cannot become the maximum
    [dummy,maxBinIdx] = max(counts(1:end-1));
    maxBinValue = bins(maxBinIdx + 1);


    % continuous histogram
    [counts,bins,sp] = optimalHistogram(data,'smooth');
    nBins = length(bins);

    % find maximum between bin 1 and the one with maxBinValue
    [maxVal, maxIdx] = max(counts(bins < maxBinValue));

else
    [maxVal, maxIdx] = max(counts);
end
%=========================


%=========================
% CUTOFF
%=========================

% for the line, we need the position of the maximum count and the position
% of the first empty bin following the last nonempty bin

% normalize counts and bins to go from 0 to 1.
countsN = counts / maxVal;
binsN = bins / max(abs(bins));

nBins = nBins - maxIdx + 1;
pointMax = [binsN(maxIdx), 1]; % maxVal = 1 now
pointEnd = [binsN(end) + median(diff(binsN)), countsN(end)];


% calculate perpendicular distance to the line
vector = pointEnd-pointMax;
[dummy,vector] = normList(vector);
distanceVector = perpVector(...
    repmat(pointMax,[nBins,1]),repmat(vector,[nBins,1]),...
    [binsN(maxIdx:end),countsN(maxIdx:end)]);
distance = normList(distanceVector);

% put distance = 0 wherever the contHistogram is above the line
distance(all(distanceVector>0,2)) = 0;

% find maximum
[maxDistance, maxDistanceIdx] = max(distance);

%========================


%========================
% ASSIGN OUTPUT
%========================

cutoffIndex = maxDistanceIdx + maxIdx - 1;
cutoffValue = bins(cutoffIndex);

if doHistogram

    % for plotting: stuff stolen from plotyy
    if verbose
        
        % check for axes (and figure!) to plot into
        if ~isempty(axesH)
            optimalHistogram(axesH,data)
            fig = get(axesH,'Parent');
        else
            fig =figure;
            set(fig,'NextPlot','add')
            optimalHistogram(data)
            axesH = gca;
        end
        ax(1) = axesH;
        ax(2) = axes('Units',get(ax(1),'Units'), ...
            'Position',get(ax(1),'Position'),...
            'ActivePositionProperty',get(ax(1),'ActivePositionProperty'),...
            'OuterPosition',get(ax(1),'OuterPosition'),...
            'Position',get(ax(1),'Position'),...
            'Parent',fig);
        plot(ax(2),bins,counts,'r',[cutoffValue,cutoffValue],[0,max(counts)],'-.r')
        set(ax(2),'YAxisLocation','right','Color','none','YColor','r')
        set(ax,'Box','off');
        y2Limits = ylim(ax(2));
        y2Limits(1) = 0;
        ylim(ax(2),y2Limits)
    end

else
    if verbose
        
        % check for axes to plot into
        if ~isempty(axesH)
            bar(axesH,bins,counts),hold on,
            plot(axesH,[cutoffValue,cutoffValue],[0,maxVal],'r')
        else
            figure,
            bar(bins,counts),hold on,
            plot([cutoffValue,cutoffValue],[0,maxVal],'r')
        end
    end

end

if ~isempty(goodDataIdx)
    cutoffIndex = goodDataIdx(cutoffIndex);
end
```
