## Supplementary material for "Adaptive-Resolution Multi-Orientation Analysis of Complex Filamentous Network Images": MATLAB code from Github: getMultiplicity.m

```
% [r, udata, sdata] = getMultiplicity(data) returns the occurrences/Multiplicity of the elements of 'data'
%
% Inputs:
%         data : n-dimensional input array
%
% Outputs: 
%          rep : # of occurrences for each element of 'data'
%        udata : sorted 1-D array of unique values in 'data'
%        sdata : sorted 1-D array of values in 'data'
%
% Note: NaN/Inf elements in input data are ignored
%
% Copyright (C) 2019, Jaqaman Lab - UT Southwestern, Goldman Lab - Northwestern 
%
% This file is part of AdaptiveResolutionOrientationSpace.
% 
% AdaptiveResolutionOrientationSpace is free software: you can redistribute it and/or modify
% it under the terms of the GNU General Public License as published by
% the Free Software Foundation, either version 3 of the License, or
% (at your option) any later version.
% 
% AdaptiveResolutionOrientationSpace is distributed in the hope that it will be useful,
% but WITHOUT ANY WARRANTY; without even the implied warranty of
% MERCHANTABILITY or FITNESS FOR A PARTICULAR PURPOSE.  See the
% GNU General Public License for more details.
% 
% You should have received a copy of the GNU General Public License
% along with AdaptiveResolutionOrientationSpace.  If not, see <http://www.gnu.org/licenses/>.
% 
% 

% Francois Aguet, 03/02/2012 (modified on 10/29/2012)
% Mark Kittisopikul, calculate udata only when requested, 09/11/2014

function [rep, udata, sdata] = getMultiplicity(data)

% fail quickly if data is empty
if(isempty(data))
    rep = [];
    udata = data;
    sdata = data;
    return;
end

if(~isinteger(data))
    data = data(isfinite(data));
end
% sort
sdata = sort(data(:));
% store as row vector
sdata = sdata(:)';

% find where the numbers change in the sorted array
isDiff = [diff(sdata)~=0 1];
idx = find(isDiff);

if(nargout > 1)
    udata = sdata(idx);
end

% count occurrences
rep = diff([0 idx]);
```
