## Supplementary material for "Adaptive-Resolution Multi-Orientation Analysis of Complex Filamentous Network Images": MATLAB code from Github: halleyft.m

```
function [ refined, refined_derivs, refined_iter, numIter ] = halleyft( v, guess, freq, deriv, TOL, maxIter, avoidNaN, uniq, varargin )
%halleyft Does Halley iteration to refine roots of Fourier series
%
% INPUT
% v - known values of Fourier series at regular intervals or it's Fourier
% transform. Each Fourier series is encoded in the first dimension as a
% column
%
% guess - guess for the root to find, see interpft_extrema. Multiple
% guesses for the same Fourier series are encoded in the first dimension as
% a column. All other dimensions must match v.
% NOTE: guess will be sorted with NaNs at the bottom if avoidNaN is true
%
% freq - logical. If true, then v is the Fourier transform of the values
%
% deriv - solve for zero of the derivative indicated
%         (optional, default = 0);
%
% TOL - tolerance for absolute distance from 0, default: 1e-12
%
% maxIter - maximum number of iterations, default: 10
%
% avoidNaN - For non-vector input, avoid processing NaN initial guesses.
%            The matrix should be sorted for this to work.
%
% uniq - Return unique zeros
%
% OUTPUT
% refined - refined zeros
% refined_derivs - derivatives of zeros
% refined_iter - iterations done per guess
% numIter - total number of iterations done
%
% Copyright (C) 2019, Jaqaman Lab - UT Southwestern, Goldman Lab - Northwestern 
%
% This file is part of AdaptiveResolutionOrientationSpace.
% 
% AdaptiveResolutionOrientationSpace is free software: you can redistribute it and/or modify
% it under the terms of the GNU General Public License as published by
% the Free Software Foundation, either version 3 of the License, or
% (at your option) any later version.
% 
% AdaptiveResolutionOrientationSpace is distributed in the hope that it will be useful,
% but WITHOUT ANY WARRANTY; without even the implied warranty of
% MERCHANTABILITY or FITNESS FOR A PARTICULAR PURPOSE.  See the
% GNU General Public License for more details.
% 
% You should have received a copy of the GNU General Public License
% along with AdaptiveResolutionOrientationSpace.  If not, see <http://www.gnu.org/licenses/>.
% 
% 

if(nargin < 3)
    freq = false;
end
if(ischar(freq))
    ip = inputParser;
    ip.addParameter('freq',false);
    ip.addParameter('deriv',0);
    ip.addParameter('TOL',1e-12);
    ip.addParameter('maxIter',10);
    ip.addParameter('avoidNaN',~isvector(guess));
    ip.addParameter('uniq',false);
    argsIn = {freq};
    if(nargin > 3)
        argsIn = [argsIn deriv];
    end
    if(nargin > 4)
        argsIn = [argsIn TOL];
    end
    if(nargin > 5)
        argsIn = [argsIn maxIter];
    end
    if(nargin > 6)
        argsIn = [argsIn avoidNaN];
    end
    if(nargin > 7)
        argsIn = [argsIn uniq];
    end
    argsIn = [argsIn varargin];
    ip.parse(argsIn{:});
    freq = ip.Results.freq;
    deriv = ip.Results.deriv;
    TOL = ip.Results.TOL;
    maxIter = ip.Results.maxIter;
    avoidNaN = ip.Results.avoidNaN;
    uniq = ip.Results.uniq;
else
    if(nargin < 4)
        deriv = 0;
    end
    if(nargin < 5)
        TOL = 1e-12;
    end
    if(nargin < 6)
        % If more than 10, probably should use interpft_extrema
        maxIter = 10;
    end
    if(nargin < 7)
        avoidNaN = ~isvector(guess);
    end
    if(nargin < 8)
        uniq = false;
    end
end

derivs = [0 1 2] + deriv;

    %% From interpft1_derivatives
    if(~freq)
        v_hat = fft(v(:,:));
    else
        v_hat = v(:,:);
    end
    
    if(avoidNaN)
        % NaN guesses slow down the algorithm
        % Group columns by number of non-nan guesses and recurse
        % Depends on the guess being sorted such that NaNs are all in the
        % bottom iterations
        guess_sz = size(guess);
        guess = guess(:,:);
        is_nan_guess = isnan(guess);
        resort = false;
        if(~issorted(is_nan_guess,1))
            [guess,sortIdx] = sort(guess,1);
            sortIdx = sub2ind(size(guess),sortIdx,repmat(1:size(guess,2),size(guess,1),1));
            resort = true;
        end
        guess_count = size(guess,1)-sum(is_nan_guess,1);
        refined = NaN(size(guess));
        if(nargout > 1)
            refined_derivs = refined(:,:,[1 1 1]);
            for i=1:size(guess,1)
                s = guess_count == i;
                if(sum(s) > 0)
                    [refined(1:i,s),refined_derivs(1:i,s,:)] = halleyft(v_hat(:,s), guess(1:i,s), true, deriv, TOL, maxIter, false, uniq);
                end
            end
            if(resort)
                for i=1:3
                    temp = refined_derivs(:,:,i);
                    temp(sortIdx) = refined_derivs(:,:,i);
                    refined_derivs(:,:,i) = temp;
                end
            end
            refined_derivs = reshape(refined_derivs,[guess_sz 3]);
        else
            for i=1:size(guess,1)
                s = guess_count == i;
                if(sum(s) > 0)
                    refined(1:i,s) = halleyft(v_hat(:,s), guess(1:i,s), true, deriv, TOL, maxIter, false, uniq);
                end
            end
        end
        if(resort)
            refined(sortIdx) = refined;
        end
        refined = reshape(refined,guess_sz);
        return;
    end
    
    K = floor(size(v,1)/2);
    freqM = ifftshift(-K:K).'*1i;
    
%     derivDim = ndims(v)+1;
    derivDim = 3;
    freqMs = arrayfun(@(x) freqM.^x,derivs,'UniformOutput',false);
    freqMs = cat(derivDim,freqMs{:});
    
    v_hat = bsxfun(@times,v_hat,freqMs);  
    
    xqrep = ones(1,derivDim);
    xqrep(derivDim) = length(derivs);
    

    %% Perform iteration
args = {};
if(isa(guess,'gpuArray'))
    args = { 'gpuArray'};
end

guess_sz = size(guess);
guess = guess(:,:);
if(nargout > 1)
%     refined_derivs = NaN([size(guess) 3]);
    % Calculate derivatives at least once
    refined_derivs = interpft1([0 2*pi],v_hat,repmat(guess,xqrep),'horner_freq');
end

% Indicates that a column still needs further iterations
columnNotDone = true(1,size(guess,2),args{:});
% Tracks whether a new guess moves the derivative closer to zero
new_guess_is_better = true(size(guess),args{:});

% Calculate initial value
guess_vals = interpft1([0 2*pi],v_hat,repmat(guess,xqrep),'horner_freq');
% do while
% while(~numIter || any(exceeds_tol) && any(new_guess_is_better(:)))
numIter = 0;
if(nargout > 2)
    refined_iter = zeros(size(guess),'uint8');
end
while(~numIter || any(columnNotDone))
%     disp('hi');
    vv = guess_vals(:,:,1);
    vd = guess_vals(:,:,2);
    vdd = guess_vals(:,:,3);
    new_guess = guess(:,columnNotDone) - 2*vv.*vd./(2*vd.^2-vv.*vdd);
    new_guess = wraparoundN(new_guess,0,2*pi);
    new_guess_vals = interpft1([0 2*pi],v_hat(:,columnNotDone,:),repmat(new_guess,xqrep),'horner_freq');
    new_guess_is_better_now = abs(new_guess_vals(:,:,1)) < abs(guess_vals(:,:,1));
    new_guess_is_better(:,columnNotDone) = new_guess_is_better_now;
    guess(new_guess_is_better) = new_guess(new_guess_is_better_now);
    if(nargout > 1)
        refined_derivs(new_guess_is_better(:,:,[1 1 1])) = new_guess_vals(new_guess_is_better_now(:,:,[1 1 1]));
    end
%     columnNotDone(columnNotDone) = any(new_guess_is_better(:,columnNotDone),1);

    % Establish new derivative value
    zero_vals = min(abs(guess_vals(:,:,1)),abs(new_guess_vals(:,:,1)));
    exceeds_tol = zero_vals > TOL;
       
    numIter = numIter + 1;
    if(numIter > maxIter)
        % Maximum number of iterations exceeded
        temp = guess(:,columnNotDone);
        % Set values whose derivative exceed tolerance to NaN
        temp( exceeds_tol ) = NaN;
        guess(:,columnNotDone) = temp;
        if(nargout > 1)
            temp = refined_derivs(:,columnNotDone,:);
            temp( exceeds_tol(:,:,[1 1 1]) ) = NaN;
            refined_derivs(:,columnNotDone,:) = temp;
        end
        if(nargout > 2)
            refined_iter(:,columnNotDone) = Inf;
        end
        % Issue a warning, use warning('off','halleyft:maxIter') to
        % deactivate
        warning('halleyft:maxIter','halleyft: maximum iteration reached');
        break;
    else
        % If new guess is not better and the value still exceeds tolerance,
        % then set the guess to NaN
        temp = guess(:,columnNotDone);
        invalid = ~new_guess_is_better_now & exceeds_tol;
        temp( invalid ) = NaN;
        guess(:,columnNotDone) = temp;
        if(nargout > 1)
            temp = refined_derivs(:,columnNotDone,:);
            temp( invalid(:,:,[1 1 1]) ) = NaN;
            refined_derivs(:,columnNotDone,:) = temp;
        end
        if(nargout > 2)
            refined_iter(:,columnNotDone) = numIter;
        end

        % New guess will only be considered further if derivative exceeds
        % tolerance
        new_guess_is_better_now = new_guess_is_better_now & exceeds_tol;
        new_guess_is_better(:,columnNotDone) = new_guess_is_better_now;
        % Find columns that are not done in the set of not done columns
        newNotDone = any(new_guess_is_better_now,1);
        % Use new guess values in the next iteration
        guess_vals = new_guess_vals(:,newNotDone,:);
        columnNotDone(columnNotDone) = newNotDone;
    end
end

refined = reshape(guess,guess_sz);
if(nargout > 1)
    refined_derivs = reshape(refined_derivs,[guess_sz 3]);
end
if(nargout > 2)
    refined_iter = reshape(refined_iter,guess_sz);
end

if(uniq && size(refined,1) > 1)
    [refined_sorted,sort_idx] = sort(refined);
    refined_sorted_diff = diff(refined_sorted);
    refined_sorted_diff = [refined_sorted(1,:)+2*pi-max(refined_sorted); refined_sorted_diff];
    if(islogical(uniq))
        uniq = 1e-6;
    end
    refined_sorted(refined_sorted_diff < uniq) = NaN;
    
    o = ones(size(sort_idx));
    idx = cell(1,ndims(sort_idx));
    idx{1} = sort_idx;
    for nd=2:ndims(sort_idx)
        idx{nd} = shiftdim((1:size(sort_idx,nd)).',nd-1).*o;
    end
    sort_idx = sub2ind(size(sort_idx),idx{:});
    refined(sort_idx) = refined_sorted;
    if(nargout > 1)
        isnan_refined = isnan(refined);
        colons = {':'};
        colons = colons(ones(1,ndims(refined)));
        for i=1:3
            temp = refined_derivs(colons{:},i);
            temp(isnan_refined) = NaN;
            refined_derivs(colons{:},i) = temp;
        end
        
    end
    if(nargout > 2)
        refined_iter(isnan_refined) = NaN;
    end
end

end
```
