## Supplementary material for "Adaptive-Resolution Multi-Orientation Analysis of Complex Filamentous Network Images": MATLAB code from Github: horner_vec_real_freq_simpler.m

```
function vq = horner_vec_real_freq_simpler(v_h,xq)
    % v represents the coefficients of the polynomial
    %   D x N
    %   D = degree of the polynomial - 1
    %   N = number of polynomials
    % xq represents the query points
    %   Q x N
    %   Q = number of query points per polynomial
    %   N = number of polynomials
    % vq will be a Q x N matrix of the value of each polynomial
    %    evaluated at Q query points
    %
    % SIMPLIFICATIONS
    % v_h is a 2-D Matrix   D x N
    % xq is a 2-D Matrix    Q x N
%
% Copyright (C) 2019, Jaqaman Lab - UT Southwestern, Goldman Lab - Northwestern 
%
% This file is part of AdaptiveResolutionOrientationSpace.
% 
% AdaptiveResolutionOrientationSpace is free software: you can redistribute it and/or modify
% it under the terms of the GNU General Public License as published by
% the Free Software Foundation, either version 3 of the License, or
% (at your option) any later version.
% 
% AdaptiveResolutionOrientationSpace is distributed in the hope that it will be useful,
% but WITHOUT ANY WARRANTY; without even the implied warranty of
% MERCHANTABILITY or FITNESS FOR A PARTICULAR PURPOSE.  See the
% GNU General Public License for more details.
% 
% You should have received a copy of the GNU General Public License
% along with AdaptiveResolutionOrientationSpace.  If not, see <http://www.gnu.org/licenses/>.
% 
% 
    
%     org_sz = size(xq);
%     v_h = v_h(:,:);
%     xq  = xq(:,:);

    nQ = size(xq,1);
    
    s = size(v_h);
    scale_factor = s(1);

    % Calculate fft and nyquist frequency
    nyquist = ceil((s(1)+1)/2);
    
    % If there is an even number of fourier coefficients, split the nyquist frequency
    if(~rem(s(1),2))
        % even number of coefficients
        % split nyquist frequency
        v_h(nyquist,:) = real(v_h(nyquist,:))/2;
%        v_h = v_h([1:nyquist nyquist nyquist+1:end],:);
%        v_h = reshape(v_h,[s(1)+1 s(2:end)]);
    end
    
    % z is Q x N
    z = exp(1i*pi*xq);
    % vq starts as 1 x N
%     colon = {':'};
%     v_h_colon = colon(ones(ndims(v_h)-1,1));
       
    vq = v_h(nyquist,:);
    vq = repmat(vq,nQ,1);
    for j = nyquist-1:-1:2
        vq = z.*vq;
        vq = repmat(v_h(j,:),nQ,1)+vq;
%         vq = bsxfun(@times,z,vq);
%         vq = bsxfun(@plus,v_h(j,v_h_colon{:}),vq);
    end
       
    % Last multiplication
%     vq = bsxfun(@times,z,vq); % We only care about the real part
    vq = z.*vq;
    vq = real(vq);
%     vq = bsxfun(@times,real(z),real(vq))-bsxfun(@times,imag(z),imag(vq));
    % Add Constant Term and Scale
%     vq = bsxfun(@plus,v_h(1,v_h_colon{:}),vq*2);
    vq = repmat(v_h(1,:),nQ,1)+vq*2;
%     vq = real(vq); % We already selected the real part above
    vq = vq./scale_factor;
    
%     vq = reshape(vq,org_sz);
end
```
