## Supplementary material for "Adaptive-Resolution Multi-Orientation Analysis of Complex Filamentous Network Images": MATLAB code from Github: interpft1.m

```
function [vq] = interpft1(varargin)
% INTERPFT1 Evaluate 1D Fourier series at off-grid points, xq
%
% INPUT
% x - (optional) interval that the periodic function is defined on
%     default: [1 size(v,1)+1]
% v - known values at regular intervals corresponding to
%     linspace(x(1),x(2)-x(1),size(v,1))
% xq - abisssa to determine value of Fourier series at
%      size(v,ii) must equal size(xq,ii) for ii > 1 if legacy is false
%      OR xq is a column vector
% method - (optional) interpolation method
%          See interp1 for methods, also available:
%          horner - Use Horner's method from chebfun
%          horner_freq - Same as above, but v already has fft applied
%          mmt - matrix multiplication method
%          default: horner
% fineGridFactor - (optional) Number of grid points to expand to using interpft
%                  default: Depends on method
%                  6 for cubic, pchip, v5cubic
%                  3 for spline
%                  10 for everything else
% legacy - (optional) logical of whether to use legacy output behavior. See
%          below. If true, output should match interp1 exactly.
%          default: false
%
% horner and mmt are very accurate methods, but are slow
% interp1 methods are typically faster but less accurate
%
% OUTPUT
% vq - values of Fourier series at xq. Similar to interp1 conventions:
% Shape of v    Shape of xq    Size of vq
% ---------------------------------------
% Vector        Vector         size(xq)
% Vector        Matrix/N-D     size(xq)
% Matrix/N-D    Row Vector     size(xq) *
% Matrix/N-D    Column Vector  [length(xq) size(v,2), ... size(v,n)]
% Matrix/N-D    Matrix/N-D     size(xq) *
% * Behavior differs from interp1. Pass xq(:) for interp1 behavior and
% reshape or set legacy to true.
%
% Copyright (C) 2019, Jaqaman Lab - UT Southwestern, Goldman Lab - Northwestern 
%
% This file is part of AdaptiveResolutionOrientationSpace.
% 
% AdaptiveResolutionOrientationSpace is free software: you can redistribute it and/or modify
% it under the terms of the GNU General Public License as published by
% the Free Software Foundation, either version 3 of the License, or
% (at your option) any later version.
% 
% AdaptiveResolutionOrientationSpace is distributed in the hope that it will be useful,
% but WITHOUT ANY WARRANTY; without even the implied warranty of
% MERCHANTABILITY or FITNESS FOR A PARTICULAR PURPOSE.  See the
% GNU General Public License for more details.
% 
% You should have received a copy of the GNU General Public License
% along with AdaptiveResolutionOrientationSpace.  If not, see <http://www.gnu.org/licenses/>.
% 
% 

% Mark Kittisopikul
% July 20th, 2016

% TODO:
% 1. Extract mapping code, consider wraparound(N)

    [x,v,xq,method,fineGridFactor,legacy] = parseinputs(varargin{:});
    
    if(legacy)
        % Use interp1 output conventions
        if(isvector(v))
            vSz = [];
        else
            vSz = size(v);
        end
        if(isvector(xq) && ~isvector(v))
            outSz = length(xq);
        else
            outSz = size(xq);
        end
        outSz = [outSz vSz(2:end)];
        xq = xq(:);
    end
    
    % Map indices from [x(1) x(2)) to [0 1)
    period = x(2) - x(1);
    xq = xq - x(1);
    xq = mod(real(xq),period)+1i*imag(xq);
    xq = xq./period;

    done = true;
    
    switch(method)
%         case 'chebfun'
%             % trigtech.horner is aligned on the domain [0,2)
%             % The output is not normalized by the number of points
%             xq = xq*2;
%             try
%                 vq = trigtech.horner(xq(:),fftshift(fft(v),1),isreal(v))./size(v,1);
%             catch err
%                 if(strcmp(err.identifier,'MATLAB:undefinedVarOrFunction'))
%                     warning('interpft1:chebfun_missing', ...
%                             'trigtech.horner in chebfun is not available. Using mmt method instead');
%                     vq = interp1(x,v,xq,'mmt',fineGridFactor);
%                     return;
%                 else
%                     rethrow(err);
%                 end
%             end
%             v_sz = size(v);
%             xq_sz = size(xq);
%             if(xq_sz(2) == 1)
%                 xq_sz = xq_sz(1);
%             end
%             vq = reshape(vq,[xq_sz v_sz(2:end)]);
        case 'horner'
            % domain [0,2)
            xq = xq*2;
            if(isreal(xq))
                vq = horner_vec_real(v,xq);
            else
                vq = horner_vec_complex(v,xq);
            end
        case 'horner_freq'
            xq = xq*2;
            if(isreal(xq))
                org_sz = size(xq);
                v = v(:,:);
                xq = xq(:,:);
                vq = horner_vec_real_freq_simpler(v,xq);
%                 vq = horner_vec_real_freq_simpler_mex(v,xq);
                vq = reshape(vq,org_sz);
%                 vq = horner_vec_real_freq(v,xq);
            else
                vq = horner_vec_complex_freq(v,xq);
            end
        case 'horner_complex'
            xq = xq*2;
            vq = horner_vec_complex(v,xq);
        case 'horner_complex_freq'
            xq = xq*2;
            vq = horner_vec_complex_freq(v,xq);
        case 'mmt'
            % domain [0,2*pi)
            xq = xq*2*pi;
            vq = matrixMultiplicationTransform(v,xq);
        case 'mmt_freq'
            xq = xq*2*pi;
            vq = matrixMultiplicationTransformFreq(v,xq);
        otherwise
            done = false;
    end
    if(~done)
        % Use interp1 methods by expanding grid points using interpft
        fineGridFactor = parseFineGridFactor(fineGridFactor,method);
        vft3 = interpft(v,size(v,1)*fineGridFactor);
        vft3 = [vft3(end-2:end,:); vft3(:,:); vft3(1:4,:)];

        % Map indices from [0 1) to [4 size(v,1)*fineGridFactor+4)
        xq = xq.*(size(v,1)*fineGridFactor);
        xq = xq+4;
        if(legacy || iscolumn(xq))
            vq = interp1(vft3,xq,method);
        else
            % break xq into columns and apply to each corresponding column in v 
            vq = cellfun(@(ii,xq) interp1(vft3(:,ii),xq,method),num2cell(1:numel(xq)/size(xq,1)),num2cell(xq(:,:),1),'UniformOutput',false);
            vq = [vq{:}];
            vSz = size(v);
            vq = reshape(vq,[size(xq,1) vSz(2:end)]);
        end
    end
    
    if(legacy)
        vq = reshape(vq,outSz);
    end
end

function [x,v,xq,method,fineGridFactor,legacy] = parseinputs(varargin)
    % Optional arguments
    method = 'horner';
    fineGridFactor = [];
    x = [];
    legacy = false;
    if(islogical(varargin{end}) && isscalar(varargin{end}))
        legacy = varargin{end};
        varargin = varargin(1:end-1);
    end
    % Decide whether optional arguments are specified, based existence of x
    switch(length(varargin))
        case 6
            [x, v,xq,method,fineGridFactor,legacy] = varargin{:};
        case 5
            % All args specified
            % x v xq method fineGridFactor
            [x, v,xq,method,fineGridFactor] = varargin{:};
        case 4
            if(ischar(varargin{4}))
                % x v xq method
                [x,v,xq,method] = varargin{:};
            else
                % v xq method fineGridFactor
                [v,xq,method,fineGridFactor] = varargin{:};
            end
        case 3
            if(ischar(varargin{3}))
                % v xq method
                [v, xq, method] = varargin{:};
            else
                % x v xq
                [x, v, xq] = varargin{:};
            end
        case 2
            % v xq

            [v, xq]  = varargin{:};
        otherwise
            error('interpft1:nargin','Incorrect number of inputs');
    end
    % Row values are transposed automatically like interp1
    if(isrow(v))
        v = v(:);
    end
    % Query points are transposed automatically like interp1
    if(isrow(xq))
        if(iscolumn(v) || legacy)
            xq = xq(:);
        end
    end
    % Default x specifying periodic boundary f(x) == f(size(v,1)+x)
    switch(numel(x))
        case 2
        case 0
            x = [1 size(v,1)+1];
        otherwise
            error('interpft1:IncorrectX', ...
            'x must either be empty or have 2 elements.');
    end
    if(~legacy)
        % Not in legacy mode
        xq_sz = size(xq);
        v_sz = size(v);
        same_sz = xq_sz(2:end) == v_sz(2:end);
        % Append ones to right side of v_sz so that
        %    length(xq_sz) == length(v_sz)
        v_sz(length(v_sz)+1:length(xq_sz)) = 1;
        % xq should either be a column such that all points are queried against all trig polynomials
        %    outSz = [length(xq) v_sz(2:end)]
        % OR be of the same size as v for every dimension except the first.
        %    outSz = size(xq)
 
        assert(iscolumn(xq) || all(xq_sz(2:end) == v_sz(2:end)) || all(xq_sz([false ~same_sz]) == 1), ...
                'interpft1:InputDimensions', ...
                'xq should either be a column or have similar dimensions as v if not in legacy mode');
    % else
        % Legacy mode
        % outSz = size(xq) % if v is a vector
        % outSz = [length(xq) v_sz(2:end)] % if xq is a vector, and v is not
        % outSz = [xq_sz v_sz(2:end)] % if xq and v both are not vectors
    end
end
function fineGridFactor = parseFineGridFactor(fineGridFactor,method)
    % Courser methods should use a finer grid if none is specified
    if(isempty(fineGridFactor))
        switch(method)
%             case 'horner'
%                 fineGridFactor = NaN;
%             case 'mmt'
%                 fineGridFactor = NaN;
%             case 'linear'
%                 fineGridFactor = 10;
%             case 'nearest'
%                 fineGridFactor = 10;
%             case 'next'
%                 fineGridFactor = 10;
%             case 'previous'
%                 fineGridFactor = 10;
            case 'pchip'
                fineGridFactor = 6;
            case 'cubic'
                fineGridFactor = 6;
            case 'v5cubic'
                fineGridFactor = 6;
            case 'spline'
                fineGridFactor = 3;
            otherwise
                fineGridFactor = 10;
        end
    end
end

function vq = matrixMultiplicationTransform(v,xq)
    vq = matrixMultiplicationTransformFreq(v_h,xq);
end
function vq = matrixMultiplicationTransformFreq(v_h,xq)
%matrixMultiplicationTransform
%
% Adapted from interpft_extrema
    s = size(v_h);
    scale_factor = s(1);

    % Calculate fft and nyquist frequency
    nyquist = ceil((s(1)+1)/2);

    % If there is an even number of fourier coefficients, split the nyquist frequency
    if(~rem(s(1),2))
        % even number of coefficients
        % split nyquist frequency
        v_h(nyquist,:) = v_h(nyquist,:)/2;
        v_h = v_h([1:nyquist nyquist nyquist+1:end],:);
        v_h = reshape(v_h,[s(1)+1 s(2:end)]);
    end
    % Wave number, unnormalized by number of points
    freq = [0:nyquist-1 -nyquist+1:1:-1]';
    
    % calculate angles multiplied by wave number
    theta = bsxfun(@times,xq,shiftdim(freq,-ndims(xq)));
    % waves
    waves = exp(1i*theta);


    % evaluate each wave by fourier coeffient
    % theta and waves have one more dimension than xq, representing
    % frequency
    ndims_waves = ndims(waves); % ndims(xq) + 1 ?
    % permute v_h such that it is a 1 by (array dim of fourier series) by
    %                               length of fourier series
    dim_permute = [ndims_waves 2:ndims_waves-1 1];
    
    % sum across waves weighted by Fourier coefficients
    % normalize by the the number of Fourier coefficients
    vq = sum(real(bsxfun(@times,waves,permute(v_h,dim_permute))),ndims_waves)/scale_factor;
end
function vq = horner_vec_real(v,xq)
    vq = horner_vec_real_freq(fft(v),xq);
end
function vq = horner_vec_complex(v,xq)
    vq = horner_vec_complex_freq(fft(v),xq);
end
function vq = horner_vec_real_freq(v_h,xq)
    % v represents the coefficients of the polynomial
    %   D x N
    %   D = degree of the polynomial - 1
    %   N = number of polynomials
    % xq represents the query points
    %   Q x N
    %   Q = number of query points per polynomial
    %   N = number of polynomials
    % vq will be a Q x N matrix of the value of each polynomial
    %    evaluated at Q query points
    s = size(v_h);
    scale_factor = s(1);

    % Calculate fft and nyquist frequency
    nyquist = ceil((s(1)+1)/2);
    
    % If there is an even number of fourier coefficients, split the nyquist frequency
    if(~rem(s(1),2))
        % even number of coefficients
        % split nyquist frequency
        v_h(nyquist,:) = real(v_h(nyquist,:))/2;
%        v_h = v_h([1:nyquist nyquist nyquist+1:end],:);
%        v_h = reshape(v_h,[s(1)+1 s(2:end)]);
    end
    
    % z is Q x N
    z = exp(1i*pi*xq);
    % vq starts as 1 x N
    colon = {':'};
    v_h_colon = colon(ones(ndims(v_h)-1,1));
       
    vq = v_h(nyquist,v_h_colon{:});
    for j = nyquist-1:-1:2
        vq = bsxfun(@times,z,vq);
        vq = bsxfun(@plus,v_h(j,v_h_colon{:}),vq);
    end
       
    % Last multiplication
    vq = bsxfun(@times,z,vq); % We only care about the real part
    vq = real(vq);
%     vq = bsxfun(@times,real(z),real(vq))-bsxfun(@times,imag(z),imag(vq));
    % Add Constant Term and Scale
    vq = bsxfun(@plus,v_h(1,v_h_colon{:}),vq*2);
%     vq = real(vq); % We already selected the real part above
    vq = vq./scale_factor;
end
function vq = horner_vec_complex_freq(v_h,xq)
    % v represents the coefficients of the polynomial
    %   D x N
    %   D = degree of the polynomial - 1
    %   N = number of polynomials
    % xq represents the query points
    %   Q x N
    %   Q = number of query points per polynomial
    %   N = number of polynomials
    % vq will be a Q x N matrix of the value of each polynomial
    %    evaluated at Q query points
    s = size(v_h);
    scale_factor = s(1);

    % Calculate fft and nyquist frequency
    nyquist = ceil((s(1)+1)/2);
    
    % If there is an even number of fourier coefficients, split the nyquist frequency
    if(~rem(s(1),2))
        % even number of coefficients
        % split nyquist frequency
        v_h(nyquist,:) = real(v_h(nyquist,:))/2;
%        v_h = v_h([1:nyquist nyquist nyquist+1:end],:);
%        v_h = reshape(v_h,[s(1)+1 s(2:end)]);
    end
    
    % z is Q x N
    z = exp(1i*pi*xq);
    % vq starts as 1 x N
    colon = {':'};
    v_h_colon = colon(ones(ndims(v_h)-1,1));
       
    vq = v_h(nyquist,v_h_colon{:});
    for j = [nyquist-1:-1:1 s(1):-1:nyquist+2]
        vq = bsxfun(@times,z,vq);
        vq = bsxfun(@plus,v_h(j,v_h_colon{:}),vq);
    end
       
    % Last multiplication
    vq = bsxfun(@times,z,vq); % We only care about the real part
%     vq = real(vq);
%     vq = bsxfun(@times,real(z),real(vq))-bsxfun(@times,imag(z),imag(vq));
    % Add Constant Term and Scale
    vq = bsxfun(@plus,v_h(nyquist+1,v_h_colon{:}),vq);
%     vq = real(vq); % We already selected the real part above
    vq = vq./scale_factor;
end
```
