## Supplementary material for "Adaptive-Resolution Multi-Orientation Analysis of Complex Filamentous Network Images": MATLAB code from Github: interpft1_derivatives.m

```
function [ vq ] = interpft1_derivatives( v, xq, derivs , period, freq, method)
%interpft1_derivatives Interpolate the derivatives of a Fourier series
%
% Copyright (C) 2019, Jaqaman Lab - UT Southwestern, Goldman Lab - Northwestern 
%
% This file is part of AdaptiveResolutionOrientationSpace.
% 
% AdaptiveResolutionOrientationSpace is free software: you can redistribute it and/or modify
% it under the terms of the GNU General Public License as published by
% the Free Software Foundation, either version 3 of the License, or
% (at your option) any later version.
% 
% AdaptiveResolutionOrientationSpace is distributed in the hope that it will be useful,
% but WITHOUT ANY WARRANTY; without even the implied warranty of
% MERCHANTABILITY or FITNESS FOR A PARTICULAR PURPOSE.  See the
% GNU General Public License for more details.
% 
% You should have received a copy of the GNU General Public License
% along with AdaptiveResolutionOrientationSpace.  If not, see <http://www.gnu.org/licenses/>.
% 
% 
if(nargin < 4)
    period = 2*pi;
    periodFactor = 1;
else
    periodFactor = 2*pi/period;
end
if(nargin < 5)
    freq = false;
end
if(nargin < 6)
    method = 'horner_freq';
end

    K = floor(size(v,1)/2);
    freqM = ifftshift(-K:K).'*1i;
    if(~freq)
        v_hat = fft(v);
    else
        v_hat = v;
    end
    derivDim = ndims(v)+1;
    derivs = shiftdim(derivs(:),-derivDim+1);
%     tic;
    freqMs = bsxfun(@power,freqM,derivs);
%     toc;
%     tic;
%         derivs_diff = diff(derivs);
%         freqMsz = ones(1,derivDim);
%         freqMsz(1) = length(freqM);
%         freqMsz(derivDim) = length(derivs);
%         freqMs = zeros(freqMsz);
%         colons = {':'};
%         colons = colons(ones(1,derivDim-1));
%         freqMs(colons{:},1) = freqM.^derivs(1);
%         for d=2:length(derivs)
%             freqMs(colons{:},d) = freqMs(colons{:},d-1).*freqM.^derivs_diff(d-1);
%         end
%     toc;
%     tic;
%     freqMs = arrayfun(@(x) freqM.^x,derivs,'UniformOutput',false);
%     freqMs = cat(derivDim,freqMs{:});
%     toc
    
    v_hat = bsxfun(@times,v_hat,freqMs);
    
    xqrep = ones(1,derivDim);
    xqrep(derivDim) = length(derivs);
    xq = repmat(xq,xqrep);
    
    vq = interpft1([0 period],v_hat, xq, method);
    if(periodFactor ~= 1)
        vq = bsxfun(@times,vq,shiftdim(periodFactor.^(derivs),-derivDim+2));
    end
    

end
```
