## Supplementary material for "Adaptive-Resolution Multi-Orientation Analysis of Complex Filamentous Network Images": MATLAB code from Github: interpft_extrema.m

```
function [maxima,minima,maxima_value,minima_value,other,other_value] = interpft_extrema(x,dim,sorted,TOL,dofft)
% interpft_extrema finds the extrema of the function when interpolated by fourier transform
% This is the equivalent of doing sinc interpolation.
%
% INPUT
% x - regularly sampled values to be interpolated.
%     Values are considered to be sampled at (0:length(x)-1)*2*pi/length(x)
% dim - dimension along which to find maxima
% sorted - logical value about whether to sort the maxima by value
% TOL - tolerance to determine if log(abs(root)) is zero to determine if
%       the root of the derivative is real.
%       If tolerance is negative, then use 10*abs(log(abs(root))) as the
%       tolerance only if no roots are found with tolerance at -TOL.
% dofft - logical. If true, transforms dim with fft. Default: true
%
% OUTPUT
% maxima - angle between 0 and 2*pi indicating location of local maxima
% minima - angle between 0 and 2*pi indicating location of local minima
% maxima_value - value of interpolated function at maxima
% minima_value - value of interpolated function at minima
% other - angle between 0 and 2*pi where second derivative is zero
% other_value - value of interpolated function at other
%
% If no output is requested, then the maxima and minima will be plotted.
% Maxima are indicated by a red vertical line and red circle.
% Minima are indicated by a green vertical line and a green circle
%
% 1D Example
%
% r = rand(7,1);
% figure;
% plot((0:length(r)-1)/length(r)*2*pi,r,'ko');
% hold on;
% plot((0:199)/200*2*pi,interpft(r,200),'k');
% interpft_extrema(r);
% hold off;
%
% 2D Example
% r = rand(11,3);
% figure;
% plot((0:size(r,1)-1)/size(r,1)*2*pi,r,'ko');
% hold on;
% plot((0:199)/200*2*pi,interpft(r,200),'k');
% interpft_extrema(r);
% hold off;
% 
% This function works by calculating a fourier series of degree length(x)
% that fits through the input points. Then the fourier series is then considered
% as a trigonometric polynomial and the roots are solved by evaluating the eigenvalues
% of a companion matrix via the builtin function roots.
%
% See also interpft, roots
%
% Author: Mark Kittisopikul, May 2016
%
% Copyright (C) 2019, Jaqaman Lab - UT Southwestern, Goldman Lab - Northwestern 
%
% This file is part of AdaptiveResolutionOrientationSpace.
% 
% AdaptiveResolutionOrientationSpace is free software: you can redistribute it and/or modify
% it under the terms of the GNU General Public License as published by
% the Free Software Foundation, either version 3 of the License, or
% (at your option) any later version.
% 
% AdaptiveResolutionOrientationSpace is distributed in the hope that it will be useful,
% but WITHOUT ANY WARRANTY; without even the implied warranty of
% MERCHANTABILITY or FITNESS FOR A PARTICULAR PURPOSE.  See the
% GNU General Public License for more details.
% 
% You should have received a copy of the GNU General Public License
% along with AdaptiveResolutionOrientationSpace.  If not, see <http://www.gnu.org/licenses/>.
% 
% 
    
%     original_size = size(x);
    if(nargin > 1 && ~isempty(dim))
        x = shiftdim(x,dim-1);
        unshift = ndims(x) - dim + 1;
    else
        if(isrow(x))
            % If the input is a row vector, transpose it without conjugation
            dim = 2;
            unshift = 1;
            x = x.';
        else
            dim = 1;
            unshift = 0;
        end
    end
    
    if(nargin < 3 || isempty(sorted))
        sorted = false;
    end
    if(nargin < 4 || isempty(TOL))
    	% Tolerance for log(abs(root)) to be near zero, in which case the root is real
        % Set negative so that tolerance adapts if no roots are found
        TOL = -eps(class(x))*1e3;
    end
    if(nargin < 5 || isempty(dofft))
        dofft = true;
    end


    output_size = size(x);
    output_size(1) = output_size(1) - 1;
    

    s = size(x);
    scale_factor = s(1);
    
    if(s(1) == 1)
        maxima = shiftdim(zeros(s),unshift);
        minima = maxima;
        other = maxima;
        maxima_value = shiftdim(x,unshift);
        minima_value = maxima_value;
        other_value = maxima_value;
        return;
    end

    % Calculate fft and nyquist frequency
    if(dofft)
        x_h = fft(x);
    else
        x_h = x;
    end
    nyquist = ceil((s(1)+1)/2);

    % If there is an even number of fourier coefficients, split the nyquist frequency
    if(~rem(s(1),2))
        % even number of coefficients
        % split nyquist frequency
        x_h(nyquist,:) = x_h(nyquist,:)/2;
        x_h = x_h([1:nyquist nyquist nyquist+1:end],:);
        x_h = reshape(x_h,[s(1)+1 s(2:end)]);
        output_size(1) = output_size(1) + 1;
    end
    % Wave number, unnormalized by number of points
    freq = [0:nyquist-1 -nyquist+1:1:-1]';

    % calculate derivatives, unscaled by 2*pi since we are only concerned with the signs of the derivatives
    dx_h = bsxfun(@times,x_h,freq * 1i);
    dx2_h = bsxfun(@times,x_h,-freq.^2);
    
    % use companion matrix approach
    dx_h = -fftshift(dx_h,1);
    dx_h = dx_h(:,:);
%     r = zeros(output_size,'like',dx_h);
    output_size1 = output_size(1);
    nProblems = prod(output_size(2:end)); 
    batchSize = min(1024,nProblems);
    nBatches = ceil(nProblems/batchSize);
    % Only use parallel workers if a pool already exists
    nWorkers = ~isempty(gcp('nocreate'))*nBatches;
    in = ones(1,nBatches)*batchSize;
    in(end) = in(end) + nProblems - sum(in);
    dx_h = mat2cell(dx_h,size(dx_h,1),in);
    r = cell(1,nBatches);
    parfor (i=1:nBatches, nWorkers)
        r{i} = zeros(output_size1,size(dx_h{i},2),'like',dx_h{i});
        for j = 1:size(dx_h{i},2);
            try
                % roots outputs only column vectors which may be shorter than
                % expected
                dx_h_roots = roots(dx_h{i}(:,j));
                dx_h_roots(end+1:output_size1) = 0;
                r{i}(:,j) = dx_h_roots;
            catch err
                switch(err.identifier)
                    case 'MATLAB:ROOTS:NonFiniteInput'
                        r{i}(:,j) = NaN(output_size1,1);
                end
            end
        end
    end
    r = [r{:}];
    r = reshape(r,output_size);
    % magnitude
    magnitude = abs(log(abs(r)));
    % keep only the real answers
    real_map = (magnitude <= abs(TOL));
    % If tolerance is negative and no roots are found, then use the
    % root that is closest to being real
    if(TOL < 0)
        no_roots = ~any(real_map);
        real_map(:,no_roots) = bsxfun(@le,magnitude(:,no_roots),min(magnitude(:,no_roots))*10);
    end
%     r(imaginary_map) = NaN;
%     real_map = ~imaginary_map;
%     clear imaginary_map
    
   
    % In the call to roots the coefficients were entered in reverse order (negative to positive)
    % rather than positive to negative. Therefore, take the negative of the angle..
    % angle will return angle between -pi and pi
    
    r = -angle(r(real_map));
    
    % Map angles to between 0 and 2 pi, moving negative values up
    % a period
    neg_extrema = r < 0;
    r(neg_extrema) = r(neg_extrema) + 2*pi;
    
    extrema = NaN(output_size);
    extrema(real_map) = r;
    
    % Use Horner's method to compute 2nd derivative value
    dx2 = interpft1([0 2*pi],dx2_h,extrema,'horner_freq');
    
    % dx2 could be equal to 0, meaning inconclusive type
    % local maxima is when dx == 0, dx2 < 0
    output_template = NaN(output_size);
    
    maxima = output_template;
    minima = output_template;
    
    maxima_map = dx2 < 0;
    minima_map = dx2 > 0;
    
    maxima(maxima_map) = extrema(maxima_map);
    % local minima is when dx == 0, dx2 > 0
    minima(minima_map) = extrema(minima_map);
    
    if(nargout > 2 || nargout == 0 || sorted)
	% calculate the value of the extrema if needed
        
        % Use Horner's method to compute value at extrema
        extrema_value = interpft1([0 2*pi],x_h,extrema,'horner_freq');
        % interpft1 will automatically scale, rescale if needed
        if(scale_factor ~= size(x_h,1))
            extrema_value = extrema_value/scale_factor*size(x_h,1);
        end

        maxima_value = output_template;
        maxima_value(maxima_map) = extrema_value(maxima_map);

        if(nargout > 3 || nargout > 1 && sorted || nargout == 0)
            minima_value = output_template;
            minima_value(minima_map) = extrema_value(minima_map);
        end
        
        if(sorted)

            
            maxima_value_inf = maxima_value;
            maxima_value_inf(isnan(maxima_value_inf)) = -Inf;
            [~,maxima,maxima_value] = sortMatrices(maxima_value_inf,maxima,maxima_value,'descend');
            clear maxima_value_inf;
            numMax = sum(maxima_map);
            numMax = max(numMax(:));
            colons = {':'};
            colons = colons(ones(1,ndims(maxima)-1));
            maxima = maxima(1:numMax,colons{:});
            maxima_value = maxima_value(1:numMax,colons{:});
            
            if(nargout > 1)
                minima_value_inf = minima_value;
                minima_value_inf(isnan(minima_value_inf)) = Inf;
                [~,minima,minima_value] = sortMatrices(minima_value_inf,minima,minima_value);
                clear minima_value_inf;
                numMin = sum(minima_map);
                numMin = max(numMin(:));
                minima = minima(1:numMin,colons{:});
                minima_value = minima_value(1:numMin,colons{:});
            end
                       
        end
        
%         maxima_value = reshape(maxima_value,output_size);
%         minima_value = reshape(minima_value,output_size);

        if(nargout > 2)
            maxima_value = shiftdim(maxima_value,unshift);
        end
        
        if(nargout > 3)
            minima_value = shiftdim(minima_value,unshift);
        end
        
        if(nargout > 4)
            % calculate roots which are not maxima or minima
            other = output_template;
            other_map = dx2 == 0;
            other(other_map) = extrema(other_map);
            
            other_value = output_template;
            other_value(other_map) = extrema_value(other_map);
            
            if(sorted)
                other_value_inf = other_value;
                other_value_inf(isnan(other_value_inf)) = -Inf;
                [~,other,other_value] = sortMatrices(other_value_inf,other,other_value,'descend');
                clear other_value_inf;
            end
            
%             other = reshape(other,output_size);
%             other_value = reshape(other_value,output_size);
            other = shiftdim(other,unshift);
            other_value = shiftdim(other_value,unshift);
        end
    end
    

    
    if(nargout == 0)
        % plot if no outputs requested
        if(sorted)
            real_maxima = maxima(:);
            real_maxima_value = maxima_value(:);
        else
            real_maxima = maxima(maxima_map);
            real_maxima_value = real(maxima_value(maxima_map));
        end
        if(~isempty(real_maxima))
        % Maxima will be green
            plot([real_maxima real_maxima]',ylim,'g');
            plot(real_maxima,real_maxima_value,'go');
        end
        if(sorted)
            real_minima = minima(:);
            real_minima_value = minima_value(:);
        else
            real_minima = minima(minima_map);
            real_minima_value = real(minima_value(minima_map));
        end
        if(~isempty(real_minima))
	    % Minima will be red
            plot([real_minima real_minima]',ylim,'r');
            plot(real_minima,real_minima_value,'ro');
        elseif(~isempty(real_maxima))
            warning('No extrema');
        end
    end
    
%     maxima = reshape(maxima,output_size);
%     minima = reshape(minima,output_size);
    maxima = shiftdim(maxima,unshift);
    if(nargout > 1)
        minima = shiftdim(minima,unshift);
    end
end
```
