## Supplementary material for "Adaptive-Resolution Multi-Orientation Analysis of Complex Filamentous Network Images": MATLAB code from Github: lap.m

```
function [x, y] = lap(cc, NONLINK_MARKER, extendedTesting, augmentCC, noLinkCost)
%LAP solves the linear assignment problem for a given cost matrix
%
% A linear assignment tries to establish links between points in two sets.
% One point in set A can only link to one point in set B and vice versa.
% The cost associated with the link from element i of A to element j of B
% is given by cc(i,j).
%
% SYNOPSIS [x, y] = lap(cc, NONLINK_MARKER, extendedTesting, augmentCC, noLinkCost)
%
% INPUT:  cc: cost matrix, which has to be square. Set cost(i,j) to the
%             value of NONLINK_MARKER, if the link is not allowed. cc can
%             be input as a sparse matrix (in which case there won't be any
%             NONLINK_MARKERs).
%             Elements of the cost matrix cannot be inf or nan!
%
%             For an unequal number of points (or, generally, if birth and
%             death is to be allowed) the cost matrix is formed as
%             a 2-by-2 catenation of four sub-matrices. If there are n
%             elements in set A and m elements in set B, sub-matrix (1,1)
%             is an n-by-m matrix with the cost for each link. The two
%             off-diagonal sub-matrices make non-links possible. They are
%             both diagonal square matrices (2,1: m-by-m; 1,2: n-by-n) with
%             the cost for not linking a point (e.g. determined by a
%             maximum search radius) in the diagonal and NONLINK_MARKER off
%             the diagonal. Sub-matrix (2,2) is a m-b-n matrix of any
%             not-NONLINK_MARKER value (suggested: one, except for sparse
%             matrix input)
%
% NONLINK_MARKER : value to indicate that two points cannot be linked.
%             Default: -1. NaN is not allowed here.
%
% extendedTesting (used to be optional, now input has no effect and
%                  extendedTesting will always be performed)
%                LAP will always make sure that:
%                  - There cannot be NaNs in cc
%                  - In every row and every column of cc there must be
%                    at least 1 element that is not a NONLINK_MARKER.
%
% augmentCC  (optional, [{0}/1]) If 1, the cost matrix will be augmented to
%            allow births and deaths.
% noLinkCost (optional, [{maximum cost + 1}]) Cost for linking a feature
%            to nothing.
%
%
% OUTPUT: x: The point A(i) links to B(x(i))
%         y: The point B(j) links to A(y(j))
%
%            Any x > m or y > n indicates that this point is not linked.
%
% EXAMPLE
%
% % Make a set of points p1 and slightly shift the points in p2
% p1 = rand(5,2)*10;
% p2 = p1+rand(5,2);
% % plot the data
% figure,plot(p1(:,1),p1(:,2),'.r',p2(:,1),p2(:,2),'.b')
% % invert the order of p2 (to make it a bit more interesting) and remove the
% % third point
% p2 = p2(end:-1:1,:);
% p2(3,:) = [];
% % calculate distance matrix between the point sets
% dm = distMat2(p1,p2);
% % remove all links that are longer than a threshold (mark with -1)
% dm(dm>5) = -1;
% % print distance matrix
% disp(dm)
% % link via LAP. nonlinkmarker=-1, augment the matrix and show
% [links12, links21] = lap(dm,-1,0,1)
% % links12 should now look something like
% % links12 =
% %
% %            4
% %            3
% %            7
% %            2
% %            1
% %            9
% %            5
% %            6
% %            8
% %
% % For links12, the first five entries are relevant (because p1 consists of
% % 5 points). The first point in p1 links to point #4 in set 2
% % (link12(1)==4). The third point in p1 links nowhere - there are only 4
% % points in links21, so an index above 4 means that a link to a virtual
% % point had to be made.
% % Links21 is analogous, except that only the first four entries count.
%
%
% lapjv is implemented through a MEX DLL.
%
% Author: Ge Yang, LCCB, TSRI, Dec. 10, 2004
% extended by jonas 6/05
%
% Copyright (C) 2019, Jaqaman Lab - UT Southwestern, Goldman Lab - Northwestern 
%
% This file is part of AdaptiveResolutionOrientationSpace.
% 
% AdaptiveResolutionOrientationSpace is free software: you can redistribute it and/or modify
% it under the terms of the GNU General Public License as published by
% the Free Software Foundation, either version 3 of the License, or
% (at your option) any later version.
% 
% AdaptiveResolutionOrientationSpace is distributed in the hope that it will be useful,
% but WITHOUT ANY WARRANTY; without even the implied warranty of
% MERCHANTABILITY or FITNESS FOR A PARTICULAR PURPOSE.  See the
% GNU General Public License for more details.
% 
% You should have received a copy of the GNU General Public License
% along with AdaptiveResolutionOrientationSpace.  If not, see <http://www.gnu.org/licenses/>.
% 
% 


% Basically, what this function does is to automatically extract the sparse matrix
% representation of cc.

% int *col = new int[size * NEIGHBOR_NUM_MAX + 1];
% int *first = new int[size + 2];
% int *x = new int[size + 1];
% int *y = new int[size + 1];
% double *u = new double[size + 1];
% double *v = new double[size + 1];
% double  *cc = new double[size * NEIGHBOR_NUM_MAX + 1];


%=====================
% TEST INPUT
%=====================

if (nargin == 1) || isempty(NONLINK_MARKER)
    NONLINK_MARKER = -1;
elseif isnan(NONLINK_MARKER)
    error('NONLINK_MARKER cannot be NaN!')
end

if nargin < 4 || isempty(augmentCC)
    augmentCC = 0;
end

% test size
scc = size(cc);
% check size only if no augmentation
if ~augmentCC

    if scc(1) ~= scc(2) || length(scc) > 2
        error('cost must be a 2D square matrix!')
    end

elseif length(scc) > 2

    error('cost must be a 2D matrix!')

else

    % if we're augmenting, sparse matrices are produced. This will be
    % problematic with cost 0
    if any(cc(:)==0) && ~issparse(cc)
        validCC = cc ~= NONLINK_MARKER;
        if any(any(cc(validCC))) < 0
            warning('LAP:negativeCosts',...
                'there are negative and zero costs. This could lead to errors')
        end
        cc(validCC) = cc(validCC) + 10;
    end

end

% extended testing has been made mandatory
%
% if nargin < 3 || isempty(extendedTesting)
%     extendedTesting = true;
% end

if nargin < 5 || isempty(noLinkCost)
    noLinkCost = max(max(cc)) + 1;
end

% if we have -1 as non-link-marker, and we get it everywhere, we get
% a segmentation fault.
if noLinkCost == NONLINK_MARKER
    noLinkCost = noLinkCost + 1;
    if noLinkCost == 0
        noLinkCost = 0.01;
    end
end


%=======================

%=================================
% AUGMENT COST MATRIX IF SELECTED
%=================================
if augmentCC

    % expand the m-by-n cost matrix to an (m+n)-by-(n+m) matrix, adding
    % diagonals with noLinkCost

    % in the lower right corner, we want the lowest cost of all - take
    % minimum cost, subtract 5 and make sure it's not 0.
    % NONLINK_MARKER is not a problem because we make the matrix sparse
    % before augmenting.
    %
    % not needed anymore since we use cc' as costLR
%     minCost = min(min(cc(cc~=NONLINK_MARKER)))-5;
%     if minCost == 0
%         minCost = -2;
%     end


    % check if sparse
    if issparse(cc)

        % mmDiag = spdiags(noLinkCost * ones(scc(1),1), 0, scc(1), scc(1));
        % nnDiag = spdiags(noLinkCost * ones(scc(2),1), 0, scc(2), scc(2));
        % nmMat  = sparse(ones(scc(2), scc(1)));
        % cc = [cc, mmDiag; nnDiag, nmMat];

        % use transposed costmatrix for LR - otherwise, nonLinkCost has no
        % effect
        %[rowIdx, colIdx] = find(cc);
        %costLR = sparse(colIdx,rowIdx,minCost*ones(length(colIdx),1),scc(2),scc(1));
        costLR = cc';

        cc = [cc, spdiags(noLinkCost * ones(scc(1),1), 0, scc(1), scc(1));...
            spdiags(noLinkCost * ones(scc(2),1), 0, scc(2), scc(2)),...
            costLR];

    else

        % mmDiag = diag(noLinkCost * ones(scc(1),1));
        % nnDiag = diag(noLinkCost * ones(scc(2),1));
        % nmMat  = sparse(ones(scc(2), scc(1)));
        % cc = [cc, mmDiag; nnDiag, nmMat];

        % use transposed costmatrix for LR - otherwise, nonLinkCost has no
        % effect
        %[rowIdx, colIdx] = find(cc ~= NONLINK_MARKER);
        %costLR = sparse(colIdx,rowIdx,minCost*ones(length(colIdx),1),scc(2),scc(1));

        % make cc sparse. Take NLM in cc into account!
        cc(cc==NONLINK_MARKER) = 0;
        cc=sparse(cc);
        costLR = cc';

        cc = [cc, diag(noLinkCost * ones(scc(1),1)); ...
            diag(noLinkCost * ones(scc(2),1)), ...
            costLR];

    end

    % remember that the size of the matrix has increased!
    scc = [sum(scc), sum(scc)];

    clear costLR colIdx rowIdx

end


%=============================
% CALCULATE SPARSE MATRIX
%=============================

% Calculate sparse representation of cost matrix with compactCC (all
% significant elements of cc), fst and kk. Read cc in rows, not cols!

% fst: for every first significant element of a row: index into compactCC
% kk : for every significant element: column

% do the work on the transposed cost matrix!
cc = cc';
% find the significant elements. If sparse input, find nonzero elements
if issparse(cc)
    [rowIdx, colIdx, val] = find(cc);
else
    [rowIdx, colIdx] = find(cc ~= NONLINK_MARKER);
    val = cc(cc ~= NONLINK_MARKER);
end


    % test that all cols and all rows are filled, and that there are no nans
    if issparse(cc)
        if (~all(sum(cc,1)) || ~all(sum(cc,2)))
            error('LAP:BadCostMatrix',...
                'Rows and columns of the cost matrix must allow at least one possible link!')
        end
    else
        if (~all(sum(cc~=NONLINK_MARKER,1)) || ~all(sum(cc~=NONLINK_MARKER,2)))
            error('LAP:BadCostMatrix',...
                'Rows and columns of the cost matrix must allow at least one possible link!')
        end
    end

    if any(~isfinite(val))
        error('LAP:NanCostMatrix','Cost matrix cannot contain NaNs or Inf!')
    end


%clear cc to save memory
clear cc;

% %I have re-written this section to save memory --Khuloud
% % write value vector, pad a zero
% compactCC = [0; val];
% % write kk, pad a zero
% kk = [0; rowIdx];
%
% % colIdx is already sorted, so we can find out the number of entries per
% % column via diff. Wherever there is a jump in the column, the deltaIdx
% % will be 1, and its rowIdx will equal fst
% deltaIdx = diff([0;colIdx]);
% % add 0 and length+1
% fst = [0;find(deltaIdx);length(val)+1];


% save some more memory: re-write values into row/col-idx. Do in the order
% row, col, val, because int32 needs less memory, and col shrinks a lot
% (but requires diff).
% Also, type the variables here already. Val should be a double, anyway,
% but better make sure, as the mex function is not forgiving.
rowIdx = int32([0;rowIdx]);
%colIdx = int32([0;find(diff([0;colIdx]));length(val)]);
val = double([0; val]);


%==================================


%==================================
% CALL MEX-FUNCTION
%==================================

% %I have re-written the following line to save memory --Khuloud
% [x, y, u, v] = mexLap(double(scc(1)), int32(length(compactCC)), double(compactCC), int32(kk), int32(fst));

% [x, y, u, v] = mexLap(double(scc(1)), int32(length(val)), double(val), ...
%     int32([0; rowIdx]), int32([0;find(diff([0;colIdx]));length(val)]));

% I saved some more memory --Jonas
% The function wants to give four outputs. u and v seem to be the costs
% associated with links in x and y, respectively.
% For some really weird reason, there is a seg-fault if colIdx is not being
% assigned above. WTF.
[x, y, u, v] = ...
    mexLap(double(scc(1)), int32(length(val)), val, ...
    rowIdx, int32([0;find(diff([0;colIdx]));length(val)])); %#ok<NASGU>

%==================================

% remove first element from output vectors, as it is a meaningless 0.
x = x(2:end);
y = y(2:end);
```
