## Supplementary material for "Adaptive-Resolution Multi-Orientation Analysis of Complex Filamentous Network Images": MATLAB code from Github: mexLap.cpp

```
/************************************************************************

   lap.cpp

   version 1.0 - 4 September 1996

   author: Roy Jonker @ MagicLogic Optimization Inc.

  


   Code for Linear Assignment Problem, according to 

   

   "A Shortest Augmenting Path Algorithm for Dense and Sparse Linear   

    Assignment Problems," Computing 38, 325-340, 1987

   

   by R. Jonker and A. Volgenant, University of Amsterdam.

*************************************************************************/
/* 
 * Compilation:
 * Mac/Linux: mex  mexLap.cpp
 * Windows: mex COMPFLAGS="$COMPFLAGS /TP /MT" -output mexLap mexLap.cpp
 */

#ifdef WINDOWS
# include <Windows.h>
#endif

#include <iostream>
#include <iomanip>
#include <time.h>     // for seconds()
#include <math.h>
#include <float.h>

#define inf DBL_MAX

/* MATLAB header files */

#include <mex.h>
#include <matrix.h>

//double lap(int n, const double cc[], const int kk[], const int first[],

//                  int x[], int y[], double u[], double v[])

/* Input Arguments */

#if !defined(N)

#define	N	            prhs[0]

#endif


#if !defined(M)

#define	M	            prhs[1]

#endif


#if !defined(CC)

#define	CC	            prhs[2]

#endif


#if !defined(KK)

#define	KK	            prhs[3]

#endif


#if !defined(FIRST)

#define	FIRST	        prhs[4]

#endif


/* Output Arguments */

#if !defined(X)

#define	X	            plhs[0]

#endif


#if !defined(Y)

#define	Y	            plhs[1]

#endif


#if !defined(U)

#define	U	            plhs[2]

#endif


#if !defined(V)

#define	V	            plhs[3]

#endif


/* Some auxilliary functions */

#if !defined(MAX)

#define	MAX(A, B)	((A) > (B) ? (A) : (B))

#endif


#if !defined(MIN)

#define	MIN(A, B)	((A) < (B) ? (A) : (B))

#endif


/////////////////////////////////////////////

double lap(int n, const double cc[], const int kk[], const int first[],

                  int x[], int y[], double u[], double v[])

{

   int h, i,j,k,l,t,last,tel,td1=0,td2,i0,j0=0,j1=0,l0;


   int *lab, *freeRow, *todo;

   bool *ok;

   double min, v0, vj, dj, tmp;

   double *d;


/* printf("LAP before computation\n");

   fflush(stdin);

   scanf(".");

*/

   ok = new bool[n + 1];

   lab = new int[n + 2];

   freeRow = new int[n + 2];

   todo = new int[n + 2];

   d = new double[n + 2];

     

   /* Initialize */

   for (j = 1; j <= n; j++) {

      v[j] = inf;

   } /* for */


   for (i = 1; i <= n; i++) {

      x[i] = 0; u[i] = 0;

      for (t = first[i]; t < first[i+1]; t++) {

         j = kk[t];

         if (cc[t] < v[j]) {

            v[j] = cc[t];

            y[j] = i;

         } /* if */

      } /* for */

   } /* for */


   for (j = n; j >= 1; j--) {

      i = y[j];

      if (x[i] == 0) {

         x[i] = j;

      } else {

         y[j] = 0;

         x[i] = -abs(x[i]);

      } /* if */

   } /* for */


   l = 0;

   for (i = 1; i <= n; i++) {

      if (x[i] < 0) {

         x[i] = -x[i];

      } else if (x[i] > 0) {

         min = inf;

         j1 = x[i];

         for (t = first[i]; t < first[i+1]; t++) {

            j = kk[t];

            if (j != j1 && cc[t] - v[j] < min) {

               min = cc[t] - v[j];

            } /* if */

         } /* for */

         u[i] = min;

         t = first[i];

         while (kk[t] != j1) {

            t++;

         } /* while */

         v[j1] = cc[t] - min;

      } else {

         freeRow[++l] = i;

      } /* if */

   } /* for */


   /* Improve initial solution */

   for (tel = 0; tel < 2; tel++) {

      h = 1;

      l0 = l;

      l = 0;

      while (h <= l0) {

         i = freeRow[h++];

         v0 = vj = inf;


         for (t = first[i]; t < first[i+1]; t++) {


            j = kk[t];

            dj = cc[t] - v[j];

            if (dj < vj) {

               if (dj >= v0) {

                  vj = dj;

                  j1 = j;

               } else {

                  vj = v0;

                  v0 = dj;

                  j1 = j0;

                  j0 = j;

               } /* if */

            } /* if */

         } /* for */


         i0 = y[j0];

         u[i] = vj;

         if (vj - v0 > FLT_EPSILON) {

            v[j0] = v[j0] - vj + v0;

         } else if (i0 > 0) {

            j0 = j1;

            i0 = y[j0];

         } /* if */


         x[i] = j0;

         y[j0] = i;


         if (i0 > 0) {

            if (vj - v0 > FLT_EPSILON) {

               freeRow[--h] = i0;

            } else {

               freeRow[++l] = i0;

            } /* if */

         } /* if */

      } /* while */

   } /* for */


   tmp = 0;

   for (i = 1; i <= n; i++) {

      tmp += u[i] + v[i];

   } /* for */


   /* Augmentation part */

   l0 = l;

   for (l = 1; l <= l0; l++) {


      for (j = 1; j <= n; j++) {

         d[j] = inf;

         ok[j] = false;

      } /* for */


      min = inf; i0 = freeRow[l];


      for (t = first[i0]; t < first[i0+1]; t++) {

         j = kk[t];

         dj = cc[t] - v[j];

         d[j] = dj;

         lab[j] = i0;


         if (dj <= min) {

            if (dj < min) {

               td1 = 0;

               min = dj;

            } /* if */

            todo[++td1] = j;

         } /* if */

      } /* for */


      for (h = 1; h <= td1; h++) {

         j = todo[h];

         if (y[j] == 0) {

            goto label2;

         } /*if */

         ok[j] = true;

      } /* for */


      td2 = n;

      last = n + 1;


      /* Repeat until a freeRow row found */

      while (true) {

         j0 = todo[td1--];

         i = y[j0];

         todo[td2--] = j0;

         t = first[i];


         for (t = first[i]; kk[t] != j0; t++) {

            /* nothing */

         } /* for */


         tmp = cc[t] - v[j0] - min;


         for (t = first[i]; t < first[i+1]; t++) {

            j = kk[t];

            if (!ok[j]) {

               vj = cc[t] - v[j] - tmp;

               if (vj < d[j]) {

                  d[j] = vj;

                  lab[j] = i;

                  if (vj == min) {

                     if (y[j] == 0) {

                        goto label1;

                     } /* if */

                     td1++;

                     todo[td1] = j;

                     ok[j] = true;

                  } /* if */

               } /* if */

            } /* if */

         } /* for */

         if (td1 == 0) {

            min = inf - 1;

            last = td2 + 1;

            for (j = 1; j <= n; j++) {

               if (d[j] <= min) {

                  if (!ok[j]) {

                     if (d[j] < min) {

                        td1 = 0;

                        min = d[j];

                     } /* if */

                     todo[++td1] = j;

                  } /* if */

               } /* if */

            } /* for */

            for (h = 1; h <= td1; h++) {

               j = todo[h];

               if (y[j] == 0) {

                  goto label1;

               } /* if */

               ok[j] = true;

            } /* for */

         } /* if */

      } /* while */

label1:

      for (k = last; k <= n; k++) {

         j0 = todo[k];

         v[j0] += d[j0] - min;

      } /* for */


label2:

      do {

         i = lab[j];

         y[j] = i;

         k = j;

         j = x[i];

         x[i] = k;

      } while (i != i0);

   } /* for */


   tmp = 0;

   for (i = 1; i <= n; i++) {

      j  = x[i];

      t = first[i];

      while (kk[t] != j) {

         t++;

      } /* while */


      u[i] = cc[t] - v[j];

      tmp += cc[t];

   } /* for */


   delete [] ok;

   delete [] lab;

   delete [] freeRow;

   delete [] todo;

   delete [] d;

   

/* printf("LAP after computation\n");

   fflush(stdin);

   scanf(".");

*/

   return(tmp);

}

/////////////////////////////////////////////


//int *kk = new int[size * NEIGHBOR_NUM_MAX + 1];

//int *first = new int[size + 2];

//int *x = new int[size + 1];

//int *y = new int[size + 1];

//double *u = new double[size + 1];

//double *v = new double[size + 1];

//double  *cc = new double[size * NEIGHBOR_NUM_MAX + 1];


// Uncomment the following for debug mode
// #define DEBUG_LAPJV


void mexFunction( int nlhs, mxArray *plhs[], int nrhs, const mxArray*prhs[])

     

{ 

	double *temp;


	

	/* The following vectors need to be allocated within this function. */


    int n;  // n is the size of the problem (the problem must be square for now) 

    int m;  // m is the size of the cost function array


	double *cc;

	int *kk;

	int *first;

	int *x;

	int *y;

	double *u;

	double *v;


// 	double *kk_temp, *first_temp;


// 	int i;

	


	/* Step 0: check input/output numbers */


	/* Check for proper number of arguments */

    if (nrhs != 5) { 

		mexErrMsgTxt("Error: five input arguments are required."); 

    } 

	else if (nlhs != 4) {

		mexErrMsgTxt("Error: four output arguments are required."); 

    } 


    

	/* Step 1: read in the dimensions */

     

	temp = mxGetPr(N);

	n = (int) (*temp);


	temp = mxGetPr(M);

	m = (int) (*temp);


#ifdef DEBUG_LAPJV

	mexPrintf("Size of question = %d  Size of cost matrix = %d\n", n, m);

#endif


	/* Step 2: Set all the input data pointers */

	cc = (double *)mxGetData(CC);

    kk = (int *)mxGetData(KK);

	first = (int *)mxGetData(FIRST);


#ifdef DEBUG_LAPJV

	for (i = 0; i < n + 2; i++)

		mexPrintf("First element = %d   KK = %d\n", first[i], kk[i]);

#endif


	/* Step 3: Set output variables */

	X = mxCreateNumericMatrix(n + 1, 1, mxINT32_CLASS, mxREAL);

	x = (int *)mxGetData(X);


	Y = mxCreateNumericMatrix(n + 1, 1, mxINT32_CLASS, mxREAL);

	y = (int *)mxGetData(Y);


	U = mxCreateNumericMatrix(n + 1, 1, mxDOUBLE_CLASS, mxREAL);

	u = (double *)mxGetData(U);


	V = mxCreateNumericMatrix(n + 1, 1, mxDOUBLE_CLASS, mxREAL);

	v = (double *)mxGetData(V);


    lap(n, cc, kk, first, x, y, u, v);


    return;

}
```
