## Supplementary material for "Adaptive-Resolution Multi-Orientation Analysis of Complex Filamentous Network Images": MATLAB code from Github: normList.m

```
function [listOfNorms,normedVectors]=normList(vectors,doCheck,correctNaN)
%calculates the norm of a list of vectors
%
%SYNOPSIS [listOfNorms,normedVectors]=normList(vectors,doCheck,correctNaN)
%
%INPUT list of vectors (nVectorsXdimension)
%      noCheck (optional, true/false, default: true): normList checks for
%           old versions of Matlab that do not have bsxfun. This check is
%           slow, so if you know that you'll have Matlab 7.6 or later, set
%           doCheck to false to speed up computation. 
%      correctNaN (optional, true/false, default: true): if true, NaNs in
%           unit vectors will be set to zero (a zero-length vector will
%           lead to all NaNs since it's divided by the norm, resulting in
%           0/0).
%
%OUTPUT listOfNorms: list (nX1) containing the norms of the vectors
%       normedVectors: list (nXdim) containing the normed vectors
%
%c: 1/03 Jonas
%%%%%%%%%%%%%%%%%%%%%%%%%%%%%%%%%%%%%%%%%%%%%%%%%%%%%%%%%%%%%%%%%%%%%%%%%%%
%
% Copyright (C) 2019, Jaqaman Lab - UT Southwestern, Goldman Lab - Northwestern 
%
% This file is part of AdaptiveResolutionOrientationSpace.
% 
% AdaptiveResolutionOrientationSpace is free software: you can redistribute it and/or modify
% it under the terms of the GNU General Public License as published by
% the Free Software Foundation, either version 3 of the License, or
% (at your option) any later version.
% 
% AdaptiveResolutionOrientationSpace is distributed in the hope that it will be useful,
% but WITHOUT ANY WARRANTY; without even the implied warranty of
% MERCHANTABILITY or FITNESS FOR A PARTICULAR PURPOSE.  See the
% GNU General Public License for more details.
% 
% You should have received a copy of the GNU General Public License
% along with AdaptiveResolutionOrientationSpace.  If not, see <http://www.gnu.org/licenses/>.
% 
% 
nVectors = size(vectors,1);
nDims = size(vectors,2);

if nargin < 2 || isempty(doCheck)
    doCheck = true;
end
if nargin < 3 || isempty(correctNaN)
    correctNaN = true;
end

%listOfNorms=zeros(nVectors,1);
listOfNorms=sqrt(sum(vectors.^2,2));

% the unit vector of length 0 is [0 0 0]
if nargout > 1
    normedVectors=zeros(size(vectors));
    if doCheck && verLessThan('matlab','7.6')
        goodVectors=find(listOfNorms);
        normedVectors(goodVectors,:)=vectors(goodVectors,:)./(repmat(listOfNorms(goodVectors),[1,nDims]));
    else
        normedVectors = bsxfun(@rdivide,vectors,listOfNorms);
        if correctNaN
            normedVectors(isnan(normedVectors)&~isnan(vectors)) = 0;
        end
    end
end
```
