## Supplementary material for "Adaptive-Resolution Multi-Orientation Analysis of Complex Filamentous Network Images": MATLAB code from Github: padarrayXT.m

```
function b = padarrayXT(varargin)
%PADARRAYXT Modified version of built-in function 'padarray'.
%   Mirroring ('symmetric' option) does not duplicate the border pixel.
%   B = PADARRAYXT(A,PADSIZE) pads array A with PADSIZE(k) number of zeros
%   along the k-th dimension of A.  PADSIZE should be a vector of
%   nonnegative integers.
%
%   B = PADARRAYXT(A,PADSIZE,PADVAL) pads array A with PADVAL (a scalar)
%   instead of with zeros.
%
%   B = PADARRAYXT(A,PADSIZE,PADVAL,DIRECTION) pads A in the direction
%   specified by the string DIRECTION.  DIRECTION can be one of the
%   following strings.
%
%       String values for DIRECTION
%       'pre'         Pads before the first array element along each
%                     dimension .
%       'post'        Pads after the last array element along each
%                     dimension.
%       'both'        Pads before the first array element and after the
%                     last array element along each dimension.
%
%   By default, DIRECTION is 'both'.
%
%   B = PADARRAYXT(A,PADSIZE,METHOD,DIRECTION) pads array A using the
%   specified METHOD.  METHOD can be one of these strings:
%
%       String values for METHOD
%       'circular'    Pads with circular repetition of elements.
%       'replicate'   Repeats border elements of A.
%       'symmetric'   Pads array with mirror reflections of itself.
%       'asymmetric'   Pads array with odd-symmetric extensions of itself. 
%
%   Class Support
%   -------------
%   When padding with a constant value, A can be numeric or logical.
%   When padding using the 'circular', 'replicate', or 'symmetric'
%   methods, A can be of any class.  B is of the same class as A.
%
%   Example
%   -------
%   Add three elements of padding to the beginning of a vector.  The
%   padding elements contain mirror copies of the array.
%
%       b = padarrayxt([1 2 3 4],3,'symmetric','pre')
%
%   Add three elements of padding to the end of the first dimension of
%   the array and two elements of padding to the end of the second
%   dimension.  Use the value of the last array element as the padding
%   value.
%
%       B = padarrayxt([1 2; 3 4],[3 2],'replicate','post')
%
%   Add three elements of padding to each dimension of a
%   three-dimensional array.  Each pad element contains the value 0.
%
%       A = [1 2; 3 4];
%       B = [5 6; 7 8];
%       C = cat(3,A,B)
%       D = padarray(C,[3 3],0,'both')
%
%   See also CIRCSHIFT, IMFILTER.
%
% Copyright (C) 2019, Jaqaman Lab - UT Southwestern, Goldman Lab - Northwestern 
%
% This file is part of AdaptiveResolutionOrientationSpace.
% 
% AdaptiveResolutionOrientationSpace is free software: you can redistribute it and/or modify
% it under the terms of the GNU General Public License as published by
% the Free Software Foundation, either version 3 of the License, or
% (at your option) any later version.
% 
% AdaptiveResolutionOrientationSpace is distributed in the hope that it will be useful,
% but WITHOUT ANY WARRANTY; without even the implied warranty of
% MERCHANTABILITY or FITNESS FOR A PARTICULAR PURPOSE.  See the
% GNU General Public License for more details.
% 
% You should have received a copy of the GNU General Public License
% along with AdaptiveResolutionOrientationSpace.  If not, see <http://www.gnu.org/licenses/>.
% 
% 

%   Copyright 1993-2010 The MathWorks, Inc.
%   $Revision: 1.11.4.13 $  $Date: 2011/08/09 17:51:33 $

%   Last modified on 04/14/2012 by Francois Aguet.

[a, method, padSize, padVal, direction] = ParseInputs(varargin{:});

if isempty(a)
    
    % treat empty matrix similar for any method
    if strcmp(direction,'both')
        sizeB = size(a) + 2*padSize;
    else
        sizeB = size(a) + padSize;
    end
    
    b = mkconstarray(class(a), padVal, sizeB);
    
elseif strcmpi(method,'constant')
    
    % constant value padding with padVal
    b = ConstantPad(a, padSize, padVal, direction);
else
    
    % compute indices then index into input image
    aSize = size(a);
    aIdx = getPaddingIndices(aSize,padSize,method,direction);
    b = a(aIdx{:});
end

if islogical(a)
    b = logical(b);
end


%%%
%%% ConstantPad
%%%
function b = ConstantPad(a, padSize, padVal, direction)

numDims = numel(padSize);

% Form index vectors to subsasgn input array into output array.
% Also compute the size of the output array.
idx   = cell(1,numDims);
sizeB = zeros(1,numDims);
for k = 1:numDims
    M = size(a,k);
    switch direction
        case 'pre'
            idx{k}   = (1:M) + padSize(k);
            sizeB(k) = M + padSize(k);
            
        case 'post'
            idx{k}   = 1:M;
            sizeB(k) = M + padSize(k);
            
        case 'both'
            idx{k}   = (1:M) + padSize(k);
            sizeB(k) = M + 2*padSize(k);
    end
end

% Initialize output array with the padding value.  Make sure the
% output array is the same type as the input.
b         = mkconstarray(class(a), padVal, sizeB);
b(idx{:}) = a;


%%%
%%% ParseInputs
%%%
function [a, method, padSize, padVal, direction] = ParseInputs(varargin)

% narginchk(2,4);
if nargin<2 || nargin>4
    error('Incompatible number of input arguments.');
end

% fixed syntax args
a         = varargin{1};
padSize   = varargin{2};

% default values
method    = 'constant';
padVal    = 0;
direction = 'both';

validateattributes(padSize, {'double'}, {'real' 'vector' 'nonnan' 'nonnegative' ...
    'integer'}, mfilename, 'PADSIZE', 2);

% Preprocess the padding size
if (numel(padSize) < ndims(a))
    padSize           = padSize(:);
    padSize(ndims(a)) = 0;
end

if nargin > 2
    
    firstStringToProcess = 3;
    
    if ~ischar(varargin{3})
        % Third input must be pad value.
        padVal = varargin{3};
        validateattributes(padVal, {'numeric' 'logical'}, {'scalar'}, ...
            mfilename, 'PADVAL', 3);
        
        firstStringToProcess = 4;
        
    end
    
    for k = firstStringToProcess:nargin
        validStrings = {'circular' 'replicate' 'symmetric' 'pre' ...
            'post' 'both'};
        string = validatestring(varargin{k}, validStrings, mfilename, ...
            'METHOD or DIRECTION', k);
        switch string
            case {'circular' 'replicate' 'symmetric'}
                method = string;
                
            case {'pre' 'post' 'both'}
                direction = string;
                
            otherwise
                error(message('images:padarray:unexpectedError'))
        end
    end
end

% Check the input array type
if strcmp(method,'constant') && ~(isnumeric(a) || islogical(a))
    error(message('images:padarray:badTypeForConstantPadding'))
end


% Internal functions called by padarray.m (modified)
function aIdx = getPaddingIndices(aSize,padSize,method,direction)
%getPaddingIndices is used by padarray and blockproc. 
%   Computes padding indices of input image.  This is function is used to
%   handle padding of in-memory images (via padarray) as well as
%   arbitrarily large images (via blockproc).
%
%   aSize : result of size(I) where I is the image to be padded
%   padSize : padding amount in each dimension.  
%             numel(padSize) can be greater than numel(aSize)
%   method : X or a 'string' padding method
%   direction : pre, post, or both.
%
%   See the help for padarray for additional information.

% Copyright 2010 The MathWorks, Inc.
% $Revision: 1.1.6.1 $ $Date: 2010/04/15 15:18:15 $

% make sure we have enough image dims for the requested padding
if numel(padSize) > numel(aSize)
    singleton_dims = numel(padSize) - numel(aSize);
    aSize = [aSize ones(1,singleton_dims)];
end

switch method
    case 'circular'
        aIdx = CircularPad(aSize, padSize, direction);
    case 'symmetric'
        aIdx = SymmetricPad(aSize, padSize, direction);
    case 'replicate' 
        aIdx = ReplicatePad(aSize, padSize, direction);
end


%%%
%%% CircularPad
%%%
function idx = CircularPad(aSize, padSize, direction)

numDims = numel(padSize);

% Form index vectors to subsasgn input array into output array.
% Also compute the size of the output array.
idx   = cell(1,numDims);
for k = 1:numDims
    M = aSize(k);
    dimNums = uint32(1:M);
    p = padSize(k);
    
    switch direction
        case 'pre'
            idx{k}   = dimNums(mod(-p:M-1, M) + 1);
            
        case 'post'
            idx{k}   = dimNums(mod(0:M+p-1, M) + 1);
            
        case 'both'
            idx{k}   = dimNums(mod(-p:M+p-1, M) + 1);
            
    end
end


%%%
%%% SymmetricPad
%%%
function idx = SymmetricPad(aSize, padSize, direction)

numDims = numel(padSize);

% Form index vectors to subsasgn input array into output array.
% Also compute the size of the output array.
idx   = cell(1,numDims);
for k = 1:numDims
    M = aSize(k);
    if M>1
        dimNums = uint32([1:M M-1:-1:2]);
        div = 2*M-2;
    else
        dimNums = [1 1];
        div = 2;
    end
    p = padSize(k);
    
    switch direction
        case 'pre'
            idx{k}   = dimNums(mod(-p:M-1, div) + 1);
            
        case 'post'
            idx{k}   = dimNums(mod(0:M+p-1, div) + 1);
            
        case 'both'
            idx{k}   = dimNums(mod(-p:M+p-1, div) + 1);
    end
end


%%%
%%% ReplicatePad
%%%
function idx = ReplicatePad(aSize, padSize, direction)

numDims = numel(padSize);

% Form index vectors to subsasgn input array into output array.
% Also compute the size of the output array.
idx   = cell(1,numDims);
for k = 1:numDims
    M = aSize(k);
    p = padSize(k);
    onesVector = uint32(ones(1,p));
    
    switch direction
        case 'pre'
            idx{k}   = [onesVector 1:M];
            
        case 'post'
            idx{k}   = [1:M M*onesVector];
            
        case 'both'
            idx{k}   = [onesVector 1:M M*onesVector];
    end
end


function out = mkconstarray(class, value, size)
%MKCONSTARRAY creates a constant array of a specified numeric class.
%   A = MKCONSTARRAY(CLASS, VALUE, SIZE) creates a constant array 
%   of value VALUE and of size SIZE.

%   Copyright 1993-2003 The MathWorks, Inc.  
%   $Revision: 1.8.4.1 $  $Date: 2003/01/26 06:00:35 $

out = repmat(feval(class, value), size);
```
