## Supplementary material for "Adaptive-Resolution Multi-Orientation Analysis of Complex Filamentous Network Images": MATLAB code from Github: perpVector.m

```
function [v_perp v_proj]=perpVector(point1,e_1,point2)
%finds the perpendicular vector v connecting the straight line defined by
%point1 and e_1 with point2
%
%SYNOPSIS v_perp=perpVector(point1,e_1,point2)
%
%INPUT point1, e_1 : point and UNIT vector defining the straight line
%      point2: point from which the new perpendicular vector should start
%      (all inputs can be lists of n-by-d, where d is the dimension)
%
%OUTPUT v_perp perpendicular vector from the straight line to the point
%           (norm(v_perp)=distance of the point from the line)
%
%c: 1/03 Jonas
%%%%%%%%%%%%%%%%%%%%%%%%%%%%%%%%%%%%%%%%%%%%%%%%%%%%%%%%%%%%%%%%%%%%%%%%%%%
%
% Copyright (C) 2019, Jaqaman Lab - UT Southwestern, Goldman Lab - Northwestern 
%
% This file is part of AdaptiveResolutionOrientationSpace.
% 
% AdaptiveResolutionOrientationSpace is free software: you can redistribute it and/or modify
% it under the terms of the GNU General Public License as published by
% the Free Software Foundation, either version 3 of the License, or
% (at your option) any later version.
% 
% AdaptiveResolutionOrientationSpace is distributed in the hope that it will be useful,
% but WITHOUT ANY WARRANTY; without even the implied warranty of
% MERCHANTABILITY or FITNESS FOR A PARTICULAR PURPOSE.  See the
% GNU General Public License for more details.
% 
% You should have received a copy of the GNU General Public License
% along with AdaptiveResolutionOrientationSpace.  If not, see <http://www.gnu.org/licenses/>.
% 
% 

%============================
% TEST INPUT
%============================

[nRows1, nCols1] = size(point1);
[nRowsE, nColsE] = size(e_1);
[nRows2, nCols2] = size(point2);

if ~all([nCols1,nCols2]==nColsE)
    error('inconsistent dimensions!');
end

nEntries = max([nRows1,nRowsE,nRows2]);
if ~all([nRows1,nRowsE,nRows2]==nEntries | [nRows1,nRowsE,nRows2]==1)
    error('inconsistent number of entries')
end

if nRows1 == 1
    point1 = repmat(point1,nEntries,1);
end
if nRows2 == 1
    point2 = repmat(point2,nEntries,1);
end
if nRowsE == 1
    e_1 = repmat(e_1,nEntries,1);
end

%===============================


%===============================
% CALCULATE DISTANCE
%===============================

%vector between point1 and point2
v_temp=point2-point1;

%projection of v_temp on e_1
v_proj=(repmat(sum(v_temp.*e_1,2),1,nCols1)).*e_1;

%v_temp - the parallel projection = the perpendicular part
v_perp=v_temp-v_proj;
```
