## Supplementary material for "Adaptive-Resolution Multi-Orientation Analysis of Complex Filamentous Network Images": MATLAB code from Github: wraparoundN.m

```
function [wrappedValues, multiplier] = wraparoundN(values,lower,upper)
%WRAPAROUNDN wraps values onto an interval [lower, upper) for N dim arrays
%
% SYNOPSIS [wrappedValues, multiplier] = wraparound(values,interval)
%
% INPUT    values      : n-by-d array of values to be wrapped
%          lower (opt) : lower bound of the wrap, singleton expanded
%          upper (opt) : upper bound of the wrap, singleton expanded
%
%          If upper is not specified, lower will be split along the first
%          dimension of size 2.
%
% OUTPUT   wrappedValues : n-by-d values as projected onto the interval
%          multiplier    : multiplier of interval to reconstitute original
%                           value
%
% REMARKS   The final value will lie within the interval [lower, upper).
%           It will never have th value of upper. To find out how
%           far away from the lower limit the value lies, you have to
%           subtract that lower limit yourself
%           Remember: If the allowed values range from 1 to 10, lower=1 and
%           upper = 11, because you want to allow a value of 10, but you
%           want to turn 11 into 1.
%
% EXAMPLES  With an interval [0;10], 19 becomes [[9], [1]]
%           With an interval [2;4],  [9.2;-2.6] becomes [[3.2;3.4], [3;-3]]
%           With an interval [-180,180], [270;360] becomes [[-90;0], [1;1]]
%
% See also bsxfun, mod
%
% Mark Kittisopikul
% August 26th, 2015
% Jaqaman Lab
% UT Southwestern
% Inspired by Jonas Dorn's wraparound
%%%%%%%%%%%%%%%%%%%%%%%%%%%%%%%%%%%%%%%%%%%%%%%%%%%%%%%%%%%%%%%%%%%%%%%%%%%
%
% Copyright (C) 2019, Jaqaman Lab - UT Southwestern, Goldman Lab - Northwestern 
%
% This file is part of AdaptiveResolutionOrientationSpace.
% 
% AdaptiveResolutionOrientationSpace is free software: you can redistribute it and/or modify
% it under the terms of the GNU General Public License as published by
% the Free Software Foundation, either version 3 of the License, or
% (at your option) any later version.
% 
% AdaptiveResolutionOrientationSpace is distributed in the hope that it will be useful,
% but WITHOUT ANY WARRANTY; without even the implied warranty of
% MERCHANTABILITY or FITNESS FOR A PARTICULAR PURPOSE.  See the
% GNU General Public License for more details.
% 
% You should have received a copy of the GNU General Public License
% along with AdaptiveResolutionOrientationSpace.  If not, see <http://www.gnu.org/licenses/>.
% 
% 

%=====================
% TEST INPUT
%=====================

if(nargin < 2)
    lower = 0;
end
if(nargin < 3)
    % upper is missing
    intervalSize = size(lower);
    % can we split lower into two halves for compatability with jonas'
    % wraparound?
    dimToSplit = find(intervalSize == 2,1);
    if(isempty(dimToSplit))
        % cannot split, default is then lower + 1
        upper = lower + 1;
    else
        % Understand lower to be an interval as in jonas' wraparound
        % Shift dim with length(2) to be the first dim
        interval = shiftdim(lower,dimToSplit-1);
        shiftedSize = size(interval);
        
        % Split interval to lower and upper, shift back so other dimensions
        % are where they were originally
        lower = interval(1,:);
        lower = reshape(lower,[1 shiftedSize(2:end)]);
        lower = shiftdim(lower,ndims(lower)-dimToSplit+1);
        
        upper = interval(2,:);
        upper = reshape(upper,[1 shiftedSize(2:end)]);
        upper = shiftdim(upper,ndims(lower)-dimToSplit+1);
    end
end

% Checks we do not need
% 1. We do not need to check number of parameters due to reasonable
% defaults
% 2. We are not limited to 2D arrays due to singleton expansion via bsxfun
% 3. We do not need to check size of interval
% 4. We do not need to check to integers because we do not scale

%=========================
% WRAP AROUND
%=========================

% Strategy: Transform interval and data to [0,U-L], Then use mod
wrappedValues = bsxfun(@minus,values,lower);
upper = bsxfun(@minus,upper,lower);
assert(all(upper(:)),'lower limits have to be strictly smaller than upper limits!')

if(nargout > 1)
    % calculate multiplier
    shiftedValues = wrappedValues;
    wrappedValues = bsxfun(@mod,wrappedValues,upper);
    multiplier = bsxfun(@rdivide,shiftedValues-wrappedValues,upper);
else
    wrappedValues = bsxfun(@mod,real(wrappedValues),upper);
end

wrappedValues = bsxfun(@plus,wrappedValues,lower);
```
