## Supplementary material for "Adaptive-Resolution Multi-Orientation Analysis of Complex Filamentous Network Images": MATLAB code from Github: joinColumns.m

```
function varargout = joinColumns(delimeter,varargin)
%joinColumns vertically concatenate columns separated by a delimeter
%
% [colVecA, colVecB, colVecC, ...] = joinColumns(delimeter, A, B, C, ...)
%
% INPUT
%
% delimeter is a scalar value that will be placed between the columns of
%    the matrices. An empty matrix indicates no delimeter.
%    (optional. A non-scalar value is considered a matrix and the delimeter
%    will be empty)
%
% A, B, C ... are matrices that can hold delimeter as a value. They need
% not be the same size or type
%
% OUTPUT
%
% columnVector - a column vector is returned for each matrix generated by
% concatenating the matrices vertically separated by the scalar delimeter
%
% joinColumns(matrix) and joinColumns([],matrix) are equivalent to matrix(:)
%
% Copyright (C) 2019, Jaqaman Lab - UT Southwestern, Goldman Lab - Northwestern 
%
% This file is part of AdaptiveResolutionOrientationSpace.
% 
% AdaptiveResolutionOrientationSpace is free software: you can redistribute it and/or modify
% it under the terms of the GNU General Public License as published by
% the Free Software Foundation, either version 3 of the License, or
% (at your option) any later version.
% 
% AdaptiveResolutionOrientationSpace is distributed in the hope that it will be useful,
% but WITHOUT ANY WARRANTY; without even the implied warranty of
% MERCHANTABILITY or FITNESS FOR A PARTICULAR PURPOSE.  See the
% GNU General Public License for more details.
% 
% You should have received a copy of the GNU General Public License
% along with AdaptiveResolutionOrientationSpace.  If not, see <http://www.gnu.org/licenses/>.
% 
% 
    
    % INPUT VALIDATION: If delimeter is not a scalar value, then assume no
    % delimeter is given and that the matrix should be turned into a column
    % vector
    if(isempty(delimeter) && isempty(varargin))
%         varargout{1} = NaN;
        varargout{1} = delimeter;
        return;
    end
    if(~isscalar(delimeter))
        % add delimeter to the cell array of matrices to linearize
        % if delimeter is empty, this should have no effect
        varargin = [ delimeter varargin];
        delimeter = [];
    elseif(isempty(varargin))
        % Only a scalar was given, just return it
        varargout{1} = delimeter;
        return;
    end
    
    % For each matrix given linearize and output. This allows for
    % vectorized syntax and is compatible with paralleization with
    % distributed objects
    
    % We only need to check if the delimeter is empty once
    if(~isempty(delimeter))
        varargin = cellfun(@appendDelimeter,varargin,'UniformOutput',false);
        varargout = cellfun(@joinColumnsSingle,varargin,'UniformOutput',false);
        varargout = cellfun(@(x) x(1:end-1),varargout,'UniformOutput',false);
    else
        varargout = cellfun(@joinColumnsSingle,varargin,'UniformOutput',false);
    end
    
    function matrix = appendDelimeter(matrix)
        % append the delimeter as the last row
        matrix(end+1,:) = delimeter;
    end
    function columnVector = joinColumnsSingle(matrix)
        % linearize
        columnVector = matrix(:);
    end
end
```
