## Supplementary material for "Adaptive-Resolution Multi-Orientation Analysis of Complex Filamentous Network Images": MATLAB code from Github: parcellfun_progress.m

```
function [ varargout ] = parcellfun_progress( func, varargin )
%parcellfun_progress is a parallelized cellfun that also has
% a progress monitor built-in.
%
% INPUT
% func - function_handle as in cellfun
% ...  - variable number of cell arrays
% PARAMETERS 
%  UniformOutput
%   true:  Combine output into an array the same size as input
%          Function output should be a scalar and can be concatenated.
%   false: Output a cell array the same size as input
%  ErrorHandler: function_handle that takes the MException error caught as an argument
%                Output will replace normal function output
%  UpdateInterval: How often to update the clock in seconds
%  DisplayFunc: function_handle to control display or char string
%    Input is a structure with the following fields
%    .in - contents of unmatched parameters
%    .nRuns - total number of iterations of func to be done
%    .nCompleted - number of completed iterations of func
%    .nCompletedTime - datetime object when the last parallel output was fetched
%    .nErrors - number of iterations that ended in an error
%    .nProgressOut - number of characters output by last call to DisplayFunc
%    .startDT - datetime object indicating when parallel calls were queued
%    .notCompleted - logical array that is true if an element is not complete
%    .completedIdx - the last index of the completed feature
%    .F parallel.FevalFuture objects
%    Output will be the number of characters printed by this function that is solely
%      passed back into the function on the next iteration
%
%    DisplayFunc can also be one the following
%      'progressText' - use @progressText to track progress
%      'progressTextMultiple' - use @progressTextMultiple to track progress
%      'parfor_progress' - use @parfor_progress to track progress
%      See @parcellfun_progress/emulate_{DisplayFunc} for more details
%  ParallelPool - parallel.pool to use with parfeval
%  Heading - optional char to be output by fprintf before iterating
%  UseErrorStruct - true: pass error struct as defined in cellfun (default)
%                  false: pass MException and same structure as DisplayFunc
%  NumOutputs - specify number of outputs
%  ReturnFutures - parallel.FevalFuture objects as first output
%                  and cell array of func outputs as second output
%  DisplayDiaries - display output of func as it's output is retrieved
%
% OUTPUT
% Variable output as in cellfun
% Each output is either:
%  cell array the same size as input if not UniformOutput
%  array the same size as input if not UniformOutput
% The number of outputs can be up to the number available from func
%
% See also cellfun, distributed.cellfun, parfor
%
% Copyright (C) 2019, Jaqaman Lab - UT Southwestern, Goldman Lab - Northwestern 
%
% This file is part of AdaptiveResolutionOrientationSpace.
% 
% AdaptiveResolutionOrientationSpace is free software: you can redistribute it and/or modify
% it under the terms of the GNU General Public License as published by
% the Free Software Foundation, either version 3 of the License, or
% (at your option) any later version.
% 
% AdaptiveResolutionOrientationSpace is distributed in the hope that it will be useful,
% but WITHOUT ANY WARRANTY; without even the implied warranty of
% MERCHANTABILITY or FITNESS FOR A PARTICULAR PURPOSE.  See the
% GNU General Public License for more details.
% 
% You should have received a copy of the GNU General Public License
% along with AdaptiveResolutionOrientationSpace.  If not, see <http://www.gnu.org/licenses/>.
% 
% 

% Mark Kittisopikul, December 2015

    %% Process Input

    ip = inputParser;
    ip.StructExpand = true;
    ip.KeepUnmatched = true;
    ip.addParameter('UniformOutput',true,@islogical);
    ip.addParameter('ErrorHandler',@defaultErrorHandler,@(x) isa(x,'function_handle'));
    ip.addParameter('UpdateInterval',1,@(x) validateattributes(x,{'numeric'},{'scalar'}));
    ip.addParameter('DisplayFunc',@defaultDisplayFunc,@(x) isa(x,'function_handle') || ischar(x));
    ip.addParameter('ParallelPool',[]);
    ip.addParameter('Heading','',@ischar);
    ip.addParameter('UseErrorStruct',true,@islogical);
    ip.addParameter('NumOutputs',[],@(x) validateattributes(x,{'numeric'},{'scalar'}));
    ip.addParameter('ReturnFutures',false,@islogical);
    ip.addParameter('DisplayDiaries',false,@islogical);
    
    narginchk(2,Inf);
        

    % Find first non-cell input to begin parsing parameters
    for inIdx = 1:length(varargin)
       if(~iscell(varargin{inIdx}))
            paramIdx = inIdx;
            break;
       elseif(inIdx == length(varargin))
           paramIdx = inIdx + 1;
       end
    end
    
    if(paramIdx == 1)
        error('parcellfun_progress:NotACell', ...
         'Input #2 expected to be a cell array.');
    end

    % Parse input parameters
    ip.parse(varargin{paramIdx:end});
    in = ip.Results;
    d.in = ip.Unmatched;
    
    % Initialize parallel pool
    if(isempty(in.ParallelPool))
        in.ParallelPool = gcp;
        if(isempty(in.ParallelPool))
            error('parcellfun_progress:CannotObtainParallelPool', ...
                'Cannot obtain parallel pool.');
        end
    end
    
    if(ischar(in.DisplayFunc))
        in.DisplayFunc = str2func(['emulate_' in.DisplayFunc]);
    end
    
    if(isequal(in.ErrorHandler,@defaultErrorHandler))
        in.UseErrorStruct = false;
    end
    
    if(~isempty(in.Heading))
        fprintf(in.Heading);
    end
   
    if(isempty(in.NumOutputs))
        in.NumOutputs = nargout - in.ReturnFutures;
    end
    
    if(nargout > in.NumOutputs + in.ReturnFutures)
        error('parcellfun_progress:InvalidNumOutputsVsNargout', ...
            'NumOutputs cannot be less than the requested number of outputs');
    end

    %% Initialize variables
    % Number of outputs
    nout = in.NumOutputs;
    % Size of cell input
    inSize = size(varargin{1});
    % Number of elements in cell input (number of parallel tasks)
    d.nRuns = numel(varargin{1});
    % Cell array to hold all output
    out = cell(d.nRuns,nout);
    % Logical array to mark which jobs have completed
    d.notComplete = true(d.nRuns,1);
    % Number of workers completed
    d.nCompleted = 0;
    % Time it took to complete d.nCompletedTime
    d.nCompletedTime = 0;
    % Number of errors
    d.nErrors = 0;  
    % Number of bytes output by progress, for backspacing
    d.nProgressOut = 0;
    % cache number of workers
    d.nWorkers = in.ParallelPool.NumWorkers;
    % Index from 1 to nRuns in case we need to figure out d.completedIdx
    idx = 1:d.nRuns;
    
    %% Start parallel execution
    d.startTimerVal = tic;
    
    % Initial display function
    d.nProgressOut = in.DisplayFunc(d);
    drawnow;
    
    d.F = cellfun(@parfunc,varargin{1:paramIdx-1},'UniformOutput',false);
    d.F = [d.F{:}];
    
    % When this function completes, user quits, or exception occurs,
    % cleanup
    cleanup = onCleanup(@()  cancel(d.F));
    
    % We will not update the estimate until we have processed all finished
    % workers, so store a copy of the structure
    dlast = d;
    
    % Main output checking loop
    while(d.nCompleted < d.nRuns)
        d.completedIdx = -1;
        % Process finished workers available at the moment.
        % Otherwise the estimate would update too frequently when multiple
        % workers return at the same time.
        while(~isempty(d.completedIdx) && d.nCompleted < d.nRuns)
            % We do not know completedIdx Yet
            % Define currentout, there is no nargout bump
            currentOut = cell(1,nout);
            try
                [d.completedIdx,currentOut{1:nout}] = fetchNext(d.F,in.UpdateInterval);
            catch err
                d.nErrors = d.nErrors + 1;
                % Find d.completedIdx
                notCompleteIdx = idx(d.notComplete);
                % Calls to parallel.Future.Read are expensive
                % Minimize calls to F.Read by using a for loop
                % Also, jobs are likely to complete sequentially, so we
                %   will probably find the answer quickly.
                for i=notCompleteIdx
                    if(d.F(i).Read)
                        d.completedIdx = i;
                        break;
                    end
                end
                % Functionally quivalent to the above, but not as fast
                % d.completedIdx = notCompleteIdx([d.F(d.notComplete).Read]);
                d.InputArguments = d.F(d.completedIdx).InputArguments;
                if(in.UseErrorStruct)
                    % For backwards compatability
                    s.identifier = err.identifier;
                    s.message = err.message;
                    s.index = d.completedIdx;
                    s.d = d;
                    if(nargout(in.ErrorHandler))
                        [currentOut{1:nout}] = in.ErrorHandler(s,d.InputArguments{:});
                    else
                        in.ErrorHandler(s,d.InputArguments{:});
                    end
                else
                    % Passes a MException and the full status data structure
                    if(nargout(in.ErrorHandler))
                        [currentOut{1:nout}] = in.ErrorHandler(err,d);
                    else
                        in.ErrorHandler(err,d);
                    end
                end
            end
            if(~isempty(d.completedIdx))
                % Worker returned output
                d.notComplete(d.completedIdx) = false;
                out(d.completedIdx,:) = currentOut;
                d.nCompleted = d.nCompleted + 1;
                d.nCompletedTime = toc(d.startTimerVal);
                if(in.DisplayDiaries)
                    fprintf('=> Diary of Index %d\n\n',d.completedIdx);
                    disp(d.F(d.completedIdx).Diary);
                    d.nProgressOut = 0;
                    dlast.nProgressOut = 0;
                    fprintf('\n\n<= Diary of Index %d\n\n',d.completedIdx);
                end
            end
            if(d.nCompleted - dlast.nCompleted > d.nWorkers)
                % Force estimate update after nWorkers have finished
                d.completedIdx = [];
            else
                d.nProgressOut = in.DisplayFunc(dlast);
                dlast.nProgressOut = d.nProgressOut;
            end
        end
        % New estimate completed, update timers
        d.nProgressOut = in.DisplayFunc(d);
        dlast = d;
    end
    
    % Convert 2D cell array into a 1D cell array of cells
    varargout = num2cell(out,1);
    if(in.UniformOutput && nout ~= 0)
        varargout = cellfun(@cell2mat,varargout,'UniformOutput',false);
        nArgOut = cellfun('prodofsize',varargout);
        assert(numel(nArgOut) == 1 && all(nArgOut(1) == nArgOut(2:end)), ...
            'parcellfun_progress:NonUniformOutput', ...
            'Non-uniform output with UniformOutput set to true');
    end
    % Make output reflect size of input
    varargout = cellfun(@(out) reshape(out,inSize),varargout,'UniformOutput',false);
    
    if(in.ReturnFutures)
        varargout = [ {d.F} varargout];
    end
        
    % Wrap func in parfeval for evaluation
    function F = parfunc(varargin)
        F = parfeval(in.ParallelPool,func,nout,varargin{:});
    end
end
function defaultErrorHandler(exception, varargin)
    rethrow(exception);
end
function nProgressOut = defaultDisplayFunc(d)
    % Prepare to erase previous output
    bp = repmat('\b',1,d.nProgressOut);
    sinceStart = toc(d.startTimerVal);
    if(~d.nCompleted)
        estDuration = 'hh:mm:ss';
        remaining = 'hh:mm:ss';
        sinceStart = duration_shim(0,0,sinceStart);
    else
        fracCompleted = d.nCompleted / ceil(d.nRuns/d.nWorkers) / d.nWorkers;
        estDuration = d.nCompletedTime / fracCompleted;
        estDuration = max(estDuration,sinceStart);
        remaining = estDuration - sinceStart;
        estDuration = duration_shim(0,0,estDuration);
        remaining = duration_shim(0,0,remaining);
        sinceStart = duration_shim(0,0,sinceStart);
    end
    if(d.nRuns ~= d.nCompleted)
        if(d.nErrors > 0)
            % Count errors
            nProgressOut = fprintf([bp ...
                ' %g / %g = %3.0f%%\n' ...
                'Errors: %g / %g = %3.0f%%\n' ...
                '  Estimated %s\n' ...
                '-   Elapsed %s\n' ...
                '---------------------\n' ...
                '= Remaining %s\n'],...
                d.nCompleted,d.nRuns,d.nCompleted/d.nRuns*100, ...
                d.nErrors,d.nCompleted,d.nErrors/d.nCompleted*100, ...
                char(estDuration), ...
                char(sinceStart), ...
                char(remaining) ...
                ) - d.nProgressOut;
        else
            nProgressOut = fprintf([bp ...
                ' %g / %g = %3.0f%%\n' ...
                '  Estimated %s\n' ...
                '-   Elapsed %s\n' ...
                '---------------------\n' ...
                '= Remaining %s\n'],...
                d.nCompleted,d.nRuns,d.nCompleted/d.nRuns*100, ...
                char(estDuration), ...
                char(sinceStart), ...
                char(remaining) ...
                ) - d.nProgressOut;
        end
    else
        % Final display when finishing
        if(d.nErrors > 0)
            % Show nErrors
            nProgressOut = fprintf([bp ...
                ' %g / %g' ...
                '; # of errors: %g (%2.0f%%).' ...
                ' Elapsed %s\n'], ...
                d.nCompleted,d.nRuns, ...
                d.nErrors,d.nErrors/d.nCompleted*100, ...
                char(sinceStart) ...
                ) - d.nProgressOut;
        else
            nProgressOut = fprintf([bp ...
                ' %g / %g' ...
                ' Elapsed %s\n'], ...
                d.nCompleted,d.nRuns, ...
                char(sinceStart) ...
                ) - d.nProgressOut;
        end
    end
end
function nProgressOut = emulate_progressTextMultiple(d) %#ok<DEFNU>
% parcellfun_progress/emulate_progressTextMultiple Emulate
% progressTextMultiple
%
% This function will call progressTextMultiple with no arguments
% for each worker the completes. Initialize with text manually
    persistent last_nCompleted;
    nProgressOut = 0;
    if(isempty(last_nCompleted))
        last_nCompleted = 0;
    end
    if(d.nCompleted ~= last_nCompleted)
        for i = 1:(d.nCompleted - last_nCompleted);
            progressTextMultiple;
        end
        last_nCompleted = d.nCompleted;
    end   
end
function nProgressOut = emulate_progressText(d) %#ok<DEFNU>
% parcellfun_progress/emulate_progressText Emulate progressText
%
% This function will call progressText with the fraction complete.
% Initialize with text as needed.
    nProgressOut = 0;
    if(d.nCompleted ~= 0)
        progressText(d.nCompleted / d.nRuns);
    end
end
% function nProgressOut = emulate_parfor_progress(d) %#ok<DEFNU>
% % parcellfun_progress/emulate_parfor_progress Emulate parfor_progress
% %
% % This function will initialize, update, and cleanup parfor_progress
%     persistent last_nCompleted;
%     nProgressOut = 1;
%     if(isempty(last_nCompleted))
%         last_nCompleted = 0;
%     end
%     if(~d.nProgressOut)
%         % initialize
%         parfor_progress(d.nRuns);
%         last_nCompleted = 0;
%     end
%     if(d.nCompleted ~= last_nCompleted)
%         for i = 1:(d.nCompleted - last_nCompleted);
%             parfor_progress;
%         end
%         last_nCompleted = d.nCompleted;
%     end
%     if(d.nCompleted == d.nRuns)
%         parfor_progress(0);
%     end
% end
function str = duration_shim(hours,minutes,seconds)
    totalSeconds = hours*3600 + minutes*60 + seconds;
    hour=uint8(floor(totalSeconds/3600.0));
    minutes=uint8(floor(mod((totalSeconds/60.0), 60.0)));
    seconds=uint8(floor(mod(totalSeconds,60.0)));

    str=sprintf('%02d:%02d:%02d',...
        hour,...
        minutes,...
        seconds ...
    );
end
```
