## Supplementary material for "Adaptive-Resolution Multi-Orientation Analysis of Complex Filamentous Network Images": MATLAB code from Github: progressText.m

```
function progressText(fractionDone, text, wtBarName)
%PROGRESSTEXT shows progress of a loop as text on the screen
%
% SYNOPSIS: progressText(fractionDone,text)
%
% INPUT fractionDone: fraction of loop done (0-1)
%		text (opt): {yourTexthere} : XX% done xx:xx:xx remaining
%                   Note: text can be changed for every call
%       popupDisp (opt) :  {waitbar fig name}  (if provided will initialize popup fig waitbar)
% OUTPUT none
%
% EXAMPLE
%   n = 1000;
%   progressText(0,'Test run', 'computation process name') % Create text & waitbar popup
%   for i = 1:n
%       pause(0.01) % Do something important
%       progressText(i/n) % Update text
%   end
%
% REMARKS progressText will set lastwarn to ''
%
% created with MATLAB ver.: 7.4.0.287 (R2007a) on Windows_NT
%
% created by: jdorn based on progressbar.m
% DATE: 29-Jun-2007
% Modified:  Mar 2017 - Andrew R. Jamieson
%
%%%%%%%%%%%%%%%%%%%%%%%%%%%%%%%%%%%%%%%%%%%%%%%%%%%%%%%%%%%%%%%%%%%%%%%%%%%
%
% Copyright (C) 2019, Jaqaman Lab - UT Southwestern, Goldman Lab - Northwestern 
%
% This file is part of AdaptiveResolutionOrientationSpace.
% 
% AdaptiveResolutionOrientationSpace is free software: you can redistribute it and/or modify
% it under the terms of the GNU General Public License as published by
% the Free Software Foundation, either version 3 of the License, or
% (at your option) any later version.
% 
% AdaptiveResolutionOrientationSpace is distributed in the hope that it will be useful,
% but WITHOUT ANY WARRANTY; without even the implied warranty of
% MERCHANTABILITY or FITNESS FOR A PARTICULAR PURPOSE.  See the
% GNU General Public License for more details.
% 
% You should have received a copy of the GNU General Public License
% along with AdaptiveResolutionOrientationSpace.  If not, see <http://www.gnu.org/licenses/>.
% 
% 

persistent starttime lastupdate clearText printText finalText warnText clearWarn progfig logMsg timeMsg popupDisp

% constants
nCharsBase = 27; % change this if changing output format

% Test input
if nargin < 1 || isempty(fractionDone) 
    fractionDone = 0;
end
if nargin < 2 || isempty(text)
    text = '';
else
    text = [text,' : '];
end


if nargin < 3 && (fractionDone == 0) && isempty(starttime)
    popupDisp = false;
end

if  nargin == 3
    popupDisp = true;
end


if fractionDone == 0 || isempty(starttime)
    % set up everything
    
    % Set time of last update to ensure calculation
    lastupdate = clock - 1;
    
    % Task starting time reference
    starttime = clock;
    
    % create fprintf-expression
    printText = sprintf('%s%%2d%%%% done %%s remaining',text);
    initialText = sprintf('%s 0%%%% done xx:xx:xx remaining',text);
    finalText = sprintf('%s100%%%% done %%s elapsed',text);
    % get length of fprintf expression
    nChars = nCharsBase + length(text);
    clearText = repmat('\b',1,nChars);
    
    % print initialText and return
    fprintf(1, initialText);
    
    % empty warning
    lastwarn('');
    warnText = '';
    clearWarn = '';

    if popupDisp


        try
            % Access progfig to see if it exists ('try' will fail if it doesn't)
            dummy = get(progfig,'UserData');
            % If progress bar needs to be reset, close figure and set handle to empty
            if fractiondone == 0
                delete(progfig) % Close progress bar
            end
        catch
            dummy_ = [];
        end

        logMsg = ['Please wait...' text];
        timeMsg = @(t) ['\nEstimated time remaining: ' t];
        progfig = waitbar(0, text, 'Name', wtBarName);        

    end

    return


elseif ~isempty(text)
    % text has been changed. Create fprintfExpressions first, then update
    % clearText
    logMsg = ['Please wait...' text];
    printText = sprintf('%s%%2d%%%% done %%s remaining',text);
    finalText = sprintf('%s100%%%% done %%s elapsed', text);
    fprintfExpression = [clearText clearWarn printText, warnText];
    fprintfExpressionFinal = [clearText, clearWarn, finalText, warnText, '\n'];
    
    nChars = nCharsBase + length(text);
    clearText = repmat('\b',1,nChars);


elseif ~isempty(lastwarn)
    
    % add warnings to the end of the progressText
    % find warning
    w = lastwarn;
    lastwarn('')
    nw = length(w);
    % erase warning
    fprintf(1,repmat('\b',1,11+nw)); %--- is this correct???
    % create new warnText
    w = regexprep(w,['(',char(10),'\s+)'],' - ');
    warnTextNew = sprintf('\n  Warning @%7.3f%%%% done : %s ',100*fractionDone,w);
    warnLength = length(warnTextNew) - 1; % subtract 2 for %%-sign
    warnText = [warnText,warnTextNew];
    
    fprintfExpression = [clearText clearWarn printText, warnText];
    fprintfExpressionFinal = [clearText, clearWarn, finalText, warnText, '\n'];
    
    % prepare cleanWarn for next time
    clearWarn = [clearWarn,repmat('\b',1,warnLength)];
else
    % all is normal. Just generate output
    fprintfExpression = [clearText clearWarn printText, warnText];
    fprintfExpressionFinal = [clearText, clearWarn, finalText, warnText, '\n'];
end

% write progress
percentDone = floor(100*fractionDone);

% get elapsed time
runTime = etime(clock,starttime);


if percentDone == 100 % Task completed
    fprintf(1,fprintfExpressionFinal,convertTime(runTime)); % finish up
    if popupDisp
        waitbar(fractionDone, progfig,  sprintf([logMsg 'Completed!']) );
        delete(progfig);
    end
    clear popupDisp progfig starttime lastupdate clearText printText finalText% Clear persistent vars
    return
end

% only update if significant time has passed
if etime(clock,lastupdate) < 0.3
    return
end

% update
timeLeft = runTime/fractionDone - runTime;
fprintf(1,fprintfExpression,percentDone,convertTime(timeLeft));
if popupDisp
    waitbar(fractionDone, progfig, sprintf([logMsg timeMsg(convertTime(timeLeft))]));
end

%%%%%%%%%%%%%%%%%%%%%%%%%%%%%%%%%%%%%%%%%%%%%%%%%%
% subfcn
function timeString = convertTime(time)

timeStruct = sec2struct(time);
if timeStruct.hour > 99
    timeString = '99:59:59';
else
    timeString = timeStruct.str(1:end-4);
end
```
