## Supplementary material for "Adaptive-Resolution Multi-Orientation Analysis of Complex Filamentous Network Images": MATLAB code from Github: progressTextMultiple.m

```
function [out] = progressTextMultiple(text_, nStep_)
%progressTextMultiple shows progress of a loop as text on the screen. Can handle multiple levels of progress
%
%SYNOPSIS
%   [] = progressTextMultiple(text_, nStep_)
%       Initializes the progressText
%   [] = progressTextMultiple(text_)
%       Changes the optional text and updates progressText
%   [] = progressTextMultiple()
%       Updates progressText
%
%INPUT
%   text        : Text in the progress display or empty
%   nSteps      : Max number of steps until completion
%
%OUTPUT
%   out     : diagnostic information about the persistent variables
%       .level      : The layer / level of prgoress display. Larger the
%                     level, smaller the increase in progress fraction
%       .iStep      : array of progress on each level
%       .nStep      : array of max number of steps needed to complete each
%                     level
%
%WARNINGS
%   Throws a progressTextMultiple:LevelZero warning if the number of steps
%   specified is exceeded and level 0 is reached. Warning is thrown only
%   once. Reset using 'clear progressTextMultiple'
%
%Tae H Kim, July 2015
%
% Copyright (C) 2019, Jaqaman Lab - UT Southwestern, Goldman Lab - Northwestern 
%
% This file is part of AdaptiveResolutionOrientationSpace.
% 
% AdaptiveResolutionOrientationSpace is free software: you can redistribute it and/or modify
% it under the terms of the GNU General Public License as published by
% the Free Software Foundation, either version 3 of the License, or
% (at your option) any later version.
% 
% AdaptiveResolutionOrientationSpace is distributed in the hope that it will be useful,
% but WITHOUT ANY WARRANTY; without even the implied warranty of
% MERCHANTABILITY or FITNESS FOR A PARTICULAR PURPOSE.  See the
% GNU General Public License for more details.
% 
% You should have received a copy of the GNU General Public License
% along with AdaptiveResolutionOrientationSpace.  If not, see <http://www.gnu.org/licenses/>.
% 
% 

%% Initialization
%persistent
persistent text iStep nStep frac weight level warned
if isempty(level)
    level = 0;
end
if isempty(text)
    text = {};
end
if isempty(warned)
    warned = false;
end

%% Input
if nargin == 0 && level ~= 0
    iStep(level) = iStep(level) + 1;
end
if nargin == 1 && level ~= 0
    iStep(level) = iStep(level) + 1;
    text{level} = text_;
end
if nargin == 2
    level = level + 1;
    iStep(level) = 0;
    nStep(level) = nStep_;
    if level == 1
        weight(1, 1) = 1;
    else
        weight(level, 1) = weight(level-1, 1) / nStep(level - 1)*0.9;
    end
    text{level} = text_;
end

if level > 0
    %% Fraction calculation
    frac(level) = iStep(level) / nStep(level);

    %% Progress Display
    if iStep(1) < nStep(1) && nargin ~= 2
        progressText(frac * weight, getFullText(text));
    elseif iStep(1) == nStep(1)
        progressText(1, getFullText(text));
    end

    %% level check
    if iStep(level) == nStep(level)
        level = level - 1;
        frac = frac(1:end-1);
        weight = weight(1:end-1);
        text = text(1:end-1);
        iStep = iStep(1:end-1);
        nStep = nStep(1:end-1);
    end

    %% Output

elseif ~warned
    warned = true;
    warning('progressTextMultiple:LevelZero','Level 0 reached. The progressText may be inaccurate');
end
if nargout > 0
    out.level = level;
    out.iStep = iStep;
    out.nStep = nStep;
end

end

%% Local function
%generates fullText
function [fullText] = getFullText(text)
text = text(~cellfun('isempty',text));
text = cellfun(@(x) [x ': '], text, 'UniformOutput', false);
fullText = [text{:}];
fullText = fullText(1:end-2);
end
```
