## Supplementary material for "Adaptive-Resolution Multi-Orientation Analysis of Complex Filamentous Network Images": MATLAB code from Github: sec2struct.m

```
function [time_strct]=sec2struct(x)
% SEC2STRUCT splits seconds into hours, minutes, seconds and milliseconds 
% INPUT:
%	x -> number of seconds (scalar value, floating point)
% OUTPUT:
%	time_strct -> structure with the following fields
%		.hour
%		.minutes
%		.seconds
%		.milliseconds
%		.str: time as HH:MM:SS:MS string
%		.vec: time as HHMMSSMS string
%
% Author: Ulrich Schmidt, April 2011
%
%
% Copyright (C) 2019, Jaqaman Lab - UT Southwestern, Goldman Lab - Northwestern 
%
% This file is part of AdaptiveResolutionOrientationSpace.
% 
% AdaptiveResolutionOrientationSpace is free software: you can redistribute it and/or modify
% it under the terms of the GNU General Public License as published by
% the Free Software Foundation, either version 3 of the License, or
% (at your option) any later version.
% 
% AdaptiveResolutionOrientationSpace is distributed in the hope that it will be useful,
% but WITHOUT ANY WARRANTY; without even the implied warranty of
% MERCHANTABILITY or FITNESS FOR A PARTICULAR PURPOSE.  See the
% GNU General Public License for more details.
% 
% You should have received a copy of the GNU General Public License
% along with AdaptiveResolutionOrientationSpace.  If not, see <http://www.gnu.org/licenses/>.
% 
% 

time_strct.hour=int8(floor(x/3600.0));
time_strct.minutes=int8(floor(mod((x/60.0), 60.0)));
time_strct.seconds=int8(floor(mod(x,60.0)));
time_strct.milliseconds=int16(mod(x,1.0)*1000);

time_strct.str=sprintf('%02d:%02d:%02d:%03d',...
    time_strct.hour,...
    time_strct.minutes,...
    time_strct.seconds,...
    time_strct.milliseconds...
);

time_strct.vec=sprintf('%02d%02d%02d.%03d',...
    time_strct.hour,...
    time_strct.minutes,...
    time_strct.seconds,...
    time_strct.milliseconds...
);
```
