## Supplementary material for "Adaptive-Resolution Multi-Orientation Analysis of Complex Filamentous Network Images": MATLAB code from Github: sortMatrices.m

```
function [ A_sorted, varargout] = sortMatrices( A, varargin )
%sortMatrices Sort first matrix and rearrange other matrices in the same order
%
% sortMatrices differs from sort in that the indices output address the whole
% matrix rather than the dimension sorted on. Additionally, other matrices
% can be specified which will be sorted in the same order as A.
%
% [sA,rB,I] = sortMatrices(A,B,3,'descend');
% assert(sA == A(I));
% assert(rB == B(I));
%
% INPUT
% A    - reference matrix to sort
% ...  - other matrices to rearrange in the same order as A
% dim  - dimension to sort along, optional (default: 1)
% mode - 'ascend' or 'descend', optional (default: 'ascend')
%
% OUTPUT
% A_sorted - A sorted
% ...      - Other matrices rearranged in the same manner as A
% I        - Linear indices describing rearrangement of A, same size as A
% 
% See also sort
%
% Copyright (C) 2019, Jaqaman Lab - UT Southwestern, Goldman Lab - Northwestern 
%
% This file is part of AdaptiveResolutionOrientationSpace.
% 
% AdaptiveResolutionOrientationSpace is free software: you can redistribute it and/or modify
% it under the terms of the GNU General Public License as published by
% the Free Software Foundation, either version 3 of the License, or
% (at your option) any later version.
% 
% AdaptiveResolutionOrientationSpace is distributed in the hope that it will be useful,
% but WITHOUT ANY WARRANTY; without even the implied warranty of
% MERCHANTABILITY or FITNESS FOR A PARTICULAR PURPOSE.  See the
% GNU General Public License for more details.
% 
% You should have received a copy of the GNU General Public License
% along with AdaptiveResolutionOrientationSpace.  If not, see <http://www.gnu.org/licenses/>.
% 
% 

% Mark Kittisopikul, 2015
% Jaqaman Lab
% UT Southwestern

defaultMode = 'ascend';
nMatrices = nargin;

% Collect information on A
size_A = size(A);
ndims_A = ndims(A);

% See if the mode was specified
extraParamStart = nMatrices;
if(nMatrices > 1 ...
        && ischar(varargin{end}) ... 
        )
%         && any(strcmp(varargin{end},{'ascend','descend'})) )...
    for nMatrices=nMatrices-1:-1:1
        if(~ischar(varargin{nMatrices}))
            nMatrices = nMatrices+1;
            break;
        end
        switch(varargin{nMatrices})
            case 'ascend'
                break;
            case 'descend'
                break;
        end
    end
    mode = varargin{nMatrices};
    extraParamStart = nMatrices+1;
%     nMatrices = nMatrices - 1;
else
    mode = defaultMode;
end

% Check if the dimension was specified
if(nMatrices > 1 ...
        && isscalar(varargin{nMatrices-1}))
    dim = varargin{nMatrices-1};
    nMatrices = nMatrices - 1;
else
    dim = find(size_A ~= 1,1,'first');
    % If no dimension was specified and the last argument is not a valid
    % mode, then treat it as a matrix
    if(~any(strcmp(mode,{'ascend','descend'})))
        nMatrices = nMatrices + 1;
        mode = defaultMode;
    end
end

% Use the builtin sort
[A_sorted,ind ] = sort(A,dim,mode,varargin{extraParamStart:end});


% Convert dimension specific indices to linear indices.
% sub2ind would do this, but using bsxfun is more efficient.
% An alternate strategy is to shift the dimension of interest to the first
% dimension
k = [1 cumprod(size_A(1:end-1))];
sizm1 = size_A-1;
% I = ones(siz);
I = 1;
for i = 1:ndims_A
    if(i ~= dim)
        I = bsxfun(@plus,I,shiftdim((0:sizm1(i))',-(i-1))*k(i));
    else
        I = bsxfun(@plus,I,(ind-1)*k(i));
    end
end

% Output the rearranged matrices
varargout = cell(1,length(varargin));
for o=2:min(nargout,nMatrices)
    assert(ndims_A == ndims(varargin{o-1}) && ...
        all(size(varargin{o-1}) == size_A), ...
        'Matrix arguments must be the same size');
    varargout{o-1} = varargin{o-1}(I);
end

% Also output the linear indices if requested
if(nargout > nMatrices)
    varargout{nargout-1} = I;
end


end
```
