## Supplementary material for "Adaptive-Resolution Multi-Orientation Analysis of Complex Filamentous Network Images": MATLAB code from Github: demo.m

```
    demo = zeros(256);
    demo(128,:) = 1;
    demo = max(imgaussfilt(demo,2),imgaussfilt(eye(256),2));
    demo = imnoise(mat2gray(demo),'gaussian',0.1,0.01);
    [res,theta,nms] = steerableAdaptiveResolutionOrientationSpaceDetector(demo);
    figure; imshow(nms,[]);
    orientationSpace.rainbowOrientationQuivers(theta,res,hsv(32));
    xlim(128+[-10 10]);
    ylim(128+[-10 10]);
%
% Copyright (C) 2019, Jaqaman Lab - UT Southwestern, Goldman Lab - Northwestern 
%
% This file is part of AdaptiveResolutionOrientationSpace.
% 
% AdaptiveResolutionOrientationSpace is free software: you can redistribute it and/or modify
% it under the terms of the GNU General Public License as published by
% the Free Software Foundation, either version 3 of the License, or
% (at your option) any later version.
% 
% AdaptiveResolutionOrientationSpace is distributed in the hope that it will be useful,
% but WITHOUT ANY WARRANTY; without even the implied warranty of
% MERCHANTABILITY or FITNESS FOR A PARTICULAR PURPOSE.  See the
% GNU General Public License for more details.
% 
% You should have received a copy of the GNU General Public License
% along with AdaptiveResolutionOrientationSpace.  If not, see <http://www.gnu.org/licenses/>.
% 
% 
```
