## Supplementary material for "Adaptive-Resolution Multi-Orientation Analysis of Complex Filamentous Network Images": MATLAB code from Github: colorcet.m

```
% COLORCET - Perceptually uniform color maps
%
% Usage 1: Generate a colour map and apply it to the current figure.
%  >> colorcet(name);
%  >> colorcet(name, keyword, arg, ...);
%
% Usage 2: Return a colour map and its description.
%  >> [map, descriptorname, description] = colorcet(name);
%
% Usage 3: Get a list of all possible colour maps.
%  >> colorcet
%
% Arguments:
%       name - A string label indicating the colour map to be generated.
%
%     names:  'L1' - 'L19'  for linear maps
%             'D1' - 'D13'  for diverging maps
%             'C1' - 'C9'   for cyclic maps
%             'R1' - 'R3'   for rainbow maps
%             'I1' - 'I3'   for isoluminant maps
%                                                    
%     maps for the red-green colour blind (Protanopia/Deuteranopia)
%             'CBL1' - 'CBL4' 
%             'CBD1' - 'CBD2' 
%             'CBC1' - 'CBC2' 
%     maps for the blue-yellow colour blind (Tritanopia)
%             'CBTL1' - 'CBTL4' 
%             'CBTD1' 
%             'CBTC1' - 'CBTC2' 
%
% Keyword - argument options
%           'N' - Number of values in the colour map. Defaults to 256.
%       'shift' - Fraction of the colour map length N that the colour map is
%                 to be cyclically rotated, may be negative.  (Should only be
%                 applied to cyclic colour maps!). Defaults to 0.
%     'reverse' - If set to 1/true reverses the colour map. Defaults to 0.
%
%
% Returns:
%            map - Nx3 rgb colour map
% descriptorname - A string giving a systematic descriptor name for the colour map.
%    description - A string giving a brief description of the colour map.
%
%
% Some colour maps have alternate names for convenience and readability.
% Run colorcet with no arguments to get a full list of names
% For example:
%   >> map = colorcet('L3');  or map = colorcet('heat');  will produce a linear heat map.
%
% Examples of use:
% Rotate a cyclic colour map by half a cycle
%   >> map = colorcet('C2', 'shift', 0.5);
%
% Generate a 64 level reversed blue-green-yellow map
%   >> map = colorcet('L9', 'N', 64, 'reverse', true);
%
% Load up the SINERAMP test image and render it with MATLAB's hot map
%   >> figure; imagesc(sineramp); colormap(hot);
%
% Now repeat using the colorcet perceptually uniform heat map
%   >> figure; imagesc(sineramp);  colorcet('heat');
%
% Notice the difference! No preceptual dead spots across the top of the
% test image and no false features across the bottom.
%
% Some colour maps likely to be of interest:
%   L1/GREY/GRAY  - Standard grey colour map 
%   L3/HEAT/FIRE  - A good heat colour map with no perceptual dead spots.
%   L16/KBGYW     - Black-Blue-Green-Yellow-White, quite nice 
%   D1/COOLWARM   - Standard blue-white-red diverging colour map with
%                   end colours matched for lightness and chroma.
%   D1A/BWRA      - Similar to D1 but with greater chroma and darker 
%                   end colours
%   D13/BWG       - Blue-White-Green diverging map, easy on the eye
%   R1/RAINBOW    - The least worst rainbow colour map I could devise,
%                   actually it's not bad.
%   R3/RAINBOW3   - Rainbow map that can also be used as a diverging map
%                   with yellow as the reference centre colour.
%   C2/PHASE4     - Good cyclic map with four distinct colours allowing
%                   four orientations or phase angles to be visualised.
%   C4/PHASE2     - A blue - white - red cyclic map for when you are only
%                   interested in phase being +ve or -ve.
%
% If you are colour blind...
% The following maps have been constructed to lie within a 2D model of
% protanopic/deuteranopic colour space. Hopefully by working within this
% colour space people who are colour blind will be able to share a common
% perceptual interpretation of data with those who have normal colour vision.
%   CBL1 to CBL3 - Linear maps for protanopes and deuteranopes
%   CBD1         - Diverging maps for protanopes and deuteranopes
%   CBC1 to CBC2 - Cyclic maps for protanopes and deuteranopes
%  There are also maps for tritanopes
%
% The colour map arrays in this function have been generated by the function CMAP.
%
% For more information about the design of these colour maps see:
% Peter Kovesi. Good Colour Maps: How to Design Them.
% https://arxiv.org/abs/1509.03700
% See also:
% https://peterkovesi.com/projects/colourmaps
%
% See also: CMAP, MAKECOLORCET, SINERAMP, SHOW

% These colour maps are released under the Creative Commons BY License.
% A summary of the conditions can be found at
% https://creativecommons.org/licenses/by/4.0/
% Basically, you are free to use these colour maps in anyway you wish
% as long as you give appropriate credit.
%
% Reference:
% Peter Kovesi. Good Colour Maps: How to Design Them.
% arXiv:1509.03700 [cs.GR] 2015
% https://arxiv.org/abs/1509.03700

% Copyright (c) 2014-2018 Peter Kovesi
% Centre for Exploration Targeting
% The University of Western Australia
% peter.kovesi at uwa edu au
%
% Permission is hereby granted, free of charge, to any person obtaining a copy
% of this software and associated documentation files (the "Software"), to deal
% in the Software without restriction, subject to the following conditions:
%
% The above copyright notice and this permission notice shall be included in
% all copies or substantial portions of the Software.
%
% The Software is provided "as is", without warranty of any kind.

% Generated by MAKECOLORCET 30-Apr-2018

function varargout = colorcet(varargin)

[mapname, reverse, shift, N] = parseinputs(varargin{:});

mapnames = {
{ 'L1'  'GREY'  'GRAY' }
{ 'L2'  'REDUCEDGREY' }
{ 'L3'  'KRYW'  'HEAT'  'FIRE' }
{ 'L4'  'KRY'  'YELLOWHEAT' }
{ 'L5'  'KGY' }
{ 'L6'  'KBC' }
{ 'L7'  'BMW' }
{ 'L8'  'BMY' }
{ 'L9'  'BGYW' }
{ 'L10'  'GEOGRAPHIC' }
{ 'L11'  'GEOGRAPHIC2' }
{ 'L12'  'DEPTH' }
{ 'L13'  'REDTERNARY' }
{ 'L14'  'GREENTERNARY' }
{ 'L15'  'BLUETERNARY' }
{ 'L16'  'KBGYW' }
{ 'L17'  'KBGYW' }
{ 'L18'  'WORB' }
{ 'L19'  'WCMR' }
{ 'D1'  'BWR'  'COOLWARM' }
{ 'D1A'  'BWRA' }
{ 'D2'  'GWV' }
{ 'D3'  'GWR' }
{ 'D4'  'BKR' }
{ 'D5'  'GKR' }
{ 'D6'  'BKY' }
{ 'D7'  'BJY'  'DIVBJY' }
{ 'D8'  'BJR' }
{ 'D9' }
{ 'D10' }
{ 'D11' }
{ 'D12' }
{ 'D13'  'BWG' }
{ 'C1' }
{ 'C2'  'PHASE4' }
{ 'C3' }
{ 'C4'  'PHASE2' }
{ 'C5'  'CYCLICGREY' }
{ 'C6' }
{ 'C7' }
{ 'C8' }
{ 'C9' }
{ 'R1'  'RAINBOW' }
{ 'R2'  'RAINBOW2' }
{ 'R3'  'RAINBOW3' }
{ 'I1' }
{ 'I2' }
{ 'I3' }
{ 'CBL1' }
{ 'CBL2' }
{ 'CBL3' }
{ 'CBL4' }
{ 'CBD1' }
{ 'CBD2' }
{ 'CBC1' }
{ 'CBC2' }
{ 'CBTL1' }
{ 'CBTL2' }
{ 'CBTL3' }
{ 'CBTL4' }
{ 'CBTD1' }
{ 'CBTC1' }
{ 'CBTC2' }
};

spaces = '                                                           ';

if ~exist('mapname', 'var') || isempty(mapname)

fprintf('\nName(s):         Description:\n');
fprintf('------------------------------------------------------\n');

for n = 1:length(mapnames)
    [map, descriptorname, description] = colorcet(mapnames{n}{1});

    len = 0;
    for m = 1:length(mapnames{n})
        len = len + fprintf('%s', upper(mapnames{n}{m}));
        if m < length(mapnames{n})
            len = len+fprintf('/');
        end
    end

    fprintf('%s', spaces(1:16-len));
    fprintf(' %s \n', description);
end

return
end

switch upper(mapname) 
case { 'L1'  'GREY'  'GRAY' }
descriptorname = 'linear_grey_0-100_c0_n256';
description = 'Grey scale';
map = [0.000000 0.000000 0.000000
       0.005561 0.005563 0.005563
       0.011212 0.011219 0.011217
       0.016877 0.016885 0.016883
       0.022438 0.022448 0.022445
       0.027998 0.028011 0.028008
       0.033540 0.033554 0.033551
       0.039316 0.039333 0.039329
       0.044700 0.044719 0.044714
       0.049695 0.049713 0.049709
       0.054322 0.054343 0.054338
       0.058797 0.058821 0.058815
       0.063041 0.063062 0.063057
       0.066996 0.067019 0.067014
       0.070827 0.070853 0.070846
       0.074487 0.074511 0.074505
       0.077967 0.077992 0.077986
       0.081449 0.081477 0.081470
       0.084728 0.084752 0.084746
       0.087913 0.087939 0.087933
       0.091003 0.091031 0.091024
       0.094004 0.094034 0.094027
       0.096981 0.097012 0.097005
       0.100036 0.100065 0.100058
       0.103113 0.103144 0.103136
       0.106180 0.106212 0.106204
       0.109243 0.109276 0.109268
       0.112317 0.112349 0.112341
       0.115378 0.115412 0.115404
       0.118518 0.118550 0.118542
       0.121566 0.121599 0.121591
       0.124708 0.124743 0.124734
       0.127866 0.127902 0.127893
       0.131020 0.131054 0.131046
       0.134177 0.134214 0.134204
       0.137342 0.137380 0.137371
       0.140512 0.140551 0.140541
       0.143696 0.143733 0.143724
       0.146874 0.146914 0.146904
       0.150110 0.150147 0.150138
       0.153293 0.153333 0.153323
       0.156556 0.156595 0.156585
       0.159750 0.159790 0.159780
       0.163036 0.163078 0.163068
       0.166276 0.166320 0.166309
       0.169568 0.169609 0.169599
       0.172813 0.172857 0.172846
       0.176080 0.176123 0.176113
       0.179399 0.179443 0.179432
       0.182689 0.182734 0.182723
       0.186036 0.186079 0.186069
       0.189342 0.189388 0.189376
       0.192671 0.192718 0.192707
       0.196020 0.196069 0.196057
       0.199344 0.199392 0.199380
       0.202714 0.202763 0.202751
       0.206110 0.206160 0.206148
       0.209462 0.209512 0.209500
       0.212840 0.212890 0.212878
       0.216264 0.216315 0.216303
       0.219652 0.219705 0.219692
       0.223068 0.223120 0.223107
       0.226487 0.226539 0.226526
       0.229881 0.229935 0.229922
       0.233319 0.233373 0.233360
       0.236797 0.236853 0.236839
       0.240217 0.240271 0.240258
       0.243676 0.243733 0.243719
       0.247158 0.247216 0.247202
       0.250627 0.250687 0.250672
       0.254124 0.254183 0.254168
       0.257603 0.257661 0.257647
       0.261107 0.261167 0.261152
       0.264609 0.264668 0.264653
       0.268119 0.268182 0.268166
       0.271637 0.271697 0.271683
       0.275184 0.275249 0.275233
       0.278724 0.278788 0.278772
       0.282274 0.282339 0.282323
       0.285801 0.285865 0.285849
       0.289381 0.289445 0.289430
       0.292935 0.293001 0.292984
       0.296520 0.296587 0.296570
       0.300095 0.300161 0.300145
       0.303679 0.303747 0.303730
       0.307277 0.307344 0.307327
       0.310897 0.310965 0.310948
       0.314490 0.314559 0.314542
       0.318115 0.318185 0.318168
       0.321735 0.321807 0.321789
       0.325361 0.325433 0.325415
       0.329019 0.329093 0.329075
       0.332668 0.332741 0.332723
       0.336301 0.336375 0.336357
       0.339973 0.340047 0.340029
       0.343630 0.343703 0.343685
       0.347314 0.347390 0.347371
       0.350992 0.351068 0.351049
       0.354679 0.354758 0.354739
       0.358360 0.358436 0.358417
       0.362066 0.362144 0.362125
       0.365769 0.365850 0.365830
       0.369478 0.369557 0.369538
       0.373195 0.373277 0.373256
       0.376930 0.377010 0.376990
       0.380661 0.380744 0.380724
       0.384389 0.384472 0.384451
       0.388136 0.388220 0.388200
       0.391889 0.391973 0.391952
       0.395652 0.395737 0.395716
       0.399400 0.399485 0.399464
       0.403182 0.403270 0.403248
       0.406955 0.407042 0.407020
       0.410744 0.410831 0.410810
       0.414525 0.414614 0.414592
       0.418314 0.418403 0.418381
       0.422113 0.422203 0.422181
       0.425924 0.426014 0.425991
       0.429741 0.429833 0.429810
       0.433552 0.433644 0.433621
       0.437391 0.437485 0.437462
       0.441212 0.441307 0.441284
       0.445054 0.445147 0.445124
       0.448900 0.448995 0.448972
       0.452749 0.452844 0.452820
       0.456594 0.456691 0.456667
       0.460449 0.460546 0.460522
       0.464332 0.464430 0.464406
       0.468190 0.468289 0.468265
       0.472077 0.472176 0.472152
       0.475957 0.476057 0.476032
       0.479848 0.479948 0.479924
       0.483732 0.483834 0.483809
       0.487644 0.487747 0.487722
       0.491552 0.491655 0.491629
       0.495453 0.495556 0.495531
       0.499365 0.499469 0.499443
       0.503286 0.503392 0.503366
       0.507220 0.507327 0.507301
       0.511151 0.511258 0.511231
       0.515092 0.515198 0.515172
       0.519043 0.519150 0.519124
       0.522990 0.523100 0.523073
       0.526948 0.527057 0.527030
       0.530903 0.531013 0.530986
       0.534868 0.534979 0.534951
       0.538843 0.538953 0.538926
       0.542819 0.542932 0.542904
       0.546800 0.546913 0.546885
       0.550785 0.550899 0.550871
       0.554777 0.554891 0.554863
       0.558768 0.558885 0.558856
       0.562782 0.562898 0.562870
       0.566788 0.566906 0.566877
       0.570793 0.570912 0.570883
       0.574810 0.574930 0.574900
       0.578839 0.578960 0.578930
       0.582863 0.582984 0.582954
       0.586898 0.587019 0.586989
       0.590937 0.591060 0.591030
       0.594982 0.595105 0.595074
       0.599033 0.599158 0.599127
       0.603086 0.603211 0.603180
       0.607150 0.607276 0.607245
       0.611211 0.611337 0.611306
       0.615273 0.615400 0.615369
       0.619357 0.619483 0.619452
       0.623434 0.623563 0.623531
       0.627513 0.627642 0.627610
       0.631602 0.631732 0.631700
       0.635693 0.635825 0.635793
       0.639795 0.639927 0.639895
       0.643892 0.644024 0.643991
       0.647999 0.648132 0.648100
       0.652118 0.652250 0.652218
       0.656238 0.656372 0.656339
       0.660354 0.660488 0.660455
       0.664479 0.664615 0.664581
       0.668616 0.668753 0.668719
       0.672755 0.672892 0.672858
       0.676895 0.677034 0.677000
       0.681032 0.681171 0.681136
       0.685189 0.685330 0.685295
       0.689347 0.689487 0.689453
       0.693501 0.693641 0.693607
       0.697664 0.697806 0.697771
       0.701831 0.701974 0.701939
       0.706011 0.706155 0.706120
       0.710188 0.710333 0.710297
       0.714366 0.714511 0.714476
       0.718555 0.718701 0.718665
       0.722745 0.722892 0.722856
       0.726944 0.727092 0.727055
       0.731143 0.731292 0.731255
       0.735352 0.735500 0.735464
       0.739567 0.739717 0.739680
       0.743776 0.743927 0.743890
       0.747997 0.748149 0.748111
       0.752216 0.752368 0.752330
       0.756445 0.756598 0.756560
       0.760678 0.760832 0.760794
       0.764916 0.765071 0.765033
       0.769151 0.769307 0.769268
       0.773396 0.773552 0.773514
       0.777652 0.777808 0.777770
       0.781905 0.782063 0.782024
       0.786164 0.786323 0.786284
       0.790422 0.790582 0.790542
       0.794691 0.794850 0.794811
       0.798963 0.799125 0.799085
       0.803235 0.803397 0.803357
       0.807517 0.807679 0.807639
       0.811800 0.811964 0.811923
       0.816091 0.816255 0.816215
       0.820377 0.820543 0.820502
       0.824674 0.824841 0.824800
       0.828981 0.829148 0.829107
       0.833283 0.833452 0.833410
       0.837595 0.837763 0.837722
       0.841907 0.842077 0.842035
       0.846220 0.846390 0.846348
       0.850542 0.850712 0.850670
       0.854871 0.855044 0.855002
       0.859199 0.859372 0.859330
       0.863537 0.863711 0.863668
       0.867874 0.868050 0.868007
       0.872217 0.872391 0.872348
       0.876557 0.876733 0.876689
       0.880910 0.881088 0.881044
       0.885265 0.885443 0.885399
       0.889623 0.889801 0.889757
       0.893986 0.894167 0.894122
       0.898350 0.898531 0.898486
       0.902724 0.902906 0.902861
       0.907099 0.907281 0.907236
       0.911475 0.911658 0.911613
       0.915854 0.916038 0.915992
       0.920242 0.920425 0.920380
       0.924631 0.924816 0.924770
       0.929025 0.929212 0.929166
       0.933422 0.933609 0.933563
       0.937828 0.938016 0.937970
       0.942236 0.942425 0.942379
       0.946642 0.946833 0.946786
       0.951059 0.951248 0.951202
       0.955477 0.955669 0.955621
       0.959896 0.960089 0.960041
       0.964319 0.964511 0.964464
       0.968750 0.968943 0.968896
       0.973184 0.973378 0.973330
       0.977620 0.977815 0.977767
       0.982064 0.982260 0.982211
       0.986507 0.986705 0.986656
       0.990954 0.991152 0.991103
       0.995403 0.995602 0.995553
       0.999862 1.000000 1.000000];

case { 'L2'  'REDUCEDGREY' }
descriptorname = 'linear_grey_10-95_c0_n256';
description = 'Grey scale with slightly reduced contrast to avoid display saturation problems';
map = [0.107674 0.107703 0.107695
       0.110324 0.110354 0.110347
       0.112948 0.112981 0.112973
       0.115542 0.115577 0.115568
       0.118203 0.118237 0.118228
       0.120794 0.120827 0.120819
       0.123442 0.123477 0.123468
       0.126152 0.126185 0.126177
       0.128786 0.128819 0.128811
       0.131486 0.131521 0.131512
       0.134177 0.134214 0.134204
       0.136844 0.136882 0.136873
       0.139515 0.139551 0.139542
       0.142258 0.142295 0.142286
       0.144991 0.145030 0.145020
       0.147699 0.147736 0.147727
       0.150420 0.150458 0.150448
       0.153132 0.153171 0.153162
       0.155910 0.155951 0.155941
       0.158658 0.158698 0.158688
       0.161424 0.161465 0.161455
       0.164185 0.164225 0.164215
       0.166949 0.166992 0.166981
       0.169727 0.169768 0.169758
       0.172485 0.172528 0.172517
       0.175286 0.175327 0.175317
       0.178109 0.178153 0.178142
       0.180871 0.180915 0.180905
       0.183690 0.183736 0.183725
       0.186516 0.186561 0.186549
       0.189342 0.189388 0.189376
       0.192168 0.192215 0.192203
       0.195017 0.195063 0.195052
       0.197849 0.197896 0.197884
       0.200684 0.200733 0.200721
       0.203572 0.203622 0.203610
       0.206452 0.206501 0.206489
       0.209293 0.209343 0.209331
       0.212187 0.212236 0.212224
       0.215043 0.215094 0.215081
       0.217948 0.218000 0.217987
       0.220859 0.220911 0.220898
       0.223735 0.223786 0.223774
       0.226655 0.226708 0.226695
       0.229544 0.229596 0.229583
       0.232479 0.232532 0.232519
       0.235416 0.235471 0.235457
       0.238324 0.238378 0.238364
       0.241270 0.241328 0.241313
       0.244194 0.244251 0.244237
       0.247158 0.247216 0.247202
       0.250094 0.250152 0.250138
       0.253075 0.253134 0.253119
       0.256026 0.256084 0.256069
       0.259018 0.259079 0.259064
       0.261983 0.262043 0.262028
       0.264955 0.265016 0.265001
       0.267938 0.268000 0.267985
       0.270954 0.271014 0.270999
       0.273949 0.274010 0.273995
       0.276954 0.277016 0.277001
       0.279957 0.280019 0.280004
       0.282981 0.283044 0.283029
       0.285979 0.286043 0.286027
       0.289025 0.289090 0.289074
       0.292051 0.292116 0.292100
       0.295078 0.295144 0.295128
       0.298117 0.298184 0.298167
       0.301164 0.301231 0.301215
       0.304221 0.304290 0.304273
       0.307277 0.307344 0.307327
       0.310358 0.310427 0.310410
       0.313418 0.313488 0.313471
       0.316492 0.316561 0.316544
       0.319567 0.319639 0.319621
       0.322656 0.322728 0.322710
       0.325724 0.325796 0.325778
       0.328835 0.328908 0.328890
       0.331927 0.332002 0.331984
       0.335043 0.335117 0.335098
       0.338129 0.338203 0.338185
       0.341251 0.341326 0.341308
       0.344358 0.344435 0.344416
       0.347497 0.347572 0.347554
       0.350626 0.350703 0.350684
       0.353742 0.353819 0.353800
       0.356893 0.356972 0.356952
       0.360024 0.360101 0.360082
       0.363168 0.363247 0.363228
       0.366331 0.366411 0.366391
       0.369478 0.369557 0.369538
       0.372634 0.372715 0.372695
       0.375804 0.375887 0.375867
       0.378974 0.379056 0.379035
       0.382143 0.382226 0.382206
       0.385322 0.385405 0.385384
       0.388515 0.388600 0.388579
       0.391704 0.391787 0.391767
       0.394893 0.394979 0.394958
       0.398096 0.398181 0.398160
       0.401298 0.401381 0.401361
       0.404494 0.404580 0.404559
       0.407710 0.407797 0.407776
       0.410931 0.411017 0.410996
       0.414146 0.414235 0.414213
       0.417377 0.417466 0.417444
       0.420595 0.420685 0.420663
       0.423828 0.423919 0.423897
       0.427077 0.427169 0.427146
       0.430314 0.430406 0.430384
       0.433552 0.433644 0.433621
       0.436813 0.436907 0.436884
       0.440066 0.440159 0.440136
       0.443327 0.443421 0.443398
       0.446588 0.446683 0.446659
       0.449857 0.449952 0.449928
       0.453129 0.453223 0.453200
       0.456401 0.456498 0.456474
       0.459685 0.459781 0.459757
       0.462959 0.463057 0.463033
       0.466255 0.466355 0.466330
       0.469559 0.469658 0.469633
       0.472842 0.472941 0.472917
       0.476150 0.476250 0.476225
       0.479457 0.479559 0.479534
       0.482762 0.482863 0.482838
       0.486073 0.486176 0.486151
       0.489388 0.489490 0.489465
       0.492709 0.492812 0.492786
       0.496035 0.496138 0.496113
       0.499365 0.499469 0.499443
       0.502700 0.502804 0.502778
       0.506035 0.506141 0.506115
       0.509384 0.509491 0.509464
       0.512727 0.512835 0.512809
       0.516071 0.516179 0.516153
       0.519433 0.519540 0.519513
       0.522790 0.522901 0.522874
       0.526154 0.526265 0.526238
       0.529520 0.529630 0.529603
       0.532887 0.532996 0.532969
       0.536259 0.536372 0.536344
       0.539629 0.539741 0.539713
       0.543018 0.543130 0.543102
       0.546405 0.546518 0.546490
       0.549788 0.549902 0.549874
       0.553174 0.553288 0.553260
       0.556574 0.556691 0.556662
       0.559977 0.560094 0.560065
       0.563383 0.563500 0.563471
       0.566788 0.566906 0.566877
       0.570190 0.570308 0.570279
       0.573610 0.573727 0.573699
       0.577030 0.577148 0.577119
       0.580451 0.580570 0.580541
       0.583873 0.583994 0.583964
       0.587300 0.587422 0.587392
       0.590734 0.590857 0.590827
       0.594173 0.594296 0.594266
       0.597614 0.597736 0.597706
       0.601056 0.601180 0.601149
       0.604509 0.604633 0.604603
       0.607962 0.608087 0.608056
       0.611413 0.611539 0.611508
       0.614865 0.614992 0.614961
       0.618334 0.618463 0.618431
       0.621792 0.621919 0.621888
       0.625265 0.625394 0.625362
       0.628741 0.628869 0.628838
       0.632216 0.632346 0.632314
       0.635693 0.635825 0.635793
       0.639178 0.639309 0.639277
       0.642665 0.642796 0.642764
       0.646152 0.646286 0.646253
       0.649651 0.649784 0.649751
       0.653139 0.653274 0.653241
       0.656649 0.656783 0.656750
       0.660149 0.660283 0.660250
       0.663662 0.663797 0.663763
       0.667166 0.667304 0.667270
       0.670686 0.670822 0.670789
       0.674198 0.674336 0.674302
       0.677724 0.677862 0.677828
       0.681239 0.681378 0.681344
       0.684770 0.684911 0.684877
       0.688305 0.688446 0.688412
       0.691840 0.691982 0.691947
       0.695376 0.695519 0.695484
       0.698913 0.699056 0.699021
       0.702460 0.702605 0.702569
       0.706011 0.706155 0.706120
       0.709563 0.709707 0.709671
       0.713112 0.713259 0.713223
       0.716676 0.716823 0.716787
       0.720233 0.720378 0.720343
       0.723795 0.723943 0.723907
       0.727364 0.727511 0.727475
       0.730934 0.731082 0.731045
       0.734512 0.734662 0.734625
       0.738084 0.738235 0.738197
       0.741665 0.741817 0.741779
       0.745250 0.745401 0.745364
       0.748837 0.748988 0.748950
       0.752426 0.752577 0.752540
       0.756022 0.756175 0.756138
       0.759614 0.759769 0.759731
       0.763216 0.763370 0.763332
       0.766819 0.766974 0.766936
       0.770427 0.770584 0.770545
       0.774033 0.774189 0.774151
       0.777652 0.777808 0.777770
       0.781265 0.781424 0.781385
       0.784885 0.785044 0.785005
       0.788505 0.788666 0.788626
       0.792129 0.792289 0.792250
       0.795755 0.795916 0.795876
       0.799392 0.799554 0.799514
       0.803020 0.803183 0.803143
       0.806663 0.806825 0.806785
       0.810297 0.810460 0.810420
       0.813941 0.814105 0.814065
       0.817588 0.817752 0.817712
       0.821239 0.821405 0.821364
       0.824889 0.825056 0.825015
       0.828548 0.828716 0.828675
       0.832208 0.832374 0.832333
       0.835867 0.836036 0.835994
       0.839535 0.839705 0.839663
       0.843198 0.843367 0.843326
       0.846867 0.847038 0.846996
       0.850542 0.850712 0.850670
       0.854219 0.854392 0.854349
       0.857898 0.858071 0.858028
       0.861583 0.861758 0.861715
       0.865266 0.865440 0.865397
       0.868961 0.869134 0.869091
       0.872650 0.872824 0.872781
       0.876341 0.876516 0.876473
       0.880038 0.880215 0.880172
       0.883743 0.883921 0.883877
       0.887441 0.887619 0.887576
       0.891149 0.891328 0.891284
       0.894862 0.895041 0.894997
       0.898569 0.898750 0.898706
       0.902286 0.902467 0.902423
       0.906005 0.906186 0.906142
       0.909724 0.909908 0.909863
       0.913449 0.913632 0.913587
       0.917176 0.917361 0.917315
       0.920898 0.921084 0.921038
       0.924631 0.924816 0.924770
       0.928365 0.928551 0.928506
       0.932103 0.932290 0.932244
       0.935848 0.936036 0.935990
       0.939591 0.939780 0.939733
       0.943337 0.943525 0.943479];

case { 'L3'  'KRYW'  'HEAT'  'FIRE' }
descriptorname = 'linear_kryw_0-100_c71_n256';
description = 'Black-Red-Yellow-White heat colour map';
map = [0.000000 0.000000 0.000000
       0.027065 0.000021 0.000000
       0.052054 0.000075 0.000000
       0.071511 0.000139 0.000000
       0.087420 0.000209 0.000000
       0.101087 0.000281 0.000000
       0.113365 0.000356 0.000000
       0.124393 0.000431 0.000000
       0.134632 0.000508 0.000000
       0.144115 0.000586 0.000000
       0.152922 0.000703 0.000000
       0.160732 0.001343 0.000000
       0.168715 0.001452 0.000000
       0.176567 0.001241 0.000000
       0.183636 0.001534 0.000000
       0.190517 0.001751 0.000000
       0.197510 0.001515 0.000000
       0.204009 0.001525 0.000000
       0.209945 0.001964 0.000000
       0.216048 0.002031 0.000000
       0.222150 0.001756 0.000000
       0.228083 0.001546 0.000019
       0.233784 0.001632 0.000035
       0.239550 0.001719 0.000033
       0.245307 0.001810 0.000019
       0.251127 0.001904 0.000019
       0.256937 0.002001 0.000035
       0.262776 0.002102 0.000033
       0.268644 0.002205 0.000020
       0.274511 0.002312 0.000022
       0.280406 0.002423 0.000036
       0.286329 0.002536 0.000030
       0.292287 0.002653 0.000020
       0.298244 0.002775 0.000028
       0.304234 0.002900 0.000036
       0.310256 0.003028 0.000023
       0.316279 0.003160 0.000013
       0.322319 0.003297 0.000033
       0.328377 0.003438 0.000033
       0.334471 0.003582 0.000021
       0.340565 0.003731 0.000024
       0.346684 0.003885 0.000035
       0.352826 0.004042 0.000024
       0.358972 0.004203 0.000011
       0.365151 0.004371 0.000033
       0.371339 0.004542 0.000030
       0.377560 0.004717 0.000015
       0.383791 0.004899 0.000028
       0.390028 0.005085 0.000033
       0.396300 0.005275 0.000019
       0.402576 0.005472 0.000023
       0.408878 0.005674 0.000033
       0.415194 0.005880 0.000022
       0.421522 0.006092 0.000018
       0.427883 0.006312 0.000033
       0.434239 0.006535 0.000022
       0.440621 0.006765 0.000016
       0.447022 0.007002 0.000032
       0.453436 0.007244 0.000021
       0.459867 0.007493 0.000016
       0.466313 0.007750 0.000031
       0.472773 0.008011 0.000019
       0.479256 0.008279 0.000019
       0.485743 0.008555 0.000030
       0.492254 0.008839 0.000014
       0.498781 0.009136 0.000023
       0.505310 0.009437 0.000028
       0.511866 0.009737 0.000006
       0.518439 0.010039 0.000026
       0.525014 0.010354 0.000024
       0.531617 0.010689 0.000016
       0.538250 0.011031 0.000027
       0.544886 0.011393 0.000016
       0.551535 0.011789 0.000023
       0.558184 0.012159 0.000025
       0.564854 0.012508 0.000015
       0.571536 0.012881 0.000025
       0.578227 0.013283 0.000016
       0.584943 0.013701 0.000023
       0.591664 0.014122 0.000023
       0.598391 0.014551 0.000019
       0.605140 0.014994 0.000024
       0.611901 0.015450 0.000014
       0.618683 0.015920 0.000022
       0.625462 0.016401 0.000016
       0.632262 0.016897 0.000021
       0.639073 0.017407 0.000020
       0.645894 0.017931 0.000021
       0.652729 0.018471 0.000021
       0.659582 0.019026 0.000021
       0.666438 0.019598 0.000021
       0.673315 0.020187 0.000020
       0.680192 0.020793 0.000020
       0.687093 0.021418 0.000020
       0.693995 0.022062 0.000019
       0.700918 0.022727 0.000020
       0.707844 0.023412 0.000018
       0.714785 0.024121 0.000018
       0.721734 0.024852 0.000015
       0.728697 0.025608 0.000002
       0.735668 0.026390 0.000000
       0.742656 0.027199 0.000000
       0.749639 0.028038 0.000000
       0.756646 0.028906 0.000000
       0.763654 0.029806 0.000000
       0.770678 0.030743 0.000000
       0.777709 0.031711 0.000000
       0.784743 0.032732 0.000000
       0.791787 0.033741 0.000000
       0.798859 0.034936 0.000000
       0.805930 0.036031 0.000000
       0.812994 0.037230 0.000000
       0.820066 0.038493 0.000000
       0.827150 0.039819 0.000000
       0.834231 0.041236 0.000000
       0.841308 0.042647 0.000000
       0.848381 0.044235 0.000000
       0.855452 0.045857 0.000000
       0.862522 0.047645 0.000000
       0.869578 0.049578 0.000000
       0.876615 0.051541 0.000000
       0.883648 0.053735 0.000000
       0.890642 0.056168 0.000000
       0.897605 0.058852 0.000000
       0.904508 0.061777 0.000000
       0.911306 0.065281 0.000000
       0.917956 0.069448 0.000000
       0.924447 0.074684 0.000000
       0.930613 0.081310 0.000000
       0.936483 0.088878 0.000000
       0.942051 0.097336 0.000000
       0.947302 0.106654 0.000000
       0.952201 0.116598 0.000000
       0.956744 0.127157 0.000000
       0.960944 0.138244 0.000000
       0.964787 0.149634 0.000000
       0.968291 0.161283 0.000000
       0.971474 0.173033 0.000000
       0.974361 0.184887 0.000000
       0.976976 0.196723 0.000000
       0.979342 0.208458 0.000000
       0.981479 0.220131 0.000000
       0.983403 0.231666 0.000000
       0.985149 0.243007 0.000000
       0.986724 0.254248 0.000000
       0.988151 0.265254 0.000000
       0.989439 0.276144 0.000000
       0.990609 0.286793 0.000000
       0.991667 0.297314 0.000000
       0.992625 0.307636 0.000000
       0.993496 0.317808 0.000000
       0.994284 0.327805 0.000000
       0.994998 0.337639 0.000000
       0.995645 0.347351 0.000000
       0.996227 0.356890 0.000000
       0.996751 0.366300 0.000000
       0.997224 0.375564 0.000000
       0.997648 0.384713 0.000000
       0.998026 0.393741 0.000000
       0.998362 0.402655 0.000000
       0.998660 0.411454 0.000000
       0.998922 0.420150 0.000000
       0.999151 0.428743 0.000000
       0.999349 0.437241 0.000000
       0.999517 0.445630 0.000000
       0.999657 0.453946 0.000000
       0.999770 0.462171 0.000000
       0.999860 0.470321 0.000000
       0.999927 0.478385 0.000000
       0.999972 0.486381 0.000000
       0.999997 0.494301 0.000000
       1.000000 0.502140 0.000000
       1.000000 0.509909 0.000013
       1.000000 0.517614 0.000045
       1.000000 0.525234 0.000097
       1.000000 0.532795 0.000169
       1.000000 0.540284 0.000258
       1.000000 0.547712 0.000365
       1.000000 0.555080 0.000493
       1.000000 0.562405 0.000640
       1.000000 0.569651 0.000804
       1.000000 0.576868 0.000989
       1.000000 0.584019 0.001194
       1.000000 0.591130 0.001419
       1.000000 0.598191 0.001663
       1.000000 0.605215 0.001928
       1.000000 0.612193 0.002215
       1.000000 0.619140 0.002521
       1.000000 0.626032 0.002850
       1.000000 0.632897 0.003201
       1.000000 0.639725 0.003574
       1.000000 0.646514 0.003970
       1.000000 0.653268 0.004390
       1.000000 0.660000 0.004834
       1.000000 0.666691 0.005303
       1.000000 0.673361 0.005797
       1.000000 0.679992 0.006317
       1.000000 0.686606 0.006865
       1.000000 0.693192 0.007441
       1.000000 0.699744 0.008043
       1.000000 0.706280 0.008676
       1.000000 0.712779 0.009349
       1.000000 0.719270 0.010023
       1.000000 0.725733 0.010724
       1.000000 0.732173 0.011565
       1.000000 0.738590 0.012339
       1.000000 0.744986 0.013160
       1.000000 0.751370 0.014042
       1.000000 0.757724 0.014955
       1.000000 0.764062 0.015913
       1.000000 0.770385 0.016915
       1.000000 0.776694 0.017964
       1.000000 0.782977 0.019062
       1.000000 0.789253 0.020212
       1.000000 0.795505 0.021417
       1.000000 0.801745 0.022680
       1.000000 0.807971 0.024005
       1.000000 0.814180 0.025396
       1.000000 0.820376 0.026858
       1.000000 0.826563 0.028394
       1.000000 0.832730 0.030013
       1.000000 0.838886 0.031717
       1.000000 0.845033 0.033480
       1.000000 0.851160 0.035488
       1.000000 0.857281 0.037452
       1.000000 0.863397 0.039592
       1.000000 0.869488 0.041898
       1.000000 0.875574 0.044392
       1.000000 0.881650 0.046958
       1.000000 0.887707 0.049770
       1.000000 0.893762 0.052828
       1.000000 0.899804 0.056209
       1.000000 0.905835 0.059919
       1.000000 0.911847 0.063925
       1.000000 0.917832 0.068579
       1.000000 0.923842 0.073948
       1.000000 0.929813 0.080899
       1.000000 0.935757 0.090648
       1.000000 0.941659 0.103767
       1.000000 0.947516 0.120514
       1.000000 0.953305 0.141489
       1.000000 0.958998 0.167199
       1.000000 0.964561 0.198230
       1.000000 0.969954 0.235141
       1.000000 0.975098 0.278605
       1.000000 0.979921 0.328833
       1.000000 0.984318 0.385714
       1.000000 0.988199 0.448662
       1.000000 0.991498 0.516532
       1.000000 0.994172 0.587541
       1.000000 0.996255 0.659848
       1.000000 0.997784 0.731944
       1.000000 0.998849 0.802589
       1.000000 0.999530 0.871151
       1.000000 0.999885 0.936834
       1.000000 1.000000 1.000000];

case { 'L4'  'KRY'  'YELLOWHEAT' }
descriptorname = 'linear_kry_0-97_c73_n256';
description = 'Black-Red-Yellow heat colour map';
map = [0.000000 0.000000 0.000000
       0.025875 0.000019 0.000000
       0.050536 0.000069 0.000000
       0.069748 0.000131 0.000000
       0.085428 0.000199 0.000000
       0.099057 0.000270 0.000000
       0.111175 0.000342 0.000000
       0.122082 0.000415 0.000000
       0.132227 0.000489 0.000000
       0.141610 0.000564 0.000000
       0.150347 0.000679 0.000000
       0.158508 0.000887 0.000000
       0.166330 0.001002 0.000000
       0.173899 0.001009 0.000000
       0.181035 0.001106 0.000000
       0.187955 0.001184 0.000000
       0.194551 0.001276 0.000000
       0.200904 0.001341 0.000000
       0.207100 0.001435 0.000000
       0.213050 0.001490 0.000000
       0.218837 0.001593 0.000000
       0.224499 0.001567 0.000000
       0.230104 0.001576 0.000009
       0.235706 0.001659 0.000007
       0.241289 0.001746 0.000009
       0.246898 0.001835 0.000007
       0.252540 0.001927 0.000009
       0.258182 0.002022 0.000008
       0.263885 0.002120 0.000009
       0.269579 0.002221 0.000008
       0.275290 0.002325 0.000009
       0.281007 0.002433 0.000009
       0.286765 0.002544 0.000009
       0.292549 0.002658 0.000009
       0.298340 0.002776 0.000009
       0.304158 0.002897 0.000010
       0.310007 0.003022 0.000009
       0.315859 0.003150 0.000010
       0.321713 0.003282 0.000008
       0.327602 0.003419 0.000008
       0.333521 0.003558 0.000008
       0.339448 0.003702 0.000008
       0.345379 0.003850 0.000010
       0.351343 0.004003 0.000009
       0.357328 0.004159 0.000010
       0.363305 0.004319 0.000009
       0.369322 0.004484 0.000008
       0.375344 0.004653 0.000009
       0.381400 0.004827 0.000008
       0.387450 0.005006 0.000010
       0.393532 0.005189 0.000009
       0.399620 0.005378 0.000008
       0.405747 0.005571 0.000010
       0.411866 0.005769 0.000008
       0.418006 0.005973 0.000010
       0.424165 0.006182 0.000008
       0.430346 0.006396 0.000006
       0.436537 0.006616 0.000010
       0.442742 0.006842 0.000008
       0.448965 0.007073 0.000006
       0.455199 0.007311 0.000010
       0.461447 0.007554 0.000008
       0.467712 0.007804 0.000006
       0.473988 0.008060 0.000010
       0.480284 0.008322 0.000008
       0.486595 0.008591 0.000005
       0.492912 0.008868 0.000009
       0.499253 0.009154 0.000009
       0.505602 0.009445 0.000005
       0.511972 0.009742 0.000008
       0.518357 0.010040 0.000009
       0.524744 0.010346 0.000007
       0.531155 0.010674 0.000004
       0.537596 0.011007 0.000008
       0.544043 0.011355 0.000009
       0.550505 0.011709 0.000006
       0.556979 0.012063 0.000003
       0.563451 0.012427 0.000007
       0.569924 0.012791 0.000008
       0.576436 0.013176 0.000007
       0.582934 0.013573 0.000005
       0.589461 0.013979 0.000002
       0.596003 0.014395 0.000005
       0.602551 0.014821 0.000007
       0.609120 0.015259 0.000007
       0.615688 0.015709 0.000006
       0.622273 0.016172 0.000004
       0.628878 0.016646 0.000001
       0.635484 0.017134 0.000002
       0.642109 0.017635 0.000004
       0.648745 0.018151 0.000005
       0.655387 0.018681 0.000006
       0.662051 0.019227 0.000006
       0.668717 0.019788 0.000006
       0.675392 0.020365 0.000006
       0.682080 0.020959 0.000006
       0.688790 0.021571 0.000007
       0.695498 0.022202 0.000007
       0.702217 0.022851 0.000007
       0.708957 0.023521 0.000007
       0.715701 0.024212 0.000007
       0.722446 0.024926 0.000006
       0.729212 0.025662 0.000002
       0.735983 0.026422 0.000000
       0.742772 0.027209 0.000000
       0.749557 0.028023 0.000000
       0.756362 0.028867 0.000000
       0.763170 0.029740 0.000000
       0.769989 0.030646 0.000000
       0.776823 0.031585 0.000000
       0.783654 0.032567 0.000000
       0.790493 0.033563 0.000000
       0.797361 0.034613 0.000000
       0.804234 0.035779 0.000000
       0.811096 0.036893 0.000000
       0.817961 0.038106 0.000000
       0.824837 0.039371 0.000000
       0.831725 0.040693 0.000000
       0.838591 0.042150 0.000000
       0.845465 0.043514 0.000000
       0.852343 0.045178 0.000000
       0.859208 0.046838 0.000000
       0.866075 0.048579 0.000000
       0.872920 0.050451 0.000000
       0.879745 0.052500 0.000000
       0.886564 0.054719 0.000000
       0.893343 0.057161 0.000000
       0.900084 0.059959 0.000000
       0.906768 0.063006 0.000000
       0.913347 0.066454 0.000000
       0.919774 0.070728 0.000000
       0.925961 0.076324 0.000000
       0.931691 0.083888 0.000000
       0.936920 0.093109 0.000000
       0.941639 0.103842 0.000000
       0.945881 0.115497 0.000000
       0.949668 0.127825 0.000000
       0.953056 0.140453 0.000000
       0.956080 0.153178 0.000000
       0.958784 0.165904 0.000000
       0.961214 0.178521 0.000000
       0.963408 0.190874 0.000000
       0.965394 0.203023 0.000000
       0.967207 0.214905 0.000000
       0.968862 0.226549 0.000000
       0.970382 0.237899 0.000000
       0.971784 0.249001 0.000000
       0.973084 0.259854 0.000000
       0.974290 0.270468 0.000000
       0.975415 0.280823 0.000000
       0.976467 0.291016 0.000000
       0.977455 0.300972 0.000000
       0.978380 0.310768 0.000000
       0.979256 0.320369 0.000000
       0.980081 0.329793 0.000000
       0.980861 0.339054 0.000000
       0.981601 0.348165 0.000000
       0.982304 0.357141 0.000000
       0.982971 0.365967 0.000000
       0.983607 0.374670 0.000000
       0.984216 0.383259 0.000000
       0.984797 0.391722 0.000000
       0.985351 0.400077 0.000000
       0.985883 0.408330 0.000000
       0.986392 0.416490 0.000000
       0.986881 0.424554 0.000000
       0.987350 0.432531 0.000000
       0.987800 0.440409 0.000000
       0.988233 0.448215 0.000000
       0.988650 0.455946 0.000000
       0.989053 0.463603 0.000000
       0.989440 0.471193 0.000000
       0.989813 0.478709 0.000000
       0.990173 0.486159 0.000000
       0.990520 0.493548 0.000000
       0.990857 0.500885 0.000000
       0.991182 0.508167 0.000000
       0.991496 0.515377 0.000000
       0.991798 0.522546 0.000000
       0.992093 0.529670 0.000000
       0.992378 0.536745 0.000000
       0.992653 0.543760 0.000000
       0.992918 0.550742 0.000000
       0.993177 0.557678 0.000000
       0.993428 0.564569 0.000000
       0.993668 0.571421 0.000000
       0.993904 0.578235 0.000000
       0.994133 0.585023 0.000000
       0.994352 0.591765 0.000000
       0.994566 0.598467 0.000000
       0.994775 0.605145 0.000000
       0.994974 0.611789 0.000000
       0.995171 0.618404 0.000000
       0.995361 0.624984 0.000000
       0.995542 0.631541 0.000000
       0.995723 0.638064 0.000000
       0.995896 0.644558 0.000000
       0.996062 0.651036 0.000000
       0.996227 0.657487 0.000000
       0.996382 0.663908 0.000000
       0.996538 0.670307 0.000000
       0.996686 0.676681 0.000000
       0.996829 0.683032 0.000000
       0.996971 0.689367 0.000000
       0.997101 0.695677 0.000000
       0.997237 0.701959 0.000000
       0.997362 0.708235 0.000000
       0.997487 0.714481 0.000000
       0.997606 0.720717 0.000000
       0.997721 0.726933 0.000000
       0.997834 0.733132 0.000000
       0.997939 0.739319 0.000000
       0.998048 0.745474 0.000000
       0.998144 0.751630 0.000000
       0.998247 0.757757 0.000000
       0.998336 0.763874 0.000000
       0.998433 0.769975 0.000000
       0.998517 0.776067 0.000000
       0.998607 0.782143 0.000000
       0.998685 0.788203 0.000000
       0.998768 0.794255 0.000000
       0.998841 0.800290 0.000000
       0.998917 0.806312 0.000000
       0.998985 0.812322 0.000000
       0.999056 0.818319 0.000000
       0.999118 0.824310 0.000000
       0.999183 0.830291 0.000000
       0.999240 0.836257 0.000000
       0.999301 0.842215 0.000000
       0.999351 0.848159 0.000000
       0.999408 0.854093 0.000000
       0.999452 0.860027 0.000000
       0.999506 0.865941 0.000000
       0.999544 0.871855 0.000000
       0.999594 0.877751 0.000000
       0.999626 0.883647 0.000000
       0.999672 0.889525 0.000000
       0.999702 0.895404 0.000000
       0.999742 0.901267 0.000000
       0.999769 0.907130 0.000000
       0.999802 0.912981 0.000000
       0.999828 0.918822 0.000000
       0.999854 0.924660 0.000000
       0.999878 0.930494 0.000000
       0.999896 0.936322 0.000000
       0.999920 0.942137 0.000000
       0.999929 0.947950 0.000000
       0.999953 0.953755 0.000000
       0.999959 0.959554 0.000000
       0.999978 0.965343 0.000000
       0.999984 0.971134 0.000000
       0.999994 0.976919 0.000000
       1.000000 0.982701 0.000000
       1.000000 0.988468 0.000000
       1.000000 0.994234 0.000000
       1.000000 1.000000 0.000000];

case { 'L5'  'KGY' }
descriptorname = 'linear_kgy_5-95_c69_n256';
description = 'Colour Map along the green edge of CIELAB space';
map = [0.002008 0.082527 0.022201
       0.004381 0.086150 0.022118
       0.006760 0.089747 0.022041
       0.009163 0.093124 0.021981
       0.011768 0.096433 0.021933
       0.014187 0.099686 0.021895
       0.016629 0.102827 0.021877
       0.019082 0.105869 0.021867
       0.021556 0.108796 0.021877
       0.024018 0.111720 0.021897
       0.026293 0.114592 0.021902
       0.028361 0.117600 0.021900
       0.030209 0.120612 0.021879
       0.031836 0.123682 0.021843
       0.033248 0.126806 0.021797
       0.034480 0.130017 0.021732
       0.035611 0.133185 0.021656
       0.036308 0.136386 0.021568
       0.036803 0.139647 0.021464
       0.037067 0.142990 0.021348
       0.037096 0.146313 0.021222
       0.036881 0.149708 0.021083
       0.036566 0.153049 0.020931
       0.036258 0.156475 0.020778
       0.035956 0.159820 0.020620
       0.035648 0.163263 0.020454
       0.035310 0.166663 0.020281
       0.034935 0.170069 0.020110
       0.034549 0.173469 0.019938
       0.034194 0.176890 0.019764
       0.033900 0.180306 0.019590
       0.033658 0.183737 0.019418
       0.033460 0.187186 0.019251
       0.033308 0.190602 0.019094
       0.033202 0.194065 0.018947
       0.033153 0.197483 0.018811
       0.033171 0.200901 0.018691
       0.033269 0.204381 0.018588
       0.033456 0.207805 0.018509
       0.033755 0.211248 0.018457
       0.034229 0.214669 0.018437
       0.034980 0.218112 0.018456
       0.035810 0.221540 0.018521
       0.036701 0.224954 0.018622
       0.037696 0.228422 0.018735
       0.038702 0.231862 0.018848
       0.039729 0.235314 0.018957
       0.040766 0.238746 0.019061
       0.041844 0.242219 0.019159
       0.042758 0.245683 0.019252
       0.043802 0.249179 0.019339
       0.044931 0.252672 0.019419
       0.045873 0.256155 0.019493
       0.047009 0.259681 0.019561
       0.047989 0.263194 0.019623
       0.049157 0.266705 0.019676
       0.050138 0.270250 0.019721
       0.051173 0.273782 0.019758
       0.052215 0.277323 0.019785
       0.053276 0.280855 0.019802
       0.054355 0.284422 0.019810
       0.055446 0.287989 0.019810
       0.056569 0.291580 0.019809
       0.057528 0.295146 0.019834
       0.058624 0.298744 0.019890
       0.059734 0.302329 0.019977
       0.060680 0.305961 0.020098
       0.061651 0.309570 0.020253
       0.062756 0.313179 0.020445
       0.063696 0.316807 0.020677
       0.064656 0.320452 0.020950
       0.065612 0.324076 0.021258
       0.066576 0.327725 0.021584
       0.067549 0.331375 0.021916
       0.068538 0.335063 0.022251
       0.069520 0.338718 0.022589
       0.070402 0.342400 0.022929
       0.071419 0.346067 0.023272
       0.072319 0.349770 0.023618
       0.073361 0.353460 0.023966
       0.074283 0.357177 0.024318
       0.075218 0.360866 0.024672
       0.076165 0.364589 0.025028
       0.077123 0.368319 0.025388
       0.078091 0.372038 0.025750
       0.079071 0.375780 0.026116
       0.080061 0.379523 0.026484
       0.081061 0.383280 0.026854
       0.082077 0.387022 0.027228
       0.082984 0.390794 0.027604
       0.084013 0.394557 0.027983
       0.084952 0.398337 0.028365
       0.085878 0.402106 0.028750
       0.086941 0.405910 0.029138
       0.087908 0.409700 0.029528
       0.088876 0.413498 0.029921
       0.089858 0.417309 0.030317
       0.090847 0.421108 0.030716
       0.091852 0.424937 0.031118
       0.092761 0.428757 0.031521
       0.093773 0.432594 0.031927
       0.094809 0.436427 0.032341
       0.095750 0.440265 0.032760
       0.096692 0.444121 0.033171
       0.097753 0.447963 0.033567
       0.098721 0.451835 0.033985
       0.099698 0.455695 0.034483
       0.100686 0.459570 0.035028
       0.101683 0.463439 0.035514
       0.102691 0.467333 0.035934
       0.103701 0.471223 0.036349
       0.104634 0.475122 0.036789
       0.105675 0.479015 0.037237
       0.106710 0.482919 0.037682
       0.107661 0.486834 0.038128
       0.108645 0.490760 0.038578
       0.109712 0.494680 0.039032
       0.110689 0.498610 0.039489
       0.111689 0.502536 0.039948
       0.112701 0.506475 0.040406
       0.113690 0.510423 0.040871
       0.114641 0.514382 0.041351
       0.115694 0.518338 0.041826
       0.116697 0.522290 0.042253
       0.117694 0.526268 0.042639
       0.118731 0.530238 0.043044
       0.119703 0.534212 0.043511
       0.120699 0.538206 0.044017
       0.121701 0.542188 0.044525
       0.122712 0.546190 0.044992
       0.123733 0.550184 0.045398
       0.124760 0.554189 0.045806
       0.125808 0.558192 0.046288
       0.126785 0.562222 0.046801
       0.127826 0.566236 0.047247
       0.128821 0.570257 0.047653
       0.129884 0.574287 0.048118
       0.130895 0.578325 0.048645
       0.131902 0.582367 0.049135
       0.132921 0.586418 0.049563
       0.133948 0.590466 0.049980
       0.134979 0.594526 0.050418
       0.135949 0.598586 0.050866
       0.137009 0.602655 0.051311
       0.138057 0.606731 0.051757
       0.139039 0.610809 0.052205
       0.140066 0.614883 0.052658
       0.141136 0.618982 0.053112
       0.142145 0.623070 0.053569
       0.143176 0.627164 0.054029
       0.144208 0.631270 0.054491
       0.145262 0.635372 0.054953
       0.146260 0.639487 0.055418
       0.147301 0.643599 0.055896
       0.148315 0.647722 0.056378
       0.149378 0.651854 0.056824
       0.150406 0.655987 0.057227
       0.151425 0.660118 0.057637
       0.152456 0.664258 0.058099
       0.153495 0.668408 0.058598
       0.154541 0.672559 0.059100
       0.155599 0.676713 0.059581
       0.156645 0.680864 0.060024
       0.157662 0.685035 0.060440
       0.158730 0.689208 0.060845
       0.159733 0.693378 0.061267
       0.160799 0.697553 0.061737
       0.161871 0.701735 0.062250
       0.162905 0.705928 0.062740
       0.163965 0.710120 0.063178
       0.164970 0.714313 0.063603
       0.166021 0.718516 0.064042
       0.167101 0.722720 0.064491
       0.168121 0.726934 0.064939
       0.169209 0.731147 0.065387
       0.170239 0.735371 0.065837
       0.171283 0.739601 0.066291
       0.172332 0.743824 0.066748
       0.173388 0.748060 0.067206
       0.174459 0.752293 0.067663
       0.175496 0.756536 0.068124
       0.176544 0.760784 0.068597
       0.177643 0.765037 0.069073
       0.178690 0.769287 0.069516
       0.179730 0.773546 0.069920
       0.180781 0.777816 0.070337
       0.181838 0.782085 0.070809
       0.182903 0.786360 0.071291
       0.183974 0.790632 0.071723
       0.185058 0.794914 0.072134
       0.186121 0.799204 0.072590
       0.187182 0.803490 0.073087
       0.188259 0.807786 0.073557
       0.189304 0.812084 0.073983
       0.190365 0.816390 0.074402
       0.191431 0.820693 0.074835
       0.192504 0.825004 0.075276
       0.193585 0.829326 0.075716
       0.194667 0.833645 0.076155
       0.195715 0.837969 0.076597
       0.196809 0.842298 0.077042
       0.197857 0.846625 0.077489
       0.198920 0.850961 0.077938
       0.199989 0.855309 0.078388
       0.201065 0.859653 0.078841
       0.202149 0.864005 0.079296
       0.203238 0.868359 0.079753
       0.204335 0.872713 0.080211
       0.205390 0.877070 0.080669
       0.206497 0.881441 0.081135
       0.207556 0.885807 0.081610
       0.208629 0.890174 0.082069
       0.209736 0.894559 0.082489
       0.211665 0.898901 0.082885
       0.215766 0.903138 0.083267
       0.222428 0.907241 0.083619
       0.231288 0.911222 0.083917
       0.241916 0.915092 0.084156
       0.253980 0.918857 0.084342
       0.267138 0.922532 0.084483
       0.281219 0.926108 0.084587
       0.296045 0.929590 0.084656
       0.311383 0.932988 0.084696
       0.327070 0.936316 0.084710
       0.343030 0.939560 0.084701
       0.359128 0.942738 0.084672
       0.375354 0.945842 0.084623
       0.391809 0.948868 0.084553
       0.408322 0.951826 0.084465
       0.424831 0.954726 0.084363
       0.441299 0.957563 0.084247
       0.457854 0.960330 0.084113
       0.474425 0.963029 0.083964
       0.490931 0.965671 0.083803
       0.507371 0.968261 0.083635
       0.523883 0.970773 0.083456
       0.540327 0.973235 0.083278
       0.556678 0.975640 0.083106
       0.573107 0.977975 0.082937
       0.589452 0.980258 0.082777
       0.605714 0.982491 0.082624
       0.622048 0.984648 0.082472
       0.638305 0.986754 0.082325
       0.654502 0.988802 0.082180
       0.670744 0.990784 0.082035
       0.686887 0.992720 0.081896
       0.703051 0.994584 0.081762
       0.719187 0.996392 0.081638
       0.735253 0.998154 0.081527
       0.751386 0.999839 0.081428
       0.767421 1.000000 0.081347
       0.783495 1.000000 0.081282
       0.799562 1.000000 0.081236
       0.815550 1.000000 0.081210
       0.831586 1.000000 0.081204
       0.847538 1.000000 0.081223];

case { 'L6'  'KBC' }
descriptorname = 'linear_kbc_5-95_c73_n256';
description = 'Blue shades running vertically up the blue edge of CIELAB space';
map = [0.000000 0.002438 0.306872
       0.000000 0.004469 0.314367
       0.000000 0.006232 0.321910
       0.000000 0.007730 0.329510
       0.000000 0.008958 0.337127
       0.000000 0.009897 0.344786
       0.000000 0.010555 0.352493
       0.000000 0.011009 0.360232
       0.000000 0.011140 0.368022
       0.001883 0.010944 0.375848
       0.004341 0.010697 0.383710
       0.006854 0.010463 0.391608
       0.009466 0.010231 0.399525
       0.012350 0.009998 0.407500
       0.015052 0.009759 0.415504
       0.017851 0.009498 0.423537
       0.020695 0.009222 0.431626
       0.023570 0.008937 0.439730
       0.026496 0.008645 0.447867
       0.029448 0.008350 0.456055
       0.032396 0.008059 0.464287
       0.035580 0.007766 0.472534
       0.038524 0.007470 0.480819
       0.041458 0.007185 0.489152
       0.044141 0.006907 0.497533
       0.046790 0.006638 0.505925
       0.049320 0.006379 0.514383
       0.051527 0.006144 0.522866
       0.053656 0.005933 0.531390
       0.055683 0.005746 0.539953
       0.057455 0.005589 0.548571
       0.059195 0.005466 0.557220
       0.060593 0.005388 0.565903
       0.061766 0.005363 0.574639
       0.062841 0.005396 0.583427
       0.063532 0.005494 0.592269
       0.063966 0.005665 0.601141
       0.064100 0.005920 0.610072
       0.063882 0.006272 0.619066
       0.063281 0.006733 0.628102
       0.062148 0.007317 0.637189
       0.060493 0.008041 0.646336
       0.058092 0.008929 0.655567
       0.055233 0.009962 0.664818
       0.052105 0.011266 0.674023
       0.049146 0.012683 0.683135
       0.046014 0.014144 0.692158
       0.043090 0.015741 0.701081
       0.040435 0.017485 0.709919
       0.037863 0.019379 0.718660
       0.035616 0.021429 0.727312
       0.033428 0.023640 0.735866
       0.031841 0.026021 0.744328
       0.030635 0.028579 0.752682
       0.029823 0.031319 0.760945
       0.029422 0.034249 0.769102
       0.029449 0.037557 0.777172
       0.029926 0.040864 0.785133
       0.030874 0.044198 0.792996
       0.032312 0.047578 0.800757
       0.034281 0.051036 0.808416
       0.036977 0.054512 0.815969
       0.040055 0.058003 0.823410
       0.043483 0.061561 0.830755
       0.047434 0.065191 0.837993
       0.051627 0.068868 0.845125
       0.056188 0.072421 0.852156
       0.060871 0.076069 0.859074
       0.065858 0.079772 0.865889
       0.071059 0.083524 0.872590
       0.076320 0.087319 0.879175
       0.081824 0.091117 0.885645
       0.087161 0.095093 0.891948
       0.092292 0.099225 0.898051
       0.097256 0.103649 0.903954
       0.102075 0.108186 0.909654
       0.106779 0.113036 0.915139
       0.111265 0.117978 0.920409
       0.115611 0.123059 0.925470
       0.119814 0.128372 0.930323
       0.123893 0.133784 0.934963
       0.127886 0.139280 0.939399
       0.131717 0.144981 0.943623
       0.135407 0.150710 0.947638
       0.138977 0.156590 0.951434
       0.142446 0.162550 0.955011
       0.145790 0.168593 0.958370
       0.148954 0.174735 0.961511
       0.152018 0.180912 0.964443
       0.154968 0.187215 0.967166
       0.157777 0.193551 0.969668
       0.160438 0.199923 0.971961
       0.163014 0.206431 0.974029
       0.165371 0.212916 0.975872
       0.167653 0.219509 0.977498
       0.169779 0.226140 0.978903
       0.171714 0.232782 0.980097
       0.173520 0.239496 0.981070
       0.175178 0.246227 0.981829
       0.176642 0.253015 0.982376
       0.178029 0.259774 0.982792
       0.179239 0.266450 0.983175
       0.180369 0.273045 0.983556
       0.181410 0.279567 0.983938
       0.182365 0.285985 0.984317
       0.183232 0.292367 0.984693
       0.184008 0.298676 0.985066
       0.184697 0.304925 0.985436
       0.185301 0.311119 0.985804
       0.185798 0.317251 0.986171
       0.186189 0.323340 0.986537
       0.186492 0.329391 0.986900
       0.186711 0.335386 0.987260
       0.186843 0.341331 0.987617
       0.186881 0.347253 0.987970
       0.186824 0.353123 0.988323
       0.186673 0.358956 0.988676
       0.186432 0.364769 0.989026
       0.186104 0.370544 0.989374
       0.185678 0.376307 0.989718
       0.185131 0.382013 0.990059
       0.184476 0.387706 0.990398
       0.183727 0.393376 0.990736
       0.182873 0.399017 0.991071
       0.181917 0.404636 0.991404
       0.180857 0.410250 0.991735
       0.179692 0.415818 0.992063
       0.178432 0.421370 0.992389
       0.177076 0.426913 0.992708
       0.175763 0.432419 0.993013
       0.174622 0.437883 0.993297
       0.173551 0.443308 0.993561
       0.172629 0.448706 0.993806
       0.171852 0.454065 0.994030
       0.171218 0.459399 0.994234
       0.170721 0.464702 0.994419
       0.170365 0.469976 0.994585
       0.170152 0.475223 0.994732
       0.170082 0.480432 0.994859
       0.170155 0.485622 0.994966
       0.170381 0.490803 0.995053
       0.170769 0.495931 0.995120
       0.171315 0.501059 0.995167
       0.172010 0.506146 0.995194
       0.172860 0.511223 0.995201
       0.173864 0.516275 0.995188
       0.175025 0.521312 0.995156
       0.176285 0.526333 0.995104
       0.177784 0.531322 0.995031
       0.179360 0.536298 0.994938
       0.181096 0.541247 0.994825
       0.182989 0.546198 0.994691
       0.185045 0.551113 0.994538
       0.187200 0.556009 0.994365
       0.189479 0.560903 0.994171
       0.191890 0.565775 0.993958
       0.194428 0.570635 0.993728
       0.196856 0.575500 0.993492
       0.199063 0.580363 0.993261
       0.201102 0.585237 0.993034
       0.202967 0.590105 0.992813
       0.204643 0.594993 0.992597
       0.206125 0.599887 0.992386
       0.207400 0.604786 0.992180
       0.208499 0.609689 0.991980
       0.209417 0.614597 0.991787
       0.210160 0.619527 0.991599
       0.210724 0.624449 0.991416
       0.211113 0.629377 0.991239
       0.211324 0.634324 0.991067
       0.211353 0.639264 0.990901
       0.211197 0.644212 0.990739
       0.210855 0.649179 0.990584
       0.210326 0.654139 0.990434
       0.209607 0.659109 0.990290
       0.208699 0.664091 0.990152
       0.207600 0.669076 0.990019
       0.206314 0.674065 0.989891
       0.204782 0.679060 0.989768
       0.203046 0.684061 0.989651
       0.201078 0.689081 0.989539
       0.198898 0.694087 0.989432
       0.196505 0.699111 0.989331
       0.193819 0.704143 0.989236
       0.190879 0.709182 0.989144
       0.187884 0.714205 0.989050
       0.184880 0.719230 0.988947
       0.181980 0.724234 0.988832
       0.179259 0.729227 0.988706
       0.176654 0.734207 0.988567
       0.174231 0.739176 0.988416
       0.171931 0.744122 0.988253
       0.169829 0.749059 0.988080
       0.167855 0.753991 0.987895
       0.166082 0.758904 0.987698
       0.164524 0.763806 0.987489
       0.163161 0.768695 0.987268
       0.162024 0.773574 0.987034
       0.161102 0.778444 0.986787
       0.160373 0.783299 0.986528
       0.159912 0.788144 0.986256
       0.159698 0.792981 0.985973
       0.159722 0.797800 0.985678
       0.159987 0.802613 0.985371
       0.160527 0.807418 0.985053
       0.161366 0.812207 0.984721
       0.162411 0.816988 0.984377
       0.163726 0.821761 0.984018
       0.165227 0.826521 0.983647
       0.167065 0.831274 0.983265
       0.169103 0.836007 0.982873
       0.171339 0.840735 0.982470
       0.173868 0.845458 0.982051
       0.177288 0.850146 0.981621
       0.182585 0.854741 0.981191
       0.190031 0.859229 0.980762
       0.199160 0.863628 0.980335
       0.209653 0.867936 0.979909
       0.221165 0.872166 0.979482
       0.233382 0.876320 0.979052
       0.246220 0.880409 0.978623
       0.259529 0.884435 0.978196
       0.273093 0.888398 0.977770
       0.286808 0.892309 0.977343
       0.300629 0.896168 0.976912
       0.314476 0.899983 0.976480
       0.328317 0.903752 0.976048
       0.342243 0.907475 0.975615
       0.356131 0.911150 0.975182
       0.369912 0.914791 0.974747
       0.383608 0.918399 0.974310
       0.397280 0.921968 0.973873
       0.410905 0.925496 0.973435
       0.424394 0.928994 0.972995
       0.437791 0.932465 0.972553
       0.451162 0.935902 0.972108
       0.464432 0.939307 0.971660
       0.477547 0.942693 0.971211
       0.490688 0.946032 0.970761
       0.503698 0.949353 0.970309
       0.516588 0.952655 0.969856
       0.529492 0.955920 0.969401
       0.542269 0.959161 0.968944
       0.554956 0.962380 0.968486
       0.567629 0.965569 0.968027
       0.580184 0.968744 0.967564
       0.592706 0.971890 0.967097
       0.605154 0.975009 0.966628
       0.617509 0.978115 0.966158
       0.629873 0.981191 0.965687
       0.642128 0.984253 0.965213
       0.654358 0.987286 0.964737
       0.666543 0.990302 0.964259
       0.678637 0.993297 0.963781
       0.690720 0.996258 0.963301
       0.702714 0.999222 0.962818];

case { 'L7'  'BMW' }
descriptorname = 'linear_bmw_5-95_c86_n256';
description = 'Blue-Pink-Light Pink colour map';
map = [0.000000 0.007869 0.295627
       0.000000 0.010180 0.303467
       0.000000 0.012490 0.311358
       0.000000 0.014296 0.319287
       0.000000 0.015847 0.327242
       0.000000 0.017130 0.335265
       0.000000 0.018143 0.343297
       0.000000 0.018877 0.351369
       0.000000 0.019332 0.359481
       0.000000 0.019537 0.367623
       0.000000 0.019696 0.375810
       0.000000 0.019848 0.384022
       0.000000 0.019989 0.392263
       0.000278 0.020121 0.400548
       0.001318 0.020246 0.408841
       0.002391 0.020361 0.417181
       0.003488 0.020469 0.425539
       0.004603 0.020572 0.433921
       0.005743 0.020668 0.442358
       0.006899 0.020759 0.450801
       0.008059 0.020848 0.459281
       0.009239 0.020933 0.467782
       0.010387 0.021014 0.476317
       0.011786 0.021099 0.484870
       0.012932 0.021185 0.493460
       0.014093 0.021272 0.502081
       0.015228 0.021363 0.510717
       0.016327 0.021464 0.519389
       0.017385 0.021575 0.528080
       0.018402 0.021697 0.536807
       0.019367 0.021832 0.545548
       0.020271 0.021984 0.554318
       0.021093 0.022157 0.563116
       0.021816 0.022358 0.571932
       0.022431 0.022589 0.580777
       0.022925 0.022854 0.589642
       0.023280 0.023157 0.598539
       0.023474 0.023505 0.607466
       0.023484 0.023905 0.616405
       0.023283 0.024365 0.625367
       0.022843 0.024892 0.634354
       0.022130 0.025497 0.643351
       0.021108 0.026192 0.652395
       0.019919 0.026976 0.661439
       0.018888 0.027836 0.670428
       0.018212 0.028767 0.679327
       0.017924 0.029768 0.688164
       0.018038 0.030843 0.696919
       0.018576 0.031990 0.705600
       0.019563 0.033222 0.714202
       0.021022 0.034609 0.722728
       0.022976 0.036126 0.731178
       0.025460 0.037612 0.739554
       0.028511 0.039192 0.747834
       0.032139 0.040867 0.756031
       0.036548 0.042546 0.764148
       0.041400 0.044350 0.772186
       0.046471 0.046119 0.780140
       0.051815 0.048019 0.788011
       0.057373 0.050007 0.795798
       0.063189 0.051954 0.803500
       0.069106 0.054002 0.811112
       0.075029 0.056167 0.818631
       0.081171 0.058240 0.826070
       0.087403 0.060492 0.833422
       0.093627 0.062774 0.840684
       0.099985 0.064997 0.847855
       0.106455 0.067338 0.854945
       0.112931 0.069742 0.861940
       0.119434 0.072103 0.868841
       0.126061 0.074573 0.875645
       0.132793 0.077004 0.882341
       0.139922 0.079441 0.888889
       0.147703 0.081844 0.895241
       0.156009 0.084044 0.901389
       0.164765 0.086070 0.907340
       0.173993 0.088131 0.913080
       0.183612 0.090027 0.918602
       0.193589 0.091859 0.923923
       0.203831 0.093506 0.929031
       0.214294 0.095185 0.933937
       0.224953 0.096634 0.938640
       0.235845 0.098158 0.943134
       0.246852 0.099485 0.947418
       0.258016 0.100759 0.951488
       0.269359 0.101958 0.955341
       0.280762 0.103098 0.958976
       0.292299 0.104102 0.962401
       0.303896 0.105031 0.965616
       0.315582 0.105968 0.968624
       0.327300 0.106791 0.971417
       0.339100 0.107487 0.973999
       0.350939 0.108108 0.976363
       0.362821 0.108698 0.978503
       0.374760 0.109246 0.980430
       0.386712 0.109699 0.982143
       0.398701 0.110057 0.983636
       0.410710 0.110342 0.984922
       0.422710 0.110562 0.985989
       0.434738 0.110719 0.986851
       0.446670 0.110837 0.987573
       0.458428 0.110942 0.988238
       0.470007 0.111043 0.988880
       0.481393 0.111140 0.989495
       0.492621 0.111235 0.990080
       0.503706 0.111325 0.990639
       0.514652 0.111412 0.991173
       0.525460 0.111493 0.991680
       0.536149 0.111570 0.992160
       0.546732 0.111639 0.992613
       0.557209 0.111703 0.993038
       0.567577 0.111761 0.993435
       0.577850 0.111813 0.993804
       0.588044 0.111859 0.994146
       0.598155 0.111899 0.994462
       0.608198 0.111931 0.994752
       0.618153 0.111956 0.995014
       0.628048 0.111971 0.995248
       0.637872 0.111978 0.995455
       0.647636 0.111977 0.995634
       0.657349 0.111967 0.995787
       0.666993 0.111948 0.995913
       0.676596 0.111919 0.996012
       0.686141 0.111881 0.996084
       0.695644 0.111832 0.996129
       0.705098 0.111771 0.996145
       0.714509 0.111698 0.996133
       0.723881 0.111623 0.996095
       0.733169 0.111644 0.996039
       0.742298 0.111939 0.995980
       0.751239 0.112622 0.995926
       0.759979 0.113675 0.995877
       0.768536 0.115101 0.995834
       0.776914 0.116982 0.995796
       0.785101 0.119253 0.995764
       0.793117 0.121843 0.995737
       0.800958 0.124848 0.995716
       0.808638 0.128194 0.995700
       0.816152 0.131836 0.995689
       0.823499 0.135750 0.995683
       0.830690 0.140000 0.995682
       0.837716 0.144563 0.995687
       0.844581 0.149347 0.995697
       0.851289 0.154341 0.995711
       0.857853 0.159563 0.995731
       0.864270 0.164994 0.995756
       0.870528 0.170624 0.995785
       0.876645 0.176385 0.995819
       0.882614 0.182336 0.995858
       0.888437 0.188472 0.995902
       0.894108 0.194717 0.995951
       0.899639 0.201042 0.996004
       0.905028 0.207544 0.996062
       0.910277 0.214128 0.996124
       0.915379 0.220820 0.996191
       0.920350 0.227573 0.996263
       0.925177 0.234440 0.996337
       0.929846 0.241408 0.996404
       0.934351 0.248511 0.996459
       0.938703 0.255690 0.996500
       0.942896 0.262992 0.996527
       0.946926 0.270385 0.996541
       0.950802 0.277844 0.996541
       0.954518 0.285374 0.996527
       0.958075 0.293005 0.996499
       0.961461 0.300706 0.996457
       0.964688 0.308489 0.996400
       0.967764 0.316321 0.996329
       0.970672 0.324190 0.996244
       0.973426 0.332136 0.996145
       0.976017 0.340116 0.996032
       0.978451 0.348139 0.995904
       0.980726 0.356214 0.995762
       0.982830 0.364323 0.995605
       0.984771 0.372480 0.995433
       0.986548 0.380685 0.995246
       0.988160 0.388909 0.995045
       0.989616 0.397153 0.994830
       0.990903 0.405446 0.994599
       0.992029 0.413752 0.994355
       0.992987 0.422082 0.994095
       0.993767 0.430460 0.993820
       0.994377 0.438845 0.993530
       0.994819 0.447259 0.993224
       0.995104 0.455685 0.992906
       0.995288 0.464073 0.992587
       0.995439 0.472352 0.992286
       0.995582 0.480518 0.992007
       0.995717 0.488587 0.991753
       0.995846 0.496541 0.991523
       0.995967 0.504424 0.991317
       0.996082 0.512190 0.991134
       0.996189 0.519878 0.990974
       0.996289 0.527492 0.990838
       0.996382 0.535022 0.990724
       0.996469 0.542482 0.990634
       0.996548 0.549865 0.990568
       0.996622 0.557187 0.990527
       0.996689 0.564431 0.990510
       0.996750 0.571618 0.990516
       0.996804 0.578748 0.990546
       0.996852 0.585825 0.990599
       0.996893 0.592842 0.990676
       0.996928 0.599803 0.990776
       0.996957 0.606716 0.990900
       0.996981 0.613571 0.991050
       0.996999 0.620399 0.991224
       0.997011 0.627163 0.991422
       0.997017 0.633895 0.991644
       0.997018 0.640577 0.991889
       0.997012 0.647216 0.992159
       0.997002 0.653811 0.992453
       0.996986 0.660383 0.992771
       0.996973 0.666916 0.993103
       0.996974 0.673408 0.993426
       0.996994 0.679851 0.993729
       0.997030 0.686268 0.994016
       0.997078 0.692652 0.994288
       0.997137 0.698989 0.994545
       0.997205 0.705308 0.994788
       0.997280 0.711599 0.995019
       0.997361 0.717858 0.995236
       0.997447 0.724089 0.995441
       0.997535 0.730302 0.995634
       0.997624 0.736484 0.995817
       0.997713 0.742656 0.995990
       0.997801 0.748795 0.996153
       0.997890 0.754924 0.996305
       0.997976 0.761025 0.996447
       0.998059 0.767108 0.996580
       0.998137 0.773177 0.996705
       0.998213 0.779230 0.996821
       0.998284 0.785269 0.996927
       0.998348 0.791286 0.997026
       0.998406 0.797290 0.997117
       0.998460 0.803281 0.997198
       0.998506 0.809262 0.997273
       0.998542 0.815223 0.997340
       0.998575 0.821174 0.997398
       0.998598 0.827114 0.997450
       0.998610 0.833039 0.997495
       0.998617 0.838954 0.997531
       0.998614 0.844860 0.997561
       0.998599 0.850750 0.997584
       0.998578 0.856635 0.997599
       0.998544 0.862514 0.997608
       0.998501 0.868379 0.997610
       0.998447 0.874227 0.997605
       0.998381 0.880073 0.997594
       0.998307 0.885914 0.997575
       0.998218 0.891742 0.997551
       0.998120 0.897564 0.997519
       0.998009 0.903386 0.997482
       0.997886 0.909182 0.997438
       0.997752 0.914978 0.997387
       0.997602 0.920768 0.997331];

case { 'L8'  'BMY' }
descriptorname = 'linear_bmy_10-95_c71_n256';
description = 'Blue-Magenta-Yellow highly saturated colour map';
map = [0.001807 0.058001 0.363987
       0.002411 0.059964 0.370477
       0.003199 0.061692 0.376968
       0.004284 0.063571 0.383411
       0.005764 0.065341 0.389792
       0.007509 0.067136 0.396170
       0.009708 0.068953 0.402477
       0.012532 0.070615 0.408753
       0.015500 0.072320 0.414992
       0.019067 0.074062 0.421158
       0.023134 0.075673 0.427294
       0.027668 0.077301 0.433365
       0.032903 0.078925 0.439375
       0.038954 0.080546 0.445323
       0.045181 0.082169 0.451225
       0.051521 0.083668 0.457045
       0.058053 0.085134 0.462783
       0.064730 0.086625 0.468469
       0.071444 0.088071 0.474086
       0.078236 0.089456 0.479603
       0.085243 0.090814 0.485020
       0.092333 0.092133 0.490377
       0.099463 0.093353 0.495635
       0.106838 0.094646 0.500785
       0.114228 0.095771 0.505813
       0.121780 0.096854 0.510755
       0.129484 0.097993 0.515579
       0.137276 0.098973 0.520292
       0.145246 0.099911 0.524867
       0.153305 0.100783 0.529301
       0.161591 0.101593 0.533576
       0.169954 0.102330 0.537726
       0.178486 0.103002 0.541697
       0.187154 0.103575 0.545519
       0.195990 0.104020 0.549154
       0.204992 0.104361 0.552592
       0.214185 0.104609 0.555818
       0.223551 0.104747 0.558836
       0.233097 0.104755 0.561622
       0.242865 0.104620 0.564150
       0.252855 0.104340 0.566400
       0.263051 0.103908 0.568355
       0.273476 0.103266 0.569997
       0.284110 0.102366 0.571302
       0.295018 0.101246 0.572244
       0.306191 0.099883 0.572810
       0.317334 0.098383 0.573123
       0.328286 0.096731 0.573342
       0.339016 0.095191 0.573505
       0.349523 0.093473 0.573615
       0.359842 0.091850 0.573668
       0.369980 0.090102 0.573663
       0.379964 0.088354 0.573601
       0.389785 0.086567 0.573485
       0.399466 0.084774 0.573315
       0.409028 0.082914 0.573086
       0.418448 0.081085 0.572797
       0.427768 0.079164 0.572450
       0.436964 0.077258 0.572046
       0.446044 0.075373 0.571586
       0.455046 0.073529 0.571068
       0.463939 0.071572 0.570492
       0.472733 0.069648 0.569860
       0.481448 0.067661 0.569175
       0.490079 0.065690 0.568432
       0.498635 0.063767 0.567632
       0.507099 0.061792 0.566773
       0.515490 0.059971 0.565848
       0.523820 0.057948 0.564872
       0.532073 0.056175 0.563851
       0.540252 0.054286 0.562758
       0.548377 0.052493 0.561604
       0.556419 0.050790 0.560397
       0.564413 0.049208 0.559130
       0.572342 0.047520 0.557808
       0.580209 0.045977 0.556427
       0.588004 0.044652 0.555015
       0.595748 0.043191 0.553580
       0.603429 0.042094 0.552141
       0.611044 0.040933 0.550673
       0.618602 0.039904 0.549197
       0.626094 0.039006 0.547699
       0.633541 0.038249 0.546196
       0.640923 0.037647 0.544665
       0.648257 0.037202 0.543126
       0.655543 0.036908 0.541559
       0.662779 0.036776 0.539990
       0.669963 0.036816 0.538413
       0.677097 0.037039 0.536808
       0.684174 0.037443 0.535177
       0.691216 0.038023 0.533537
       0.698213 0.038789 0.531892
       0.705166 0.039756 0.530222
       0.712073 0.040926 0.528539
       0.718937 0.042269 0.526839
       0.725763 0.043681 0.525114
       0.732545 0.045402 0.523382
       0.739287 0.047268 0.521621
       0.745979 0.049331 0.519851
       0.752636 0.051385 0.518073
       0.759259 0.053656 0.516259
       0.765847 0.056145 0.514450
       0.772388 0.058663 0.512604
       0.778895 0.061273 0.510746
       0.785368 0.064105 0.508880
       0.791795 0.067003 0.506977
       0.798150 0.070170 0.505032
       0.804401 0.073831 0.503004
       0.810538 0.077734 0.500912
       0.816578 0.082182 0.498735
       0.822505 0.086838 0.496464
       0.828316 0.091840 0.494132
       0.834021 0.097033 0.491711
       0.839615 0.102552 0.489193
       0.845098 0.108207 0.486611
       0.850471 0.114097 0.483938
       0.855734 0.120158 0.481182
       0.860879 0.126390 0.478346
       0.865920 0.132721 0.475434
       0.870848 0.139132 0.472424
       0.875666 0.145738 0.469338
       0.880360 0.152361 0.466143
       0.884949 0.159124 0.462871
       0.889425 0.165922 0.459530
       0.893786 0.172817 0.456085
       0.898030 0.179779 0.452562
       0.902157 0.186809 0.448935
       0.906173 0.193900 0.445214
       0.910075 0.200980 0.441411
       0.913858 0.208168 0.437520
       0.917520 0.215382 0.433508
       0.921057 0.222652 0.429423
       0.924487 0.229912 0.425240
       0.927798 0.237271 0.420945
       0.930992 0.244589 0.416567
       0.934054 0.252001 0.412094
       0.937013 0.259394 0.407599
       0.939859 0.266747 0.403164
       0.942597 0.274106 0.398789
       0.945222 0.281443 0.394493
       0.947736 0.288781 0.390263
       0.950147 0.296089 0.386098
       0.952454 0.303382 0.382003
       0.954656 0.310693 0.377978
       0.956742 0.317967 0.374028
       0.958730 0.325230 0.370123
       0.960612 0.332515 0.366305
       0.962386 0.339764 0.362532
       0.964055 0.347010 0.358827
       0.965616 0.354245 0.355220
       0.967078 0.361481 0.351637
       0.968435 0.368712 0.348141
       0.969684 0.375928 0.344699
       0.970825 0.383152 0.341337
       0.971862 0.390357 0.338027
       0.972797 0.397569 0.334809
       0.973623 0.404764 0.331614
       0.974343 0.411972 0.328517
       0.974952 0.419174 0.325472
       0.975457 0.426361 0.322523
       0.975859 0.433549 0.319615
       0.976153 0.440735 0.316779
       0.976336 0.447921 0.314018
       0.976408 0.455117 0.311327
       0.976373 0.462289 0.308718
       0.976242 0.469470 0.306166
       0.976068 0.476579 0.303627
       0.975914 0.483598 0.301116
       0.975790 0.490539 0.298612
       0.975694 0.497378 0.296099
       0.975628 0.504135 0.293580
       0.975594 0.510796 0.291077
       0.975593 0.517395 0.288558
       0.975623 0.523904 0.286028
       0.975684 0.530346 0.283522
       0.975777 0.536724 0.280985
       0.975904 0.543024 0.278481
       0.976066 0.549261 0.275968
       0.976259 0.555428 0.273431
       0.976486 0.561551 0.270898
       0.976748 0.567609 0.268351
       0.977047 0.573607 0.265822
       0.977381 0.579556 0.263283
       0.977747 0.585450 0.260726
       0.978147 0.591286 0.258163
       0.978586 0.597082 0.255620
       0.979067 0.602820 0.253067
       0.979584 0.608524 0.250488
       0.980134 0.614164 0.247943
       0.980721 0.619782 0.245339
       0.981351 0.625340 0.242765
       0.982021 0.630867 0.240179
       0.982725 0.636346 0.237620
       0.983466 0.641773 0.235029
       0.984253 0.647176 0.232415
       0.985070 0.652558 0.229822
       0.985855 0.657930 0.227478
       0.986562 0.663308 0.225449
       0.987190 0.668697 0.223746
       0.987744 0.674097 0.222336
       0.988234 0.679506 0.221191
       0.988671 0.684928 0.220260
       0.989056 0.690349 0.219516
       0.989392 0.695780 0.218964
       0.989677 0.701205 0.218598
       0.989916 0.706641 0.218405
       0.990116 0.712067 0.218347
       0.990278 0.717503 0.218414
       0.990401 0.722924 0.218618
       0.990485 0.728356 0.218956
       0.990535 0.733785 0.219398
       0.990553 0.739216 0.219946
       0.990533 0.744634 0.220617
       0.990483 0.750054 0.221352
       0.990400 0.755478 0.222179
       0.990285 0.760895 0.223133
       0.990141 0.766308 0.224114
       0.989964 0.771723 0.225221
       0.989757 0.777135 0.226420
       0.989522 0.782537 0.227655
       0.989255 0.787940 0.228978
       0.988960 0.793346 0.230331
       0.988632 0.798742 0.231820
       0.988278 0.804139 0.233299
       0.987896 0.809533 0.234914
       0.987488 0.814920 0.236539
       0.987049 0.820306 0.238209
       0.986580 0.825692 0.239950
       0.986083 0.831079 0.241755
       0.985559 0.836457 0.243582
       0.985008 0.841836 0.245471
       0.984427 0.847206 0.247432
       0.983815 0.852585 0.249403
       0.983176 0.857950 0.251454
       0.982514 0.863322 0.253514
       0.981819 0.868690 0.255622
       0.981095 0.874047 0.257764
       0.980345 0.879407 0.259965
       0.979565 0.884769 0.262191
       0.978754 0.890124 0.264447
       0.977917 0.895481 0.266749
       0.977055 0.900831 0.269086
       0.976155 0.906185 0.271441
       0.975233 0.911531 0.273841
       0.974279 0.916881 0.276284
       0.973300 0.922226 0.278730
       0.972289 0.927565 0.281202
       0.971243 0.932907 0.283727
       0.970173 0.938250 0.286250
       0.969069 0.943591 0.288848
       0.967943 0.948925 0.291435
       0.966774 0.954269 0.294053
       0.965579 0.959605 0.296711
       0.964356 0.964928 0.299378
       0.963103 0.970262 0.302058
       0.961816 0.975591 0.304771];

case { 'L9'  'BGYW' }
descriptorname = 'linear_bgyw_20-98_c66_n256';
description = 'Blue-green-yellow colour map';
map = [0.019721 0.000000 0.673123
       0.021792 0.009320 0.674573
       0.023764 0.020375 0.676023
       0.025723 0.031533 0.677444
       0.027593 0.042972 0.678843
       0.029434 0.053399 0.680216
       0.031200 0.062887 0.681577
       0.032926 0.071508 0.682918
       0.034668 0.079503 0.684227
       0.036359 0.087188 0.685529
       0.037918 0.094420 0.686790
       0.039376 0.101305 0.688043
       0.040814 0.107970 0.689257
       0.042148 0.114406 0.690448
       0.043258 0.120677 0.691614
       0.044531 0.126775 0.692760
       0.045545 0.132741 0.693866
       0.046606 0.138556 0.694948
       0.047504 0.144249 0.696016
       0.048422 0.149855 0.697032
       0.049286 0.155330 0.698026
       0.049957 0.160733 0.698993
       0.050622 0.166065 0.699920
       0.051213 0.171335 0.700825
       0.051756 0.176525 0.701681
       0.052246 0.181670 0.702510
       0.052659 0.186776 0.703310
       0.053041 0.191816 0.704056
       0.053359 0.196841 0.704768
       0.053600 0.201763 0.705450
       0.053810 0.206723 0.706079
       0.053957 0.211602 0.706667
       0.054028 0.216462 0.707214
       0.054057 0.221280 0.707708
       0.054033 0.226087 0.708155
       0.053937 0.230848 0.708561
       0.053775 0.235613 0.708916
       0.053580 0.240320 0.709204
       0.053321 0.245038 0.709438
       0.052992 0.249739 0.709619
       0.052597 0.254448 0.709744
       0.052169 0.259124 0.709792
       0.051684 0.263787 0.709775
       0.051133 0.268426 0.709693
       0.050513 0.273070 0.709541
       0.049841 0.277706 0.709315
       0.049147 0.282346 0.709000
       0.048250 0.286949 0.708594
       0.047380 0.291595 0.708096
       0.046417 0.296209 0.707507
       0.045403 0.300826 0.706823
       0.044335 0.305471 0.706025
       0.043076 0.310101 0.705111
       0.041991 0.314716 0.704076
       0.040637 0.319369 0.702911
       0.039245 0.324013 0.701593
       0.037801 0.328685 0.700131
       0.036303 0.333363 0.698496
       0.034665 0.338039 0.696686
       0.033000 0.342763 0.694672
       0.031386 0.347493 0.692448
       0.029770 0.352238 0.689951
       0.028127 0.357040 0.687188
       0.026459 0.361850 0.684122
       0.024810 0.366706 0.680696
       0.023175 0.371612 0.676862
       0.022142 0.376567 0.672638
       0.022449 0.381458 0.668154
       0.024367 0.386284 0.663474
       0.027722 0.391084 0.658599
       0.032448 0.395820 0.653549
       0.038520 0.400517 0.648330
       0.045286 0.405157 0.642941
       0.052276 0.409775 0.637391
       0.059484 0.414349 0.631687
       0.066532 0.418894 0.625826
       0.073577 0.423406 0.619832
       0.080364 0.427906 0.613680
       0.087130 0.432375 0.607409
       0.093557 0.436810 0.600990
       0.099853 0.441224 0.594451
       0.105987 0.445620 0.587776
       0.111858 0.450010 0.580988
       0.117453 0.454381 0.574078
       0.122861 0.458727 0.567057
       0.128129 0.463053 0.559908
       0.133113 0.467390 0.552646
       0.137893 0.471706 0.545279
       0.142403 0.476005 0.537807
       0.146724 0.480295 0.530212
       0.150863 0.484571 0.522508
       0.154770 0.488851 0.514709
       0.158487 0.493115 0.506793
       0.161968 0.497389 0.498785
       0.165186 0.501646 0.490669
       0.168265 0.505884 0.482435
       0.171135 0.510137 0.474094
       0.173792 0.514387 0.465640
       0.176217 0.518626 0.457091
       0.178500 0.522858 0.448431
       0.180501 0.527094 0.439652
       0.182320 0.531321 0.430761
       0.183948 0.535545 0.421739
       0.185386 0.539773 0.412622
       0.186558 0.544004 0.403359
       0.187572 0.548235 0.393960
       0.188341 0.552458 0.384425
       0.189035 0.556660 0.374907
       0.189889 0.560809 0.365600
       0.190954 0.564902 0.356547
       0.192251 0.568948 0.347725
       0.193793 0.572950 0.339131
       0.195533 0.576895 0.330729
       0.197538 0.580789 0.322560
       0.199758 0.584645 0.314548
       0.202249 0.588451 0.306744
       0.204983 0.592235 0.299098
       0.207945 0.595960 0.291610
       0.211141 0.599659 0.284255
       0.214556 0.603317 0.277069
       0.218209 0.606943 0.270007
       0.222058 0.610529 0.263058
       0.226180 0.614076 0.256234
       0.230491 0.617597 0.249560
       0.235054 0.621086 0.242989
       0.239755 0.624542 0.236541
       0.244639 0.627969 0.230125
       0.249711 0.631366 0.223833
       0.254986 0.634734 0.217627
       0.260432 0.638064 0.211539
       0.266047 0.641365 0.205496
       0.271769 0.644646 0.199506
       0.277667 0.647900 0.193617
       0.283675 0.651131 0.187776
       0.289872 0.654321 0.181979
       0.296184 0.657497 0.176260
       0.302586 0.660633 0.170604
       0.309130 0.663761 0.164943
       0.315789 0.666842 0.159383
       0.322561 0.669914 0.153861
       0.329418 0.672954 0.148352
       0.336338 0.675967 0.142883
       0.343412 0.678952 0.137484
       0.350573 0.681911 0.132085
       0.357790 0.684855 0.126671
       0.365068 0.687773 0.121265
       0.372482 0.690653 0.116032
       0.379957 0.693520 0.110723
       0.387478 0.696364 0.105363
       0.395102 0.699172 0.100096
       0.402791 0.701958 0.094912
       0.410539 0.704728 0.089608
       0.418327 0.707468 0.084378
       0.426219 0.710182 0.079119
       0.434143 0.712869 0.073963
       0.442136 0.715545 0.068745
       0.450197 0.718176 0.063586
       0.458290 0.720790 0.058350
       0.466436 0.723383 0.053154
       0.474669 0.725950 0.048139
       0.482912 0.728487 0.043005
       0.491223 0.731003 0.038082
       0.499578 0.733492 0.033323
       0.507987 0.735953 0.029083
       0.516420 0.738395 0.025269
       0.524931 0.740804 0.021961
       0.533456 0.743199 0.018981
       0.542041 0.745555 0.016456
       0.550674 0.747895 0.014369
       0.559335 0.750201 0.012644
       0.568048 0.752481 0.011320
       0.576796 0.754746 0.010293
       0.585535 0.756978 0.009763
       0.594250 0.759203 0.009522
       0.602908 0.761428 0.009379
       0.611494 0.763652 0.009384
       0.620009 0.765882 0.009549
       0.628459 0.768101 0.009832
       0.636840 0.770331 0.010273
       0.645147 0.772558 0.010973
       0.653404 0.774785 0.011934
       0.661602 0.777023 0.012885
       0.669729 0.779252 0.014059
       0.677802 0.781491 0.015391
       0.685807 0.783731 0.016941
       0.693746 0.785975 0.018714
       0.701638 0.788218 0.020670
       0.709477 0.790469 0.022856
       0.717243 0.792730 0.025320
       0.724951 0.794991 0.027999
       0.732606 0.797257 0.030918
       0.740194 0.799540 0.034117
       0.747721 0.801818 0.037833
       0.755191 0.804112 0.041619
       0.762597 0.806412 0.045386
       0.769942 0.808721 0.049415
       0.777225 0.811040 0.053367
       0.784442 0.813368 0.057479
       0.791592 0.815713 0.061666
       0.798682 0.818062 0.065966
       0.805691 0.820431 0.070321
       0.812633 0.822818 0.074765
       0.819503 0.825213 0.079243
       0.826307 0.827627 0.083887
       0.833031 0.830061 0.088519
       0.839682 0.832508 0.093204
       0.846238 0.834979 0.098097
       0.852714 0.837475 0.103017
       0.859094 0.839990 0.107993
       0.865385 0.842529 0.113165
       0.871585 0.845091 0.118365
       0.877678 0.847681 0.123615
       0.883670 0.850305 0.129043
       0.889545 0.852960 0.134620
       0.895306 0.855649 0.140239
       0.900939 0.858369 0.146038
       0.906444 0.861133 0.151930
       0.911805 0.863943 0.158007
       0.917021 0.866786 0.164237
       0.922074 0.869684 0.170588
       0.926941 0.872640 0.177165
       0.931621 0.875652 0.183920
       0.936105 0.878722 0.190889
       0.940370 0.881862 0.198073
       0.944398 0.885072 0.205502
       0.948129 0.888373 0.213229
       0.951580 0.891758 0.221264
       0.954701 0.895240 0.229571
       0.957433 0.898838 0.238318
       0.959791 0.902557 0.247542
       0.961985 0.906295 0.257711
       0.964250 0.909980 0.269181
       0.966576 0.913609 0.281810
       0.968964 0.917178 0.295751
       0.971400 0.920681 0.310838
       0.973879 0.924134 0.327055
       0.976396 0.927514 0.344535
       0.978934 0.930837 0.363141
       0.981479 0.934086 0.382912
       0.984012 0.937273 0.403922
       0.986513 0.940382 0.426212
       0.988951 0.943416 0.449799
       0.991297 0.946373 0.474734
       0.993515 0.949257 0.501089
       0.995550 0.952058 0.528967
       0.997358 0.954781 0.558446
       0.998872 0.957417 0.589673
       0.999999 0.959969 0.622765
       1.000000 0.962433 0.657860
       1.000000 0.964811 0.695105
       1.000000 0.967109 0.734701
       0.998411 0.969313 0.776944
       0.995642 0.971437 0.822026
       0.991455 0.973485 0.870116
       0.985442 0.975455 0.921874
       0.977176 0.977372 0.977324];

case { 'L10'  'GEOGRAPHIC' }
descriptorname = 'linear_gow_60-85_c27_n256';
description = 'A "geographical" colour map.  Best used with relief shading';
map = [0.398723 0.606241 0.564505
       0.401565 0.607207 0.562660
       0.404398 0.608160 0.560801
       0.407237 0.609111 0.558952
       0.410093 0.610046 0.557124
       0.412934 0.610993 0.555275
       0.415773 0.611923 0.553442
       0.418626 0.612850 0.551624
       0.421478 0.613776 0.549799
       0.424349 0.614696 0.547983
       0.427230 0.615613 0.546175
       0.430095 0.616530 0.544360
       0.432982 0.617430 0.542562
       0.435879 0.618332 0.540762
       0.438765 0.619231 0.538972
       0.441687 0.620117 0.537191
       0.444612 0.620995 0.535401
       0.447523 0.621863 0.533624
       0.450472 0.622738 0.531863
       0.453424 0.623605 0.530102
       0.456387 0.624459 0.528346
       0.459366 0.625306 0.526600
       0.462354 0.626147 0.524852
       0.465356 0.626982 0.523120
       0.468375 0.627815 0.521382
       0.471421 0.628637 0.519661
       0.474474 0.629443 0.517954
       0.477536 0.630254 0.516235
       0.480613 0.631054 0.514544
       0.483721 0.631839 0.512848
       0.486852 0.632618 0.511162
       0.489989 0.633391 0.509490
       0.493144 0.634159 0.507826
       0.496331 0.634907 0.506157
       0.499541 0.635648 0.504522
       0.502772 0.636388 0.502868
       0.506018 0.637111 0.501246
       0.509307 0.637822 0.499614
       0.512610 0.638524 0.498021
       0.515935 0.639216 0.496403
       0.519302 0.639900 0.494825
       0.522680 0.640568 0.493235
       0.526100 0.641219 0.491682
       0.529549 0.641867 0.490118
       0.533024 0.642501 0.488578
       0.536538 0.643116 0.487048
       0.540071 0.643719 0.485522
       0.543648 0.644310 0.484016
       0.547260 0.644886 0.482532
       0.550916 0.645446 0.481050
       0.554610 0.645991 0.479600
       0.558333 0.646522 0.478149
       0.562120 0.647035 0.476720
       0.565924 0.647527 0.475315
       0.569780 0.648000 0.473908
       0.573689 0.648459 0.472533
       0.577637 0.648901 0.471177
       0.581641 0.649320 0.469840
       0.585704 0.649715 0.468510
       0.589801 0.650087 0.467217
       0.593972 0.650436 0.465929
       0.598192 0.650761 0.464687
       0.602480 0.651063 0.463444
       0.606831 0.651338 0.462242
       0.611242 0.651586 0.461062
       0.615719 0.651804 0.459908
       0.620282 0.651990 0.458789
       0.624906 0.652143 0.457690
       0.629611 0.652263 0.456627
       0.634408 0.652348 0.455600
       0.639278 0.652396 0.454611
       0.644236 0.652406 0.453640
       0.649300 0.652375 0.452728
       0.654446 0.652301 0.451843
       0.659709 0.652180 0.451001
       0.665080 0.652007 0.450208
       0.670579 0.651782 0.449464
       0.676122 0.651521 0.448757
       0.681606 0.651261 0.448051
       0.686988 0.651022 0.447357
       0.692257 0.650806 0.446672
       0.697405 0.650611 0.445975
       0.702452 0.650440 0.445291
       0.707385 0.650294 0.444613
       0.712209 0.650172 0.443924
       0.716941 0.650074 0.443237
       0.721551 0.650000 0.442556
       0.726070 0.649954 0.441874
       0.730480 0.649933 0.441181
       0.734796 0.649937 0.440499
       0.739012 0.649968 0.439824
       0.743122 0.650027 0.439137
       0.747134 0.650114 0.438462
       0.751059 0.650227 0.437793
       0.754882 0.650369 0.437113
       0.758605 0.650541 0.436433
       0.762232 0.650742 0.435759
       0.765778 0.650972 0.435084
       0.769219 0.651232 0.434399
       0.772572 0.651524 0.433718
       0.775832 0.651845 0.433051
       0.779004 0.652193 0.432386
       0.782087 0.652571 0.431712
       0.785073 0.652982 0.431032
       0.787965 0.653430 0.430355
       0.790776 0.653909 0.429680
       0.793502 0.654415 0.429006
       0.796124 0.654954 0.428333
       0.798667 0.655533 0.427665
       0.801121 0.656143 0.426989
       0.803487 0.656780 0.426305
       0.805762 0.657454 0.425636
       0.807953 0.658157 0.424966
       0.810055 0.658888 0.424285
       0.812074 0.659664 0.423606
       0.813999 0.660467 0.422929
       0.815844 0.661310 0.422252
       0.817598 0.662187 0.421577
       0.819266 0.663092 0.420902
       0.820849 0.664031 0.420229
       0.822348 0.665001 0.419558
       0.823755 0.666011 0.418873
       0.825080 0.667051 0.418191
       0.826318 0.668135 0.417522
       0.827466 0.669244 0.416841
       0.828534 0.670391 0.416155
       0.829517 0.671561 0.415473
       0.830406 0.672780 0.414792
       0.831232 0.674012 0.414137
       0.832023 0.675259 0.413553
       0.832791 0.676509 0.413067
       0.833550 0.677761 0.412680
       0.834290 0.679005 0.412386
       0.835011 0.680254 0.412186
       0.835718 0.681507 0.412085
       0.836408 0.682765 0.412089
       0.837084 0.684017 0.412192
       0.837744 0.685280 0.412392
       0.838383 0.686534 0.412690
       0.839008 0.687794 0.413088
       0.839621 0.689057 0.413582
       0.840215 0.690311 0.414174
       0.840790 0.691574 0.414869
       0.841350 0.692842 0.415668
       0.841894 0.694095 0.416570
       0.842423 0.695366 0.417566
       0.842931 0.696632 0.418652
       0.843421 0.697896 0.419858
       0.843898 0.699163 0.421148
       0.844360 0.700437 0.422541
       0.844803 0.701702 0.424040
       0.845226 0.702981 0.425643
       0.845630 0.704250 0.427349
       0.846018 0.705525 0.429135
       0.846388 0.706804 0.431033
       0.846740 0.708076 0.433026
       0.847072 0.709358 0.435130
       0.847387 0.710635 0.437320
       0.847682 0.711914 0.439604
       0.847956 0.713193 0.442006
       0.848211 0.714474 0.444497
       0.848447 0.715767 0.447073
       0.848661 0.717053 0.449762
       0.848854 0.718333 0.452556
       0.849026 0.719626 0.455433
       0.849176 0.720910 0.458407
       0.849303 0.722201 0.461487
       0.849406 0.723496 0.464667
       0.849486 0.724796 0.467931
       0.849543 0.726091 0.471304
       0.849576 0.727386 0.474769
       0.849582 0.728686 0.478324
       0.849564 0.729991 0.481987
       0.849520 0.731289 0.485730
       0.849450 0.732601 0.489574
       0.849351 0.733905 0.493523
       0.849224 0.735215 0.497572
       0.849069 0.736520 0.501699
       0.848886 0.737833 0.505913
       0.848682 0.739156 0.510219
       0.848478 0.740459 0.514541
       0.848284 0.741760 0.518854
       0.848097 0.743064 0.523156
       0.847919 0.744352 0.527452
       0.847749 0.745639 0.531746
       0.847586 0.746925 0.536032
       0.847427 0.748205 0.540317
       0.847274 0.749473 0.544598
       0.847125 0.750753 0.548879
       0.846980 0.752018 0.553140
       0.846838 0.753278 0.557418
       0.846698 0.754546 0.561680
       0.846562 0.755801 0.565935
       0.846427 0.757055 0.570191
       0.846293 0.758310 0.574444
       0.846161 0.759553 0.578701
       0.846030 0.760806 0.582947
       0.845899 0.762047 0.587194
       0.845769 0.763290 0.591443
       0.845638 0.764533 0.595683
       0.845507 0.765773 0.599925
       0.845375 0.767005 0.604167
       0.845242 0.768238 0.608404
       0.845108 0.769472 0.612628
       0.844973 0.770707 0.616868
       0.844836 0.771936 0.621098
       0.844696 0.773160 0.625325
       0.844554 0.774386 0.629552
       0.844409 0.775612 0.633786
       0.844261 0.776841 0.638008
       0.844110 0.778061 0.642235
       0.843957 0.779280 0.646458
       0.843801 0.780500 0.650681
       0.843641 0.781722 0.654899
       0.843479 0.782935 0.659122
       0.843313 0.784159 0.663352
       0.843144 0.785372 0.667570
       0.842972 0.786589 0.671786
       0.842795 0.787797 0.676007
       0.842613 0.789015 0.680221
       0.842426 0.790223 0.684441
       0.842235 0.791434 0.688668
       0.842039 0.792647 0.692888
       0.841837 0.793861 0.697101
       0.841631 0.795065 0.701319
       0.841420 0.796272 0.705539
       0.841204 0.797480 0.709760
       0.840983 0.798691 0.713976
       0.840756 0.799901 0.718195
       0.840525 0.801103 0.722411
       0.840288 0.802308 0.726635
       0.840045 0.803515 0.730854
       0.839796 0.804722 0.735078
       0.839540 0.805925 0.739302
       0.839276 0.807133 0.743520
       0.839006 0.808335 0.747743
       0.838732 0.809540 0.751963
       0.838452 0.810740 0.756184
       0.838166 0.811945 0.760408
       0.837875 0.813145 0.764632
       0.837576 0.814348 0.768853
       0.837268 0.815554 0.773079
       0.836954 0.816757 0.777312
       0.836632 0.817955 0.781537
       0.836303 0.819156 0.785765
       0.835969 0.820358 0.789991
       0.835627 0.821564 0.794224
       0.835277 0.822765 0.798451
       0.834921 0.823963 0.802681
       0.834558 0.825164 0.806918
       0.834187 0.826369 0.811149
       0.833808 0.827568 0.815384
       0.833417 0.828773 0.819618
       0.833020 0.829973 0.823855
       0.832617 0.831175 0.828091
       0.832208 0.832374 0.832333];

case { 'L11'  'GEOGRAPHIC2' }
descriptorname = 'linear_gow_65-90_c35_n256';
description = 'A lighter "geographical" colour map.  Best used with relief shading';
map = [0.437651 0.678142 0.360542
       0.443142 0.678442 0.360996
       0.448595 0.678738 0.361458
       0.453990 0.679033 0.361916
       0.459344 0.679324 0.362367
       0.464653 0.679613 0.362817
       0.469909 0.679901 0.363272
       0.475130 0.680184 0.363728
       0.480292 0.680466 0.364186
       0.485421 0.680746 0.364645
       0.490523 0.681023 0.365105
       0.495569 0.681297 0.365568
       0.500589 0.681569 0.366035
       0.505559 0.681839 0.366499
       0.510515 0.682106 0.366955
       0.515419 0.682373 0.367412
       0.520298 0.682636 0.367878
       0.525147 0.682897 0.368352
       0.529962 0.683155 0.368821
       0.534743 0.683410 0.369283
       0.539489 0.683661 0.369746
       0.544218 0.683910 0.370213
       0.548915 0.684158 0.370682
       0.553573 0.684404 0.371152
       0.558206 0.684651 0.371624
       0.562832 0.684894 0.372097
       0.567410 0.685138 0.372572
       0.571968 0.685376 0.373048
       0.576506 0.685613 0.373528
       0.581006 0.685847 0.374010
       0.585496 0.686078 0.374487
       0.589945 0.686308 0.374958
       0.594385 0.686537 0.375433
       0.598797 0.686765 0.375921
       0.603189 0.686990 0.376409
       0.607559 0.687215 0.376889
       0.611898 0.687437 0.377367
       0.616222 0.687659 0.377848
       0.620529 0.687878 0.378333
       0.624802 0.688097 0.378818
       0.629061 0.688315 0.379305
       0.633299 0.688532 0.379794
       0.637517 0.688747 0.380285
       0.641709 0.688961 0.380780
       0.645886 0.689173 0.381271
       0.650045 0.689383 0.381755
       0.654180 0.689592 0.382242
       0.658298 0.689800 0.382742
       0.662400 0.690007 0.383243
       0.666468 0.690213 0.383736
       0.670536 0.690419 0.384227
       0.674570 0.690624 0.384722
       0.678589 0.690829 0.385220
       0.682591 0.691035 0.385719
       0.686573 0.691242 0.386220
       0.690536 0.691448 0.386723
       0.694476 0.691656 0.387228
       0.698407 0.691862 0.387733
       0.702313 0.692070 0.388244
       0.706208 0.692276 0.388756
       0.710075 0.692482 0.389261
       0.713923 0.692688 0.389762
       0.717761 0.692893 0.390272
       0.721566 0.693099 0.390793
       0.725372 0.693304 0.391308
       0.729139 0.693512 0.391817
       0.732901 0.693719 0.392328
       0.736633 0.693928 0.392843
       0.740354 0.694139 0.393361
       0.744054 0.694352 0.393879
       0.747735 0.694569 0.394399
       0.751399 0.694787 0.394923
       0.755037 0.695007 0.395451
       0.758654 0.695230 0.395974
       0.762253 0.695454 0.396491
       0.765836 0.695682 0.397014
       0.769391 0.695911 0.397549
       0.772931 0.696141 0.398083
       0.776451 0.696375 0.398609
       0.779946 0.696611 0.399133
       0.783422 0.696850 0.399665
       0.786877 0.697094 0.400211
       0.790304 0.697341 0.400758
       0.793717 0.697594 0.401288
       0.797092 0.697851 0.401815
       0.800459 0.698112 0.402357
       0.803794 0.698379 0.402907
       0.807107 0.698651 0.403457
       0.810389 0.698929 0.403999
       0.813653 0.699212 0.404539
       0.816890 0.699503 0.405093
       0.820093 0.699801 0.405653
       0.823279 0.700105 0.406206
       0.826433 0.700418 0.406756
       0.829556 0.700739 0.407312
       0.832645 0.701066 0.407871
       0.835710 0.701399 0.408432
       0.838740 0.701742 0.408996
       0.841739 0.702100 0.409565
       0.844701 0.702470 0.410138
       0.847626 0.702850 0.410708
       0.850520 0.703238 0.411272
       0.853375 0.703637 0.411847
       0.856192 0.704051 0.412434
       0.858961 0.704480 0.413014
       0.861692 0.704924 0.413592
       0.864378 0.705383 0.414176
       0.867013 0.705860 0.414765
       0.869604 0.706356 0.415357
       0.872146 0.706868 0.415953
       0.874626 0.707396 0.416555
       0.877050 0.707950 0.417163
       0.879416 0.708531 0.417765
       0.881719 0.709134 0.418368
       0.883951 0.709760 0.418990
       0.886107 0.710417 0.419615
       0.888182 0.711107 0.420234
       0.890174 0.711827 0.420861
       0.892080 0.712578 0.421496
       0.893883 0.713377 0.422137
       0.895582 0.714213 0.422787
       0.897159 0.715104 0.423445
       0.898597 0.716048 0.424113
       0.899905 0.717044 0.424794
       0.901042 0.718101 0.425476
       0.901995 0.719237 0.426163
       0.902744 0.720450 0.426879
       0.903249 0.721762 0.427607
       0.903554 0.723156 0.428357
       0.903775 0.724582 0.429170
       0.903992 0.726002 0.430057
       0.904204 0.727415 0.431018
       0.904412 0.728829 0.432058
       0.904615 0.730245 0.433150
       0.904814 0.731652 0.434325
       0.905007 0.733061 0.435588
       0.905194 0.734467 0.436910
       0.905375 0.735864 0.438304
       0.905550 0.737268 0.439775
       0.905718 0.738669 0.441315
       0.905881 0.740068 0.442936
       0.906038 0.741455 0.444629
       0.906188 0.742855 0.446381
       0.906333 0.744240 0.448215
       0.906471 0.745628 0.450125
       0.906602 0.747017 0.452111
       0.906727 0.748401 0.454157
       0.906844 0.749777 0.456277
       0.906955 0.751168 0.458474
       0.907058 0.752537 0.460736
       0.907154 0.753918 0.463074
       0.907242 0.755294 0.465486
       0.907322 0.756664 0.467974
       0.907393 0.758037 0.470537
       0.907456 0.759400 0.473151
       0.907511 0.760773 0.475860
       0.907556 0.762134 0.478624
       0.907593 0.763499 0.481462
       0.907619 0.764865 0.484368
       0.907636 0.766224 0.487362
       0.907643 0.767579 0.490411
       0.907640 0.768936 0.493525
       0.907626 0.770295 0.496716
       0.907601 0.771652 0.499976
       0.907565 0.773000 0.503308
       0.907517 0.774350 0.506715
       0.907457 0.775702 0.510186
       0.907385 0.777056 0.513734
       0.907300 0.778399 0.517347
       0.907203 0.779743 0.521020
       0.907091 0.781089 0.524769
       0.906967 0.782433 0.528593
       0.906828 0.783777 0.532478
       0.906675 0.785117 0.536434
       0.906507 0.786459 0.540456
       0.906323 0.787791 0.544548
       0.906124 0.789133 0.548715
       0.905909 0.790465 0.552935
       0.905678 0.791801 0.557243
       0.905436 0.793139 0.561587
       0.905197 0.794466 0.565944
       0.904968 0.795784 0.570297
       0.904747 0.797101 0.574645
       0.904535 0.798414 0.578998
       0.904331 0.799726 0.583333
       0.904134 0.801023 0.587667
       0.903944 0.802319 0.592008
       0.903760 0.803614 0.596330
       0.903581 0.804904 0.600651
       0.903406 0.806192 0.604973
       0.903234 0.807474 0.609293
       0.903066 0.808755 0.613596
       0.902899 0.810027 0.617910
       0.902735 0.811303 0.622214
       0.902572 0.812570 0.626522
       0.902412 0.813837 0.630830
       0.902252 0.815104 0.635123
       0.902093 0.816370 0.639422
       0.901935 0.817625 0.643712
       0.901778 0.818882 0.648004
       0.901620 0.820138 0.652299
       0.901461 0.821396 0.656592
       0.901303 0.822649 0.660872
       0.901143 0.823895 0.665158
       0.900983 0.825143 0.669447
       0.900820 0.826393 0.673727
       0.900656 0.827635 0.678011
       0.900491 0.828884 0.682285
       0.900324 0.830124 0.686568
       0.900154 0.831367 0.690841
       0.899983 0.832602 0.695122
       0.899809 0.833843 0.699396
       0.899632 0.835077 0.703675
       0.899451 0.836313 0.707948
       0.899267 0.837550 0.712221
       0.899079 0.838778 0.716501
       0.898887 0.840016 0.720768
       0.898692 0.841245 0.725047
       0.898493 0.842477 0.729315
       0.898291 0.843701 0.733588
       0.898086 0.844933 0.737856
       0.897877 0.846157 0.742131
       0.897664 0.847384 0.746401
       0.897447 0.848613 0.750677
       0.897224 0.849841 0.754949
       0.896996 0.851060 0.759214
       0.896763 0.852292 0.763486
       0.896524 0.853508 0.767756
       0.896281 0.854736 0.772031
       0.896033 0.855958 0.776304
       0.895780 0.857177 0.780574
       0.895522 0.858398 0.784851
       0.895258 0.859622 0.789124
       0.894989 0.860841 0.793398
       0.894715 0.862065 0.797666
       0.894433 0.863285 0.801940
       0.894145 0.864502 0.806218
       0.893850 0.865719 0.810489
       0.893549 0.866936 0.814766
       0.893244 0.868161 0.819041
       0.892934 0.869375 0.823321
       0.892617 0.870589 0.827598
       0.892294 0.871813 0.831882
       0.891962 0.873027 0.836157
       0.891623 0.874243 0.840440
       0.891278 0.875462 0.844719
       0.890927 0.876674 0.849001
       0.890568 0.877895 0.853280
       0.890203 0.879109 0.857564
       0.889831 0.880325 0.861853
       0.889452 0.881544 0.866137
       0.889067 0.882756 0.870419
       0.888674 0.883977 0.874709
       0.888270 0.885191 0.878999
       0.887859 0.886409 0.883287
       0.887441 0.887619 0.887576];

case { 'L12'  'DEPTH' }
descriptorname = 'linear_blue_95-50_c20_n256';
description = 'A "water depth" colour map';
map = [0.943337 0.943525 0.943479
       0.940360 0.941658 0.943250
       0.937378 0.939796 0.943020
       0.934395 0.937930 0.942785
       0.931415 0.936069 0.942549
       0.928432 0.934201 0.942307
       0.925452 0.932339 0.942064
       0.922472 0.930482 0.941815
       0.919485 0.928617 0.941566
       0.916502 0.926757 0.941313
       0.913520 0.924897 0.941059
       0.910535 0.923043 0.940802
       0.907550 0.921181 0.940543
       0.904560 0.919324 0.940279
       0.901573 0.917474 0.940012
       0.898585 0.915610 0.939741
       0.895599 0.913762 0.939467
       0.892611 0.911904 0.939190
       0.889620 0.910053 0.938909
       0.886632 0.908200 0.938627
       0.883642 0.906346 0.938339
       0.880647 0.904492 0.938051
       0.877657 0.902643 0.937757
       0.874664 0.900791 0.937461
       0.871675 0.898942 0.937162
       0.868684 0.897095 0.936859
       0.865685 0.895246 0.936553
       0.862696 0.893394 0.936242
       0.859698 0.891549 0.935927
       0.856701 0.889701 0.935607
       0.853702 0.887854 0.935285
       0.850708 0.886013 0.934958
       0.847711 0.884168 0.934629
       0.844718 0.882320 0.934296
       0.841722 0.880477 0.933958
       0.838720 0.878635 0.933618
       0.835725 0.876790 0.933273
       0.832725 0.874952 0.932925
       0.829730 0.873112 0.932572
       0.826731 0.871272 0.932216
       0.823728 0.869432 0.931856
       0.820729 0.867594 0.931491
       0.817728 0.865755 0.931124
       0.814729 0.863922 0.930751
       0.811732 0.862084 0.930372
       0.808733 0.860246 0.929989
       0.805729 0.858406 0.929599
       0.802729 0.856572 0.929207
       0.799735 0.854740 0.928810
       0.796729 0.852905 0.928408
       0.793736 0.851067 0.928002
       0.790730 0.849240 0.927591
       0.787730 0.847403 0.927174
       0.784736 0.845572 0.926754
       0.781736 0.843740 0.926329
       0.778734 0.841914 0.925898
       0.775735 0.840086 0.925460
       0.772736 0.838253 0.925016
       0.769739 0.836424 0.924567
       0.766742 0.834597 0.924116
       0.763746 0.832766 0.923659
       0.760754 0.830942 0.923193
       0.757759 0.829117 0.922721
       0.754768 0.827287 0.922244
       0.751775 0.825459 0.921759
       0.748779 0.823634 0.921266
       0.745789 0.821813 0.920767
       0.742805 0.819983 0.920265
       0.739820 0.818159 0.919756
       0.736828 0.816341 0.919237
       0.733844 0.814513 0.918713
       0.730859 0.812690 0.918184
       0.727879 0.810868 0.917647
       0.724906 0.809051 0.917100
       0.721922 0.807228 0.916543
       0.718952 0.805402 0.915976
       0.715987 0.803583 0.915405
       0.713011 0.801761 0.914828
       0.710050 0.799946 0.914243
       0.707088 0.798122 0.913646
       0.704127 0.796301 0.913038
       0.701173 0.794485 0.912421
       0.698220 0.792665 0.911794
       0.695276 0.790844 0.911159
       0.692340 0.789029 0.910515
       0.689401 0.787209 0.909857
       0.686464 0.785390 0.909186
       0.683537 0.783571 0.908509
       0.680612 0.781754 0.907820
       0.677708 0.779931 0.907114
       0.674796 0.778115 0.906396
       0.671896 0.776297 0.905668
       0.669006 0.774474 0.904922
       0.666118 0.772656 0.904161
       0.663250 0.770841 0.903391
       0.660377 0.769016 0.902602
       0.657528 0.767197 0.901795
       0.654674 0.765384 0.900972
       0.651848 0.763558 0.900134
       0.649028 0.761737 0.899276
       0.646216 0.759915 0.898394
       0.643421 0.758098 0.897499
       0.640646 0.756273 0.896578
       0.637887 0.754453 0.895632
       0.635144 0.752621 0.894664
       0.632425 0.750803 0.893663
       0.629722 0.748970 0.892642
       0.627045 0.747146 0.891589
       0.624398 0.745313 0.890502
       0.621768 0.743485 0.889382
       0.619187 0.741647 0.888226
       0.616627 0.739821 0.887028
       0.614099 0.737975 0.885788
       0.611622 0.736132 0.884498
       0.609177 0.734294 0.883167
       0.606742 0.732452 0.881822
       0.604322 0.730605 0.880462
       0.601901 0.728760 0.879093
       0.599501 0.726917 0.877716
       0.597106 0.725079 0.876323
       0.594714 0.723229 0.874924
       0.592345 0.721385 0.873512
       0.589965 0.719548 0.872093
       0.587607 0.717706 0.870654
       0.585267 0.715865 0.869215
       0.582918 0.714015 0.867760
       0.580592 0.712172 0.866293
       0.578266 0.710332 0.864815
       0.575967 0.708492 0.863333
       0.573658 0.706650 0.861834
       0.571363 0.704803 0.860326
       0.569080 0.702963 0.858804
       0.566816 0.701118 0.857274
       0.564548 0.699273 0.855738
       0.562304 0.697432 0.854183
       0.560057 0.695594 0.852628
       0.557820 0.693747 0.851049
       0.555592 0.691911 0.849473
       0.553381 0.690066 0.847875
       0.551185 0.688226 0.846272
       0.548997 0.686382 0.844662
       0.546813 0.684537 0.843036
       0.544639 0.682698 0.841403
       0.542480 0.680852 0.839760
       0.540326 0.679014 0.838102
       0.538193 0.677180 0.836435
       0.536054 0.675333 0.834760
       0.533931 0.673494 0.833071
       0.531829 0.671650 0.831379
       0.529733 0.669815 0.829670
       0.527641 0.667974 0.827947
       0.525567 0.666127 0.826221
       0.523503 0.664288 0.824481
       0.521441 0.662453 0.822735
       0.519402 0.660604 0.820974
       0.517373 0.658764 0.819202
       0.515340 0.656931 0.817425
       0.513335 0.655083 0.815638
       0.511329 0.653242 0.813834
       0.509345 0.651409 0.812026
       0.507366 0.649571 0.810203
       0.505390 0.647725 0.808377
       0.503436 0.645885 0.806538
       0.501502 0.644044 0.804685
       0.499557 0.642210 0.802822
       0.497650 0.640372 0.800952
       0.495727 0.638530 0.799074
       0.493831 0.636696 0.797178
       0.491949 0.634855 0.795280
       0.490066 0.633016 0.793374
       0.488203 0.631179 0.791451
       0.486310 0.629337 0.789553
       0.484386 0.627507 0.787688
       0.482434 0.625678 0.785868
       0.480433 0.623861 0.784086
       0.478400 0.622030 0.782334
       0.476336 0.620224 0.780617
       0.474235 0.618409 0.778937
       0.472108 0.616600 0.777293
       0.469947 0.614786 0.775668
       0.467747 0.612983 0.774076
       0.465517 0.611192 0.772517
       0.463268 0.609395 0.770985
       0.460998 0.607604 0.769470
       0.458697 0.605802 0.767985
       0.456364 0.604026 0.766528
       0.454013 0.602235 0.765095
       0.451637 0.600453 0.763675
       0.449234 0.598673 0.762280
       0.446802 0.596902 0.760912
       0.444357 0.595124 0.759556
       0.441882 0.593356 0.758230
       0.439374 0.591588 0.756913
       0.436861 0.589814 0.755619
       0.434311 0.588050 0.754344
       0.431761 0.586296 0.753076
       0.429165 0.584536 0.751839
       0.426554 0.582776 0.750612
       0.423922 0.581024 0.749394
       0.421268 0.579278 0.748206
       0.418596 0.577521 0.747024
       0.415906 0.575784 0.745856
       0.413198 0.574029 0.744705
       0.410465 0.572292 0.743570
       0.407696 0.570545 0.742446
       0.404913 0.568807 0.741328
       0.402108 0.567079 0.740236
       0.399286 0.565333 0.739154
       0.396448 0.563614 0.738071
       0.393583 0.561884 0.737013
       0.390701 0.560154 0.735958
       0.387785 0.558418 0.734927
       0.384853 0.556703 0.733896
       0.381902 0.554978 0.732881
       0.378928 0.553254 0.731874
       0.375931 0.551544 0.730875
       0.372902 0.549827 0.729895
       0.369855 0.548117 0.728916
       0.366794 0.546407 0.727950
       0.363691 0.544691 0.726994
       0.360566 0.542986 0.726051
       0.357446 0.541271 0.725115
       0.354264 0.539567 0.724181
       0.351077 0.537876 0.723258
       0.347854 0.536170 0.722345
       0.344597 0.534471 0.721443
       0.341326 0.532778 0.720550
       0.338013 0.531081 0.719669
       0.334698 0.529393 0.718786
       0.331312 0.527695 0.717918
       0.327927 0.526011 0.717058
       0.324500 0.524320 0.716201
       0.321054 0.522634 0.715352
       0.317561 0.520951 0.714505
       0.314040 0.519273 0.713675
       0.310494 0.517596 0.712844
       0.306890 0.515904 0.712027
       0.303249 0.514242 0.711217
       0.299595 0.512559 0.710409
       0.295877 0.510886 0.709610
       0.292134 0.509220 0.708819
       0.288339 0.507551 0.708026
       0.284501 0.505874 0.707245
       0.280612 0.504224 0.706474
       0.276707 0.502550 0.705702
       0.272717 0.500893 0.704937
       0.268704 0.499227 0.704179
       0.264608 0.497581 0.703429
       0.260473 0.495911 0.702683
       0.256265 0.494266 0.701937
       0.252025 0.492603 0.701205
       0.247705 0.490969 0.700476
       0.243288 0.489303 0.699749
       0.238828 0.487668 0.699029
       0.234308 0.486013 0.698315
       0.229665 0.484366 0.697605];

case { 'L13'  'REDTERNARY' }
descriptorname = 'linear_ternary-red_0-50_c52_n256';
description = 'red colour map for ternary images';
map = [0.000000 0.000000 0.000000
       0.008025 0.001515 0.000000
       0.016166 0.003054 0.000000
       0.024126 0.004557 0.000000
       0.032164 0.006081 0.000000
       0.040345 0.007616 0.000000
       0.047917 0.009057 0.000000
       0.055209 0.010404 0.000000
       0.062360 0.011803 0.000000
       0.068888 0.012988 0.000000
       0.074917 0.014153 0.000000
       0.080792 0.015256 0.000000
       0.086386 0.016319 0.000000
       0.091832 0.017341 0.000000
       0.096947 0.018317 0.000000
       0.101991 0.019262 0.000000
       0.106882 0.020182 0.000000
       0.111562 0.021071 0.000000
       0.116123 0.021929 0.000000
       0.120506 0.022761 0.000000
       0.124822 0.023572 0.000000
       0.129014 0.024369 0.000000
       0.133121 0.025144 0.000000
       0.137116 0.025899 0.000000
       0.141037 0.026637 0.000000
       0.144836 0.027362 0.000000
       0.148518 0.028076 0.000000
       0.152181 0.028765 0.000000
       0.155795 0.029444 0.000000
       0.159289 0.030110 0.000000
       0.162745 0.030760 0.000000
       0.166118 0.031401 0.000000
       0.169474 0.032031 0.000000
       0.172713 0.032655 0.000000
       0.175901 0.033257 0.000000
       0.179087 0.033824 0.000000
       0.182228 0.034436 0.000000
       0.185421 0.035067 0.000000
       0.188468 0.035630 0.000000
       0.191419 0.036170 0.000000
       0.194369 0.036729 0.000000
       0.197251 0.037267 0.000001
       0.200116 0.037802 0.000001
       0.203028 0.038349 0.000001
       0.205942 0.038898 0.000001
       0.208838 0.039449 0.000001
       0.211764 0.039999 0.000001
       0.214662 0.040546 0.000001
       0.217610 0.041100 0.000001
       0.220569 0.041669 0.000001
       0.223463 0.042219 0.000001
       0.226347 0.042741 0.000001
       0.229245 0.043284 0.000001
       0.232226 0.043860 0.000001
       0.235221 0.044439 0.000001
       0.238148 0.045002 0.000001
       0.241064 0.045522 0.000001
       0.244011 0.046076 0.000001
       0.247008 0.046672 0.000001
       0.249954 0.047232 0.000001
       0.252930 0.047749 0.000001
       0.255910 0.048339 0.000001
       0.258943 0.048928 0.000001
       0.261906 0.049479 0.000001
       0.264862 0.050025 0.000001
       0.267829 0.050590 0.000001
       0.270843 0.051156 0.000002
       0.273831 0.051721 0.000001
       0.276825 0.052289 0.000001
       0.279830 0.052858 0.000001
       0.282842 0.053424 0.000001
       0.285840 0.053993 0.000001
       0.288879 0.054565 0.000002
       0.291905 0.055134 0.000002
       0.294935 0.055708 0.000001
       0.297985 0.056296 0.000001
       0.301006 0.056866 0.000001
       0.304010 0.057414 0.000001
       0.307051 0.057986 0.000001
       0.310132 0.058580 0.000002
       0.313201 0.059170 0.000002
       0.316275 0.059754 0.000001
       0.319309 0.060320 0.000001
       0.322337 0.060876 0.000001
       0.325377 0.061448 0.000001
       0.328485 0.062046 0.000001
       0.331574 0.062650 0.000001
       0.334666 0.063220 0.000002
       0.337720 0.063788 0.000002
       0.340804 0.064374 0.000001
       0.343876 0.064956 0.000001
       0.346984 0.065538 0.000001
       0.350077 0.066125 0.000001
       0.353178 0.066712 0.000001
       0.356288 0.067299 0.000001
       0.359393 0.067883 0.000002
       0.362521 0.068473 0.000002
       0.365650 0.069079 0.000002
       0.368758 0.069658 0.000001
       0.371846 0.070223 0.000001
       0.374984 0.070835 0.000001
       0.378125 0.071435 0.000001
       0.381246 0.072005 0.000001
       0.384378 0.072592 0.000002
       0.387536 0.073217 0.000002
       0.390689 0.073807 0.000002
       0.393819 0.074384 0.000002
       0.396954 0.074981 0.000001
       0.400109 0.075576 0.000001
       0.403271 0.076172 0.000001
       0.406426 0.076769 0.000001
       0.409593 0.077367 0.000001
       0.412770 0.077967 0.000001
       0.415937 0.078566 0.000002
       0.419122 0.079167 0.000002
       0.422300 0.079769 0.000002
       0.425503 0.080371 0.000002
       0.428701 0.080974 0.000001
       0.431921 0.081588 0.000001
       0.435119 0.082195 0.000001
       0.438283 0.082780 0.000001
       0.441496 0.083388 0.000001
       0.444722 0.084012 0.000001
       0.447914 0.084617 0.000002
       0.451115 0.085205 0.000002
       0.454327 0.085801 0.000002
       0.457558 0.086427 0.000002
       0.460799 0.087056 0.000002
       0.464040 0.087652 0.000002
       0.467267 0.088257 0.000001
       0.470494 0.088871 0.000001
       0.473727 0.089482 0.000001
       0.476974 0.090095 0.000001
       0.480225 0.090707 0.000001
       0.483481 0.091327 0.000001
       0.486734 0.091949 0.000002
       0.489964 0.092545 0.000002
       0.493214 0.093154 0.000002
       0.496489 0.093784 0.000002
       0.499773 0.094411 0.000002
       0.503048 0.095031 0.000002
       0.506311 0.095640 0.000002
       0.509570 0.096244 0.000001
       0.512849 0.096865 0.000001
       0.516155 0.097503 0.000001
       0.519451 0.098123 0.000001
       0.522735 0.098737 0.000001
       0.526027 0.099359 0.000001
       0.529324 0.099983 0.000001
       0.532624 0.100606 0.000001
       0.535921 0.101230 0.000002
       0.539230 0.101854 0.000002
       0.542554 0.102479 0.000002
       0.545879 0.103114 0.000002
       0.549190 0.103745 0.000002
       0.552493 0.104350 0.000002
       0.555803 0.104980 0.000002
       0.559150 0.105620 0.000002
       0.562508 0.106255 0.000001
       0.565833 0.106885 0.000001
       0.569154 0.107506 0.000001
       0.572486 0.108128 0.000001
       0.575845 0.108769 0.000001
       0.579206 0.109409 0.000001
       0.582555 0.110041 0.000001
       0.585905 0.110669 0.000001
       0.589252 0.111304 0.000001
       0.592623 0.111938 0.000001
       0.595985 0.112574 0.000002
       0.599357 0.113219 0.000002
       0.602706 0.113846 0.000002
       0.606061 0.114470 0.000002
       0.609450 0.115116 0.000002
       0.612837 0.115764 0.000002
       0.616225 0.116406 0.000002
       0.619604 0.117026 0.000002
       0.622993 0.117673 0.000002
       0.626392 0.118330 0.000002
       0.629782 0.118961 0.000002
       0.633173 0.119597 0.000002
       0.636573 0.120241 0.000001
       0.639973 0.120883 0.000001
       0.643369 0.121525 0.000001
       0.646787 0.122170 0.000001
       0.650195 0.122814 0.000001
       0.653616 0.123460 0.000001
       0.657039 0.124107 0.000001
       0.660462 0.124754 0.000001
       0.663903 0.125405 0.000001
       0.667336 0.126054 0.000001
       0.670749 0.126694 0.000000
       0.674188 0.127345 0.000000
       0.677636 0.128002 0.000000
       0.681057 0.128643 0.000000
       0.684508 0.129292 0.000000
       0.687974 0.129955 0.000000
       0.691419 0.130605 0.000000
       0.694864 0.131251 0.000000
       0.698318 0.131904 0.000000
       0.701774 0.132558 0.000000
       0.705239 0.133211 0.000000
       0.708711 0.133866 0.000000
       0.712179 0.134526 0.000000
       0.715647 0.135181 0.000000
       0.719101 0.135823 0.000000
       0.722576 0.136484 0.000000
       0.726074 0.137148 0.000000
       0.729563 0.137812 0.000000
       0.733042 0.138466 0.000000
       0.736511 0.139115 0.000000
       0.740008 0.139770 0.000000
       0.743513 0.140443 0.000000
       0.747014 0.141109 0.000000
       0.750510 0.141761 0.000000
       0.754010 0.142423 0.000000
       0.757513 0.143086 0.000000
       0.761022 0.143748 0.000000
       0.764536 0.144410 0.000000
       0.768052 0.145079 0.000000
       0.771571 0.145747 0.000000
       0.775075 0.146396 0.000000
       0.778601 0.147068 0.000000
       0.782130 0.147743 0.000000
       0.785650 0.148393 0.000000
       0.789189 0.149067 0.000000
       0.792727 0.149745 0.000000
       0.796254 0.150404 0.000000
       0.799795 0.151070 0.000000
       0.803329 0.151740 0.000000
       0.806876 0.152409 0.000000
       0.810418 0.153079 0.000000
       0.813968 0.153750 0.000000
       0.817524 0.154421 0.000000
       0.821083 0.155092 0.000000
       0.824651 0.155768 0.000000
       0.828210 0.156447 0.000000
       0.831768 0.157107 0.000000
       0.835337 0.157782 0.000000
       0.838911 0.158468 0.000000
       0.842479 0.159136 0.000000
       0.846038 0.159803 0.000000
       0.849626 0.160479 0.000000
       0.853215 0.161166 0.000000
       0.856801 0.161843 0.000000
       0.860383 0.162515 0.000000
       0.863979 0.163194 0.000000
       0.867570 0.163876 0.000000
       0.871152 0.164554 0.000000
       0.874735 0.165225 0.000000
       0.878339 0.165907 0.000000
       0.881956 0.166591 0.000000
       0.885556 0.167274 0.000000
       0.889154 0.167950 0.000000
       0.892773 0.168634 0.000000
       0.896392 0.169319 0.000000
       0.900000 0.170000 0.000000];

case { 'L14'  'GREENTERNARY' }
descriptorname = 'linear_ternary-green_0-46_c42_n256';
description = 'green colour map for ternary images';
map = [0.000000 0.000000 0.000000
       0.000000 0.003593 0.000000
       0.000000 0.007143 0.000000
       0.000000 0.010757 0.000000
       0.000000 0.014524 0.000000
       0.000000 0.018105 0.000000
       0.000000 0.021687 0.000000
       0.000000 0.025284 0.000000
       0.000000 0.028821 0.000000
       0.000000 0.032444 0.000000
       0.000000 0.036221 0.000000
       0.000000 0.039766 0.000000
       0.000000 0.043161 0.000000
       0.000000 0.046597 0.000000
       0.000000 0.049768 0.000000
       0.000000 0.052749 0.000000
       0.000000 0.055759 0.000000
       0.000000 0.058542 0.000000
       0.000000 0.061269 0.000000
       0.000000 0.063951 0.000000
       0.000000 0.066548 0.000000
       0.000000 0.069095 0.000000
       0.000000 0.071498 0.000000
       0.000000 0.073870 0.000000
       0.000000 0.076086 0.000000
       0.000000 0.078320 0.000000
       0.000000 0.080566 0.000000
       0.000000 0.082735 0.000000
       0.000000 0.084825 0.000000
       0.000000 0.086923 0.000000
       0.000000 0.088913 0.000000
       0.000000 0.090908 0.000000
       0.000000 0.092798 0.000000
       0.000000 0.094766 0.000000
       0.000000 0.096572 0.000000
       0.000000 0.098465 0.000000
       0.000000 0.100250 0.000000
       0.000000 0.102040 0.000000
       0.000000 0.103822 0.000000
       0.000000 0.105523 0.000000
       0.000000 0.107251 0.000000
       0.000000 0.108919 0.000000
       0.000000 0.110591 0.000000
       0.000000 0.112220 0.000000
       0.000004 0.113814 0.000001
       0.000016 0.115393 0.000002
       0.000017 0.116984 0.000002
       0.000013 0.118628 0.000002
       0.000010 0.120178 0.000001
       0.000013 0.121763 0.000002
       0.000016 0.123381 0.000002
       0.000017 0.125016 0.000002
       0.000016 0.126623 0.000002
       0.000011 0.128260 0.000001
       0.000008 0.129902 0.000001
       0.000014 0.131502 0.000002
       0.000017 0.133121 0.000002
       0.000018 0.134773 0.000002
       0.000016 0.136352 0.000002
       0.000012 0.138026 0.000002
       0.000008 0.139617 0.000001
       0.000014 0.141302 0.000002
       0.000018 0.142931 0.000002
       0.000018 0.144584 0.000002
       0.000017 0.146213 0.000002
       0.000012 0.147872 0.000002
       0.000009 0.149540 0.000001
       0.000013 0.151162 0.000002
       0.000017 0.152811 0.000002
       0.000019 0.154479 0.000002
       0.000018 0.156177 0.000002
       0.000015 0.157816 0.000002
       0.000011 0.159468 0.000001
       0.000009 0.161179 0.000001
       0.000015 0.162841 0.000002
       0.000018 0.164516 0.000002
       0.000010 0.166179 0.000002
       0.000000 0.167860 0.000002
       0.000000 0.169573 0.000002
       0.000000 0.171228 0.000001
       0.000000 0.172923 0.000001
       0.000000 0.174622 0.000002
       0.000000 0.176286 0.000002
       0.000000 0.178036 0.000002
       0.000000 0.179697 0.000002
       0.000000 0.181385 0.000002
       0.000000 0.183095 0.000002
       0.000000 0.184820 0.000001
       0.000000 0.186526 0.000001
       0.000000 0.188256 0.000002
       0.000000 0.189940 0.000002
       0.000000 0.191654 0.000002
       0.000000 0.193387 0.000002
       0.000000 0.195105 0.000002
       0.000000 0.196844 0.000002
       0.000000 0.198536 0.000001
       0.000000 0.200250 0.000001
       0.000000 0.201993 0.000002
       0.000000 0.203754 0.000002
       0.000000 0.205472 0.000002
       0.000000 0.207218 0.000002
       0.000000 0.208945 0.000002
       0.000000 0.210693 0.000002
       0.000000 0.212438 0.000002
       0.000000 0.214188 0.000001
       0.000000 0.215944 0.000001
       0.000000 0.217685 0.000002
       0.000000 0.219438 0.000002
       0.000000 0.221200 0.000002
       0.000000 0.222957 0.000002
       0.000000 0.224682 0.000002
       0.000000 0.226480 0.000002
       0.000000 0.228240 0.000002
       0.000000 0.229985 0.000001
       0.000000 0.231771 0.000001
       0.000000 0.233518 0.000001
       0.000000 0.235327 0.000002
       0.000000 0.237102 0.000002
       0.000000 0.238852 0.000002
       0.000000 0.240623 0.000002
       0.000000 0.242413 0.000002
       0.000000 0.244186 0.000002
       0.000000 0.245978 0.000002
       0.000000 0.247788 0.000002
       0.000000 0.249556 0.000001
       0.000000 0.251368 0.000001
       0.000000 0.253159 0.000001
       0.000000 0.254962 0.000002
       0.000000 0.256746 0.000002
       0.000000 0.258543 0.000002
       0.000000 0.260349 0.000002
       0.000000 0.262153 0.000002
       0.000000 0.263970 0.000002
       0.000000 0.265771 0.000002
       0.000000 0.267558 0.000002
       0.000000 0.269406 0.000002
       0.000000 0.271203 0.000002
       0.000000 0.273019 0.000001
       0.000000 0.274836 0.000001
       0.000000 0.276671 0.000001
       0.000000 0.278481 0.000001
       0.000000 0.280299 0.000002
       0.000000 0.282145 0.000002
       0.000000 0.283955 0.000002
       0.000000 0.285779 0.000002
       0.000000 0.287614 0.000002
       0.000000 0.289464 0.000002
       0.000000 0.291304 0.000002
       0.000000 0.293126 0.000002
       0.000000 0.294967 0.000002
       0.000000 0.296822 0.000002
       0.000000 0.298661 0.000002
       0.000000 0.300498 0.000001
       0.000000 0.302337 0.000001
       0.000000 0.304208 0.000001
       0.000000 0.306063 0.000001
       0.000000 0.307906 0.000001
       0.000000 0.309772 0.000002
       0.000000 0.311612 0.000002
       0.000000 0.313489 0.000002
       0.000000 0.315345 0.000001
       0.000000 0.317208 0.000000
       0.000000 0.319072 0.000000
       0.000000 0.320945 0.000000
       0.000000 0.322818 0.000000
       0.000000 0.324668 0.000000
       0.000000 0.326541 0.000000
       0.000000 0.328429 0.000000
       0.000000 0.330298 0.000000
       0.000000 0.332188 0.000000
       0.000000 0.334067 0.000000
       0.000000 0.335937 0.000000
       0.000000 0.337822 0.000000
       0.000000 0.339716 0.000000
       0.000000 0.341600 0.000000
       0.000000 0.343483 0.000000
       0.000000 0.345376 0.000000
       0.000000 0.347274 0.000000
       0.000000 0.349165 0.000000
       0.000000 0.351062 0.000000
       0.000000 0.352958 0.000000
       0.000000 0.354869 0.000000
       0.000000 0.356762 0.000000
       0.000000 0.358648 0.000000
       0.000000 0.360553 0.000000
       0.000000 0.362468 0.000000
       0.000000 0.364372 0.000000
       0.000000 0.366296 0.000000
       0.000000 0.368200 0.000000
       0.000000 0.370104 0.000000
       0.000000 0.372018 0.000000
       0.000000 0.373950 0.000000
       0.000000 0.375862 0.000000
       0.000000 0.377782 0.000000
       0.000000 0.379703 0.000000
       0.000000 0.381629 0.000000
       0.000000 0.383558 0.000000
       0.000000 0.385472 0.000000
       0.000000 0.387403 0.000000
       0.000000 0.389340 0.000000
       0.000000 0.391278 0.000000
       0.000000 0.393200 0.000000
       0.000000 0.395142 0.000000
       0.000000 0.397072 0.000000
       0.000000 0.399010 0.000000
       0.000000 0.400966 0.000000
       0.000000 0.402899 0.000000
       0.000000 0.404837 0.000000
       0.000000 0.406787 0.000000
       0.000000 0.408735 0.000000
       0.000000 0.410691 0.000000
       0.000000 0.412643 0.000000
       0.000000 0.414584 0.000000
       0.000000 0.416542 0.000000
       0.000000 0.418490 0.000000
       0.000000 0.420447 0.000000
       0.000000 0.422401 0.000000
       0.000000 0.424370 0.000000
       0.000000 0.426327 0.000000
       0.000000 0.428296 0.000000
       0.000000 0.430262 0.000000
       0.000000 0.432234 0.000000
       0.000000 0.434190 0.000000
       0.000000 0.436168 0.000000
       0.000000 0.438139 0.000000
       0.000000 0.440108 0.000000
       0.000000 0.442092 0.000000
       0.000000 0.444064 0.000000
       0.000000 0.446029 0.000000
       0.000000 0.448011 0.000000
       0.000000 0.449995 0.000000
       0.000000 0.451985 0.000000
       0.000000 0.453961 0.000000
       0.000000 0.455945 0.000000
       0.000000 0.457932 0.000000
       0.000000 0.459917 0.000000
       0.000000 0.461912 0.000000
       0.000000 0.463909 0.000000
       0.000000 0.465889 0.000000
       0.000000 0.467889 0.000000
       0.000000 0.469893 0.000000
       0.000000 0.471891 0.000000
       0.000000 0.473880 0.000000
       0.000000 0.475887 0.000000
       0.000000 0.477883 0.000000
       0.000000 0.479896 0.000000
       0.000000 0.481897 0.000000
       0.000000 0.483893 0.000000
       0.000000 0.485907 0.000000
       0.000000 0.487925 0.000000
       0.000000 0.489924 0.000000
       0.000000 0.491950 0.000000
       0.000000 0.493950 0.000000
       0.000000 0.495969 0.000000
       0.000000 0.497998 0.000000
       0.000000 0.500000 0.000000];

case { 'L15'  'BLUETERNARY' }
descriptorname = 'linear_ternary-blue_0-44_c57_n256';
description = 'blue colour map for ternary images';
map = [0.000000 0.000000 0.000000
       0.000743 0.002453 0.007434
       0.001492 0.004924 0.014921
       0.002224 0.007339 0.022238
       0.002973 0.009819 0.029727
       0.003759 0.012430 0.037588
       0.004496 0.014837 0.045061
       0.005217 0.017216 0.052174
       0.005923 0.019547 0.059280
       0.006612 0.021819 0.066122
       0.007284 0.024037 0.072890
       0.007942 0.026211 0.079443
       0.008587 0.028340 0.085862
       0.009227 0.030440 0.092265
       0.009844 0.032460 0.098364
       0.010477 0.034486 0.104584
       0.011109 0.036556 0.110783
       0.011710 0.038495 0.116668
       0.012273 0.040403 0.122424
       0.012818 0.042245 0.127904
       0.013348 0.043976 0.133264
       0.013879 0.045660 0.138500
       0.014395 0.047394 0.143575
       0.014889 0.049082 0.148578
       0.015355 0.050605 0.153349
       0.015827 0.052141 0.158011
       0.016294 0.053656 0.162613
       0.016752 0.055160 0.167171
       0.017191 0.056648 0.171533
       0.017621 0.057987 0.175845
       0.018044 0.059476 0.180120
       0.018452 0.060774 0.184207
       0.018859 0.062143 0.188304
       0.019258 0.063465 0.192290
       0.019638 0.064737 0.196182
       0.019991 0.066008 0.199990
       0.020369 0.067270 0.203831
       0.020743 0.068517 0.207566
       0.021108 0.069731 0.211228
       0.021466 0.070890 0.214792
       0.021817 0.072042 0.218328
       0.022172 0.073262 0.221839
       0.022518 0.074345 0.225285
       0.022855 0.075456 0.228641
       0.023186 0.076553 0.231960
       0.023512 0.077642 0.235257
       0.023840 0.078716 0.238500
       0.024178 0.079786 0.241793
       0.024507 0.080865 0.245060
       0.024831 0.081970 0.248322
       0.025156 0.082990 0.251560
       0.025480 0.084106 0.254803
       0.025801 0.085132 0.257985
       0.026127 0.086192 0.261265
       0.026456 0.087333 0.264558
       0.026783 0.088377 0.267805
       0.027109 0.089458 0.271082
       0.027438 0.090542 0.274378
       0.027766 0.091648 0.277662
       0.028090 0.092669 0.280876
       0.028421 0.093789 0.284198
       0.028753 0.094903 0.287528
       0.029077 0.095942 0.290776
       0.029408 0.097031 0.294074
       0.029741 0.098170 0.297423
       0.030072 0.099229 0.300719
       0.030403 0.100326 0.304016
       0.030735 0.101425 0.307330
       0.031072 0.102536 0.310723
       0.031403 0.103644 0.314028
       0.031734 0.104697 0.317340
       0.032072 0.105849 0.320738
       0.032408 0.106963 0.324078
       0.032743 0.108016 0.327392
       0.033082 0.109178 0.330769
       0.033414 0.110275 0.334179
       0.033742 0.111382 0.337529
       0.034077 0.112500 0.340899
       0.034420 0.113618 0.344240
       0.034765 0.114687 0.347627
       0.035114 0.115863 0.351044
       0.035456 0.116946 0.354445
       0.035791 0.118104 0.357839
       0.036121 0.119202 0.361222
       0.036456 0.120312 0.364591
       0.036796 0.121431 0.367975
       0.037139 0.122554 0.371374
       0.037483 0.123687 0.374812
       0.037828 0.124824 0.378263
       0.038169 0.125976 0.381685
       0.038511 0.127073 0.385105
       0.038856 0.128228 0.388561
       0.039200 0.129350 0.392000
       0.039546 0.130519 0.395463
       0.039890 0.131629 0.398890
       0.040235 0.132771 0.402332
       0.040581 0.133916 0.405818
       0.040922 0.135056 0.409233
       0.041266 0.136172 0.412685
       0.041619 0.137353 0.416196
       0.041970 0.138486 0.419638
       0.042315 0.139599 0.423081
       0.042655 0.140792 0.426583
       0.042998 0.141916 0.430062
       0.043353 0.143069 0.433531
       0.043707 0.144222 0.437051
       0.044054 0.145395 0.440537
       0.044401 0.146517 0.444043
       0.044755 0.147690 0.447524
       0.045109 0.148833 0.451029
       0.045452 0.150015 0.454550
       0.045792 0.151147 0.458032
       0.046147 0.152305 0.461541
       0.046509 0.153468 0.465061
       0.046871 0.154638 0.468602
       0.047221 0.155821 0.472157
       0.047563 0.156961 0.475683
       0.047916 0.158149 0.479207
       0.048273 0.159297 0.482723
       0.048631 0.160463 0.486278
       0.048997 0.161667 0.489859
       0.049352 0.162826 0.493419
       0.049698 0.164012 0.496956
       0.050046 0.165142 0.500484
       0.050404 0.166344 0.504062
       0.050762 0.167514 0.507630
       0.051121 0.168704 0.511196
       0.051479 0.169878 0.514786
       0.051833 0.171050 0.518336
       0.052190 0.172226 0.521895
       0.052551 0.173416 0.525509
       0.052909 0.174608 0.529096
       0.053266 0.175769 0.532666
       0.053628 0.176973 0.536283
       0.053990 0.178179 0.539894
       0.054349 0.179346 0.543487
       0.054708 0.180533 0.547071
       0.055067 0.181721 0.550673
       0.055429 0.182915 0.554294
       0.055793 0.184117 0.557934
       0.056155 0.185326 0.561570
       0.056521 0.186508 0.565196
       0.056888 0.187723 0.568838
       0.057244 0.188906 0.572447
       0.057600 0.190100 0.576066
       0.057967 0.191297 0.579694
       0.058336 0.192504 0.583348
       0.058702 0.193715 0.587014
       0.059067 0.194922 0.590661
       0.059436 0.196132 0.594323
       0.059804 0.197330 0.597974
       0.060167 0.198531 0.601607
       0.060527 0.199735 0.605254
       0.060887 0.200943 0.608925
       0.061253 0.202155 0.612590
       0.061628 0.203380 0.616299
       0.062000 0.204595 0.619993
       0.062370 0.205814 0.623678
       0.062743 0.207033 0.627362
       0.063109 0.208240 0.631039
       0.063471 0.209455 0.634720
       0.063840 0.210677 0.638419
       0.064212 0.211911 0.642128
       0.064582 0.213103 0.645808
       0.064951 0.214348 0.649512
       0.065324 0.215565 0.653234
       0.065695 0.216795 0.656960
       0.066066 0.218017 0.660658
       0.066438 0.219245 0.664381
       0.066811 0.220486 0.668116
       0.067184 0.221694 0.671834
       0.067556 0.222943 0.675556
       0.067927 0.224150 0.679263
       0.068301 0.225400 0.683018
       0.068677 0.226635 0.686779
       0.069052 0.227875 0.690523
       0.069427 0.229100 0.694243
       0.069800 0.230333 0.697992
       0.070170 0.231583 0.701750
       0.070545 0.232810 0.705491
       0.070927 0.234052 0.709266
       0.071310 0.235310 0.713043
       0.071685 0.236545 0.716810
       0.072052 0.237792 0.720560
       0.072426 0.239022 0.724321
       0.072809 0.240262 0.728088
       0.073193 0.241532 0.731881
       0.073573 0.242763 0.735655
       0.073946 0.244010 0.739436
       0.074319 0.245258 0.743217
       0.074700 0.246514 0.747018
       0.075082 0.247778 0.750834
       0.075461 0.249022 0.754617
       0.075841 0.250264 0.758408
       0.076222 0.251540 0.762219
       0.076602 0.252787 0.766025
       0.076984 0.254048 0.769846
       0.077365 0.255306 0.773648
       0.077744 0.256561 0.777448
       0.078128 0.257813 0.781281
       0.078511 0.259093 0.785110
       0.078893 0.260346 0.788934
       0.079275 0.261608 0.792754
       0.079660 0.262879 0.796600
       0.080044 0.264143 0.800434
       0.080427 0.265411 0.804270
       0.080810 0.266670 0.808094
       0.081193 0.267938 0.811936
       0.081582 0.269226 0.815835
       0.081967 0.270490 0.819662
       0.082351 0.271751 0.823495
       0.082734 0.273028 0.827356
       0.083120 0.274302 0.831221
       0.083510 0.275585 0.835102
       0.083897 0.276855 0.838956
       0.084288 0.278136 0.842840
       0.084673 0.279411 0.846691
       0.085055 0.280676 0.850551
       0.085441 0.281969 0.854446
       0.085828 0.283248 0.858321
       0.086218 0.284524 0.862196
       0.086607 0.285804 0.866076
       0.087001 0.287091 0.869974
       0.087391 0.288378 0.873876
       0.087778 0.289670 0.877775
       0.088167 0.290952 0.881683
       0.088557 0.292240 0.885573
       0.088947 0.293524 0.889468
       0.089338 0.294815 0.893377
       0.089730 0.296106 0.897298
       0.090121 0.297405 0.901213
       0.090513 0.298691 0.905136
       0.090904 0.299987 0.909044
       0.091296 0.301277 0.912961
       0.091690 0.302571 0.916883
       0.092085 0.303870 0.920814
       0.092476 0.305173 0.924766
       0.092866 0.306470 0.928684
       0.093261 0.307756 0.932611
       0.093659 0.309078 0.936589
       0.094052 0.310371 0.940523
       0.094445 0.311667 0.944450
       0.094842 0.312971 0.948399
       0.095238 0.314279 0.952351
       0.095633 0.315585 0.956322
       0.096025 0.316887 0.960265
       0.096419 0.318192 0.964218
       0.096817 0.319502 0.968191
       0.097216 0.320816 0.972160
       0.097613 0.322120 0.976129
       0.098012 0.323433 0.980098
       0.098407 0.324739 0.984058
       0.098802 0.326049 0.988028
       0.099201 0.327365 0.992016
       0.099601 0.328683 0.996011
       0.100000 0.330000 1.000000];

case { 'L16'  'KBGYW' }
descriptorname = 'linear_kbgyw_5-98_c62_n256';
description = 'Black-Blue-Green-Yellow-White colour map';
map = [0.066001 0.066024 0.066019
       0.073531 0.067157 0.081516
       0.080354 0.068289 0.094843
       0.086320 0.069502 0.107225
       0.091205 0.070686 0.119692
       0.095166 0.071969 0.132228
       0.098937 0.073058 0.144842
       0.102637 0.073887 0.157440
       0.106253 0.074435 0.170149
       0.109698 0.074836 0.182855
       0.112892 0.075212 0.195602
       0.115865 0.075547 0.208393
       0.118636 0.075871 0.221163
       0.121136 0.076152 0.233959
       0.123471 0.076440 0.246716
       0.125651 0.076696 0.259527
       0.127551 0.076951 0.272264
       0.129268 0.077199 0.285016
       0.130822 0.077434 0.297773
       0.132122 0.077693 0.310467
       0.133238 0.077937 0.323151
       0.134167 0.078217 0.335754
       0.134893 0.078507 0.348328
       0.135390 0.078813 0.360855
       0.135697 0.079175 0.373298
       0.135824 0.079553 0.385703
       0.135764 0.079992 0.398024
       0.135522 0.080484 0.410253
       0.135097 0.081016 0.422418
       0.134465 0.081668 0.434446
       0.133610 0.082336 0.446390
       0.132576 0.083014 0.458260
       0.131361 0.083921 0.469954
       0.129968 0.084835 0.481536
       0.128329 0.085789 0.493021
       0.126529 0.087046 0.504316
       0.124527 0.088286 0.515455
       0.122323 0.089654 0.526498
       0.119984 0.091206 0.537303
       0.117438 0.092805 0.547927
       0.114676 0.094674 0.558386
       0.111808 0.096524 0.568659
       0.108683 0.098699 0.578672
       0.105418 0.100955 0.588468
       0.101969 0.103416 0.598063
       0.098375 0.106004 0.607414
       0.094641 0.108758 0.616441
       0.090708 0.111754 0.625196
       0.086641 0.114830 0.633685
       0.082459 0.118193 0.641872
       0.078079 0.121626 0.649731
       0.073814 0.125368 0.657199
       0.069419 0.129244 0.664292
       0.064888 0.133341 0.671013
       0.060456 0.137629 0.677331
       0.056065 0.142072 0.683206
       0.051735 0.146711 0.688627
       0.047663 0.151577 0.693544
       0.043895 0.156669 0.697933
       0.040589 0.161953 0.701753
       0.037760 0.167425 0.704968
       0.035653 0.173115 0.707533
       0.033912 0.179013 0.709499
       0.032749 0.184938 0.711164
       0.031656 0.190799 0.712688
       0.030664 0.196668 0.714107
       0.029761 0.202442 0.715411
       0.028951 0.208227 0.716588
       0.028232 0.213979 0.717641
       0.027604 0.219702 0.718558
       0.027065 0.225395 0.719356
       0.026613 0.231086 0.720012
       0.026245 0.236764 0.720529
       0.025983 0.242399 0.720895
       0.025814 0.248069 0.721109
       0.025721 0.253699 0.721171
       0.025696 0.259342 0.721077
       0.025732 0.264950 0.720824
       0.025825 0.270603 0.720404
       0.026001 0.276234 0.719801
       0.026230 0.281860 0.719008
       0.026481 0.287483 0.718037
       0.026742 0.293128 0.716883
       0.027045 0.298789 0.715511
       0.027353 0.304447 0.713922
       0.027623 0.310133 0.712130
       0.027866 0.315817 0.710113
       0.028066 0.321502 0.707848
       0.028168 0.327214 0.705354
       0.028179 0.332966 0.702597
       0.028061 0.338707 0.699568
       0.027763 0.344473 0.696277
       0.027280 0.350282 0.692685
       0.026539 0.356100 0.688793
       0.025532 0.361939 0.684568
       0.024173 0.367803 0.680018
       0.022675 0.373712 0.675113
       0.022239 0.379575 0.669893
       0.024007 0.385356 0.664371
       0.027800 0.391083 0.658593
       0.033577 0.396711 0.652549
       0.041196 0.402291 0.646270
       0.049455 0.407821 0.639753
       0.057803 0.413298 0.633006
       0.066263 0.418717 0.626053
       0.074632 0.424100 0.618891
       0.082773 0.429448 0.611519
       0.090615 0.434763 0.603973
       0.098247 0.440039 0.596219
       0.105537 0.445286 0.588285
       0.112543 0.450514 0.580198
       0.119172 0.455715 0.571930
       0.125565 0.460891 0.563494
       0.131625 0.466051 0.554885
       0.137368 0.471205 0.546134
       0.142767 0.476332 0.537222
       0.147875 0.481446 0.528148
       0.152681 0.486550 0.518928
       0.157212 0.491653 0.509543
       0.161469 0.496722 0.500009
       0.165331 0.501807 0.490341
       0.168995 0.506869 0.480509
       0.172284 0.511931 0.470536
       0.175317 0.516995 0.460393
       0.178078 0.522035 0.450111
       0.180490 0.527095 0.439656
       0.182645 0.532137 0.429036
       0.184524 0.537183 0.418242
       0.186099 0.542215 0.407285
       0.187365 0.547251 0.396142
       0.188348 0.552293 0.384809
       0.189209 0.557301 0.373496
       0.190292 0.562236 0.362475
       0.191679 0.567090 0.351791
       0.193398 0.571869 0.341447
       0.195421 0.576592 0.331384
       0.197790 0.581233 0.321620
       0.200480 0.585829 0.312106
       0.203556 0.590349 0.302854
       0.206965 0.594822 0.293843
       0.210734 0.599238 0.285076
       0.214837 0.603597 0.276535
       0.219281 0.607907 0.268143
       0.223998 0.612161 0.259956
       0.229040 0.616381 0.251920
       0.234349 0.620555 0.244001
       0.239911 0.624677 0.236273
       0.245806 0.628755 0.228701
       0.251992 0.632786 0.221259
       0.258366 0.636783 0.213920
       0.264964 0.640732 0.206695
       0.271763 0.644646 0.199500
       0.278834 0.648522 0.192491
       0.286086 0.652354 0.185610
       0.293526 0.656155 0.178741
       0.301109 0.659913 0.171916
       0.308884 0.663641 0.165160
       0.316848 0.667315 0.158598
       0.324933 0.670963 0.151986
       0.333161 0.674574 0.145469
       0.341505 0.678153 0.138912
       0.350037 0.681681 0.132530
       0.358642 0.685191 0.126131
       0.367360 0.688664 0.119690
       0.376231 0.692095 0.113405
       0.385188 0.695486 0.107108
       0.394233 0.698845 0.100738
       0.403387 0.702173 0.094490
       0.412667 0.705463 0.088268
       0.421977 0.708726 0.082038
       0.431402 0.711945 0.075694
       0.440928 0.715125 0.069656
       0.450517 0.718275 0.063424
       0.460164 0.721389 0.057182
       0.469946 0.724468 0.051150
       0.479750 0.727511 0.045105
       0.489615 0.730523 0.039074
       0.499594 0.733489 0.033395
       0.509615 0.736422 0.028381
       0.519697 0.739331 0.023990
       0.529868 0.742185 0.020282
       0.540069 0.745010 0.017058
       0.550357 0.747804 0.014496
       0.560694 0.750555 0.012516
       0.571073 0.753264 0.010898
       0.581505 0.755946 0.010052
       0.591911 0.758608 0.009587
       0.602240 0.761258 0.009395
       0.612465 0.763911 0.009430
       0.622607 0.766564 0.009633
       0.632649 0.769216 0.010052
       0.642594 0.771876 0.010748
       0.652452 0.774528 0.011847
       0.662225 0.777197 0.012989
       0.671897 0.779855 0.014429
       0.681491 0.782523 0.016102
       0.690999 0.785198 0.018089
       0.700422 0.787873 0.020399
       0.709768 0.790558 0.022980
       0.719025 0.793253 0.025904
       0.728183 0.795950 0.029244
       0.737269 0.798662 0.032918
       0.746273 0.801379 0.037138
       0.755190 0.804113 0.041627
       0.764003 0.806859 0.046159
       0.772738 0.809614 0.050930
       0.781394 0.812381 0.055784
       0.789949 0.815169 0.060722
       0.798397 0.817975 0.065847
       0.806756 0.820801 0.071087
       0.815012 0.823646 0.076312
       0.823173 0.826515 0.081795
       0.831232 0.829407 0.087295
       0.839167 0.832322 0.092849
       0.846977 0.835269 0.098673
       0.854674 0.838247 0.104528
       0.862245 0.841258 0.110615
       0.869673 0.844302 0.116753
       0.876965 0.847384 0.123019
       0.884115 0.850510 0.129515
       0.891095 0.853679 0.136097
       0.897909 0.856902 0.142931
       0.904539 0.860180 0.149929
       0.910975 0.863513 0.157084
       0.917202 0.866902 0.164501
       0.923198 0.870364 0.172111
       0.928942 0.873906 0.179994
       0.934398 0.877534 0.188220
       0.939549 0.881259 0.196732
       0.944369 0.885081 0.205552
       0.948819 0.889016 0.214744
       0.952801 0.893087 0.224423
       0.956312 0.897303 0.234607
       0.959271 0.901690 0.245394
       0.961901 0.906152 0.257327
       0.964607 0.910539 0.271106
       0.967400 0.914847 0.286504
       0.970272 0.919067 0.303770
       0.973215 0.923210 0.322671
       0.976201 0.927257 0.343147
       0.979231 0.931213 0.365397
       0.982267 0.935070 0.389336
       0.985271 0.938832 0.414973
       0.988207 0.942492 0.442413
       0.991031 0.946038 0.471745
       0.993679 0.949476 0.503159
       0.996062 0.952801 0.536754
       0.998098 0.956010 0.572657
       0.999659 0.959096 0.611088
       1.000000 0.962060 0.652275
       1.000000 0.964901 0.696592
       0.999737 0.967624 0.744287
       0.997389 0.970219 0.795696
       0.993265 0.972710 0.851178
       0.986752 0.975079 0.911686
       0.977176 0.977372 0.977324];

case { 'L17'  'KBGYW' }
descriptorname = 'linear_worb_100-25_c53_n256';
description = 'White-Orange-Red-Blue, decreasing lightness with increasing saturation';
map = [0.999862 1.000000 1.000000
       0.998126 0.997011 0.988965
       0.996350 0.993970 0.977925
       0.994541 0.990931 0.966900
       0.992697 0.987895 0.955886
       0.990809 0.984867 0.944879
       0.988887 0.981839 0.933876
       0.986930 0.978811 0.922893
       0.984933 0.975792 0.911914
       0.982898 0.972782 0.900944
       0.980829 0.969767 0.889982
       0.978721 0.966762 0.879028
       0.976580 0.963758 0.868089
       0.974403 0.960763 0.857145
       0.972194 0.957768 0.846214
       0.969942 0.954777 0.835294
       0.967663 0.951788 0.824375
       0.965339 0.948806 0.813464
       0.962988 0.945828 0.802559
       0.960601 0.942857 0.791660
       0.958179 0.939887 0.780765
       0.955721 0.936921 0.769872
       0.953229 0.933953 0.758985
       0.950701 0.930999 0.748103
       0.950414 0.927081 0.740722
       0.950607 0.922960 0.734135
       0.950773 0.918833 0.727547
       0.950914 0.914716 0.720966
       0.951028 0.910599 0.714391
       0.951117 0.906482 0.707823
       0.951181 0.902368 0.701260
       0.951220 0.898254 0.694699
       0.951234 0.894147 0.688151
       0.951223 0.890037 0.681596
       0.951188 0.885935 0.675057
       0.951128 0.881831 0.668522
       0.951044 0.877729 0.661991
       0.950936 0.873629 0.655456
       0.950804 0.869532 0.648935
       0.950649 0.865437 0.642414
       0.950469 0.861345 0.635898
       0.950266 0.857254 0.629383
       0.950040 0.853166 0.622880
       0.949792 0.849083 0.616378
       0.949520 0.844998 0.609875
       0.949226 0.840917 0.603385
       0.948910 0.836837 0.596895
       0.949179 0.832471 0.591523
       0.949773 0.827939 0.586791
       0.950346 0.823410 0.582067
       0.950897 0.818875 0.577350
       0.951422 0.814343 0.572643
       0.951925 0.809809 0.567928
       0.952408 0.805270 0.563229
       0.952872 0.800733 0.558517
       0.953313 0.796191 0.553827
       0.953730 0.791650 0.549143
       0.954126 0.787109 0.544453
       0.954501 0.782559 0.539766
       0.954856 0.778013 0.535092
       0.955190 0.773457 0.530422
       0.955504 0.768904 0.525755
       0.955797 0.764352 0.521090
       0.956069 0.759791 0.516427
       0.956319 0.755237 0.511774
       0.956549 0.750675 0.507126
       0.956760 0.746105 0.502480
       0.956951 0.741534 0.497851
       0.957124 0.736966 0.493198
       0.957277 0.732395 0.488574
       0.957527 0.727713 0.484781
       0.957898 0.722908 0.482021
       0.958252 0.718103 0.479253
       0.958589 0.713288 0.476488
       0.958909 0.708470 0.473723
       0.959214 0.703639 0.470976
       0.959503 0.698800 0.468217
       0.959776 0.693956 0.465465
       0.960034 0.689116 0.462720
       0.960277 0.684248 0.459983
       0.960503 0.679385 0.457246
       0.960714 0.674514 0.454523
       0.960909 0.669636 0.451791
       0.961088 0.664739 0.449065
       0.961252 0.659845 0.446335
       0.961401 0.654931 0.443622
       0.961535 0.650020 0.440897
       0.961654 0.645088 0.438196
       0.961758 0.640158 0.435493
       0.961848 0.635206 0.432786
       0.961923 0.630253 0.430084
       0.961983 0.625281 0.427395
       0.962029 0.620307 0.424699
       0.961984 0.615316 0.422506
       0.961725 0.610348 0.421671
       0.961456 0.605363 0.420836
       0.961176 0.600371 0.420005
       0.960886 0.595363 0.419167
       0.960585 0.590340 0.418318
       0.960272 0.585315 0.417488
       0.959948 0.580264 0.416642
       0.959614 0.575197 0.415793
       0.959270 0.570116 0.414948
       0.958917 0.565021 0.414102
       0.958554 0.559922 0.413259
       0.958180 0.554793 0.412412
       0.957794 0.549651 0.411550
       0.957397 0.544490 0.410708
       0.956991 0.539308 0.409853
       0.956577 0.534109 0.408992
       0.956154 0.528903 0.408136
       0.955720 0.523661 0.407278
       0.955274 0.518406 0.406424
       0.954817 0.513122 0.405566
       0.954352 0.507824 0.404693
       0.953878 0.502492 0.403840
       0.953329 0.497179 0.403124
       0.952114 0.492175 0.403893
       0.950901 0.487158 0.404644
       0.949675 0.482118 0.405405
       0.948444 0.477054 0.406151
       0.947208 0.471981 0.406879
       0.945960 0.466877 0.407603
       0.944712 0.461750 0.408321
       0.943452 0.456599 0.409031
       0.942188 0.451434 0.409736
       0.940912 0.446232 0.410434
       0.939634 0.441008 0.411110
       0.938347 0.435771 0.411784
       0.937055 0.430491 0.412463
       0.935753 0.425189 0.413122
       0.934442 0.419851 0.413769
       0.933127 0.414480 0.414413
       0.931807 0.409081 0.415050
       0.930483 0.403651 0.415680
       0.929144 0.398179 0.416305
       0.927802 0.392664 0.416925
       0.926455 0.387112 0.417532
       0.925098 0.381533 0.418124
       0.923740 0.375898 0.418718
       0.921399 0.370998 0.420456
       0.918955 0.366174 0.422293
       0.916510 0.361302 0.424128
       0.914060 0.356422 0.425945
       0.911597 0.351496 0.427762
       0.909129 0.346551 0.429553
       0.906659 0.341578 0.431345
       0.904177 0.336564 0.433111
       0.901693 0.331519 0.434882
       0.899204 0.326440 0.436633
       0.896706 0.321335 0.438374
       0.894201 0.316196 0.440108
       0.891688 0.311002 0.441838
       0.889171 0.305773 0.443548
       0.886648 0.300473 0.445248
       0.884117 0.295145 0.446943
       0.881578 0.289773 0.448635
       0.879030 0.284336 0.450308
       0.876478 0.278864 0.451981
       0.873921 0.273317 0.453631
       0.871358 0.267679 0.455287
       0.868790 0.262019 0.456923
       0.866207 0.256256 0.458559
       0.862669 0.251647 0.460773
       0.858756 0.247425 0.463203
       0.854842 0.243169 0.465628
       0.850911 0.238910 0.468049
       0.846976 0.234642 0.470463
       0.843034 0.230288 0.472853
       0.839079 0.225966 0.475257
       0.835115 0.221565 0.477631
       0.831143 0.217155 0.480011
       0.827156 0.212688 0.482377
       0.823159 0.208199 0.484728
       0.819149 0.203674 0.487090
       0.815135 0.199067 0.489426
       0.811108 0.194481 0.491778
       0.807071 0.189779 0.494105
       0.803016 0.185067 0.496420
       0.798957 0.180253 0.498750
       0.794880 0.175410 0.501061
       0.790792 0.170498 0.503357
       0.786699 0.165481 0.505654
       0.782582 0.160417 0.507963
       0.778457 0.155287 0.510242
       0.774316 0.150057 0.512523
       0.769385 0.146290 0.514971
       0.763788 0.143876 0.517558
       0.758175 0.141468 0.520123
       0.752532 0.139023 0.522702
       0.746879 0.136621 0.525271
       0.741197 0.134263 0.527832
       0.735501 0.131866 0.530394
       0.729783 0.129483 0.532951
       0.724036 0.127087 0.535496
       0.718269 0.124741 0.538056
       0.712475 0.122372 0.540594
       0.706666 0.120054 0.543137
       0.700822 0.117745 0.545675
       0.694949 0.115427 0.548210
       0.689061 0.113168 0.550735
       0.683132 0.110862 0.553256
       0.677183 0.108540 0.555777
       0.671193 0.106343 0.558299
       0.665176 0.104054 0.560821
       0.659127 0.101818 0.563343
       0.653044 0.099603 0.565846
       0.646936 0.097418 0.568355
       0.640780 0.095268 0.570860
       0.634096 0.094454 0.573284
       0.626476 0.095920 0.575546
       0.618823 0.097420 0.577796
       0.611107 0.098865 0.580064
       0.603340 0.100272 0.582313
       0.595515 0.101663 0.584572
       0.587632 0.103050 0.586823
       0.579700 0.104335 0.589071
       0.571688 0.105671 0.591328
       0.563621 0.106980 0.593578
       0.555465 0.108169 0.595823
       0.547249 0.109470 0.598067
       0.538954 0.110659 0.600315
       0.530569 0.111840 0.602559
       0.522090 0.113019 0.604804
       0.513536 0.114104 0.607049
       0.504872 0.115229 0.609288
       0.496095 0.116370 0.611525
       0.487226 0.117412 0.613756
       0.478222 0.118524 0.615998
       0.469106 0.119531 0.618236
       0.459838 0.120545 0.620475
       0.450437 0.121547 0.622703
       0.440672 0.122888 0.624791
       0.429847 0.125757 0.626220
       0.418817 0.128432 0.627652
       0.407579 0.131021 0.629080
       0.396112 0.133476 0.630513
       0.384379 0.135804 0.631938
       0.372369 0.138122 0.633366
       0.360053 0.140266 0.634793
       0.347395 0.142361 0.636221
       0.334334 0.144370 0.637643
       0.320829 0.146279 0.639068
       0.306812 0.148125 0.640494
       0.292214 0.149927 0.641916
       0.276946 0.151601 0.643337
       0.260860 0.153235 0.644759
       0.243795 0.154822 0.646184
       0.225566 0.156367 0.647606
       0.205801 0.157795 0.649032
       0.184014 0.159198 0.650450
       0.159438 0.160526 0.651874
       0.130653 0.161862 0.653286
       0.094242 0.163093 0.654706
       0.038143 0.164291 0.656134
       0.000000 0.165388 0.657553];

case { 'L18'  'WORB' }
descriptorname = 'linear_wyor_100-45_c55_n256';
description = 'White-Yellow-Orange-Red, decreasing lightness with increasing saturation';
map = [0.999862 1.000000 1.000000
       0.997648 0.998314 0.989833
       0.995406 0.996565 0.979659
       0.993140 0.994821 0.969489
       0.990835 0.993082 0.959329
       0.988502 0.991337 0.949172
       0.986143 0.989598 0.939023
       0.983750 0.987855 0.928875
       0.981330 0.986117 0.918735
       0.978877 0.984381 0.908604
       0.976397 0.982642 0.898471
       0.973888 0.980905 0.888347
       0.971350 0.979170 0.878223
       0.968782 0.977436 0.868107
       0.966183 0.975700 0.857984
       0.963557 0.973970 0.847869
       0.960903 0.972243 0.837763
       0.958219 0.970510 0.827648
       0.955505 0.968782 0.817539
       0.952760 0.967055 0.807433
       0.949986 0.965325 0.797322
       0.947187 0.963600 0.787218
       0.944356 0.961873 0.777111
       0.941494 0.960153 0.766994
       0.940225 0.957714 0.760227
       0.939295 0.955118 0.754207
       0.938354 0.952521 0.748186
       0.937402 0.949928 0.742165
       0.936438 0.947340 0.736144
       0.935456 0.944753 0.730137
       0.934464 0.942166 0.724121
       0.933460 0.939581 0.718112
       0.932445 0.937000 0.712103
       0.931420 0.934415 0.706100
       0.930383 0.931837 0.700092
       0.929328 0.929261 0.694085
       0.928264 0.926687 0.688090
       0.927188 0.924118 0.682084
       0.926104 0.921548 0.676090
       0.925001 0.918977 0.670097
       0.923894 0.916414 0.664096
       0.922771 0.913854 0.658102
       0.921635 0.911290 0.652106
       0.920484 0.908731 0.646107
       0.919327 0.906175 0.640118
       0.918160 0.903619 0.634126
       0.916980 0.901065 0.628130
       0.916279 0.898329 0.622644
       0.915853 0.895495 0.617456
       0.915418 0.892658 0.612268
       0.914972 0.889822 0.607091
       0.914514 0.886990 0.601899
       0.914042 0.884161 0.596720
       0.913557 0.881330 0.591536
       0.913059 0.878501 0.586353
       0.912549 0.875677 0.581168
       0.912029 0.872852 0.575996
       0.911496 0.870026 0.570803
       0.910953 0.867207 0.565620
       0.910399 0.864392 0.560445
       0.909830 0.861573 0.555258
       0.909248 0.858756 0.550081
       0.908658 0.855945 0.544898
       0.908059 0.853130 0.539709
       0.907445 0.850321 0.534530
       0.906818 0.847510 0.529351
       0.906182 0.844707 0.524159
       0.905536 0.841903 0.518978
       0.904875 0.839098 0.513787
       0.904201 0.836297 0.508599
       0.903760 0.833404 0.503684
       0.903591 0.830399 0.499122
       0.903411 0.827394 0.494567
       0.903219 0.824389 0.489997
       0.903015 0.821390 0.485432
       0.902799 0.818385 0.480867
       0.902570 0.815389 0.476309
       0.902331 0.812388 0.471749
       0.902079 0.809395 0.467181
       0.901817 0.806398 0.462601
       0.901544 0.803398 0.458035
       0.901259 0.800406 0.453463
       0.900964 0.797409 0.448899
       0.900657 0.794421 0.444324
       0.900339 0.791426 0.439740
       0.900011 0.788436 0.435166
       0.899672 0.785448 0.430578
       0.899321 0.782457 0.425985
       0.898958 0.779468 0.421394
       0.898583 0.776484 0.416813
       0.898199 0.773493 0.412217
       0.897807 0.770512 0.407611
       0.897405 0.767524 0.403009
       0.897090 0.764500 0.398574
       0.897031 0.761353 0.394596
       0.896960 0.758211 0.390622
       0.896879 0.755066 0.386634
       0.896788 0.751919 0.382660
       0.896685 0.748771 0.378676
       0.896572 0.745625 0.374695
       0.896448 0.742483 0.370696
       0.896314 0.739341 0.366716
       0.896169 0.736185 0.362714
       0.896014 0.733045 0.358710
       0.895849 0.729899 0.354726
       0.895674 0.726750 0.350720
       0.895489 0.723601 0.346703
       0.895293 0.720455 0.342695
       0.895088 0.717316 0.338666
       0.894872 0.714160 0.334655
       0.894647 0.711018 0.330609
       0.894410 0.707865 0.326569
       0.894163 0.704718 0.322546
       0.893905 0.701567 0.318491
       0.893638 0.698418 0.314430
       0.893362 0.695272 0.310388
       0.893100 0.692115 0.306383
       0.893056 0.688835 0.302984
       0.893002 0.685551 0.299620
       0.892939 0.682262 0.296232
       0.892866 0.678978 0.292841
       0.892784 0.675694 0.289463
       0.892692 0.672410 0.286050
       0.892591 0.669118 0.282679
       0.892481 0.665825 0.279275
       0.892361 0.662536 0.275874
       0.892231 0.659234 0.272434
       0.892093 0.655943 0.269043
       0.891945 0.652636 0.265609
       0.891787 0.649343 0.262179
       0.891621 0.646033 0.258749
       0.891446 0.642731 0.255314
       0.891261 0.639424 0.251867
       0.891069 0.636116 0.248419
       0.890867 0.632802 0.244924
       0.890656 0.629484 0.241476
       0.890437 0.626168 0.237995
       0.890209 0.622851 0.234519
       0.889972 0.619534 0.231012
       0.889727 0.616205 0.227507
       0.889614 0.612783 0.224683
       0.889507 0.609361 0.221973
       0.889392 0.605919 0.219257
       0.889268 0.602486 0.216538
       0.889137 0.599048 0.213787
       0.888997 0.595599 0.211048
       0.888849 0.592157 0.208302
       0.888693 0.588691 0.205554
       0.888528 0.585244 0.202795
       0.888355 0.581773 0.200023
       0.888173 0.578304 0.197289
       0.887984 0.574831 0.194539
       0.887787 0.571356 0.191737
       0.887582 0.567876 0.188969
       0.887371 0.564391 0.186191
       0.887152 0.560894 0.183379
       0.886926 0.557405 0.180579
       0.886692 0.553894 0.177815
       0.886450 0.550387 0.174988
       0.886199 0.546874 0.172147
       0.885940 0.543352 0.169353
       0.885673 0.539819 0.166496
       0.885399 0.536293 0.163667
       0.885170 0.532708 0.161478
       0.884954 0.529101 0.159492
       0.884731 0.525476 0.157551
       0.884502 0.521841 0.155629
       0.884265 0.518218 0.153659
       0.884022 0.514575 0.151717
       0.883771 0.510917 0.149805
       0.883512 0.507260 0.147843
       0.883246 0.503588 0.145907
       0.882973 0.499906 0.143947
       0.882694 0.496219 0.141996
       0.882409 0.492524 0.140018
       0.882119 0.488821 0.138112
       0.881822 0.485100 0.136102
       0.881517 0.481376 0.134198
       0.881205 0.477639 0.132232
       0.880885 0.473888 0.130305
       0.880558 0.470137 0.128316
       0.880226 0.466359 0.126356
       0.879887 0.462569 0.124365
       0.879542 0.458782 0.122384
       0.879190 0.454978 0.120431
       0.878833 0.451152 0.118515
       0.878468 0.447298 0.117052
       0.878096 0.443435 0.116137
       0.877717 0.439547 0.115141
       0.877331 0.435653 0.114188
       0.876938 0.431741 0.113307
       0.876543 0.427804 0.112353
       0.876143 0.423845 0.111415
       0.875737 0.419887 0.110488
       0.875323 0.415900 0.109577
       0.874902 0.411900 0.108583
       0.874477 0.407879 0.107679
       0.874046 0.403845 0.106809
       0.873609 0.399774 0.105871
       0.873167 0.395708 0.104907
       0.872720 0.391604 0.104038
       0.872268 0.387471 0.103162
       0.871809 0.383337 0.102238
       0.871342 0.379159 0.101339
       0.870868 0.374964 0.100448
       0.870390 0.370739 0.099568
       0.869908 0.366507 0.098702
       0.869424 0.362228 0.097851
       0.868934 0.357931 0.096914
       0.868411 0.353600 0.096399
       0.867836 0.349284 0.096516
       0.867253 0.344929 0.096632
       0.866671 0.340545 0.096748
       0.866085 0.336116 0.096864
       0.865490 0.331660 0.096979
       0.864892 0.327166 0.097092
       0.864294 0.322652 0.097204
       0.863688 0.318072 0.097312
       0.863074 0.313463 0.097418
       0.862458 0.308814 0.097519
       0.861834 0.304094 0.097617
       0.861203 0.299356 0.097711
       0.860574 0.294538 0.097800
       0.859939 0.289692 0.097885
       0.859295 0.284771 0.097966
       0.858648 0.279815 0.098043
       0.857997 0.274772 0.098117
       0.857341 0.269689 0.098187
       0.856681 0.264508 0.098254
       0.856018 0.259280 0.098319
       0.855350 0.253950 0.098381
       0.854673 0.248554 0.098440
       0.853966 0.243057 0.098636
       0.853154 0.237651 0.099470
       0.852341 0.232114 0.100299
       0.851512 0.226483 0.101115
       0.850685 0.220749 0.101921
       0.849859 0.214853 0.102720
       0.849024 0.208870 0.103510
       0.848182 0.202734 0.104219
       0.847338 0.196478 0.104939
       0.846491 0.190003 0.105709
       0.845641 0.183361 0.106456
       0.844789 0.176515 0.107140
       0.843927 0.169491 0.107783
       0.843065 0.162164 0.108439
       0.842201 0.154525 0.109146
       0.841328 0.146577 0.109818
       0.840454 0.138289 0.110433
       0.839576 0.129502 0.111048
       0.838688 0.120174 0.111656
       0.837805 0.110272 0.112254
       0.836912 0.099467 0.112853
       0.836016 0.087701 0.113425
       0.835117 0.074450 0.113949
       0.834216 0.059213 0.114471];

case { 'L19'  'WCMR' }
descriptorname = 'linear_wcmr_100-45_c42_n256';
description = 'White-Cyan-Magenta-Red, decreasing lightness with increasing saturation';
map = [0.999862 1.000000 1.000000
       0.994940 0.998133 0.999618
       0.990018 0.996204 0.999221
       0.985086 0.994280 0.998823
       0.980146 0.992356 0.998424
       0.975197 0.990428 0.998023
       0.970245 0.988501 0.997623
       0.965284 0.986578 0.997222
       0.960322 0.984653 0.996821
       0.955348 0.982726 0.996419
       0.950365 0.980800 0.996017
       0.945377 0.978872 0.995615
       0.940381 0.976949 0.995212
       0.935373 0.975022 0.994808
       0.930361 0.973102 0.994405
       0.925337 0.971175 0.994001
       0.920306 0.969250 0.993596
       0.915267 0.967329 0.993191
       0.910225 0.965401 0.992783
       0.905169 0.963479 0.992374
       0.900101 0.961554 0.991965
       0.895027 0.959634 0.991555
       0.889939 0.957712 0.991145
       0.884845 0.955791 0.990735
       0.880659 0.953567 0.990836
       0.876670 0.951276 0.991051
       0.872681 0.948987 0.991264
       0.868683 0.946701 0.991475
       0.864669 0.944418 0.991685
       0.860653 0.942136 0.991895
       0.856627 0.939854 0.992102
       0.852598 0.937572 0.992309
       0.848553 0.935292 0.992515
       0.844506 0.933012 0.992719
       0.840449 0.930740 0.992921
       0.836381 0.928461 0.993122
       0.832307 0.926191 0.993322
       0.828220 0.923921 0.993519
       0.824127 0.921650 0.993715
       0.820023 0.919379 0.993910
       0.815917 0.917117 0.994103
       0.811794 0.914847 0.994296
       0.807663 0.912584 0.994487
       0.803521 0.910323 0.994677
       0.799374 0.908063 0.994866
       0.795208 0.905802 0.995053
       0.791036 0.903542 0.995239
       0.787957 0.901005 0.995458
       0.785507 0.898310 0.995695
       0.783046 0.895621 0.995930
       0.780583 0.892929 0.996163
       0.778117 0.890240 0.996396
       0.775641 0.887551 0.996627
       0.773161 0.884869 0.996856
       0.770682 0.882184 0.997084
       0.768187 0.879501 0.997311
       0.765700 0.876818 0.997536
       0.763195 0.874142 0.997760
       0.760693 0.871468 0.997983
       0.758184 0.868795 0.998204
       0.755666 0.866118 0.998424
       0.753139 0.863448 0.998643
       0.750619 0.860774 0.998860
       0.748086 0.858104 0.999075
       0.745542 0.855441 0.999289
       0.743003 0.852776 0.999501
       0.740448 0.850110 0.999711
       0.737887 0.847445 0.999919
       0.735329 0.844789 1.000000
       0.732762 0.842132 1.000000
       0.731152 0.839264 1.000000
       0.730711 0.836147 0.999945
       0.730266 0.833029 0.999602
       0.729811 0.829916 0.999257
       0.729349 0.826799 0.998909
       0.728881 0.823680 0.998560
       0.728410 0.820563 0.998210
       0.727933 0.817448 0.997859
       0.727450 0.814332 0.997507
       0.726963 0.811220 0.997155
       0.726470 0.808104 0.996802
       0.725973 0.804987 0.996447
       0.725470 0.801872 0.996092
       0.724958 0.798759 0.995737
       0.724439 0.795641 0.995380
       0.723915 0.792528 0.995022
       0.723387 0.789415 0.994664
       0.722853 0.786301 0.994305
       0.722314 0.783181 0.993945
       0.721771 0.780066 0.993585
       0.721222 0.776956 0.993223
       0.720668 0.773833 0.992859
       0.720110 0.770723 0.992493
       0.720127 0.767485 0.991827
       0.721681 0.763940 0.990366
       0.723222 0.760392 0.988905
       0.724750 0.756837 0.987445
       0.726254 0.753277 0.985983
       0.727738 0.749715 0.984524
       0.729209 0.746155 0.983060
       0.730662 0.742592 0.981601
       0.732101 0.739023 0.980140
       0.733517 0.735444 0.978675
       0.734921 0.731868 0.977218
       0.736300 0.728283 0.975754
       0.737670 0.724701 0.974294
       0.739030 0.721106 0.972838
       0.740363 0.717520 0.971375
       0.741680 0.713917 0.969914
       0.742992 0.710317 0.968456
       0.744276 0.706713 0.966995
       0.745549 0.703101 0.965532
       0.746809 0.699481 0.964072
       0.748055 0.695867 0.962614
       0.749276 0.692246 0.961155
       0.750495 0.688615 0.959695
       0.751847 0.684947 0.958100
       0.754710 0.680972 0.955129
       0.757530 0.677003 0.952157
       0.760320 0.673014 0.949192
       0.763072 0.669017 0.946226
       0.765798 0.665007 0.943268
       0.768479 0.660993 0.940309
       0.771139 0.656976 0.937349
       0.773754 0.652935 0.934389
       0.776347 0.648900 0.931436
       0.778902 0.644841 0.928481
       0.781431 0.640781 0.925534
       0.783928 0.636714 0.922587
       0.786392 0.632627 0.919638
       0.788825 0.628535 0.916696
       0.791226 0.624428 0.913755
       0.793608 0.620315 0.910813
       0.795948 0.616181 0.907877
       0.798267 0.612037 0.904939
       0.800559 0.607892 0.902003
       0.802818 0.603726 0.899072
       0.805054 0.599547 0.896142
       0.807265 0.595351 0.893211
       0.809447 0.591147 0.890285
       0.812460 0.586809 0.885788
       0.815521 0.582443 0.881134
       0.818531 0.578061 0.876483
       0.821505 0.573672 0.871846
       0.824427 0.569256 0.867203
       0.827312 0.564827 0.862577
       0.830153 0.560388 0.857943
       0.832949 0.555915 0.853320
       0.835708 0.551444 0.848705
       0.838424 0.546943 0.844091
       0.841103 0.542425 0.839488
       0.843738 0.537890 0.834884
       0.846338 0.533321 0.830288
       0.848903 0.528751 0.825693
       0.851423 0.524143 0.821108
       0.853910 0.519519 0.816530
       0.856364 0.514874 0.811951
       0.858778 0.510200 0.807382
       0.861157 0.505499 0.802813
       0.863507 0.500789 0.798254
       0.865813 0.496036 0.793705
       0.868093 0.491282 0.789154
       0.870328 0.486469 0.784611
       0.872652 0.481809 0.778812
       0.874968 0.477180 0.772556
       0.877237 0.472532 0.766310
       0.879465 0.467856 0.760072
       0.881652 0.463150 0.753845
       0.883793 0.458434 0.747629
       0.885889 0.453682 0.741412
       0.887941 0.448912 0.735216
       0.889956 0.444105 0.729022
       0.891931 0.439257 0.722838
       0.893861 0.434391 0.716673
       0.895756 0.429495 0.710507
       0.897610 0.424562 0.704351
       0.899425 0.419598 0.698205
       0.901198 0.414589 0.692078
       0.902940 0.409551 0.685950
       0.904638 0.404464 0.679829
       0.906304 0.399340 0.673729
       0.907934 0.394184 0.667637
       0.909523 0.388986 0.661551
       0.911080 0.383732 0.655472
       0.912601 0.378427 0.649410
       0.914091 0.373074 0.643347
       0.915228 0.368133 0.636498
       0.916075 0.363529 0.628974
       0.916888 0.358896 0.621464
       0.917658 0.354244 0.613968
       0.918384 0.349555 0.606493
       0.919075 0.344826 0.599030
       0.919733 0.340070 0.591578
       0.920349 0.335278 0.584135
       0.920930 0.330425 0.576718
       0.921480 0.325537 0.569296
       0.921992 0.320637 0.561908
       0.922467 0.315667 0.554520
       0.922906 0.310656 0.547147
       0.923313 0.305592 0.539786
       0.923684 0.300457 0.532452
       0.924021 0.295285 0.525119
       0.924323 0.290055 0.517808
       0.924593 0.284761 0.510499
       0.924832 0.279425 0.503206
       0.925039 0.273992 0.495934
       0.925214 0.268491 0.488679
       0.925358 0.262913 0.481430
       0.925471 0.257249 0.474200
       0.925142 0.252198 0.466598
       0.924073 0.248334 0.458331
       0.922972 0.244401 0.450085
       0.921841 0.240474 0.441857
       0.920674 0.236555 0.433620
       0.919485 0.232562 0.425422
       0.918267 0.228570 0.417230
       0.917021 0.224493 0.409040
       0.915737 0.220464 0.400881
       0.914438 0.216357 0.392706
       0.913104 0.212208 0.384555
       0.911743 0.208013 0.376431
       0.910358 0.203803 0.368298
       0.908939 0.199500 0.360171
       0.907501 0.195205 0.352059
       0.906033 0.190832 0.343955
       0.904536 0.186423 0.335868
       0.903019 0.181926 0.327774
       0.901470 0.177425 0.319698
       0.899901 0.172795 0.311602
       0.898302 0.168115 0.303524
       0.896686 0.163392 0.295448
       0.895041 0.158552 0.287365
       0.893101 0.154270 0.279211
       0.889946 0.153025 0.270600
       0.886773 0.151806 0.261956
       0.883577 0.150610 0.253309
       0.880359 0.149433 0.244613
       0.877124 0.148227 0.235934
       0.873873 0.147070 0.227188
       0.870599 0.145942 0.218403
       0.867310 0.144811 0.209562
       0.864007 0.143679 0.200651
       0.860676 0.142573 0.191705
       0.857330 0.141495 0.182654
       0.853966 0.140408 0.173527
       0.850587 0.139315 0.164302
       0.847190 0.138317 0.154890
       0.843776 0.137283 0.145381
       0.840351 0.136221 0.135588
       0.836902 0.135254 0.125642
       0.833439 0.134282 0.115345
       0.829961 0.133294 0.104675
       0.826465 0.132336 0.093595
       0.822954 0.131393 0.081950
       0.819425 0.130485 0.069425
       0.815891 0.129538 0.055606];

case { 'D1'  'BWR'  'COOLWARM' }
descriptorname = 'diverging_bwr_40-95_c42_n256';
description = 'Classic diverging blue - white - red colour map. End colours are matched in lightness and chroma';
map = [0.127567 0.316187 0.858358
       0.146748 0.320479 0.859229
       0.163863 0.324762 0.860098
       0.179359 0.329098 0.860959
       0.193703 0.333427 0.861824
       0.207083 0.337762 0.862685
       0.219673 0.342127 0.863540
       0.231620 0.346493 0.864392
       0.242974 0.350890 0.865238
       0.253887 0.355292 0.866090
       0.264359 0.359690 0.866932
       0.274452 0.364108 0.867779
       0.284220 0.368554 0.868622
       0.293689 0.372991 0.869454
       0.302889 0.377456 0.870284
       0.311844 0.381923 0.871119
       0.320606 0.386402 0.871951
       0.329136 0.390910 0.872774
       0.337471 0.395415 0.873596
       0.345632 0.399922 0.874415
       0.353641 0.404451 0.875234
       0.361507 0.408995 0.876048
       0.369226 0.413547 0.876855
       0.376821 0.418101 0.877667
       0.384277 0.422669 0.878472
       0.391636 0.427265 0.879273
       0.398873 0.431860 0.880072
       0.406027 0.436454 0.880870
       0.413074 0.441058 0.881665
       0.420024 0.445677 0.882452
       0.426897 0.450317 0.883241
       0.433674 0.454963 0.884029
       0.440389 0.459609 0.884808
       0.447027 0.464277 0.885586
       0.453592 0.468942 0.886363
       0.460089 0.473609 0.887132
       0.466530 0.478303 0.887899
       0.472897 0.482998 0.888668
       0.479220 0.487715 0.889429
       0.485472 0.492419 0.890187
       0.491690 0.497148 0.890943
       0.497845 0.501883 0.891698
       0.503942 0.506618 0.892449
       0.509994 0.511368 0.893191
       0.515998 0.516123 0.893936
       0.521964 0.520897 0.894679
       0.527893 0.525675 0.895414
       0.533772 0.530458 0.896147
       0.539616 0.535247 0.896879
       0.545429 0.540048 0.897604
       0.551196 0.544860 0.898324
       0.556933 0.549678 0.899047
       0.562633 0.554503 0.899765
       0.568289 0.559334 0.900476
       0.573923 0.564179 0.901185
       0.579533 0.569020 0.901892
       0.585104 0.573879 0.902597
       0.590639 0.578745 0.903297
       0.596157 0.583615 0.903990
       0.601643 0.588491 0.904684
       0.607111 0.593386 0.905377
       0.612536 0.598275 0.906061
       0.617948 0.603183 0.906742
       0.623340 0.608097 0.907423
       0.628701 0.613005 0.908098
       0.634042 0.617933 0.908766
       0.639354 0.622868 0.909434
       0.644644 0.627808 0.910102
       0.649924 0.632754 0.910762
       0.655172 0.637707 0.911418
       0.660406 0.642669 0.912072
       0.665622 0.647634 0.912722
       0.670820 0.652608 0.913370
       0.675992 0.657599 0.914014
       0.681145 0.662587 0.914651
       0.686291 0.667582 0.915283
       0.691412 0.672586 0.915913
       0.696522 0.677597 0.916545
       0.701607 0.682607 0.917173
       0.706689 0.687633 0.917793
       0.711747 0.692669 0.918406
       0.716793 0.697697 0.919017
       0.721813 0.702746 0.919628
       0.726831 0.707793 0.920233
       0.731833 0.712848 0.920832
       0.736818 0.717917 0.921434
       0.741790 0.722982 0.922030
       0.746752 0.728061 0.922620
       0.751703 0.733147 0.923206
       0.756633 0.738233 0.923789
       0.761554 0.743335 0.924364
       0.766465 0.748436 0.924937
       0.771368 0.753542 0.925510
       0.776252 0.758661 0.926078
       0.781127 0.763779 0.926638
       0.785994 0.768906 0.927196
       0.790844 0.774040 0.927751
       0.795687 0.779183 0.928301
       0.800522 0.784336 0.928849
       0.805342 0.789486 0.929392
       0.810155 0.794644 0.929933
       0.814959 0.799813 0.930469
       0.819751 0.804979 0.930999
       0.824536 0.810155 0.931524
       0.829317 0.815341 0.932045
       0.834083 0.820528 0.932562
       0.838838 0.825727 0.933077
       0.843588 0.830929 0.933580
       0.848332 0.836123 0.934071
       0.853065 0.841312 0.934546
       0.857785 0.846483 0.934997
       0.862500 0.851636 0.935418
       0.867188 0.856754 0.935799
       0.871868 0.861824 0.936124
       0.876516 0.866816 0.936376
       0.881143 0.871721 0.936533
       0.885730 0.876488 0.936572
       0.890267 0.881098 0.936462
       0.894749 0.885498 0.936168
       0.899150 0.889639 0.935653
       0.903460 0.893470 0.934878
       0.907659 0.896941 0.933803
       0.911719 0.899981 0.932391
       0.915622 0.902543 0.930604
       0.919352 0.904571 0.928397
       0.922884 0.906025 0.925760
       0.926191 0.906861 0.922662
       0.929260 0.907058 0.919090
       0.932081 0.906601 0.915055
       0.934645 0.905495 0.910562
       0.936951 0.903744 0.905624
       0.938989 0.901385 0.900267
       0.940776 0.898451 0.894532
       0.942321 0.894994 0.888448
       0.943634 0.891054 0.882060
       0.944738 0.886701 0.875413
       0.945647 0.881981 0.868548
       0.946383 0.876952 0.861491
       0.946973 0.871679 0.854293
       0.947427 0.866192 0.846979
       0.947763 0.860544 0.839587
       0.947999 0.854768 0.832124
       0.948148 0.848895 0.824612
       0.948223 0.842950 0.817077
       0.948232 0.836954 0.809523
       0.948182 0.830920 0.801955
       0.948080 0.824855 0.794392
       0.947930 0.818776 0.786829
       0.947736 0.812689 0.779263
       0.947497 0.806596 0.771714
       0.947219 0.800497 0.764172
       0.946903 0.794403 0.756647
       0.946550 0.788306 0.749129
       0.946161 0.782215 0.741627
       0.945739 0.776120 0.734141
       0.945282 0.770027 0.726661
       0.944786 0.763936 0.719196
       0.944254 0.757849 0.711744
       0.943688 0.751761 0.704299
       0.943088 0.745667 0.696871
       0.942454 0.739586 0.689459
       0.941781 0.733491 0.682046
       0.941077 0.727400 0.674658
       0.940343 0.721309 0.667280
       0.939570 0.715225 0.659914
       0.938765 0.709139 0.652556
       0.937928 0.703046 0.645214
       0.937059 0.696952 0.637889
       0.936156 0.690857 0.630578
       0.935217 0.684767 0.623274
       0.934247 0.678674 0.615978
       0.933247 0.672584 0.608708
       0.932216 0.666479 0.601435
       0.931156 0.660382 0.594184
       0.930061 0.654282 0.586943
       0.928932 0.648177 0.579725
       0.927775 0.642076 0.572511
       0.926589 0.635969 0.565300
       0.925371 0.629855 0.558115
       0.924124 0.623748 0.550948
       0.922846 0.617623 0.543787
       0.921536 0.611505 0.536646
       0.920195 0.605375 0.529510
       0.918825 0.599253 0.522380
       0.917435 0.593119 0.515273
       0.916000 0.586974 0.508189
       0.914548 0.580832 0.501104
       0.913063 0.574678 0.494034
       0.911547 0.568525 0.486979
       0.910007 0.562374 0.479941
       0.908435 0.556191 0.472902
       0.906835 0.550023 0.465886
       0.905209 0.543839 0.458893
       0.903551 0.537653 0.451909
       0.901867 0.531448 0.444932
       0.900157 0.525239 0.437972
       0.898415 0.519026 0.431021
       0.896654 0.512790 0.424078
       0.894861 0.506548 0.417165
       0.893037 0.500297 0.410259
       0.891190 0.494040 0.403358
       0.889318 0.487769 0.396467
       0.887415 0.481471 0.389598
       0.885491 0.475177 0.382744
       0.883539 0.468854 0.375901
       0.881559 0.462509 0.369072
       0.879551 0.456162 0.362252
       0.877521 0.449796 0.355455
       0.875466 0.443409 0.348652
       0.873382 0.437002 0.341871
       0.871276 0.430570 0.335113
       0.869145 0.424113 0.328340
       0.866982 0.417644 0.321588
       0.864800 0.411141 0.314849
       0.862598 0.404607 0.308127
       0.860363 0.398063 0.301409
       0.858102 0.391479 0.294708
       0.855824 0.384851 0.288017
       0.853513 0.378206 0.281334
       0.851183 0.371513 0.274662
       0.848834 0.364793 0.267990
       0.846453 0.358037 0.261348
       0.844052 0.351227 0.254714
       0.841630 0.344361 0.248074
       0.839181 0.337468 0.241428
       0.836709 0.330505 0.234812
       0.834217 0.323493 0.228182
       0.831700 0.316425 0.221546
       0.829158 0.309293 0.214916
       0.826593 0.302056 0.208310
       0.824003 0.294773 0.201675
       0.821397 0.287402 0.195077
       0.818762 0.279956 0.188466
       0.816115 0.272389 0.181804
       0.813434 0.264735 0.175193
       0.810737 0.256973 0.168544
       0.808021 0.249074 0.161909
       0.805278 0.241026 0.155209
       0.802516 0.232832 0.148495
       0.799739 0.224457 0.141803
       0.796928 0.215945 0.135061
       0.794109 0.207176 0.128267
       0.791258 0.198154 0.121393
       0.788392 0.188894 0.114504
       0.785506 0.179300 0.107595
       0.782595 0.169376 0.100561
       0.779666 0.158980 0.093435
       0.776723 0.148083 0.086208
       0.773748 0.136563 0.078844
       0.770765 0.124287 0.071410
       0.767752 0.111069 0.063657
       0.764730 0.096422 0.055653
       0.761680 0.080002 0.047336
       0.758619 0.060659 0.038618
       0.755534 0.036080 0.029827
       0.752427 0.008418 0.021805];

case { 'D1A'  'BWRA' }
descriptorname = 'diverging_bwr_20-95_c54_n256';
description = 'Diverging blue - white - red colour map. Similar to D1 but with darker end point colours';
map = [0.091874 0.160934 0.446902
       0.093320 0.165181 0.458168
       0.094888 0.169522 0.469498
       0.096269 0.173806 0.480851
       0.097771 0.178162 0.492271
       0.099148 0.182462 0.503735
       0.100522 0.186837 0.515243
       0.101881 0.191205 0.526810
       0.103239 0.195599 0.538413
       0.104476 0.199990 0.550056
       0.105793 0.204451 0.561752
       0.107055 0.208868 0.573487
       0.108206 0.213311 0.585273
       0.109463 0.217795 0.597096
       0.110610 0.222270 0.608963
       0.111756 0.226779 0.620859
       0.112913 0.231296 0.632785
       0.114011 0.235825 0.644733
       0.115169 0.240343 0.656710
       0.116414 0.244907 0.668683
       0.117659 0.249496 0.680642
       0.119053 0.254113 0.692594
       0.120548 0.258734 0.704479
       0.122270 0.263389 0.716302
       0.124294 0.268040 0.728007
       0.126662 0.272753 0.739584
       0.129463 0.277501 0.750967
       0.132772 0.282286 0.762116
       0.136649 0.287082 0.772998
       0.141252 0.291964 0.783561
       0.146503 0.296886 0.793753
       0.152542 0.301840 0.803524
       0.159357 0.306886 0.812846
       0.166938 0.311963 0.821694
       0.175186 0.317138 0.830031
       0.184077 0.322370 0.837855
       0.193556 0.327650 0.845158
       0.203495 0.333024 0.851959
       0.213813 0.338429 0.858267
       0.224397 0.343911 0.864127
       0.235244 0.349464 0.869554
       0.246144 0.355074 0.874605
       0.257133 0.360699 0.879317
       0.268081 0.366416 0.883742
       0.279006 0.372142 0.887911
       0.289802 0.377928 0.891879
       0.300479 0.383747 0.895676
       0.311032 0.389588 0.899339
       0.321403 0.395471 0.902896
       0.331620 0.401371 0.906370
       0.341684 0.407294 0.909783
       0.351572 0.413248 0.913151
       0.361309 0.419216 0.916484
       0.370889 0.425207 0.919795
       0.380339 0.431216 0.923091
       0.389638 0.437242 0.926371
       0.398815 0.443282 0.929645
       0.407873 0.449344 0.932912
       0.416822 0.455418 0.936183
       0.425656 0.461512 0.939438
       0.434396 0.467622 0.942689
       0.443057 0.473742 0.945921
       0.451631 0.479897 0.949139
       0.460123 0.486052 0.952334
       0.468563 0.492235 0.955502
       0.476941 0.498445 0.958625
       0.485264 0.504664 0.961691
       0.493552 0.510898 0.964696
       0.501811 0.517174 0.967624
       0.510031 0.523464 0.970443
       0.518244 0.529784 0.973150
       0.526440 0.536131 0.975711
       0.534623 0.542517 0.978117
       0.542815 0.548935 0.980348
       0.551002 0.555376 0.982380
       0.559194 0.561877 0.984198
       0.567399 0.568398 0.985792
       0.575603 0.574966 0.987156
       0.583800 0.581577 0.988277
       0.592008 0.588224 0.989171
       0.600192 0.594919 0.989832
       0.608373 0.601647 0.990272
       0.616522 0.608422 0.990507
       0.624642 0.615213 0.990552
       0.632728 0.622046 0.990426
       0.640769 0.628918 0.990148
       0.648768 0.635811 0.989738
       0.656715 0.642732 0.989213
       0.664595 0.649679 0.988590
       0.672433 0.656647 0.987890
       0.680193 0.663636 0.987128
       0.687908 0.670642 0.986308
       0.695556 0.677666 0.985446
       0.703144 0.684699 0.984551
       0.710669 0.691759 0.983623
       0.718134 0.698826 0.982678
       0.725548 0.705916 0.981711
       0.732898 0.713012 0.980726
       0.740195 0.720129 0.979729
       0.747439 0.727255 0.978713
       0.754632 0.734398 0.977693
       0.761766 0.741545 0.976658
       0.768857 0.748713 0.975612
       0.775904 0.755893 0.974558
       0.782901 0.763081 0.973493
       0.789855 0.770285 0.972418
       0.796765 0.777509 0.971326
       0.803643 0.784731 0.970212
       0.810477 0.791950 0.969076
       0.817278 0.799168 0.967911
       0.824042 0.806365 0.966700
       0.830777 0.813529 0.965440
       0.837478 0.820650 0.964115
       0.844143 0.827701 0.962706
       0.850780 0.834656 0.961185
       0.857391 0.841470 0.959523
       0.863975 0.848099 0.957688
       0.870510 0.854497 0.955636
       0.877011 0.860595 0.953322
       0.883468 0.866324 0.950696
       0.889858 0.871609 0.947710
       0.896175 0.876361 0.944310
       0.902396 0.880507 0.940448
       0.908503 0.883968 0.936077
       0.914470 0.886659 0.931157
       0.920268 0.888519 0.925664
       0.925882 0.889494 0.919577
       0.931275 0.889550 0.912895
       0.936433 0.888671 0.905623
       0.941320 0.886851 0.897774
       0.945934 0.884116 0.889388
       0.950261 0.880495 0.880500
       0.954292 0.876054 0.871165
       0.958028 0.870850 0.861428
       0.961469 0.864961 0.851345
       0.964633 0.858468 0.840981
       0.967537 0.851449 0.830374
       0.970182 0.843988 0.819573
       0.972603 0.836157 0.808636
       0.974801 0.828019 0.797582
       0.976811 0.819641 0.786461
       0.978639 0.811077 0.775275
       0.980309 0.802360 0.764067
       0.981828 0.793538 0.752841
       0.983205 0.784624 0.741617
       0.984461 0.775642 0.730404
       0.985591 0.766617 0.719199
       0.986611 0.757556 0.708010
       0.987522 0.748467 0.696845
       0.988324 0.739357 0.685707
       0.989033 0.730224 0.674594
       0.989644 0.721077 0.663518
       0.990160 0.711926 0.652456
       0.990584 0.702757 0.641425
       0.990918 0.693569 0.630432
       0.991162 0.684364 0.619460
       0.991318 0.675149 0.608515
       0.991386 0.665911 0.597591
       0.991365 0.656658 0.586700
       0.991256 0.647378 0.575850
       0.991058 0.638080 0.565010
       0.990769 0.628764 0.554225
       0.990388 0.619429 0.543468
       0.989912 0.610063 0.532753
       0.989337 0.600687 0.522069
       0.988652 0.591296 0.511447
       0.987858 0.581887 0.500883
       0.986947 0.572479 0.490377
       0.985901 0.563061 0.479943
       0.984718 0.553636 0.469594
       0.983376 0.544233 0.459329
       0.981877 0.534844 0.449175
       0.980196 0.525482 0.439125
       0.978323 0.516149 0.429221
       0.976255 0.506875 0.419454
       0.973979 0.497660 0.409834
       0.971489 0.488493 0.400372
       0.968779 0.479402 0.391072
       0.965848 0.470385 0.381929
       0.962706 0.461443 0.372962
       0.959353 0.452587 0.364161
       0.955803 0.443795 0.355533
       0.952057 0.435082 0.347039
       0.948145 0.426418 0.338679
       0.944075 0.417826 0.330457
       0.939863 0.409277 0.322361
       0.935522 0.400774 0.314351
       0.931075 0.392268 0.306459
       0.926532 0.383793 0.298618
       0.921914 0.375305 0.290861
       0.917226 0.366823 0.283168
       0.912475 0.358305 0.275524
       0.907681 0.349762 0.267886
       0.902842 0.341172 0.260329
       0.897965 0.332538 0.252787
       0.893059 0.323816 0.245249
       0.888125 0.315036 0.237779
       0.883165 0.306196 0.230273
       0.878183 0.297238 0.222836
       0.873176 0.288171 0.215371
       0.868153 0.279021 0.207943
       0.863099 0.269734 0.200498
       0.858014 0.260303 0.193115
       0.852907 0.250732 0.185750
       0.847760 0.241005 0.178376
       0.842582 0.231126 0.171004
       0.837350 0.221063 0.163715
       0.832061 0.210781 0.156432
       0.826701 0.200295 0.149181
       0.821255 0.189645 0.142011
       0.815711 0.178777 0.134933
       0.810039 0.167675 0.127924
       0.804231 0.156402 0.121008
       0.798246 0.144909 0.114278
       0.792071 0.133219 0.107747
       0.785678 0.121347 0.101406
       0.779035 0.109456 0.095344
       0.772129 0.097351 0.089503
       0.764938 0.085214 0.084009
       0.757441 0.073154 0.078743
       0.749632 0.060978 0.073995
       0.741514 0.049092 0.069570
       0.733089 0.037259 0.065437
       0.724356 0.026823 0.061723
       0.715339 0.018618 0.058443
       0.706052 0.012283 0.055532
       0.696523 0.007339 0.052912
       0.686776 0.004037 0.050680
       0.676849 0.001944 0.048770
       0.666751 0.000853 0.047012
       0.656541 0.000571 0.045448
       0.646215 0.000925 0.044119
       0.635819 0.001762 0.042871
       0.625365 0.002950 0.041932
       0.614873 0.004382 0.040902
       0.604377 0.005967 0.039904
       0.593863 0.007634 0.038882
       0.583358 0.009337 0.037833
       0.572880 0.011077 0.036734
       0.562419 0.012852 0.035630
       0.551981 0.014445 0.034215
       0.541566 0.015957 0.032974
       0.531204 0.017388 0.031658
       0.520872 0.018728 0.030292
       0.510579 0.019973 0.028861
       0.500328 0.021123 0.027367
       0.490123 0.022180 0.025811
       0.479964 0.023147 0.024193
       0.469845 0.024025 0.022515
       0.459760 0.024816 0.020778
       0.449730 0.025523 0.018981
       0.439744 0.026148 0.017126
       0.429804 0.026693 0.015214
       0.419914 0.027159 0.013240
       0.410078 0.027551 0.011092
       0.400282 0.027868 0.008951];

case { 'D2'  'GWV' }
descriptorname = 'diverging_gwv_55-95_c39_n256';
description = 'Diverging green - white - violet colour map';
map = [0.221573 0.591068 0.055073
       0.229654 0.593887 0.068993
       0.237600 0.596706 0.081291
       0.245312 0.599521 0.092565
       0.252923 0.602329 0.103154
       0.260374 0.605140 0.113125
       0.267681 0.607957 0.122534
       0.274907 0.610762 0.131675
       0.282020 0.613561 0.140449
       0.289020 0.616374 0.148940
       0.295919 0.619183 0.157212
       0.302734 0.621973 0.165276
       0.309501 0.624780 0.173201
       0.316165 0.627579 0.180945
       0.322761 0.630380 0.188595
       0.329293 0.633174 0.196096
       0.335751 0.635970 0.203492
       0.342161 0.638762 0.210790
       0.348512 0.641551 0.218015
       0.354823 0.644342 0.225143
       0.361055 0.647138 0.232239
       0.367262 0.649926 0.239245
       0.373426 0.652707 0.246191
       0.379544 0.655497 0.253110
       0.385615 0.658282 0.259969
       0.391666 0.661062 0.266776
       0.397669 0.663850 0.273568
       0.403641 0.666622 0.280290
       0.409575 0.669408 0.286998
       0.415477 0.672186 0.293681
       0.421346 0.674960 0.300337
       0.427207 0.677741 0.306971
       0.433019 0.680506 0.313582
       0.438805 0.683285 0.320172
       0.444586 0.686057 0.326717
       0.450323 0.688832 0.333283
       0.456041 0.691597 0.339815
       0.461742 0.694360 0.346318
       0.467421 0.697132 0.352832
       0.473066 0.699897 0.359318
       0.478709 0.702666 0.365804
       0.484322 0.705428 0.372267
       0.489923 0.708192 0.378734
       0.495508 0.710956 0.385184
       0.501078 0.713714 0.391642
       0.506620 0.716478 0.398084
       0.512154 0.719234 0.404509
       0.517680 0.721986 0.410950
       0.523183 0.724749 0.417379
       0.528674 0.727500 0.423790
       0.534141 0.730258 0.430217
       0.539603 0.733011 0.436636
       0.545060 0.735759 0.443051
       0.550498 0.738513 0.449465
       0.555916 0.741259 0.455873
       0.561339 0.744013 0.462282
       0.566749 0.746761 0.468697
       0.572141 0.749504 0.475116
       0.577519 0.752254 0.481518
       0.582893 0.755004 0.487938
       0.588256 0.757746 0.494348
       0.593618 0.760490 0.500759
       0.598965 0.763229 0.507174
       0.604307 0.765974 0.513596
       0.609631 0.768710 0.520006
       0.614949 0.771456 0.526440
       0.620273 0.774189 0.532861
       0.625572 0.776934 0.539284
       0.630877 0.779666 0.545721
       0.636168 0.782404 0.552155
       0.641445 0.785141 0.558580
       0.646730 0.787872 0.565020
       0.652003 0.790606 0.571469
       0.657271 0.793344 0.577913
       0.662530 0.796070 0.584370
       0.667785 0.798804 0.590824
       0.673032 0.801532 0.597290
       0.678273 0.804265 0.603756
       0.683509 0.806994 0.610219
       0.688746 0.809720 0.616698
       0.693965 0.812444 0.623176
       0.699190 0.815172 0.629653
       0.704411 0.817895 0.636147
       0.709628 0.820620 0.642639
       0.714833 0.823345 0.649140
       0.720041 0.826068 0.655641
       0.725246 0.828794 0.662153
       0.730441 0.831517 0.668664
       0.735631 0.834237 0.675181
       0.740820 0.836955 0.681701
       0.746008 0.839676 0.688238
       0.751198 0.842394 0.694769
       0.756372 0.845109 0.701313
       0.761547 0.847824 0.707860
       0.766721 0.850541 0.714411
       0.771894 0.853256 0.720971
       0.777065 0.855974 0.727538
       0.782227 0.858685 0.734111
       0.787388 0.861400 0.740685
       0.792547 0.864117 0.747272
       0.797702 0.866823 0.753859
       0.802857 0.869536 0.760455
       0.808011 0.872250 0.767052
       0.813158 0.874957 0.773659
       0.818306 0.877667 0.780275
       0.823453 0.880374 0.786899
       0.828602 0.883084 0.793528
       0.833747 0.885791 0.800161
       0.838881 0.888490 0.806800
       0.844011 0.891176 0.813434
       0.849136 0.893851 0.820073
       0.854241 0.896508 0.826712
       0.859331 0.899136 0.833335
       0.864396 0.901723 0.839945
       0.869422 0.904261 0.846519
       0.874400 0.906735 0.853063
       0.879316 0.909115 0.859551
       0.884155 0.911388 0.865963
       0.888885 0.913523 0.872278
       0.893481 0.915478 0.878464
       0.897924 0.917243 0.884498
       0.902176 0.918753 0.890338
       0.906208 0.919998 0.895957
       0.909983 0.920927 0.901310
       0.913471 0.921520 0.906368
       0.916643 0.921741 0.911093
       0.919475 0.921567 0.915455
       0.921955 0.920984 0.919441
       0.924059 0.919995 0.923031
       0.925791 0.918590 0.926210
       0.927149 0.916791 0.928985
       0.928150 0.914605 0.931369
       0.928808 0.912060 0.933372
       0.929148 0.909188 0.935027
       0.929196 0.906026 0.936361
       0.928984 0.902601 0.937398
       0.928542 0.898954 0.938182
       0.927905 0.895121 0.938746
       0.927101 0.891128 0.939125
       0.926163 0.887014 0.939352
       0.925109 0.882796 0.939454
       0.923970 0.878507 0.939456
       0.922757 0.874155 0.939382
       0.921487 0.869760 0.939248
       0.920175 0.865334 0.939070
       0.918830 0.860886 0.938859
       0.917468 0.856422 0.938624
       0.916072 0.851948 0.938371
       0.914671 0.847462 0.938105
       0.913252 0.842976 0.937827
       0.911823 0.838487 0.937545
       0.910390 0.834003 0.937260
       0.908943 0.829515 0.936973
       0.907498 0.825022 0.936683
       0.906040 0.820534 0.936390
       0.904573 0.816052 0.936094
       0.903105 0.811563 0.935794
       0.901623 0.807076 0.935492
       0.900139 0.802584 0.935188
       0.898644 0.798096 0.934882
       0.897148 0.793614 0.934573
       0.895638 0.789124 0.934262
       0.894123 0.784635 0.933949
       0.892602 0.780141 0.933633
       0.891070 0.775651 0.933315
       0.889534 0.771166 0.932995
       0.887989 0.766670 0.932672
       0.886442 0.762177 0.932347
       0.884882 0.757689 0.932020
       0.883316 0.753192 0.931691
       0.881744 0.748701 0.931360
       0.880160 0.744207 0.931026
       0.878574 0.739718 0.930690
       0.876976 0.735218 0.930350
       0.875379 0.730718 0.930006
       0.873767 0.726222 0.929661
       0.872154 0.721717 0.929313
       0.870524 0.717226 0.928963
       0.868900 0.712714 0.928611
       0.867255 0.708216 0.928257
       0.865609 0.703710 0.927901
       0.863961 0.699200 0.927542
       0.862300 0.694692 0.927181
       0.860628 0.690185 0.926819
       0.858950 0.685677 0.926454
       0.857266 0.681156 0.926088
       0.855580 0.676649 0.925718
       0.853876 0.672130 0.925344
       0.852177 0.667612 0.924967
       0.850459 0.663092 0.924590
       0.848740 0.658561 0.924213
       0.847009 0.654036 0.923834
       0.845276 0.649511 0.923450
       0.843530 0.644971 0.923063
       0.841784 0.640442 0.922675
       0.840028 0.635903 0.922285
       0.838258 0.631364 0.921892
       0.836485 0.626816 0.921495
       0.834705 0.622266 0.921094
       0.832914 0.617717 0.920694
       0.831122 0.613160 0.920294
       0.829319 0.608614 0.919891
       0.827503 0.604050 0.919484
       0.825683 0.599482 0.919074
       0.823856 0.594908 0.918664
       0.822026 0.590330 0.918254
       0.820178 0.585761 0.917841
       0.818327 0.581170 0.917425
       0.816475 0.576591 0.917005
       0.814605 0.571991 0.916581
       0.812731 0.567394 0.916154
       0.810851 0.562793 0.915727
       0.808966 0.558172 0.915300
       0.807069 0.553557 0.914873
       0.805159 0.548946 0.914443
       0.803247 0.544313 0.914010
       0.801327 0.539672 0.913573
       0.799403 0.535035 0.913133
       0.797461 0.530389 0.912691
       0.795517 0.525734 0.912248
       0.793570 0.521069 0.911804
       0.791604 0.516393 0.911357
       0.789637 0.511715 0.910909
       0.787658 0.507030 0.910459
       0.785676 0.502334 0.910006
       0.783684 0.497637 0.909548
       0.781683 0.492907 0.909090
       0.779669 0.488195 0.908632
       0.777656 0.483449 0.908174
       0.775625 0.478707 0.907711
       0.773589 0.473949 0.907244
       0.771553 0.469192 0.906776
       0.769496 0.464414 0.906307
       0.767437 0.459614 0.905837
       0.765376 0.454814 0.905364
       0.763294 0.449990 0.904887
       0.761211 0.445157 0.904406
       0.759116 0.440309 0.903927
       0.757016 0.435456 0.903449
       0.754910 0.430574 0.902966
       0.752783 0.425678 0.902480
       0.750663 0.420764 0.901991
       0.748525 0.415840 0.901502
       0.746378 0.410900 0.901011
       0.744225 0.405938 0.900518
       0.742063 0.400958 0.900024
       0.739897 0.395946 0.899529
       0.737709 0.390919 0.899029
       0.735522 0.385856 0.898525
       0.733326 0.380798 0.898023
       0.731115 0.375690 0.897522
       0.728901 0.370563 0.897017
       0.726676 0.365417 0.896507
       0.724443 0.360233 0.895996
       0.722195 0.355043 0.895484
       0.719948 0.349793 0.894972];

case { 'D3'  'GWR' }
descriptorname = 'diverging_gwr_55-95_c38_n256';
description = 'Diverging green - white - red colour map';
map = [0.221573 0.591068 0.055073
       0.229654 0.593887 0.068993
       0.237600 0.596706 0.081291
       0.245312 0.599521 0.092565
       0.252923 0.602329 0.103154
       0.260374 0.605140 0.113125
       0.267681 0.607957 0.122534
       0.274907 0.610762 0.131675
       0.282020 0.613561 0.140449
       0.289020 0.616374 0.148940
       0.295919 0.619183 0.157212
       0.302734 0.621973 0.165276
       0.309501 0.624780 0.173201
       0.316165 0.627579 0.180945
       0.322761 0.630380 0.188595
       0.329293 0.633174 0.196096
       0.335751 0.635970 0.203492
       0.342161 0.638762 0.210790
       0.348512 0.641551 0.218015
       0.354823 0.644342 0.225143
       0.361055 0.647138 0.232239
       0.367262 0.649926 0.239245
       0.373426 0.652707 0.246191
       0.379544 0.655497 0.253110
       0.385615 0.658282 0.259969
       0.391666 0.661062 0.266776
       0.397669 0.663850 0.273568
       0.403641 0.666622 0.280290
       0.409575 0.669408 0.286998
       0.415477 0.672186 0.293681
       0.421346 0.674960 0.300337
       0.427207 0.677741 0.306971
       0.433019 0.680506 0.313582
       0.438805 0.683285 0.320172
       0.444586 0.686057 0.326717
       0.450323 0.688832 0.333283
       0.456041 0.691597 0.339815
       0.461742 0.694360 0.346318
       0.467421 0.697132 0.352832
       0.473066 0.699897 0.359318
       0.478709 0.702666 0.365804
       0.484322 0.705428 0.372267
       0.489923 0.708192 0.378734
       0.495508 0.710956 0.385184
       0.501078 0.713714 0.391642
       0.506620 0.716478 0.398084
       0.512154 0.719234 0.404509
       0.517680 0.721986 0.410950
       0.523183 0.724749 0.417379
       0.528674 0.727500 0.423790
       0.534141 0.730258 0.430217
       0.539603 0.733011 0.436636
       0.545060 0.735759 0.443051
       0.550498 0.738513 0.449465
       0.555916 0.741259 0.455873
       0.561339 0.744013 0.462282
       0.566749 0.746761 0.468697
       0.572141 0.749504 0.475116
       0.577519 0.752254 0.481518
       0.582893 0.755004 0.487938
       0.588256 0.757746 0.494348
       0.593618 0.760490 0.500759
       0.598965 0.763229 0.507174
       0.604307 0.765974 0.513596
       0.609631 0.768710 0.520006
       0.614949 0.771456 0.526440
       0.620273 0.774189 0.532861
       0.625572 0.776934 0.539284
       0.630877 0.779666 0.545721
       0.636168 0.782404 0.552155
       0.641445 0.785141 0.558580
       0.646730 0.787872 0.565020
       0.652003 0.790606 0.571469
       0.657271 0.793344 0.577913
       0.662530 0.796070 0.584370
       0.667785 0.798804 0.590824
       0.673032 0.801532 0.597290
       0.678273 0.804265 0.603756
       0.683509 0.806994 0.610219
       0.688746 0.809720 0.616698
       0.693965 0.812444 0.623176
       0.699190 0.815172 0.629653
       0.704411 0.817895 0.636147
       0.709628 0.820620 0.642639
       0.714833 0.823345 0.649140
       0.720041 0.826068 0.655641
       0.725246 0.828794 0.662153
       0.730441 0.831517 0.668664
       0.735631 0.834237 0.675181
       0.740820 0.836955 0.681701
       0.746008 0.839676 0.688238
       0.751198 0.842394 0.694769
       0.756372 0.845109 0.701313
       0.761547 0.847824 0.707860
       0.766721 0.850541 0.714411
       0.771894 0.853256 0.720971
       0.777065 0.855974 0.727538
       0.782227 0.858685 0.734111
       0.787388 0.861400 0.740685
       0.792547 0.864117 0.747272
       0.797702 0.866823 0.753859
       0.802857 0.869536 0.760455
       0.808011 0.872250 0.767052
       0.813158 0.874957 0.773659
       0.818306 0.877667 0.780275
       0.823453 0.880374 0.786899
       0.828602 0.883084 0.793528
       0.833749 0.885791 0.800156
       0.838888 0.888489 0.806785
       0.844028 0.891174 0.813399
       0.849169 0.893848 0.820005
       0.854298 0.896503 0.826594
       0.859426 0.899127 0.833140
       0.864544 0.901709 0.839641
       0.869646 0.904240 0.846059
       0.874733 0.906704 0.852390
       0.879795 0.909070 0.858575
       0.884827 0.911325 0.864595
       0.889811 0.913434 0.870391
       0.894743 0.915359 0.875923
       0.899598 0.917082 0.881122
       0.904360 0.918545 0.885933
       0.909017 0.919727 0.890287
       0.913548 0.920582 0.894132
       0.917924 0.921083 0.897404
       0.922129 0.921200 0.900049
       0.926142 0.920907 0.902031
       0.929950 0.920195 0.903323
       0.933534 0.919050 0.903899
       0.936898 0.917487 0.903768
       0.940019 0.915492 0.902943
       0.942905 0.913113 0.901443
       0.945558 0.910358 0.899319
       0.947990 0.907261 0.896612
       0.950211 0.903856 0.893379
       0.952236 0.900187 0.889690
       0.954089 0.896286 0.885602
       0.955781 0.892189 0.881175
       0.957328 0.887927 0.876465
       0.958757 0.883540 0.871537
       0.960079 0.879043 0.866422
       0.961308 0.874469 0.861170
       0.962461 0.869831 0.855818
       0.963549 0.865149 0.850383
       0.964577 0.860436 0.844898
       0.965558 0.855695 0.839374
       0.966496 0.850930 0.833826
       0.967396 0.846159 0.828257
       0.968257 0.841383 0.822689
       0.969081 0.836594 0.817109
       0.969876 0.831809 0.811538
       0.970643 0.827016 0.805971
       0.971383 0.822225 0.800413
       0.972097 0.817427 0.794859
       0.972783 0.812631 0.789316
       0.973437 0.807836 0.783777
       0.974066 0.803035 0.778241
       0.974668 0.798235 0.772713
       0.975245 0.793437 0.767193
       0.975795 0.788630 0.761680
       0.976319 0.783824 0.756177
       0.976819 0.779010 0.750680
       0.977293 0.774196 0.745183
       0.977739 0.769382 0.739705
       0.978158 0.764569 0.734221
       0.978554 0.759745 0.728746
       0.978927 0.754930 0.723279
       0.979275 0.750098 0.717826
       0.979599 0.745271 0.712368
       0.979896 0.740441 0.706930
       0.980169 0.735604 0.701486
       0.980417 0.730766 0.696062
       0.980642 0.725929 0.690630
       0.980844 0.721077 0.685220
       0.981022 0.716236 0.679803
       0.981177 0.711382 0.674404
       0.981308 0.706525 0.669011
       0.981417 0.701656 0.663626
       0.981502 0.696795 0.658241
       0.981565 0.691928 0.652860
       0.981605 0.687050 0.647497
       0.981623 0.682166 0.642139
       0.981618 0.677291 0.636789
       0.981591 0.672400 0.631441
       0.981542 0.667504 0.626099
       0.981471 0.662605 0.620774
       0.981378 0.657700 0.615442
       0.981264 0.652777 0.610127
       0.981128 0.647860 0.604822
       0.980971 0.642938 0.599521
       0.980793 0.638008 0.594223
       0.980594 0.633070 0.588931
       0.980374 0.628128 0.583655
       0.980134 0.623175 0.578382
       0.979873 0.618213 0.573126
       0.979591 0.613239 0.567864
       0.979288 0.608274 0.562622
       0.978963 0.603286 0.557380
       0.978619 0.598288 0.552144
       0.978256 0.593293 0.546915
       0.977875 0.588274 0.541688
       0.977474 0.583256 0.536488
       0.977051 0.578225 0.531280
       0.976608 0.573193 0.526087
       0.976146 0.568137 0.520895
       0.975665 0.563082 0.515708
       0.975166 0.558000 0.510541
       0.974647 0.552916 0.505371
       0.974110 0.547823 0.500217
       0.973555 0.542717 0.495076
       0.972981 0.537596 0.489928
       0.972388 0.532458 0.484793
       0.971773 0.527306 0.479679
       0.971140 0.522132 0.474562
       0.970490 0.516963 0.469457
       0.969823 0.511761 0.464356
       0.969137 0.506549 0.459258
       0.968435 0.501329 0.454172
       0.967714 0.496072 0.449097
       0.966972 0.490824 0.444029
       0.966212 0.485527 0.438956
       0.965436 0.480231 0.433902
       0.964643 0.474914 0.428866
       0.963834 0.469571 0.423825
       0.963008 0.464205 0.418799
       0.962160 0.458808 0.413783
       0.961300 0.453392 0.408771
       0.960423 0.447950 0.403773
       0.959525 0.442499 0.398772
       0.958613 0.437006 0.393785
       0.957683 0.431490 0.388816
       0.956733 0.425929 0.383844
       0.955774 0.420351 0.378880
       0.954792 0.414741 0.373934
       0.953796 0.409097 0.368990
       0.952782 0.403421 0.364046
       0.951750 0.397701 0.359118
       0.950707 0.391943 0.354204
       0.949643 0.386138 0.349298
       0.948566 0.380309 0.344387
       0.947475 0.374429 0.339513
       0.946361 0.368497 0.334629
       0.945240 0.362510 0.329752
       0.944098 0.356483 0.324865
       0.942942 0.350396 0.320024
       0.941767 0.344231 0.315162
       0.940581 0.338024 0.310335
       0.939377 0.331752 0.305500
       0.938158 0.325400 0.300657
       0.936927 0.318991 0.295844
       0.935676 0.312480 0.291045
       0.934408 0.305924 0.286229
       0.933128 0.299245 0.281447
       0.931836 0.292464 0.276678
       0.930530 0.285582 0.271880
       0.929203 0.278617 0.267119];

case { 'D4'  'BKR' }
descriptorname = 'diverging_bkr_55-10_c35_n256';
description = 'Diverging blue - black - red colour map';
map = [0.097481 0.507361 0.982050
       0.103297 0.503886 0.974284
       0.108633 0.500409 0.966535
       0.113690 0.496942 0.958799
       0.118405 0.493479 0.951074
       0.122753 0.490022 0.943359
       0.126920 0.486570 0.935656
       0.130887 0.483118 0.927961
       0.134599 0.479683 0.920283
       0.138107 0.476239 0.912619
       0.141429 0.472799 0.904966
       0.144583 0.469382 0.897324
       0.147583 0.465942 0.889689
       0.150429 0.462521 0.882070
       0.153122 0.459115 0.874463
       0.155737 0.455700 0.866867
       0.158196 0.452303 0.859287
       0.160509 0.448899 0.851717
       0.162773 0.445491 0.844160
       0.164876 0.442113 0.836617
       0.166931 0.438712 0.829089
       0.168858 0.435343 0.821568
       0.170678 0.431970 0.814057
       0.172420 0.428587 0.806569
       0.174093 0.425225 0.799088
       0.175656 0.421850 0.791617
       0.177184 0.418495 0.784167
       0.178617 0.415148 0.776724
       0.179947 0.411801 0.769289
       0.181219 0.408460 0.761874
       0.182432 0.405123 0.754478
       0.183587 0.401786 0.747087
       0.184682 0.398469 0.739716
       0.185722 0.395149 0.732352
       0.186652 0.391833 0.725003
       0.187577 0.388524 0.717669
       0.188410 0.385209 0.710345
       0.189179 0.381913 0.703038
       0.189904 0.378621 0.695745
       0.190575 0.375327 0.688466
       0.191197 0.372043 0.681192
       0.191771 0.368778 0.673945
       0.192295 0.365498 0.666704
       0.192771 0.362235 0.659485
       0.193200 0.358963 0.652276
       0.193585 0.355724 0.645078
       0.193926 0.352463 0.637901
       0.194219 0.349221 0.630741
       0.194462 0.345970 0.623590
       0.194658 0.342756 0.616451
       0.194810 0.339530 0.609331
       0.194920 0.336290 0.602220
       0.194991 0.333090 0.595129
       0.195023 0.329878 0.588048
       0.195016 0.326654 0.580991
       0.194972 0.323465 0.573945
       0.194891 0.320283 0.566922
       0.194773 0.317084 0.559906
       0.194618 0.313906 0.552901
       0.194424 0.310739 0.545926
       0.194189 0.307547 0.538955
       0.193914 0.304390 0.532003
       0.193601 0.301227 0.525066
       0.193255 0.298076 0.518150
       0.192877 0.294933 0.511239
       0.192466 0.291808 0.504364
       0.192021 0.288671 0.497491
       0.191543 0.285530 0.490638
       0.191034 0.282437 0.483788
       0.190494 0.279321 0.476972
       0.189924 0.276217 0.470174
       0.189322 0.273106 0.463376
       0.188694 0.270019 0.456611
       0.188042 0.266912 0.449863
       0.187332 0.263853 0.443130
       0.186584 0.260764 0.436412
       0.185849 0.257690 0.429711
       0.185050 0.254650 0.423021
       0.184211 0.251594 0.416365
       0.183353 0.248549 0.409723
       0.182467 0.245479 0.403094
       0.181558 0.242452 0.396479
       0.180624 0.239435 0.389886
       0.179667 0.236424 0.383328
       0.178690 0.233381 0.376776
       0.177679 0.230380 0.370229
       0.176586 0.227400 0.363716
       0.175513 0.224381 0.357237
       0.174425 0.221422 0.350760
       0.173274 0.218445 0.344285
       0.172110 0.215468 0.337853
       0.170926 0.212510 0.331431
       0.169731 0.209549 0.325035
       0.168466 0.206628 0.318668
       0.167215 0.203677 0.312297
       0.165883 0.200711 0.305991
       0.164583 0.197803 0.299665
       0.163243 0.194903 0.293360
       0.161874 0.191977 0.287084
       0.160423 0.189084 0.280829
       0.159022 0.186198 0.274609
       0.157547 0.183291 0.268409
       0.156101 0.180414 0.262223
       0.154562 0.177580 0.256049
       0.153019 0.174711 0.249901
       0.151466 0.171838 0.243786
       0.149912 0.169022 0.237716
       0.148272 0.166158 0.231642
       0.146647 0.163361 0.225586
       0.145053 0.160515 0.219569
       0.143390 0.157733 0.213573
       0.141750 0.154961 0.207634
       0.140091 0.152202 0.201710
       0.138507 0.149518 0.195876
       0.136925 0.146804 0.190082
       0.135409 0.144194 0.184372
       0.133999 0.141626 0.178763
       0.132675 0.139097 0.173238
       0.131502 0.136699 0.167865
       0.130527 0.134442 0.162677
       0.129726 0.132263 0.157636
       0.129157 0.130284 0.152835
       0.128909 0.128416 0.148300
       0.128989 0.126759 0.144071
       0.129445 0.125362 0.140114
       0.130306 0.124151 0.136519
       0.131506 0.123206 0.133323
       0.133152 0.122526 0.130511
       0.135227 0.122112 0.128042
       0.137745 0.121960 0.125997
       0.140645 0.122064 0.124289
       0.143939 0.122412 0.122986
       0.147617 0.122992 0.122050
       0.151605 0.123789 0.121448
       0.155954 0.124779 0.121147
       0.160502 0.125959 0.121113
       0.165315 0.127202 0.121310
       0.170362 0.128627 0.121709
       0.175534 0.130182 0.122277
       0.180848 0.131741 0.122988
       0.186297 0.133389 0.123817
       0.191796 0.135097 0.124738
       0.197384 0.136809 0.125750
       0.203005 0.138582 0.126745
       0.208675 0.140352 0.127838
       0.214374 0.142144 0.128910
       0.220102 0.143939 0.130082
       0.225835 0.145753 0.131190
       0.231583 0.147532 0.132325
       0.237347 0.149325 0.133473
       0.243077 0.151094 0.134644
       0.248867 0.152867 0.135743
       0.254647 0.154654 0.136919
       0.260415 0.156455 0.138097
       0.266210 0.158212 0.139199
       0.271982 0.159930 0.140378
       0.277808 0.161741 0.141543
       0.283615 0.163485 0.142685
       0.289442 0.165184 0.143839
       0.295259 0.166966 0.145010
       0.301095 0.168689 0.146141
       0.306952 0.170411 0.147295
       0.312802 0.172124 0.148424
       0.318674 0.173848 0.149617
       0.324542 0.175552 0.150745
       0.330433 0.177285 0.151888
       0.336324 0.178980 0.153037
       0.342239 0.180658 0.154197
       0.348152 0.182347 0.155364
       0.354070 0.184047 0.156536
       0.360007 0.185760 0.157661
       0.365958 0.187433 0.158838
       0.371902 0.189089 0.159950
       0.377875 0.190746 0.161151
       0.383854 0.192414 0.162305
       0.389833 0.194097 0.163465
       0.395843 0.195726 0.164605
       0.401837 0.197381 0.165733
       0.407863 0.199005 0.166929
       0.413894 0.200640 0.168055
       0.419937 0.202284 0.169249
       0.425981 0.203944 0.170381
       0.432059 0.205552 0.171530
       0.438125 0.207183 0.172685
       0.444211 0.208786 0.173848
       0.450299 0.210398 0.175019
       0.456400 0.212023 0.176139
       0.462512 0.213606 0.177339
       0.468644 0.215196 0.178500
       0.474788 0.216813 0.179636
       0.480922 0.218394 0.180784
       0.487089 0.219985 0.181940
       0.493249 0.221553 0.183104
       0.499428 0.223141 0.184275
       0.505616 0.224677 0.185462
       0.511822 0.226282 0.186595
       0.518041 0.227839 0.187786
       0.524258 0.229379 0.188923
       0.530491 0.230944 0.190075
       0.536741 0.232492 0.191232
       0.542991 0.234032 0.192399
       0.549254 0.235589 0.193572
       0.555519 0.237134 0.194744
       0.561815 0.238634 0.195890
       0.568104 0.240160 0.197064
       0.574405 0.241704 0.198205
       0.580727 0.243204 0.199359
       0.587051 0.244716 0.200520
       0.593392 0.246235 0.201689
       0.599740 0.247762 0.202864
       0.606090 0.249252 0.204054
       0.612458 0.250750 0.205195
       0.618847 0.252252 0.206389
       0.625227 0.253742 0.207535
       0.631627 0.255236 0.208694
       0.638034 0.256706 0.209859
       0.644449 0.258174 0.211031
       0.650882 0.259669 0.212209
       0.657324 0.261129 0.213352
       0.663772 0.262597 0.214528
       0.670231 0.264069 0.215706
       0.676697 0.265517 0.216877
       0.683172 0.266961 0.218039
       0.689663 0.268422 0.219209
       0.696160 0.269878 0.220391
       0.702666 0.271307 0.221542
       0.709187 0.272744 0.222721
       0.715714 0.274183 0.223874
       0.722242 0.275629 0.225037
       0.728791 0.277046 0.226238
       0.735349 0.278468 0.227400
       0.741913 0.279885 0.228581
       0.748490 0.281289 0.229719
       0.755077 0.282721 0.230912
       0.761666 0.284110 0.232093
       0.768268 0.285507 0.233238
       0.774882 0.286907 0.234440
       0.781510 0.288311 0.235614
       0.788138 0.289708 0.236792
       0.794781 0.291101 0.237955
       0.801430 0.292477 0.239123
       0.808092 0.293852 0.240283
       0.814759 0.295231 0.241481
       0.821440 0.296617 0.242641
       0.828124 0.297972 0.243811
       0.834824 0.299354 0.244987
       0.841530 0.300700 0.246170
       0.848243 0.302055 0.247360
       0.854970 0.303412 0.248549
       0.861703 0.304772 0.249698
       0.868449 0.306134 0.250892
       0.875195 0.307454 0.252075
       0.881956 0.308815 0.253251
       0.888727 0.310149 0.254437
       0.895504 0.311465 0.255606
       0.902290 0.312793 0.256787];

case { 'D5'  'GKR' }
descriptorname = 'diverging_gkr_60-10_c40_n256';
description = 'Diverging green - black - red colour map';
map = [0.210902 0.651143 0.087950
       0.211132 0.646445 0.089674
       0.211341 0.641753 0.091371
       0.211529 0.637073 0.092907
       0.211694 0.632393 0.094540
       0.211837 0.627721 0.095984
       0.211958 0.623057 0.097463
       0.212058 0.618398 0.098858
       0.212137 0.613735 0.100202
       0.212195 0.609100 0.101514
       0.212234 0.604457 0.102800
       0.212254 0.599819 0.104014
       0.212255 0.595188 0.105171
       0.212237 0.590564 0.106383
       0.212201 0.585955 0.107459
       0.212146 0.581339 0.108494
       0.212072 0.576744 0.109602
       0.211980 0.572143 0.110582
       0.211868 0.567553 0.111546
       0.211738 0.562973 0.112485
       0.211588 0.558383 0.113398
       0.211419 0.553817 0.114210
       0.211233 0.549258 0.115057
       0.211029 0.544698 0.115916
       0.210809 0.540144 0.116689
       0.210573 0.535596 0.117425
       0.210321 0.531064 0.118200
       0.210052 0.526539 0.118899
       0.209766 0.522000 0.119552
       0.209464 0.517495 0.120198
       0.209145 0.512978 0.120821
       0.208812 0.508482 0.121423
       0.208462 0.503980 0.122004
       0.208097 0.499479 0.122566
       0.207716 0.495005 0.123107
       0.207322 0.490531 0.123629
       0.206916 0.486051 0.124130
       0.206493 0.481592 0.124611
       0.206041 0.477134 0.125075
       0.205562 0.472684 0.125526
       0.205081 0.468240 0.125950
       0.204605 0.463811 0.126332
       0.204105 0.459385 0.126683
       0.203571 0.454966 0.127026
       0.203021 0.450548 0.127371
       0.202462 0.446135 0.127708
       0.201890 0.441747 0.128021
       0.201304 0.437356 0.128300
       0.200705 0.432966 0.128553
       0.200093 0.428587 0.128790
       0.199469 0.424214 0.129018
       0.198834 0.419857 0.129239
       0.198184 0.415498 0.129450
       0.197525 0.411152 0.129646
       0.196860 0.406811 0.129823
       0.196155 0.402472 0.129982
       0.195427 0.398158 0.130121
       0.194725 0.393833 0.130241
       0.193985 0.389525 0.130345
       0.193215 0.385218 0.130433
       0.192441 0.380939 0.130506
       0.191656 0.376653 0.130564
       0.190860 0.372359 0.130608
       0.190054 0.368094 0.130639
       0.189236 0.363828 0.130656
       0.188415 0.359577 0.130660
       0.187572 0.355343 0.130650
       0.186681 0.351098 0.130627
       0.185829 0.346866 0.130591
       0.184926 0.342645 0.130543
       0.184004 0.338416 0.130481
       0.183078 0.334223 0.130407
       0.182141 0.330022 0.130319
       0.181197 0.325814 0.130217
       0.180245 0.321637 0.130102
       0.179284 0.317468 0.129971
       0.178322 0.313305 0.129824
       0.177314 0.309160 0.129662
       0.176263 0.304998 0.129487
       0.175265 0.300846 0.129300
       0.174228 0.296730 0.129104
       0.173165 0.292594 0.128904
       0.172098 0.288483 0.128701
       0.171024 0.284371 0.128490
       0.169947 0.280271 0.128266
       0.168846 0.276206 0.128024
       0.167710 0.272095 0.127759
       0.166590 0.268027 0.127476
       0.165408 0.263990 0.127186
       0.164298 0.259929 0.126898
       0.163121 0.255871 0.126613
       0.161944 0.251853 0.126322
       0.160715 0.247828 0.126011
       0.159495 0.243783 0.125674
       0.158302 0.239787 0.125318
       0.157042 0.235800 0.124952
       0.155812 0.231813 0.124582
       0.154522 0.227827 0.124204
       0.153235 0.223838 0.123816
       0.151947 0.219891 0.123418
       0.150653 0.215944 0.123011
       0.149348 0.212000 0.122596
       0.148003 0.208049 0.122172
       0.146641 0.204143 0.121740
       0.145328 0.200184 0.121299
       0.143930 0.196317 0.120852
       0.142539 0.192398 0.120395
       0.141163 0.188528 0.119930
       0.139704 0.184645 0.119460
       0.138351 0.180770 0.118993
       0.136935 0.176947 0.118516
       0.135534 0.173128 0.118002
       0.134202 0.169377 0.117468
       0.132860 0.165579 0.116969
       0.131584 0.161930 0.116496
       0.130405 0.158283 0.116003
       0.129247 0.154698 0.115493
       0.128285 0.151227 0.114995
       0.127436 0.147888 0.114535
       0.126802 0.144683 0.114131
       0.126430 0.141629 0.113778
       0.126315 0.138755 0.113463
       0.126495 0.136086 0.113188
       0.127011 0.133735 0.112967
       0.127976 0.131616 0.112816
       0.129294 0.129806 0.112746
       0.131090 0.128277 0.112762
       0.133298 0.127052 0.112870
       0.135926 0.126235 0.113069
       0.139048 0.125709 0.113354
       0.142606 0.125498 0.113710
       0.146528 0.125610 0.114137
       0.150855 0.126019 0.114665
       0.155528 0.126658 0.115308
       0.160469 0.127581 0.116026
       0.165702 0.128711 0.116746
       0.171188 0.130083 0.117519
       0.176847 0.131540 0.118395
       0.182657 0.133154 0.119230
       0.188635 0.134897 0.120102
       0.194697 0.136671 0.121002
       0.200792 0.138570 0.121923
       0.207026 0.140501 0.122863
       0.213244 0.142461 0.123818
       0.219545 0.144458 0.124783
       0.225861 0.146442 0.125771
       0.232187 0.148452 0.126694
       0.238513 0.150497 0.127663
       0.244868 0.152502 0.128614
       0.251268 0.154537 0.129587
       0.257629 0.156584 0.130564
       0.264031 0.158601 0.131493
       0.270427 0.160578 0.132441
       0.276822 0.162611 0.133388
       0.283231 0.164603 0.134350
       0.289648 0.166602 0.135276
       0.296074 0.168584 0.136184
       0.302505 0.170564 0.137166
       0.308982 0.172534 0.138119
       0.315433 0.174527 0.139015
       0.321899 0.176464 0.139942
       0.328386 0.178466 0.140925
       0.334894 0.180392 0.141839
       0.341388 0.182339 0.142766
       0.347907 0.184303 0.143696
       0.354436 0.186256 0.144629
       0.360965 0.188209 0.145571
       0.367520 0.190115 0.146449
       0.374099 0.192042 0.147391
       0.380677 0.193990 0.148287
       0.387255 0.195892 0.149224
       0.393861 0.197799 0.150148
       0.400489 0.199693 0.151039
       0.407107 0.201601 0.151945
       0.413749 0.203526 0.152851
       0.420399 0.205411 0.153762
       0.427074 0.207313 0.154675
       0.433739 0.209188 0.155597
       0.440436 0.211076 0.156511
       0.447140 0.212938 0.157378
       0.453859 0.214814 0.158308
       0.460583 0.216707 0.159189
       0.467337 0.218562 0.160056
       0.474087 0.220436 0.160990
       0.480851 0.222269 0.161898
       0.487645 0.224108 0.162778
       0.494436 0.225986 0.163679
       0.501242 0.227823 0.164551
       0.508064 0.229635 0.165402
       0.514890 0.231500 0.166313
       0.521727 0.233299 0.167213
       0.528595 0.235161 0.168067
       0.535456 0.236984 0.168978
       0.542345 0.238773 0.169850
       0.549241 0.240579 0.170711
       0.556138 0.242400 0.171580
       0.563071 0.244195 0.172452
       0.569993 0.246002 0.173325
       0.576947 0.247822 0.174206
       0.583899 0.249593 0.175079
       0.590869 0.251407 0.175911
       0.597851 0.253184 0.176794
       0.604849 0.254977 0.177690
       0.611854 0.256739 0.178544
       0.618882 0.258510 0.179386
       0.625905 0.260285 0.180236
       0.632951 0.262050 0.181087
       0.640009 0.263827 0.181941
       0.647076 0.265573 0.182798
       0.654154 0.267310 0.183658
       0.661246 0.269093 0.184518
       0.668354 0.270837 0.185390
       0.675466 0.272561 0.186224
       0.682593 0.274307 0.187066
       0.689737 0.276063 0.187934
       0.696886 0.277784 0.188757
       0.704049 0.279520 0.189590
       0.711229 0.281225 0.190425
       0.718410 0.282967 0.191262
       0.725615 0.284666 0.192102
       0.732824 0.286377 0.192943
       0.740047 0.288095 0.193791
       0.747279 0.289808 0.194631
       0.754526 0.291523 0.195435
       0.761776 0.293208 0.196286
       0.769043 0.294905 0.197111
       0.776327 0.296612 0.197919
       0.783618 0.298288 0.198737
       0.790917 0.299983 0.199554
       0.798230 0.301657 0.200375
       0.805553 0.303338 0.201198
       0.812889 0.305028 0.202023
       0.820235 0.306704 0.202848
       0.827595 0.308369 0.203683
       0.834966 0.310045 0.204502
       0.842348 0.311686 0.205295
       0.849738 0.313362 0.206132
       0.857137 0.315009 0.206939
       0.864552 0.316671 0.207734
       0.871978 0.318318 0.208538
       0.879408 0.319973 0.209341
       0.886857 0.321603 0.210147
       0.894314 0.323252 0.210954
       0.901778 0.324876 0.211771
       0.909255 0.326509 0.212556
       0.916746 0.328147 0.213343
       0.924242 0.329786 0.214158
       0.931750 0.331390 0.214936
       0.939272 0.333032 0.215752
       0.946801 0.334651 0.216553
       0.954343 0.336247 0.217333
       0.961890 0.337860 0.218123
       0.969454 0.339483 0.218912
       0.977027 0.341079 0.219702
       0.984612 0.342688 0.220501
       0.992203 0.344268 0.221276];

case { 'D6'  'BKY' }
descriptorname = 'diverging_bky_60-10_c30_n256';
description = 'Diverging blue - black - yellow colour map';
map = [0.056674 0.579589 0.981211
       0.066107 0.575466 0.973449
       0.074463 0.571347 0.965697
       0.081994 0.567245 0.957964
       0.088742 0.563143 0.950235
       0.094990 0.559037 0.942524
       0.100668 0.554947 0.934817
       0.106026 0.550862 0.927125
       0.110979 0.546784 0.919448
       0.115599 0.542710 0.911783
       0.119927 0.538643 0.904126
       0.124011 0.534573 0.896487
       0.127903 0.530516 0.888856
       0.131547 0.526469 0.881237
       0.135004 0.522413 0.873629
       0.138270 0.518381 0.866036
       0.141367 0.514348 0.858452
       0.144295 0.510311 0.850882
       0.147074 0.506285 0.843327
       0.149754 0.502273 0.835785
       0.152239 0.498272 0.828252
       0.154646 0.494261 0.820733
       0.156937 0.490260 0.813227
       0.159112 0.486263 0.805734
       0.161195 0.482281 0.798256
       0.163157 0.478296 0.790788
       0.164994 0.474325 0.783335
       0.166804 0.470358 0.775894
       0.168476 0.466392 0.768463
       0.170089 0.462431 0.761053
       0.171597 0.458486 0.753650
       0.173040 0.454550 0.746262
       0.174423 0.450607 0.738893
       0.175690 0.446675 0.731527
       0.176930 0.442755 0.724180
       0.178112 0.438826 0.716852
       0.179174 0.434926 0.709530
       0.180186 0.431019 0.702218
       0.181139 0.427123 0.694926
       0.182037 0.423217 0.687651
       0.182879 0.419343 0.680381
       0.183668 0.415458 0.673139
       0.184401 0.411584 0.665901
       0.185089 0.407720 0.658676
       0.185715 0.403866 0.651475
       0.186261 0.400007 0.644276
       0.186764 0.396167 0.637104
       0.187242 0.392324 0.629937
       0.187671 0.388497 0.622789
       0.188037 0.384665 0.615652
       0.188349 0.380860 0.608542
       0.188614 0.377046 0.601429
       0.188837 0.373235 0.594342
       0.189018 0.369440 0.587265
       0.189156 0.365652 0.580213
       0.189253 0.361874 0.573171
       0.189307 0.358098 0.566139
       0.189321 0.354327 0.559125
       0.189295 0.350575 0.552136
       0.189229 0.346817 0.545152
       0.189124 0.343075 0.538193
       0.188981 0.339338 0.531238
       0.188801 0.335602 0.524305
       0.188585 0.331872 0.517395
       0.188335 0.328153 0.510486
       0.188048 0.324439 0.503602
       0.187718 0.320755 0.496733
       0.187341 0.317052 0.489884
       0.186924 0.313369 0.483051
       0.186487 0.309698 0.476239
       0.186034 0.306032 0.469447
       0.185539 0.302342 0.462648
       0.184988 0.298699 0.455889
       0.184404 0.295046 0.449148
       0.183793 0.291424 0.442419
       0.183151 0.287776 0.435705
       0.182479 0.284157 0.429005
       0.181777 0.280534 0.422320
       0.181047 0.276949 0.415668
       0.180290 0.273354 0.409028
       0.179504 0.269768 0.402399
       0.178696 0.266184 0.395809
       0.177856 0.262607 0.389222
       0.176942 0.259054 0.382651
       0.176002 0.255492 0.376111
       0.175093 0.251950 0.369576
       0.174112 0.248421 0.363067
       0.173099 0.244859 0.356589
       0.172066 0.241360 0.350119
       0.171010 0.237854 0.343661
       0.169936 0.234353 0.337239
       0.168820 0.230852 0.330817
       0.167659 0.227374 0.324429
       0.166491 0.223885 0.318067
       0.165257 0.220449 0.311707
       0.164079 0.216983 0.305401
       0.162808 0.213522 0.299093
       0.161537 0.210090 0.292796
       0.160168 0.206688 0.286525
       0.158869 0.203249 0.280287
       0.157476 0.199820 0.274080
       0.156120 0.196464 0.267864
       0.154668 0.193059 0.261706
       0.153211 0.189680 0.255556
       0.151742 0.186320 0.249417
       0.150264 0.182945 0.243305
       0.148707 0.179608 0.237253
       0.147176 0.176253 0.231178
       0.145637 0.172953 0.225121
       0.144027 0.169679 0.219133
       0.142425 0.166382 0.213140
       0.140840 0.163143 0.207226
       0.139188 0.159878 0.201301
       0.137646 0.156737 0.195467
       0.136019 0.153582 0.189683
       0.134550 0.150536 0.183961
       0.133060 0.147565 0.178357
       0.131674 0.144687 0.172798
       0.130417 0.141919 0.167412
       0.129217 0.139275 0.162165
       0.128252 0.136863 0.157060
       0.127432 0.134664 0.152174
       0.126854 0.132648 0.147558
       0.126555 0.130927 0.143171
       0.126522 0.129463 0.139054
       0.126776 0.128336 0.135297
       0.127372 0.127517 0.131860
       0.128329 0.127042 0.128745
       0.129609 0.126937 0.126043
       0.131230 0.127180 0.123616
       0.133180 0.127798 0.121587
       0.135446 0.128700 0.119925
       0.138066 0.129975 0.118611
       0.140933 0.131471 0.117510
       0.144037 0.133248 0.116741
       0.147386 0.135262 0.116219
       0.150916 0.137499 0.115879
       0.154626 0.139843 0.115718
       0.158506 0.142419 0.115715
       0.162469 0.145108 0.115839
       0.166523 0.147853 0.116062
       0.170648 0.150690 0.116353
       0.174854 0.153587 0.116684
       0.179077 0.156581 0.117050
       0.183330 0.159546 0.117466
       0.187660 0.162609 0.117928
       0.191938 0.165628 0.118394
       0.196284 0.168741 0.118836
       0.200582 0.171816 0.119267
       0.204950 0.174945 0.119708
       0.209296 0.178078 0.120148
       0.213651 0.181161 0.120581
       0.218015 0.184313 0.121006
       0.222373 0.187482 0.121423
       0.226750 0.190608 0.121835
       0.231121 0.193791 0.122239
       0.235506 0.196965 0.122636
       0.239871 0.200107 0.123026
       0.244248 0.203320 0.123409
       0.248672 0.206539 0.123784
       0.253063 0.209717 0.124151
       0.257465 0.212921 0.124508
       0.261879 0.216175 0.124859
       0.266303 0.219390 0.125205
       0.270736 0.222629 0.125546
       0.275157 0.225882 0.125873
       0.279601 0.229123 0.126178
       0.284032 0.232387 0.126462
       0.288486 0.235664 0.126735
       0.292940 0.238922 0.127005
       0.297415 0.242211 0.127281
       0.301871 0.245491 0.127558
       0.306373 0.248815 0.127825
       0.310858 0.252118 0.128074
       0.315341 0.255432 0.128303
       0.319846 0.258752 0.128517
       0.324339 0.262076 0.128720
       0.328866 0.265410 0.128918
       0.333387 0.268767 0.129113
       0.337902 0.272090 0.129305
       0.342448 0.275477 0.129491
       0.346991 0.278837 0.129666
       0.351529 0.282213 0.129830
       0.356094 0.285567 0.129980
       0.360639 0.288972 0.130116
       0.365218 0.292356 0.130238
       0.369793 0.295753 0.130348
       0.374389 0.299173 0.130446
       0.378972 0.302566 0.130533
       0.383577 0.306013 0.130608
       0.388177 0.309438 0.130672
       0.392788 0.312853 0.130725
       0.397410 0.316307 0.130768
       0.402030 0.319751 0.130799
       0.406676 0.323203 0.130819
       0.411315 0.326649 0.130827
       0.415965 0.330129 0.130824
       0.420617 0.333603 0.130810
       0.425290 0.337079 0.130784
       0.429957 0.340563 0.130746
       0.434635 0.344042 0.130695
       0.439315 0.347557 0.130633
       0.444016 0.351060 0.130558
       0.448715 0.354570 0.130471
       0.453414 0.358083 0.130370
       0.458126 0.361604 0.130254
       0.462840 0.365126 0.130124
       0.467578 0.368670 0.129976
       0.472314 0.372198 0.129810
       0.477051 0.375749 0.129627
       0.481802 0.379302 0.129429
       0.486554 0.382863 0.129218
       0.491328 0.386418 0.128998
       0.496079 0.389989 0.128774
       0.500863 0.393569 0.128542
       0.505635 0.397148 0.128296
       0.510432 0.400754 0.128026
       0.515228 0.404331 0.127726
       0.520028 0.407934 0.127403
       0.524844 0.411536 0.127072
       0.529664 0.415149 0.126745
       0.534484 0.418763 0.126413
       0.539312 0.422381 0.126058
       0.544153 0.426011 0.125666
       0.549001 0.429650 0.125245
       0.553845 0.433285 0.124809
       0.558698 0.436937 0.124361
       0.563574 0.440577 0.123897
       0.568436 0.444245 0.123414
       0.573320 0.447891 0.122914
       0.578196 0.451567 0.122397
       0.583088 0.455239 0.121862
       0.587983 0.458915 0.121309
       0.592897 0.462588 0.120740
       0.597801 0.466280 0.120150
       0.602719 0.469981 0.119541
       0.607649 0.473667 0.118927
       0.612569 0.477375 0.118276
       0.617508 0.481079 0.117550
       0.622449 0.484792 0.116850
       0.627401 0.488520 0.116147
       0.632358 0.492240 0.115356
       0.637321 0.495965 0.114544
       0.642290 0.499697 0.113783
       0.647264 0.503438 0.112978
       0.652245 0.507188 0.112101
       0.657236 0.510935 0.111213
       0.662231 0.514696 0.110307
       0.667225 0.518458 0.109363
       0.672234 0.522211 0.108320
       0.677250 0.525989 0.107358
       0.682260 0.529764 0.106343
       0.687288 0.533535 0.105206
       0.692325 0.537331 0.104115
       0.697354 0.541110 0.103000
       0.702399 0.544909 0.101796];

case { 'D7'  'BJY'  'DIVBJY' }
descriptorname = 'diverging-linear_bjy_30-90_c45_n256';
description = 'Linear-diverging blue - grey - yellow colour map. This kind of diverging map has no perceptual dead spot at the centre';
map = [0.077854 0.193434 0.758268
       0.095213 0.196286 0.756899
       0.110071 0.199096 0.755533
       0.123169 0.201942 0.754165
       0.135136 0.204807 0.752787
       0.146079 0.207651 0.751425
       0.156256 0.210493 0.750044
       0.165720 0.213331 0.748672
       0.174718 0.216212 0.747300
       0.183181 0.219046 0.745919
       0.191264 0.221884 0.744542
       0.198972 0.224721 0.743167
       0.206408 0.227604 0.741781
       0.213482 0.230440 0.740401
       0.220352 0.233290 0.739022
       0.226948 0.236174 0.737630
       0.233316 0.239017 0.736242
       0.239529 0.241884 0.734862
       0.245521 0.244723 0.733470
       0.251386 0.247608 0.732082
       0.257059 0.250450 0.730686
       0.262589 0.253330 0.729293
       0.267969 0.256179 0.727897
       0.273256 0.259069 0.726502
       0.278398 0.261923 0.725108
       0.283426 0.264783 0.723702
       0.288342 0.267642 0.722298
       0.293158 0.270547 0.720897
       0.297879 0.273413 0.719498
       0.302504 0.276292 0.718089
       0.307055 0.279164 0.716685
       0.311507 0.282038 0.715273
       0.315902 0.284897 0.713859
       0.320203 0.287777 0.712443
       0.324412 0.290661 0.711035
       0.328578 0.293540 0.709618
       0.332677 0.296434 0.708197
       0.336686 0.299324 0.706780
       0.340652 0.302190 0.705354
       0.344545 0.305094 0.703930
       0.348395 0.307970 0.702504
       0.352174 0.310880 0.701076
       0.355920 0.313767 0.699643
       0.359593 0.316658 0.698211
       0.363219 0.319553 0.696781
       0.366808 0.322454 0.695345
       0.370332 0.325334 0.693902
       0.373829 0.328242 0.692473
       0.377271 0.331132 0.691020
       0.380675 0.334053 0.689584
       0.384026 0.336952 0.688136
       0.387337 0.339863 0.686686
       0.390621 0.342771 0.685237
       0.393856 0.345663 0.683779
       0.397054 0.348581 0.682324
       0.400228 0.351488 0.680864
       0.403358 0.354406 0.679406
       0.406450 0.357329 0.677951
       0.409509 0.360227 0.676485
       0.412543 0.363146 0.675016
       0.415526 0.366075 0.673548
       0.418484 0.368994 0.672077
       0.421415 0.371905 0.670607
       0.424321 0.374836 0.669127
       0.427202 0.377761 0.667649
       0.430038 0.380694 0.666162
       0.432854 0.383621 0.664676
       0.435644 0.386539 0.663197
       0.438397 0.389477 0.661707
       0.441125 0.392409 0.660210
       0.443840 0.395351 0.658712
       0.446516 0.398287 0.657222
       0.449177 0.401228 0.655720
       0.451808 0.404160 0.654212
       0.454416 0.407102 0.652703
       0.456990 0.410054 0.651200
       0.459554 0.412999 0.649689
       0.462087 0.415936 0.648167
       0.464610 0.418884 0.646654
       0.467099 0.421828 0.645128
       0.469573 0.424791 0.643604
       0.472019 0.427746 0.642081
       0.474443 0.430697 0.640551
       0.476846 0.433643 0.639016
       0.479235 0.436612 0.637480
       0.481599 0.439568 0.635941
       0.483939 0.442536 0.634401
       0.486269 0.445488 0.632850
       0.488580 0.448460 0.631302
       0.490877 0.451427 0.629744
       0.493131 0.454398 0.628193
       0.495389 0.457359 0.626627
       0.497630 0.460325 0.625065
       0.499830 0.463298 0.623503
       0.502038 0.466278 0.621921
       0.504223 0.469265 0.620358
       0.506374 0.472239 0.618780
       0.508533 0.475222 0.617192
       0.510652 0.478194 0.615602
       0.512766 0.481174 0.614010
       0.514866 0.484158 0.612418
       0.516948 0.487153 0.610827
       0.519015 0.490137 0.609227
       0.521058 0.493121 0.607622
       0.523095 0.496112 0.606001
       0.525109 0.499110 0.604398
       0.527116 0.502108 0.602775
       0.529106 0.505100 0.601151
       0.531072 0.508108 0.599530
       0.533031 0.511097 0.597893
       0.534973 0.514109 0.596260
       0.536910 0.517111 0.594620
       0.538822 0.520104 0.592981
       0.540719 0.523121 0.591328
       0.542610 0.526129 0.589669
       0.544480 0.529142 0.588011
       0.546345 0.532150 0.586355
       0.548190 0.535158 0.584688
       0.550019 0.538182 0.583010
       0.551841 0.541187 0.581334
       0.553639 0.544210 0.579660
       0.555431 0.547229 0.577961
       0.557224 0.550253 0.576283
       0.558981 0.553270 0.574571
       0.560738 0.556295 0.572879
       0.562489 0.559327 0.571163
       0.564216 0.562363 0.569448
       0.565927 0.565379 0.567735
       0.568832 0.568090 0.565696
       0.572924 0.570471 0.563357
       0.576982 0.572868 0.560993
       0.581013 0.575253 0.558628
       0.585030 0.577643 0.556260
       0.589010 0.580050 0.553890
       0.592988 0.582444 0.551513
       0.596932 0.584856 0.549126
       0.600851 0.587258 0.546731
       0.604759 0.589669 0.544326
       0.608647 0.592096 0.541912
       0.612500 0.594509 0.539495
       0.616351 0.596936 0.537083
       0.620180 0.599362 0.534645
       0.623987 0.601786 0.532209
       0.627773 0.604226 0.529762
       0.631545 0.606658 0.527307
       0.635296 0.609102 0.524842
       0.639035 0.611540 0.522367
       0.642757 0.613979 0.519886
       0.646462 0.616437 0.517410
       0.650152 0.618894 0.514908
       0.653824 0.621341 0.512399
       0.657488 0.623807 0.509886
       0.661125 0.626262 0.507364
       0.664752 0.628732 0.504833
       0.668374 0.631200 0.502287
       0.671972 0.633672 0.499727
       0.675558 0.636146 0.497173
       0.679130 0.638621 0.494605
       0.682692 0.641097 0.492020
       0.686241 0.643582 0.489417
       0.689779 0.646070 0.486819
       0.693301 0.648561 0.484198
       0.696810 0.651055 0.481577
       0.700309 0.653547 0.478939
       0.703796 0.656053 0.476291
       0.707272 0.658548 0.473621
       0.710739 0.661052 0.470957
       0.714188 0.663566 0.468264
       0.717637 0.666070 0.465561
       0.721062 0.668587 0.462848
       0.724487 0.671101 0.460127
       0.727896 0.673619 0.457391
       0.731297 0.676140 0.454649
       0.734694 0.678663 0.451881
       0.738069 0.681185 0.449100
       0.741441 0.683718 0.446295
       0.744807 0.686253 0.443490
       0.748164 0.688792 0.440655
       0.751511 0.691323 0.437827
       0.754846 0.693862 0.434967
       0.758172 0.696412 0.432097
       0.761486 0.698952 0.429194
       0.764800 0.701501 0.426279
       0.768097 0.704055 0.423351
       0.771397 0.706613 0.420409
       0.774677 0.709170 0.417454
       0.777960 0.711727 0.414464
       0.781230 0.714283 0.411459
       0.784496 0.716855 0.408439
       0.787746 0.719419 0.405399
       0.790994 0.721982 0.402323
       0.794239 0.724559 0.399236
       0.797468 0.727131 0.396138
       0.800697 0.729711 0.393000
       0.803918 0.732292 0.389841
       0.807133 0.734873 0.386660
       0.810334 0.737453 0.383469
       0.813535 0.740045 0.380234
       0.816733 0.742633 0.376981
       0.819916 0.745220 0.373694
       0.823101 0.747819 0.370373
       0.826277 0.750413 0.367037
       0.829450 0.753006 0.363662
       0.832609 0.755613 0.360258
       0.835768 0.758218 0.356839
       0.838919 0.760822 0.353363
       0.842069 0.763427 0.349867
       0.845208 0.766041 0.346322
       0.848342 0.768649 0.342765
       0.851471 0.771271 0.339156
       0.854599 0.773882 0.335507
       0.857717 0.776508 0.331807
       0.860833 0.779126 0.328076
       0.863948 0.781755 0.324297
       0.867047 0.784383 0.320497
       0.870146 0.787011 0.316622
       0.873245 0.789639 0.312694
       0.876334 0.792272 0.308747
       0.879420 0.794907 0.304717
       0.882501 0.797543 0.300635
       0.885582 0.800188 0.296514
       0.888656 0.802826 0.292316
       0.891723 0.805469 0.288056
       0.894790 0.808119 0.283738
       0.897847 0.810765 0.279360
       0.900903 0.813415 0.274880
       0.903954 0.816074 0.270356
       0.907005 0.818723 0.265725
       0.910050 0.821384 0.261011
       0.913089 0.824041 0.256201
       0.916121 0.826706 0.251329
       0.919154 0.829372 0.246310
       0.922188 0.832037 0.241211
       0.925209 0.834703 0.235999
       0.928230 0.837375 0.230635
       0.931251 0.840048 0.225149
       0.934264 0.842720 0.219543
       0.937280 0.845393 0.213759
       0.940289 0.848071 0.207813
       0.943293 0.850750 0.201663
       0.946291 0.853432 0.195346
       0.949290 0.856120 0.188807
       0.952285 0.858804 0.181989
       0.955281 0.861494 0.174956
       0.958271 0.864188 0.167579
       0.961255 0.866875 0.159837
       0.964236 0.869570 0.151746
       0.967221 0.872270 0.143204
       0.970196 0.874965 0.134118
       0.973173 0.877666 0.124344
       0.976142 0.880367 0.113845
       0.979114 0.883071 0.102329
       0.982084 0.885780 0.089538
       0.985048 0.888488 0.074910
       0.988006 0.891196 0.057534
       0.990967 0.893909 0.035647];

case { 'D8'  'BJR' }
descriptorname = 'diverging-linear_bjr_30-55_c53_n256';
description = 'Linear-diverging blue - grey - red';
map = [0.000000 0.165400 0.843837
       0.000000 0.167452 0.840236
       0.000000 0.169477 0.836628
       0.000000 0.171449 0.833023
       0.000000 0.173444 0.829426
       0.000000 0.175433 0.825821
       0.000000 0.177441 0.822228
       0.000000 0.179397 0.818626
       0.025922 0.181352 0.815035
       0.052828 0.183324 0.811445
       0.073384 0.185317 0.807854
       0.089868 0.187262 0.804267
       0.103966 0.189199 0.800679
       0.116407 0.191133 0.797092
       0.127542 0.193080 0.793516
       0.137761 0.195019 0.789931
       0.147117 0.196957 0.786356
       0.155861 0.198849 0.782774
       0.164034 0.200761 0.779199
       0.171680 0.202688 0.775625
       0.178949 0.204618 0.772055
       0.185848 0.206535 0.768481
       0.192370 0.208410 0.764919
       0.198609 0.210304 0.761349
       0.204610 0.212211 0.757786
       0.210322 0.214092 0.754225
       0.215835 0.215986 0.750664
       0.221125 0.217853 0.747103
       0.226217 0.219731 0.743546
       0.231121 0.221593 0.739992
       0.235866 0.223468 0.736430
       0.240421 0.225322 0.732883
       0.244857 0.227192 0.729330
       0.249169 0.229048 0.725784
       0.253334 0.230898 0.722228
       0.257367 0.232740 0.718684
       0.261283 0.234608 0.715143
       0.265078 0.236446 0.711602
       0.268795 0.238266 0.708057
       0.272364 0.240098 0.704517
       0.275890 0.241945 0.700980
       0.279285 0.243756 0.697440
       0.282598 0.245584 0.693902
       0.285796 0.247424 0.690371
       0.288957 0.249239 0.686840
       0.292015 0.251065 0.683310
       0.294989 0.252875 0.679776
       0.297899 0.254696 0.676253
       0.300736 0.256484 0.672730
       0.303503 0.258286 0.669202
       0.306227 0.260100 0.665677
       0.308862 0.261897 0.662159
       0.311417 0.263709 0.658629
       0.313936 0.265488 0.655108
       0.316391 0.267264 0.651596
       0.318778 0.269083 0.648072
       0.321117 0.270866 0.644555
       0.323399 0.272631 0.641040
       0.325617 0.274415 0.637530
       0.327800 0.276213 0.634022
       0.329939 0.277979 0.630510
       0.332012 0.279755 0.626993
       0.334044 0.281517 0.623489
       0.336015 0.283293 0.619980
       0.337949 0.285045 0.616470
       0.339853 0.286808 0.612956
       0.341693 0.288582 0.609457
       0.343495 0.290332 0.605944
       0.345255 0.292101 0.602444
       0.346981 0.293846 0.598943
       0.348654 0.295602 0.595438
       0.350298 0.297364 0.591943
       0.351882 0.299115 0.588431
       0.353449 0.300846 0.584939
       0.354988 0.302588 0.581434
       0.356467 0.304338 0.577931
       0.357919 0.306097 0.574432
       0.359322 0.307811 0.570936
       0.360692 0.309570 0.567445
       0.362053 0.311292 0.563951
       0.363354 0.313023 0.560449
       0.364631 0.314747 0.556957
       0.365882 0.316490 0.553450
       0.367088 0.318212 0.549959
       0.368271 0.319945 0.546469
       0.369415 0.321654 0.542970
       0.370525 0.323382 0.539466
       0.371609 0.325091 0.535971
       0.372666 0.326809 0.532482
       0.373699 0.328536 0.528989
       0.374695 0.330250 0.525486
       0.375655 0.331962 0.521981
       0.376604 0.333677 0.518497
       0.377504 0.335391 0.514994
       0.378383 0.337090 0.511492
       0.379236 0.338794 0.508003
       0.380062 0.340498 0.504505
       0.380867 0.342202 0.501000
       0.381632 0.343890 0.497503
       0.382369 0.345591 0.493997
       0.383098 0.347303 0.490498
       0.383787 0.348991 0.486991
       0.384445 0.350693 0.483479
       0.385083 0.352373 0.479982
       0.385696 0.354062 0.476471
       0.386285 0.355764 0.472955
       0.386851 0.357455 0.469460
       0.387391 0.359123 0.465933
       0.387909 0.360802 0.462420
       0.388405 0.362497 0.458912
       0.388875 0.364174 0.455394
       0.389314 0.365862 0.451878
       0.389726 0.367530 0.448350
       0.390120 0.369216 0.444831
       0.390497 0.370883 0.441297
       0.390851 0.372559 0.437777
       0.391179 0.374245 0.434231
       0.391480 0.375911 0.430704
       0.391757 0.377580 0.427170
       0.392011 0.379248 0.423615
       0.392244 0.380925 0.420076
       0.392455 0.382580 0.416529
       0.392644 0.384245 0.412982
       0.392812 0.385903 0.409420
       0.392957 0.387568 0.405865
       0.393080 0.389238 0.402285
       0.393182 0.390899 0.398722
       0.393262 0.392550 0.395151
       0.396690 0.393142 0.392138
       0.403430 0.392662 0.389690
       0.410070 0.392170 0.387243
       0.416616 0.391670 0.384798
       0.423076 0.391156 0.382351
       0.429475 0.390621 0.379909
       0.435797 0.390071 0.377461
       0.442047 0.389522 0.375008
       0.448216 0.388962 0.372554
       0.454339 0.388377 0.370102
       0.460380 0.387778 0.367648
       0.466384 0.387170 0.365197
       0.472328 0.386549 0.362742
       0.478210 0.385916 0.360280
       0.484042 0.385271 0.357838
       0.489829 0.384612 0.355381
       0.495572 0.383944 0.352911
       0.501273 0.383261 0.350453
       0.506917 0.382549 0.347983
       0.512526 0.381833 0.345509
       0.518102 0.381112 0.343053
       0.523630 0.380361 0.340577
       0.529128 0.379596 0.338095
       0.534578 0.378820 0.335634
       0.539997 0.378029 0.333161
       0.545392 0.377226 0.330668
       0.550748 0.376409 0.328192
       0.556066 0.375558 0.325699
       0.561369 0.374711 0.323229
       0.566640 0.373842 0.320748
       0.571876 0.372947 0.318247
       0.577090 0.372041 0.315765
       0.582270 0.371120 0.313261
       0.587431 0.370185 0.310773
       0.592578 0.369237 0.308257
       0.597683 0.368266 0.305770
       0.602773 0.367269 0.303239
       0.607846 0.366273 0.300730
       0.612882 0.365237 0.298215
       0.617910 0.364191 0.295702
       0.622918 0.363128 0.293179
       0.627904 0.362054 0.290656
       0.632869 0.360938 0.288128
       0.637816 0.359827 0.285589
       0.642747 0.358677 0.283074
       0.647657 0.357536 0.280511
       0.652551 0.356348 0.277985
       0.657438 0.355152 0.275443
       0.662302 0.353909 0.272876
       0.667144 0.352671 0.270333
       0.671978 0.351402 0.267737
       0.676799 0.350118 0.265179
       0.681595 0.348802 0.262614
       0.686390 0.347473 0.260036
       0.691164 0.346097 0.257446
       0.695935 0.344723 0.254871
       0.700685 0.343323 0.252263
       0.705423 0.341890 0.249641
       0.710152 0.340434 0.247041
       0.714865 0.338956 0.244409
       0.719574 0.337446 0.241806
       0.724265 0.335907 0.239168
       0.728947 0.334357 0.236542
       0.733622 0.332772 0.233872
       0.738282 0.331132 0.231235
       0.742941 0.329510 0.228576
       0.747584 0.327820 0.225901
       0.752213 0.326109 0.223215
       0.756839 0.324379 0.220533
       0.761454 0.322630 0.217815
       0.766065 0.320832 0.215085
       0.770664 0.318995 0.212382
       0.775252 0.317132 0.209637
       0.779837 0.315234 0.206911
       0.784419 0.313307 0.204156
       0.788986 0.311335 0.201345
       0.793549 0.309348 0.198568
       0.798097 0.307287 0.195781
       0.802644 0.305221 0.192971
       0.807189 0.303086 0.190139
       0.811722 0.300929 0.187317
       0.816252 0.298739 0.184442
       0.820769 0.296500 0.181548
       0.825285 0.294202 0.178685
       0.829800 0.291885 0.175740
       0.834305 0.289508 0.172820
       0.838800 0.287067 0.169888
       0.843293 0.284596 0.166918
       0.847782 0.282090 0.163930
       0.852272 0.279515 0.160884
       0.856745 0.276880 0.157832
       0.861219 0.274187 0.154755
       0.865690 0.271435 0.151649
       0.870152 0.268647 0.148510
       0.874615 0.265779 0.145400
       0.879071 0.262846 0.142176
       0.883527 0.259852 0.138928
       0.887972 0.256779 0.135649
       0.892420 0.253643 0.132370
       0.896862 0.250411 0.128991
       0.901296 0.247130 0.125627
       0.905733 0.243740 0.122129
       0.910162 0.240276 0.118692
       0.914587 0.236756 0.115085
       0.919007 0.233078 0.111512
       0.923433 0.229337 0.107793
       0.927845 0.225494 0.104048
       0.932259 0.221527 0.100205
       0.936677 0.217450 0.096258
       0.941082 0.213225 0.092292
       0.945490 0.208896 0.088179
       0.949892 0.204434 0.083966
       0.954295 0.199762 0.079537
       0.958695 0.194994 0.075045
       0.963090 0.190011 0.070373
       0.967486 0.184862 0.065518
       0.971877 0.179480 0.060460
       0.976264 0.173886 0.055081
       0.980655 0.168034 0.049511
       0.985043 0.161951 0.043373
       0.989428 0.155509 0.037065
       0.993809 0.148717 0.030440
       0.998188 0.141588 0.024101
       1.000000 0.133978 0.017854
       1.000000 0.125852 0.011691
       1.000000 0.117060 0.005434
       1.000000 0.107566 0.000000
       1.000000 0.097091 0.000000];

case { 'D9' }
descriptorname = 'diverging_bwr_55-98_c37_n256';
description = 'Diverging low contrast blue - white - red colour map.  Good in conjunction with relief shading';
map = [0.141243 0.500924 0.998106
       0.162377 0.504255 0.998107
       0.181058 0.507583 0.998107
       0.198013 0.510919 0.998105
       0.213606 0.514278 0.998100
       0.228129 0.517635 0.998093
       0.241742 0.520995 0.998085
       0.254627 0.524368 0.998074
       0.266852 0.527748 0.998061
       0.278562 0.531140 0.998045
       0.289778 0.534536 0.998028
       0.300572 0.537947 0.998008
       0.311013 0.541347 0.997986
       0.321096 0.544773 0.997962
       0.330868 0.548204 0.997936
       0.340387 0.551637 0.997908
       0.349641 0.555073 0.997877
       0.358654 0.558517 0.997844
       0.367464 0.561986 0.997809
       0.376089 0.565436 0.997771
       0.384510 0.568908 0.997731
       0.392771 0.572392 0.997689
       0.400888 0.575880 0.997645
       0.408829 0.579368 0.997598
       0.416646 0.582857 0.997549
       0.424326 0.586366 0.997498
       0.431898 0.589870 0.997444
       0.439328 0.593395 0.997388
       0.446666 0.596920 0.997329
       0.453895 0.600446 0.997268
       0.461025 0.603990 0.997205
       0.468062 0.607533 0.997140
       0.475022 0.611081 0.997071
       0.481881 0.614628 0.997001
       0.488661 0.618196 0.996928
       0.495366 0.621758 0.996853
       0.502001 0.625337 0.996775
       0.508567 0.628920 0.996695
       0.515052 0.632506 0.996612
       0.521478 0.636101 0.996527
       0.527846 0.639699 0.996439
       0.534151 0.643299 0.996349
       0.540405 0.646914 0.996256
       0.546605 0.650528 0.996161
       0.552740 0.654149 0.996063
       0.558825 0.657782 0.995962
       0.564866 0.661412 0.995859
       0.570861 0.665047 0.995754
       0.576818 0.668697 0.995646
       0.582712 0.672348 0.995535
       0.588570 0.676001 0.995422
       0.594394 0.679658 0.995306
       0.600174 0.683329 0.995187
       0.605909 0.687001 0.995066
       0.611618 0.690674 0.994942
       0.617284 0.694356 0.994816
       0.622917 0.698047 0.994687
       0.628517 0.701739 0.994555
       0.634082 0.705443 0.994420
       0.639611 0.709152 0.994283
       0.645106 0.712853 0.994143
       0.650581 0.716575 0.994001
       0.656026 0.720290 0.993856
       0.661436 0.724014 0.993708
       0.666815 0.727744 0.993557
       0.672178 0.731479 0.993403
       0.677513 0.735222 0.993247
       0.682813 0.738970 0.993087
       0.688096 0.742719 0.992924
       0.693352 0.746469 0.992758
       0.698581 0.750228 0.992589
       0.703794 0.753995 0.992417
       0.708986 0.757764 0.992243
       0.714145 0.761536 0.992065
       0.719293 0.765320 0.991885
       0.724415 0.769097 0.991702
       0.729519 0.772886 0.991516
       0.734604 0.776685 0.991327
       0.739671 0.780478 0.991135
       0.744709 0.784286 0.990941
       0.749732 0.788088 0.990743
       0.754750 0.791900 0.990542
       0.759733 0.795715 0.990339
       0.764712 0.799542 0.990132
       0.769665 0.803362 0.989923
       0.774605 0.807195 0.989711
       0.779532 0.811027 0.989495
       0.784446 0.814865 0.989276
       0.789338 0.818706 0.989053
       0.794217 0.822559 0.988826
       0.799081 0.826410 0.988597
       0.803928 0.830267 0.988365
       0.808765 0.834130 0.988130
       0.813582 0.837994 0.987893
       0.818389 0.841863 0.987653
       0.823187 0.845733 0.987410
       0.827967 0.849615 0.987164
       0.832738 0.853492 0.986913
       0.837499 0.857380 0.986658
       0.842244 0.861271 0.986400
       0.846971 0.865165 0.986140
       0.851694 0.869070 0.985876
       0.856404 0.872970 0.985609
       0.861102 0.876873 0.985339
       0.865791 0.880788 0.985066
       0.870464 0.884705 0.984789
       0.875134 0.888626 0.984509
       0.879788 0.892550 0.984225
       0.884437 0.896477 0.983936
       0.889073 0.900407 0.983644
       0.893696 0.904342 0.983350
       0.898312 0.908285 0.983053
       0.902923 0.912226 0.982754
       0.907521 0.916171 0.982451
       0.912106 0.920124 0.982144
       0.916687 0.924081 0.981832
       0.921255 0.928037 0.981516
       0.925820 0.932001 0.981197
       0.930374 0.935972 0.980875
       0.934917 0.939944 0.980550
       0.939456 0.943917 0.980222
       0.943992 0.947901 0.979890
       0.948534 0.951881 0.979532
       0.953113 0.955842 0.979095
       0.957764 0.959670 0.978430
       0.962497 0.963118 0.977259
       0.967204 0.965705 0.975174
       0.971575 0.966854 0.971868
       0.975245 0.966168 0.967291
       0.978010 0.963715 0.961721
       0.979942 0.959958 0.955596
       0.981296 0.955461 0.949236
       0.982343 0.950638 0.942834
       0.983244 0.945708 0.936433
       0.984082 0.940749 0.930039
       0.984882 0.935790 0.923664
       0.985646 0.930832 0.917296
       0.986381 0.925874 0.910930
       0.987086 0.920914 0.904574
       0.987755 0.915958 0.898224
       0.988395 0.911007 0.891887
       0.989008 0.906054 0.885553
       0.989590 0.901097 0.879224
       0.990139 0.896146 0.872908
       0.990659 0.891191 0.866596
       0.991150 0.886242 0.860296
       0.991612 0.881288 0.853995
       0.992045 0.876334 0.847707
       0.992450 0.871386 0.841430
       0.992826 0.866433 0.835156
       0.993173 0.861482 0.828893
       0.993491 0.856531 0.822634
       0.993781 0.851580 0.816385
       0.994043 0.846629 0.810137
       0.994277 0.841683 0.803905
       0.994485 0.836732 0.797675
       0.994666 0.831785 0.791457
       0.994819 0.826833 0.785248
       0.994946 0.821883 0.779041
       0.995047 0.816930 0.772843
       0.995121 0.811979 0.766655
       0.995170 0.807030 0.760477
       0.995192 0.802074 0.754307
       0.995188 0.797121 0.748141
       0.995159 0.792170 0.741981
       0.995105 0.787219 0.735829
       0.995025 0.782265 0.729693
       0.994920 0.777312 0.723555
       0.994791 0.772351 0.717437
       0.994636 0.767392 0.711318
       0.994457 0.762434 0.705204
       0.994254 0.757478 0.699100
       0.994027 0.752514 0.693014
       0.993776 0.747559 0.686924
       0.993502 0.742596 0.680841
       0.993203 0.737627 0.674776
       0.992879 0.732666 0.668718
       0.992531 0.727693 0.662667
       0.992160 0.722721 0.656622
       0.991766 0.717757 0.650581
       0.991349 0.712776 0.644547
       0.990910 0.707803 0.638530
       0.990448 0.702829 0.632519
       0.989964 0.697844 0.626512
       0.989458 0.692870 0.620523
       0.988926 0.687880 0.614524
       0.988372 0.682891 0.608556
       0.987800 0.677904 0.602576
       0.987206 0.672910 0.596616
       0.986588 0.667914 0.590656
       0.985947 0.662913 0.584714
       0.985287 0.657909 0.578773
       0.984605 0.652892 0.572848
       0.983899 0.647882 0.566926
       0.983172 0.642870 0.561005
       0.982429 0.637851 0.555100
       0.981661 0.632828 0.549209
       0.980872 0.627800 0.543318
       0.980064 0.622765 0.537441
       0.979234 0.617725 0.531564
       0.978380 0.612680 0.525701
       0.977513 0.607641 0.519839
       0.976621 0.602581 0.514002
       0.975709 0.597523 0.508165
       0.974778 0.592463 0.502326
       0.973829 0.587380 0.496496
       0.972862 0.582301 0.490697
       0.971870 0.577220 0.484877
       0.970858 0.572129 0.479086
       0.969828 0.567032 0.473287
       0.968780 0.561926 0.467516
       0.967715 0.556807 0.461742
       0.966624 0.551683 0.455974
       0.965516 0.546553 0.450219
       0.964390 0.541400 0.444477
       0.963248 0.536256 0.438722
       0.962082 0.531094 0.432996
       0.960904 0.525926 0.427277
       0.959703 0.520740 0.421545
       0.958486 0.515543 0.415842
       0.957245 0.510343 0.410150
       0.955993 0.505126 0.404443
       0.954717 0.499893 0.398760
       0.953426 0.494664 0.393080
       0.952111 0.489398 0.387407
       0.950787 0.484130 0.381749
       0.949438 0.478852 0.376099
       0.948075 0.473546 0.370439
       0.946693 0.468235 0.364801
       0.945297 0.462898 0.359165
       0.943879 0.457556 0.353539
       0.942447 0.452200 0.347927
       0.940992 0.446807 0.342314
       0.939526 0.441402 0.336699
       0.938040 0.435980 0.331089
       0.936540 0.430531 0.325494
       0.935015 0.425064 0.319923
       0.933477 0.419571 0.314327
       0.931925 0.414048 0.308765
       0.930359 0.408504 0.303170
       0.928768 0.402935 0.297612
       0.927165 0.397333 0.292052
       0.925549 0.391711 0.286478
       0.923914 0.386041 0.280922
       0.922263 0.380360 0.275397
       0.920589 0.374635 0.269850
       0.918905 0.368874 0.264299
       0.917212 0.363064 0.258758
       0.915489 0.357239 0.253217
       0.913763 0.351346 0.247684
       0.912013 0.345416 0.242136
       0.910252 0.339454 0.236613
       0.908472 0.333425 0.231066
       0.906678 0.327333 0.225524
       0.904868 0.321205 0.219989
       0.903044 0.315005 0.214434];

case { 'D10' }
descriptorname = 'diverging_cwm_80-100_c22_n256';
description = 'Diverging low contrast cyan - white - magenta colour map.  Good in conjunction with relief shading';
map = [0.000000 0.850714 1.000000
       0.000000 0.851941 1.000000
       0.056211 0.853159 1.000000
       0.100291 0.854380 1.000000
       0.131679 0.855603 1.000000
       0.157115 0.856818 1.000000
       0.179018 0.858036 1.000000
       0.198457 0.859254 1.000000
       0.216153 0.860472 1.000000
       0.232414 0.861688 1.000000
       0.247584 0.862905 1.000000
       0.261829 0.864119 1.000000
       0.275321 0.865326 1.000000
       0.288134 0.866539 1.000000
       0.300395 0.867753 1.000000
       0.312148 0.868965 1.000000
       0.323487 0.870167 1.000000
       0.334426 0.871381 1.000000
       0.344993 0.872589 1.000000
       0.355262 0.873794 1.000000
       0.365212 0.875001 1.000000
       0.374914 0.876206 1.000000
       0.384368 0.877410 1.000000
       0.393601 0.878614 1.000000
       0.402625 0.879816 1.000000
       0.411458 0.881020 1.000000
       0.420113 0.882220 1.000000
       0.428599 0.883422 1.000000
       0.436930 0.884622 1.000000
       0.445107 0.885821 1.000000
       0.453148 0.887017 1.000000
       0.461056 0.888214 1.000000
       0.468849 0.889410 1.000000
       0.476516 0.890605 1.000000
       0.484071 0.891801 1.000000
       0.491541 0.892993 1.000000
       0.498889 0.894188 1.000000
       0.506141 0.895380 1.000000
       0.513319 0.896571 1.000000
       0.520397 0.897760 1.000000
       0.527405 0.898949 1.000000
       0.534325 0.900138 1.000000
       0.541170 0.901325 1.000000
       0.547952 0.902513 1.000000
       0.554657 0.903698 1.000000
       0.561294 0.904885 1.000000
       0.567871 0.906070 1.000000
       0.574380 0.907253 1.000000
       0.580840 0.908435 1.000000
       0.587236 0.909616 1.000000
       0.593584 0.910798 1.000000
       0.599876 0.911977 1.000000
       0.606109 0.913157 1.000000
       0.612301 0.914336 1.000000
       0.618452 0.915508 1.000000
       0.624547 0.916690 1.000000
       0.630603 0.917867 1.000000
       0.636613 0.919036 1.000000
       0.642578 0.920212 1.000000
       0.648502 0.921386 1.000000
       0.654390 0.922561 1.000000
       0.660242 0.923734 1.000000
       0.666056 0.924899 1.000000
       0.671834 0.926073 1.000000
       0.677584 0.927239 1.000000
       0.683289 0.928407 1.000000
       0.688971 0.929576 1.000000
       0.694606 0.930746 1.000000
       0.700225 0.931908 1.000000
       0.705811 0.933072 1.000000
       0.711369 0.934236 1.000000
       0.716897 0.935402 1.000000
       0.722387 0.936569 1.000000
       0.727862 0.937728 1.000000
       0.733310 0.938888 1.000000
       0.738732 0.940050 1.000000
       0.744124 0.941205 1.000000
       0.749490 0.942368 1.000000
       0.754846 0.943523 1.000000
       0.760166 0.944681 1.000000
       0.765471 0.945834 1.000000
       0.770749 0.946991 1.000000
       0.776004 0.948144 1.000000
       0.781242 0.949296 1.000000
       0.786462 0.950450 1.000000
       0.791652 0.951601 1.000000
       0.796827 0.952753 1.000000
       0.801985 0.953904 1.000000
       0.807128 0.955053 1.000000
       0.812244 0.956202 1.000000
       0.817346 0.957346 1.000000
       0.822435 0.958496 1.000000
       0.827499 0.959639 1.000000
       0.832550 0.960786 1.000000
       0.837588 0.961925 1.000000
       0.842606 0.963072 1.000000
       0.847603 0.964211 1.000000
       0.852597 0.965353 1.000000
       0.857563 0.966495 1.000000
       0.862527 0.967639 1.000000
       0.867465 0.968775 1.000000
       0.872396 0.969912 1.000000
       0.877306 0.971050 1.000000
       0.882208 0.972189 1.000000
       0.887097 0.973324 1.000000
       0.891973 0.974456 1.000000
       0.896836 0.975590 1.000000
       0.901682 0.976724 1.000000
       0.906520 0.977857 1.000000
       0.911344 0.978988 1.000000
       0.916154 0.980122 1.000000
       0.920954 0.981251 1.000000
       0.925747 0.982382 1.000000
       0.930527 0.983506 1.000000
       0.935291 0.984638 1.000000
       0.940051 0.985763 1.000000
       0.944797 0.986890 1.000000
       0.949529 0.988011 1.000000
       0.954256 0.989138 1.000000
       0.958971 0.990260 1.000000
       0.963674 0.991382 1.000000
       0.968372 0.992506 1.000000
       0.973059 0.993626 1.000000
       0.977733 0.994742 1.000000
       0.982404 0.995860 1.000000
       0.987251 0.997024 1.000000
       0.991200 0.997975 1.000000
       0.996849 0.999338 1.000000
       0.999882 0.998202 0.999687
       0.999905 0.995746 0.999256
       0.999926 0.993298 0.998823
       0.999943 0.990841 0.998389
       0.999957 0.988385 0.997955
       0.999968 0.985932 0.997521
       0.999976 0.983475 0.997087
       0.999981 0.981021 0.996652
       0.999983 0.978561 0.996217
       0.999982 0.976105 0.995782
       0.999977 0.973651 0.995347
       0.999970 0.971193 0.994912
       0.999960 0.968737 0.994477
       0.999947 0.966277 0.994041
       0.999930 0.963819 0.993606
       0.999911 0.961361 0.993170
       0.999889 0.958903 0.992731
       0.999863 0.956443 0.992292
       0.999835 0.953984 0.991852
       0.999804 0.951522 0.991413
       0.999770 0.949060 0.990973
       0.999732 0.946598 0.990533
       0.999692 0.944138 0.990093
       0.999649 0.941674 0.989654
       0.999603 0.939212 0.989212
       0.999554 0.936752 0.988769
       0.999502 0.934281 0.988325
       0.999446 0.931818 0.987884
       0.999388 0.929352 0.987443
       0.999327 0.926886 0.987001
       0.999263 0.924421 0.986556
       0.999196 0.921958 0.986111
       0.999126 0.919487 0.985666
       0.999054 0.917023 0.985222
       0.998978 0.914553 0.984777
       0.998899 0.912082 0.984331
       0.998817 0.909613 0.983883
       0.998733 0.907144 0.983434
       0.998645 0.904671 0.982988
       0.998555 0.902199 0.982543
       0.998461 0.899729 0.982096
       0.998365 0.897257 0.981646
       0.998266 0.894783 0.981196
       0.998164 0.892307 0.980747
       0.998059 0.889829 0.980298
       0.997952 0.887352 0.979849
       0.997841 0.884877 0.979398
       0.997728 0.882397 0.978944
       0.997612 0.879919 0.978491
       0.997493 0.877441 0.978041
       0.997372 0.874962 0.977591
       0.997247 0.872483 0.977138
       0.997120 0.869996 0.976684
       0.996990 0.867518 0.976230
       0.996858 0.865032 0.975776
       0.996722 0.862555 0.975322
       0.996584 0.860068 0.974867
       0.996443 0.857578 0.974413
       0.996299 0.855096 0.973958
       0.996152 0.852610 0.973504
       0.996003 0.850119 0.973049
       0.995851 0.847626 0.972594
       0.995696 0.845139 0.972136
       0.995539 0.842650 0.971677
       0.995379 0.840159 0.971218
       0.995216 0.837666 0.970760
       0.995050 0.835169 0.970301
       0.994882 0.832672 0.969842
       0.994711 0.830179 0.969383
       0.994537 0.827678 0.968923
       0.994361 0.825180 0.968465
       0.994182 0.822685 0.968005
       0.994001 0.820180 0.967544
       0.993817 0.817679 0.967081
       0.993630 0.815178 0.966618
       0.993441 0.812673 0.966155
       0.993248 0.810168 0.965692
       0.993053 0.807665 0.965228
       0.992854 0.805155 0.964765
       0.992653 0.802647 0.964301
       0.992449 0.800142 0.963837
       0.992242 0.797626 0.963374
       0.992032 0.795115 0.962910
       0.991821 0.792603 0.962442
       0.991606 0.790088 0.961975
       0.991389 0.787572 0.961510
       0.991170 0.785057 0.961046
       0.990947 0.782536 0.960580
       0.990723 0.780015 0.960111
       0.990495 0.777498 0.959643
       0.990266 0.774969 0.959174
       0.990034 0.772446 0.958707
       0.989799 0.769920 0.958239
       0.989562 0.767392 0.957769
       0.989321 0.764868 0.957296
       0.989077 0.762333 0.956825
       0.988830 0.759800 0.956357
       0.988581 0.757270 0.955888
       0.988329 0.754740 0.955416
       0.988076 0.752198 0.954942
       0.987821 0.749656 0.954470
       0.987564 0.747122 0.953997
       0.987304 0.744578 0.953525
       0.987041 0.742036 0.953052
       0.986774 0.739498 0.952575
       0.986504 0.736944 0.952099
       0.986232 0.734398 0.951626
       0.985958 0.731847 0.951153
       0.985681 0.729293 0.950677
       0.985403 0.726739 0.950200
       0.985122 0.724182 0.949723
       0.984838 0.721621 0.949246
       0.984552 0.719066 0.948769
       0.984262 0.716508 0.948292
       0.983969 0.713939 0.947815
       0.983673 0.711378 0.947338
       0.983375 0.708811 0.946857
       0.983076 0.706239 0.946376
       0.982776 0.703664 0.945898
       0.982473 0.701090 0.945421
       0.982167 0.698510 0.944941
       0.981857 0.695938 0.944460
       0.981544 0.693355 0.943978
       0.981229 0.690766 0.943496
       0.980912 0.688187 0.943016
       0.980592 0.685599 0.942535
       0.980271 0.683007 0.942050
       0.979948 0.680409 0.941565];

case { 'D11' }
descriptorname = 'diverging-isoluminant_cjo_70_c25_n256';
description = 'Diverging isoluminant lightblue - lightgrey - orange colour map';
map = [0.000000 0.715266 1.000000
       0.000000 0.714897 1.000000
       0.000000 0.714529 1.000000
       0.000000 0.714166 1.000000
       0.024434 0.713803 1.000000
       0.068274 0.713439 0.999119
       0.097664 0.713073 0.996395
       0.120619 0.712707 0.993681
       0.139973 0.712345 0.990962
       0.156930 0.711985 0.988243
       0.172105 0.711625 0.985530
       0.185976 0.711263 0.982814
       0.198720 0.710900 0.980100
       0.210636 0.710537 0.977386
       0.221799 0.710175 0.974670
       0.232354 0.709814 0.971961
       0.242343 0.709455 0.969248
       0.251880 0.709096 0.966537
       0.260971 0.708735 0.963827
       0.269705 0.708372 0.961120
       0.278083 0.708009 0.958414
       0.286141 0.707649 0.955707
       0.293942 0.707291 0.952999
       0.301478 0.706935 0.950291
       0.308801 0.706577 0.947587
       0.315882 0.706218 0.944884
       0.322759 0.705858 0.942180
       0.329450 0.705499 0.939476
       0.335947 0.705141 0.936776
       0.342299 0.704783 0.934068
       0.348484 0.704426 0.931370
       0.354529 0.704069 0.928667
       0.360416 0.703714 0.925971
       0.366203 0.703359 0.923274
       0.371835 0.703004 0.920571
       0.377372 0.702646 0.917880
       0.382787 0.702288 0.915179
       0.388093 0.701930 0.912485
       0.393298 0.701576 0.909792
       0.398412 0.701224 0.907098
       0.403430 0.700872 0.904401
       0.408350 0.700518 0.901709
       0.413196 0.700162 0.899018
       0.417945 0.699807 0.896327
       0.422617 0.699453 0.893633
       0.427233 0.699100 0.890944
       0.431764 0.698747 0.888255
       0.436214 0.698395 0.885567
       0.440594 0.698043 0.882875
       0.444926 0.697691 0.880188
       0.449185 0.697340 0.877502
       0.453375 0.696990 0.874816
       0.457510 0.696641 0.872134
       0.461595 0.696292 0.869447
       0.465607 0.695942 0.866760
       0.469592 0.695590 0.864083
       0.473492 0.695238 0.861395
       0.477363 0.694885 0.858712
       0.481173 0.694534 0.856033
       0.484938 0.694184 0.853348
       0.488664 0.693836 0.850668
       0.492334 0.693491 0.847989
       0.495959 0.693146 0.845311
       0.499542 0.692800 0.842635
       0.503085 0.692453 0.839959
       0.506590 0.692103 0.837280
       0.510053 0.691753 0.834603
       0.513478 0.691402 0.831928
       0.516862 0.691052 0.829253
       0.520199 0.690706 0.826576
       0.523517 0.690361 0.823898
       0.526794 0.690018 0.821226
       0.530027 0.689674 0.818549
       0.533228 0.689330 0.815882
       0.536404 0.688985 0.813205
       0.539534 0.688638 0.810533
       0.542649 0.688292 0.807866
       0.545723 0.687946 0.805192
       0.548770 0.687601 0.802523
       0.551781 0.687256 0.799859
       0.554762 0.686912 0.797184
       0.557717 0.686568 0.794519
       0.560642 0.686224 0.791849
       0.563548 0.685882 0.789185
       0.566416 0.685540 0.786520
       0.569255 0.685197 0.783852
       0.572079 0.684851 0.781183
       0.574866 0.684506 0.778517
       0.577634 0.684162 0.775851
       0.580384 0.683822 0.773185
       0.583097 0.683483 0.770524
       0.585800 0.683143 0.767856
       0.588459 0.682801 0.765200
       0.591112 0.682458 0.762531
       0.593738 0.682115 0.759868
       0.596341 0.681774 0.757208
       0.598922 0.681433 0.754551
       0.601478 0.681093 0.751888
       0.604022 0.680753 0.749221
       0.606533 0.680413 0.746564
       0.609035 0.680074 0.743905
       0.611506 0.679736 0.741240
       0.613954 0.679397 0.738586
       0.616398 0.679059 0.735922
       0.618818 0.678722 0.733268
       0.621209 0.678385 0.730609
       0.623592 0.678049 0.727949
       0.625946 0.677713 0.725297
       0.628293 0.677376 0.722632
       0.630620 0.677037 0.719981
       0.632926 0.676696 0.717331
       0.635219 0.676355 0.714670
       0.637503 0.676014 0.712024
       0.639775 0.675671 0.709382
       0.642033 0.675326 0.706741
       0.644281 0.674978 0.704102
       0.646534 0.674626 0.701474
       0.648782 0.674269 0.698859
       0.651033 0.673906 0.696263
       0.653285 0.673534 0.693673
       0.655561 0.673154 0.691105
       0.657852 0.672762 0.688571
       0.660156 0.672353 0.686053
       0.662496 0.671927 0.683566
       0.664855 0.671485 0.681111
       0.667260 0.671029 0.678700
       0.669703 0.670553 0.676328
       0.672180 0.670053 0.673992
       0.674697 0.669529 0.671699
       0.677263 0.668984 0.669453
       0.679852 0.668421 0.667243
       0.682489 0.667837 0.665071
       0.685161 0.667227 0.662948
       0.687858 0.666600 0.660844
       0.690576 0.665964 0.658780
       0.693323 0.665308 0.656752
       0.696083 0.664637 0.654732
       0.698845 0.663963 0.652742
       0.701619 0.663275 0.650774
       0.704400 0.662576 0.648817
       0.707178 0.661871 0.646871
       0.709952 0.661153 0.644927
       0.712713 0.660435 0.643000
       0.715480 0.659718 0.641072
       0.718224 0.658989 0.639155
       0.720961 0.658266 0.637239
       0.723689 0.657540 0.635319
       0.726408 0.656805 0.633409
       0.729110 0.656072 0.631496
       0.731803 0.655328 0.629579
       0.734482 0.654587 0.627671
       0.737146 0.653850 0.625758
       0.739804 0.653103 0.623853
       0.742441 0.652363 0.621931
       0.745066 0.651621 0.620031
       0.747687 0.650871 0.618115
       0.750287 0.650122 0.616205
       0.752875 0.649373 0.614288
       0.755462 0.648616 0.612380
       0.758031 0.647857 0.610474
       0.760589 0.647106 0.608570
       0.763132 0.646344 0.606655
       0.765673 0.645581 0.604746
       0.768190 0.644818 0.602834
       0.770710 0.644053 0.600923
       0.773208 0.643288 0.599017
       0.775703 0.642524 0.597110
       0.778187 0.641750 0.595196
       0.780657 0.640977 0.593291
       0.783118 0.640209 0.591380
       0.785574 0.639431 0.589465
       0.788012 0.638652 0.587558
       0.790444 0.637872 0.585659
       0.792869 0.637092 0.583743
       0.795279 0.636310 0.581834
       0.797682 0.635518 0.579933
       0.800081 0.634734 0.578014
       0.802460 0.633946 0.576120
       0.804838 0.633151 0.574199
       0.807206 0.632356 0.572298
       0.809563 0.631561 0.570383
       0.811911 0.630766 0.568476
       0.814248 0.629961 0.566574
       0.816584 0.629157 0.564660
       0.818900 0.628358 0.562761
       0.821218 0.627548 0.560845
       0.823522 0.626737 0.558935
       0.825820 0.625927 0.557037
       0.828109 0.625114 0.555120
       0.830393 0.624303 0.553210
       0.832664 0.623487 0.551308
       0.834931 0.622662 0.549401
       0.837190 0.621836 0.547488
       0.839439 0.621020 0.545585
       0.841680 0.620197 0.543674
       0.843911 0.619369 0.541760
       0.846137 0.618537 0.539852
       0.848358 0.617697 0.537954
       0.850569 0.616867 0.536038
       0.852778 0.616027 0.534126
       0.854975 0.615183 0.532225
       0.857161 0.614340 0.530314
       0.859346 0.613496 0.528406
       0.861522 0.612650 0.526501
       0.863695 0.611805 0.524585
       0.865853 0.610958 0.522675
       0.868012 0.610098 0.520764
       0.870153 0.609249 0.518862
       0.872301 0.608392 0.516949
       0.874433 0.607531 0.515035
       0.876560 0.606664 0.513125
       0.878684 0.605791 0.511210
       0.880800 0.604929 0.509304
       0.882908 0.604063 0.507392
       0.885014 0.603186 0.505471
       0.887110 0.602308 0.503564
       0.889202 0.601429 0.501661
       0.891285 0.600549 0.499736
       0.893361 0.599670 0.497841
       0.895437 0.598781 0.495913
       0.897504 0.597891 0.494007
       0.899565 0.597006 0.492093
       0.901616 0.596108 0.490178
       0.903666 0.595212 0.488269
       0.905712 0.594314 0.486347
       0.907749 0.593415 0.484428
       0.909779 0.592516 0.482520
       0.911803 0.591605 0.480600
       0.913827 0.590690 0.478686
       0.915835 0.589776 0.476770
       0.917851 0.588860 0.474861
       0.919851 0.587944 0.472930
       0.921853 0.587026 0.471021
       0.923846 0.586109 0.469107
       0.925833 0.585185 0.467186
       0.927812 0.584250 0.465255
       0.929792 0.583317 0.463334
       0.931763 0.582382 0.461423
       0.933729 0.581446 0.459503
       0.935696 0.580509 0.457575
       0.937653 0.579571 0.455656
       0.939606 0.578618 0.453731
       0.941551 0.577666 0.451815
       0.943496 0.576725 0.449887
       0.945435 0.575770 0.447955
       0.947368 0.574801 0.446030
       0.949293 0.573844 0.444116
       0.951219 0.572887 0.442191
       0.953139 0.571915 0.440254
       0.955052 0.570942 0.438328
       0.956959 0.569967 0.436401
       0.958867 0.568992 0.434466
       0.960768 0.568016 0.432545
       0.962662 0.567039 0.430608
       0.964551 0.566048 0.428674
       0.966440 0.565054 0.426741];

case { 'D12' }
descriptorname = 'diverging-isoluminant_cjm_75_c23_n256';
description = 'Diverging isoluminant lightblue - lightgrey - pink colour map';
map = [0.000000 0.786626 1.000000
       0.000000 0.786161 0.999757
       0.000000 0.785695 0.997551
       0.000000 0.785230 0.995348
       0.000000 0.784765 0.993152
       0.000000 0.784301 0.990947
       0.000000 0.783834 0.988747
       0.051263 0.783363 0.986548
       0.088344 0.782895 0.984349
       0.115241 0.782430 0.982150
       0.137228 0.781965 0.979949
       0.156125 0.781496 0.977749
       0.172849 0.781026 0.975549
       0.188052 0.780556 0.973355
       0.201941 0.780087 0.971156
       0.214876 0.779618 0.968960
       0.226996 0.779148 0.966764
       0.238382 0.778678 0.964566
       0.249192 0.778209 0.962372
       0.259479 0.777740 0.960181
       0.269293 0.777271 0.957988
       0.278688 0.776799 0.955795
       0.287708 0.776324 0.953601
       0.296421 0.775850 0.951407
       0.304820 0.775376 0.949213
       0.312941 0.774902 0.947023
       0.320825 0.774428 0.944834
       0.328456 0.773954 0.942644
       0.335875 0.773480 0.940453
       0.343104 0.773006 0.938261
       0.350134 0.772532 0.936074
       0.356995 0.772059 0.933879
       0.363675 0.771585 0.931692
       0.370212 0.771110 0.929504
       0.376620 0.770631 0.927315
       0.382869 0.770152 0.925129
       0.388998 0.769673 0.922946
       0.394995 0.769195 0.920755
       0.400893 0.768716 0.918570
       0.406655 0.768237 0.916386
       0.412329 0.767759 0.914205
       0.417887 0.767280 0.912018
       0.423352 0.766801 0.909836
       0.428735 0.766323 0.907654
       0.434016 0.765845 0.905471
       0.439222 0.765365 0.903287
       0.444356 0.764882 0.901102
       0.449395 0.764398 0.898921
       0.454364 0.763914 0.896742
       0.459257 0.763431 0.894561
       0.464086 0.762948 0.892380
       0.468838 0.762464 0.890197
       0.473519 0.761980 0.888017
       0.478154 0.761497 0.885840
       0.482722 0.761014 0.883662
       0.487236 0.760530 0.881482
       0.491691 0.760041 0.879301
       0.496072 0.759553 0.877122
       0.500418 0.759069 0.874947
       0.504717 0.758585 0.872772
       0.508956 0.758098 0.870592
       0.513142 0.757608 0.868424
       0.517291 0.757119 0.866244
       0.521379 0.756631 0.864072
       0.525434 0.756142 0.861895
       0.529446 0.755653 0.859720
       0.533401 0.755165 0.857542
       0.537335 0.754676 0.855373
       0.541207 0.754184 0.853195
       0.545056 0.753689 0.851021
       0.548865 0.753194 0.848853
       0.552627 0.752701 0.846677
       0.556352 0.752212 0.844509
       0.560055 0.751723 0.842340
       0.563717 0.751231 0.840169
       0.567339 0.750735 0.837997
       0.570924 0.750236 0.835825
       0.574480 0.749737 0.833657
       0.578008 0.749242 0.831489
       0.581507 0.748749 0.829320
       0.584978 0.748256 0.827148
       0.588404 0.747760 0.824977
       0.591820 0.747261 0.822812
       0.595191 0.746761 0.820641
       0.598539 0.746261 0.818472
       0.601861 0.745762 0.816311
       0.605156 0.745263 0.814139
       0.608431 0.744763 0.811975
       0.611667 0.744263 0.809809
       0.614878 0.743764 0.807644
       0.618074 0.743265 0.805475
       0.621240 0.742763 0.803311
       0.624386 0.742258 0.801147
       0.627502 0.741752 0.798985
       0.630602 0.741246 0.796816
       0.633673 0.740744 0.794656
       0.636725 0.740245 0.792492
       0.639751 0.739743 0.790327
       0.642756 0.739238 0.788164
       0.645737 0.738728 0.786006
       0.648705 0.738217 0.783844
       0.651651 0.737710 0.781681
       0.654567 0.737205 0.779515
       0.657479 0.736696 0.777359
       0.660356 0.736184 0.775191
       0.663227 0.735677 0.773030
       0.666067 0.735172 0.770874
       0.668897 0.734662 0.768708
       0.671702 0.734150 0.766554
       0.674492 0.733637 0.764403
       0.677270 0.733125 0.762254
       0.680012 0.732613 0.760119
       0.682749 0.732097 0.757997
       0.685467 0.731576 0.755888
       0.688160 0.731055 0.753804
       0.690827 0.730537 0.751753
       0.693483 0.730014 0.749732
       0.696114 0.729484 0.747778
       0.698710 0.728949 0.745875
       0.701287 0.728411 0.744057
       0.703834 0.727867 0.742334
       0.706350 0.727317 0.740716
       0.708828 0.726761 0.739239
       0.711266 0.726197 0.737890
       0.713660 0.725625 0.736712
       0.716019 0.725042 0.735706
       0.718320 0.724446 0.734892
       0.720580 0.723841 0.734266
       0.722791 0.723228 0.733837
       0.724962 0.722606 0.733608
       0.727073 0.721977 0.733573
       0.729141 0.721339 0.733725
       0.731162 0.720695 0.734053
       0.733148 0.720046 0.734545
       0.735090 0.719389 0.735182
       0.736994 0.718721 0.735944
       0.738872 0.718054 0.736828
       0.740712 0.717386 0.737802
       0.742539 0.716708 0.738865
       0.744337 0.716029 0.739988
       0.746120 0.715344 0.741154
       0.747894 0.714651 0.742376
       0.749643 0.713966 0.743623
       0.751401 0.713275 0.744891
       0.753131 0.712579 0.746180
       0.754872 0.711890 0.747485
       0.756595 0.711196 0.748791
       0.758320 0.710496 0.750106
       0.760034 0.709795 0.751433
       0.761748 0.709096 0.752747
       0.763459 0.708389 0.754078
       0.765173 0.707678 0.755402
       0.766873 0.706974 0.756723
       0.768574 0.706265 0.758049
       0.770277 0.705549 0.759368
       0.771976 0.704834 0.760696
       0.773667 0.704118 0.762016
       0.775361 0.703402 0.763340
       0.777057 0.702681 0.764666
       0.778739 0.701953 0.765990
       0.780423 0.701232 0.767309
       0.782109 0.700507 0.768631
       0.783789 0.699775 0.769956
       0.785465 0.699043 0.771284
       0.787140 0.698311 0.772602
       0.788812 0.697576 0.773923
       0.790478 0.696842 0.775247
       0.792146 0.696106 0.776575
       0.793816 0.695362 0.777897
       0.795472 0.694614 0.779216
       0.797131 0.693868 0.780539
       0.798792 0.693128 0.781865
       0.800448 0.692381 0.783183
       0.802098 0.691623 0.784511
       0.803751 0.690864 0.785831
       0.805396 0.690112 0.787154
       0.807048 0.689359 0.788474
       0.808691 0.688597 0.789796
       0.810328 0.687831 0.791116
       0.811970 0.687065 0.792440
       0.813605 0.686297 0.793766
       0.815242 0.685530 0.795083
       0.816876 0.684754 0.796403
       0.818503 0.683977 0.797725
       0.820132 0.683205 0.799051
       0.821764 0.682423 0.800372
       0.823385 0.681640 0.801690
       0.825006 0.680856 0.803012
       0.826630 0.680071 0.804337
       0.828245 0.679284 0.805654
       0.829864 0.678497 0.806981
       0.831480 0.677709 0.808301
       0.833087 0.676913 0.809623
       0.834699 0.676112 0.810942
       0.836306 0.675311 0.812263
       0.837914 0.674508 0.813583
       0.839519 0.673705 0.814906
       0.841119 0.672901 0.816231
       0.842720 0.672089 0.817548
       0.844316 0.671275 0.818867
       0.845910 0.670467 0.820189
       0.847504 0.669647 0.821515
       0.849101 0.668827 0.822836
       0.850687 0.668007 0.824154
       0.852282 0.667178 0.825475
       0.853861 0.666348 0.826800
       0.855453 0.665522 0.828117
       0.857031 0.664686 0.829444
       0.858612 0.663859 0.830764
       0.860196 0.663022 0.832086
       0.861773 0.662184 0.833405
       0.863350 0.661337 0.834727
       0.864919 0.660488 0.836047
       0.866493 0.659644 0.837371
       0.868068 0.658789 0.838687
       0.869631 0.657944 0.840014
       0.871200 0.657089 0.841332
       0.872766 0.656231 0.842655
       0.874327 0.655364 0.843971
       0.875891 0.654499 0.845294
       0.877449 0.653634 0.846613
       0.879007 0.652762 0.847935
       0.880564 0.651898 0.849260
       0.882120 0.651021 0.850577
       0.883677 0.650144 0.851902
       0.885227 0.649267 0.853220
       0.886778 0.648377 0.854543
       0.888326 0.647494 0.855866
       0.889872 0.646607 0.857184
       0.891417 0.645711 0.858505
       0.892961 0.644816 0.859830
       0.894507 0.643919 0.861148
       0.896045 0.643022 0.862475
       0.897585 0.642121 0.863796
       0.899122 0.641209 0.865111
       0.900656 0.640308 0.866435
       0.902191 0.639394 0.867758
       0.903724 0.638478 0.869081
       0.905259 0.637561 0.870396
       0.906786 0.636645 0.871723
       0.908316 0.635715 0.873042
       0.909842 0.634791 0.874363
       0.911366 0.633863 0.875687
       0.912890 0.632927 0.877004
       0.914414 0.631991 0.878329
       0.915929 0.631055 0.879649
       0.917456 0.630110 0.880972
       0.918967 0.629162 0.882292
       0.920484 0.628218 0.883618
       0.922005 0.627262 0.884938
       0.923519 0.626306 0.886262
       0.925026 0.625349 0.887579
       0.926538 0.624391 0.888905
       0.928046 0.623428 0.890225
       0.929556 0.622452 0.891548
       0.931065 0.621481 0.892870];

case { 'D13'  'BWG' }
descriptorname = 'diverging_bwg_20-95_c41_n256';
description = 'Diverging blue - white - green colour map';
map = [0.065559 0.176330 0.407628
       0.069881 0.181110 0.415946
       0.074129 0.185945 0.424281
       0.078264 0.190742 0.432638
       0.082501 0.195596 0.440988
       0.086610 0.200459 0.449369
       0.090704 0.205386 0.457741
       0.094774 0.210325 0.466124
       0.098735 0.215285 0.474519
       0.102698 0.220308 0.482901
       0.106653 0.225313 0.491303
       0.110513 0.230363 0.499671
       0.114289 0.235479 0.508070
       0.118135 0.240568 0.516439
       0.121825 0.245725 0.524817
       0.125578 0.250925 0.533183
       0.129213 0.256126 0.541538
       0.132849 0.261385 0.549896
       0.136388 0.266664 0.558229
       0.139912 0.271962 0.566567
       0.143422 0.277330 0.574876
       0.146831 0.282717 0.583179
       0.150235 0.288111 0.591471
       0.153535 0.293557 0.599742
       0.156830 0.299052 0.607997
       0.160014 0.304559 0.616226
       0.163224 0.310123 0.624439
       0.166310 0.315707 0.632627
       0.169379 0.321316 0.640789
       0.172324 0.326968 0.648933
       0.175249 0.332682 0.657046
       0.178117 0.338393 0.665123
       0.180847 0.344157 0.673180
       0.183551 0.349975 0.681191
       0.186190 0.355822 0.689186
       0.188725 0.361700 0.697127
       0.191159 0.367610 0.705037
       0.193535 0.373579 0.712901
       0.195792 0.379578 0.720727
       0.197952 0.385615 0.728506
       0.199998 0.391710 0.736234
       0.201955 0.397839 0.743917
       0.203818 0.404008 0.751547
       0.205512 0.410231 0.759108
       0.207104 0.416484 0.766617
       0.208530 0.422783 0.774060
       0.209820 0.429145 0.781444
       0.210960 0.435554 0.788753
       0.211943 0.442007 0.795982
       0.212709 0.448504 0.803137
       0.213299 0.455061 0.810207
       0.213704 0.461664 0.817193
       0.213884 0.468320 0.824083
       0.213827 0.475048 0.830880
       0.213520 0.481815 0.837570
       0.212962 0.488643 0.844144
       0.212179 0.495526 0.850607
       0.211096 0.502467 0.856950
       0.209765 0.509465 0.863170
       0.208208 0.516508 0.869250
       0.206469 0.523607 0.875194
       0.204539 0.530741 0.880995
       0.202505 0.537917 0.886653
       0.200478 0.545110 0.892158
       0.198568 0.552330 0.897511
       0.196895 0.559553 0.902709
       0.195510 0.566782 0.907755
       0.194652 0.573991 0.912639
       0.194378 0.581191 0.917375
       0.194819 0.588362 0.921949
       0.196092 0.595509 0.926366
       0.198261 0.602616 0.930633
       0.201399 0.609682 0.934740
       0.205545 0.616708 0.938697
       0.210649 0.623686 0.942500
       0.216720 0.630615 0.946139
       0.223667 0.637493 0.949630
       0.231495 0.644322 0.952966
       0.240073 0.651113 0.956145
       0.249402 0.657855 0.959164
       0.259402 0.664536 0.962025
       0.269977 0.671182 0.964733
       0.281067 0.677780 0.967286
       0.292670 0.684318 0.969674
       0.304705 0.690818 0.971909
       0.317127 0.697274 0.973979
       0.329909 0.703682 0.975891
       0.343000 0.710041 0.977647
       0.356376 0.716355 0.979240
       0.370005 0.722612 0.980671
       0.383891 0.728831 0.981943
       0.397992 0.735005 0.983048
       0.412293 0.741120 0.983995
       0.426773 0.747200 0.984780
       0.441428 0.753220 0.985399
       0.456238 0.759199 0.985855
       0.471203 0.765135 0.986150
       0.486288 0.771016 0.986282
       0.501509 0.776851 0.986253
       0.516827 0.782633 0.986065
       0.532244 0.788373 0.985720
       0.547735 0.794071 0.985223
       0.563296 0.799722 0.984580
       0.578889 0.805324 0.983791
       0.594504 0.810897 0.982873
       0.610117 0.816437 0.981833
       0.625705 0.821942 0.980677
       0.641239 0.827417 0.979424
       0.656711 0.832873 0.978078
       0.672077 0.838313 0.976660
       0.687334 0.843737 0.975175
       0.702463 0.849159 0.973640
       0.717472 0.854574 0.972059
       0.732311 0.859986 0.970429
       0.746964 0.865384 0.968760
       0.761408 0.870772 0.967043
       0.775602 0.876138 0.965267
       0.789488 0.881463 0.963422
       0.802970 0.886718 0.961474
       0.815940 0.891866 0.959394
       0.828224 0.896859 0.957133
       0.839639 0.901625 0.954644
       0.849937 0.906097 0.951863
       0.858862 0.910179 0.948737
       0.866154 0.913780 0.945213
       0.871555 0.916807 0.941234
       0.874846 0.919178 0.936786
       0.875884 0.920837 0.931838
       0.874584 0.921751 0.926416
       0.870967 0.921899 0.920538
       0.865129 0.921306 0.914259
       0.857249 0.920029 0.907621
       0.847554 0.918135 0.900687
       0.836313 0.915703 0.893525
       0.823790 0.912836 0.886196
       0.810249 0.909611 0.878736
       0.795925 0.906112 0.871198
       0.781011 0.902403 0.863610
       0.765662 0.898541 0.855987
       0.749985 0.894576 0.848349
       0.734085 0.890521 0.840711
       0.718005 0.886413 0.833067
       0.701786 0.882249 0.825424
       0.685466 0.878048 0.817782
       0.669069 0.873808 0.810139
       0.652611 0.869536 0.802487
       0.636120 0.865223 0.794820
       0.619597 0.860871 0.787131
       0.603061 0.856471 0.779404
       0.586543 0.852024 0.771644
       0.570061 0.847516 0.763830
       0.553646 0.842958 0.755965
       0.537326 0.838336 0.748038
       0.521099 0.833656 0.740040
       0.505006 0.828909 0.731965
       0.489058 0.824092 0.723807
       0.473273 0.819212 0.715577
       0.457675 0.814271 0.707251
       0.442273 0.809268 0.698839
       0.427058 0.804195 0.690343
       0.412052 0.799061 0.681755
       0.397263 0.793866 0.673088
       0.382713 0.788606 0.664320
       0.368404 0.783288 0.655469
       0.354334 0.777917 0.646529
       0.340528 0.772484 0.637500
       0.326980 0.766998 0.628387
       0.313741 0.761461 0.619185
       0.300775 0.755873 0.609885
       0.288142 0.750231 0.600508
       0.275868 0.744541 0.591044
       0.263925 0.738807 0.581493
       0.252359 0.733021 0.571860
       0.241209 0.727186 0.562148
       0.230492 0.721309 0.552338
       0.220285 0.715397 0.542447
       0.210542 0.709436 0.532473
       0.201347 0.703431 0.522406
       0.192774 0.697386 0.512264
       0.184818 0.691304 0.502044
       0.177529 0.685190 0.491743
       0.170900 0.679031 0.481348
       0.165000 0.672847 0.470897
       0.159841 0.666615 0.460360
       0.155460 0.660363 0.449783
       0.151748 0.654079 0.439141
       0.148732 0.647763 0.428477
       0.146358 0.641423 0.417790
       0.144553 0.635066 0.407105
       0.143192 0.628689 0.396447
       0.142215 0.622285 0.385841
       0.141535 0.615878 0.375325
       0.141044 0.609461 0.364919
       0.140639 0.603033 0.354659
       0.140251 0.596604 0.344534
       0.139832 0.590168 0.334620
       0.139356 0.583741 0.324866
       0.138793 0.577316 0.315354
       0.138098 0.570894 0.306061
       0.137210 0.564485 0.296971
       0.136137 0.558075 0.288094
       0.135006 0.551687 0.279468
       0.133662 0.545298 0.271035
       0.132189 0.538922 0.262812
       0.130605 0.532557 0.254807
       0.128827 0.526201 0.246965
       0.126973 0.519845 0.239334
       0.125041 0.513519 0.231889
       0.122966 0.507192 0.224569
       0.120836 0.500882 0.217480
       0.118671 0.494584 0.210518
       0.116372 0.488295 0.203730
       0.113973 0.482017 0.197077
       0.111572 0.475753 0.190549
       0.109072 0.469505 0.184182
       0.106551 0.463247 0.177979
       0.103914 0.457025 0.171840
       0.101241 0.450814 0.165851
       0.098563 0.444619 0.159986
       0.095803 0.438422 0.154269
       0.092982 0.432257 0.148640
       0.090206 0.426079 0.143157
       0.087373 0.419938 0.137780
       0.084463 0.413802 0.132468
       0.081522 0.407680 0.127248
       0.078467 0.401572 0.122146
       0.075481 0.395489 0.117173
       0.072443 0.389407 0.112320
       0.069456 0.383351 0.107502
       0.066270 0.377299 0.102797
       0.063173 0.371257 0.098194
       0.060006 0.365246 0.093619
       0.056764 0.359243 0.089201
       0.053428 0.353262 0.084837
       0.050177 0.347301 0.080536
       0.046870 0.341340 0.076306
       0.043388 0.335411 0.072205
       0.040070 0.329491 0.068208
       0.036650 0.323572 0.064214
       0.033173 0.317685 0.060346
       0.029999 0.311801 0.056535
       0.026949 0.305975 0.052698
       0.024019 0.300121 0.049141
       0.021208 0.294297 0.045442
       0.018516 0.288504 0.041997
       0.015942 0.282735 0.038495
       0.013480 0.276965 0.035242
       0.010993 0.271214 0.031988
       0.008814 0.265485 0.028960
       0.006907 0.259783 0.025879
       0.005222 0.254088 0.022744
       0.003752 0.248423 0.019551
       0.002493 0.242739 0.016299
       0.001437 0.237131 0.012975
       0.000581 0.231503 0.009410
       0.000000 0.225900 0.005946];

case { 'C1' }
descriptorname = 'cyclic_mrybm_35-75_c68_n256';
description = 'Cyclic: magenta - red - yellow - blue. Alows four orientations/phase angles to be visulaised.';
map = [0.975609 0.520118 0.970963
       0.980325 0.517215 0.963189
       0.984195 0.513568 0.954482
       0.987253 0.509216 0.944879
       0.989542 0.504200 0.934424
       0.991117 0.498555 0.923199
       0.992044 0.492333 0.911264
       0.992386 0.485608 0.898711
       0.992208 0.478440 0.885626
       0.991576 0.470885 0.872092
       0.990553 0.462981 0.858186
       0.989194 0.454827 0.843992
       0.987544 0.446415 0.829579
       0.985648 0.437830 0.814990
       0.983537 0.429071 0.800291
       0.981245 0.420188 0.785511
       0.978786 0.411201 0.770684
       0.976182 0.402116 0.755835
       0.973444 0.392971 0.740978
       0.970571 0.383765 0.726138
       0.967575 0.374503 0.711311
       0.964440 0.365189 0.696498
       0.961184 0.355864 0.681701
       0.957791 0.346503 0.666936
       0.954249 0.337156 0.652191
       0.950551 0.327802 0.637451
       0.946681 0.318488 0.622713
       0.942637 0.309226 0.607977
       0.938393 0.299997 0.593222
       0.933944 0.290868 0.578444
       0.929287 0.281847 0.563657
       0.924409 0.272939 0.548831
       0.919305 0.264182 0.533960
       0.913979 0.255566 0.519077
       0.908420 0.247117 0.504149
       0.902640 0.238833 0.489176
       0.896646 0.230734 0.474192
       0.890441 0.222811 0.459182
       0.884048 0.215011 0.444155
       0.877467 0.207399 0.429109
       0.870720 0.199872 0.414066
       0.863829 0.192502 0.399025
       0.856787 0.185227 0.384010
       0.849624 0.178009 0.369000
       0.842350 0.170816 0.354007
       0.834978 0.163729 0.339075
       0.827528 0.156645 0.324159
       0.820013 0.149618 0.309330
       0.812451 0.142614 0.294521
       0.804858 0.135667 0.279821
       0.797250 0.128854 0.265183
       0.789662 0.122170 0.250661
       0.782115 0.115783 0.236278
       0.774642 0.109705 0.222009
       0.767296 0.104001 0.207949
       0.760123 0.098945 0.194118
       0.753173 0.094721 0.180491
       0.746518 0.091421 0.167231
       0.740217 0.089258 0.154263
       0.734335 0.088445 0.141736
       0.728940 0.089069 0.129628
       0.724103 0.091195 0.117995
       0.719878 0.094767 0.106894
       0.716312 0.099636 0.096236
       0.713433 0.105801 0.086225
       0.711275 0.113024 0.076774
       0.709830 0.121038 0.068001
       0.709098 0.129894 0.059810
       0.709044 0.139202 0.052084
       0.709630 0.149017 0.045170
       0.710808 0.159138 0.038855
       0.712514 0.169499 0.033292
       0.714696 0.179947 0.028947
       0.717291 0.190504 0.025470
       0.720213 0.201067 0.022733
       0.723417 0.211657 0.020622
       0.726850 0.222137 0.019034
       0.730457 0.232565 0.017876
       0.734193 0.242879 0.017071
       0.738018 0.253114 0.016547
       0.741915 0.263227 0.016249
       0.745847 0.273221 0.016125
       0.749795 0.283111 0.016137
       0.753751 0.292884 0.016250
       0.757692 0.302570 0.016440
       0.761605 0.312172 0.016684
       0.765495 0.321712 0.016989
       0.769322 0.331165 0.017296
       0.773097 0.340580 0.017600
       0.776812 0.349937 0.017896
       0.780442 0.359250 0.018177
       0.784000 0.368555 0.018435
       0.787459 0.377827 0.018665
       0.790825 0.387090 0.018861
       0.794094 0.396362 0.019014
       0.797244 0.405638 0.019122
       0.800294 0.414903 0.019179
       0.803223 0.424179 0.019183
       0.806046 0.433460 0.019132
       0.808758 0.442759 0.019026
       0.811360 0.452053 0.018866
       0.813856 0.461340 0.018654
       0.816265 0.470631 0.018394
       0.818571 0.479917 0.018090
       0.820801 0.489183 0.017747
       0.822954 0.498459 0.017369
       0.825032 0.507709 0.016963
       0.827042 0.516942 0.016575
       0.828978 0.526150 0.016225
       0.830837 0.535317 0.015949
       0.832616 0.544456 0.015792
       0.834313 0.553536 0.015820
       0.835903 0.562570 0.016118
       0.837376 0.571505 0.016798
       0.838696 0.580351 0.018004
       0.839847 0.589056 0.019918
       0.840771 0.597613 0.022763
       0.841431 0.605951 0.026810
       0.841773 0.614040 0.032376
       0.841735 0.621817 0.040011
       0.841252 0.629224 0.049120
       0.840261 0.636189 0.059305
       0.838684 0.642636 0.070471
       0.836472 0.648502 0.082640
       0.833559 0.653724 0.095596
       0.829894 0.658244 0.109262
       0.825436 0.662013 0.123432
       0.820169 0.664981 0.138185
       0.814079 0.667153 0.153209
       0.807174 0.668516 0.168557
       0.799467 0.669078 0.184047
       0.790984 0.668874 0.199610
       0.781782 0.667950 0.215201
       0.771898 0.666350 0.230722
       0.761389 0.664160 0.246105
       0.750316 0.661435 0.261333
       0.738731 0.658248 0.276357
       0.726679 0.654667 0.291131
       0.714212 0.650774 0.305673
       0.701369 0.646617 0.319948
       0.688182 0.642255 0.333971
       0.674664 0.637733 0.347744
       0.660836 0.633096 0.361267
       0.646712 0.628377 0.374577
       0.632276 0.623598 0.387655
       0.617525 0.618782 0.400558
       0.602450 0.613930 0.413255
       0.587033 0.609090 0.425781
       0.571260 0.604231 0.438168
       0.555110 0.599363 0.450404
       0.538576 0.594493 0.462504
       0.521599 0.589621 0.474508
       0.504201 0.584753 0.486396
       0.486316 0.579863 0.498214
       0.467958 0.574931 0.509934
       0.449120 0.569964 0.521598
       0.429782 0.564934 0.533218
       0.409982 0.559832 0.544799
       0.389720 0.554613 0.556346
       0.369082 0.549264 0.567890
       0.348105 0.543747 0.579434
       0.326877 0.538047 0.590979
       0.305561 0.532112 0.602556
       0.284236 0.525924 0.614165
       0.263159 0.519446 0.625830
       0.242495 0.512650 0.637551
       0.222572 0.505507 0.649339
       0.203679 0.498022 0.661190
       0.186189 0.490122 0.673126
       0.170514 0.481834 0.685136
       0.157117 0.473118 0.697232
       0.146378 0.464018 0.709410
       0.138585 0.454497 0.721643
       0.133804 0.444580 0.733949
       0.131904 0.434263 0.746297
       0.132531 0.423594 0.758686
       0.135191 0.412602 0.771094
       0.139304 0.401284 0.783498
       0.144495 0.389694 0.795878
       0.150242 0.377897 0.808211
       0.156303 0.365938 0.820451
       0.162459 0.353867 0.832570
       0.168638 0.341794 0.844516
       0.174839 0.329789 0.856233
       0.181049 0.317937 0.867668
       0.187461 0.306394 0.878752
       0.194078 0.295237 0.889426
       0.201013 0.284653 0.899619
       0.208478 0.274783 0.909260
       0.216496 0.265785 0.918299
       0.225063 0.257783 0.926677
       0.234299 0.250981 0.934357
       0.244088 0.245443 0.941312
       0.254476 0.241339 0.947531
       0.265294 0.238660 0.953012
       0.276520 0.237496 0.957776
       0.287984 0.237734 0.961852
       0.299643 0.239327 0.965296
       0.311348 0.242176 0.968161
       0.323045 0.246128 0.970496
       0.334642 0.251060 0.972381
       0.346059 0.256764 0.973864
       0.357329 0.263129 0.975020
       0.368354 0.269998 0.975904
       0.379155 0.277244 0.976570
       0.389731 0.284764 0.977064
       0.400095 0.292503 0.977423
       0.410262 0.300363 0.977682
       0.420229 0.308302 0.977868
       0.430058 0.316272 0.978004
       0.439768 0.324207 0.978110
       0.449401 0.332129 0.978200
       0.458996 0.339971 0.978287
       0.468590 0.347722 0.978377
       0.478245 0.355355 0.978492
       0.488019 0.362824 0.978640
       0.497946 0.370147 0.978828
       0.508073 0.377305 0.979061
       0.518451 0.384258 0.979347
       0.529120 0.391021 0.979691
       0.540096 0.397551 0.980097
       0.551427 0.403865 0.980569
       0.563108 0.409948 0.981113
       0.575134 0.415789 0.981731
       0.587510 0.421405 0.982418
       0.600220 0.426814 0.983172
       0.613223 0.432013 0.983998
       0.626503 0.437003 0.984886
       0.640016 0.441823 0.985827
       0.653712 0.446473 0.986823
       0.667569 0.450995 0.987859
       0.681527 0.455396 0.988937
       0.695577 0.459695 0.990045
       0.709665 0.463920 0.991178
       0.723765 0.468069 0.992333
       0.737851 0.472183 0.993503
       0.751904 0.476241 0.994667
       0.765890 0.480264 0.995824
       0.779793 0.484245 0.996963
       0.793606 0.488206 0.998069
       0.807288 0.492107 0.999125
       0.820827 0.495947 1.000000
       0.834201 0.499717 1.000000
       0.847361 0.503390 1.000000
       0.860287 0.506933 1.000000
       0.872915 0.510299 1.000000
       0.885202 0.513454 1.000000
       0.897090 0.516325 1.000000
       0.908507 0.518885 1.000000
       0.919391 0.521035 1.000000
       0.929679 0.522735 0.998520
       0.939301 0.523911 0.995999
       0.948193 0.524505 0.992736
       0.956308 0.524462 0.988644
       0.963596 0.523740 0.983677
       0.970035 0.522293 0.977792];

case { 'C2'  'PHASE4' }
descriptorname = 'cyclic_mygbm_30-95_c78_n256';
description = 'Cyclic: magenta - yellow - green - blue. Alows four orientations/phase angles to be visulaised.  Really good!';
map = [0.937688 0.333524 0.948091
       0.943832 0.342828 0.942393
       0.949390 0.352750 0.936128
       0.954392 0.363232 0.929311
       0.958860 0.374216 0.921979
       0.962829 0.385575 0.914155
       0.966340 0.397272 0.905883
       0.969436 0.409209 0.897208
       0.972159 0.421299 0.888172
       0.974539 0.433497 0.878829
       0.976626 0.445734 0.869226
       0.978446 0.457972 0.859397
       0.980040 0.470168 0.849387
       0.981421 0.482281 0.839222
       0.982617 0.494300 0.828938
       0.983645 0.506196 0.818543
       0.984527 0.517989 0.808076
       0.985266 0.529642 0.797533
       0.985878 0.541157 0.786940
       0.986372 0.552554 0.776288
       0.986756 0.563819 0.765594
       0.987036 0.574940 0.754857
       0.987225 0.585954 0.744073
       0.987329 0.596830 0.733261
       0.987359 0.607585 0.722406
       0.987323 0.618208 0.711527
       0.987233 0.628715 0.700603
       0.987102 0.639094 0.689645
       0.986942 0.649357 0.678645
       0.986769 0.659488 0.667616
       0.986595 0.669503 0.656545
       0.986436 0.679384 0.645426
       0.986305 0.689155 0.634284
       0.986211 0.698787 0.623088
       0.986165 0.708312 0.611844
       0.986172 0.717716 0.600553
       0.986236 0.727001 0.589203
       0.986356 0.736180 0.577794
       0.986531 0.745264 0.566320
       0.986753 0.754257 0.554758
       0.987015 0.763154 0.543105
       0.987306 0.771982 0.531342
       0.987613 0.780738 0.519460
       0.987915 0.789439 0.507436
       0.988199 0.798073 0.495260
       0.988446 0.806659 0.482912
       0.988635 0.815182 0.470388
       0.988746 0.823650 0.457646
       0.988752 0.832060 0.444700
       0.988628 0.840393 0.431506
       0.988343 0.848634 0.418045
       0.987865 0.856767 0.404327
       0.987155 0.864764 0.390333
       0.986163 0.872591 0.376069
       0.984851 0.880197 0.361516
       0.983160 0.887543 0.346700
       0.981048 0.894578 0.331631
       0.978455 0.901226 0.316372
       0.975339 0.907443 0.300918
       0.971656 0.913155 0.285373
       0.967365 0.918306 0.269826
       0.962437 0.922845 0.254315
       0.956863 0.926719 0.238952
       0.950639 0.929902 0.223876
       0.943771 0.932362 0.209203
       0.936285 0.934099 0.195058
       0.928203 0.935119 0.181532
       0.919585 0.935443 0.168836
       0.910473 0.935102 0.156970
       0.900913 0.934145 0.146103
       0.890971 0.932627 0.136225
       0.880697 0.930612 0.127473
       0.870144 0.928148 0.119769
       0.859367 0.925314 0.113150
       0.848400 0.922162 0.107415
       0.837288 0.918740 0.102569
       0.826051 0.915112 0.098502
       0.814723 0.911316 0.095091
       0.803324 0.907388 0.092162
       0.791866 0.903355 0.089659
       0.780359 0.899246 0.087518
       0.768811 0.895079 0.085510
       0.757233 0.890864 0.083837
       0.745619 0.886624 0.082243
       0.733976 0.882352 0.080673
       0.722295 0.878067 0.079194
       0.710601 0.873765 0.077792
       0.698863 0.869457 0.076415
       0.687099 0.865139 0.075063
       0.675294 0.860818 0.073757
       0.663455 0.856488 0.072319
       0.651565 0.852155 0.071005
       0.639628 0.847804 0.069678
       0.627641 0.843452 0.068313
       0.615598 0.839095 0.066946
       0.603505 0.834729 0.065602
       0.591341 0.830355 0.064284
       0.579114 0.825970 0.063016
       0.566812 0.821583 0.061599
       0.554426 0.817184 0.060374
       0.541951 0.812777 0.059088
       0.529396 0.808369 0.057695
       0.516724 0.803948 0.056522
       0.503949 0.799524 0.055189
       0.491057 0.795083 0.053903
       0.478012 0.790640 0.052644
       0.464840 0.786193 0.051424
       0.451514 0.781731 0.050257
       0.438032 0.777261 0.049220
       0.424375 0.772770 0.048120
       0.410564 0.768269 0.047322
       0.396557 0.763754 0.046740
       0.382394 0.759216 0.046427
       0.368078 0.754661 0.046596
       0.353614 0.750060 0.047299
       0.339062 0.745430 0.048740
       0.324426 0.740751 0.050897
       0.309837 0.736016 0.054069
       0.295322 0.731217 0.058336
       0.281049 0.726343 0.063783
       0.267165 0.721371 0.070322
       0.253870 0.716308 0.077992
       0.241338 0.711121 0.086870
       0.229810 0.705805 0.096608
       0.219614 0.700353 0.107413
       0.210918 0.694751 0.118989
       0.203999 0.689009 0.131291
       0.198943 0.683096 0.144224
       0.195928 0.677034 0.157682
       0.194864 0.670809 0.171614
       0.195597 0.664426 0.185943
       0.197953 0.657910 0.200497
       0.201632 0.651248 0.215333
       0.206391 0.644457 0.230304
       0.211828 0.637562 0.245381
       0.217714 0.630567 0.260517
       0.223814 0.623480 0.275653
       0.229923 0.616316 0.290713
       0.235925 0.609097 0.305738
       0.241618 0.601816 0.320644
       0.246935 0.594504 0.335432
       0.251836 0.587158 0.350083
       0.256216 0.579800 0.364592
       0.260111 0.572415 0.378970
       0.263461 0.565010 0.393203
       0.266243 0.557617 0.407304
       0.268465 0.550209 0.421268
       0.270134 0.542804 0.435128
       0.271218 0.535387 0.448859
       0.271734 0.527982 0.462471
       0.271701 0.520573 0.476000
       0.271122 0.513171 0.489416
       0.269969 0.505756 0.502754
       0.268234 0.498366 0.516003
       0.265977 0.490956 0.529190
       0.263162 0.483520 0.542278
       0.259815 0.476095 0.555286
       0.255941 0.468642 0.568216
       0.251617 0.461165 0.581055
       0.246802 0.453652 0.593797
       0.241605 0.446094 0.606425
       0.236047 0.438486 0.618943
       0.230168 0.430809 0.631314
       0.224104 0.423032 0.643534
       0.217927 0.415169 0.655603
       0.211702 0.407184 0.667494
       0.205504 0.399058 0.679192
       0.199449 0.390795 0.690701
       0.193673 0.382343 0.702012
       0.188179 0.373723 0.713124
       0.182998 0.364881 0.724042
       0.178289 0.355845 0.734770
       0.173922 0.346566 0.745302
       0.169985 0.337070 0.755664
       0.166387 0.327320 0.765861
       0.163120 0.317346 0.775902
       0.160049 0.307119 0.785807
       0.157238 0.296666 0.795575
       0.154570 0.285952 0.805220
       0.152024 0.275051 0.814744
       0.149664 0.263945 0.824140
       0.147444 0.252636 0.833404
       0.145543 0.241182 0.842515
       0.144017 0.229604 0.851446
       0.143121 0.217998 0.860186
       0.143053 0.206386 0.868690
       0.144045 0.194808 0.876912
       0.146305 0.183361 0.884837
       0.150065 0.172187 0.892412
       0.155365 0.161400 0.899600
       0.162302 0.151032 0.906369
       0.170751 0.141359 0.912696
       0.180618 0.132437 0.918560
       0.191735 0.124462 0.923961
       0.203887 0.117652 0.928885
       0.216808 0.112135 0.933354
       0.230285 0.107942 0.937390
       0.244173 0.105248 0.941008
       0.258283 0.104034 0.944254
       0.272456 0.104173 0.947160
       0.286585 0.105597 0.949767
       0.300591 0.108073 0.952118
       0.314397 0.111532 0.954258
       0.327961 0.115651 0.956218
       0.341257 0.120310 0.958033
       0.354259 0.125435 0.959731
       0.366979 0.130836 0.961339
       0.379418 0.136365 0.962880
       0.391595 0.142096 0.964362
       0.403518 0.147862 0.965807
       0.415202 0.153628 0.967221
       0.426686 0.159414 0.968604
       0.437993 0.165152 0.969962
       0.449134 0.170885 0.971293
       0.460135 0.176498 0.972606
       0.471061 0.182036 0.973886
       0.481915 0.187511 0.975137
       0.492732 0.192804 0.976352
       0.503569 0.197982 0.977523
       0.514456 0.203022 0.978633
       0.525406 0.207899 0.979689
       0.536474 0.212587 0.980667
       0.547669 0.217088 0.981567
       0.559028 0.221368 0.982381
       0.570568 0.225422 0.983096
       0.582302 0.229249 0.983716
       0.594235 0.232843 0.984237
       0.606358 0.236227 0.984653
       0.618678 0.239354 0.984965
       0.631160 0.242265 0.985178
       0.643789 0.244961 0.985296
       0.656562 0.247499 0.985325
       0.669430 0.249819 0.985273
       0.682370 0.252030 0.985147
       0.695371 0.254083 0.984956
       0.708386 0.256007 0.984709
       0.721386 0.257863 0.984407
       0.734366 0.259680 0.984054
       0.747284 0.261432 0.983653
       0.760121 0.263191 0.983207
       0.772859 0.264941 0.982712
       0.785479 0.266761 0.982159
       0.797943 0.268669 0.981533
       0.810234 0.270687 0.980822
       0.822330 0.272847 0.980006
       0.834182 0.275246 0.979052
       0.845756 0.277908 0.977932
       0.857023 0.280889 0.976608
       0.867940 0.284304 0.975035
       0.878448 0.288192 0.973174
       0.888519 0.292629 0.970965
       0.898101 0.297682 0.968371
       0.907167 0.303395 0.965334
       0.915672 0.309845 0.961820
       0.923611 0.316999 0.957797
       0.930949 0.324889 0.953227];

case { 'C3' }
descriptorname = 'cyclic_rwbkr_15-85_c43_n256';
description = 'Cyclic: red - white - blue - black - red.  Allows four orientations/phase angles to be visulaised.';
map = [0.865005 0.233578 0.147089
       0.870855 0.240570 0.152427
       0.876134 0.248328 0.158474
       0.880862 0.256692 0.165078
       0.885070 0.265654 0.172323
       0.888797 0.275091 0.180097
       0.892080 0.284910 0.188385
       0.894967 0.295046 0.197070
       0.897499 0.305419 0.206140
       0.899722 0.315925 0.215496
       0.901674 0.326497 0.225125
       0.903398 0.337139 0.235015
       0.904920 0.347762 0.245012
       0.906270 0.358339 0.255213
       0.907471 0.368870 0.265482
       0.908539 0.379309 0.275868
       0.909492 0.389664 0.286275
       0.910344 0.399927 0.296787
       0.911093 0.410107 0.307307
       0.911754 0.420170 0.317885
       0.912327 0.430149 0.328488
       0.912816 0.440036 0.339123
       0.913226 0.449836 0.349769
       0.913557 0.459552 0.360429
       0.913805 0.469192 0.371124
       0.913970 0.478744 0.381845
       0.914051 0.488242 0.392582
       0.914047 0.497673 0.403352
       0.913956 0.507031 0.414133
       0.913777 0.516338 0.424950
       0.913507 0.525604 0.435788
       0.913146 0.534812 0.446646
       0.912692 0.543976 0.457532
       0.912145 0.553091 0.468449
       0.911502 0.562183 0.479398
       0.910763 0.571214 0.490368
       0.909924 0.580226 0.501367
       0.908980 0.589188 0.512385
       0.907941 0.598131 0.523444
       0.906791 0.607051 0.534521
       0.905538 0.615928 0.545633
       0.904170 0.624785 0.556768
       0.902700 0.633621 0.567931
       0.901104 0.642427 0.579139
       0.899392 0.651202 0.590365
       0.897551 0.659940 0.601628
       0.895575 0.668638 0.612915
       0.893457 0.677285 0.624240
       0.891193 0.685861 0.635576
       0.888770 0.694350 0.646941
       0.886170 0.702747 0.658315
       0.883379 0.711002 0.669698
       0.880382 0.719079 0.681064
       0.877162 0.726938 0.692426
       0.873704 0.734529 0.703725
       0.869980 0.741780 0.714956
       0.865987 0.748638 0.726084
       0.861698 0.755027 0.737061
       0.857103 0.760865 0.747852
       0.852208 0.766088 0.758399
       0.846992 0.770624 0.768649
       0.841483 0.774408 0.778566
       0.835679 0.777404 0.788086
       0.829606 0.779550 0.797168
       0.823277 0.780844 0.805775
       0.816732 0.781277 0.813877
       0.809988 0.780858 0.821461
       0.803084 0.779621 0.828513
       0.796046 0.777616 0.835044
       0.788904 0.774884 0.841073
       0.781672 0.771517 0.846621
       0.774368 0.767562 0.851735
       0.767018 0.763115 0.856450
       0.759622 0.758246 0.860812
       0.752194 0.753013 0.864867
       0.744726 0.747508 0.868670
       0.737223 0.741761 0.872247
       0.729681 0.735842 0.875647
       0.722086 0.729797 0.878901
       0.714448 0.723644 0.882045
       0.706761 0.717436 0.885098
       0.698997 0.711173 0.888080
       0.691171 0.704880 0.891008
       0.683277 0.698573 0.893895
       0.675297 0.692273 0.896750
       0.667235 0.685959 0.899574
       0.659087 0.679654 0.902370
       0.650848 0.673374 0.905157
       0.642505 0.667101 0.907929
       0.634058 0.660845 0.910687
       0.625490 0.654607 0.913435
       0.616816 0.648386 0.916167
       0.608017 0.642184 0.918890
       0.599085 0.635996 0.921607
       0.590015 0.629820 0.924307
       0.580813 0.623672 0.926996
       0.571456 0.617527 0.929677
       0.561950 0.611411 0.932344
       0.552267 0.605304 0.935004
       0.542406 0.599226 0.937655
       0.532359 0.593161 0.940294
       0.522097 0.587108 0.942922
       0.511636 0.581082 0.945538
       0.500946 0.575073 0.948145
       0.489998 0.569084 0.950745
       0.478791 0.563125 0.953335
       0.467299 0.557180 0.955914
       0.455491 0.551248 0.958477
       0.443358 0.545344 0.961014
       0.430869 0.539457 0.963520
       0.417993 0.533597 0.965985
       0.404710 0.527768 0.968399
       0.391007 0.521956 0.970733
       0.376830 0.516176 0.972973
       0.362160 0.510430 0.975074
       0.346989 0.504717 0.977011
       0.331274 0.499024 0.978726
       0.315055 0.493367 0.980171
       0.298309 0.487764 0.981270
       0.281070 0.482185 0.981958
       0.263412 0.476648 0.982152
       0.245328 0.471163 0.981768
       0.227020 0.465708 0.980723
       0.208557 0.460312 0.978938
       0.190164 0.454977 0.976338
       0.172092 0.449673 0.972866
       0.154666 0.444427 0.968461
       0.138317 0.439211 0.963097
       0.123443 0.434051 0.956759
       0.110797 0.428947 0.949457
       0.100790 0.423873 0.941217
       0.094074 0.418847 0.932084
       0.090801 0.413860 0.922124
       0.090856 0.408907 0.911400
       0.093726 0.403989 0.900003
       0.098788 0.399093 0.888013
       0.105260 0.394237 0.875526
       0.112642 0.389407 0.862623
       0.120287 0.384593 0.849379
       0.128047 0.379811 0.835869
       0.135554 0.375042 0.822165
       0.142782 0.370293 0.808309
       0.149595 0.365569 0.794358
       0.155939 0.360846 0.780338
       0.161827 0.356167 0.766291
       0.167238 0.351478 0.752231
       0.172187 0.346818 0.738178
       0.176731 0.342166 0.724150
       0.180866 0.337526 0.710152
       0.184645 0.332913 0.696191
       0.188062 0.328292 0.682268
       0.191080 0.323694 0.668408
       0.193837 0.319117 0.654586
       0.196271 0.314541 0.640825
       0.198398 0.310007 0.627114
       0.200276 0.305469 0.613451
       0.201920 0.300926 0.599853
       0.203336 0.296429 0.586302
       0.204520 0.291938 0.572809
       0.205457 0.287445 0.559363
       0.206228 0.283001 0.545983
       0.206772 0.278549 0.532652
       0.207118 0.274114 0.519378
       0.207283 0.269706 0.506156
       0.207273 0.265289 0.492999
       0.207093 0.260909 0.479915
       0.206750 0.256535 0.466877
       0.206237 0.252193 0.453896
       0.205541 0.247863 0.440976
       0.204721 0.243517 0.428133
       0.203758 0.239225 0.415330
       0.202639 0.234958 0.402594
       0.201414 0.230669 0.389929
       0.200091 0.226446 0.377349
       0.198692 0.222216 0.364833
       0.197241 0.218044 0.352411
       0.195718 0.213904 0.340089
       0.194226 0.209803 0.327859
       0.192713 0.205781 0.315778
       0.191294 0.201796 0.303814
       0.190015 0.197936 0.292051
       0.188925 0.194210 0.280470
       0.188096 0.190558 0.269181
       0.187558 0.187114 0.258128
       0.187397 0.183833 0.247458
       0.187700 0.180780 0.237178
       0.188495 0.178030 0.227312
       0.189857 0.175483 0.217960
       0.191857 0.173281 0.209151
       0.194556 0.171408 0.200926
       0.197894 0.169897 0.193375
       0.201969 0.168718 0.186452
       0.206807 0.167880 0.180194
       0.212316 0.167425 0.174643
       0.218523 0.167291 0.169736
       0.225390 0.167458 0.165417
       0.232876 0.167899 0.161800
       0.240938 0.168626 0.158679
       0.249511 0.169577 0.156080
       0.258544 0.170674 0.153891
       0.267949 0.171938 0.152129
       0.277717 0.173330 0.150716
       0.287730 0.174829 0.149595
       0.297982 0.176332 0.148635
       0.308426 0.177961 0.147921
       0.318999 0.179525 0.147325
       0.329695 0.181108 0.146811
       0.340457 0.182692 0.146397
       0.351291 0.184266 0.146052
       0.362175 0.185821 0.145738
       0.373089 0.187303 0.145432
       0.384045 0.188745 0.145125
       0.395023 0.190135 0.144814
       0.406028 0.191486 0.144508
       0.417039 0.192794 0.144189
       0.428063 0.194064 0.143857
       0.439096 0.195235 0.143510
       0.450162 0.196393 0.143145
       0.461239 0.197459 0.142763
       0.472338 0.198471 0.142364
       0.483446 0.199432 0.141951
       0.494593 0.200341 0.141527
       0.505737 0.201195 0.141086
       0.516929 0.201993 0.140606
       0.528127 0.202732 0.140086
       0.539349 0.203414 0.139563
       0.550602 0.204038 0.139052
       0.561878 0.204577 0.138533
       0.573172 0.205042 0.137985
       0.584489 0.205457 0.137391
       0.595832 0.205818 0.136755
       0.607209 0.206109 0.136104
       0.618602 0.206324 0.135477
       0.630017 0.206465 0.134841
       0.641457 0.206534 0.134138
       0.652919 0.206529 0.133411
       0.664400 0.206457 0.132679
       0.675890 0.206320 0.131935
       0.687382 0.206119 0.131186
       0.698863 0.205862 0.130455
       0.710334 0.205563 0.129692
       0.721761 0.205247 0.128936
       0.733142 0.204943 0.128287
       0.744434 0.204676 0.127667
       0.755617 0.204479 0.127130
       0.766647 0.204403 0.126762
       0.777494 0.204511 0.126577
       0.788095 0.204870 0.126609
       0.798415 0.205584 0.126916
       0.808398 0.206754 0.127610
       0.817988 0.208409 0.128694
       0.827146 0.210701 0.130312
       0.835815 0.213691 0.132391
       0.843960 0.217449 0.135098
       0.851553 0.221996 0.138432
       0.858572 0.227395 0.142421];

case { 'C4'  'PHASE2' }
descriptorname = 'cyclic_wrwbw_40-90_c42_n256';
description = 'Cyclic: white - red - white - blue. Good if you just want to visualise +ve and -ve phase';
map = [0.872996 0.835925 0.848636
       0.877699 0.833901 0.840706
       0.881937 0.830747 0.831998
       0.885721 0.826513 0.822535
       0.889064 0.821259 0.812361
       0.891986 0.815066 0.801538
       0.894512 0.808025 0.790125
       0.896664 0.800232 0.778192
       0.898468 0.791781 0.765812
       0.899963 0.782777 0.753046
       0.901163 0.773309 0.739986
       0.902101 0.763466 0.726668
       0.902799 0.753318 0.713164
       0.903274 0.742941 0.699522
       0.903546 0.732373 0.685788
       0.903632 0.721658 0.671985
       0.903546 0.710853 0.658151
       0.903297 0.699962 0.644295
       0.902893 0.689025 0.630457
       0.902340 0.678038 0.616625
       0.901646 0.667017 0.602817
       0.900817 0.655984 0.589038
       0.899861 0.644920 0.575308
       0.898773 0.633862 0.561618
       0.897565 0.622779 0.547972
       0.896231 0.611687 0.534367
       0.894778 0.600578 0.520811
       0.893199 0.589450 0.507301
       0.891508 0.578308 0.493835
       0.889699 0.567152 0.480413
       0.887774 0.555951 0.467042
       0.885741 0.544743 0.453703
       0.883594 0.533492 0.440415
       0.881335 0.522213 0.427182
       0.878966 0.510901 0.413975
       0.876492 0.499543 0.400832
       0.873914 0.488157 0.387699
       0.871233 0.476704 0.374636
       0.868451 0.465197 0.361603
       0.865557 0.453639 0.348608
       0.862575 0.442023 0.335656
       0.859484 0.430317 0.322739
       0.856299 0.418531 0.309852
       0.853017 0.406664 0.296986
       0.849646 0.394701 0.284149
       0.846179 0.382650 0.271372
       0.842639 0.370502 0.258650
       0.839013 0.358273 0.245967
       0.835322 0.345942 0.233361
       0.831575 0.333560 0.220891
       0.827774 0.321100 0.208479
       0.823953 0.308621 0.196258
       0.820128 0.296143 0.184203
       0.816336 0.283748 0.172419
       0.812593 0.271498 0.160987
       0.808961 0.259528 0.149965
       0.805465 0.247916 0.139396
       0.802174 0.236832 0.129550
       0.799139 0.226445 0.120401
       0.796404 0.216970 0.112235
       0.794045 0.208617 0.105036
       0.792093 0.201620 0.099146
       0.790606 0.196252 0.094653
       0.789622 0.192610 0.091622
       0.789161 0.190892 0.090163
       0.789234 0.191163 0.090390
       0.789838 0.193419 0.092261
       0.790962 0.197538 0.095704
       0.792579 0.203391 0.100618
       0.794645 0.210759 0.106933
       0.797111 0.219434 0.114310
       0.799935 0.229166 0.122776
       0.803042 0.239754 0.132158
       0.806397 0.251005 0.142219
       0.809927 0.262728 0.152861
       0.813593 0.274787 0.164057
       0.817351 0.287069 0.175562
       0.821159 0.299505 0.187446
       0.824982 0.311958 0.199498
       0.828802 0.324443 0.211814
       0.832583 0.336893 0.224203
       0.836319 0.349271 0.236771
       0.839997 0.361569 0.249367
       0.843593 0.373780 0.262059
       0.847116 0.385885 0.274795
       0.850555 0.397923 0.287580
       0.853904 0.409864 0.300420
       0.857161 0.421695 0.313295
       0.860323 0.433454 0.326179
       0.863385 0.445140 0.339129
       0.866343 0.456744 0.352079
       0.869205 0.468288 0.365086
       0.871964 0.479783 0.378131
       0.874616 0.491223 0.391223
       0.877165 0.502595 0.404337
       0.879611 0.513947 0.417515
       0.881950 0.525246 0.430724
       0.884181 0.536520 0.443981
       0.886299 0.547752 0.457272
       0.888303 0.558956 0.470622
       0.890194 0.570138 0.484001
       0.891974 0.581300 0.497446
       0.893632 0.592447 0.510915
       0.895179 0.603563 0.524441
       0.896601 0.614657 0.538017
       0.897899 0.625748 0.551626
       0.899078 0.636830 0.565276
       0.900123 0.647885 0.578997
       0.901032 0.658923 0.592753
       0.901799 0.669940 0.606537
       0.902416 0.680902 0.620368
       0.902874 0.691831 0.634220
       0.903157 0.702683 0.648081
       0.903252 0.713443 0.661969
       0.903140 0.724079 0.675831
       0.902797 0.734556 0.689670
       0.902196 0.744811 0.703444
       0.901310 0.754800 0.717128
       0.900107 0.764433 0.730663
       0.898547 0.773639 0.744014
       0.896604 0.782335 0.757114
       0.894232 0.790411 0.769896
       0.891401 0.797775 0.782295
       0.888086 0.804331 0.794228
       0.884268 0.809974 0.805614
       0.879921 0.814628 0.816400
       0.875053 0.818222 0.826500
       0.869659 0.820706 0.835868
       0.863763 0.822052 0.844461
       0.857367 0.822244 0.852254
       0.850519 0.821301 0.859226
       0.843249 0.819266 0.865397
       0.835598 0.816209 0.870788
       0.827602 0.812187 0.875443
       0.819306 0.807307 0.879404
       0.810755 0.801654 0.882741
       0.801980 0.795339 0.885524
       0.793021 0.788462 0.887815
       0.783900 0.781113 0.889692
       0.774635 0.773380 0.891217
       0.765267 0.765358 0.892455
       0.755789 0.757092 0.893452
       0.746222 0.748657 0.894271
       0.736572 0.740101 0.894938
       0.726848 0.731449 0.895489
       0.717053 0.722744 0.895952
       0.707174 0.714010 0.896348
       0.697220 0.705260 0.896692
       0.687192 0.696508 0.896995
       0.677088 0.687757 0.897261
       0.666893 0.679021 0.897497
       0.656625 0.670320 0.897716
       0.646252 0.661634 0.897920
       0.635788 0.652966 0.898109
       0.625218 0.644335 0.898284
       0.614535 0.635732 0.898443
       0.603750 0.627151 0.898589
       0.592833 0.618605 0.898719
       0.581776 0.610071 0.898835
       0.570587 0.601573 0.898936
       0.559254 0.593109 0.899023
       0.547759 0.584666 0.899096
       0.536092 0.576261 0.899155
       0.524243 0.567870 0.899199
       0.512194 0.559519 0.899230
       0.499930 0.551197 0.899247
       0.487451 0.542909 0.899250
       0.474710 0.534647 0.899241
       0.461689 0.526426 0.899218
       0.448371 0.518231 0.899182
       0.434727 0.510066 0.899133
       0.420726 0.501957 0.899071
       0.406366 0.493892 0.898997
       0.391589 0.485884 0.898911
       0.376378 0.477955 0.898813
       0.360670 0.470116 0.898703
       0.344488 0.462367 0.898584
       0.327773 0.454769 0.898455
       0.310525 0.447311 0.898317
       0.292687 0.440071 0.898173
       0.274320 0.433070 0.898025
       0.255438 0.426364 0.897873
       0.236086 0.420024 0.897721
       0.216377 0.414099 0.897572
       0.196465 0.408665 0.897429
       0.176587 0.403793 0.897294
       0.157222 0.399529 0.897173
       0.138935 0.395984 0.897068
       0.122604 0.393164 0.896984
       0.109587 0.391157 0.896922
       0.101101 0.389956 0.896885
       0.098617 0.389623 0.896875
       0.102465 0.390141 0.896891
       0.112045 0.391521 0.896933
       0.125956 0.393705 0.897000
       0.142768 0.396680 0.897089
       0.161373 0.400407 0.897198
       0.180846 0.404784 0.897322
       0.200725 0.409798 0.897459
       0.220656 0.415338 0.897604
       0.240270 0.421352 0.897754
       0.259559 0.427791 0.897906
       0.278335 0.434557 0.898057
       0.296588 0.441618 0.898205
       0.314279 0.448912 0.898347
       0.331417 0.456388 0.898483
       0.348031 0.464036 0.898610
       0.364101 0.471801 0.898728
       0.379687 0.479664 0.898835
       0.394806 0.487615 0.898930
       0.409495 0.495622 0.899014
       0.423773 0.503697 0.899085
       0.437703 0.511820 0.899144
       0.451270 0.519984 0.899191
       0.464526 0.528190 0.899224
       0.477471 0.536428 0.899244
       0.490158 0.544691 0.899251
       0.502595 0.552984 0.899244
       0.514813 0.561314 0.899224
       0.526821 0.569670 0.899191
       0.538629 0.578058 0.899143
       0.550252 0.586482 0.899082
       0.561715 0.594928 0.899006
       0.573020 0.603408 0.898916
       0.584171 0.611908 0.898811
       0.595191 0.620447 0.898692
       0.606077 0.629000 0.898559
       0.616855 0.637586 0.898410
       0.627506 0.646196 0.898247
       0.638052 0.654831 0.898070
       0.648495 0.663506 0.897878
       0.658840 0.672192 0.897670
       0.669102 0.680898 0.897447
       0.679265 0.689646 0.897208
       0.689359 0.698403 0.896951
       0.699353 0.707189 0.896678
       0.709279 0.715984 0.896380
       0.719114 0.724772 0.896052
       0.728867 0.733550 0.895688
       0.738537 0.742304 0.895279
       0.748117 0.751012 0.894812
       0.757591 0.759633 0.894267
       0.766954 0.768152 0.893621
       0.776196 0.776522 0.892856
       0.785292 0.784676 0.891931
       0.794222 0.792552 0.890802
       0.802953 0.800083 0.889428
       0.811472 0.807174 0.887747
       0.819731 0.813733 0.885706
       0.827711 0.819667 0.883231
       0.835372 0.824874 0.880258
       0.842682 0.829258 0.876720
       0.849606 0.832709 0.872561
       0.856118 0.835162 0.867711
       0.862203 0.836544 0.862130
       0.867831 0.836807 0.855777];

case { 'C5'  'CYCLICGREY' }
descriptorname = 'cyclic_grey_15-85_c0_n256';
description = 'Cyclic: greyscale';
map = [0.468775 0.468876 0.468851
       0.473809 0.473910 0.473885
       0.478873 0.478974 0.478949
       0.483952 0.484053 0.484028
       0.489066 0.489168 0.489143
       0.494219 0.494323 0.494298
       0.499390 0.499494 0.499468
       0.504619 0.504723 0.504698
       0.509864 0.509971 0.509945
       0.515152 0.515259 0.515233
       0.520472 0.520582 0.520555
       0.525832 0.525943 0.525915
       0.531213 0.531324 0.531297
       0.536628 0.536741 0.536713
       0.542052 0.542166 0.542138
       0.547505 0.547619 0.547591
       0.552974 0.553088 0.553060
       0.558458 0.558575 0.558546
       0.563974 0.564090 0.564061
       0.569479 0.569597 0.569568
       0.575009 0.575130 0.575100
       0.580553 0.580673 0.580643
       0.586108 0.586229 0.586199
       0.591671 0.591794 0.591764
       0.597244 0.597367 0.597337
       0.602822 0.602947 0.602916
       0.608420 0.608546 0.608515
       0.614009 0.614136 0.614105
       0.619630 0.619758 0.619726
       0.625244 0.625373 0.625341
       0.630879 0.631008 0.630976
       0.636517 0.636648 0.636616
       0.642161 0.642293 0.642261
       0.647813 0.647945 0.647913
       0.653479 0.653614 0.653581
       0.659154 0.659290 0.659256
       0.664836 0.664973 0.664939
       0.670538 0.670675 0.670641
       0.676235 0.676374 0.676340
       0.681941 0.682080 0.682046
       0.687665 0.687806 0.687771
       0.693393 0.693533 0.693499
       0.699127 0.699270 0.699235
       0.704874 0.705017 0.704982
       0.710619 0.710765 0.710729
       0.716361 0.716507 0.716471
       0.722082 0.722228 0.722192
       0.727794 0.727941 0.727905
       0.733474 0.733623 0.733586
       0.739110 0.739261 0.739224
       0.744669 0.744820 0.744783
       0.750140 0.750294 0.750256
       0.755493 0.755645 0.755608
       0.760680 0.760834 0.760797
       0.765666 0.765820 0.765782
       0.770392 0.770549 0.770511
       0.774812 0.774969 0.774930
       0.778871 0.779028 0.778989
       0.782504 0.782661 0.782622
       0.785655 0.785815 0.785775
       0.788265 0.788426 0.788386
       0.790292 0.790452 0.790412
       0.791694 0.791854 0.791815
       0.792444 0.792605 0.792565
       0.792525 0.792685 0.792646
       0.791934 0.792094 0.792055
       0.790685 0.790845 0.790806
       0.788809 0.788968 0.788929
       0.786333 0.786493 0.786454
       0.783303 0.783463 0.783424
       0.779781 0.779939 0.779900
       0.775822 0.775980 0.775941
       0.771487 0.771643 0.771604
       0.766822 0.766977 0.766938
       0.761895 0.762049 0.762011
       0.756755 0.756908 0.756870
       0.751447 0.751599 0.751561
       0.745996 0.746147 0.746110
       0.740452 0.740601 0.740565
       0.734839 0.734988 0.734951
       0.729167 0.729316 0.729279
       0.723462 0.723609 0.723573
       0.717744 0.717888 0.717853
       0.712003 0.712146 0.712111
       0.706263 0.706408 0.706372
       0.700517 0.700660 0.700625
       0.694773 0.694916 0.694881
       0.689051 0.689192 0.689157
       0.683325 0.683464 0.683430
       0.677617 0.677755 0.677721
       0.671908 0.672047 0.672012
       0.666211 0.666346 0.666313
       0.660525 0.660659 0.660626
       0.654846 0.654980 0.654947
       0.649187 0.649320 0.649288
       0.643521 0.643653 0.643620
       0.637875 0.638006 0.637973
       0.632235 0.632365 0.632333
       0.626601 0.626730 0.626699
       0.620984 0.621110 0.621079
       0.615365 0.615492 0.615460
       0.609764 0.609889 0.609858
       0.604177 0.604301 0.604271
       0.598585 0.598709 0.598679
       0.593017 0.593138 0.593109
       0.587443 0.587564 0.587535
       0.581889 0.582009 0.581979
       0.576355 0.576474 0.576444
       0.570806 0.570925 0.570896
       0.565277 0.565394 0.565365
       0.559771 0.559887 0.559858
       0.554263 0.554378 0.554350
       0.548771 0.548884 0.548856
       0.543280 0.543392 0.543364
       0.537807 0.537919 0.537891
       0.532338 0.532450 0.532422
       0.526885 0.526994 0.526967
       0.521429 0.521537 0.521510
       0.515988 0.516096 0.516070
       0.510568 0.510675 0.510648
       0.505151 0.505255 0.505230
       0.499743 0.499848 0.499822
       0.494361 0.494465 0.494439
       0.488970 0.489071 0.489046
       0.483594 0.483696 0.483671
       0.478236 0.478336 0.478312
       0.472880 0.472979 0.472954
       0.467547 0.467644 0.467620
       0.462213 0.462309 0.462285
       0.456893 0.456990 0.456966
       0.451591 0.451688 0.451664
       0.446288 0.446384 0.446360
       0.441000 0.441095 0.441071
       0.435737 0.435829 0.435806
       0.430470 0.430562 0.430539
       0.425221 0.425311 0.425289
       0.419977 0.420066 0.420044
       0.414746 0.414835 0.414813
       0.409532 0.409621 0.409599
       0.404321 0.404407 0.404385
       0.399124 0.399209 0.399188
       0.393946 0.394031 0.394010
       0.388785 0.388869 0.388848
       0.383627 0.383709 0.383689
       0.378476 0.378558 0.378537
       0.373342 0.373424 0.373403
       0.368225 0.368306 0.368286
       0.363110 0.363189 0.363170
       0.358024 0.358100 0.358081
       0.352939 0.353015 0.352997
       0.347868 0.347944 0.347925
       0.342815 0.342890 0.342872
       0.337758 0.337831 0.337813
       0.332740 0.332813 0.332795
       0.327703 0.327776 0.327758
       0.322711 0.322782 0.322765
       0.317708 0.317778 0.317761
       0.312720 0.312791 0.312773
       0.307750 0.307819 0.307802
       0.302797 0.302865 0.302848
       0.297862 0.297928 0.297911
       0.292937 0.293003 0.292986
       0.288025 0.288090 0.288074
       0.283141 0.283204 0.283189
       0.278255 0.278318 0.278303
       0.273391 0.273454 0.273438
       0.268533 0.268596 0.268580
       0.263702 0.263762 0.263748
       0.258870 0.258931 0.258916
       0.254058 0.254117 0.254103
       0.249254 0.249310 0.249296
       0.244464 0.244520 0.244506
       0.239728 0.239782 0.239769
       0.235024 0.235078 0.235065
       0.230301 0.230356 0.230342
       0.225677 0.225732 0.225718
       0.221091 0.221141 0.221129
       0.216565 0.216615 0.216603
       0.212121 0.212170 0.212158
       0.207772 0.207821 0.207809
       0.203575 0.203625 0.203613
       0.199506 0.199554 0.199543
       0.195680 0.195729 0.195717
       0.192073 0.192120 0.192109
       0.188758 0.188804 0.188793
       0.185768 0.185812 0.185801
       0.183084 0.183130 0.183118
       0.180841 0.180885 0.180874
       0.179059 0.179103 0.179092
       0.177753 0.177798 0.177787
       0.176891 0.176937 0.176926
       0.176557 0.176602 0.176591
       0.176761 0.176807 0.176796
       0.177502 0.177547 0.177536
       0.178705 0.178748 0.178737
       0.180371 0.180415 0.180404
       0.182504 0.182549 0.182538
       0.185089 0.185135 0.185124
       0.188024 0.188069 0.188058
       0.191243 0.191290 0.191278
       0.194809 0.194855 0.194843
       0.198570 0.198617 0.198605
       0.202568 0.202618 0.202606
       0.206765 0.206813 0.206801
       0.211053 0.211104 0.211091
       0.215466 0.215520 0.215507
       0.219987 0.220040 0.220027
       0.224532 0.224585 0.224572
       0.229189 0.229240 0.229228
       0.233860 0.233917 0.233903
       0.238574 0.238630 0.238616
       0.243318 0.243374 0.243360
       0.248117 0.248175 0.248161
       0.252897 0.252955 0.252941
       0.257693 0.257750 0.257736
       0.262523 0.262583 0.262568
       0.267341 0.267401 0.267386
       0.272193 0.272256 0.272241
       0.277080 0.277143 0.277127
       0.281962 0.282027 0.282011
       0.286836 0.286901 0.286885
       0.291763 0.291828 0.291812
       0.296685 0.296752 0.296736
       0.301604 0.301671 0.301654
       0.306573 0.306640 0.306623
       0.311521 0.311588 0.311571
       0.316510 0.316579 0.316562
       0.321490 0.321561 0.321543
       0.326488 0.326561 0.326543
       0.331506 0.331581 0.331562
       0.336540 0.336615 0.336596
       0.341589 0.341664 0.341645
       0.346643 0.346720 0.346701
       0.351704 0.351779 0.351761
       0.356798 0.356876 0.356857
       0.361885 0.361964 0.361944
       0.366981 0.367059 0.367040
       0.372084 0.372165 0.372145
       0.377213 0.377294 0.377274
       0.382332 0.382415 0.382394
       0.387464 0.387548 0.387527
       0.392604 0.392689 0.392668
       0.397749 0.397835 0.397814
       0.402882 0.402970 0.402949
       0.408016 0.408103 0.408082
       0.413150 0.413237 0.413216
       0.418255 0.418344 0.418322
       0.423360 0.423451 0.423429
       0.428459 0.428550 0.428528
       0.433528 0.433620 0.433597
       0.438592 0.438684 0.438662
       0.443646 0.443741 0.443717
       0.448682 0.448778 0.448754
       0.453699 0.453796 0.453773
       0.458724 0.458822 0.458798
       0.463745 0.463845 0.463821];

case { 'C6' }
descriptorname = 'cyclic-isoluminant colour circle_mgbm_67_c31_n256';
description = 'Cyclic: isoluminant';
map = [0.852580 0.567041 0.521037
       0.851095 0.568174 0.516638
       0.849462 0.569374 0.512338
       0.847659 0.570638 0.508155
       0.845706 0.571964 0.504068
       0.843601 0.573341 0.500086
       0.841355 0.574758 0.496229
       0.838961 0.576244 0.492495
       0.836434 0.577747 0.488883
       0.833771 0.579315 0.485395
       0.830981 0.580904 0.482039
       0.828070 0.582528 0.478790
       0.825052 0.584185 0.475672
       0.821933 0.585870 0.472659
       0.818702 0.587563 0.469780
       0.815386 0.589281 0.467003
       0.811976 0.591023 0.464347
       0.808482 0.592780 0.461789
       0.804904 0.594535 0.459347
       0.801250 0.596306 0.457002
       0.797524 0.598079 0.454775
       0.793736 0.599867 0.452636
       0.789872 0.601649 0.450585
       0.785954 0.603440 0.448636
       0.781972 0.605222 0.446771
       0.777929 0.607017 0.445007
       0.773827 0.608805 0.443322
       0.769680 0.610580 0.441722
       0.765489 0.612353 0.440188
       0.761194 0.614143 0.438715
       0.756770 0.615968 0.437305
       0.753155 0.617435 0.436202
       0.747997 0.619510 0.434741
       0.743802 0.621159 0.433639
       0.739337 0.622898 0.432553
       0.734820 0.624630 0.431523
       0.730260 0.626351 0.430560
       0.725654 0.628071 0.429682
       0.720989 0.629775 0.428896
       0.716289 0.631480 0.428198
       0.711532 0.633171 0.427591
       0.706726 0.634855 0.427073
       0.701859 0.636534 0.426646
       0.696952 0.638195 0.426326
       0.691991 0.639855 0.426116
       0.686969 0.641494 0.426021
       0.681892 0.643133 0.426044
       0.676772 0.644756 0.426189
       0.671584 0.646373 0.426465
       0.666345 0.647970 0.426880
       0.661054 0.649566 0.427431
       0.655714 0.651138 0.428115
       0.650314 0.652691 0.428951
       0.644860 0.654236 0.429943
       0.639367 0.655766 0.431095
       0.633824 0.657275 0.432408
       0.628234 0.658754 0.433863
       0.622597 0.660223 0.435518
       0.616931 0.661672 0.437326
       0.611229 0.663091 0.439306
       0.605489 0.664478 0.441478
       0.599738 0.665847 0.443818
       0.593957 0.667185 0.446319
       0.588157 0.668499 0.449003
       0.582354 0.669779 0.451844
       0.576554 0.671025 0.454848
       0.570733 0.672241 0.458006
       0.564930 0.673420 0.461339
       0.559141 0.674565 0.464814
       0.553357 0.675678 0.468421
       0.547596 0.676760 0.472174
       0.541845 0.677805 0.476042
       0.536124 0.678809 0.480032
       0.530423 0.679782 0.484123
       0.524745 0.680723 0.488338
       0.519103 0.681632 0.492632
       0.513480 0.682510 0.497033
       0.507888 0.683356 0.501517
       0.502312 0.684164 0.506059
       0.496759 0.684953 0.510686
       0.491249 0.685710 0.515371
       0.485726 0.686434 0.520113
       0.480242 0.687135 0.524918
       0.474770 0.687810 0.529766
       0.469307 0.688462 0.534654
       0.463843 0.689090 0.539579
       0.458391 0.689690 0.544553
       0.452953 0.690264 0.549556
       0.447495 0.690820 0.554588
       0.442057 0.691360 0.559648
       0.436586 0.691885 0.564724
       0.431105 0.692389 0.569828
       0.425597 0.692873 0.574953
       0.420065 0.693337 0.580105
       0.414506 0.693782 0.585273
       0.408934 0.694213 0.590438
       0.403324 0.694633 0.595625
       0.397660 0.695041 0.600822
       0.391949 0.695435 0.606030
       0.386190 0.695814 0.611261
       0.380429 0.696172 0.616497
       0.374627 0.696512 0.621735
       0.368798 0.696835 0.626995
       0.362932 0.697140 0.632262
       0.357073 0.697428 0.637535
       0.351186 0.697695 0.642816
       0.345298 0.697943 0.648098
       0.339438 0.698168 0.653389
       0.333584 0.698371 0.658684
       0.327760 0.698550 0.663985
       0.322002 0.698704 0.669278
       0.316320 0.698831 0.674565
       0.310723 0.698931 0.679844
       0.305227 0.699001 0.685123
       0.299868 0.699042 0.690379
       0.294667 0.699050 0.695625
       0.289678 0.699026 0.700844
       0.284891 0.698968 0.706038
       0.280370 0.698874 0.711200
       0.276171 0.698743 0.716325
       0.272243 0.698575 0.721397
       0.268736 0.698367 0.726432
       0.265599 0.698120 0.731404
       0.262911 0.697832 0.736315
       0.260695 0.697501 0.741157
       0.259061 0.697125 0.745919
       0.257926 0.696707 0.750598
       0.257385 0.696245 0.755183
       0.257403 0.695737 0.759665
       0.257990 0.695183 0.764055
       0.259197 0.694585 0.768332
       0.260942 0.693948 0.772502
       0.263335 0.693271 0.776545
       0.266257 0.692550 0.780460
       0.269703 0.691780 0.784261
       0.273631 0.690968 0.787926
       0.278011 0.690129 0.791471
       0.282824 0.689252 0.794890
       0.287995 0.688332 0.798179
       0.293535 0.687376 0.801343
       0.299404 0.686388 0.804385
       0.305542 0.685371 0.807298
       0.311895 0.684311 0.810086
       0.318505 0.683235 0.812758
       0.325275 0.682118 0.815316
       0.332233 0.680977 0.817754
       0.339309 0.679811 0.820083
       0.346480 0.678621 0.822311
       0.353753 0.677409 0.824423
       0.361102 0.676161 0.826443
       0.368521 0.674893 0.828361
       0.375967 0.673605 0.830189
       0.383448 0.672297 0.831926
       0.390944 0.670964 0.833574
       0.398445 0.669611 0.835138
       0.405954 0.668239 0.836624
       0.413450 0.666838 0.838033
       0.420949 0.665428 0.839355
       0.428449 0.663997 0.840606
       0.435914 0.662548 0.841787
       0.443334 0.661075 0.842901
       0.450731 0.659591 0.843949
       0.458104 0.658089 0.844934
       0.465467 0.656566 0.845832
       0.472817 0.655015 0.846661
       0.480153 0.653455 0.847415
       0.487469 0.651881 0.848090
       0.494770 0.650279 0.848683
       0.502060 0.648659 0.849189
       0.509346 0.647023 0.849602
       0.516625 0.645359 0.849918
       0.523910 0.643682 0.850129
       0.531191 0.641989 0.850235
       0.538476 0.640275 0.850233
       0.545750 0.638538 0.850118
       0.553015 0.636789 0.849887
       0.560292 0.635014 0.849533
       0.567557 0.633228 0.849050
       0.574810 0.631425 0.848434
       0.582063 0.629601 0.847683
       0.589301 0.627772 0.846792
       0.596536 0.625924 0.845751
       0.603749 0.624073 0.844559
       0.610934 0.622198 0.843208
       0.618087 0.620336 0.841705
       0.625208 0.618457 0.840041
       0.632288 0.616577 0.838210
       0.639318 0.614690 0.836220
       0.646288 0.612814 0.834070
       0.653192 0.610952 0.831754
       0.660034 0.609095 0.829264
       0.666784 0.607249 0.826604
       0.673454 0.605408 0.823785
       0.680017 0.603604 0.820814
       0.686487 0.601808 0.817691
       0.692847 0.600044 0.814424
       0.699075 0.598300 0.811017
       0.705192 0.596598 0.807471
       0.711178 0.594922 0.803786
       0.717031 0.593286 0.799983
       0.722738 0.591682 0.796058
       0.728320 0.590108 0.792033
       0.733763 0.588579 0.787903
       0.739069 0.587093 0.783686
       0.744225 0.585654 0.779373
       0.749244 0.584240 0.774981
       0.754135 0.582870 0.770519
       0.758880 0.581546 0.765986
       0.763490 0.580267 0.761385
       0.767969 0.579023 0.756730
       0.772321 0.577812 0.752023
       0.776547 0.576662 0.747267
       0.780642 0.575534 0.742466
       0.784624 0.574439 0.737620
       0.788481 0.573400 0.732747
       0.792226 0.572390 0.727832
       0.795859 0.571409 0.722889
       0.799392 0.570467 0.717927
       0.802809 0.569561 0.712930
       0.806130 0.568692 0.707916
       0.809346 0.567860 0.702881
       0.812462 0.567061 0.697817
       0.815495 0.566284 0.692750
       0.818432 0.565533 0.687655
       0.821284 0.564821 0.682562
       0.824046 0.564148 0.677456
       0.826722 0.563500 0.672327
       0.829309 0.562881 0.667185
       0.831807 0.562297 0.662042
       0.834212 0.561741 0.656880
       0.836522 0.561220 0.651709
       0.838735 0.560744 0.646526
       0.840851 0.560313 0.641331
       0.842856 0.559924 0.636140
       0.844756 0.559576 0.630937
       0.846544 0.559274 0.625720
       0.848225 0.559020 0.620513
       0.849792 0.558817 0.615286
       0.851231 0.558668 0.610070
       0.852561 0.558574 0.604859
       0.853747 0.558538 0.599650
       0.854815 0.558562 0.594444
       0.855743 0.558648 0.589248
       0.856527 0.558798 0.584075
       0.857170 0.559014 0.578920
       0.857666 0.559299 0.573783
       0.858011 0.559653 0.568674
       0.858201 0.560075 0.563610
       0.858222 0.560569 0.558565
       0.858079 0.561138 0.553581
       0.857772 0.561790 0.548657
       0.857299 0.562510 0.543773
       0.856658 0.563300 0.538962
       0.855851 0.564162 0.534214
       0.854873 0.565086 0.529553
       0.853721 0.566097 0.524967];

case { 'C7' }
descriptorname = 'cyclic_mygbm_50-90_c46_n256';
description = 'Cyclic: low contrast colour ellipse';
map = [0.912976 0.580896 0.585646
       0.915613 0.584998 0.580934
       0.918123 0.589141 0.576268
       0.920488 0.593354 0.571612
       0.922727 0.597608 0.567002
       0.924822 0.601909 0.562419
       0.926787 0.606257 0.557855
       0.928616 0.610657 0.553328
       0.930317 0.615088 0.548848
       0.931877 0.619574 0.544383
       0.933307 0.624088 0.539952
       0.934595 0.628642 0.535562
       0.935751 0.633229 0.531214
       0.936777 0.637849 0.526900
       0.937668 0.642500 0.522606
       0.938433 0.647178 0.518359
       0.939072 0.651882 0.514140
       0.939586 0.656608 0.509947
       0.939974 0.661351 0.505787
       0.940233 0.666116 0.501684
       0.940356 0.670908 0.497611
       0.940358 0.675708 0.493558
       0.940240 0.680520 0.489545
       0.940003 0.685356 0.485570
       0.939647 0.690191 0.481631
       0.939175 0.695038 0.477717
       0.938583 0.699893 0.473837
       0.937863 0.704758 0.470011
       0.937032 0.709628 0.466197
       0.936089 0.714491 0.462421
       0.935032 0.719368 0.458682
       0.933869 0.724236 0.454972
       0.932591 0.729106 0.451289
       0.931193 0.733992 0.447627
       0.929634 0.739034 0.443899
       0.928205 0.743366 0.440689
       0.926356 0.748617 0.436853
       0.924545 0.753479 0.433298
       0.922640 0.758256 0.429867
       0.920597 0.763087 0.426403
       0.918448 0.767913 0.422977
       0.916177 0.772733 0.419607
       0.913783 0.777547 0.416270
       0.911245 0.782342 0.413002
       0.908564 0.787129 0.409788
       0.905725 0.791897 0.406650
       0.902715 0.796651 0.403606
       0.899532 0.801390 0.400658
       0.896159 0.806109 0.397803
       0.892579 0.810798 0.395084
       0.888775 0.815463 0.392509
       0.884732 0.820087 0.390097
       0.880434 0.824674 0.387879
       0.875869 0.829217 0.385864
       0.871009 0.833696 0.384095
       0.865809 0.838099 0.382619
       0.860258 0.842415 0.381477
       0.854343 0.846621 0.380676
       0.848004 0.850691 0.380323
       0.841260 0.854610 0.380425
       0.834063 0.858317 0.381083
       0.826430 0.861804 0.382281
       0.818356 0.864995 0.384139
       0.809893 0.867873 0.386632
       0.801088 0.870395 0.389760
       0.792003 0.872526 0.393540
       0.782724 0.874251 0.397894
       0.773326 0.875593 0.402729
       0.763896 0.876528 0.408018
       0.754476 0.877123 0.413643
       0.745119 0.877368 0.419564
       0.735844 0.877325 0.425680
       0.726686 0.877007 0.431975
       0.717648 0.876452 0.438365
       0.708713 0.875696 0.444847
       0.699880 0.874751 0.451370
       0.691175 0.873631 0.457929
       0.682572 0.872369 0.464518
       0.674055 0.870973 0.471090
       0.665618 0.869465 0.477653
       0.657255 0.867855 0.484201
       0.648948 0.866146 0.490746
       0.640692 0.864356 0.497244
       0.632485 0.862490 0.503716
       0.624314 0.860548 0.510160
       0.616164 0.858544 0.516572
       0.608036 0.856487 0.522954
       0.599906 0.854378 0.529300
       0.591784 0.852223 0.535599
       0.583652 0.850008 0.541881
       0.575522 0.847756 0.548129
       0.567366 0.845474 0.554326
       0.559167 0.843159 0.560491
       0.550933 0.840812 0.566631
       0.542685 0.838418 0.572736
       0.534393 0.835994 0.578801
       0.526071 0.833539 0.584841
       0.517705 0.831050 0.590840
       0.509294 0.828524 0.596817
       0.500857 0.825963 0.602752
       0.492395 0.823364 0.608662
       0.483896 0.820731 0.614519
       0.475370 0.818065 0.620366
       0.466787 0.815369 0.626160
       0.458160 0.812634 0.631928
       0.449505 0.809870 0.637659
       0.440826 0.807066 0.643348
       0.432134 0.804225 0.649009
       0.423368 0.801349 0.654620
       0.414584 0.798443 0.660203
       0.405763 0.795503 0.665745
       0.396878 0.792533 0.671246
       0.387973 0.789529 0.676711
       0.379034 0.786493 0.682123
       0.370077 0.783414 0.687501
       0.361084 0.780307 0.692837
       0.352055 0.777175 0.698115
       0.343003 0.773998 0.703361
       0.333894 0.770804 0.708559
       0.324734 0.767570 0.713704
       0.315566 0.764314 0.718802
       0.306350 0.761028 0.723853
       0.297087 0.757711 0.728855
       0.287806 0.754367 0.733803
       0.278531 0.750993 0.738697
       0.269216 0.747589 0.743538
       0.259849 0.744157 0.748323
       0.250435 0.740701 0.753049
       0.241010 0.737225 0.757729
       0.231563 0.733723 0.762345
       0.222035 0.730199 0.766907
       0.212503 0.726647 0.771416
       0.202913 0.723073 0.775857
       0.193318 0.719487 0.780241
       0.183722 0.715873 0.784565
       0.174115 0.712230 0.788823
       0.164476 0.708581 0.793021
       0.154772 0.704903 0.797152
       0.145088 0.701212 0.801229
       0.135309 0.697502 0.805240
       0.125529 0.693775 0.809196
       0.115702 0.690038 0.813079
       0.105845 0.686281 0.816908
       0.096034 0.682506 0.820663
       0.086314 0.678718 0.824355
       0.076594 0.674917 0.827984
       0.066987 0.671103 0.831559
       0.057363 0.667274 0.835061
       0.047913 0.663438 0.838503
       0.038783 0.659581 0.841889
       0.030423 0.655719 0.845207
       0.024144 0.651842 0.848457
       0.019383 0.647945 0.851646
       0.016091 0.644045 0.854778
       0.014241 0.640142 0.857846
       0.013802 0.636222 0.860859
       0.014758 0.632288 0.863817
       0.017308 0.628349 0.866701
       0.021411 0.624394 0.869529
       0.026822 0.620434 0.872303
       0.033498 0.616458 0.875016
       0.041669 0.612468 0.877678
       0.050034 0.608484 0.880283
       0.058959 0.604471 0.882825
       0.067796 0.600451 0.885316
       0.076491 0.596432 0.887746
       0.085201 0.592403 0.890132
       0.093901 0.588342 0.892465
       0.102751 0.584286 0.894729
       0.111716 0.580216 0.896925
       0.120706 0.576134 0.899053
       0.129928 0.572025 0.901108
       0.139192 0.567907 0.903089
       0.148704 0.563781 0.904982
       0.158460 0.559629 0.906782
       0.168397 0.555456 0.908483
       0.178641 0.551281 0.910077
       0.189142 0.547081 0.911547
       0.199940 0.542870 0.912891
       0.211116 0.538640 0.914095
       0.222618 0.534386 0.915137
       0.234560 0.530129 0.916002
       0.246955 0.525863 0.916669
       0.259798 0.521585 0.917123
       0.273045 0.517325 0.917346
       0.286909 0.513068 0.917274
       0.301260 0.508847 0.916913
       0.316190 0.504671 0.916207
       0.331573 0.500555 0.915149
       0.347516 0.496550 0.913685
       0.363815 0.492687 0.911805
       0.380474 0.489012 0.909484
       0.397306 0.485561 0.906728
       0.414164 0.482370 0.903566
       0.430919 0.479476 0.900008
       0.447391 0.476889 0.896110
       0.463493 0.474621 0.891927
       0.479141 0.472668 0.887490
       0.494276 0.471018 0.882866
       0.508875 0.469663 0.878085
       0.522934 0.468557 0.873178
       0.536482 0.467693 0.868186
       0.549513 0.467075 0.863109
       0.562077 0.466654 0.857974
       0.574178 0.466416 0.852806
       0.585883 0.466350 0.847598
       0.597185 0.466445 0.842376
       0.608123 0.466689 0.837131
       0.618715 0.467071 0.831875
       0.628979 0.467575 0.826606
       0.638951 0.468194 0.821334
       0.648641 0.468938 0.816061
       0.658071 0.469781 0.810780
       0.667248 0.470713 0.805503
       0.676198 0.471743 0.800233
       0.684931 0.472843 0.794957
       0.693464 0.474036 0.789686
       0.701807 0.475311 0.784417
       0.709981 0.476638 0.779140
       0.717946 0.478044 0.773885
       0.725756 0.479528 0.768628
       0.733409 0.481059 0.763375
       0.740909 0.482668 0.758127
       0.748269 0.484331 0.752873
       0.755469 0.486075 0.747638
       0.762509 0.487901 0.742400
       0.769412 0.489773 0.737167
       0.776177 0.491738 0.731943
       0.782798 0.493751 0.726721
       0.789286 0.495840 0.721506
       0.795620 0.498028 0.716310
       0.801814 0.500263 0.711115
       0.807876 0.502586 0.705928
       0.813798 0.504985 0.700752
       0.819589 0.507456 0.695586
       0.825248 0.509997 0.690426
       0.830776 0.512615 0.685287
       0.836156 0.515313 0.680150
       0.841399 0.518100 0.675039
       0.846505 0.520949 0.669943
       0.851482 0.523879 0.664847
       0.856329 0.526887 0.659780
       0.861038 0.529959 0.654718
       0.865615 0.533107 0.649686
       0.870056 0.536334 0.644658
       0.874365 0.539624 0.639661
       0.878527 0.543009 0.634682
       0.882551 0.546459 0.629717
       0.886448 0.549972 0.624782
       0.890201 0.553552 0.619869
       0.893823 0.557217 0.614966
       0.897312 0.560927 0.610098
       0.900659 0.564711 0.605252
       0.903872 0.568559 0.600432
       0.906953 0.572476 0.595637
       0.909895 0.576450 0.590868];

case { 'C8' }
descriptorname = 'cyclic_mybm_20-100_c48_n256';
description = 'Cyclic: colour ellipse';
map = [0.761632 0.498431 0.458066
       0.765168 0.505669 0.455815
       0.768590 0.512957 0.453582
       0.771920 0.520255 0.451380
       0.775137 0.527583 0.449199
       0.778256 0.534924 0.447034
       0.781267 0.542282 0.444905
       0.784175 0.549654 0.442795
       0.786973 0.557044 0.440699
       0.789662 0.564437 0.438640
       0.792246 0.571848 0.436611
       0.794725 0.579275 0.434597
       0.797094 0.586705 0.432615
       0.799366 0.594152 0.430648
       0.801519 0.601609 0.428707
       0.803572 0.609086 0.426793
       0.805515 0.616562 0.424905
       0.807358 0.624052 0.423026
       0.809091 0.631548 0.421182
       0.810709 0.639054 0.419373
       0.812226 0.646570 0.417576
       0.813632 0.654092 0.415798
       0.814932 0.661626 0.414051
       0.816127 0.669159 0.412334
       0.817203 0.676701 0.410635
       0.818172 0.684238 0.408951
       0.819031 0.691794 0.407299
       0.819780 0.699341 0.405681
       0.820418 0.706901 0.404074
       0.820943 0.714449 0.402493
       0.821355 0.722003 0.400963
       0.821652 0.729564 0.399420
       0.821831 0.737116 0.397942
       0.821892 0.744666 0.396471
       0.821833 0.752215 0.395042
       0.821650 0.759757 0.393638
       0.821340 0.767298 0.392275
       0.820900 0.774832 0.390955
       0.820327 0.782362 0.389659
       0.819615 0.789879 0.388423
       0.818756 0.797383 0.387225
       0.817745 0.804878 0.386090
       0.816573 0.812353 0.385024
       0.815219 0.819807 0.384035
       0.813679 0.827240 0.383131
       0.811941 0.834636 0.382308
       0.809981 0.841990 0.381617
       0.807789 0.849290 0.381053
       0.805336 0.856521 0.380629
       0.802609 0.863678 0.380375
       0.799587 0.870723 0.380316
       0.796228 0.877654 0.380479
       0.792530 0.884434 0.380890
       0.788456 0.891030 0.381566
       0.783990 0.897419 0.382544
       0.779099 0.903553 0.383864
       0.773778 0.909395 0.385524
       0.768012 0.914907 0.387574
       0.761795 0.920043 0.390015
       0.755129 0.924757 0.392870
       0.748012 0.929013 0.396137
       0.740462 0.932768 0.399799
       0.732511 0.935996 0.403880
       0.724175 0.938654 0.408326
       0.715511 0.940731 0.413136
       0.706547 0.942212 0.418248
       0.697331 0.943090 0.423655
       0.687929 0.943369 0.429307
       0.678380 0.943064 0.435157
       0.668741 0.942192 0.441140
       0.659051 0.940778 0.447241
       0.649375 0.938857 0.453412
       0.639728 0.936468 0.459614
       0.630156 0.933635 0.465798
       0.620687 0.930422 0.471973
       0.611331 0.926847 0.478069
       0.602106 0.922968 0.484093
       0.593034 0.918803 0.490035
       0.584094 0.914408 0.495871
       0.575304 0.909799 0.501612
       0.566655 0.905011 0.507226
       0.558126 0.900065 0.512732
       0.549735 0.894990 0.518132
       0.541443 0.889797 0.523414
       0.533266 0.884511 0.528589
       0.525180 0.879137 0.533646
       0.517179 0.873696 0.538624
       0.509242 0.868197 0.543485
       0.501374 0.862639 0.548261
       0.493551 0.857028 0.552931
       0.485783 0.851378 0.557529
       0.478059 0.845695 0.562036
       0.470374 0.839981 0.566452
       0.462703 0.834228 0.570786
       0.455084 0.828444 0.575047
       0.447463 0.822638 0.579236
       0.439886 0.816802 0.583336
       0.432327 0.810941 0.587367
       0.424772 0.805057 0.591329
       0.417245 0.799155 0.595212
       0.409728 0.793226 0.599027
       0.402216 0.787274 0.602765
       0.394741 0.781303 0.606433
       0.387270 0.775308 0.610030
       0.379825 0.769295 0.613554
       0.372385 0.763265 0.617019
       0.364970 0.757217 0.620410
       0.357583 0.751154 0.623727
       0.350192 0.745061 0.626968
       0.342827 0.738962 0.630150
       0.335476 0.732840 0.633259
       0.328131 0.726698 0.636302
       0.320826 0.720543 0.639270
       0.313526 0.714372 0.642174
       0.306256 0.708191 0.645003
       0.298990 0.701987 0.647770
       0.291753 0.695780 0.650471
       0.284520 0.689554 0.653095
       0.277336 0.683310 0.655664
       0.270168 0.677062 0.658158
       0.263007 0.670796 0.660578
       0.255862 0.664512 0.662944
       0.248772 0.658229 0.665229
       0.241680 0.651932 0.667457
       0.234628 0.645617 0.669614
       0.227580 0.639302 0.671699
       0.220581 0.632975 0.673724
       0.213567 0.626637 0.675680
       0.206634 0.620300 0.677575
       0.199656 0.613937 0.679391
       0.192770 0.607589 0.681147
       0.185917 0.601216 0.682844
       0.179054 0.594845 0.684467
       0.172228 0.588462 0.686034
       0.165429 0.582082 0.687530
       0.158727 0.575703 0.688967
       0.151988 0.569300 0.690328
       0.145365 0.562912 0.691634
       0.138704 0.556504 0.692880
       0.132127 0.550102 0.694047
       0.125595 0.543693 0.695165
       0.119102 0.537287 0.696220
       0.112695 0.530867 0.697205
       0.106335 0.524450 0.698131
       0.099987 0.518034 0.698997
       0.093771 0.511605 0.699804
       0.087677 0.505180 0.700552
       0.081652 0.498762 0.701234
       0.075664 0.492327 0.701856
       0.069923 0.485895 0.702425
       0.064270 0.479470 0.702936
       0.058824 0.473024 0.703384
       0.053521 0.466595 0.703774
       0.048609 0.460148 0.704107
       0.043840 0.453709 0.704385
       0.039571 0.447261 0.704606
       0.035783 0.440810 0.704771
       0.032406 0.434361 0.704880
       0.029883 0.427911 0.704935
       0.027994 0.421439 0.704934
       0.026730 0.414978 0.704878
       0.026090 0.408505 0.704767
       0.026078 0.402017 0.704601
       0.026701 0.395546 0.704379
       0.027974 0.389050 0.704101
       0.029920 0.382538 0.703765
       0.032576 0.376031 0.703372
       0.036166 0.369499 0.702917
       0.040364 0.362959 0.702394
       0.045202 0.356417 0.701806
       0.050548 0.349848 0.701154
       0.056478 0.343269 0.700425
       0.062852 0.336664 0.699612
       0.069668 0.330061 0.698715
       0.076852 0.323430 0.697727
       0.084667 0.316794 0.696642
       0.092813 0.310156 0.695445
       0.101512 0.303486 0.694124
       0.110689 0.296854 0.692693
       0.120261 0.290197 0.691103
       0.130402 0.283588 0.689386
       0.140936 0.277009 0.687498
       0.151895 0.270483 0.685451
       0.163358 0.264016 0.683226
       0.175186 0.257625 0.680817
       0.187421 0.251389 0.678237
       0.199942 0.245255 0.675463
       0.212816 0.239347 0.672512
       0.225952 0.233621 0.669374
       0.239248 0.228201 0.666064
       0.252705 0.223045 0.662603
       0.266249 0.218224 0.658983
       0.279818 0.213789 0.655242
       0.293355 0.209774 0.651396
       0.306838 0.206253 0.647453
       0.320194 0.203176 0.643441
       0.333367 0.200630 0.639390
       0.346319 0.198659 0.635305
       0.359042 0.197263 0.631220
       0.371494 0.196426 0.627137
       0.383667 0.196131 0.623087
       0.395523 0.196405 0.619075
       0.407057 0.197193 0.615096
       0.418273 0.198474 0.611192
       0.429183 0.200239 0.607344
       0.439775 0.202463 0.603555
       0.450060 0.205084 0.599832
       0.460046 0.208064 0.596173
       0.469768 0.211376 0.592588
       0.479205 0.214952 0.589044
       0.488396 0.218828 0.585583
       0.497347 0.222924 0.582155
       0.506061 0.227213 0.578792
       0.514586 0.231701 0.575476
       0.522900 0.236339 0.572200
       0.531026 0.241111 0.568961
       0.538980 0.246021 0.565766
       0.546768 0.251075 0.562617
       0.554397 0.256199 0.559484
       0.561882 0.261452 0.556383
       0.569212 0.266780 0.553314
       0.576430 0.272188 0.550281
       0.583494 0.277708 0.547265
       0.590443 0.283283 0.544279
       0.597279 0.288932 0.541309
       0.603993 0.294635 0.538381
       0.610587 0.300422 0.535453
       0.617073 0.306290 0.532568
       0.623453 0.312173 0.529695
       0.629714 0.318160 0.526850
       0.635881 0.324185 0.524017
       0.641938 0.330281 0.521211
       0.647891 0.336418 0.518436
       0.653748 0.342631 0.515663
       0.659501 0.348876 0.512931
       0.665153 0.355193 0.510214
       0.670714 0.361538 0.507524
       0.676166 0.367940 0.504852
       0.681519 0.374407 0.502200
       0.686784 0.380914 0.499563
       0.691952 0.387454 0.496960
       0.697016 0.394059 0.494381
       0.701986 0.400725 0.491819
       0.706870 0.407407 0.489264
       0.711653 0.414150 0.486747
       0.716342 0.420934 0.484239
       0.720928 0.427780 0.481766
       0.725435 0.434656 0.479306
       0.729838 0.441582 0.476862
       0.734148 0.448550 0.474445
       0.738362 0.455558 0.472048
       0.742487 0.462603 0.469669
       0.746509 0.469710 0.467305
       0.750440 0.476831 0.464961
       0.754273 0.483991 0.462635
       0.758003 0.491206 0.460337];

case { 'C9' }
descriptorname = 'cyclic_bgrmb_35-70_c75_n256';
description = 'Cyclic: blue - green - red - magenta. Allows four orientations/phase angles to be visulaised.';
map = [0.279654 0.191839 0.947123
       0.278959 0.197983 0.941022
       0.280276 0.204885 0.934325
       0.283454 0.212452 0.927052
       0.288180 0.220627 0.919229
       0.294173 0.229270 0.910899
       0.301121 0.238361 0.902097
       0.308749 0.247825 0.892880
       0.316744 0.257521 0.883298
       0.324909 0.267417 0.873402
       0.333094 0.277489 0.863247
       0.341099 0.287608 0.852870
       0.348852 0.297786 0.842321
       0.356274 0.307958 0.831632
       0.363293 0.318119 0.820828
       0.369902 0.328222 0.809947
       0.376083 0.338257 0.799006
       0.381804 0.348236 0.788010
       0.387082 0.358128 0.776989
       0.391934 0.367932 0.765932
       0.396349 0.377665 0.754857
       0.400353 0.387309 0.743760
       0.403934 0.396880 0.732651
       0.407115 0.406386 0.721522
       0.409918 0.415802 0.710393
       0.412336 0.425163 0.699240
       0.414378 0.434447 0.688079
       0.416065 0.443686 0.676898
       0.417405 0.452865 0.665690
       0.418376 0.461986 0.654457
       0.419020 0.471068 0.643203
       0.419319 0.480101 0.631921
       0.419272 0.489085 0.620608
       0.418881 0.498051 0.609252
       0.418151 0.506953 0.597852
       0.417095 0.515825 0.586417
       0.415675 0.524679 0.574925
       0.413917 0.533488 0.563395
       0.411805 0.542279 0.551794
       0.409339 0.551036 0.540124
       0.406508 0.559768 0.528393
       0.403304 0.568468 0.516575
       0.399702 0.577154 0.504683
       0.395738 0.585821 0.492661
       0.391362 0.594449 0.480547
       0.386569 0.603050 0.468310
       0.381393 0.611609 0.455945
       0.375783 0.620125 0.443438
       0.369762 0.628572 0.430769
       0.363337 0.636942 0.417923
       0.356533 0.645203 0.404891
       0.349353 0.653339 0.391673
       0.341864 0.661311 0.378233
       0.334133 0.669063 0.364579
       0.326229 0.676545 0.350723
       0.318331 0.683688 0.336625
       0.310577 0.690430 0.322349
       0.303131 0.696694 0.307869
       0.296302 0.702389 0.293266
       0.290302 0.707439 0.278577
       0.285446 0.711766 0.263846
       0.282030 0.715287 0.249130
       0.280241 0.717941 0.234555
       0.280322 0.719678 0.220176
       0.282389 0.720457 0.206111
       0.286370 0.720270 0.192429
       0.292261 0.719114 0.179287
       0.299835 0.717014 0.166767
       0.308869 0.713994 0.154905
       0.319056 0.710129 0.143839
       0.330155 0.705473 0.133573
       0.341875 0.700106 0.124114
       0.353974 0.694103 0.115548
       0.366272 0.687561 0.107755
       0.378561 0.680538 0.100738
       0.390739 0.673134 0.094469
       0.402686 0.665391 0.088741
       0.414348 0.657391 0.083562
       0.425666 0.649169 0.078779
       0.436617 0.640770 0.074448
       0.447176 0.632242 0.070334
       0.457354 0.623605 0.066462
       0.467159 0.614864 0.062800
       0.476581 0.606057 0.059179
       0.485645 0.597195 0.055600
       0.494381 0.588258 0.052090
       0.502774 0.579283 0.048789
       0.510863 0.570247 0.045323
       0.518663 0.561175 0.041976
       0.526171 0.552060 0.038511
       0.533408 0.542885 0.035214
       0.540407 0.533645 0.031867
       0.547162 0.524363 0.028877
       0.553688 0.515013 0.026036
       0.560008 0.505587 0.023343
       0.566110 0.496100 0.020796
       0.572021 0.486542 0.018393
       0.577734 0.476896 0.016132
       0.583275 0.467169 0.014011
       0.588636 0.457334 0.012051
       0.593840 0.447402 0.009965
       0.598875 0.437379 0.008255
       0.603760 0.427225 0.006667
       0.608493 0.416943 0.005204
       0.613067 0.406520 0.003866
       0.617516 0.395955 0.002651
       0.621824 0.385210 0.001556
       0.626015 0.374312 0.000601
       0.630083 0.363200 0.000000
       0.634036 0.351894 0.000000
       0.637873 0.340383 0.000000
       0.641612 0.328626 0.000000
       0.645263 0.316617 0.000000
       0.648843 0.304332 0.000000
       0.652351 0.291782 0.000000
       0.655818 0.278929 0.000000
       0.659246 0.265763 0.000716
       0.662680 0.252284 0.002273
       0.666120 0.238467 0.004328
       0.669616 0.224333 0.006956
       0.673182 0.209907 0.010213
       0.676854 0.195153 0.014464
       0.680650 0.180081 0.019341
       0.684620 0.164737 0.025185
       0.688783 0.149117 0.032125
       0.693148 0.133244 0.040511
       0.697733 0.117107 0.049454
       0.702561 0.100784 0.058636
       0.707621 0.084326 0.068259
       0.712913 0.067632 0.078128
       0.718427 0.050818 0.088512
       0.724144 0.034099 0.099205
       0.730042 0.020769 0.110339
       0.736082 0.011276 0.121741
       0.742260 0.005477 0.133608
       0.748529 0.003229 0.145782
       0.754872 0.004142 0.158227
       0.761252 0.007923 0.170939
       0.767656 0.014473 0.183900
       0.774067 0.023110 0.197089
       0.780466 0.033735 0.210421
       0.786843 0.045986 0.223905
       0.793177 0.057778 0.237550
       0.799470 0.069155 0.251264
       0.805706 0.079760 0.265054
       0.811897 0.090028 0.278962
       0.818026 0.099812 0.292907
       0.824099 0.109281 0.306940
       0.830114 0.118373 0.321031
       0.836062 0.127122 0.335186
       0.841948 0.135671 0.349370
       0.847762 0.144002 0.363612
       0.853516 0.152070 0.377917
       0.859210 0.159966 0.392274
       0.864834 0.167747 0.406691
       0.870390 0.175367 0.421155
       0.875886 0.182854 0.435697
       0.881309 0.190258 0.450280
       0.886666 0.197564 0.464926
       0.891952 0.204790 0.479636
       0.897170 0.211946 0.494400
       0.902314 0.219012 0.509221
       0.907392 0.226042 0.524101
       0.912394 0.232980 0.539043
       0.917330 0.239904 0.554042
       0.922187 0.246778 0.569096
       0.926969 0.253622 0.584215
       0.931682 0.260417 0.599394
       0.936325 0.267170 0.614618
       0.940885 0.273930 0.629913
       0.945369 0.280622 0.645271
       0.949762 0.287309 0.660687
       0.954066 0.293962 0.676159
       0.958268 0.300578 0.691674
       0.962351 0.307156 0.707233
       0.966311 0.313690 0.722822
       0.970124 0.320157 0.738442
       0.973765 0.326521 0.754069
       0.977205 0.332829 0.769674
       0.980406 0.338983 0.785245
       0.983321 0.344978 0.800727
       0.985909 0.350792 0.816086
       0.988104 0.356351 0.831258
       0.989858 0.361623 0.846170
       0.991098 0.366557 0.860762
       0.991769 0.371076 0.874938
       0.991812 0.375159 0.888611
       0.991172 0.378736 0.901682
       0.989810 0.381756 0.914069
       0.987688 0.384179 0.925676
       0.984795 0.385972 0.936438
       0.981119 0.387127 0.946281
       0.976677 0.387629 0.955180
       0.971500 0.387485 0.963099
       0.965620 0.386714 0.970043
       0.959096 0.385350 0.976034
       0.951977 0.383446 0.981118
       0.944337 0.381026 0.985347
       0.936233 0.378145 0.988792
       0.927723 0.374881 0.991537
       0.918881 0.371274 0.993671
       0.909763 0.367394 0.995271
       0.900405 0.363286 0.996431
       0.890858 0.358990 0.997227
       0.881156 0.354570 0.997729
       0.871325 0.350027 0.997997
       0.861386 0.345402 0.998084
       0.851357 0.340735 0.998033
       0.841257 0.336022 0.997879
       0.831084 0.331287 0.997649
       0.820841 0.326553 0.997364
       0.810538 0.321822 0.997040
       0.800181 0.317095 0.996687
       0.789762 0.312358 0.996305
       0.779283 0.307657 0.995919
       0.768739 0.302971 0.995529
       0.758137 0.298307 0.995135
       0.747460 0.293660 0.994737
       0.736704 0.289048 0.994335
       0.725878 0.284427 0.993929
       0.714962 0.279853 0.993520
       0.703965 0.275294 0.993106
       0.692881 0.270758 0.992686
       0.681685 0.266234 0.992261
       0.670407 0.261734 0.991833
       0.659002 0.257267 0.991402
       0.647493 0.252824 0.990967
       0.635861 0.248419 0.990528
       0.624101 0.243996 0.990086
       0.612191 0.239650 0.989641
       0.600144 0.235331 0.989191
       0.587930 0.231016 0.988735
       0.575561 0.226747 0.988275
       0.563001 0.222502 0.987816
       0.550239 0.218297 0.987353
       0.537282 0.214130 0.986885
       0.524101 0.210003 0.986405
       0.510700 0.205964 0.985912
       0.497074 0.201950 0.985401
       0.483207 0.198061 0.984865
       0.469123 0.194311 0.984293
       0.454803 0.190637 0.983671
       0.440272 0.187189 0.982988
       0.425580 0.183927 0.982222
       0.410775 0.180946 0.981341
       0.395919 0.178352 0.980321
       0.381127 0.176078 0.979121
       0.366521 0.174375 0.977702
       0.352262 0.173189 0.976015
       0.338575 0.172666 0.974015
       0.325675 0.172869 0.971648
       0.313857 0.173868 0.968860
       0.303350 0.175689 0.965603
       0.294471 0.178440 0.961833
       0.287449 0.182005 0.957517
       0.282495 0.186507 0.952617];

case { 'R1'  'RAINBOW' }
descriptorname = 'rainbow_bgyrm_35-85_c69_n256';
description = 'The least worst rainbow colour map I can devise.  Note there are small perceptual blind spots at yellow and red';
map = [0.000000 0.186555 0.962645
       0.000000 0.202474 0.948892
       0.000000 0.217314 0.935162
       0.000000 0.231251 0.921454
       0.011192 0.244416 0.907769
       0.036140 0.257001 0.894099
       0.055678 0.269036 0.880450
       0.069388 0.280559 0.866824
       0.079257 0.291717 0.853222
       0.086705 0.302461 0.839647
       0.092118 0.312895 0.826088
       0.095864 0.323023 0.812557
       0.098322 0.332857 0.799061
       0.099569 0.342411 0.785592
       0.099878 0.351700 0.772158
       0.099398 0.360748 0.758769
       0.098334 0.369568 0.745424
       0.096762 0.378143 0.732142
       0.095150 0.386472 0.718915
       0.093450 0.394578 0.705765
       0.092187 0.402438 0.692700
       0.091506 0.410081 0.679709
       0.091673 0.417471 0.666827
       0.092844 0.424627 0.654053
       0.095394 0.431565 0.641381
       0.099143 0.438266 0.628834
       0.104166 0.444763 0.616395
       0.110368 0.451048 0.604073
       0.117428 0.457142 0.591857
       0.125293 0.463058 0.579746
       0.133667 0.468835 0.567725
       0.142349 0.474458 0.555783
       0.151155 0.479959 0.543930
       0.159927 0.485344 0.532131
       0.168595 0.490665 0.520364
       0.176962 0.495890 0.508647
       0.185000 0.501082 0.496924
       0.192599 0.506214 0.485209
       0.199744 0.511329 0.473481
       0.206472 0.516422 0.461738
       0.212630 0.521505 0.449946
       0.218319 0.526594 0.438106
       0.223499 0.531668 0.426188
       0.228195 0.536761 0.414206
       0.232391 0.541842 0.402124
       0.236143 0.546950 0.389957
       0.239440 0.552064 0.377691
       0.242321 0.557185 0.365302
       0.244805 0.562312 0.352799
       0.246945 0.567440 0.340162
       0.248755 0.572576 0.327380
       0.250235 0.577700 0.314474
       0.251536 0.582828 0.301434
       0.252620 0.587940 0.288280
       0.253608 0.593041 0.275015
       0.254581 0.598096 0.261656
       0.255602 0.603125 0.248249
       0.256812 0.608103 0.234795
       0.258301 0.613005 0.221337
       0.260222 0.617848 0.207945
       0.262662 0.622604 0.194693
       0.265748 0.627264 0.181569
       0.269580 0.631819 0.168769
       0.274215 0.636261 0.156288
       0.279745 0.640574 0.144203
       0.286146 0.644760 0.132651
       0.293465 0.648825 0.121677
       0.301631 0.652749 0.111525
       0.310620 0.656561 0.102116
       0.320300 0.660235 0.093657
       0.330589 0.663803 0.086199
       0.341426 0.667253 0.079852
       0.352683 0.670611 0.074629
       0.364265 0.673870 0.070490
       0.376108 0.677059 0.067449
       0.388103 0.680163 0.065346
       0.400212 0.683221 0.064139
       0.412371 0.686221 0.063672
       0.424524 0.689182 0.063795
       0.436657 0.692099 0.064392
       0.448738 0.694977 0.065356
       0.460735 0.697834 0.066603
       0.472667 0.700671 0.068066
       0.484504 0.703483 0.069670
       0.496254 0.706277 0.071332
       0.507931 0.709054 0.073060
       0.519500 0.711809 0.074792
       0.530991 0.714547 0.076552
       0.542399 0.717285 0.078345
       0.553721 0.719995 0.080170
       0.564973 0.722691 0.082034
       0.576175 0.725387 0.083799
       0.587287 0.728054 0.085523
       0.598352 0.730714 0.087403
       0.609370 0.733361 0.089162
       0.620333 0.735986 0.090964
       0.631243 0.738609 0.092702
       0.642110 0.741207 0.094561
       0.652930 0.743801 0.096272
       0.663728 0.746374 0.098130
       0.674473 0.748933 0.099882
       0.685194 0.751487 0.101673
       0.695880 0.754017 0.103500
       0.706537 0.756534 0.105240
       0.717167 0.759037 0.107085
       0.727763 0.761527 0.108841
       0.738339 0.764004 0.110662
       0.748898 0.766467 0.112456
       0.759436 0.768908 0.114198
       0.769954 0.771339 0.116039
       0.780447 0.773731 0.117807
       0.790912 0.776103 0.119578
       0.801346 0.778436 0.121333
       0.811747 0.780723 0.123100
       0.822102 0.782952 0.124876
       0.832394 0.785113 0.126614
       0.842617 0.787180 0.128339
       0.852741 0.789132 0.130043
       0.862744 0.790937 0.131648
       0.872587 0.792575 0.133222
       0.882234 0.794006 0.134750
       0.891645 0.795182 0.136117
       0.900764 0.796075 0.137499
       0.909542 0.796642 0.138691
       0.917926 0.796843 0.139742
       0.925856 0.796642 0.140735
       0.933283 0.796010 0.141513
       0.940177 0.794922 0.142116
       0.946481 0.793364 0.142555
       0.952185 0.791318 0.142816
       0.957272 0.788805 0.142895
       0.961738 0.785825 0.142796
       0.965601 0.782401 0.142526
       0.968884 0.778563 0.142096
       0.971619 0.774345 0.141528
       0.973850 0.769789 0.140823
       0.975625 0.764936 0.139944
       0.977003 0.759813 0.139019
       0.978027 0.754482 0.138042
       0.978754 0.748953 0.136925
       0.979235 0.743287 0.135752
       0.979510 0.737489 0.134613
       0.979616 0.731601 0.133368
       0.979590 0.725642 0.132121
       0.979457 0.719617 0.130869
       0.979240 0.713547 0.129580
       0.978956 0.707442 0.128297
       0.978620 0.701311 0.126979
       0.978244 0.695158 0.125740
       0.977837 0.688992 0.124422
       0.977396 0.682798 0.123131
       0.976930 0.676600 0.121859
       0.976445 0.670392 0.120607
       0.975943 0.664163 0.119372
       0.975425 0.657931 0.118152
       0.974888 0.651681 0.116883
       0.974335 0.645412 0.115665
       0.973765 0.639140 0.114396
       0.973178 0.632850 0.113254
       0.972574 0.626542 0.112029
       0.971949 0.620228 0.110831
       0.971307 0.613880 0.109664
       0.970649 0.607537 0.108406
       0.969974 0.601158 0.107281
       0.969283 0.594769 0.106128
       0.968577 0.588355 0.104902
       0.967854 0.581929 0.103810
       0.967111 0.575486 0.102663
       0.966350 0.569008 0.101520
       0.965575 0.562526 0.100399
       0.964783 0.555995 0.099296
       0.963976 0.549464 0.098231
       0.963155 0.542899 0.097093
       0.962313 0.536304 0.096016
       0.961457 0.529684 0.095014
       0.960588 0.523034 0.093919
       0.959698 0.516344 0.092832
       0.958794 0.509636 0.091874
       0.957875 0.502885 0.090828
       0.956935 0.496103 0.089807
       0.955987 0.489287 0.088805
       0.955018 0.482438 0.087830
       0.954035 0.475547 0.086854
       0.953037 0.468597 0.085824
       0.952023 0.461618 0.084968
       0.951006 0.454601 0.084163
       0.949974 0.447528 0.083344
       0.948941 0.440440 0.082697
       0.947910 0.433319 0.082226
       0.946884 0.426179 0.081913
       0.945871 0.419041 0.081847
       0.944889 0.411908 0.082109
       0.943940 0.404805 0.082728
       0.943048 0.397784 0.083928
       0.942227 0.390858 0.085608
       0.941494 0.384075 0.088120
       0.940882 0.377503 0.091363
       0.940411 0.371185 0.095416
       0.940102 0.365211 0.100349
       0.939981 0.359638 0.106304
       0.940073 0.354545 0.113159
       0.940396 0.349992 0.120891
       0.940963 0.346044 0.129653
       0.941791 0.342805 0.139220
       0.942881 0.340263 0.149720
       0.944224 0.338473 0.160949
       0.945815 0.337469 0.172917
       0.947642 0.337226 0.185612
       0.949675 0.337722 0.198848
       0.951894 0.338940 0.212690
       0.954272 0.340808 0.227018
       0.956771 0.343280 0.241759
       0.959367 0.346267 0.256857
       0.962023 0.349739 0.272244
       0.964720 0.353586 0.287904
       0.967432 0.357782 0.303759
       0.970124 0.362223 0.319786
       0.972793 0.366884 0.335916
       0.975406 0.371704 0.352141
       0.977964 0.376679 0.368454
       0.980452 0.381728 0.384800
       0.982857 0.386853 0.401211
       0.985177 0.392048 0.417642
       0.987402 0.397274 0.434095
       0.989529 0.402534 0.450593
       0.991552 0.407823 0.467113
       0.993475 0.413137 0.483640
       0.995282 0.418440 0.500200
       0.996983 0.423767 0.516792
       0.998575 0.429107 0.533400
       1.000000 0.434449 0.550052
       1.000000 0.439806 0.566736
       1.000000 0.445169 0.583447
       1.000000 0.450544 0.600206
       1.000000 0.455927 0.617007
       1.000000 0.461325 0.633848
       1.000000 0.466733 0.650727
       1.000000 0.472152 0.667657
       1.000000 0.477574 0.684622
       1.000000 0.483011 0.701639
       1.000000 0.488468 0.718705
       1.000000 0.493928 0.735815
       1.000000 0.499394 0.752973
       1.000000 0.504887 0.770184
       1.000000 0.510380 0.787441
       1.000000 0.515883 0.804745
       1.000000 0.521408 0.822100
       1.000000 0.526950 0.839502
       1.000000 0.532492 0.856949
       1.000000 0.538050 0.874449
       1.000000 0.543612 0.892002
       1.000000 0.549195 0.909598
       0.998480 0.554784 0.927242
       0.996596 0.560390 0.944944
       0.994532 0.566003 0.962686
       0.992279 0.571631 0.980481];

case { 'R2'  'RAINBOW2' }
descriptorname = 'rainbow_bgyr_35-85_c72_n256';
description = 'Reasonable rainbow colour map from blue to red.  Note there is a small perceptual blind spot at yellow';
map = [0.000000 0.203873 0.962505
       0.000000 0.215239 0.951401
       0.000000 0.226130 0.940312
       0.000000 0.236541 0.929229
       0.000000 0.246539 0.918169
       0.000000 0.256203 0.907120
       0.000000 0.265574 0.896082
       0.000000 0.274646 0.885058
       0.000000 0.283483 0.874045
       0.000000 0.292086 0.863051
       0.000000 0.300474 0.852064
       0.000000 0.308697 0.841088
       0.000000 0.316722 0.830126
       0.000000 0.324583 0.819168
       0.000000 0.332324 0.808231
       0.000000 0.339905 0.797297
       0.000000 0.347361 0.786382
       0.000000 0.354696 0.775463
       0.000000 0.361910 0.764564
       0.000000 0.369018 0.753670
       0.000000 0.376017 0.742795
       0.000000 0.382910 0.731928
       0.000000 0.389688 0.721075
       0.000000 0.396363 0.710256
       0.000000 0.402916 0.699459
       0.000000 0.409338 0.688715
       0.000000 0.415611 0.678016
       0.000000 0.421721 0.667385
       0.000000 0.427676 0.656839
       0.000000 0.433415 0.646385
       0.000000 0.438964 0.636050
       0.000000 0.444319 0.625832
       0.000000 0.449447 0.615746
       0.000000 0.454378 0.605790
       0.000000 0.459111 0.595971
       0.004338 0.463666 0.586271
       0.029615 0.468070 0.576680
       0.055795 0.472354 0.567173
       0.077065 0.476524 0.557739
       0.095292 0.480614 0.548369
       0.111187 0.484648 0.539027
       0.125303 0.488652 0.529714
       0.137988 0.492620 0.520402
       0.149372 0.496582 0.511097
       0.159631 0.500548 0.501787
       0.169000 0.504521 0.492444
       0.177466 0.508492 0.483085
       0.185172 0.512463 0.473695
       0.192171 0.516453 0.464289
       0.198564 0.520457 0.454827
       0.204434 0.524478 0.445315
       0.209743 0.528510 0.435773
       0.214606 0.532552 0.426161
       0.219048 0.536606 0.416512
       0.223088 0.540663 0.406791
       0.226742 0.544738 0.396996
       0.230020 0.548829 0.387135
       0.232998 0.552915 0.377201
       0.235675 0.557030 0.367159
       0.238015 0.561136 0.357035
       0.240060 0.565256 0.346782
       0.241853 0.569393 0.336400
       0.243339 0.573542 0.325884
       0.244577 0.577688 0.315230
       0.245562 0.581851 0.304391
       0.246300 0.586027 0.293358
       0.246799 0.590194 0.282138
       0.247074 0.594379 0.270674
       0.247143 0.598560 0.258956
       0.247035 0.602746 0.246963
       0.246790 0.606929 0.234727
       0.246470 0.611094 0.222163
       0.246155 0.615232 0.209355
       0.245952 0.619356 0.196319
       0.245995 0.623422 0.183036
       0.246445 0.627422 0.169693
       0.247481 0.631352 0.156300
       0.249253 0.635178 0.142994
       0.251961 0.638897 0.130010
       0.255696 0.642489 0.117414
       0.260573 0.645938 0.105575
       0.266592 0.649255 0.094696
       0.273717 0.652417 0.084904
       0.281824 0.655447 0.076489
       0.290780 0.658349 0.069753
       0.300426 0.661134 0.064513
       0.310611 0.663825 0.060865
       0.321123 0.666420 0.058721
       0.331861 0.668960 0.057692
       0.342715 0.671437 0.057693
       0.353565 0.673877 0.058443
       0.364386 0.676284 0.059738
       0.375123 0.678664 0.061142
       0.385749 0.681022 0.062974
       0.396267 0.683375 0.064759
       0.406655 0.685711 0.066664
       0.416923 0.688033 0.068644
       0.427067 0.690343 0.070512
       0.437095 0.692656 0.072423
       0.447010 0.694942 0.074359
       0.456832 0.697229 0.076211
       0.466567 0.699505 0.078090
       0.476213 0.701772 0.079998
       0.485774 0.704034 0.081943
       0.495271 0.706288 0.083778
       0.504697 0.708529 0.085565
       0.514055 0.710760 0.087502
       0.523348 0.712977 0.089316
       0.532587 0.715195 0.091171
       0.541761 0.717404 0.092931
       0.550901 0.719596 0.094839
       0.559990 0.721776 0.096566
       0.569022 0.723955 0.098445
       0.578019 0.726127 0.100225
       0.586980 0.728282 0.102035
       0.595902 0.730435 0.103855
       0.604790 0.732578 0.105640
       0.613634 0.734709 0.107440
       0.622456 0.736828 0.109246
       0.631250 0.738946 0.111025
       0.640011 0.741042 0.112818
       0.648742 0.743145 0.114525
       0.657454 0.745226 0.116361
       0.666130 0.747307 0.118133
       0.674791 0.749367 0.119864
       0.683430 0.751435 0.121611
       0.692053 0.753477 0.123379
       0.700648 0.755521 0.125168
       0.709231 0.757551 0.126907
       0.717792 0.759568 0.128678
       0.726335 0.761583 0.130479
       0.734867 0.763586 0.132206
       0.743382 0.765586 0.133961
       0.751882 0.767564 0.135676
       0.760367 0.769541 0.137469
       0.768838 0.771513 0.139171
       0.777308 0.773464 0.140965
       0.785757 0.775413 0.142689
       0.794199 0.777358 0.144438
       0.802624 0.779283 0.146171
       0.811047 0.781205 0.147914
       0.819455 0.783114 0.149672
       0.827860 0.785020 0.151378
       0.836256 0.786914 0.153109
       0.844650 0.788797 0.154858
       0.853036 0.790662 0.156615
       0.861415 0.792515 0.158345
       0.869782 0.794342 0.160020
       0.878140 0.796125 0.161782
       0.886474 0.797859 0.163460
       0.894770 0.799521 0.165067
       0.903007 0.801060 0.166695
       0.911155 0.802450 0.168191
       0.919171 0.803636 0.169639
       0.927005 0.804555 0.170896
       0.934588 0.805135 0.171997
       0.941846 0.805316 0.172894
       0.948695 0.805036 0.173545
       0.955064 0.804241 0.173921
       0.960881 0.802887 0.173992
       0.966095 0.800973 0.173747
       0.970687 0.798499 0.173191
       0.974650 0.795494 0.172336
       0.978013 0.792008 0.171212
       0.980823 0.788098 0.169857
       0.983139 0.783838 0.168251
       0.985042 0.779280 0.166516
       0.986600 0.774500 0.164627
       0.987886 0.769553 0.162650
       0.988970 0.764486 0.160560
       0.989900 0.759324 0.158484
       0.990719 0.754111 0.156344
       0.991460 0.748847 0.154141
       0.992144 0.743562 0.151958
       0.992789 0.738248 0.149806
       0.993396 0.732926 0.147601
       0.993978 0.727586 0.145427
       0.994541 0.722236 0.143205
       0.995087 0.716888 0.141029
       0.995616 0.711519 0.138803
       0.996126 0.706139 0.136595
       0.996620 0.700748 0.134440
       0.997096 0.695344 0.132228
       0.997555 0.689930 0.130058
       0.997997 0.684498 0.127831
       0.998421 0.679060 0.125641
       0.998829 0.673610 0.123396
       0.999219 0.668147 0.121195
       0.999590 0.662666 0.119036
       0.999944 0.657170 0.116820
       1.000000 0.651659 0.114580
       1.000000 0.646127 0.112437
       1.000000 0.640588 0.110240
       1.000000 0.635028 0.107973
       1.000000 0.629450 0.105798
       1.000000 0.623864 0.103599
       1.000000 0.618249 0.101348
       1.000000 0.612613 0.099135
       1.000000 0.606972 0.096882
       1.000000 0.601295 0.094743
       1.000000 0.595606 0.092465
       1.000000 0.589890 0.090257
       1.000000 0.584160 0.088032
       1.000000 0.578400 0.085726
       1.000000 0.572627 0.083542
       1.000000 0.566822 0.081316
       1.000000 0.560983 0.079004
       1.000000 0.555126 0.076745
       1.000000 0.549247 0.074530
       1.000000 0.543332 0.072245
       1.000000 0.537393 0.070004
       1.000000 0.531413 0.067732
       1.000000 0.525408 0.065424
       1.000000 0.519370 0.063180
       1.000000 0.513296 0.060810
       1.000000 0.507185 0.058502
       1.000000 0.501042 0.056232
       1.000000 0.494857 0.053826
       1.000000 0.488627 0.051494
       1.000000 0.482359 0.049242
       1.000000 0.476046 0.046828
       1.000000 0.469692 0.044447
       1.000000 0.463268 0.042093
       1.000000 0.456815 0.039648
       1.000000 0.450307 0.037261
       1.000000 0.443742 0.034882
       1.000000 0.437117 0.032495
       1.000000 0.430426 0.030303
       1.000000 0.423666 0.028180
       1.000000 0.416857 0.026121
       1.000000 0.409966 0.024126
       1.000000 0.402993 0.022194
       1.000000 0.395949 0.020325
       1.000000 0.388816 0.018517
       0.999937 0.381591 0.016771
       0.999612 0.374278 0.015085
       0.999273 0.366853 0.013457
       0.998920 0.359319 0.011916
       0.998553 0.351672 0.010169
       0.998173 0.343900 0.008744
       0.997780 0.336002 0.007354
       0.997376 0.327956 0.006020
       0.996958 0.319762 0.004743
       0.996528 0.311383 0.003522
       0.996086 0.302821 0.002356
       0.995632 0.294070 0.001245
       0.995165 0.285096 0.000187
       0.994686 0.275912 0.000000
       0.994195 0.266419 0.000000
       0.993693 0.256636 0.000000
       0.993178 0.246521 0.000000
       0.992649 0.236050 0.000000
       0.992106 0.225108 0.000000
       0.991553 0.213705 0.000000
       0.990988 0.201692 0.000000
       0.990411 0.189028 0.000000];

case { 'R3'  'RAINBOW3' }
descriptorname = 'diverging-rainbow_bgymr_45-85_c67_n256';
description = 'Diverging-rainbow colourmap. Yellow is the central reference colour. The blue and red end points are matched in lightness and chroma';
map = [0.032081 0.361818 0.973476
       0.058360 0.370921 0.958455
       0.075961 0.379844 0.943453
       0.088784 0.388582 0.928471
       0.098335 0.397136 0.913519
       0.105384 0.405552 0.898579
       0.110498 0.413803 0.883666
       0.113804 0.421912 0.868770
       0.115569 0.429905 0.853881
       0.115925 0.437769 0.839018
       0.114869 0.445489 0.824162
       0.112656 0.453108 0.809318
       0.109175 0.460590 0.794469
       0.104545 0.467961 0.779610
       0.099069 0.475206 0.764736
       0.092835 0.482295 0.749820
       0.086330 0.489228 0.734863
       0.080053 0.496001 0.719825
       0.074782 0.502594 0.704688
       0.071457 0.508994 0.689437
       0.070850 0.515169 0.674030
       0.073695 0.521132 0.658462
       0.079835 0.526880 0.642713
       0.089012 0.532399 0.626765
       0.100260 0.537716 0.610623
       0.112876 0.542839 0.594280
       0.126002 0.547793 0.577744
       0.139098 0.552610 0.561037
       0.151827 0.557316 0.544167
       0.163878 0.561929 0.527144
       0.174975 0.566476 0.509970
       0.185078 0.570978 0.492662
       0.194117 0.575468 0.475242
       0.202038 0.579943 0.457659
       0.208951 0.584410 0.439954
       0.214835 0.588880 0.422096
       0.219794 0.593370 0.404111
       0.223837 0.597853 0.385966
       0.227109 0.602342 0.367691
       0.229637 0.606831 0.349303
       0.231627 0.611301 0.330812
       0.233107 0.615739 0.312296
       0.234366 0.620145 0.293851
       0.235532 0.624477 0.275594
       0.236859 0.628729 0.257607
       0.238557 0.632872 0.240116
       0.240938 0.636893 0.223328
       0.244180 0.640760 0.207400
       0.248503 0.644469 0.192557
       0.253926 0.648012 0.179018
       0.260516 0.651390 0.166938
       0.268210 0.654588 0.156415
       0.276904 0.657645 0.147487
       0.286384 0.660543 0.140122
       0.296536 0.663333 0.134288
       0.307117 0.666005 0.129714
       0.317998 0.668600 0.126226
       0.329039 0.671124 0.123560
       0.340113 0.673597 0.121585
       0.351161 0.676031 0.120101
       0.362123 0.678435 0.118954
       0.372963 0.680813 0.117981
       0.383688 0.683184 0.117106
       0.394258 0.685538 0.116375
       0.404694 0.687877 0.115628
       0.415001 0.690206 0.114863
       0.425178 0.692536 0.114138
       0.435230 0.694841 0.113445
       0.445159 0.697149 0.112685
       0.454996 0.699445 0.111904
       0.464730 0.701733 0.111122
       0.474371 0.704018 0.110337
       0.483922 0.706295 0.109542
       0.493403 0.708560 0.108658
       0.502809 0.710815 0.107827
       0.512149 0.713058 0.107028
       0.521421 0.715303 0.106166
       0.530636 0.717537 0.105234
       0.539785 0.719757 0.104341
       0.548895 0.721965 0.103492
       0.557936 0.724175 0.102546
       0.566945 0.726374 0.101601
       0.575904 0.728562 0.100647
       0.584812 0.730744 0.099684
```
