## Supplementary material for "Adaptive-Resolution Multi-Orientation Analysis of Complex Filamentous Network Images": MATLAB code from Github: partitions.m

```
function plist = partitions(total_sum,candidate_set,max_count,fixed_count)
% extracts the list of all partitions of a number as integer sums of a list of candidates
% usage: plist = partitions(total_sum,candidate_set)
% usage: plist = partitions(total_sum,candidate_set,max_count,fixed_count)
%
% PARTITIONS solves the money changing problem. E.g.,
% how can you make change for one dollar given coins
% of a given set of denominations. A good reference on
% the general problem is found here:
%
% http://en.wikipedia.org/wiki/Integer_partition
%
% PARTITIONS uses a recursive strategy to enumerate all
% possible partitions of the total_sum. This may be
% highly intensive for large sums or large sets of
% candidates.
%
% arguments: (input)
%  total_sum - scalar positive integer (to be partitioned)
%
%              BEWARE! a large total_sum can easily cause
%              stack problems. For example, the number of
%              partitions of 40 is 37338, a set that took 24
%              seconds to completely enumerate on my cpu.
%
%  candidate_set - (OPTIONAL) vector of (distinct) candidate
%              positive integers for the partitions.
%
%              Efficiency considerations force me to require
%              that the candidates be sorted in non-decreasing
%              order. An error is produced otherwise.
%
%              DEFAULT: candidate_set = 1:total_sum
%
%              BEWARE! large candidate sets can easily cause
%              stack problems
%
%  max_count - (OPTIONAL) the maximum quantity of any
%              candidate in the final sum.
%
%              max_count must be either a vector of the
%              same length as candidate_set, or a scalar
%              that applies to all elements in that set.
%
%              DEFAULT = floor(total_sum./candidate_set)
%
%  fixed_count - (OPTIONAL) Allows you to specify a fixed
%              number of terms in the partitioned sum.
%
%              fixed_count must be a positive integer if
%              supplied.
%
%              DEFAULT = []
%
% arguments: (output)
%  plist - array of partitions of total_sum. This is a list
%              of the quantity of each element such that
%              plist*candidate_set(:) yields total_sum
%
%
% example: Write 9 as an integer combination of the set [1 2 4 7]
%
%  partitions(9,[1 2 4 7])
%
% ans =
%    9     0     0     0
%    7     1     0     0
%    5     2     0     0
%    3     3     0     0
%    1     4     0     0
%    5     0     1     0
%    3     1     1     0
%    1     2     1     0
%    1     0     2     0
%    2     0     0     1
%    0     1     0     1
%
% Thus, we can write 9 = 9*1
% or 9 = 1*1 + 4*2
% or 9 = 1*2 + 1*7
% or any of 8 distinct other ways.
%
% There are 11 such ways to write 9 in terms of these
% candidates.
%
%
% example: Change a 1 dollar bill (100 cents) as an integer
%  combination of the set [1 5 10 25 50], using no more than
%  4 of any one coin denomination. Note that no pennies will
%  be allowed by the maximum constraint.
%
%  partitions(100,[1 5 10 25 50],4)
%
% ans =
%    0     4     3     2     0
%    0     2     4     2     0
%    0     3     1     3     0
%    0     1     2     3     0
%    0     0     0     4     0
%    0     4     3     0     1
%    0     2     4     0     1
%    0     3     1     1     1
%    0     1     2     1     1
%    0     0     0     2     1
%    0     0     0     0     2
%
% example: Write 13 as an integer combination of the set [2 4 6 8 10 12]
%  (Note that no such combination exists.)
%
%  partitions(13,[2 4 6 8 10 12])
%
% ans =
%   Empty matrix: 0-by-6
%
%
% example: find all possible ways to write 100 as a sum of EXACTLY 4
%  squares of the integers 1:9.
%
%  partitions(100,(1:9).^2,[],4)
% ans =
%   0    0    0    0    4    0    0    0    0
%   1    0    0    0    2    0    1    0    0
%   2    0    0    0    0    0    2    0    0
%   0    1    0    2    0    0    0    1    0
%   1    0    2    0    0    0    0    0    1
%
%
% Author: John D'Errico
%
% Release: 2
% Release date: 7/15/08

% default for candidate_set
if (nargin<2) || isempty(candidate_set)
  candidate_set = 1:total_sum;
end

% how many candidates are there
n = length(candidate_set);

% error checks
if any(candidate_set<=0)
  error('All members of candidate_set must be > 0')
end
% candidates must be sorted in increasng order
if any(diff(candidate_set)<0)
  error('Efficiency requires that candidate_set be sorted')
end

% check for a max_count. do we supply a default?
if (nargin<3) || isempty(max_count)
  % how high do we need look?
  max_count = floor(total_sum./candidate_set);
elseif length(max_count)==1
  % if a scalar was provided, then turn it into a vector
  max_count = repmat(max_count,1,n);
end

% check for a fixed_count
if (nargin<4) || isempty(fixed_count)
  fixed_count = [];
elseif (fixed_count<0) || (fixed_count~=round(fixed_count))
  error('fixed_count must be a positive integer if supplied')
end

% check for degenerate cases
if isempty(fixed_count)
  if total_sum == 0
    plist = zeros(1,n);
    return
  elseif (n == 0)
    plist = [];
    return
  elseif (n == 1)
    % only one element in the set. can we form
    % total_sum from it as an integer multiple?
    p = total_sum/candidate_set;
    if (p==fix(p)) && (p<=max_count)
      plist = p;
    else
      plist = [];
    end
    return
 end
else
  % there was a fixed_count supplied
  if (total_sum == 0) && (fixed_count == 0)
    plist = zeros(1,n);
    return
  elseif (n == 0) || (fixed_count <= 0)
    plist = [];
    return
  elseif (n==1)
    % there must be a non-zero fixed_count, since
    % we did not trip the last test. since there
    % is only one candidate in the set, will it work?
    if (fixed_count*candidate_set) == total_sum
      plist = fixed_count;
    else
      plist = [];
    end
    return
  end
end

% finally, we can do some work. start with the
% largest element and work backwards
m = max_count(end);
% do we need to back off on m?
c = candidate_set(end);
m = min([m,floor(total_sum/c),fixed_count]);

plist = zeros(0,n);
for i = 0:m
  temp = partitions(total_sum - i*c, ...
      candidate_set(1:(end-1)), ...
      max_count(1:(end-1)),fixed_count-i);
  plist = [plist;[temp,repmat(i,size(temp,1),1)]];  %#ok
end
```
