## Supplementary material for "Adaptive-Resolution Multi-Orientation Analysis of Complex Filamentous Network Images": MATLAB code from Github: nonLocalMaximaSuppression.m

```
function [ nlms ] = nonLocalMaximaSuppression( rotationResponse, theta , suppressionValue)
%nonLocalMaximaSuppression Suppress pixels which are not local maxima in
%both orientation and filter response
%
% INPUT
% rotationResponse: matrix Y by X by R, R = # of rotation angles
%             (4th output of steerableDetector or steerableVanGinkelFilter)
% theta: (optional) rotation angles. Default: (0:R-1)*pi/R
% suppressionValue: (optional) value to give suppressed pixels (def: 0)
%
% OUTPUT
% nlms: rotationResponse with non-local maxima set to suppressedValue
%
% Copyright (C) 2019, Jaqaman Lab - UT Southwestern, Goldman Lab - Northwestern 
%
% This file is part of AdaptiveResolutionOrientationSpace.
% 
% AdaptiveResolutionOrientationSpace is free software: you can redistribute it and/or modify
% it under the terms of the GNU General Public License as published by
% the Free Software Foundation, either version 3 of the License, or
% (at your option) any later version.
% 
% AdaptiveResolutionOrientationSpace is distributed in the hope that it will be useful,
% but WITHOUT ANY WARRANTY; without even the implied warranty of
% MERCHANTABILITY or FITNESS FOR A PARTICULAR PURPOSE.  See the
% GNU General Public License for more details.
% 
% You should have received a copy of the GNU General Public License
% along with AdaptiveResolutionOrientationSpace.  If not, see <http://www.gnu.org/licenses/>.
% 
% 

% Mark Kittisopikul, 2015
% UT Southwestern

% TODO: full-backwards compatability with nonMaximumSuppression?

nO = size(rotationResponse,3);

if(nargin < 2 || isempty(theta))
    % default value is the rotation planes correspond to angles that
    % equally divide pi
    theta = 0:nO-1;
    theta = theta*pi/nO;
    %TODO: allow for theta of different sizes
end
if(nargin < 3)
    suppressionValue = 0;
end
% if(nargin < 4)
%     distance = 1;
% end

nlms = rotationResponse;

% TODO: optimize later
for o = 1:nO
    nlms(:,:,o) = nonMaximumSuppression(rotationResponse(:,:,o),theta(o));
end
% suppression response if less than the previous or next orientation
% response
% nlms(nlms < nlms(:,:,[2:end 1]) | nlms < nlms(:,:,[end 1:end-1])) = suppressionValue;
nlms(rotationResponse < rotationResponse(:,:,[2:end 1]) | rotationResponse < rotationResponse(:,:,[end 1:end-1])) = suppressionValue;
% what if equal on either side or both sides?

end % end of function
```
