## Supplementary material for "Adaptive-Resolution Multi-Orientation Analysis of Complex Filamentous Network Images": MATLAB code from Github: nonLocalMaximaSuppressionPrecise.m

```
function [ nlms, offset, X, Y ] = nonLocalMaximaSuppressionPrecise( rotationResponse, theta , suppressionValue, interpMethod, mask, offsetAngle, angleMultiplier)
%nonLocalMaximaSuppression Suppress pixels which are not local maxima in
%both orientation and filter response
%
% INPUT
% rotationResponse: matrix Y by X by R, R = # of rotation angles
%             (4th output of steerableDetector or steerableVanGinkelFilter)
% theta: (optional) rotation angles. Default: (0:R-1)*pi/R
% suppressionValue: (optional) value to give suppressed pixels (def: 0)
% interpMethod: (optional) see interp3, default: cubic
% mask: (optional) 
%
% OUTPUT
% nlms: rotationResponse with non-local maxima set to suppressedValue
% offset: offset from center of pixel for sub-pixel localization
% X: x-coordinate of sub-pixel localization
% Y: y-coordinate of sub-pixel localization
%
% Copyright (C) 2019, Jaqaman Lab - UT Southwestern, Goldman Lab - Northwestern 
%
% This file is part of AdaptiveResolutionOrientationSpace.
% 
% AdaptiveResolutionOrientationSpace is free software: you can redistribute it and/or modify
% it under the terms of the GNU General Public License as published by
% the Free Software Foundation, either version 3 of the License, or
% (at your option) any later version.
% 
% AdaptiveResolutionOrientationSpace is distributed in the hope that it will be useful,
% but WITHOUT ANY WARRANTY; without even the implied warranty of
% MERCHANTABILITY or FITNESS FOR A PARTICULAR PURPOSE.  See the
% GNU General Public License for more details.
% 
% You should have received a copy of the GNU General Public License
% along with AdaptiveResolutionOrientationSpace.  If not, see <http://www.gnu.org/licenses/>.
% 
% 

% Mark Kittisopikul, 2015
% UT Southwestern

% Sub-pixel localization added October 2017
% Mark Kittisopikul
% Northwestern

% TODO: full-backwards compatability with nonMaximumSuppression?

nO = size(theta,3);

if(nargin < 2 || isempty(theta))
    % default value is the rotation planes correspond to angles that
    % equally divide pi
    theta = 0:nO-1;
    theta = theta*pi/nO;
    %TODO: allow for theta of different sizes
end
if(nargin < 3 || isempty(suppressionValue))
    suppressionValue = 0;
end
if(nargin < 4 || isempty(interpMethod))
    interpMethod = 'cubic';
end
if(nargin < 5)
    mask = [];
else
% mask should be dilated beyond the minimal area    
    mask = imdilate(mask,strel('disk',3));
end
% if(nargin < 4)
%     distance = 1;
% end
if(nargin < 6 || isempty(offsetAngle))
    offsetAngle = theta;
end
if(nargin < 7 || isempty(angleMultiplier))
    angleMultiplier = 3;
end

% nlms = theta;
rotationResponseSize = size(rotationResponse);
if(ndims(rotationResponse) < 3)
    rotationResponseSize(3) = 1;
end
ny = rotationResponseSize(1);
nx = rotationResponseSize(2);
nAngles = rotationResponseSize(3);
period = nAngles*angleMultiplier;

% See Boyd, Chebyshev and Fourier Spectral Methods, Second Edition
% (Revised), Dover, 2001. Toronto. ISBN 0-486-41183-4, page 198
if(isempty(mask))
    if(angleMultiplier ~= 1)
        rotationResponse = interpft(rotationResponse,period,3);
    end
else
    rotationResponse = shiftdim(rotationResponse,2);
    rotationResponseTemp = interpft(rotationResponse(:,mask),period);
    rotationResponse = NaN([period size(mask)]);
    rotationResponse(:,mask) = rotationResponseTemp;
    rotationResponse = shiftdim(rotationResponse,1);
    clear rotationResponseTemp;
end
rotationResponse = padarrayXT(rotationResponse,[1 1 0] ,'symmetric');
if(angleMultiplier ~= 1)
    rotationResponse = padarrayXT(rotationResponse,[0 0 1] ,'circular');
end

% Map angles in radians (0:2*pi) to angle index (1:nAngles)
% angleIdx = theta/(2*pi)*(nAngles*3-1)+1;

% Map angles in from [-pi/2:pi/2) to (0:pi]
angleIdx = theta;
if(angleMultiplier ~= 1)
    angleIdx(angleIdx < 0) = angleIdx(angleIdx < 0) + pi; % now (0:pi)
    % Map angles in radians (0:pi) to angle index (2:nAngles*3+1)
    angleIdx = angleIdx/pi*(period)+2;
    % Negative values should be advanced by one period
    % angleIdx(angleIdx < 0) = angleIdx(angleIdx < 0)+period;
end

% Offset by 1 due to padding
[x,y] = meshgrid(2:nx+1,2:ny+1);

x_offset = cos(offsetAngle);
y_offset = sin(offsetAngle);

Xplus = bsxfun(@plus,x,x_offset);
Yplus = bsxfun(@plus,y,y_offset);

Xminus = bsxfun(@minus,x,x_offset);
Yminus = bsxfun(@minus,y,y_offset);

if(nargout > 1)

% Extra Chebfun points
m = sqrt(2)/2;
XplusCheb = bsxfun(@plus,x,x_offset.*m);
YplusCheb = bsxfun(@plus,y,y_offset.*m);

XminusCheb = bsxfun(@minus,x,x_offset.*m);
YminusCheb = bsxfun(@minus,y,y_offset.*m);

end

x = cat(4,Xminus,repmat(x,[1 1 nO]),Xplus);
y = cat(4,Yminus,repmat(y,[1 1 nO]),Yplus);
% if(angleMultiplier ~= 1)
    angleIdx = repmat(angleIdx,[1 1 1 3]);
% end

if(nargout > 1)

% Extra Chebfun points
x = cat(4,x,XplusCheb,XminusCheb);
y = cat(4,y,YplusCheb,YminusCheb);
%     if(angleMultiplier ~= 1)
        angleIdx(:,:,:,4:5) = angleIdx(:,:,:,1:2);
%     end

clear XplusCheb YplusCheb XminusCheb YminusCheb

end


clear Xplus Yplus Xminus Yminus x_offset y_offset;

if(angleMultiplier ~= 1)
    A = interp3(rotationResponse,x,y,angleIdx,interpMethod,0);
else
    %% Use Fourier interpolation
    A = zeros(size(x));
%     parfor d=1:size(x,4)
    parfor j=1:size(x,4)*nO
        G = zeros(rotationResponseSize);
%         for o=1:nO
            for a=1:nAngles
                G(:,:,a) = interp2(rotationResponse(:,:,a),squeeze(x(:,:,j)),squeeze(y(:,:,j)),interpMethod,0);
            end
            A(:,:,j) = interpft1([0 pi],shiftdim(G,2),shiftdim(angleIdx(:,:,j),-1));
%         end
    end
end

nlms = A(:,:,:,2);
nlms(nlms < A(:,:,:,1) | nlms < A(:,:,:,3)) = suppressionValue;

% TODO: optimize later
% for o = 1:nO
% %     nlms(:,:,o) = nonMaximumSuppression(rotationResponse(:,:,o),theta(o));
% 
% %     res = padarrayXT(rrotationResponse(:,:,o), [1 1 0], 'symmetric');
% 
%     x_offset = cos(theta(:,:,o));
%     y_offset = sin(theta(:,:,o));
% 
%     % +1 interp
%     A1 = interp3(rotationResponse, x+x_offset, y+y_offset,angleIdx(:,:,o),interpMethod,0);
% 
%     % -1 interp
%     A2 = interp3(rotationResponse, x-x_offset, y-y_offset,angleIdx(:,:,o),interpMethod,0);
% 
%     % TODO: We only need to interpolate where not NaN
%     temp = interp3(rotationResponse, x, y, angleIdx(:,:,o),interpMethod,0);
%     temp(temp < A1 | temp < A2) = suppressionValue;
%     nlms(:,:,o) = temp;
% 
% %     res(res<A1 | res<A2) = 0;
% end

%% since the input theta already should represent the local maxima in
%% in orientation, we no longer need to suppress in the orientation dimension
% suppression response if less than the previous or next orientation
% response
% nlms(nlms < nlms(:,:,[2:end 1]) | nlms < nlms(:,:,[end 1:end-1])) = suppressionValue;
% nlms(rotationResponse < rotationResponse(:,:,[2:end 1]) | rotationResponse < rotationResponse(:,:,[end 1:end-1])) = suppressionValue;
% what if equal on either side or both sides?

if(nargout > 1)
    % Calculate sub-pixel offset
    notSuppressed = nlms ~= suppressionValue & ~isnan(nlms);
    
    % Use extra points in order
    A = A(:,:,:,[1 5 2 4 3]);
    
    A = reshape(A,nx*ny*nO,size(A,4));
    
%     A = A(notSuppressed,[3 2 1 2]);
    A = A(notSuppressed,[5 4 3 2 1 2 3 4]);

    nS_offset = interpft_extrema(A,2,true);
    nS_offset = nS_offset(:,1);
    nS_offset = cos(nS_offset);
    offset = NaN(size(nlms));
    offset(notSuppressed) = nS_offset;
end

if(nargout > 2)

    [Xp,Yp] = meshgrid(1:size(nlms,2),1:size(nlms,1));

    % Get sub-pixel NLMS points
    X = joinColumns(Xp+cos(theta).*offset);
    Y = joinColumns(Yp+sin(theta).*offset);

end

end % end of function
```
