## Supplementary material for "Adaptive-Resolution Multi-Orientation Analysis of Complex Filamentous Network Images": MATLAB code from Github: OrientationSpaceEdgeFilter.m

```
classdef OrientationSpaceEdgeFilter < OrientationSpaceFilter
    %OrientationSpaceRidgeFilter OrientationSpace filter that does edge
    %filtering only
%
% Copyright (C) 2019, Jaqaman Lab - UT Southwestern, Goldman Lab - Northwestern 
%
% This file is part of AdaptiveResolutionOrientationSpace.
% 
% AdaptiveResolutionOrientationSpace is free software: you can redistribute it and/or modify
% it under the terms of the GNU General Public License as published by
% the Free Software Foundation, either version 3 of the License, or
% (at your option) any later version.
% 
% AdaptiveResolutionOrientationSpace is distributed in the hope that it will be useful,
% but WITHOUT ANY WARRANTY; without even the implied warranty of
% MERCHANTABILITY or FITNESS FOR A PARTICULAR PURPOSE.  See the
% GNU General Public License for more details.
% 
% You should have received a copy of the GNU General Public License
% along with AdaptiveResolutionOrientationSpace.  If not, see <http://www.gnu.org/licenses/>.
% 
% 
    
    properties
    end
    
    methods
        function obj = OrientationSpaceEdgeFilter(f_c,b_f,K,varargin)
            obj@OrientationSpaceFilter(f_c,b_f,K,varargin{:});
        end
        function R = getResponse(obj,I)
            If = fft2(I);
            angularResponse = obj.applyEdgeFilter(If);
            ns = [0 cumsum([obj.n])];
            R(numel(obj)) = OrientationSpaceResponse;
            for o=1:numel(obj)
                R(o) = OrientationSpaceResponse(obj(o),angularResponse(:,:,ns(o)+1:ns(o+1)));
            end
            R = reshape(R,size(obj));
        end
    end
    
end
```
