## Supplementary material for "Adaptive-Resolution Multi-Orientation Analysis of Complex Filamentous Network Images": MATLAB code from Github: OrientationSpaceFilter.m

```
classdef OrientationSpaceFilter < handle & matlab.mixin.Heterogeneous
    %OrientationSpaceFilter is a class object that represents a polar
    %separable frequency domain filter
    %
    % Filter object used by
    % steerableAdaptiveResolutionOrientationSpaceDetector.m
    % 
    % f_c: maximum frequency for the radial filter
    % b_f: frequency bandwidth for the radial filter
    % K: number of rotation angles through 360 degrees
%
% Copyright (C) 2019, Jaqaman Lab - UT Southwestern, Goldman Lab - Northwestern 
%
% This file is part of AdaptiveResolutionOrientationSpace.
% 
% AdaptiveResolutionOrientationSpace is free software: you can redistribute it and/or modify
% it under the terms of the GNU General Public License as published by
% the Free Software Foundation, either version 3 of the License, or
% (at your option) any later version.
% 
% AdaptiveResolutionOrientationSpace is distributed in the hope that it will be useful,
% but WITHOUT ANY WARRANTY; without even the implied warranty of
% MERCHANTABILITY or FITNESS FOR A PARTICULAR PURPOSE.  See the
% GNU General Public License for more details.
% 
% You should have received a copy of the GNU General Public License
% along with AdaptiveResolutionOrientationSpace.  If not, see <http://www.gnu.org/licenses/>.
% 
% 

    % Mark Kittisopikul, August 22nd, 2015
    % Jaqaman Lab
    % UT Southwestern
    
    properties (SetAccess = immutable)
        % radial central frequency
        f_c
        % radial frequency bandwidth
        b_f
        % angular order
        K
        % normilization setting
        normEnergy
        % number of angular filter templates
        n
        % Sample factor, multiplier to calculate n from K
        sampleFactor = 1
    end
    properties (SetAccess = protected, Dependent)
        % basis angles
        angles
    end
    
    properties (Transient)
        % filter size, should correspond with image size
        size
        % Filter itself
        F
        % Angular gaussians useful for manipulating response
        angularGaussians
    end
    
    properties (Dependent = true)
        % f_c
        centralFrequency
        % b_f
        frequencyBandwidth
        % K
        order
    end
    
    methods
        function obj = OrientationSpaceFilter(f_c,b_f,K,normEnergy)
            if(nargin == 0)
                return;
            end
            if(~isscalar(f_c) || ~isscalar(b_f) || ~isscalar(K))
                s = [length(f_c) length(b_f) length(K)];
                s(2) = max(1,s(2));
                f_c = repmat(f_c(:),1,s(2),s(3));
                if(isempty(b_f))
                    b_f = 1/sqrt(2) * f_c;
                else
                    b_f = repmat(b_f(:)',s(1),1,s(3));
                end
                K   = repmat(shiftdim(K(:),-2),s(1),s(2),1);
                if(nargin < 4)
                        normEnergy = [];
                end
                normEnergy = repmat({normEnergy},size(K));
                constructor = str2func(class(obj));
                obj = arrayfun(constructor,f_c,b_f,K,normEnergy,'UniformOutput',false);
                obj = reshape([obj{:}],size(obj));
                return;
            end
            if(isempty(b_f))
                % Set the bandwidth to be 0.8 of the central frequency by
                % default
                b_f = 1/sqrt(2) * f_c;
            end
            if(nargin < 4 || isempty(normEnergy))
                normEnergy = 'none';
            else
                if(iscell(normEnergy))
                    normEnergy = normEnergy{1};
                end
            end
            obj.f_c = f_c;
            obj.b_f = b_f;
            obj.K = K;
            obj.normEnergy = normEnergy;
            
%             obj.n = 2*ceil(K) + 1;
            obj.n = 2*obj.sampleFactor*ceil(K) + 1;
%             obj.angles = (0:obj.n-1)/obj.n*pi;
%             obj.angles = 0:pi/obj.n:pi-pi/obj.n;
            
        end
        function ridgeFilter = real(obj)
            ridgeFilter = OrientationSpaceRidgeFilter(obj.f_c,obj.b_f,obj.K);
            % The filter itself does not change (for the moment)
            ridgeFilter.F = obj.F;
            ridgeFilter.size = obj.size;
        end
        function edgeFilter = imag(obj)
            edgeFilter = OrientationSpaceEdgeFilter(obj.f_c,obj.b_f,obj.K);
            % The filter itself does not change (for the moment)
            edgeFilter.F = obj.F;
            edgeFilter.size = obj.size;
        end
        function angles = get.angles(obj)
            angles = (0:obj.n-1)/obj.n*pi;
        end
        function f_c = get.centralFrequency(obj)
            f_c = obj.f_c;
        end
        function b_f = get.frequencyBandwidth(obj)
            b_f = obj.b_f;
        end
        function K = get.order(obj)
            K = obj.K;
        end
        function n = get.n(obj)
            if(isempty(obj.n))
                obj.n = 2*obj.sampleFactor*ceil(obj.K) + 1;
            end
            n = obj.n;
        end
        function R = mtimes(obj,I)
            % Convolution
            if(isa(obj,'OrientationSpaceFilter'))
                R = getResponse(obj,I);
            elseif(isa(I,'OrientationSpaceFilter'))
                % The convolution is commutative, swap the parameters
                R = getResponse(I,obj);
            end
        end
        function R = getResponse(obj,I)
            If = fft2(I);
            ridgeResponse = obj.applyRidgeFilter(If);
            edgeResponse = obj.applyEdgeFilter(If);
            angularResponse = ridgeResponse + edgeResponse;
            ns = [0 cumsum([obj.n])];
            R(numel(obj)) = OrientationSpaceResponse;
            for o=1:numel(obj)
                R(o) = OrientationSpaceResponse(obj(o),angularResponse(:,:,ns(o)+1:ns(o+1)));
            end
            R = reshape(R,size(obj));
        end
        function R = getRidgeResponse(obj,I)
            If = fft2(I);
            ridgeResponse = obj.applyRidgeFilter(If);
            R = OrientationSpaceResponse(obj,ridgeResponse);
        end
        function R = getEdgeResponse(obj,I)
            If = fft2(I);
            edgeResponse = obj.applyEdgeFilter(If);
            R = OrientationSpaceResponse(obj,edgeResponse);
        end
        function A = getAngularGaussians(obj)
            if(isempty(obj.angularGaussians))
                N = obj.n;
                x = 0:N-1;
                xx = bsxfun(@minus,x,x');
                xx = wraparoundN(xx,-N/2,N/2);
                obj.angularGaussians = exp(-xx.^2/2);
            end
            A = obj.angularGaussians;
        end
        function imshow(obj,n,varargin)
            if(nargin < 2 || isempty(n))
                n = 1;
            end
            if(nargin < 3)
                varargin{1} = [];
            end
            imshow(fftshift(obj.F(:,:,n)),varargin{:});
        end
        function suppress(obj,tol)
            obj.F(abs(obj.F) < tol) = 0;
        end
        function E = getEnergy(obj)
            if(~isscalar(obj))
                E = complex(zeros(numel(obj),max([obj.n])),0);
                for o=1:numel(obj)
                    E(o,1:obj(o).n) = obj(o).getEnergy();
                end
                E = reshape(E,[size(obj) max([obj.n])]);
                return;
            end
            requireSetup(obj);
            s = size(obj.F);
            s(end+1:3) = 1;
            F = reshape(obj.F,s(1)*s(2),s(3));
            E = sqrt(sum(real(F).^2)) + 1j*sqrt(sum(imag(F).^2));
            E = E ./ sqrt(s(1)*s(2));
        end
        function clearTransients(obj)
            for o=1:numel(obj)
                obj(o).size = [];
                obj(o).F = [];
                obj(o).angularGaussians = [];
            end
        end
        function filter = getFilterAtIndex(obj,ind)
            requireSetup(obj);
            coeffs = diric(bsxfun(@minus,(1:obj.n).',ind)*pi/obj.n*2,obj.n);
            filter = reshape(obj.F,prod(obj.size),obj.n)*coeffs;
            filter = reshape(filter,[obj.size length(ind)]);
        end
        function filter = getFilterAtAngle(obj,theta)
            requireSetup(obj);
            % Force theta to be a row vector, 1xT
            theta = theta(:).';
            % Use periodic sinc function to interpolate values
            coeffs = diric(bsxfun(@minus,(0:obj.n-1).'*pi/obj.n*2,theta*2),obj.n);
            % Reshape YxXxN to PxN, then PxN x NxT = PxT
            filter = reshape(obj.F,prod(obj.size),obj.n)*coeffs;

            % If theta in [pi,2*pi), invert imaginary component
            imag_sign = sign(sin(theta+eps*10));
            filter = real(filter) + 1i*bsxfun(@times,imag_sign,imag(filter));

            % Resize PxT to YxXxT
            filter = reshape(filter,[obj.size length(theta)]);
        end
        function h = objshow(obj,varargin)
            requireSetup(obj);
            h = imshow(fftshift(ifft2(real(obj.F(:,:,1)))),varargin{:});
        end
        function circshiftAngles(obj,Kshift)
            for ii=1:numel(obj)
                obj(ii).angles = circshift(obj(ii).angles,Kshift,2);
                if(~isempty(obj(ii).F))
                    obj(ii).F = circshift(obj(ii).F,Kshift,3);
                end
            end
        end
        function flhm = getFullLengthatHalfMaximum(obj,percent)
            if(nargin < 2)
                percent = 0.5;
            end
            if(~isscalar(obj))
                flhm = arrayfun(@(o) getFullLengthatHalfMaximum(o,percent),obj);
                return;
            end
            requireSetup(obj);
            flhm = real(ifft(sum(real(obj.F(:,:,1)),2)));
            flhm = flhm./flhm(1);
            guess = find(flhm < percent,1,'first');
            % Multiply by 2 to get full length
            flhm = interp1(flhm(1:guess),(1:guess)-1,percent,'pchip')*2;
        end
        function fwhm = getFullWidthatHalfMaximum(obj,percent)
            if(nargin < 2)
                percent = 0.5;
            end
            if(~isscalar(obj))
                fwhm = arrayfun(@(o) getFullWidthatHalfMaximum(o,percent),obj);
                return;
            end
            requireSetup(obj);
            fwhm = real(ifft(sum(real(obj.F(:,:,1)),1)));
            fwhm = fwhm./fwhm(1);
            guess = find(fwhm < percent,1,'first');
            % Multiply by 2 to get full width
            fwhm = interp1(fwhm(1:guess),(1:guess)-1,percent,'pchip')*2;
        end
    end
    methods
        function A = getAngularKernel(obj,coords)
            if(nargin < 2)
                coords = orientationSpace.getFrequencySpaceCoordinates(obj.size);
            end
            A = orientationSpace.angularKernel(obj(1).K,obj(1).angles,coords);
        end
        function R = getRadialKernel(obj,coords)
            if(nargin < 2)
                coords = orientationSpace.getFrequencySpaceCoordinates(obj.size);
            end
            R = orientationSpace.radialKernel([obj.f_c], [obj.b_f],coords);
        end
        function setupFilter(obj,siz)
            if(isscalar(siz))
                siz = siz([1 1]);
            end
            coords = orientationSpace.getFrequencySpaceCoordinates(siz);
            notSetup = ~cellfun(@(x) isequal(siz,x),{obj.size});
            notSetup = notSetup | cellfun('isempty',{obj.F});
            obj = obj(notSetup);
            if(isempty(obj))
                return;
            end
            [obj.size] = deal(siz);
            if( all(obj(1).K == [obj.K]) )
                % angular component is all the same
                
%                 A = orientationSpace.angularKernel(obj(1).K,obj(1).angles,coords);
%                 R = orientationSpace.radialKernel([obj.f_c], [obj.b_f],coords);
                A = obj.getAngularKernel(coords);
                R = obj.getRadialKernel(coords);
                for o=1:numel(obj)
                    obj(o).F = bsxfun(@times,A, R(:,:,o));
                end
            else
                for o=1:numel(obj)
                    obj(o).F = orientationSpace.kernel(obj(o).f_c, obj(o).b_f, obj(o).K, obj(o).angles, coords);
                end
            end
            for o=1:numel(obj)
                if(isempty(obj(o).normEnergy))
                    break;
                end
                switch(obj(o).normEnergy)
                    case 'energy'
                        % E is complex
                        E = shiftdim(obj(o).getEnergy(),-1);
                        F = obj(o).F;
                        obj(o).F = bsxfun(@rdivide,real(F),real(E)) +1j*bsxfun(@rdivide,imag(F),imag(E));
                    case 'peak'
                        F = obj(o).F;
                        sumF = sum(F(:))./numel(F);
                        obj(o).F = real(F)./real(sumF) + 1j*imag(F)./imag(sumF);
                    case 'scale'
                        obj(o).F = obj(o).F ./ obj(o).f_c ./ sqrt(siz(1)*siz(2));
                    case 'sqrtscale'
                        obj(o).F = obj(o).F ./ sqrt(obj(o).f_c) ./ sqrt(siz(1)*siz(2));
                    case 'n'
                        obj(o).F = obj(o).F ./ obj(o).n;
                    case 'none'
                    otherwise
                        error('OrientationSpaceFilter:setupFilterNormEnergy', ...
                            'Invalid normEnergy property');
                end
            end
        end
        function ridgeResponse = applyRidgeFilter(obj,If)
            obj.setupFilter(size(If)); %#ok<CPROP>
            ridgeResponse = real(ifft2(bsxfun(@times,If,real(cat(3,obj.F)))));
        end
        function edgeResponse = applyEdgeFilter(obj,If)
            obj.setupFilter(size(If)); %#ok<CPROP>
            edgeResponse = 1j*real(ifft2(bsxfun(@times,If.*-1j,imag(cat(3,obj.F)))));
        end
        function requireSetup(obj)
            if(isempty(obj.F))
                error('OrientationSpaceFilter:NotSetup','Filter must be setup in order for this operation to succeed.');
            end
        end
    end
    methods (Static)
        function F = constructEqualLengthFilters(f_c, b_f, K, normEnergy, constructor)
            if(nargin < 5)
                constructor = @OrientationSpaceFilter;
            end
            %% Approximate cone by height of triangle
            % Largest central frequency or smallest scale
            f_c_max = max(f_c(:));
%             height = sin(pi/(2*K+1))*f_c_max;
            arcLength = pi/(2*K+1)*f_c_max;
            
            assert(isscalar(K));
            
            %% Normal constructor
            s = [length(f_c) length(b_f) 1];
            s(2) = max(1,s(2));
            f_c = repmat(f_c(:),1,s(2),s(3));
            if(isempty(b_f))
                b_f = 1/sqrt(2) * f_c;
            else
                b_f = repmat(b_f(:)',s(1),1,s(3));
            end
            
            if(nargin < 4)
                    normEnergy = [];
            end
            normEnergy = repmat({normEnergy},size(f_c));
            
%             K = (pi./asin(height ./ f_c) - 1)./2;
            K = (pi/arcLength*f_c-1)/2;
            
            F = arrayfun(constructor,f_c,b_f,K,normEnergy,'UniformOutput',false);
            F = reshape([F{:}],size(F));

        end
        function F = constructByRadialOrder(f_c, K_f, K, normEnergy, constructor)
            if(nargin < 4)
                normEnergy = [];
            end
            if(nargin < 5)
                constructor = @OrientationSpaceFilter;
            end
            b_f = f_c ./ sqrt(K_f);
            F = constructor(f_c, b_f, K, normEnergy);
            F = F(logical(eye(length(f_c))));
        end
    end
end
```
