## Supplementary material for "Adaptive-Resolution Multi-Orientation Analysis of Complex Filamentous Network Images": MATLAB code from Github: OrientationSpaceNMS.m

```
classdef OrientationSpaceNMS < handle
    %OrientationSpaceNMS Summary of this class goes here
    %   Detailed explanation goes here
%
% Copyright (C) 2019, Jaqaman Lab - UT Southwestern, Goldman Lab - Northwestern 
%
% This file is part of AdaptiveResolutionOrientationSpace.
% 
% AdaptiveResolutionOrientationSpace is free software: you can redistribute it and/or modify
% it under the terms of the GNU General Public License as published by
% the Free Software Foundation, either version 3 of the License, or
% (at your option) any later version.
% 
% AdaptiveResolutionOrientationSpace is distributed in the hope that it will be useful,
% but WITHOUT ANY WARRANTY; without even the implied warranty of
% MERCHANTABILITY or FITNESS FOR A PARTICULAR PURPOSE.  See the
% GNU General Public License for more details.
% 
% You should have received a copy of the GNU General Public License
% along with AdaptiveResolutionOrientationSpace.  If not, see <http://www.gnu.org/licenses/>.
% 
% 
    
    properties
        response
    end
    properties (Dependent = true)
        nms;
    end
    
    methods
        function obj = OrientationSpaceNMS(response)
            obj.response = response;
        end
        function h = imshow(obj,varargin)
            h = imshow(obj.nms,varargin{:});
        end
        function h = imshowpair(A,B)
            if(nargin > 1)
                if(isa(B,'OrientationSpaceNMS'))
                    B = real(B.nms);
                end
            else
                B = imag(A.nms);
            end
            if(isa(A,'OrientationSpaceNMS'))
                A = real(A.nms);
            end
            h = imshowpair(A,B);
        end
        function nms = get.nms(obj)
            nms = obj.response.nms;
        end
    end
    
end
```
