## Supplementary material for "Adaptive-Resolution Multi-Orientation Analysis of Complex Filamentous Network Images": MATLAB code from Github: README.md

# Adaptive Resolution Orientation Space

## Abstract
Microscopy images of cytoskeletal, nucleoskeletal, and other filamentous
structures contain complex junctions where multiple filaments with distinct
orientations overlap, yet state-of-the-art software generally uses single
orientation analysis to segment these structures.
We describe an image analysis approach to simultaneously segment both
filamentous structures and their intersections (of arbitrary geometry) in microscopy images,
based on analytically resolving coincident multiple orientations upon
image filtering in a manner that balances orientation resolution and
spatial localization.


## Manuscript

"Adaptive-Resolution Multi-Orientation Analysis of Complex Filamentous Network Images"

Mark Kittisopikul<sup>1,2</sup>, Amir Vahabikashi<sup>2</sup>, Takeshi Shimi<sup>2,3</sup>, Robert D. Goldman<sup>2</sup>, and Khuloud Jaqaman<sup>1,4</sup>

### Affiliations

1. Department of Biophysics, UT Southwestern Medical Center, Dallas, TX 75390
2. Department of Cell and Developmental Biology, Feinberg School of Medicine, Northwestern University, Chicago, IL 60611
3. Institute of Innovative Research, Tokyo Institute of Technology, Yokohama, Japan
4. Department of Bioinformatics, UT Southwestern Medical Center, Dallas, TX 75390

## Requirements

MATLAB 2017a or later
* Optimization Toolbox
* Signal Processing Toolbox
* Image Processing Toolbox
* Statistics and Machine Learning Toolbox
* Curve Fitting Toolbox
* Bioinformatics Toolbox
* Parallel Computing Toolbox
* MATLAB Distributed Computing Server

See https://www.mathworks.com/support/requirements/matlab-system-requirements.html for operating system and hardware requirements.

Git (version 2.2 or later) for installation (Alternatively, download a zip file)


## Installation
Typical Install Time: 5 minutes

In MATLAB execute:

    !git clone https://github.com/mkitti/AdaptiveResolutionOrientationSpace 
    addpath(genpath('AdaptiveResolutionOrientationSpace'))

## steerableAdaptiveResolutionOrientationSpaceDetector
Perform adaptive resolution orientation space segmentation. This is the main
driver function that performs the full segmentation as described in Section S6
and illustrated in Figure S4.

### BASIC INPUT (ORDERED ARGUMENTS)
The only required input is a 2D image. The other two basic inputs, which are optional, allow analysis customization.

    I - (required) image
        Type: 2D numeric matrix, non-empty, N x M
    order - (optional), K_h parameter that determines the highest *K* value used for initial filtering via OrientationSpaceFilter
        Type: Numeric, scalar
        Default: 8
    sigma - (optional), scale parameter setting the radial bandpass in pixels
            central frequency, f_c, of the bandpass filter will be 1/(2*pi*sigma) 
        Type: Numeric, scalar
        Default: 2 (pixels)

### ADVANCED INPUTS (NAMED PARAMETERS)
    adaptLengthInRegime - Adapt the resolution with the highest regime by searching for the maxima with the smallest derivative with respect to *K*;
        Type: logical
        Default: true
    meanThresholdMethod - Function to determine threshold of mean response
        Type: char, function_handle
        Default: @thresholdOtsu
    mask - Binary mask the same size as I to limit the area of processing
        Type: logical
        Default: []
    nlmsMask - Override mask for NLMS processing. N x M
        Type: logical
        Default: [] (Calculate mask using mean filter response)
    nlmsThreshold - Override attenuated mean response threshold to apply to NLMS
        Type: numeric, 2D
        Default: [] (Use AMOR)
    useParallelPool - Logical if parallel pool should be used
        Type: logical
        Default: true
    maskDilationDiskRadius - Disc structure element radius in pixels to dilate the mask calculated from the mean response
        Type: numeric
        Default: 3
    maskFillHoles - Logical indicating if holes should be filled in the nlmsMask. True indicates to holes should be filled.
        Type: logical
        Default: false
    diagnosticMode - True if diagnostic figures should be shown
        Type: logical, scalar
        Default: false
    K_sampling_delta - Interval to sample K when using adaptLengthInRegime
        Type: numeric, scalar
        Default: 0.1
    responseOrder - K_m, orientation filter resolution at which to calculate the response values;
        Type: numeric, scalar
        Default: 3
    bridgingLevels - Number of bridging steps to complete. A value of 1 or 2 is valid.
    	Type: numeric, scalar
    	Default: 2
    suppressionValue - Value to assign to pixels that are suppressed in the NMS/NLMS steps
    	Type: numeric, scalar
    	Default: 0
    filter - OrientationSpaceFilter object instance to use, overrides order and sigma parameters; Used to share filter initialization between many function calls
    	Type: OrientationSpaceFilter
    	Default: Create new filter based on order and sigma inputs
    response - OrientationSpaceResponse object to use, overrides order, sigma, and filter; used to share filter response between many function calls.
    	Type: OrientationSpaceResponse
    	Default: Convolve filter with the response to calculate the response

### FURTHER ADVANCED INPUTS: UNSERIALIZATION INPUTS (NAMED PARAMETERS)
These parameters allow some of the output in the struct *other*, below,
to be fed back into the function in order to obtain the full output of the
function. The purpose of this is so that the full output can be regenerated
from a subset of the output that has been saved to disk, or otherwise serialized,
without the need for complete re-computation.

    maxima_highest - numeric 3D array
    K_highest - numeric 3D array
    bridging - struct array
    nlms_highest - numeric 3D array
    nlms_single - numeric 2D array

See OUTPUT for detailed descriptions of the above.

Also note that the *StructExpand* option of the builtin *inputParser* is set to
true, meaning that the named parameters can be passed in using a struct.

### OUTPUT

The main output of the function is the response-weighted segmentation as outlined in Section S6.
This is the 3rd output, *nms*, as described below.
For a binary segmentation, the 2D map *nms* can be thresholded by a single threshold (e.g. `nms > 0`)
or by a threshold map such as *meanResponse* or *attenuatedMeanResponse* in the output struct *other*.

Along with this, the orientations, *theta*, and corresponding *response* values
at K = K<sub>m</sub> are provided as the 2nd and 1st outputs, respectively.
These facilitate use of the function as an orientation detector.
This is meant to mimic the outputs provided by the steerableDetector MATLAB function
available from [François Aguet](http://www.francoisaguet.net/software.html) or
as part of [Filament Analysis Software](http://dx.doi.org/10.17632/xycvj95pw9.1).


    response - Orientation filter response values at resolution K = K_m corresponding to the maxima in theta
        Type: 3D numeric array of dimensions N x M x T. T corresponds to the largest number of maxima found at any pixel in the image.
    theta    - Contains the orientation local maxima detected at each pixel.
        Type: 3D numeric array of dimensions N x M x T. T corresponds to the largest number of maxima found at any pixel in the image.
    nms      - Response-weighted segmentation output (analogous to non-maximum suppression output of previous analyses)
        Type: 2D numeric array of dimensions N x M
        
The fourth output, *angularResponse*, is the sampled orientation responses
at K = K<sub>h</sub> and is again meant for compatibility with steerableDetector.
The *angularResponse* can be used to construct
an *OrientationSpaceResponse* below. This can be used to perform further
analysis of orientation space including for lower resolutions (K < K<sub>h</sub>).

    angularResponse - Filter responses corresponding to equiangular 2K_h+1 samples at resolution K = K_h
        Type: 3D array of dimensions N x M x 2K_h+1
        
The fifth output, *other*, is a structure that contains fields referring to intermediate
results created throughout the analysis process. Importantly, this contains information
about the three AR-NLMS branches employed for segmentation. Because of the maximum response
projections performed at various stages of the algorithm, this information is not readily extracted
from the prior outputs. The information in *other* can be used to determine in which step
of the procedure a pixel was added or excluded from the final output.
Additionally, the orientation information could be used for more precise
localization operations.

    other    - Struct containing the following fields for lower-level analysis and serialization
        .nlms_highest -  AR-NLMS using highest regime maxima using K = K_m responses
            Type: 3D numeric array of dimensions N x M x (T-1)
        .nlms_highest_mip -  Maximum response projection of nlms_highest
            Type: 2D numeric array of dimensions N x M
        .maxima_highest -  Orientation local maxima at highest regime
            Type: 3D numeric array of dimensions N x M x (T-1)
        .K_highest - K values corresponding to maxima in maxima_highest
            Type: 3D numeric array of dimensions N x M x (T-1)
        .maxima_single_angle - Orientation maximum at Regime 0
            Type: 2D numeric array of dimensions N x M
        .nlms_single - NLMS using maximum_single_angle and K = K_m responses
            Type: 2D numeric array of dimensions N x M
        .nlms_single_binary nlms_single thresholded using the attenuatedMeanResponse
            Type: 2D logical array of dimensions N x M
        .meanResponse - Mean orientation filter response
            Type: 2D numeric array of dimensions N x M
        .attenuatedMeanResponse - meanResponse attenuated by neighborhood occupancy
            Type: 2D numeric array of dimensions N x M
        .nlmsMask - Logical mask of the area where the segmentation was analyzed
            Type: 2D logical array of dimensions N x M
        .params - Struct containing the input parameters
            Type: Struct
        .nlmsR NLMS using maxima from the highest regime and from regime 0 using the filter response at K = K_h
            Type: 3D numeric array of dimensions N x M x T
        .nlmsR_mip_binary Maximum response projection of nlmsR thresholded by the attenuatedMeanResponse
            Type: 2D logical array of dimensions N x M
        .bridging A structure array with a length of 2. First element of the array corresponds with the first bridging step. The second element of the array corresponds with the second bridging step.
            .full_binary - (Top input) Array with true values indicating a superset of pixels in the final segmentation
                Type: 2D logical array, N x M
            .consensus_binary - (Left input) 2D logical array containing a subset of pixels used in bridging
                Type: 2D logical array, N x M
            .segments - Connected components to connect together with bridges
                Type: 2D integer array, N x M
            .fragments - Pixels in which to search for bridges between segments
                Type: 2D integer array, N x M
            .bridges - Pixels added to connect segments
                Type: 2D logical array, N x M
            .bridgedSkeleton - 2D logical array, output of the bridging procedure, where the segments have been connected with the bridges and have been subjected to morphological skeletonization


In summary, the outputs allow for basic usage as 
a segmentation scheme (*nms*) and orientation detector
(*response*, *theta*, *angularResponse*),
and for advanced usage as an intermediate routine for further analysis of
the identified multi-resolution features (*other*).


### EXAMPLES
    % Create demo image
    demo = zeros(256);
    demo(128,:) = 1;
    demo = max(imgaussfilt(demo,2),imgaussfilt(eye(256),2));
    demo = imnoise(mat2gray(demo),'gaussian',0.1,0.01);
    % Run basic segmentation analysis
    [res,theta,nms] = steerableAdaptiveResolutionOrientationSpaceDetector(demo);
    figure; imshow(nms,[]);
    % Overlay orientations
    orientationSpace.rainbowOrientationQuivers(theta,res,hsv(32));
    xlim(128+[-10 10]);
    ylim(128+[-10 10]);

![Zoom in Demonstration of Adaptive Resolution Orientation Space and NLMS Analysis](demo/demo.png)

## OrientationSpaceFilter
OrientationSpaceFilter represents a filter that is polar separable in the Fourier domain. It is used internally by steerableAdaptiveResolutionOrientationSpaceDetector but can be used independently.

### Construction

    F = OrientationSpaceFilter(f_c,b_f,K);
    F = OrientationSpaceFilter.constructByRadialOrder(f_c,K_f,K);

* f_c is the central frequency of the filter
* b_f is the frequency bandwidth
* K is the order of the filter that determines orientation resolution
* K_f is the radial frequency order, f_c^2/b_f^2. This is 1 in this work.

### Example

The following two examples are equivalent with f_c = b_f = 1/(4*pi), K_f = 1, and K = 8

    F = OrientationSpaceFilter(1./2/pi./2,1./2./pi./2,8);
    F = OrientationSpaceFilter.constructByRadialOrder(1/2/pi./2,1,8);

### Application to Image

    %% I is just an ordinary image loaded via imread or via BioFormats
    % I = imread('image.tif')
    I = demo;
    R = F*I;

*R* is an *OrientationSpaceResponse* object instance


### Setup and Visualization without an Image

    % Setup filter for 256 x 256 image
    F.setupFilter(256);
    % Show Fourier domain representation
    figure; imshow(F,[]);
    % Show image domain representation
    figure; objshow(F,[]);

## OrientationSpaceResponse
OrientationSpaceResponse represents a filter response. It is used internally by steerableAdaptiveResolutionOrientationSpaceDetector, but can be used independently to examine the filter response.

### Construction

A response object is usually created by an OrientationSpaceFilter as above.

    R = OrientationSpaceResponse(filter,angularResponse)

* filter is an OrientationSpaceFilter
* angularResponse is a 3D array, N x M x 2K+1

### Get ridge orientation local maxima, response, and non-local maxima suppression

    maxima_highest = R.getRidgeOrientationLocalMaxima();
    res_highest_K8 = R.interpft1(maxima_highest);
    nlms_highest_K8 = R.nonLocalMaximaSuppressionPrecise(maxima_highest);

    figure; imshow(max(res_highest_K8,[],3),[]);
    figure; imshow(max(nlms_highest_K8,[],3),[]);

### Get response at K = 3, and non-local maxima suppression

    R3 = R.getResponseAtOrderFT(3,2);
    res_highest_K3 = R3.interpft1(maxima_highest);
    nlms_highest_K3 = R3.nonLocalMaximaSuppressionPrecise(maxima_highest);

    figure; imshow(max(res_highest_K3,[],3),[]);
    figure; imshow(max(nlms_highest_K3,[],3),[]);

### Get derivative of response with respect to orientation

    Rd = R.getDerivativeResponse(1);
    % Should close to zero
    der_highest_K8 = Rd.interpft1(maxima_highest);

    figure; imshow(der_highest_K8(:,:,1),[]);

## LICENSE
Copyright (C) 2019, Jaqaman Lab - UT Southwestern, Goldman Lab - Northwestern 

This file is part of AdaptiveResolutionOrientationSpace.

AdaptiveResolutionOrientationSpace is free software: you can redistribute it and/or modify
it under the terms of the GNU General Public License as published by
the Free Software Foundation, either version 3 of the License, or
(at your option) any later version.

AdaptiveResolutionOrientationSpace is distributed in the hope that it will be useful,
but WITHOUT ANY WARRANTY; without even the implied warranty of
MERCHANTABILITY or FITNESS FOR A PARTICULAR PURPOSE.  See the
GNU General Public License for more details.
 
You should have received a copy of the GNU General Public License
along with AdaptiveResolutionOrientationSpace.  If not, see <http://www.gnu.org/licenses/>.
