## Supplementary material for "Adaptive-Resolution Multi-Orientation Analysis of Complex Filamentous Network Images": MATLAB code from Github: steerableAdaptiveResolutionOrientationSpaceDetector.m

```
function [ response, theta, nms, angularResponse, other ] = steerableAdaptiveResolutionOrientationSpaceDetector( I, varargin )
%steerableAdaptiveResolutionOrientationSpaceDetector Perform adaptive resolution orientation
%space segmentation
% 
% ### BASIC INPUT (ORDERED ARGUMENTS)
% The only required input is a 2D image. The other two basic inputs, which are optional, allow analysis customization.
% 
%     I - (required) image
%         Type: 2D numeric matrix, non-empty, N x M
%     order - (optional), K_h parameter that determines the highest *K* value used for initial filtering via OrientationSpaceFilter
%         Type: Numeric, scalar
%         Default: 8
%     sigma - (optional), scale parameter setting the radial bandpass in pixels
%             central frequency, f_c, of the bandpass filter will be 1/(2*pi*sigma) 
%         Type: Numeric, scalar
%         Default: 2 (pixels)
% 
% ### ADVANCED INPUTS (NAMED PARAMETERS)
%     adaptLengthInRegime - Adapt the resolution with the highest regime by searching for the maxima with the smallest derivative with respect to *K*;
%         Type: logical
%         Default: true
%     meanThresholdMethod - Function to determine threshold of mean response
%         Type: char, function_handle
%         Default: @thresholdOtsu
%     mask - Binary mask the same size as I to limit the area of processing
%         Type: logical
%         Default: []
%     nlmsMask - Override mask for NLMS processing. N x M
%         Type: logical
%         Default: [] (Calculate mask using mean filter response)
%     nlmsThreshold - Override attenuated mean response threshold to apply to NLMS
%         Type: numeric, 2D
%         Default: [] (Use AMOR)
%     useParallelPool - Logical if parallel pool should be used
%         Type: logical
%         Default: true
%     maskDilationDiskRadius - Disc structure element radius in pixels to dilate the mask calculated from the mean response
%         Type: numeric
%         Default: 3
%     maskFillHoles - Logical indicating if holes should be filled in the nlmsMask. True indicates to holes should be filled.
%         Type: logical
%         Default: false
%     diagnosticMode - True if diagnostic figures should be shown
%         Type: logical, scalar
%         Default: false
%     K_sampling_delta - Interval to sample K when using adaptLengthInRegime
%         Type: numeric, scalar
%         Default: 0.1
%     responseOrder - K_m, orientation filter resolution at which to calculate the response values;
%         Type: numeric, scalar
%         Default: 3
%     bridgingLevels - Number of bridging steps to complete. A value of 1 or 2 is valid.
%     	Type: numeric, scalar
%     	Default: 2
%     suppressionValue - Value to assign to pixels that are suppressed in the NMS/NLMS steps
%     	Type: numeric, scalar
%     	Default: 0
%     filter - OrientationSpaceFilter object instance to use, overrides order and sigma parameters; Used to share filter initialization between many function calls
%     	Type: OrientationSpaceFilter
%     	Default: Create new filter based on order and sigma inputs
%     response - OrientationSpaceResponse object to use, overrides order, sigma, and filter; used to share filter response between many function calls.
%     	Type: OrientationSpaceResponse
%     	Default: Convolve filter with the response to calculate the response
% 
% ### FURTHER ADVANCED INPUTS: UNSERIALIZATION INPUTS (NAMED PARAMETERS)
% These parameters allow some of the output in the struct *other*, below,
% to be fed back into the function in order to obtain the full output of the
% function. The purpose of this is so that the full output can be regenerated
% from a subset of the output that has been saved to disk, or otherwise serialized,
% without the need for complete re-computation.
% 
%     maxima_highest - numeric 3D array
%     K_highest - numeric 3D array
%     bridging - struct array
%     nlms_highest - numeric 3D array
%     nlms_single - numeric 2D array
% 
% See OUTPUT for detailed descriptions of the above.
% 
% Also note that the *StructExpand* option of the builtin *inputParser* is set to
% true, meaning that the named parameters can be passed in using a struct.
% 
% ### OUTPUT
% 
% The main output of the function is the response-weighted segmentation as outlined in Section S6.
% This is the 3rd output, *nms*, as described below.
% For a binary segmentation, the 2D map *nms* can be thresholded by a single threshold (e.g. `nms > 0`)
% or by a threshold map such as *meanResponse* or *attenuatedMeanResponse* in the output struct *other*.
% 
% Along with this, the orientations, *theta*, and corresponding *response* values
% at K = K<sub>m</sub> are provided as the 2nd and 1st outputs, respectively.
% These facilitate use of the function as an orientation detector.
% This is meant to mimic the outputs provided by the steerableDetector MATLAB function
% available from [François Aguet](http://www.francoisaguet.net/software.html) or
% as part of [Filament Analysis Software](http://dx.doi.org/10.17632/xycvj95pw9.1).
% 
% 
%     response - Orientation filter response values at resolution K = K_m corresponding to the maxima in theta
%         Type: 3D numeric array of dimensions N x M x T. T corresponds to the largest number of maxima found at any pixel in the image.
%     theta    - Contains the orientation local maxima detected at each pixel.
%         Type: 3D numeric array of dimensions N x M x T. T corresponds to the largest number of maxima found at any pixel in the image.
%     nms      - Response-weighted segmentation output (analogous to non-maximum suppression output of previous analyses)
%         Type: 2D numeric array of dimensions N x M
%         
% The fourth output, *angularResponse*, is the sampled orientation responses
% at K = K<sub>h</sub> and is again meant for compatibility with steerableDetector.
% The *angularResponse* can be used to construct
% an *OrientationSpaceResponse* below. This can be used to perform further
% analysis of orientation space including for lower resolutions (K < K<sub>h</sub>).
% 
%     angularResponse - Filter responses corresponding to equiangular 2K_h+1 samples at resolution K = K_h
%         Type: 3D array of dimensions N x M x 2K_h+1
%         
% The fifth output, *other*, is a structure that contains fields referring to intermediate
% results created throughout the analysis process. Importantly, this contains information
% about the three AR-NLMS branches employed for segmentation. Because of the maximum response
% projections performed at various stages of the algorithm, this information is not readily extracted
% from the prior outputs. The information in *other* can be used to determine in which step
% of the procedure a pixel was added or excluded from the final output.
% Additionally, the orientation information could be used for more precise
% localization operations.
% 
%     other    - Struct containing the following fields for lower-level analysis and serialization
%         .nlms_highest -  AR-NLMS using highest regime maxima using K = K_m responses
%             Type: 3D numeric array of dimensions N x M x (T-1)
%         .nlms_highest_mip -  Maximum response projection of nlms_highest
%             Type: 2D numeric array of dimensions N x M
%         .maxima_highest -  Orientation local maxima at highest regime
%             Type: 3D numeric array of dimensions N x M x (T-1)
%         .K_highest - K values corresponding to maxima in maxima_highest
%             Type: 3D numeric array of dimensions N x M x (T-1)
%         .maxima_single_angle - Orientation maximum at Regime 0
%             Type: 2D numeric array of dimensions N x M
%         .nlms_single - NLMS using maximum_single_angle and K = K_m responses
%             Type: 2D numeric array of dimensions N x M
%         .nlms_single_binary nlms_single thresholded using the attenuatedMeanResponse
%             Type: 2D logical array of dimensions N x M
%         .meanResponse - Mean orientation filter response
%             Type: 2D numeric array of dimensions N x M
%         .attenuatedMeanResponse - meanResponse attenuated by neighborhood occupancy
%             Type: 2D numeric array of dimensions N x M
%         .nlmsMask - Logical mask of the area where the segmentation was analyzed
%             Type: 2D logical array of dimensions N x M
%         .params - Struct containing the input parameters
%             Type: Struct
%         .nlmsR NLMS using maxima from the highest regime and from regime 0 using the filter response at K = K_h
%             Type: 3D numeric array of dimensions N x M x T
%         .nlmsR_mip_binary Maximum response projection of nlmsR thresholded by the attenuatedMeanResponse
%             Type: 2D logical array of dimensions N x M
%         .bridging A structure array with a length of 2. First element of the array corresponds with the first bridging step. The second element of the array corresponds with the second bridging step.
%             .full_binary - (Top input) Array with true values indicating a superset of pixels in the final segmentation
%                 Type: 2D logical array, N x M
%             .consensus_binary - (Left input) 2D logical array containing a subset of pixels used in bridging
%                 Type: 2D logical array, N x M
%             .segments - Connected components to connect together with bridges
%                 Type: 2D integer array, N x M
%             .fragments - Pixels in which to search for bridges between segments
%                 Type: 2D integer array, N x M
%             .bridges - Pixels added to connect segments
%                 Type: 2D logical array, N x M
%             .bridgedSkeleton - 2D logical array, output of the bridging procedure, where the segments have been connected with the bridges and have been subjected to morphological skeletonization
% 
% 
% In summary, the outputs allow for basic usage as 
% a segmentation scheme (*nms*) and orientation detector
% (*response*, *theta*, *angularResponse*),
% and for advanced usage as an intermediate routine for further analysis of
% the identified multi-resolution features (*other*).
% 
% 
% ### EXAMPLES
%     % Create demo image
%     demo = zeros(256);
%     demo(128,:) = 1;
%     demo = max(imgaussfilt(demo,2),imgaussfilt(eye(256),2));
%     demo = imnoise(mat2gray(demo),'gaussian',0.1,0.01);
%     % Run basic segmentation analysis
%     [res,theta,nms] = steerableAdaptiveResolutionOrientationSpaceDetector(demo);
%     figure; imshow(nms,[]);
%     % Overlay orientations
%     orientationSpace.rainbowOrientationQuivers(theta,res,hsv(32));
%     xlim(128+[-10 10]);
%     ylim(128+[-10 10]);
%
%
% Copyright (C) 2019, Jaqaman Lab - UT Southwestern, Goldman Lab - Northwestern 
%
% This file is part of AdaptiveResolutionOrientationSpace.
% 
% AdaptiveResolutionOrientationSpace is free software: you can redistribute it and/or modify
% it under the terms of the GNU General Public License as published by
% the Free Software Foundation, either version 3 of the License, or
% (at your option) any later version.
% 
% AdaptiveResolutionOrientationSpace is distributed in the hope that it will be useful,
% but WITHOUT ANY WARRANTY; without even the implied warranty of
% MERCHANTABILITY or FITNESS FOR A PARTICULAR PURPOSE.  See the
% GNU General Public License for more details.
% 
% You should have received a copy of the GNU General Public License
% along with AdaptiveResolutionOrientationSpace.  If not, see <http://www.gnu.org/licenses/>.
% 
% 

% Mark Kittisopikul, Ph.D.
% Goldman Lab
% Northwestern University
% 2018
    
ip = inputParser;
ip.addRequired('I',@(x) validateattributes(x,{'numeric'},{'2d','nonempty'}));
ip.addOptional('order',8,@(x) validateattributes(x,{'numeric'},{'scalar'}));
ip.addOptional('sigma',2,@(x) validateattributes(x,{'numeric'},{'scalar'}));
ip.addParamValue('filter',[],@(x) validateattributes(x,{'OrientationSpaceFilter'},{'scalar'}));
ip.addParamValue('response',[],@(x) validateattributes(x,{'OrientationSpaceResponse'},{'scalar'}));
ip.addParamValue('adaptLengthInRegime',true,@islogical);
ip.addParamValue('meanThresholdMethod','otsu',@(x) validateattributes(x,{'char','function_handle'},{})); % @(x) validatestring(x,{'otsu','rosin'}));
ip.addParamValue('meanThreshold',[],@(x) validateattributes(x,{'numeric'},{'scalar'}));
ip.addParamValue('mask',[],@islogical);
ip.addParamValue('nlmsMask',[],@islogical);
ip.addParamValue('nlmsThreshold',[],@(x) validateattributes(x,{'numeric'},{'2d'}));
ip.addParamValue('useParallelPool',true,@islogical);
ip.addParamValue('maskDilationDiskRadius',3,@(x) validateattributes(x,{'numeric'},{'scalar'}));
ip.addParamValue('maskFillHoles',false,@(x) validateattributes(x,{'logical'},{'scalar'}));
ip.addParamValue('diagnosticMode',false,@(x) validateattributes(x,{'logical'},{'scalar'}));
ip.addParamValue('K_sampling_delta',0.1,@(x) validateattributes(x,{'numeric'},{'scalar'}));
ip.addParamValue('responseOrder',3,@(x) validateattributes(x,{'numeric'},{'scalar'}));

ip.parse(I,varargin{:});

if( isempty(ip.Results.response) )
    if( isempty(ip.Results.filter) )
        F = OrientationSpaceFilter.constructByRadialOrder(1/2/pi./ip.Results.sigma,1,ip.Results.order,'none');
    else
        F = ip.Results.filter;
    end

    R = F*I;
else
    R = ip.Results.response;
    F = ip.Results.filter;
end

if(ip.Results.useParallelPool)
    pool = gcp;
end

% Obtain Fourier transform
a_hat = fft(real(R.a),[],3);

%% Evaluate and threshold mean first to quickly reduce the problem
meanResponse = a_hat(:,:,1)./size(a_hat,3);
if(ip.Results.diagnosticMode)
    oldPrefImshowBorder = iptgetpref('ImshowBorder');
    iptsetpref('ImshowBorder','loose');
    other.diagFig(1) = figure('Name','meanResponse');
    imshow(meanResponse,[]);
    title(other.diagFig(end).Children,other.diagFig(end).Name, ...
        'Interpreter','none')
end

%% Thresholding
if(isempty(ip.Results.meanThreshold))
    % If method is char, convert to a function
    if(ischar(ip.Results.meanThresholdMethod))
        switch(ip.Results.meanThresholdMethod)
            case 'otsu'
                meanThresholdMethod = @thresholdOtsu;
            case 'rosin'
                meanThresholdMethod = @thresholdRosin;
            otherwise
                meanThresholdMethod = str2func(ip.Results.meanThresholdMethod);
        end
    else
        % Assume this is a function
        meanThresholdMethod = ip.Results.meanThresholdMethod;
    end

    % We could mask the thresholding function, but ...
%     if(~isempty(ip.Results.mask))
%         meanThreshold = meanThresholdMethod(meanResponse(ip.Results.mask));
%     else
%         meanThreshold = meanThresholdMethod(meanResponse);
%     end

    % It is more flexible to leave it up to the user
    meanThreshold = meanThresholdMethod(meanResponse);
else
    % User can also set a fixed threshold directly
    % This can a single threshold or the size of the image
    meanThreshold = ip.Results.meanThreshold;
end

if(ip.Results.diagnosticMode)
    other.diagFig(end+1) = figure('Name','meanResponse Histogram');
    histogram(meanResponse(:));
    xlabel('meanResponse');
    ylabel('count');
    line([1 1].*meanThreshold,ylim,'Color','r','LineWidth',2);
    title(other.diagFig(end).Children,other.diagFig(end).Name, ...
        'Interpreter','none')
end

%% Masking

if(isempty(ip.Results.nlmsMask))
    meanResponseMask = meanResponse > meanThreshold;
    if(ip.Results.diagnosticMode)
        other.diagFig(end+1) = figure('Name','meanResponse > meanThreshold');
        imshow(meanResponseMask,[]);
        title(other.diagFig(end).Children,other.diagFig(end).Name, ...
            'Interpreter','none')
    end
    if(~isempty(ip.Results.mask))
        meanResponseMask = meanResponseMask & ip.Results.mask;
    end
    if(ip.Results.diagnosticMode)
        other.diagFig(end+1) = figure('Name','meanResponseMask');
        imshow(meanResponseMask,[]);
        title(other.diagFig(end).Children,other.diagFig(end).Name, ...
            'Interpreter','none')
    end

    % For NLMS (nlmsMask)
    meanResponseMaskDilated = imdilate(meanResponseMask,strel('disk',ip.Results.maskDilationDiskRadius));
    if(ip.Results.maskFillHoles)
        meanResponseMaskDilated = imfill(meanResponseMaskDilated,'holes');
    end
    if(~isempty(ip.Results.mask))
        meanResponseMaskDilated = meanResponseMaskDilated & ip.Results.mask;
    end
    if(ip.Results.diagnosticMode)
        other.diagFig(end+1) = figure('Name','meanResponseMaskDilated');
        imshow(meanResponseMaskDilated,[]);
        title(other.diagFig(end).Children,other.diagFig(end).Name, ...
            'Interpreter','none')
        diag_rp = regionprops(meanResponseMaskDilated,'BoundingBox','Area');
        [~,diag_rp_max_idx] = max([diag_rp.Area]);
        diag_rp = diag_rp(diag_rp_max_idx);
        rectangle('Position',diag_rp.BoundingBox,'EdgeColor','r');
    end

    nlmsMask = meanResponseMaskDilated;
else
    % User defined nlmsMask
    nlmsMask = ip.Results.nlmsMask;
    if(ip.Results.diagnosticMode)
        diag_rp = regionprops(nlmsMask,'BoundingBox','Area');
    end
end

%% Setup orientation analysis problem
nanTemplate = NaN(size(nlmsMask));

a_hat = shiftdim(a_hat,2);
a_hat = a_hat(:,nlmsMask);

%% Evaluate single orientation, fast easy case

R_res = R.getResponseAtOrderFT(ip.Results.responseOrder,2);
maximum_single_angle = nanTemplate;
maximum_single_angle(nlmsMask) = wraparoundN(-angle(a_hat(2,:))/2,0,pi);
nlms_single = nonLocalMaximaSuppressionPrecise(real(R_res.a),maximum_single_angle,[],[],nlmsMask);
nlms_single_binary = nlms_single > meanResponse;

if(ip.Results.diagnosticMode)
    other.diagFig(end+1) = figure('Name','maximum_single_angle');
    
    cm = colorcet('CBC2','N',256);
    
    
    maximum_single_angle_map = orientationSpace.blendOrientationMap(maximum_single_angle,R_res.interpft1(maximum_single_angle),cm);
    imshow(maximum_single_angle_map,[]);
    caxis([-1/256 pi]);
    title(other.diagFig(end).Children,other.diagFig(end).Name, ...
        'Interpreter','none')
    xlim([0 diag_rp.BoundingBox(3)]+diag_rp.BoundingBox(1));
    ylim([0 diag_rp.BoundingBox(4)]+diag_rp.BoundingBox(2));
    
    other.diagFig(end+1) = figure('Name','nlms_single');
    imshow(nlms_single,[]);
    title(other.diagFig(end).Children,other.diagFig(end).Name, ...
        'Interpreter','none')
    xlim([0 diag_rp.BoundingBox(3)]+diag_rp.BoundingBox(1));
    ylim([0 diag_rp.BoundingBox(4)]+diag_rp.BoundingBox(2));
    
   other.diagFig(end+1) = figure('Name','nlms_single_binary');
   imshow(nlms_single_binary,[]);
   title(other.diagFig(end).Children,other.diagFig(end).Name, ...
        'Interpreter','none');
    xlim([0 diag_rp.BoundingBox(3)]+diag_rp.BoundingBox(1));
    ylim([0 diag_rp.BoundingBox(4)]+diag_rp.BoundingBox(2));
end

%% Determine nlmsThreshold
if(isempty(ip.Results.nlmsThreshold))
    %% Attenuate meanResponse by neighbor occupancy
    nhood_filter = [1 1 1; 1 0 1; 1 1 1];
    nhood_occupancy = imfilter(double(nlms_single_binary),nhood_filter)/8;
    % double the occupancy for accelerated attenuation
    nhood_occupancy = nhood_occupancy*2; 
    attenuatedMeanResponse = (1-nhood_occupancy).*meanResponse;
    attenuatedMeanResponse = max(attenuatedMeanResponse,0);

    nlmsThreshold = attenuatedMeanResponse;
else
    % User defined nlmsThreshold
    nlmsThreshold = ip.Results.nlmsThreshold;
end

if(ip.Results.diagnosticMode)
    other.diagFig(end+1) = figure('Name','nlmsThreshold');
    imshow(nlmsThreshold,[]);
    title(other.diagFig(end).Children,other.diagFig(end).Name, ...
        'Interpreter','none');
    xlim([0 diag_rp.BoundingBox(3)]+diag_rp.BoundingBox(1));
    ylim([0 diag_rp.BoundingBox(4)]+diag_rp.BoundingBox(2));
end


%% Calculate high resolution maxima

% Adapt length
if(ip.Results.adaptLengthInRegime)
    % Find orientation maxima with nlmsMask only
    [maxima_highest_temp,minima_highest_temp] = interpft_extrema(a_hat,1,true,[],false);
    % Count
    n_maxima_highest_temp = ...
        size(maxima_highest_temp,1) - sum(isnan(maxima_highest_temp));
    K_high = F.K;
    K_low = max(n_maxima_highest_temp - 1,ip.Results.responseOrder);
    warning('off','halleyft:maxIter');
    [K_high,K_low] = ...
        orientationSpace.diffusion.findRegimeBifurcation( ...
            a_hat,F.K, ...
            K_high,K_low, ...
            maxima_highest_temp,minima_highest_temp, ...
            [],0.1,true);
    best_derivs = orientationSpace.diffusion.orientationMaximaFirstDerivative(a_hat,F.K,maxima_highest_temp);
    best_abs_derivs = abs(best_derivs);
    best_K = repmat(F.K,size(best_derivs));
    best_maxima = maxima_highest_temp;
    maxima_working = maxima_highest_temp;
    for K=F.K:-ip.Results.K_sampling_delta:1
        s = K > K_high;
        lower_a_hat = orientationSpace.getResponseAtOrderVecHat(a_hat(:,s),F.K,K);
        [new_derivs(:,s),~,maxima_working(:,s)] = orientationSpace.diffusion.orientationMaximaFirstDerivative(lower_a_hat,K,maxima_working(:,s),[],true);
        new_abs_derivs(:,s) = abs(new_derivs(:,s));
        better(:,s) = new_abs_derivs(:,s) < best_abs_derivs(:,s);
        
        % Update better
        best_abs_derivs(better) = new_abs_derivs(better);
        best_derivs(better) = new_derivs(better);
        best_K(better) = K;
        best_maxima(better) = maxima_working(better);
    end
    
    maxima_highest_temp = best_maxima / 2;
else
    % Find orientation maxima with nlmsMask only
    maxima_highest_temp = interpft_extrema(a_hat,1,true,[],false)/2;
    best_K = repmat(F.K,size(maxima_highest_temp));
end

maxima_highest = nanTemplate(:,:,ones(size(maxima_highest_temp,1),1));
maxima_highest = shiftdim(maxima_highest,2);
for i=1:size(maxima_highest_temp,1)
    maxima_highest(i,nlmsMask) = maxima_highest_temp(i,:);
end
maxima_highest = shiftdim(maxima_highest,1);
if(ip.Results.diagnosticMode)
    other.diagFig(end+1) = figure('Name','maxima_highest(:,:,1)');
    imshow(maxima_highest(:,:,1),[]);
    caxis([0 pi]); colormap(hsv);
    title(other.diagFig(end).Children,other.diagFig(end).Name, ...
        'Interpreter','none')
    xlim([0 diag_rp.BoundingBox(3)]+diag_rp.BoundingBox(1));
    ylim([0 diag_rp.BoundingBox(4)]+diag_rp.BoundingBox(2));
    
    other.diagFig(end+1) = figure('Name','maxima_highest');
    cm = colorcet('CBC2','N',256);
    maxima_highest_response = R_res.interpft1(maxima_highest);
    imshow(max(maxima_highest_response,[],3),[]);
    orientationSpace.rainbowOrientationQuivers(maxima_highest,maxima_highest_response,cm);
    title(other.diagFig(end).Children,other.diagFig(end).Name, ...
        'Interpreter','none');
    xlim([0 diag_rp.BoundingBox(3)]+diag_rp.BoundingBox(1));
    ylim([0 diag_rp.BoundingBox(4)]+diag_rp.BoundingBox(2));
end

%% Perform NLMS

% Perform nlms within nlmsMask and apply nlmsThreshold for highest
% resolution maxima at lower response level
nlms_highest = nonLocalMaximaSuppressionPrecise(real(R_res.a),maxima_highest,[],[],nlmsMask);
nlms_highest_mip = max(nlms_highest,[],3);
nlms_highest_mip_binary = nlms_highest_mip > nlmsThreshold;
if(ip.Results.diagnosticMode)
    other.diagFig(end+1) = figure('Name','nlms_highest_mip');
    imshow(nlms_highest_mip,[]);
    title(other.diagFig(end).Children,other.diagFig(end).Name, ...
        'Interpreter','none')
    xlim([0 diag_rp.BoundingBox(3)]+diag_rp.BoundingBox(1));
    ylim([0 diag_rp.BoundingBox(4)]+diag_rp.BoundingBox(2));
    
    other.diagFig(end+1) = figure('Name','nlms_highest_mip_binary');
    imshow(nlms_highest_mip_binary,[]);
    title(other.diagFig(end).Children,other.diagFig(end).Name, ...
        'Interpreter','none');
    xlim([0 diag_rp.BoundingBox(3)]+diag_rp.BoundingBox(1));
    ylim([0 diag_rp.BoundingBox(4)]+diag_rp.BoundingBox(2));
end

% Perform nlms within nlmsMask and apply nlmsThreshold for highest
% resolution maxima and single maxima at high response level
nlmsR = nonLocalMaximaSuppressionPrecise(real(R.a),cat(3,maxima_highest,maximum_single_angle),[],[],nlmsMask);
nlmsR_mip = max(nlmsR,[],3);
nlmsR_mip_binary = nlmsR_mip > nlmsThreshold;
if(ip.Results.diagnosticMode)
    other.diagFig(end+1) = figure('Name','nlmsR_mip');
    imshow(nlmsR_mip,[]);
    title(other.diagFig(end).Children,other.diagFig(end).Name, ...
        'Interpreter','none');
    xlim([0 diag_rp.BoundingBox(3)]+diag_rp.BoundingBox(1));
    ylim([0 diag_rp.BoundingBox(4)]+diag_rp.BoundingBox(2));
    
    other.diagFig(end+1) = figure('Name','nlmsR_mip_binary');
    imshow(nlmsR_mip_binary,[]);
    title(other.diagFig(end).Children,other.diagFig(end).Name, ...
        'Interpreter','none');
    xlim([0 diag_rp.BoundingBox(3)]+diag_rp.BoundingBox(1));
    ylim([0 diag_rp.BoundingBox(4)]+diag_rp.BoundingBox(2));
end

% Calculate the maximum projected response at the lower response level
% using highest and lowest resolution maxima
combinedMaxima = cat(3,maxima_highest,maximum_single_angle);
combinedResR = R_res.interpft1(combinedMaxima);
combinedResR_mip = max(combinedResR,[],3);
if(ip.Results.diagnosticMode)
    other.diagFig(end+1) = figure('Name','combinedResR_mip');
    imshow(combinedResR_mip,[]);
    title(other.diagFig(end).Children,other.diagFig(end).Name, ...
        'Interpreter','none');
    xlim([0 diag_rp.BoundingBox(3)]+diag_rp.BoundingBox(1));
    ylim([0 diag_rp.BoundingBox(4)]+diag_rp.BoundingBox(2));
end

%% Calculate consensus and determine pixels for potential improvement
consensus_binary = nlms_highest_mip_binary & nlms_single_binary;
if(ip.Results.diagnosticMode)
    other.diagFig(end+1) = figure('Name','consensus_binary');
    imshow(consensus_binary,[]);
    title(other.diagFig(end).Children,other.diagFig(end).Name, ...
        'Interpreter','none');
    xlim([0 diag_rp.BoundingBox(3)]+diag_rp.BoundingBox(1));
    ylim([0 diag_rp.BoundingBox(4)]+diag_rp.BoundingBox(2));
end

%% Do minimal bridging

if(~ip.Results.diagnosticMode)
    [bridgedSkeletonOne, other.bridging] = ...
        bridging.minimalBridgingFromBinary( ...
        I, ...
        nlms_highest_mip_binary, ...
        consensus_binary, ...
        5, ...
        ip.Results.diagnosticMode);
else

    [bridgedSkeletonOne, other.bridging, other.diagFig(end+1:end+2)] = ...
        bridging.minimalBridgingFromBinary( ...
        I, ...
        nlms_highest_mip_binary, ...
        consensus_binary, ...
        5, ...
        ip.Results.diagnosticMode, ...
        diag_rp.BoundingBox);
    
end

%% Repeat bridging with nlmsR

if(~ip.Results.diagnosticMode)
    [bridgedSkeletonTwo, other.bridging(2)] = ...
            bridging.minimalBridgingFromBinary( ...
            I, ...
            nlmsR_mip_binary | bridgedSkeletonOne, ...
            bridgedSkeletonOne, ...
            10, ...
            ip.Results.diagnosticMode);
else

    [bridgedSkeletonTwo, other.bridging(2), other.diagFig(end+1:end+2)] = ...
        bridging.minimalBridgingFromBinary( ...
        I, ...
        nlmsR_mip_binary | bridgedSkeletonOne, ...
        bridgedSkeletonOne, ...
        10, ...
        ip.Results.diagnosticMode, ...
        diag_rp.BoundingBox);

    
end


response = combinedResR;
theta = combinedMaxima;
response_nan = isnan(response);
response(response_nan) = -Inf;
[response,theta,response_nan] = sortMatrices(response,theta,response_nan,3,'descend');
response(response_nan) = NaN;

nms = bridgedSkeletonTwo.*combinedResR_mip;
angularResponse = R.a;

if(nargout > 4)
    % Process and output NLMS / maxima from highest K
    nlms_highest_nan = isnan(nlms_highest);
    nlms_highest(nlms_highest_nan) = -Inf;
    [nlms_highest,maxima_highest,nlms_highest_nan,combinedResR(:,:,1:end-1)] = sortMatrices(nlms_highest,maxima_highest,nlms_highest_nan,combinedResR(:,:,1:end-1),3,'descend');
    nlms_highest(nlms_highest_nan) = NaN;
    
    other.nlms_highest = nlms_highest;
    other.maxima_highest = maxima_highest;
    other.nlms_highest_mip = nlms_highest_mip;
    
    % Process and output NLMS / maxima from single regime
    other.maximum_single_angle = maximum_single_angle;
    other.nlms_single = nlms_single;
    other.nlms_single_binary = nlms_single_binary;
    other.attenuatedMeanResponse = attenuatedMeanResponse;
    other.meanResponse = meanResponse;
    other.nlmsMask = nlmsMask;
    other.params = ip.Results;
    other.combinedResR = combinedResR;
    other.nlmsR = nlmsR;
    other.nlmsR_mip_binary = nlmsR_mip_binary;
    
    % format K array
    other.K_highest = nanTemplate(:,:,ones(size(best_K,1),1));
    other.K_highest = shiftdim(other.K_highest,2);
    for i=1:size(best_K,1)
        other.K_highest(i,nlmsMask) = best_K(i,:);
    end
    other.K_highest = shiftdim(other.K_highest,1);
    other.K_highest(isnan(maxima_highest)) = NaN;
end

if(ip.Results.diagnosticMode)
    other.diagFig(end+1) = figure('Name','response');
    imshow(max(response,[],3),[]);
    h = orientationSpace.rainbowOrientationQuivers(theta,response);
    set(h,'LineWidth',2);
    title(other.diagFig(end).Children,other.diagFig(end).Name, ...
        'Interpreter','none');
    xlim([0 diag_rp.BoundingBox(3)]+diag_rp.BoundingBox(1));
    ylim([0 diag_rp.BoundingBox(4)]+diag_rp.BoundingBox(2));
    
    other.diagFig(end+1) = figure('Name','nms');
    imshow(nms,[]);    
    title(other.diagFig(end).Children,other.diagFig(end).Name, ...
        'Interpreter','none')
    xlim([0 diag_rp.BoundingBox(3)]+diag_rp.BoundingBox(1));
    ylim([0 diag_rp.BoundingBox(4)]+diag_rp.BoundingBox(2));
    
    other.diagFig(end+1) = figure('Name','nms > 0');
    imshow(nms > 0,[]);    
    title(other.diagFig(end).Children,other.diagFig(end).Name, ...
        'Interpreter','none')
    xlim([0 diag_rp.BoundingBox(3)]+diag_rp.BoundingBox(1));
    ylim([0 diag_rp.BoundingBox(4)]+diag_rp.BoundingBox(2));
    
    iptsetpref('ImshowBorder',oldPrefImshowBorder);
end


end
```
