## Supplementary figures and images for "Adaptive-Resolution Multi-Orientation Analysis of Complex Filamentous Network Images"

### demo.png

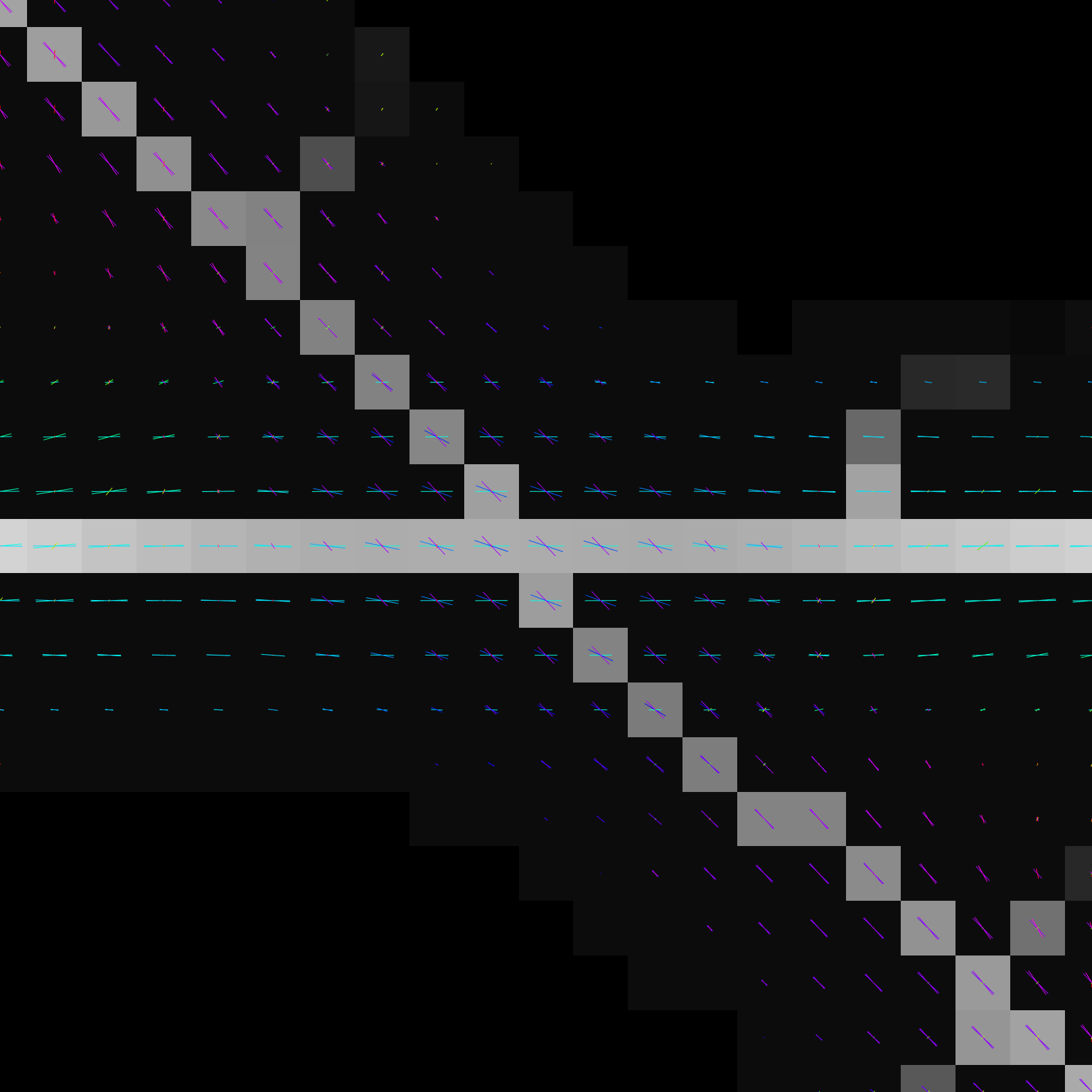
