## Supplementary material for "Adaptive-Resolution Multi-Orientation Analysis of Complex Filamentous Network Images": URL for Uncompressed Movies

### Supplemental Movies

The supplemental movies have been compressed to fit the 40 MB bioRxiv file size limit. The original movies can be found at this URL:

https://cloud.biohpc.swmed.edu/index.php/s/bYNxPPHtE5xkGFC

The legend for the movies can be found at the end of the supplemental material.
